# Enzymatic bromination of native peptides for late-stage structural diversification via Suzuki-Miyaura coupling

**DOI:** 10.64898/2025.12.17.694899

**Authors:** Haley N. Bridge, Chase L. Radziej, Amy M. Weeks

**Affiliations:** Department of Biochemistry, University of Wisconsin – Madison, Madison, WI, USA 53706; Department of Chemistry, University of Wisconsin – Madison, Madison, WI, USA 53706

## Abstract

Flavin-dependent halogenases provide a biocatalytic approach for site-selective halogenation of aromatic compounds, but their use in late-stage functionalization of peptides has remained limited. Here, we show that the tryptophan (Trp) 7-halogenase RebH and an engineered variant (4V) originally optimized for larger small-molecule scaffolds can brominate peptidyl-Trp residues across a broad range of sequence and positional contexts. Through extensive analysis of diverse substrate sequences, we define features that enable RebH activity and reveal 4V’s expanded sequence tolerance. We applied 4V for enzymatic bromination of diverse bioactive peptide scaffolds, including an antimicrobial peptide, a cell-penetrating peptide, and a G protein-coupled receptor agonist, without the need for sequence modification. These brominated peptides served as substrates for Suzuki-Miyaura coupling, enabling installation of functional groups that conferred new functional properties or tuned the biological activity of these peptides. Our results expand the substrate landscape of FDHs and establish bromination-enabled cross-coupling as a general approach for late-stage diversification of bioactive peptides.

## Introduction

Residue-specific modification of proteins and peptides enables diverse applications in chemistry and biology, including the development of covalent inhibitors, functional probes, antibody-drug conjugates, and chemoproteomics tools^1–3^. Tryptophan represents an attractive target for selective modification based on its unique reactivity^4–12^, relative rarity in the proteome^13^, and occurrence in many bioactive peptide classes^14–17^. However, selective modification of Trp in unprotected peptides and proteins remains challenging. Most chemical approaches for Trp modification rely on the nucleophilicity of Trp and suffer from cross-reactivity with Cys, His, and/or Tyr^1^. A recent strategy based on oxidative cyclization demonstrated that Trp residues can be selectively modified under mild, biocompatible conditions to install valuable bioorthogonal handles^12^. However, a key limitation of current approaches is that they cannot introduce versatile synthetic handles, such as aryl halides, that enable modular C–C, C–N, and C–O bond formation for downstream functionalization^18–20^. Aryl bromides and aryl iodides, in particular, are attractive intermediates for late-stage structural diversification via Pd-catalyzed cross-coupling, a powerful and widely used platform in medicinal chemistry and bioconjugation^21,22^.

Enzymatic halogenation by flavin-dependent halogenases (FDHs) offers a complementary strategy for generating aryl halides as cross-coupling substrates under mild, aqueous conditions^23–27^. Numerous FDHs that halogenate free Trp have been discovered^28–33^. A subset of these can modify Trp residues in peptides, typically when Trp is positioned at one of the termini or in the presence of a leader sequence that enables substrate recognition^34–37^. For example, SrpI can modify Trp in the context of a C-terminal heptapeptide motif (LTVPW)^34^, PyrH can modify C-terminal Trp in a (G/S)GW motif^36^, and ThaI can modify peptides and proteins with an engineered C-terminal Trp-containing sequence^37^. A recently characterized peptide FDH from the chlorolassin biosynthetic gene cluster, ChlH, expands this scope by halogenating internal Trp residues in diverse peptide substrates^38^. However, ChlH is highly selective for chlorination and does not efficiently catalyze bromination or iodination, limiting its utility for installing aryl bromides or aryl iodides as synthetic handles for structural diversification.

To expand the synthetic potential of FDHs, both directed evolution and genome mining approaches have been applied to identify biocatalysts with desirable properties^39^. In particular, the tryptophan 7-halogenase RebH has been extensively engineered to improve its thermal stability^40^, alter its regioselectivity^41^, and expand its substrate scope to include Trp analogs^42^, larger aromatic scaffolds^43^, and aromatic fragment compounds used in drug discovery (**Fig. 1**)^44^.

**Figure 1.**
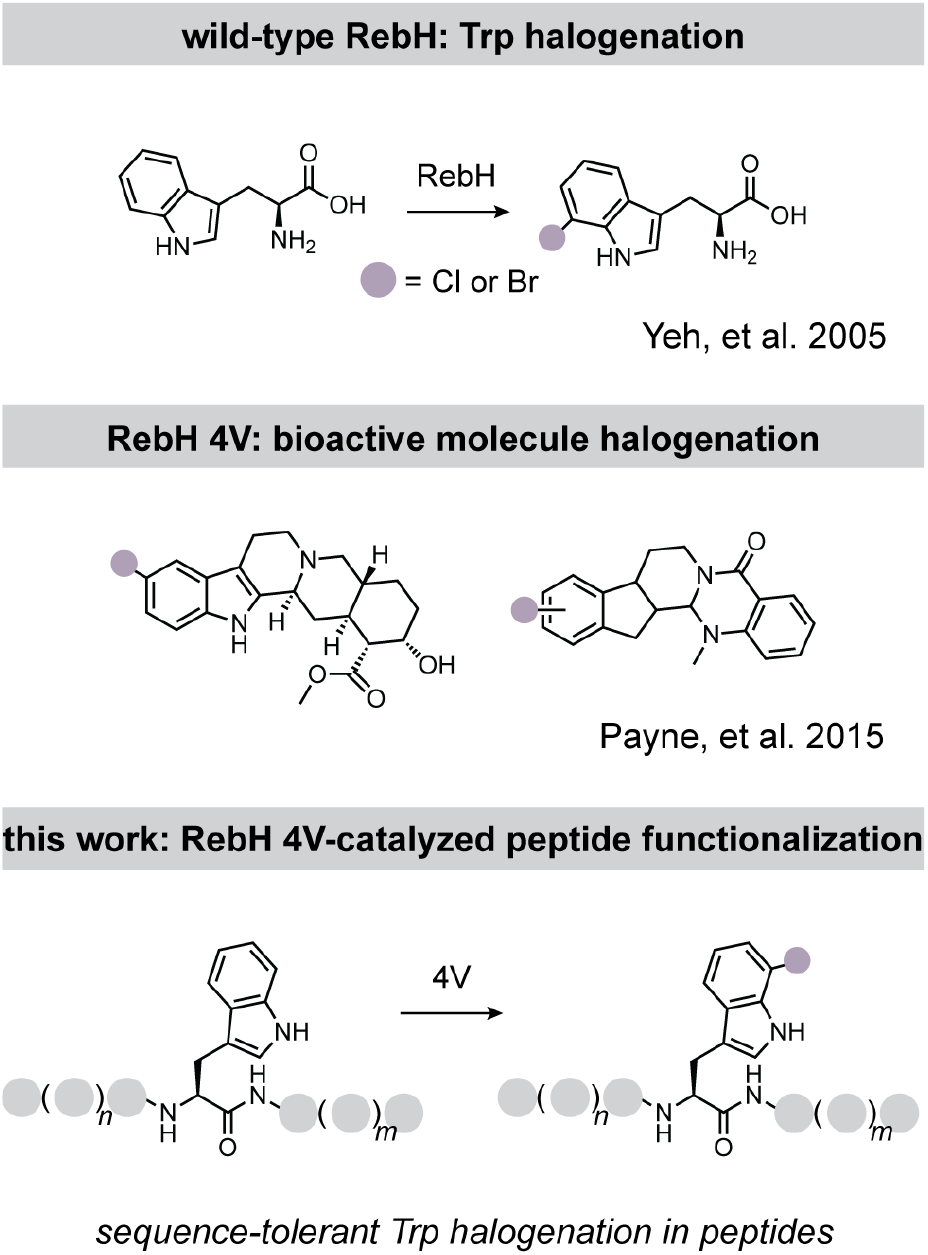
Application of the Trp 7-halogenase RebH and its variants for late-stage peptide functionalization. Top, wild-type RebH catalyzes chlorination or bromination of free Trp. Middle, the RebH variant 4V was engineered to halogenate larger aromatic scaffolds. Bottom, in this work, we apply RebH 4V for sequence-tolerant Trp halogenation in the context of peptides.

RebH is highly amenable to protein engineering based on foundational work from the Lewis lab that established a strategy for high-yield expression in *E. coli*, enabling LC-MS-based screening in lysate, and the development of a robust system for regeneration of the FADH_2_ cofactor required for enzyme activity^44^. However, most engineering efforts have focused on small molecules, and the ability of RebH variants to act on peptides has remained underexplored.

Here, we report a chemoenzymatic approach for late-stage diversification of Trp-containing peptides that integrates RebH-catalyzed bromination and Suzuki-Miyaura cross-coupling (**Fig. 1**). We leveraged the previously developed LC-MS-based screening platform^45^ to systematically evaluate the activity of RebH and a variant with expanded substrate scope (4V, engineered in the Lewis lab^43^) on peptides with varying length, sequence, and Trp position. We found that RebH displays robust peptide halogenation activity, while 4V exhibits enhanced activity and expanded substrate scope. Based on these results, we used 4V in a modular strategy for generating peptidyl-aryl bromides from native Trp-containing peptides that can be diversified via Suzuki-Miyaura coupling. This approach enabled us to generate structurally complex peptide derivatives that are otherwise challenging to synthesize. Using this strategy, we synthesized analogs of diverse bioactive peptides, including a fluorescent derivative of the cell-penetrating peptide transportan, analogs of the antimicrobial peptide (RW)_3_ with enhanced potency, and analogs of the endogenous opioid peptide endomorphin-1 with altered μ-opioid receptor agonist activity. Together, these results establish enzymatic halogenation as a generalizable approach for structural diversification of native bioactive peptides via Pd-catalyzed cross-coupling.

## Results

### RebH and 4V possess robust bromination activity on peptide substrates

RebH bromination activity has been measured previously on dozens of small molecule substrates, and extensive rational design and directed evolution campaigns have been undertaken to modify its substrate scope (**Fig. 1**)^39,42,43^. However, wild-type RebH was previously reported to lack activity on peptide substrates^36^, and the peptide activity of RebH variants remains largely unexplored. We hypothesized that 4V^43^, a variant of RebH engineered to have activity on small molecule substrates larger than Trp, might have activity on Trp-containing peptides (**Fig. 1**, bottom). Recent discoveries that other FDHs can catalyze protein and peptide bromination suggest that such activity is possible^35–37^, although these systems often rely on engineered recognition elements such as leader sequences or tags.

To test the ability of RebH and 4V to brominate peptides, we synthesized a panel of Trp-containing sequences in which Trp was placed at the N terminus, C terminus, or center, with varying numbers of glycine residues (Gly_n_, n = 1–5) appended to one or both sides (**Fig. 2**). We then incubated each peptide with RebH, 4V, or the inactive RebH mutant K79A for 20 h and determined percent conversion to brominated product using an LC-HRMS-based assay (**Supplementary Fig. 1-3**). Surprisingly, we found that both 4V and RebH had substantial activity on all peptides tested (**Fig. 2, Supplementary Fig. 4**). The brominated products exhibited the expected 1:1 distribution of ^79^Br and ^81^Br (**Fig. 2a-c**, center), and their structures were confirmed using 1D ^1^H and 2D COSY, TOCSY, and NOESY NMR experiments (**Supplementary Fig. 5-16)**.

**Figure 2.**
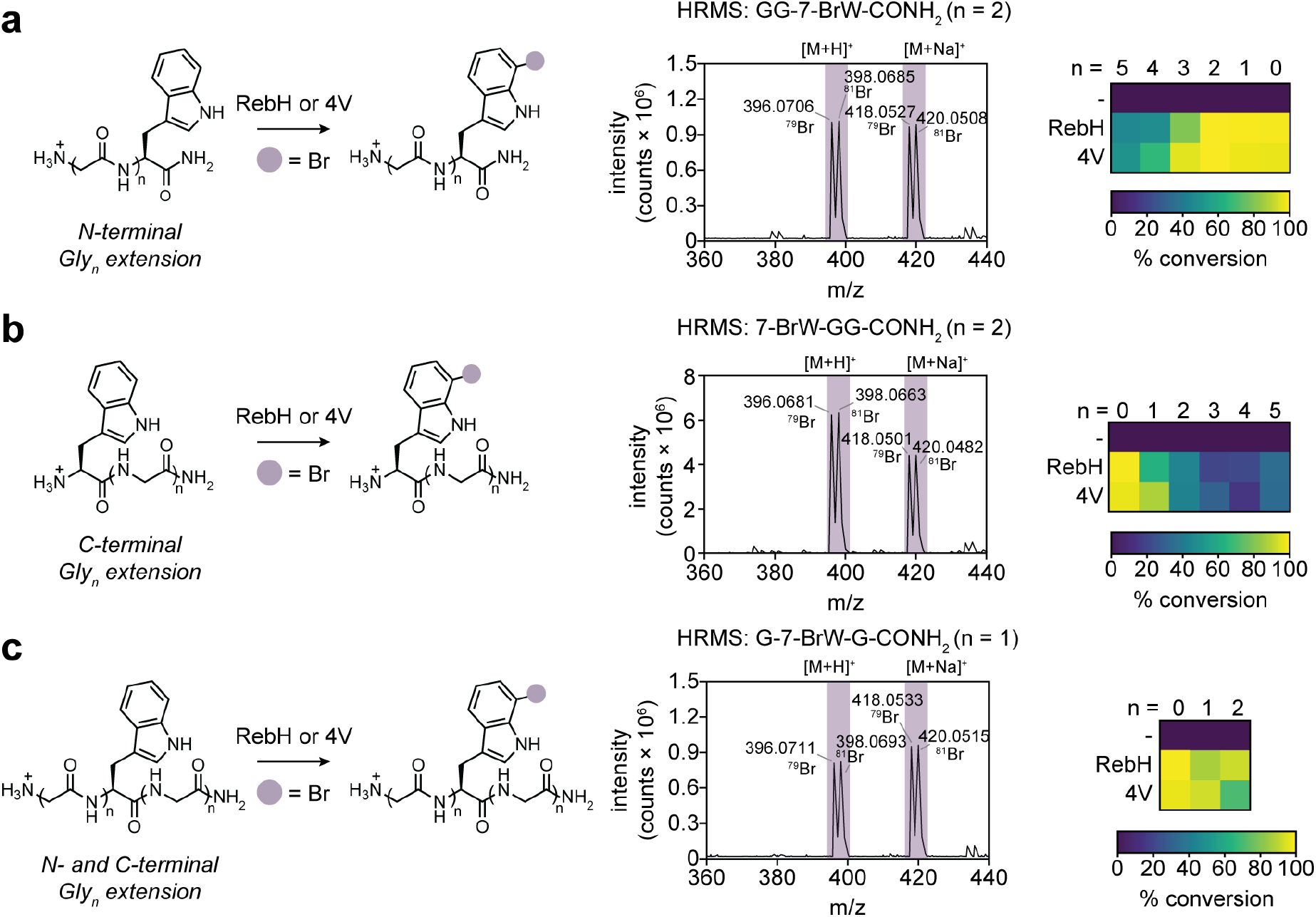
RebH and 4V catalyze bromination of Trp-containing peptides. (a) Left, RebH and 4V activity on Trp-containing peptides with an N-terminal Gly_n_ extension (n = 1-5 glycines). Center, high-resolution mass spectrum of GG-7-BrW-CONH_2_ generated by 4V-catalyzed bromination. Right, heatmap summarizing percent conversion of Gly_n_-Trp peptides to Gly_n_-Br-Trp peptides. (b) Left, RebH and 4V activity on Trp-containing peptides with a C-terminal Gly_n_ extension (n = 1-5 glycines). Center, high-resolution mass spectrum of 7-BrW-GG-CONH_2_ generated by 4V-catalyzed bromination. Right, heatmap summarizing percent conversion of Trp-Gly_n_ peptides to Br-Trp-Gly_n_ peptides. (c) Left, RebH and 4V activity on Trp-containing peptides with a central Trp residue flanked by one or two glycines on each side. Center, high-resolution mass spectrum of G-7-BrW-G-CONH_2_ generated by 4V-catalyzed bromination. Right, heatmap summarizing percent conversion of central Trp Gly peptides to the brominated form.

**Figure 3.**
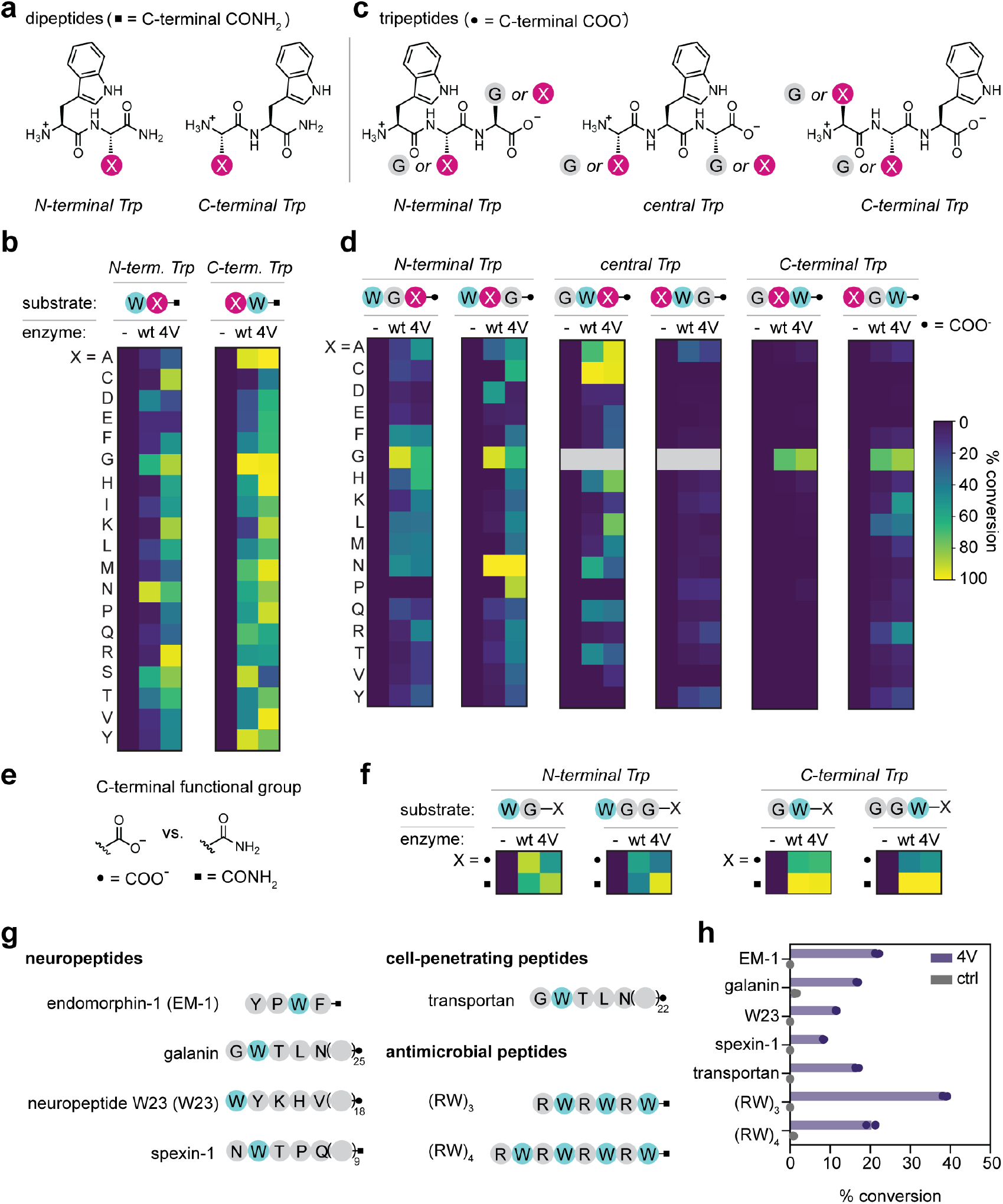
Sequence and structural determinants of RebH and 4V activity on peptides. (a) Design of dipeptide libraries with Trp at the N (left) or C (right) terminus. (b) Heatmap summarizing RebH and 4V activity on N- and C-terminal Trp dipeptide libraries. Heatmap colors represent the mean value (n=3). (c) Tripeptide library designs. Tripeptides had Trp at the N-terminal, central, or C-terminal position, with one position fixed as Gly and the other position varied to 17 other amino acids. (d) Heatmap summarizing RebH and 4V activity on tripeptide libraries. Heatmap colors represent the mean value (n=3). (e) C-terminal functional groups present in the di- and tri-peptide libraries. (f) Dependence of RebH and 4V activity on C-terminal functional group. Heatmap colors represent the mean value (n=3). (g) Trp-containing biologically active peptides that were tested as 4V substrates. (h) 4V activity on biologically active peptides (n =3 independent experiments).

**Figure 4.**
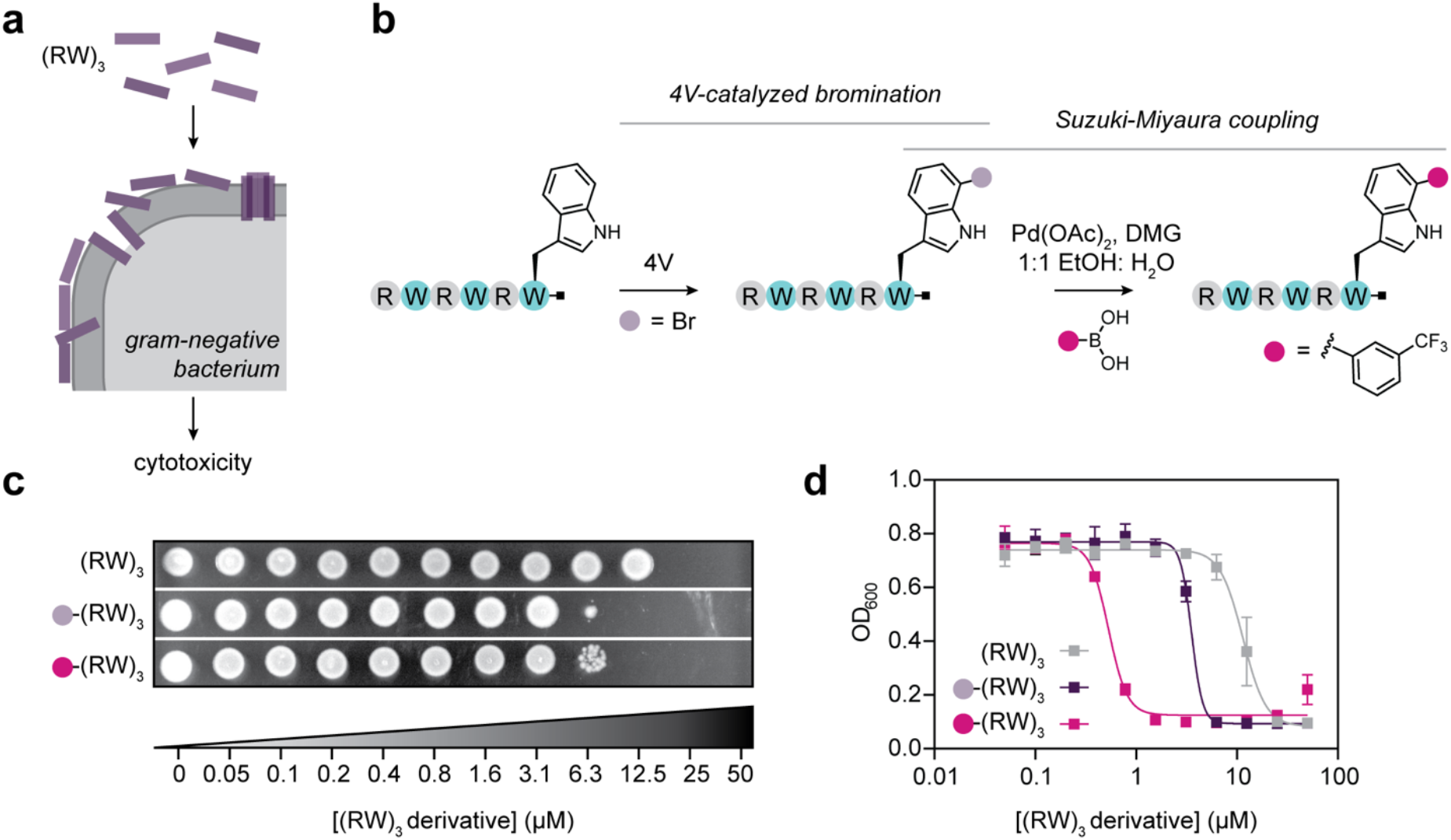
4V enables late-stage diversification of antimicrobial peptides to tune their activity. (a) (RW)_3_ is an antimicrobial peptide that disrupts the membrane of gram-negative bacteria. (b) Strategy for late-stage modification of (RW)_3_ via 4V-catalyzed bromination followed by Suzuki-Miyaura coupling. (c) Minimum bactericidal concentration assays for (RW)_3_ (top), Br-(RW)_3_ (middle), and 3-trifluoromethylphenyl (RW)_3_ (bottom). (d) IC_50_ measurements for (RW)_3_ (grey), Br-(RW)_3_ (purple), and 3-trifluoromethylphenyl-(RW)_3_ (magenta).

**Figure 5.**
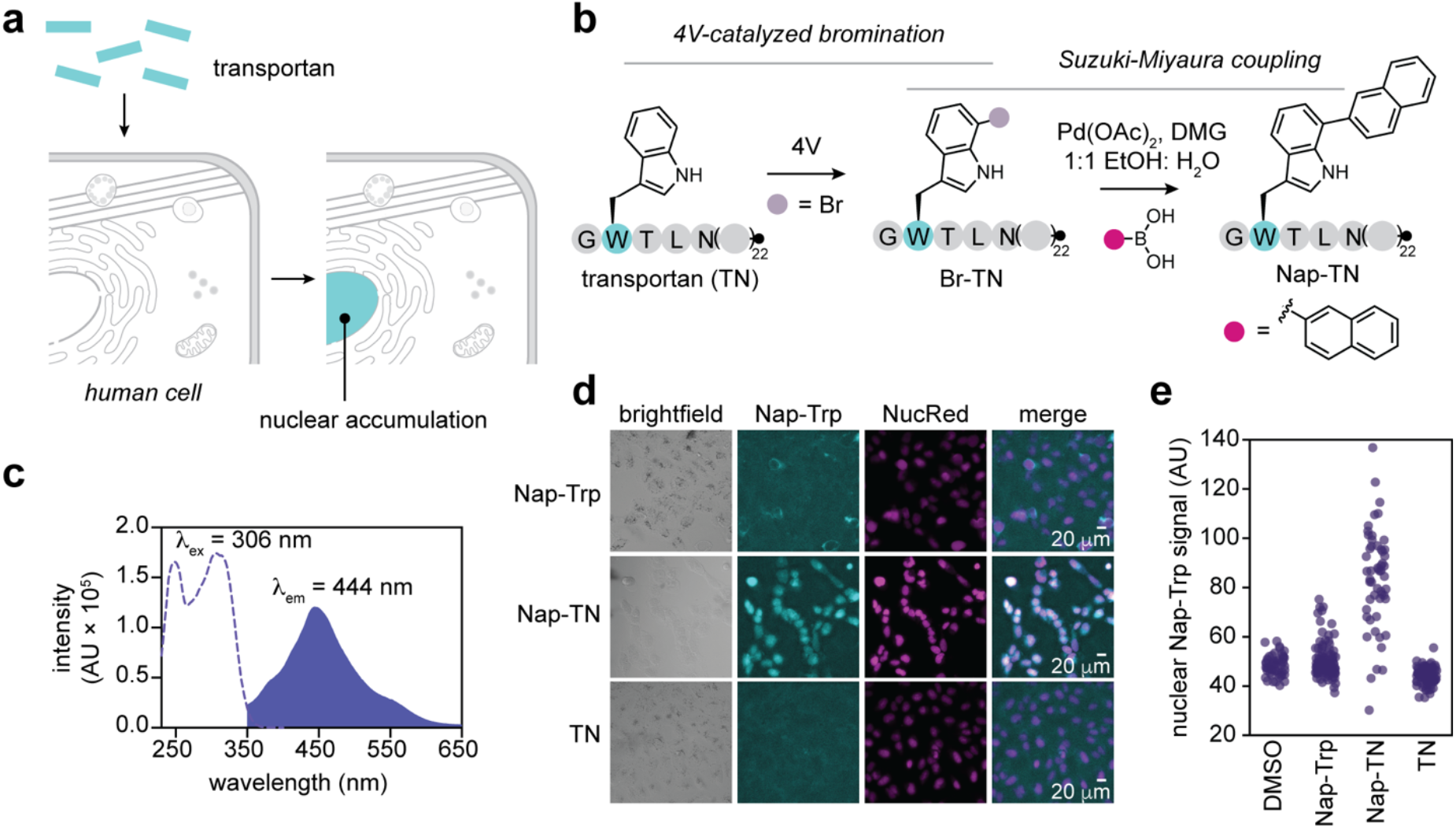
4V-catalyzed peptide functionalization to generate a fluorescent probe for cellular peptide uptake. (a) Transportan is a cell-penetrating peptide that accumulates in the nucleus. (b) Strategy for synthesis of naphthyl-transportan (Nap-TN) via 4V-catalyzed bromination and Suzuki-Miyaura coupling. (c) Excitation (dotted line) and emission (filled curve) spectra for the Nap-Trp fluorophore. (d) Fluorescence imaging of Nap-TN uptake into mammalian cells. Nuclear accumulation of Nap-Trp signal is only observed when the Nap-Trp fluorophore is incorporated into transportan (second row, cyan). (e) Quantification of nuclear Nap-Trp signal intensity for DMSO, Nap-Trp, Nap-TN, and TN.

When Trp was placed at the peptide C terminus, RebH and 4V exhibited similar trends in activity (**Fig. 2a**). Both enzymes gave approximately 100% conversion for GW and GGW peptides, but 4V produced a substantial amount of dibrominated product (17±7% for GW and 26 ±13% for GGW), while RebH produced <5% dibrominated product (**Supplementary Fig. 17**). Activity decreased on peptides with 3-5 Gly residues appended at the N terminus, leveling off at 50% conversion for Gly_5_-W for both enzymes, with little (<5%) dibromination activity observed for these peptides. However, 4V retained higher activity on Gly_3_-W and Gly_4_-W peptides compared to RebH, suggesting that its ability to accommodate larger substrates improves its activity on longer C-terminal Trp peptides. When Trp was placed at the peptide N terminus (**Fig. 2b, Supplementary Fig. 17**), both 4V and RebH had generally lower activity, with a trend of decreasing activity on longer peptides. While 4V gave higher conversion on the WG peptide, no substantial differences in activity between RebH and 4V were observed on longer peptides. For both enzymes, there was a steep dropoff in activity when two or three glycines were appended to the C terminus, but no further diminishment in activity was observed when four or five glycines were added.

The observation that RebH and 4V activity level off after addition of three glycine residues on the C terminus or four residues on the N terminus suggests that these glycines must be accommodated within the enzyme, and that additional residues protrude from the active site and do not affect activity. To explore this hypothesis, we performed flexible peptide docking using FlexPepDock^46^ with a crystal structure of RebH (PDB ID: 2E4G)^47^. The results from docking WGGG show the C terminus near the edge of the RebH structure, providing support for this hypothesis (**Supplementary Fig. 18**). The modeled peptide pose closely resembles that observed in a crystal structure of D-WS bound to the tryptophan 6-halogenase ThaI^35^, supporting the validity of the model.

We next sought to evaluate RebH and 4V activity on peptides containing a central Trp residue (**Fig. 2c**). We treated peptides with Trp flanked by one or two glycine residues on each side with RebH or 4V and measured conversion to brominated product by LC-HRMS. Both enzymes efficiently brominated these substrates, with conversions exceeding 80% and monobromination >95% (**Supplementary Fig. 4, Supplementary Fig. 19**). These results extend the known substrate scope of RebH and 4V to include non-terminal Trp residues, a capability that remains rare among FDHs.

Our results demonstrate that both RebH and 4V possess substantial bromination activity on a variety of Trp-containing peptide substrates as measured by high-resolution LC-MS and confirmed by 1D ^1^H and 2D COSY, TOCSY, and NOESY NMR experiments. While a previous study reported that RebH lacks detectable activity on peptide substrates^36^, differences in the experimental design may account for this variation. For example, the prior work did not coexpress RebH with the GroES/GroEL chaperonin system, used different buffer conditions, and evaluated chlorination rather than bromination. Our findings indicate that under optimized conditions, both RebH and 4V exhibit robust peptide halogenation activity, potentially expanding their utility for late-stage peptide modification.

### Impact of substrate peptide sequence and structural features on RebH and 4V activity

To examine how peptide substrate features influence RebH and 4V activity, we synthesized a panel of 38 dipeptides containing Trp at either the N or C terminus, paired with each of the 19 other standard amino acids (**Fig. 3a**). For synthetic convenience, all peptides were prepared with C-terminal carboxamide groups. Trp-Trp was excluded from the dipeptide panel. We treated each of the peptides individually with RebH or 4V for 20 h and then assessed conversion to the brominated product using high-resolution LC-MS (**Fig. 3b, Supplementary Fig. 20, Supplementary Fig. 21**).

For the N-terminal Trp dipeptides, RebH exhibited >20% conversion on only 5 of 19 potential substrates (**Fig. 3b**, left). These dipeptides (WD, WG, WN, WS, and WT) all have a small hydrophilic side chain in the C-terminal position, except Gly. These data suggest that there are constraints on the side chain size and polarity that can be accommodated by RebH. In contrast, 4V had much broader activity on N-terminal Trp dipeptides, with >20% conversion observed on all sequences except WE (**Fig. 3b**, left). For peptides with Trp at the C terminus (**Fig. 3b**, right), RebH catalyzed >20% conversion on 17 of the 19 potential substrates, with little to no activity observed on CW and KW. 4V brominated all 19 dipeptides at >20% conversion and displayed increased activity on 15 of 19 dipeptides compared to their N-terminal Trp counterparts and to RebH. Both RebH and 4V harbor two hydrogen bonding residues (N467 and N470 in RebH and T467 and S470 in 4V) that do not interact with free Trp but are ideally positioned to engage the C-terminal residue of dipeptide substrates, providing a structural rationale for the broader sequence scope of RebH and 4V for C-terminal Trp peptides (**Supplementary Fig. 22**).

We next sought to further probe the substrate scope of RebH and 4V using a panel of 98 Trp-containing tripeptides to more comprehensively determine the sequences that are preferred by each enzyme (**Fig. 3c**). To reflect the most common C-terminal functional group in naturally occurring peptides and proteins, these peptides were synthesized with a C-terminal carboxylate group. The tripeptide panel consisted of sequences with Trp fixed at either terminus or in the center of the peptide. A second position was held constant as Gly, while the third position was varied to 17 other amino acids, with Trp omitted to avoid ambiguity in localizing enzyme-catalyzed modifications, Ile omitted based on its similarity to Leu, and Ser omitted based on its chemical similarity to Thr. Based on its sequence diversity, this panel enabled us to assess the impact of residue size, charge, and hydrogen bonding ability of the substrate on RebH and 4V activity. A general analysis of substrate halogenation by each enzyme revealed that RebH had detectable activity (≥0.08%) on 76/98 (77.6%) of tripeptides, while 4V had detectable activity (≥0.18%) on 84/98 (85.7%) of the tripeptides (**Fig. 3d, Supplementary Fig. 23-25**). The amount of conversion observed for each peptide varied based on sequence features as described below.

We first examined the group of peptides in which Trp was fixed at the N terminus (**Fig. 3d**, left). In the first set of N-terminal Trp peptides, Gly was fixed at the second position, and the C-terminal position was varied to 17 other amino acids. Both RebH and 4V displayed broad but low-to-moderate (2.5-56% for RebH; 2.0-40% for 4V) conversion of 16/17 peptides in this group. For N-terminal Trp peptides with a variable second position and a fixed C-terminal Gly residue (**Fig. 3d**, left), there was a similar trend, with broad activity but low-to-moderate (1.3-98% for RebH; 2.8-68% for 4V) conversion observed for 13/17 peptides with RebH and 17/17 peptides with 4V, with elevated activity observed on select WXG tripeptides. Notably, 4V catalyzed 68% conversion of WNG to the brominated peptide, while RebH converted >98% of this peptide to the brominated form. The increased activity of both enzymes on this substrate can be rationalized based on hydrogen bonding residues near the active site that are positioned to interact with the substrate peptide. In a model of RebH bound to WNG, N467 and N470 are poised to hydrogen bond with the Asn side chain of the substrate. In contrast to RebH, 4V harbors N467T and N470S mutations and makes only one hydrogen bond to the Asn side chain of the substrate, potentially explaining the lower extent of 4V-catalyzed WNG bromination. RebH also exhibited higher activity on the WDG peptide compared to other N-terminal Trp peptides, suggesting that similar hydrogen bonding interactions might support higher activity on this peptide. Extension of the central side chain by one carbon in the WQG and WEG peptides dramatically reduced the observed conversion, suggesting that both side chain length and polarity are important for peptide substrate recognition.

We next evaluated the group of tripeptides with Trp in the center, with Gly fixed at either the N terminus or C terminus and the remaining position varied (**Fig. 3d**, center). This group is of particular interest because ThaI is the only Trp halogenase tool among the many characterized enzymes for which activity on peptides with a non-terminal Trp has been examined. For the peptides with a GWX motif, RebH displayed >20% activity on 6/16 peptides, while 4V halogenated 10/16 peptides to >20% conversion. Only GWP was not detectably halogenated by either enzyme. Peptides with an XWG motif are poor substrates for both RebH and 4V, which had nearly identical activity profiles for substrates in this group. Both RebH and 4V showed detectable activity on 14/16 peptides, but only AWG, KWG, PWG, RWG, and YWG showed >5% conversion, with 4V also catalyzing 5.5% bromination of HWG.

The final group of tripeptides that we examined had Trp fixed at the C terminus, with Gly fixed in either the first or second position and the remaining position varied to 17 other amino acids (**Fig. 3d**, right). Peptides with GXW sequences were the worst group of substrates for both RebH and 4V, with <3% conversion observed for all sequences except GGW. This result is consistent with a prior study of ThaI activity on peptides that showed that C-terminal Trp peptides were not favorable substrates for this enzyme. Peptides with Gly fixed at the second position with a variable first position were also generally poor substrates for both RebH and 4V. RebH catalyzed detectable (0.1-18%) conversion of 14/17 peptides, but only 3 peptides reached >5% conversion. While 4V also had low activity on peptides within this series, it had an expanded substrate scope compared to RebH, catalyzing detectable (0.2-31%) conversion of 16/17 peptides, with 11/17 peptides brominated at >5% conversion.

Dipeptides with C-terminal Trp exhibited the highest conversion as a group among all of the peptides that we tested. We were therefore surprised to find that extending these peptides by one N-terminal Gly residue severely diminished RebH and 4V bromination activity. In contrast, extending dipeptide sequences with N-terminal Trp by one C-terminal Gly residue had a more modest impact on activity. We wondered whether the difference in C-terminal functional group between the dipeptide panel (C-terminal carboxamides) and the tripeptide panel (C-terminal carboxylates) might account for this difference. To examine the effect of the C-terminal functional group, we tested a panel of sequence-matched Gly-containing Trp peptides with either a carboxylate or a carboxamide at the C terminus (**Fig. 3e, f, Supplementary Fig. 26**). Our data show that when Trp is in the C-terminal position, RebH prefers the carboxamide over the sequence-matched carboxylate peptide (**Fig. 3f**, right). Conversely, for peptides with N-terminal Trp, RebH prefers the C-terminal carboxylate over the carboxamide (**Fig. 3f**, left). In contrast to RebH, 4V prefers peptides with a carboxamide over a carboxylate regardless of the location of Trp within the peptide (**Fig. 3f**). This strong bias of 4V for carboxamide-containing Trp peptides could be due to the presence of two substituted residues within the 4V substrate binding site. In 4V, two Asn residues within 7Å of the carboxylate group of L-Trp (N467 and N470) are substituted to a Thr and Ser residue, respectively (**Supplementary Fig. 22**). These mutations alter the hydrogen bonding characteristics of the substrate binding pocket, potentially leading to a more favorable interaction with peptides containing a C-terminal carboxamide.

### 4V catalyzes late-stage bromination of unmodified bioactive peptides

The breadth of activity that we observed across short peptide substrates suggested that 4V may be able to tolerate additional sequence and structural complexity. Trp residues are common in bioactive peptides and selective Trp bromination could provide a handle for Pd-catalyzed cross-coupling reactions that are already commonly deployed in pharmaceutical synthesis and drug discovery^21,22^. We therefore evaluated whether these enzymes could act on a set of more complex bioactive peptides containing intrinsic Trp residues without any sequence redesign or modification (**Fig. 3g**). Informed by the results of our tripeptide substrate screen, we selected seven commercially available bioactive peptides varying in length from 4-30 amino acids that contained one or more Trp residues in N-terminal, C-terminal, or internal positions. These included GPCR ligands (endomorphin-1, galanin, spexin-1, and neuropeptide W23), a cell-penetrating peptide (transportan), and two antimicrobial peptides ((RW)_3_ and (RW)_4_).

Despite the challenging nature of these peptides, 4V had activity on all of the peptides tested, catalyzing modest 8-38% conversion to the monobrominated product depending on peptide identity (**Fig. 3h**). In line with our previous results that suggested that N- and C-terminal extensions longer than four amino acids protrude from the enzyme active site, there was no clear length dependence to the extent of bromination catalyzed by 4V. Our model peptide results also suggested that 4V generally has higher activity on peptides with a C-terminal carboxamide, and the bioactive peptides generally followed this trend, with the highest conversion observed for (RW)_3_ (38.7±0.6%), endomorphin-1 (21.7±0.5%), and (RW)_4_ (21±1%), all of which have a C-terminal carboxamide. However, the presence of a C-terminal carboxamide was not sufficient to explain the trends in 4V activity, which also displayed substantial dependence on peptide sequence.

The highest conversion (38.7±0.6%), was observed on (RW)_3_, which contains both the RW and WR motifs that were favorable substrates in the dipeptide activity screen and the WXG/XWG tripeptide activity screens. (RW)_4_, which has the same sequence motifs, exhibited somewhat lower conversion (21±1%), but was among the more favored substrates among the bioactive peptide panel. Endomorphin-1, which was converted to a similar extent (21.7±0.5%), also contains motifs that we identified as favorable in our dipeptide (WF, PW) and tripeptide (XWF, PWX) screens. Galanin and transportan share the same 12 N-terminal residues with a favorable N-terminal GWT motif and displayed the same extent of conversion (16.8±0.2% for galanin and 16.6±0.5% for transportan) despite different overall lengths and C-terminal sequences. This result suggests that the sequence proximal to the modified Trp is a key determinant of 4V activity. In agreement with this sequence dependence, neuropeptide W (N-terminal WYK) and spexin-1 (N-terminal NWT), which contain the least favorable motifs, had the lowest 4V-catalyzed conversion at 11.5±0.1% and 8.3±0.1%, respectively.

Evaluating the transformation that 4V catalyzed on the (RW)_3_ and (RW)_4_ peptides enabled us to examine another key parameter: Trp site selectivity in a peptide with multiple Trp residues. Both peptides were converted by 4V to monobrominated products, raising the question of which Trp residue is modified. To localize the bromine modification, we performed a trypsin digestion of the (RW)_3_ 4V bromination reactions and analyzed the digest fragments using LC-HRMS (**Supplementary Fig. 27**). The digestion produced a 204.1131 [M+H^+^] ion that can only be assigned to the C-terminal tryptophanamide residue, which differs in mass from Trp by 0.98 Da. This single-Trp selectivity based on sequence and structural features highlights the potential utility of application of 4V for site-selective bromination.

### 4V enables late-stage diversification of antimicrobial peptides to tune their activity

Arg- and Trp-rich peptides have broad-spectrum antimicrobial activity due to their direct interaction with the plasma membrane (**Fig. 4a**). Based on 4V’s ability to brominate (RW)_3_, we set out to use 4V-catalyzed bromination followed by Suzuki-Miyaura coupling to structurally diversify this peptide (**Fig. 4b**). We hypothesized that addition of a bromine atom to these peptides would lead to increased antimicrobial activity as halogens are commonly introduced into drug molecules to enhance membrane permeability^48^. We also predicted that introduction of a 3-trifluoromethylphenyl group would further improve activity based on its hydrophobicity and prevalence in drugs^49^. To test these hypotheses, we prepared the monobrominated (RW)_3_ peptide and the 3-trifluoromethylphenyl derivative, purified them to >90% by HPLC (**Supplementary Fig. 28, 29**), and tested their ability to inhibit the growth of *E. coli*.

To evaluate the effects of (RW)_3_ bromination and 3-trifluoromethylphenyl modification on antimicrobial activity, we performed an 18 h *E. coli* XL10 growth inhibition assay across varying peptide concentrations and determined the minimal inhibitory concentration (MIC), minimal bactericidal concentration (MBC), and half-maximal inhibitory concentration (IC_50_) for each derivative (**Fig. 4c, d, Supplementary Fig. 30, 31**). Unmodified (RW)_3_ displayed an MIC of 25 µM, an MBC of 50 µM (**Fig. 4c**), and an IC_50_ of 11.5 ± 0.5 µM (**Fig. 4d**), consistent with the previously reported IC_50_ of 16 µM. Bromination of (RW)_3_ led to approximately four-fold decreases in MIC (6.25 µM), MBC (12.5 µM), and IC_50_ (3.5 ± 0.3 µM) (**Fig. 4c, d, Supplementary Fig. 30, 31**). Subsequent conversion of Br-(RW)_3_ to CF_3_(-RW)_3_ resulted in a 16-fold decrease in MIC (1.6 µM) (**Supplementary Fig. 30, 31**) and IC_50_ (0.5 ± 0.3 µM) (**Fig. 4d**) and a four-fold decrease in MBC (12.5 µM) (**Fig. 4c**) relative to the parent peptide. These marked enhancements in antimicrobial potency demonstrate the effectiveness of this late-stage functionalization strategy and align with previous observations that hydrophobic modification of (RW)_3_ increases its bioactivity.

To test whether the same trends of improved antimicrobial activity extended to longer Arg/Trp-rich peptides, we assessed the impact of (RW)_4_ and Br-(RW)_4_ on *E. coli* growth (**Supplementary Fig. 32**). For (RW)_4_, we observed an MIC of 0.8 µM and an MBC of 0.8 µM, consistent with previous reports showing greater potency of (RW)_4_ relative to (RW)_350_. Bromination of (RW)_4_ resulted in approximately a two-fold increase in antimicrobial activity, with an MIC of 0.4 µM and an MBC of 0.4 µM. Together, these results demonstrate the utility of 4V-catalyzed bromination followed by Suzuki-Miyaura coupling as a general strategy for late-stage diversification to tune the activity of bioactive peptides.

### 4V-catalyzed peptide functionalization to generate a fluorescent probe for cellular peptide uptake

Cell-penetrating peptides (CPPs) including transportan are widely used to deliver cargo into mammalian cells (**Fig. 5a**)^51,52^. Fluorescent labeling is commonly used to visualize CPP uptake, but conventional labeling strategies typically target Lys residues or the N terminus, sites whose positive charge is important for membrane translocation^53^. We therefore tested whether 4V-catalyzed bromination of transportan followed by Suzuki-Miyaura coupling could enable direct fluorescent labeling of a Trp residue without altering the peptide’s charge (**Fig. 5b**).

Using this approach, we brominated Trp 2 of transportan and coupled it to naphthalene-2-boronic acid to generate a naphthyl-modified peptide (Nap-transportan, Nap-TN, **Supplementary Fig. 33**). The Nap-Trp fluorophore exhibited an excitation maximum of 306 nm and an emission maximum of 444 nm (**Fig. 5c**), allowing Nap-TN imaging using standard DAPI (blue) channel filter sets. To assess whether this derivative could be used to visualize cellular uptake, we incubated human U2OS cells with Nap-TN for 20 min in the presence of a far-red nuclear stain (NucRed) (**Fig. 5d**). Under these conditions, we observed intracellular blue fluorescence that colocalized with the nucleus. No intracellular blue fluorescence was observed for DMSO or unmodified transportan, and a control compound lacking CPP properties (Nap-Trp) also showed no detectable uptake. Quantification of nuclear Nap-Trp signal demonstrated a substantial increase when cells were incubated with Nap-TN, but not DMSO, TN, or Nap-Trp (**Fig. 5e**). These results demonstrate that 4V-enabled Trp functionalization can convert a native CPP into an imaging probe without compromising its biological function. Notably, 432 of 1855 CPPs deposited in the CPP database CPPsite 2.0^54^ contain at least one Trp residue, suggesting that this approach has the potential for broad applicability.

### 4V-catalyzed diversification of a µ opioid receptor agonist to modulate its activity

Endomorphin-1 (YPWF) is a natural neuropeptide agonist of the µ opioid receptor, a G protein-coupled receptor (GPCR) that is the target of most opioids used clinically and non-medically (**Fig. 6a**)^17,55^. We hypothesized that structural diversification at Trp 3 of endomorphin-1 via 4V-catalyzed bromination followed by Suzuki-Miyaura coupling to a panel of aryl boronic acids would enable modulation of endomorphin-1 activity. Using this approach, we prepared a panel of six endomorphin-1 derivatives (**Supplementary Fig. 34, 35**). We found that, similar to the FDH ThaI^35^, 4V retains activity on D-Trp, enabling us to prepare a corresponding panel of six stereochemical controls from an endomorphin-1 variant in which Trp 3 was replaced with D-Trp ([D-Trp 3]endomorphin-1) (**Fig. 6b, Supplementary Fig. 36, 37**).

**Figure 6.**
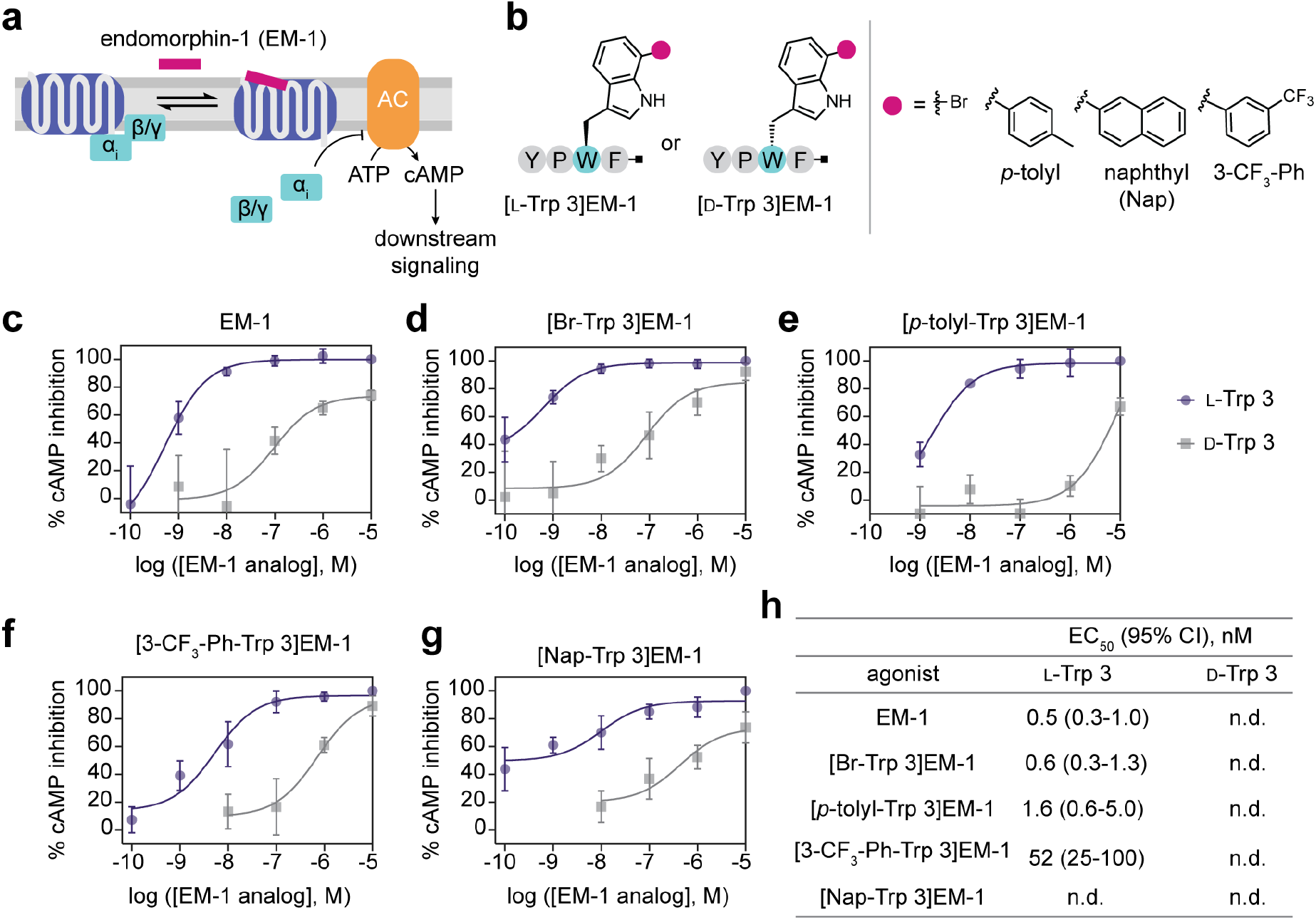
4V-catalyzed diversification of a µ opioid receptor agonist to modulate its activity. (a) Endomorphin-1 (EM-1) is an agonist of µ opioid receptor, which signals through adenylyl-cyclase inhibitory G proteins (G_i_) to inhibit cAMP production. (b) [L-Trp 3]EM-1 and [D-Trp 3]EM-1 analog designs. (c) Dose-response curve for EM-1. (d) Dose-response curve for [Br-Trp 3]EM-1. (e) Dose-response curve for [p-tolyl-Trp 3]EM-1. (f) Dose-response curve for [3-trifluoromethylphenyl-Trp 3]EM-1. (g) Dose-response curve for [naphthyl-Trp 3]EM-1. (h) Summary of EC_50_ values for EM-1 analogs. EC_50_s were determined by fitting the inhibitory portion of the dose response curve for four independent experiments.

The µOR signals primarily through adenylyl-cyclase inhibitory G proteins (G_i_) (**Fig. 6a**)^55^. We therefore assessed the bioactivity of endomorphin-1 derivatives by evaluating their ability to inhibit cyclic AMP (cAMP) production in HEK293T cells expressing µOR (**Fig. 6c-h, Supplementary Fig. 38-40**). Native [L-Trp 3]endomorphin-1 exhibited an EC_50_ value of 0.5 nM (95% CI: 0.3-1.0 nM) (**Fig. 6c**). All L-Trp 3-based analogs generated quantifiable dose-response curves, whereas the corresponding D-Trp 3 analogs produced weak signaling insufficient for reliable EC_50_ determination. While the [Br-Trp 3]EM-1 analog displayed similar potency to unmodified EM-1 (**Fig. 6d**), analogs that were further functionalized by Suzuki-Miyaura coupling displayed reduced potency that appeared to correlate with the size of the functional group that was introduced (**Fig. 6e-h**). Overall, our results demonstrate that functionalization of EM-1 at Trp 3 can modulate its GPCR agonist activity and highlight a facile way to access an analog series of bioactive peptides along with matched stereochemical controls.

## Discussion

Peptides are an important class of therapeutic molecules whose bioactivity can be tuned through alterations in chemical structure^56,57^. While side chain modification of peptide therapeutics is typically achieved by incorporation of modified building blocks during solid-phase peptide synthesis, enzymatic late-stage modification represents an attractive complementary strategy. Although several natural FDHs function on peptide substrates, peptide activity often requires a specific leader sequence or tag, limiting the utility of these enzymes for late-stage functionalization of peptides^35–37^. Here, we explored the peptide activity of the Trp 7-halogenase RebH, among the most extensively engineered FDHs, on peptides. Contrary to previous reports, we find that RebH has robust activity on peptides, with its efficiency modulated by peptide sequence and structural features. Because many RebH variants have been engineered previously^39^, the newly revealed peptide activity of this enzyme provides a uniquely tractable platform for the engineering of peptide-selective halogenases and opens up the application of existing RebH variants to peptides.

The discovery of peptide activity in RebH motivated us to examine the RebH variant 4V, which was engineered by the Lewis lab to accommodate larger substrates^43^. We reasoned that 4V might have enhanced activity on peptides based on its expanded substrate scope. By performing a large-scale screen of 4V activity on di- and tripeptide substrates, we defined the sequence tolerance of both RebH and 4V, finding that 4V generally produced higher conversion and had an expanded sequence scope compared to RebH. These results establish RebH and its engineered variants as a practical platform for extending FDH catalysis to peptide substrates despite the parent enzyme’s native role in free amino acid halogenation.

Comparison of the sequences and structures of RebH and 4V suggests that a substrate-binding lid and adjacent loops are important determinants of sequence tolerance. Two hydrogen-bonding residues, N467 and N470 in RebH and T467 and S470 in 4V, are positioned to interact with the C terminus of an extended substrate. In RebH, these residues also participate in a hydrogen-bonding network that appears to stabilize the substrate-binding lid in a closed conformation^47^. Substitutions at these positions in 4V may therefore permit additional conformational flexibility that could explain the expanded substrate tolerance we observe. More broadly, conformational changes within the active site are likely important for binding peptide substrates as RebH possesses a relatively restricted substrate pocket (calculated volume of 44.3 Å^3^ by CASTpFold^58^) compared to FDHs that natively act on linear peptides such as ChlH (442 Å^3^)^38^.

We applied 4V in combination with Suzuki-Miyaura coupling for late-stage chemoenzymatic functionalization of unmodified bioactive peptides. Although conversion was modest on many of the substrates, this strategy did not require the introduction of protecting groups or tags and provided sufficient material for downstream biological assays. Using this chemoenzymatic strategy, we tuned the biological activity of antimicrobial peptides and GPCR agonists and produced a functional cell-penetrating peptide that was modified to enable fluorescence imaging. Although we focused on Suzuki-Miyaura coupling, previous work has shown that aryl bromides generated by RebH are versatile chemical handles that enable functionalization using diverse cross-coupling strategies including Buchwald-Hartwig amination and alkoxylation^27,59,60^. Building on this foundation, our strategy therefore has the potential to enable peptide diversification via formation of C-C-, C-N and C-O bonds. Together, our results expand the conceptual and practical scope of FDH-catalyzed halogenation for peptide chemistry and chemical biology.

## Supporting Information

Supplementary Information (PDF): Methods, Supplementary Figures 1 – 41, Supplementary Tables 1-3.

Supplementary Appendix (PDF): LC-MS data for glycine peptide, dipeptide, and tripeptide bioconversion reactions.

Supplementary Dataset (XLSX): LC-MS product and reactant peak areas for enzymatic bromination of glycine peptides, dipeptides, tripeptides, and bioactive peptides.

## Acknowledgements

We thank S. Coyle, D. Sashital, T. Galateo, and members of the Weeks lab for helpful discussions. This work was supported in part by startup funds from the University of Wisconsin-Madison Department of Biochemistry and by an NIH Director’s New Innovator Award (DP2GM149548) to A.M.W. H.N.B was supported in part by a William R. & Dorothy E. Sullivan Wisconsin Distinguished Graduate Fellowship. This study made use of the National Magnetic Resonance Facility at Madison, which is supported by NIH grant R24GM141526. Helium recovery equipment, computers, and infrastructure for data archive were funded by the University of Wisconsin-Madison, NIH R24GM141526, and National Science Foundation NSF 1946970 (NSF Mid-Scale Research Infrastructure Big Idea). Instrumentation was funded by an NIH Shared Instrumentation Grant (S10RR023438). We thank Paulo Falco Cobra for assistance with NMR data collection and analysis.

