## Supporting Information for "Enzymatic bromination of native peptides for late-stage structural diversification via Suzuki-Miyaura coupling"

**Affiliation**

   1.4 Protein expression and purification…………………………………………………...8
   1.5 Solid-phase peptide synthesis………………………………………………………..10
   1.6 Analytical scale RebH reactions……………………………………………………...10
   1.7 Preparative scale RebH reaction……………………………………………………..10
   1.8 Trypsin digestion of brominated bioactive peptides…………………………………...11
   1.9 Suzuki-Miyaura coupling of brominated bioactive peptides…………………………...11
   1.10. μOR GloSensor assay…………………………………………………………………11
   1.11. Immunofluorescence for μOR transfected HEK293T cells…………………………..12
   1.12 Transportan localization assay in U2OS cells………………………………………...12
   1.13 Bacterial minimal inhibitory concentration measurements…………………………..13
   1.14 Bacterial $IC_{50}$ determination……………………………………………………….…13
   1.15 Minimal bactericidal concentration on petri dishes………………………………………13
   1.16 Minimal bactericidal concentration spot assay…………………………………………...13
2. **Supplementary Figures and Tables**…………………………………………………………15
3. **References**…………………………………………………………………………………..59

### 1. Methods

#### 1.1 Materials and reagents.

| REAGENT or RESOURCE | SOURCE | IDENTIFIER |
|---|---|---|
| **Chemical Reagents** | | |
| Sodium bromide | Sigma Aldrich | S-9756 |
| Sodium chloride | Fisher Scientific | S640-10 |
| HEPES | Fisher Scientific | AC172570010 |
| Imidazole | Sigma Aldrich | I5513 |
| Sodium phosphate monobasic dihydrate | Fisher Scientific | S381-500 |
| Sodium phosphate dibasic heptahydrate | Fisher Scientific | S373-500 |
| Dimethyl Sulfoxide | Fisher Scientific | BP2311 |
| Chaperone Plasmid Set | TaKaRa Bio | 3340 |
| Glycerol | Fisher Scientific | G33-4 |
| L-(+)-Arabinose | Sigma-Aldrich | A91906-100G-A |
| Isopropyl β-d-1-thiogalactopyranoside (IPTG) | Goldbio | I2481C100 |
| Kanamycin | Goldbio | K-120-50 |
| Chloramphenicol | Goldbio | C-105-25 |
| Carbenicillin | Goldbio | C-103-100 |
| Sodium hydroxide | Sigma-Aldrich | 415413-500ML |
| Hydrochloric acid | Sigma-Aldrich | 320331-500ml |
| cOmplete protease inhibitor cocktail | Sigma-Aldrich | 11873580001 |
| Flavin adenine dinucleotide (FAD) | Sigma Aldrich | F8384-100MG |
| beta-Nicotinamide adenine dinucleotide, reduced disodium salt hydrate | Sigma Aldrich | N8129-100MG |
| L-Tryptophan | Sigma Aldrich | T0254-1G |
| D-Tryptophan | Sigma Aldrich | T9753-25G |
| P-Tolylboronic Acid | Chem Impex International | CMX-30418-5G |
| 2-Naphthaleneboronic acid | VWR International | 77085-354 (EA) |
| Phenylboronic acid | Sigma Aldrich | P20009-10G |
| Pyrene-1-boronic acid | VWR International | AAH64211-03 (EA) |
| 3-(Trifluoromethyl)phenylboronic acid | Sigma Aldrich | 432032-1G |
| Rink Amide AM Resin (100-200 MESH) | Sigma Aldrich | 8551300025 |
| Diethyl Ether | Sigma Aldrich | 676845-1L |
| N,N-Dimethylformamide | Sigma Aldrich | 648531-4X4L |
| Methylene Chloride | Fisher Scientific | D37-4 |
| Triisopropylsilane | Sigma Aldrich | 233781 |
| N,N-diisopropylcarbodiimide | VWR International | D3183 |
| 1-Hydroxybenzotriazole hydrate | Chem-Impex | 24755 |
| Nalpha-Fmoc-Nin-Boc-D-Tryptophan | Chem Impex International | 2484 |
| Fmoc-L-alanine | Chem Impex International | 02369 |
| Fmoc-S-trityl-L-cysteine | Chem Impex International | 00177 |
| Fmoc-L-aspartic acid beta-tert-butyl ester | Chem Impex International | 00494 |
| Fmoc-L-glutamic acid gamma-tert-butyl ester hydrate | Chem Impex International | 02413 |
| Fmoc-glycine | Chem Impex International | 02416 |

| | | |
|---|---|---|
| Fmoc-L-phenylalanine | Chem Impex International | 02443 |
| Nalpha-Fmoc-Nim-trityl-L-histidine | Chem Impex International | 00713 |
| Nalpha-Fmoc-Nepsilon-Boc-L-lysine | Chem Impex International | 00493 |
| Fmoc-L-leucine | Chem Impex International | 00145 |
| Fmoc-L-methionine | Chem Impex International | 02436 |
| Fmoc-L-isoleucine | Chem Impex International | 02425 |
| Nalpha-Fmoc-Ngamma-trityl-L-asparagine | Chem Impex International | 02386 |
| Fmoc-L-proline | Chem Impex International | 02448 |
| Fmoc-O-tert-butyl-L-serine | Chem Impex International | 02454 |
| Nalpha-Fmoc- Ndelta-trityl-L-glutamine | Chem Impex International | 02411 |
| Nalpha-Fmoc-Nomega-(2,2,4,6,7-pentamethyldihydro-benzofuran-5-sulfonyl)-L-arginine | Chem Impex International | 01964 |
| Fmoc-O-tert-butyl-L-threonine | Chem Impex International | 02460 |
| Fmoc-L-valine | Chem Impex International | 02470 |
| Nalpha-Fmoc-Nin-Boc-L-tryptophan | Chem Impex International | 01954 |
| Fmoc-O-tert-butyl-L-tyrosine | Chem Impex International | 00495 |
| Fmoc-L-methionine 4-alkoxybenzyl alcohol resin | Chem Impex International | 01913 |
| Fmoc-L-valine 4-alkoxybenzyl alcohol resin, (0.3-0.8 meq/g, 100-200 mesh) | Chem Impex International | 01770 |
| Fmoc-L-isoleucine 4-alkoxybenzyl alcohol resin, (0.3-0.8 meq/g, 100-200 mesh) | Chem Impex International | 01910 |
| Fmoc-L-leucine 4-alkoxybenzyl alcohol resin, (100-200 mesh, 0.3-0.8 meq/g) | Chem Impex International | 01911 |
| Fmoc-L-phenylalanine 4-alkoxybenzyl alcohol resin, (0.3-0.8 meq/g, 100-200 mesh) | Chem Impex International | 01914 |
| Nalpha-Fmoc-L-tryptophan 4-alkoxybenzyl alcohol resin, (0.3-0.8 meq/g, 100-200 mesh) | Chem Impex International | 01918 |
| Nalpha-Fmoc-Nepsilon-Boc-L-lysine 4-alkoxybenzyl alcohol resin, (0.3-0.8 meq/g, 100-200 mesh) | Chem Impex International | 01912 |
| Fmoc-O-tert-butyl-L-serine 4-alkoxybenzyl alcohol resin, (0.3- 0.8 meq/g, 100-200 mesh) | Chem Impex International | 01916 |
| Fmoc-O-tert-butyl-L-tyrosine 4-alkoxybenzyl alcohol resin | Chem Impex International | 01915 |
| Fmoc-O-tert-butyl-L-threonine 4-alkoxybenzyl alcohol resin, (0.3-0.8 meq/g, 100-200 mesh) | Chem Impex International | 01917 |
| Nalpha-Fmoc-Nim-trityl-L-histidine 4-alkoxybenzyl alcohol resin, (0.3-0.8 meq/g, 100-200 mesh) | Chem Impex International | 01909 |
| Na-Fmoc-L-glutamine 4-alkoxybenzyl alcohol resin | Chem Impex International | 01768 |

| | | |
|---|---|---|
| Fmoc-L-glutamic acid gamma-tert-butyl ester 4-alkoxybenzyl alcohol resin, (0.3-0.8 meq/g, 100-200 mesh) | Chem Impex International | 01769 |
| Nalpha-Fmoc-Nomega-(2,2,4,6,7-Pbf-5-sulfonyl)-L-arginine 4-alkoxy-benzyl-alcohol resin, (100-200 mesh, 0.2-0.8 meq/g) | Chem Impex International | 04168 |
| Fmoc-Ala-Wang resin (100-200 mesh) | Chem Impex International | 01903-1G |
| O-(Benzotriazol-1-yl)-N,N,N',N'-tetramethyluronium hexafluorophosphate | Chem Impex International | 02011 |
| Fmoc-Asp(OtBu)-Wang resin (100-200 mesh) | Chem Impex International | 01906-1G |
| Fmoc-L-proline 4-alkoxybenzyl alcohol resin | Chem Impex International | 01749-1G |
| Fmoc-Cys(Trt)-Wang resin (100-200 mesh) | Chem Impex International | 01701-1G |
| Fmoc-Gly-Wang resin (100-200 mesh) | Chem Impex International | 01908-1G |
| Palladium(II) acetate | Sigma Aldrich | 520764-1G |
| Optima Formic acid | Fisher Scientific | A11750 |
| Optima methanol | Fisher Scientific | A4564 |
| Optima acetonitrile | Fisher Scientific | A9554 |
| Optima water | Fisher Scientific | W64 |
| Trifluoroacetic acid | Sigma Aldrich | T6508-1L |
| Agarose LE (Molecular Biology Grade) | Goldbio | A-201-100 |
| 1 kb Plus DNA Ladder | VWR International | 101228-486 |
| GelGreen | Fisher Scientific | NC9728313 |
| Acetic Acid | Sigma Aldrich | A6283-2.5L |
| Coomassie Brilliant Blue G-250 | MP Biomedicals | 808274 |
| PageRuler Plus Protein Ladder | Life Technologies Corporation | 26619 |
| Invitrogen NuPAGE 4 to 12% Bis-Tris Protein Gel | Fisher Scientific | NP0321BOX |
| MES SDS Running Buffer (20X) | Fisher Scientific | NP0002 |
| Pierce™ Quantitative Fluorometric Peptide Assay | Fisher Scientific | PI23290 (EA) |
| Thermo Scientific HisPur Ni-NTA Resin | Life Technologies Corporation | 88222 |
| Deuterium Oxide | Fisher Scientific | NC2958348 |
| 3-(Trimethylsilyl)-1-propanesulfonic acid sodium salt | Sigma Aldrich | sc-238479 |
| **Peptides** | | |
| Endomorphin 1 | Cayman chemical | CAYM-23280-5 |
| Galanin (Human) | Cayman chemical | CAYM-33139-1 |
| Wkymvm | Cayman chemical | CAYM-33589-1 |
| Transportan | Cayman chemical | CAYM-37497-5 |
| Spexin 1 | Cayman chemical | CAYM-36866-1 |
| Neuropeptide W-23 | Cayman chemical | CAYM-24551-1 |
| RW3 | Cayman chemical | CAYM-37493-25 |
| RW4 | Cayman chemical | CAYM-37494-10 |

| | | |
|---|---|---|
| **Recombinant Proteins** | | |
| Catalase From Bovine Liver Lyophilized Powder | Sigma Aldrich | C40-100MG |
| Glucose dehydrogenase | Sigma Aldrich | 19359-10MG-F |
| Dpn1 | NEB | R0176S |
| HindIII-HF | NEB | R0104S |
| NcoI-HF | NEB | R3193S |
| BamHI-HF | NEB | R3136S |
| NotI-HF | NEB | R3189S |
| KOD Hot Start DNA Polymerase | Sigma Aldrich | 71086-3 |
| Trypsin/lys-c mix mass spec grade | Promega | V5113 |
| **Bacterial Cell Culture Reagents** | | |
| Lb Broth Miller | EMD Millipore | 71753-6 |
| Lb agar Miller | Sigma Aldrich | L3027-1KG |
| Terrific broth EZMix powder | Sigma Aldrich | T9179-1KG |
| Mueller Hinton II Broth | Fisher Scientific | B12322 |
| M9 Minimal Salts, 5x | Fisher Scientific | DF0485-17 |
| Glucose | Chem Impex International | 805 |
| $MgSO_4$ | Fisher Scientific | BP213-1 |
| $CaCl_2$ | Fisher Scientific | C79-500 |
| Bacto Dehydrated Agar | Fisher Scientific | 214010 |
| **Mammalian Tissue Culture Reagents** | | |
| SuperBlock PBS blocking buffer | Life Technologies Corporation | 37580 |
| Heparin sulfate | Sigma Aldrich | H3149-100KU |
| NucRed Live 647 ReadyProbes Reagent | Life Technologies Corporation | R37106 |
| NucBlue Fixed Cell ReadyProbes Reagent (DAPI) | Life Technologies Corporation | R37606 |
| Rhodopsin Chimeric Recombinant Mouse Monoclonal Antibody (Rho 1D4) | Life Technologies Corporation | MA547924 |
| Goat anti-Mouse IgG (H+L) Highly Cross-Adsorbed Secondary Antibody, Alexa Fluor™ Plus 488 | Life Technologies Corporation | A32723 |
| pGloSensor-22F cAMP Plasmid | Promega | E2301 |
| GloSensor cAMP Reagent, 25mg | Promega | E1290 |
| Forskolin | Sigma Aldrich | F3917-10MG |
| Dulbecco's Phosphate Buffered Saline | Sigma Aldrich | D8537-6X1L |
| Dulbecco's Modified Eagle Medium with high glucose | Cytiva | SH30243.02 |
| Fetal Bovine Serum | Life Technologies Corporation | A4736401 |
| Penicillin-streptomycin | Cytiva | SV30010 |
| CO2 Independent Medium | Life Technologies Corporation | 18045088 |
| FluoroBrite™ DMEM | Life Technologies Corporation | A1896701 |
| ORPM-1 Duet | Addgene | 213361 |
| PCDNA3-muOR-GFP11-T2A-GFP10-bArrestin1-T2A mCherry | Addgene | 113611 |
| Trypsin-EDTA solution | Sigma Aldrich | T4049-500ML |
| Poly-D-lysine | Life Technologies Corporation | A3890401 |
| Fugene 4K Transfection Reagent | Promega | E5911 |
| **Model Organisms/Cell Lines** | | |

| | | |
|---|---|---|
| HEK293T cell line | ATCC | CRL-3216 |
| U2OS cell line | ATCC | HTB-96 |
| E. coli XL-10 Gold: endA1 glnV44 recA1 thi-1 gyrA96 relA1 lac Hte Δ(mcrA)183 Δ(mcrCB-hsdSMR-mrr)173 $tet^R$ F'[proAB $lacI^qZ\Delta M15$ Tn10($Tet^R$ Amy $Cm^R$)] | Agilent | 200315 |
| **Instrumentation/Hardware** | | |
| Nikon Ti2e | Nikon | |
| Agilent 1260 HPLC | Agilent | |
| Agilent 6230B LC-MS TOF | Agilent | |
| Tecan Spark microplate reader | Tecan | |
| Avestin EmulsiFlex-C3 | Avestin | C321220 |
| Agilent Zorbax Extend-C18, 4.6mm, 1.8m, UHPLC guard column | Agilent | 820750-906 |
| Zorbax C-18 Extend 50 mm HPLC column | Agilent | 727700-902 |
| Zorbax C-18 Extend 150 mm HPLC column | Agilent | 763973-902 |
| Agilent Eclipse 150 mm XDB-C18 | Agilent | 963954-302 |
| Agilent Eclipse 250 mm XDB-C18 | Agilent | 990967-202 |
| Sealing mat, 384 wells, square, preslitted | NETA Scientific INC | 5043-9320 |
| Agilent 0.19 mL 384 well plates | Agilent | 5043-9315 |
| Greiner UV-Star Microplate 96 well plates | Greiner | 655801 |
| Corning 96-Well Solid Black Plates | Corning | 3915 |
| Corning® Costar® Ultra-Low Attachment Multiple Well Plate | Sigma Aldrich | CLS3471-24EA |
| 96-Well Solid White Plates | VWR International | 29444-041 (CS) |
| MilliporeSigma Amicon Ultra-4 Centrifugal Filter Units | Fisher Scientific | UFC800324 |
| SnakeSkin Dialysis Tubing, 10K MWCO, 35 mm | Life Technologies Corporation | PI88245 |

##### 1.2 General analytical methods.

**Semipreparative peptide HPLC method.** Sample (1 mL) was injected into an Agilent 1260 HPLC equipped with a ZORBAX Eclipse XDB-C18 (9.4 × 250 mm, 5 μm) column. The solvent system was A: water with 0.1% trifluoroacetic acid (TFA) and B: 100% acetonitrile. Samples were separated at a flow rate of 2 mL/min using the following gradient: hold at 100% A for 5 min, gradient from 0-100% B from 5.1-40 min; hold 100% B for 5 min, followed by a 15 min re-equilibration to 100% A for a total gradient time of 60 minutes.

**Analytical peptide HPLC method.** Sample (10 μL) was injected into an Agilent 1260 HPLC equipped with a 150 mm Zorbax C18 Extend (4.6 × 150 mm, 3.5 μm) column. The solvent system was A: water with 0.1% TFA and B: 100% acetonitrile with 0.1% TFA. Samples were separated at a flow rate of 0.6 mL/min using the following gradient: hold at 100% 0-100% B over 20 min; hold 100% B for 5 min, followed by a 10 min re-equilibration to 100% A for a total gradient time of 35 minutes.

**Peptide LC-MS method.** This LC-MS method was adapted from a previously described method[1]. Samples were run on an Agilent 6230B Time of Flight (TOF) LC/MS equipped with an Agilent Jet

Stream Ionization Source. An Agilent Extend C18 (2.1 × 50mm, 1.8 μm) column with a C18 guard column was used for peptide analysis and individual amino acid analysis. Solvent A: water with 0.1% formic acid. Solvent B: 100% acetonitrile. Peptides were eluted at a flow rate of 0.6 mL/min with the following gradient: 98% A for 0.1 min; gradient from 98% A to 95% B from 0.1-1.5 min; hold at 95% B from 1.5-2 min, re-equilibration to 98% A from 2.1-5 min.

**LC-MS data analysis for bromination reactions.** The extracted ion chromatogram for the major isotope of the reactant was extracted within a window of 100 ppm. For mono-brominated products, the ion chromatograms for the two major bromine isotopes ($^{79}$Br and $^{81}$Br) were extracted within a window of 100 ppm and summed together. For di-brominated products, the ion chromatograms for the two major bromine isotopes (resulting in 3 major peaks) were extracted within a window of 100 ppm and summed together. To determine percent conversion, the sum of all the brominated extracted ion chromatogram areas (product peak area) was divided by the total of all the extracted ion chromatogram areas (product peak area + substrate peak area) and multiplied by 100 according to the formula:

$$percent\ conversion = \frac{product\ peak\ area}{product\ peak\ area + substrate\ peak\ area} \times 100$$

##### 1.3 Molecular biology methods.

**Cloning.** *E. coli* codon-optimized genes encoding RebH and 4V were purchased from Twist Bioscience and Integrated DNA Technologies, respectively. Each gene had a 20-30 base pair overhang on the 3' and 5' ends for Gibson cloning into pET28a linearized by digestion with HindIII and NcoI. The RebH-K79A mutant was generated by site-directed mutagenesis using the primers 5'-cttaatcgccactgcatatgatgcattgcactcgcgcatc-3' (forward) and 5'-gatgcgcgagtgcaatgcatcatatgcagtggcgattaag-3' (reverse).

**RebH**
ttgtttaactttaagaaggagatataccatgagtggaaaaattgacaagatccttatcgtaggaggtgggacagccggctggatggca
gcctcttacttgggaaaagcattgcagggaacagcggatattacacttttgcaggcgcccgacatccctacgttaggagttggcgagg
caaccattcccaatttacagactgcgtttttcgactttttaggtatccccgaagacgaatggatgcgcgagtgcaatgcatcatataaagt
ggcgattaagttcattaattggcgcacagcaggtgaaggaacgtccgaggcccgtgagttggatggtggcccggaccatttttatcatt
ctttcggccttttaaagtatcatgagcagatccctctgagccattactggtttgaccgctcttatcgtggaaagaccgtcgaacccttcgatt
atgcttgctataaggaacctgttatcttagacgcaaatcgtagcccccgccgtttagatgggtctaaagttacgaactacgcctggcactt
tgatgcgcacttggtcgcggacttcttgcgccgtttcgctaccgagaagttgggagttcgccacgttgaggatcgtgtagagcacgttca
gcgcgatgctaacggtaatatcgaaagtgtccgcacagctaccggccgcgtgtttgacgcagatttgttcgttgactgtagtggatttcgt
ggactgttgattaataaggctatggaggaaccatttctggacatgtctgaccatttattgaatgatagcgcagtggcaacgcaggtccct
cacgatgatgacgcgaacggtgtcgaaccttttacttctgcaattgctatgaaatcggggtggacctggaagatccctatgcttggccgc
tttggtacaggctatgtgtacagcagtcgttttgctacagaggatgaagccgttcgtgaattctgcgaaatgtggcacttagacccggag
acccagccactgaaccgtatccgttttcgtgtaggtcgtaaccgccgcgcctgggtgggaaattgtgttagtattgggacgtcctcgtgct
tcgtggaacccccttgaatcaactggaatctacttcgtgtacgccgccttgtaccaattggttaaacactttcccgataagtccttgaatcctg
tcttaacagctcgttttaatcgcgaaatcgagactatgtttgacgatactcgcgattttattcaggcacatttctacttctcgccacgcactga
tacacctttctggcgcgctaataaagaattacgtttggctgatggcatgcaagagaaaatcgacatgtaccgcgcagggatggcgatc
aacgctcccgccagtgacgatgcgcagctgtactatgggaactttgaggaagaattccgcaacttttggaataattcgaattattattgc
gtccttgcaggattgggtttggtgccggacgcgccgtcaccccgtttagctcatatgccccaggcaactgagagcgtggatgaagtcttt

ggcgctgtcaaagatcgccaacgcaatcttcttgaaacacttccttcgcttcacgaatttttgcgccaacaacacggccgccatcaccat
caccaccattaaaagcttgcggccgcactcgagcaccaccacc

**4V**
cccctctagaaataattttgtttaactttaagaaggagatataccatgccgggaaagattgataaaatcttaatcgtcgggggtggcacg
gcggggtggatggccgcgtcatacttgggcaaggccttgcaagggactgccgacattacgttgttacaagcgcccgacatcccaacc
ttgggggtgggtgaagccacgattcccaacttacaaaccgcgtttttcgactttttgggtattccggaggatgagtgggtacgtgaatgta
acgcctcctacaaagttgcaattaaatttatcaattggcgtacagcaggagaaggtacgtcggaggcgcgcgagttagacgggga
ccagaccacttctatcactccttcggccttctgaaataccatgaacaaattcccctgtcgcattattggtttgaccgctcttatcgtggcaag
acggtcgaacccttcgattacgcgtgctacatggaacctgttattttggacgctaaccgcagtccccgtcgcctggacggcagcaaagt
aactaactatgcgtggcattttgatgcccaccttgtcgctgattttcttcgccgtttcgcaaccgaaaaattgggcgtccgccatgtcgagg
accgtgtagaacatgtccagcgcgacgctaacggtaatatcgagtctgtacgcacagccactggtcgtgttttcgatgcggatcttttcgt
tgattgttcgggatttcgtggacttcttattaacaaagcaatggaggagcctttttttagacatgagcgatcatttgctgaatgattcagccgtg
gcgacacaggttccacatgatgatgatgccaatggcgtagaaccgttcacctccgcgatcgccatgaaatctgggtggacatggaaa
attccgatgttaggtcgctttggcacaggttacgtgtactcatcgcgctttgccaccgaagatgaagcagtgcgtgagttctgtgagatgt
ggcatctggatcccgaaacccaaccactgaatcgtattcgtttccgtgtaggccgtaatcgtcgcgcatgggtggggaattgcgtatcta
ttggaacgagtagctgctttgtggagccgctggagtctaccggaatttatttcgtttatgctgcattgtaccagctggtaaaacatttccctg
acaagtccttgaaccctgtattgactgcccgtttcaatcgtgaaattgaaacaatgttcgacgacacacgtgactttatccaggctcatttc
tacttcagtcctcgtacagatactccgttttggcgtgcaaacaaggagctgcgccttgccgatggaatgcaagagaagattgacatgta
ccgtgcaggtatggtcattaacgccccagccagcgatgacgcacagctgtattacggcaactttgaggaagagtttcgcaacttctgg
acaaattctagttattattgtgttcttgcaggattagggttggttcctgacgctccttcgccacgtttagcacacatgccccaagctacggaa
agtgttgacgaggtttttggggccgttaaggaccgccaacgcaatttattagagacgttacccagcttgcacgaatttctgcgtcaacag
cacggacgtcaccatcatcaccaccattaaaagcttgcggccgcactcgagcacc

##### 1.4 Protein expression and purification

Protein sequences purified in this study are given below. Purification schemes were adapted from previously described methods[2].

**RebH**
MSGKIDKILIVGGGTAGWMAASYLGKALQGTADITLLQAPDIPTLGVGEATIPNLQTAFFDFLGIP
EDEWMRECNASYKVAIKFINWRTAGEGTSEARELDGGPDHFYHSFGLLKYHEQIPLSHYWFDR
SYRGKTVEPFDYACYKEPVILDANRSPRRLDGSKVTNYAWHFDAHLVADFLRRFATEKLGVRHV
EDRVEHVQRDANGNIESVRTATGRVFDADLFVDCSGFRGLLINKAMEEPFLDMSDHLLNDSAV
ATQVPHDDDANGVEPFTSAIAMKSGWTWKIPMLGRFGTGYVYSSRFATEDEAVREFCEMWHL
DPETQPLNRIRFRVGRNRRAWVGNCVSIGTSSCFVEPLESTGIYFVYAALYQLVKHFPDKSLNP
VLTARFNREIETMFDDTRDFIQAHFYFSPRTDTPFWRANKELRLADGMQEKIDMYRAGMAINAP
ASDDAQLYYGNFEEEFRNFWNNSNYYCVLAGLGLVPDAPSPRLAHMPQATESVDEVFGAVKD
RQRNLLETLPSLHEFLRQQHGRHHHHHH

**RebH-K79A**
MSGKIDKILIVGGGTAGWMAASYLGKALQGTADITLLQAPDIPTLGVGEATIPNLQTAFFDFLGIP
EDEWMRECNASYAVAIKFINWRTAGEGTSEARELDGGPDHFYHSFGLLKYHEQIPLSHYWFDR
SYRGKTVEPFDYACYKEPVILDANRSPRRLDGSKVTNYAWHFDAHLVADFLRRFATEKLGVRHV
EDRVEHVQRDANGNIESVRTATGRVFDADLFVDCSGFRGLLINKAMEEPFLDMSDHLLNDSAV
ATQVPHDDDANGVEPFTSAIAMKSGWTWKIPMLGRFGTGYVYSSRFATEDEAVREFCEMWHL
DPETQPLNRIRFRVGRNRRAWVGNCVSIGTSSCFVEPLESTGIYFVYAALYQLVKHFPDKSLNP
VLTARFNREIETMFDDTRDFIQAHFYFSPRTDTPFWRANKELRLADGMQEKIDMYRAGMAINAP
ASDDAQLYYGNFEEEFRNFWNNSNYYCVLAGLGLVPDAPSPRLAHMPQATESVDEVFGAVKD
RQRNLLETLPSLHEFLRQQHGRHHHHHH

**4V**
MPGKIDKILIVGGGTAGWMAASYLGKALQGTADITLLQAPDIPTLGVGEATIPNLQTAFFDFLGIP
EDEWVRECNASYKVAIKFINWRTAGEGTSEARELDGGPDHFYHSFGLLKYHEQIPLSHYWFDR
SYRGKTVEPFDYACYMEPVILDANRSPRRLDGSKVTNYAWHFDAHLVADFLRRFATEKLGVRH
VEDRVEHVQRDANGNIESVRTATGRVFDADLFVDCSGFRGLLINKAMEEPFLDMSDHLLNDSA
VATQVPHDDDANGVEPFTSAIAMKSGWTWKIPMLGRFGTGYVYSSRFATEDEAVREFCEMWH
LDPETQPLNRIRFRVGRNRRAWVGNCVSIGTSSCFVEPLESTGIYFVYAALYQLVKHFPDKSLN
PVLTARFNREIETMFDDTRDFIQAHFYFSPRTDTPFWRANKELRLADGMQEKIDMYRAGMVINA
PASDDAQLYYGNFEEEFRNFWTNSSYYCVLAGLGLVPDAPSPRLAHMPQATESVDEVFGAVKD
RQRNLLETLPSLHEFLRQQHGRHHHHHH

For large scale 1 L expression, 15 mL of LB media supplemented with 25 µg/mL chloramphenicol and 50 µg/mL kanamycin was inoculated with a single colony of BL21 (de3) *E. coli* transformed with pGro7 and pET28a-RebH variant. Cultures were incubated at 37°C with shaking at 200 rpm for 16 h. The entire 15 mL *E. coli* culture was then added to 1 L of autoclaved TB media supplemented with 25 µg/mL chloramphenicol and 50 µg/mL kanamycin. Flasks were shaken at 37°C at 250 rpm until an OD of 0.7-0.8 was reached at which point protein expression was induced by addition of 2 g of solid arabinose (2 mg/mL final concentration) and IPTG (100 µM final concentration). Flasks were then shaken at 25°C at 180 rpm for 16 hours. After 16 hours of expression, cell pellets were harvested by centrifugation at 5000 × g for 15 minutes at 4°C. Cell pellets were resuspended in 30 mL of lysis buffer (25 mM HEPES, pH 7.4) and lysed by three passes through a cell homogenizer at 10,000 psi. The lysate was clarified by centrifugation at 10,000 × g for 40 min. Ni-NTA resin (1 mL) was equilibrated with 10 column volumes wash buffer (20 mM sodium phosphate, 25 mM imidazole, 300 mM NaCl, pH 7.4). The clarified lysate was transferred to a new 50 mL Falcon tube and the equilibrated Ni-NTA resin was added. The lysate and resin were incubated on a rocker for 1 h. After 1 h, the Ni-NTA resin was pelleted by centrifugation at 500 × g for 2 min. The lysate was removed to a waste container and the Ni-NTA resin was resuspended and transferred to a protein column and the remaining lysate was allowed to drip through by gravity flow. The column was washed by addition of 20 column volumes of wash buffer. After washing, the protein was eluted from the Ni-NTA by addition of 3 mL of elution buffer (20 mM sodium phosphate buffer, 300 mM NaCl, 300 mM imidazole, pH 7.4). The eluted protein was transferred to 10,000 MWCO dialysis tubing and dialyzed against storage buffer (25 mM HEPES, 10% glycerol, pH 7.4) for 20 hours. Protein by purity was assessed by SDS-PAGE (**Supplementary Fig. 41**).

For large scale 1 L MBP-RebF expression, 15 mL of LB media was inoculated with a single colony of BL21 *E. coli* containing pBH4-6xHis-MBP-TEV-RebF variant and grown for 16 hours. The next day, the entire 15 mL *E. coli* culture was added to 1 L of autoclaved LB media supplemented with 1 mL of 50 mg/mL carbenicillin. 2.8 L flasks were shaken at 37°C at 250 rpm until an OD of 0.6-0.7 was reached at which point protein expression was induced by addition of 1 mL of 1 M IPTG (1 mM final concentration). Flasks were shaken at 16°C at 180 rpm for 16 hours. After 16 hours of expression, cell pellets were harvested by centrifugation at 5000 x g for 15 minutes. Cell pellets were resuspended in 30 mL of lysis buffer (25 mM HEPES, pH 7.4) and lysed by three passes through a cell homogenizer set to10,000 psi. The lysate was clarified by centrifugation at 10,000 x g for 40 minutes. Meanwhile, 1 mL of Ni-NTA resin was equilibrated with 10 column volumes wash buffer (20 mM phosphate, 25 mM imidazole, 300 mM NaCl, pH 7.4). The clarified lysate was transferred to a new 50 mL Falcon tube and the equilibrated Ni-NTA resin was added. The lysate and resin was incubated on a rocker for 1 hour. After 1 hour, the Ni-NTA resin was pelleted by centrifugation at 500 x g for 2 minutes. The lysate was removed to a waste container and the Ni-NTA resin was resuspended and transferred to a protein column and the remaining lysate was allowed to drip through by gravity. The column was washed by addition of 20 column volumes of wash buffer. After washing, the protein was eluted from the Ni-NTA by addition of 3 mL of elution buffer (20 mM phosphate buffer, 300 mM NaCl, 300 mM imidazole, pH 7.4). The eluted protein was transferred to 10,000 MWCO dialysis tubing and dialyzed against storage buffer (25 mM HEPES, 10% glycerol, pH 7.4) for 20 hours.

##### 1.5 Solid-phase peptide synthesis

Tripeptides were ordered from peptide2.0 as crude peptide product in a 96 well plate. Peptides with a carboxamide group at the C-terminus were synthesized using Rink amide resin (Chem Impex International). Peptides with a carboxylate at the C-terminus were synthesized using Wang resin pre-loaded with the C terminal amino acid residue (Chem Impex International). Fmoc amino acids with reactive side chains were protected with the following acid-labile protecting groups: Asp (OtBu); Glu (OtBu); His(Trt); Lys(Boc); Asn(Trt); Gln(Trt); Arg(Pbf); Ser(tBu); Thr(tBu); Trp(Boc); Tyr(tBu). Fmoc groups were deprotected by incubating for 30 min with 4-methylpiperidine (20% v/v in DMF). After deprotecting, resin was washed with 5 volumes of DMF. Coupling steps were then performed using 5 molar equivalents of Fmoc amino acid, 5 equivalents of diisopropylcarbodiimide (DIC), and 5 equivalents of 1-hydroxybenzotriazole (HOBt). After synthesis, peptides were cleaved from the resin using a cocktail containing 95% TFA, 2.5% triisopropylsilane, and 2.5% water. The volume of TFA was reduced under a stream of nitrogen and peptide were precipitated with 10 volumes of diethyl ether. Pellets were washed with 10 additional volumes of diethyl ether and allowed to dry. The crude product was purified using an Agilent 1260 Infinity II HPLC fitted with a semi-preparative ZORBAX Eclipse XDB-C18, 9.4 × 250 mm, 5 μm column according to semipreparative HPLC method described above. ESI-MS data for synthetic peptides are shown in the Supplementary Appendix.

##### 1.6 Analytical-scale RebH reactions

Reactions were set up in triplicate in 384 well plates. Substrate (1 mM final concentration) was added to each reaction well. Master mix (14.4 μL) containing MBP-RebF (2.5 μM final concentration), RebH variant (25 μM final concentration), glucose dehydrogenase (10 U/mL final concentration), NAD (100 μM final concentration), FAD (100 μM final concentration), glucose (20 mM final concentration), NaBr (10 mM final concentration) in 25 mM HEPES, pH 7.4 was added by multichannel pipette to each reaction well. The reactions were incubated at room temperature for 24 hours without shaking. After 24 hours, the reactions were quenched with an equal volume of mass spec grade methanol (15 μL) and then centrifuged at 1,500 × g for 20 minutes to remove the precipitate. Supernatant (15 μL) was transferred to a clean 384 well plate using a multichannel pipette. The plates were sealed with a silicon mat and 0.25 μL was analyzed by the peptide LC-MS method described above and ESI-MS data are shown the Supplementary Appendix.

##### 1.7 Preparative-scale RebH reactions

Reactions were set up in a 50 mL Falcon tube. Solid peptide substrate was added to a final concentration of 10 mM. NAD and FAD were added as solids to a final concentration of 100 μM. Glucose dehydrogenase was added as a lyophilized powder to a final concentration of 10 U/mL. MBP-RebF and 4V enzymes were added to final concentrations of 2.5 μM and 10 μM, respectively. Glucose dissolved in 25 mM HEPES, pH 7.4 was added to a final concentration of 20 mM. The total reaction volume was brought up to 15 mL by addition of 25 mM HEPES, pH 7.4. The 15 mL reaction was placed on a rocker for 24 hours. After 24 hours, the reaction was quenched by addition of an equal volume (15 mL) methanol and then centrifuged at 8,000 × g for 15 minutes to remove precipitate. The reaction was concentrated down to 2 mL using a vacuum centrifuge before injecting on a the semipreparative HPLC and purifiying according to the semipreparative HPLC method described above. Purified peptides were analyzed by HPLC and LC-MS. ESI-MS data are shown in the Supplementary Appendix.

##### 1.8 Trypsin digestion of brominated bioactive peptides

Analytical scale 4V bromination reactions (30 µL) were quenched with an equal volume of mass spec grade methanol and centrifuged at 10,000 × g for 10 min to remove any precipitate. The soluble fraction (30 µL) was removed to a new 1.5 mL reaction tube and dried in a vacuum concentrator. The dried pellet was resuspended in 15 µL of 50 mM Tris, pH 8.0 by vortexing. Sequencing grade trypsin was added to a w/w of 1:10 peptide:trypsin (0.1 µg trypsin). The reaction was incubated for 16 hours at 37°C on a thermomixer shaking at 500 rpm. The reaction was quenched by adding an equal volume of mass spec grade methanol and centrifuging at 10,000 × g for 10 min to remove any precipitate. Reactions were analyzed using the peptide analytical LC-MS method described above.

##### 1.9 Suzuki-Miyaura coupling of brominated bioactive peptides

Suzuki-Miyaura couplings were performed using a method adapted from previous work[3,4]. Purified 4V brominated peptides were resuspended to 20 mM in 50% water/ethanol. Suzuki-Miyaura coupling reactions (40 µL) were set up with 5 mM aryl boronic acid (final concentration), 0.25 mM brominated peptide (final concentration), 2.5 mM DMG/Pd catalyst (final concentration), and 75 mM $NaH_2PO_4$ (final concentration). Water was added to bring the volume up to 40 µL. The reactions were incubated at 45°C for 4 hours on a thermomixer set to shake at 500 rpm. Reactions were quenched with an equal volume of 5 µl/mL 3-mercaptopropionic acid in water and centrifuged to remove insoluble material. Reactions were analyzed using the peptide analytical LC-MS method described above.

##### 1.10 µOR GloSensor assay

The GloSensor cAMP Assay (Promega) was performed according to the manufacturer's protocol. Unless otherwise stated, HEK293T cells were cultured in DMEM+10% FBS+1% penicillin-streptomycin. In brief, HEK293T cells (passage 15-20) were seeded at a density of 1-1.5 x $10^5$ cells per mL in a 6-well culture dish. For each well, 2 µg of GloSensor 22F plasmid and 2 µg of MOR-duet plasmid was mixed with 200 µL pre-warmed serum-free DMEM and incubated for 5 min at room temperature in a 1.5 mL tube. After 5 minutes, 12 µL Fugene 4K transfection reagent (Promega, 3:1 ratio) was added to the plasmid solution and mixed thoroughly. The transfection solution was incubated for 20 min at room temperature, after which, the entirety was added dropwise onto a single well of cells. The µOR transfected cells were allowed to grow for 24 h at 37°C with 5% $CO_2$. The next day, the µOR transfected cells were trypsinized and re-seeded in a solid white cell treated 96 well plate coated with poly-D-lysine at a density of 1-1.5 x $10^4$ cells per well. The re-seeded cells were grown for another 24 h at 37°C with 5% $CO_2$. The next day, the cells were rinsed once with 100 µL of pre-warmed $CO_2$-independent medium+10% FBS, taking care to avoid detaching the cells. The rinsed cells were then incubated for 30 min in the dark at 37°C with 5% $CO_2$ with 98 µL of pre-warmed $CO_2$-independent medium + 10% FBS +2% GloSensor reagent. After 30 min, the cells were removed from the incubator and allowed to cool to room temperature for 15 min in the dark. A 10-min baseline luminescence reading was taken using a Tecan Spark microplate reader set to operate at room temperature. A 10-fold serial dilution of agonist compounds was performed in PBS. After 10 min, 2 µL of 50X µOR agonist or DMSO control was added in quadruplicate to each well of the 96 well plate using a multichannel pipette. The cells were incubated with compound in the dark for 10 min at room temperature. After incubation, 500 nM forskolin (final concentration) or PBS control was added to each well. Immediately upon addition of forskolin, luminescence values were recorded for 2 h using a 400 ms integration value.

For each agonist, data were normalized per-well according to the following equation:

$$\%\, cAMP\ inhibition = \left(1 - \left(\frac{Y - Y_{min}}{Y_{max} - Y_{min}}\right)\right) \times 100$$

Where Y is the luminescence value at a given agonist concentration, $Y_{min}$ is the luminescence value at the highest agonist concentration, and $Y_{max}$ is the luminescence value for the forskolin only control. For agonists that gave biphasic responses, only the inhibitory portion ($Y < Y_{max}$) of the dose–response curve was used for fitting.

The dose-response curve was fit to the following equation:

$$\%\, cAMP\ inhibition = Bottom + \left(\frac{Top - Bottom}{1 + 10^{(\log EC_{50}) - x}}\right)$$

#### 1.11 Immunofluorescence for µOR transfected HEK293T cells

HEK293T cells (passage 15-20) were seeded at a density of 1-1.5 × $10^5$ cells per mL in a 6-well culture dish coated with poly-D-lysine. The cells were transfected with 2 µg of MOR-duet plasmid using a 3:1 Fugene 4K transfection reagent ratio and allowed to grow for 24 h. The next day, the cells were fixed using 4% formaldehyde in PBS for 10 minutes at room temperature. The fixative solution was removed and the cells were washed 3 × 10 min with PBS with gentle rocking. The cells were then permeabilized with 0.1% Triton X-100 in PBS for 15 min at room temperature. The permeabilization solution was removed and the cells were washed 3 × 10 min with PBS with gentle rocking. Cells were then blocked using SuperBlock buffer for 1 h at room temperature. 1D4 primary antibody (Life Technologies) was diluted 1:1000 into SuperBlock buffer. 1D4 was chosen to target the 1D4 epitope tag incorporated at the C terminus of the µOR construct. The cells were incubated with the primary antibody overnight at 4°C. The primary antibody solution was removed the following morning and the cells were washed 3 × 10 min with PBS with gentle rocking. During all subsequent steps, care was taken to protect the cells from exposure to light. AlexaFluor 488 conjugated goat anti-mouse secondary antibody was diluted 1:500 in SuperBlock buffer and incubated with the cells for 1 h at room temperature. The secondary antibody solution was removed and the cells were washed 3 × 10 min with PBS + 0.1% Tween-20 with gentle rocking. A final wash in PBS was performed to remove Tween-20 prior to imaging. After washing was completed, 2 drops of NucBlue were added to counterstain the cell nuclei. The cells were imaged using a Nikon microscope equipped with an Orca-Fire camera.

#### 1.12 Transportan localization assay in U2OS cells

U2OS (passage 10-15) cells were trypsinized and seeded at a density of 1.5 × $10^4$ cells per mL in a 96-well culture dish in FluoroBrite DMEM. The following day, the cells were incubated with 5 µM (final concentration) of 7-naphthalene-L-tryptophan, transportan, naphthalene-transportan, or DMSO as a negative control for 20 min at 37°C + 5% $CO_2$. The final concentration of DMSO was 0.5%. After incubation, the cells were washed 2 × 0.5 mg/mL heparin sulfate in PBS and then washed with 2 × FluoroBrite DMEM. FluoroBrite DMEM (200 µL) was added to each well for imaging. 1 drop of NucRed nuclear counterstain was added to visualize the cell nuclei. The cells were imaged using a Nikon microscope equipped with an Orca-Fire camera. Fiji was used for quantification of nuclear transportan signals in Figure 5e. Fluorescence images were background corrected and individual nuclei were segmented based on a threshold intensity in the NucRed channel. The resulting nuclear masks were used to query the mean intensity in the transportan channel.

##### 1.13 Bacterial minimal inhibitory concentration measurements

Mueller-Hinton II cation-adjusted broth (10 mL) was inoculated with *E. coli* XL10 and incubated overnight with shaking at 37°C. The next day, the overnight culture was diluted to an $OD_{600}$ of 0.5 with Mueller-Hinton II cation-adjusted broth (equivalent to ~1 × $10^8$ colony forming units). The $OD_{600}$ 0.5 culture was further diluted 20-fold into fresh Mueller-Hinton II cation-adjusted broth (equivalent to ~1 × $10^6$ colony forming units). A 2-fold serial dilution series was performed for each antimicrobial peptide to achieve a 100X stock solution. For $(RW)_3$ peptides, the highest stock concentration was 625 µM for $(RW)_3$ peptides. Pre-warmed M9 media (178 µL) was added to each well of a sterile 96-well plate. Using a multichannel pipette, 2 µL of the 100X stock peptide was added to the appropriate wells. Each compound was assayed in triplicate and was arrayed in every other row to avoid errors associated with uneven evaporation across the plate. The assay was initiated by addition of 20 µL of the ~1 × $10^6$ colony forming unit culture. M9 without added bacteria was included as a control for contamination. M9 media with bacteria, but without antibiotics, was spotted as a positive control for growth. Bacterial growth was monitored using a Tecan Spark microplate reader set to 37°C and measuring absorbance at 600 nm every 10 min with shaking for 540 seconds (amplitude 3) for 18 h.

##### 1.14 Bacterial $IC_{50}$ determination

$IC_{50}$ calculations were performed on the optical density measurements obtained from the minimal inhibitory concentration growth assay (see above). Data were fit in GraphPad Prism version 10.5.0 using a [inhibitor] vs. response –Variable slope (four parameters) dose-response curve.

##### 1.15 Minimal bactericidal concentration on petri dishes

From each triplicate experiment, 100 µL of the 18-hour minimal inhibitory growth culture (see above) was plated on a petri dish containing Mueller-Hinton II cation-adjusted agar. The plates incubated at 37°C overnight. The next morning, the bacterial growth on each triplicate plate was assessed. M9 media (100 µL) without bacteria was plated as a control for contamination. M9 media (100 µL) with bacteria, but without antibiotics, was spotted as a positive control for growth.

##### 1.16 Minimal bactericidal concentration spot assay

Using a multichannel pipette, 3 µL from each triplicate of the 18-hour minimal inhibitory growth culture (see above) was spotted on a square petri dish containing Mueller-Hinton II cation-adjusted agar. The plates were incubated at 37°C overnight. The next morning, the bacterial growth on each triplicate plate was assessed. M9 media (3 µL) without bacteria was plated as a control for contamination. M9 media (3 µL) with bacteria, but without antibiotics, was spotted as a positive control for growth.

#### 2. Supplementary Figures and Tables

**Supplementary Figure 1. Representative extracted ion chromatograms (EICs) for GGW-$CONH_2$ and GG-7-BrW-$CONH_2$.**

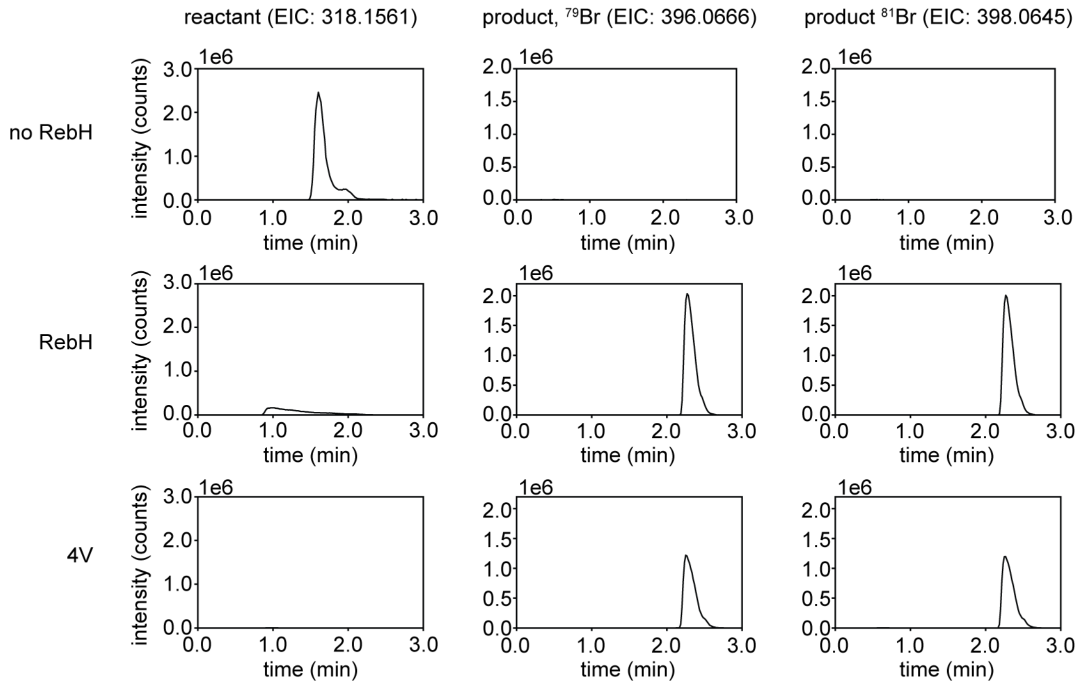

**Supplementary Figure 2. Representative extracted ion chromatograms (EICs) for WGG-CONH$_2$ and 7-BrW-GG-CONH$_2$.**

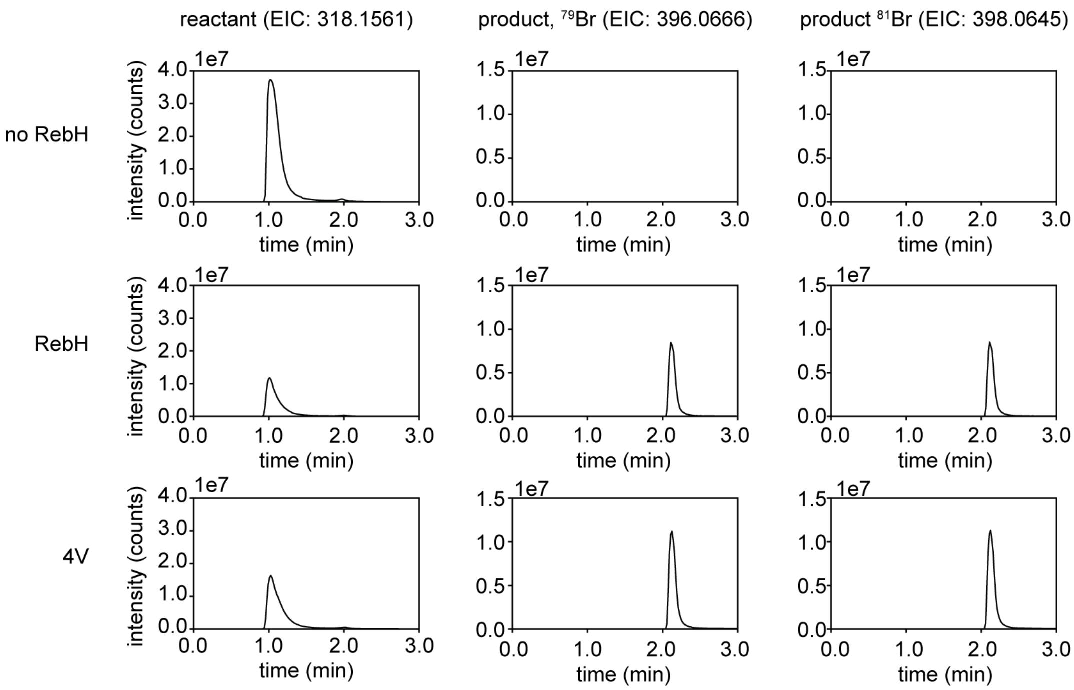

**Supplementary Figure 3. Representative extracted ion chromatograms (EICs) for GWG-CONH$_2$ and G7-BrW-G-CONH$_2$.**

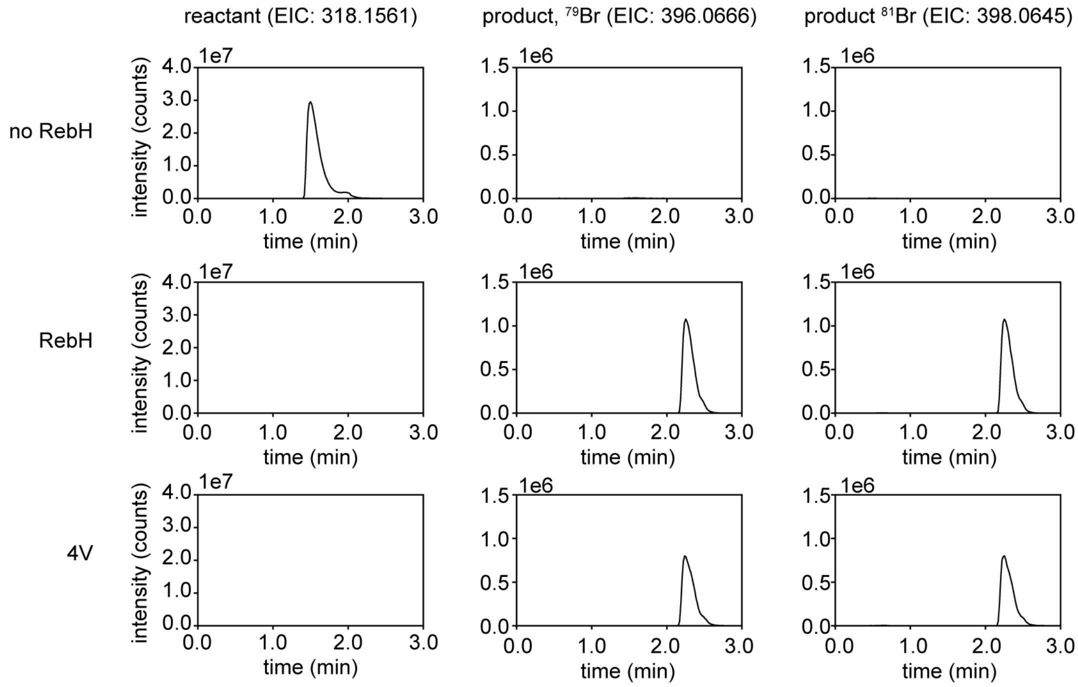

**Supplementary Figure 4. RebH and 4V activity on peptides with $Gly_n$ extensions.** (a) Peptides with C-terminal Trp and N-terminal $Gly_n$. (b) Peptides with N-terminal Trp and C-terminal $Gly_n$. (c) Peptides with central Trp and Gly residues on the N- and C-terminal sides. Percent conversion was calculated using LC-HRMS extracted ion chromatograms for the reactants and products (n=3 replicates).

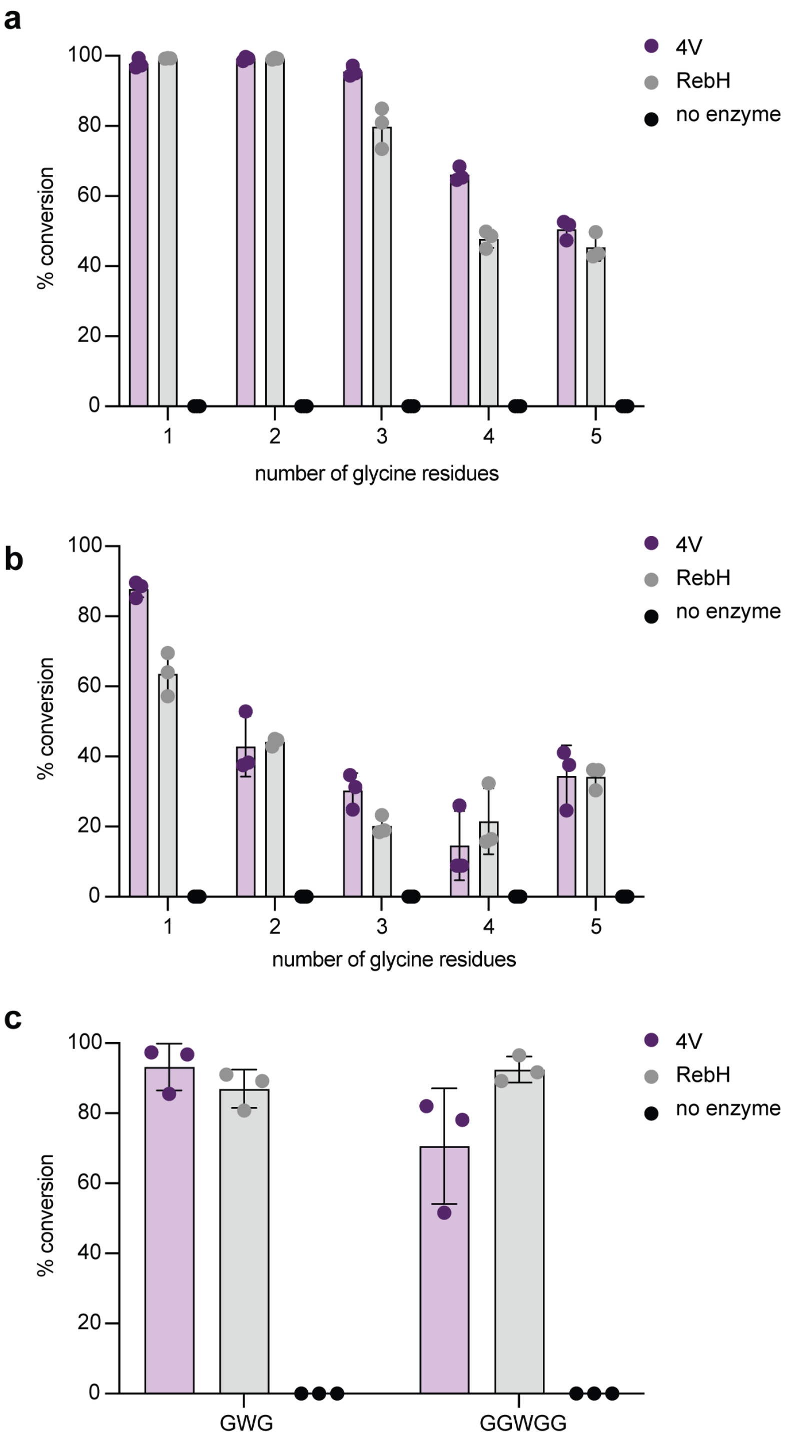

**Supplementary Figure 5. $^{1}H$ NMR spectrum for 7-Br-Trp ($D_2O$, 500 MHz):** δ 7.69 (dd, *J* = 10.0 Hz, small coupling unresolved, 1H), 7.38 (s, 1H), 7.10 (t, *J* = 7.5 Hz, 1H), 4.16 (dd, *J* = 10.0, 5.0 Hz, 1H), 3.41 (ABX, 2H).

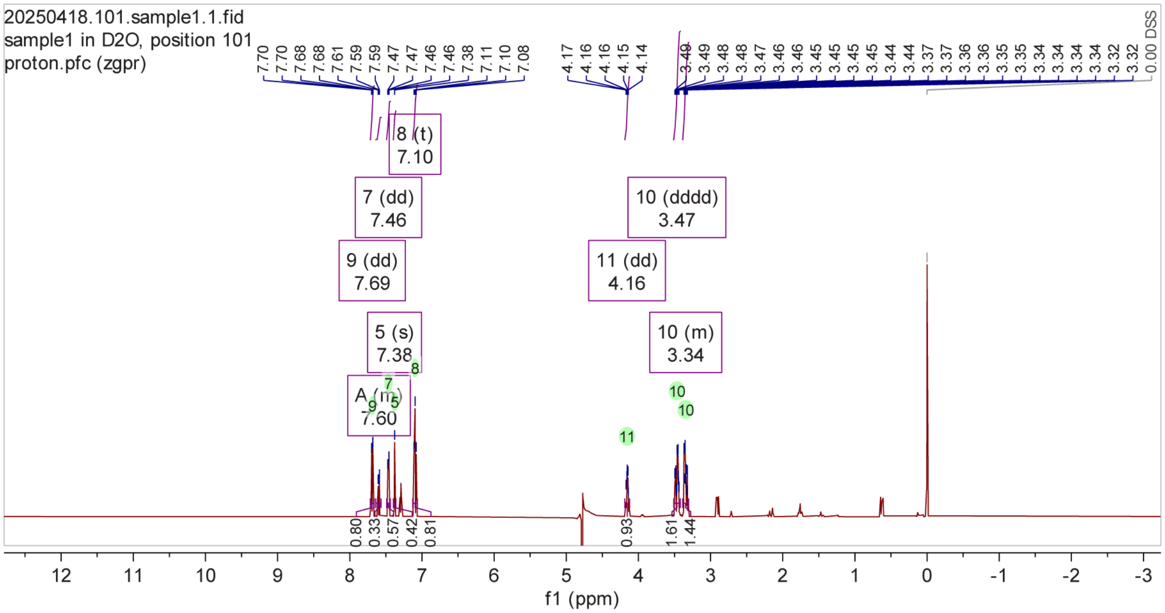

$^{1}H$ NMR (500 MHz, $D_2O$) δ 7.70, 7.70, 7.68, 7.68, 7.47, 7.47, 7.46, 7.46, 7.38, 7.11, 7.10, 7.08, 4.17, 4.16, 4.16, 4.15, 4.14, 3.49, 3.49, 3.48, 3.48, 3.48, 3.48, 3.47, 3.47, 3.46, 3.46, 3.45, 3.45, 3.45, 3.45, 3.44, 3.44, 3.37, 3.37, 3.36, 3.36, 3.35, 3.35, 3.34, 3.34, 3.34, 3.34, 3.33, 3.33, 3.32, 3.32.

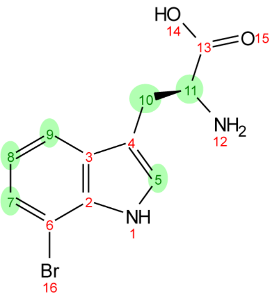

**Supplementary Figure 6. $^1$H-$^1$H COSY NMR spectrum for 7-Br-Trp ($D_2O$, 500 MHz).**

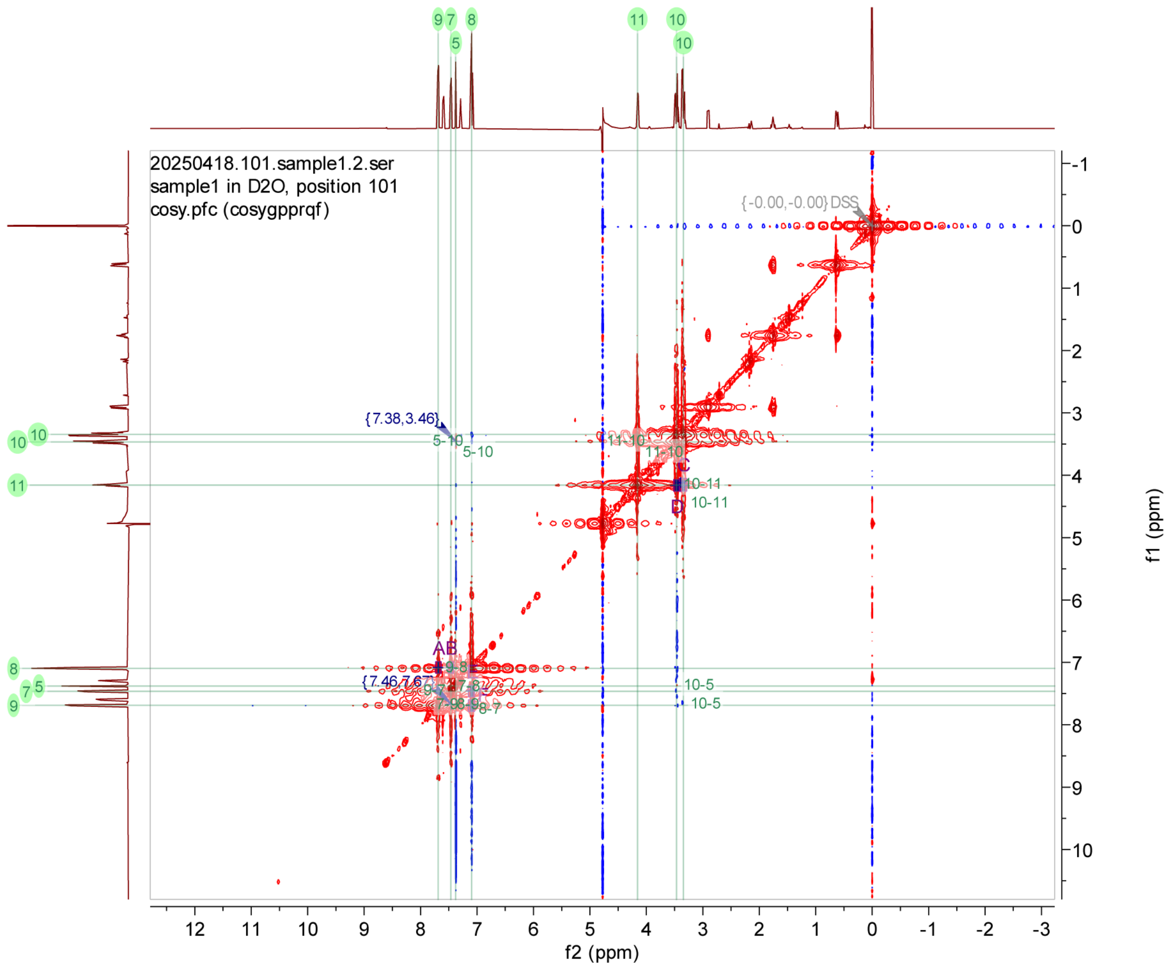

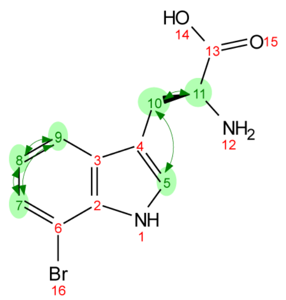

**Supplementary Figure 7. $^{1}H$-$^{1}H$ TOCSY NMR spectrum for 7-Br-Trp ($D_2O$, 500 MHz).**

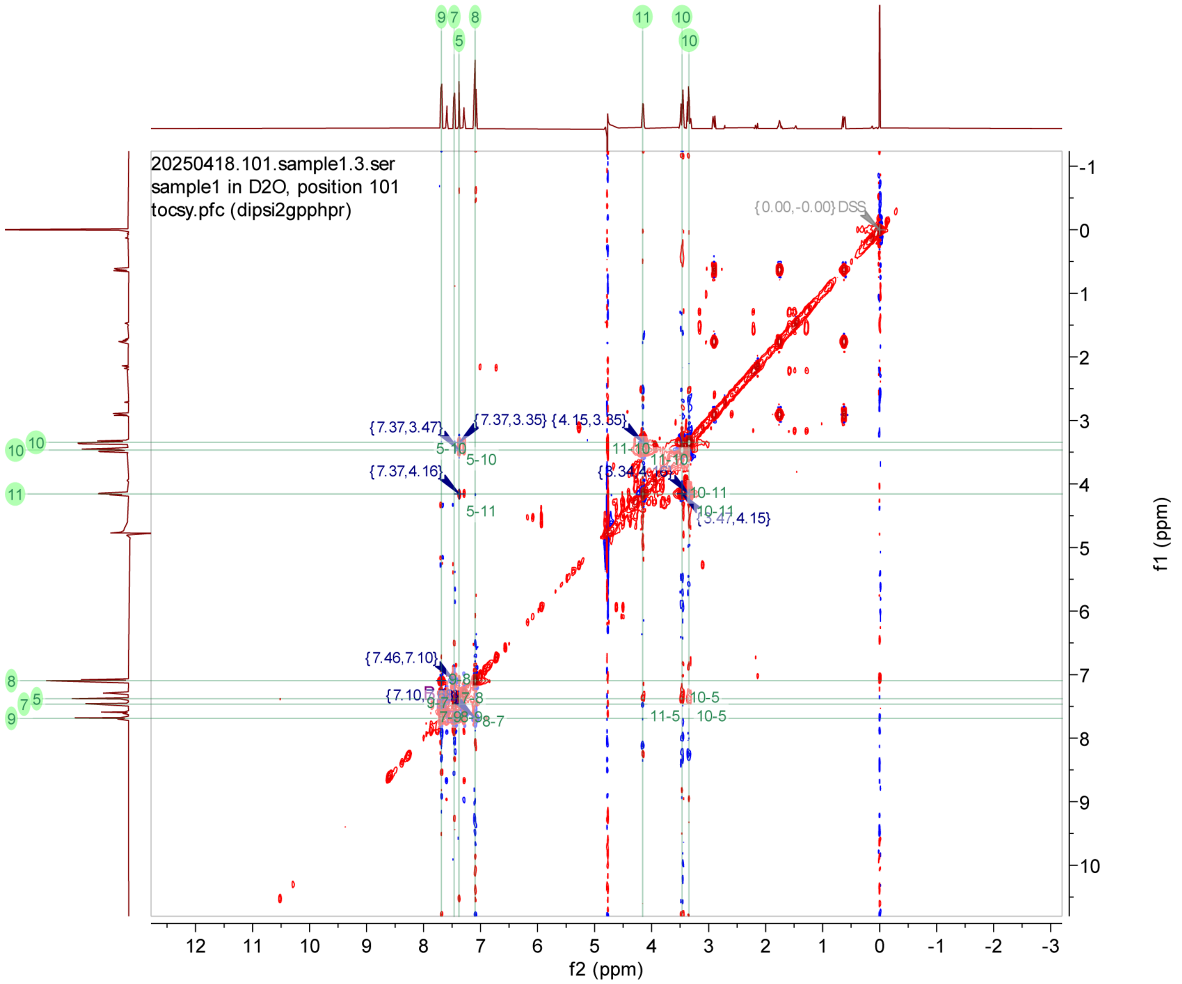

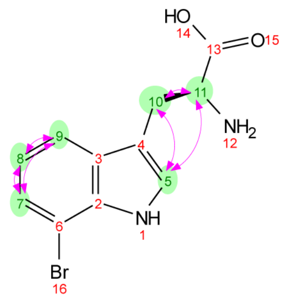

**Supplementary Figure 8. $^{1}H$-$^{1}H$ NOESY NMR spectrum for 7-Br-Trp ($D_2O$, 500 MHz).**

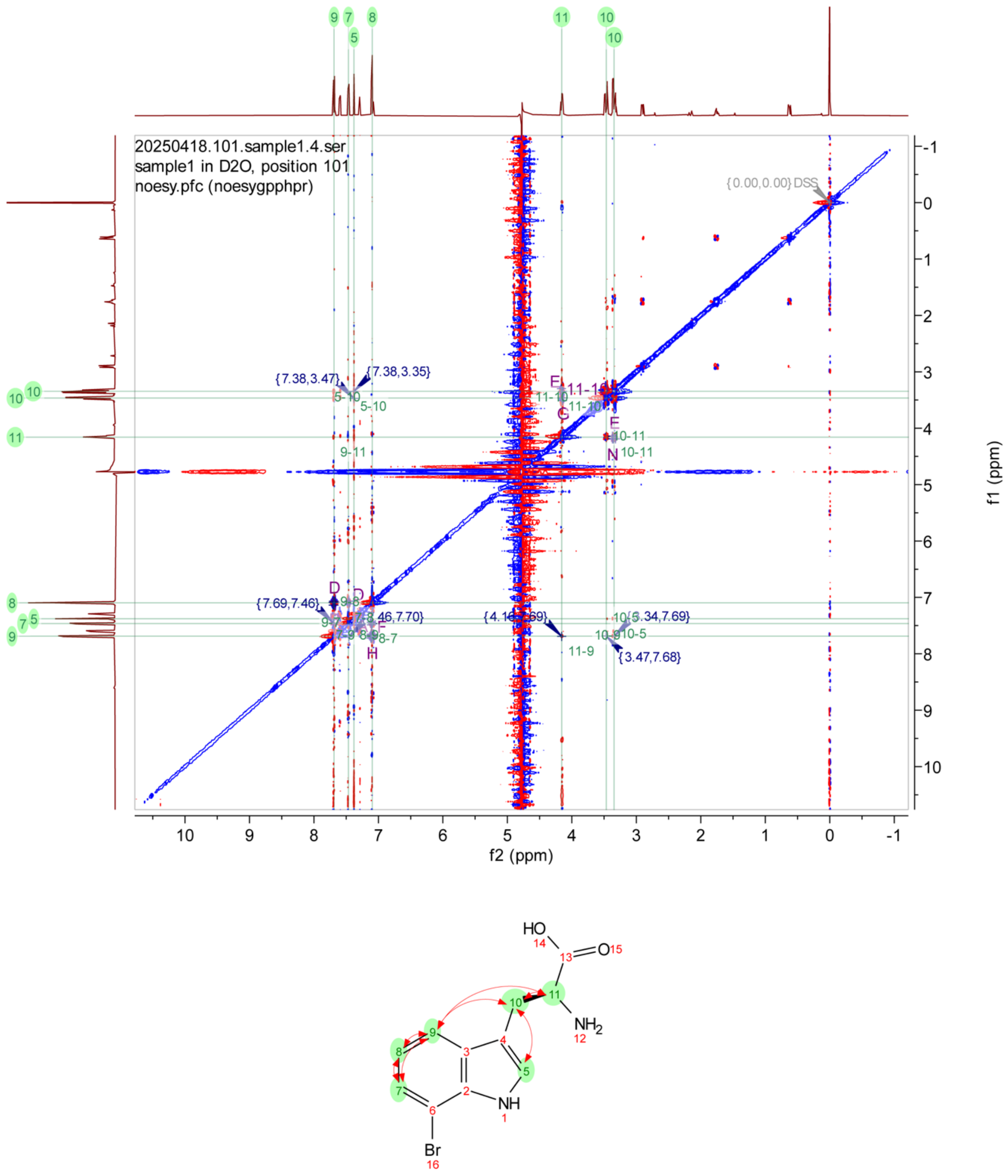

#### Supplementary Figure 9. $^1$H NMR spectrum for GG-7-BrW-CONH$_2$ ($D_2O$, 500 MHz).

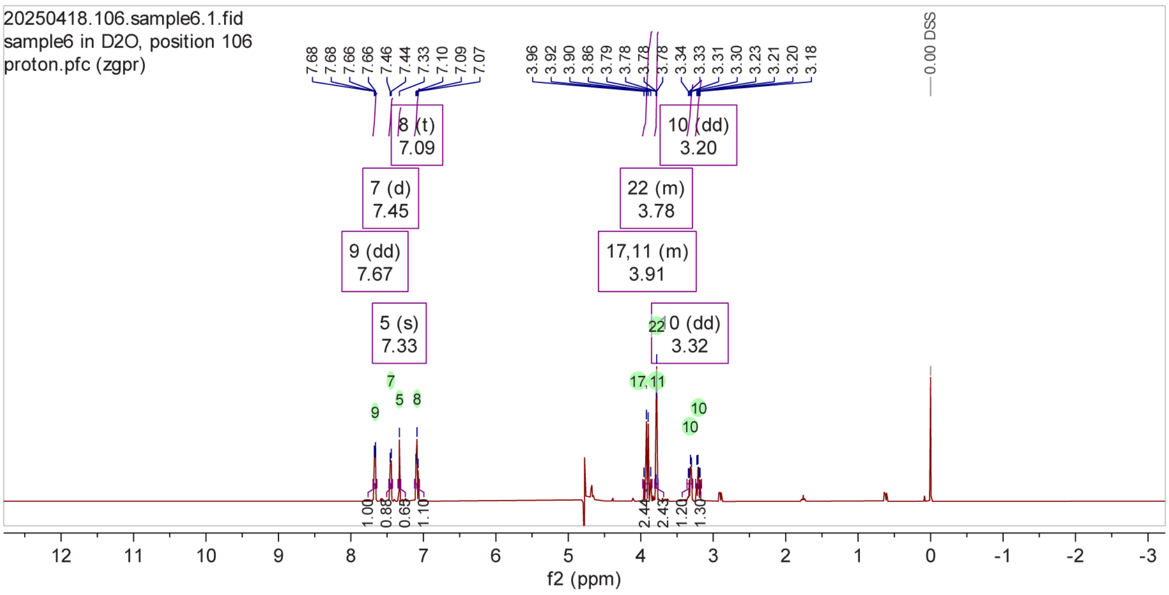

$^1$H NMR (500 MHz, $D_2O$) δ 7.68, 7.68, 7.66, 7.66, 7.46, 7.44, 7.33, 7.10, 7.09, 7.07, 3.96, 3.92, 3.90, 3.86, 3.79, 3.78, 3.78, 3.78, 3.34, 3.33, 3.31, 3.30, 3.23, 3.21, 3.20, 3.18.

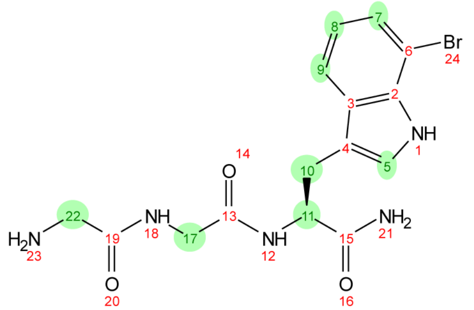

**Supplementary Figure 10. $^1H$-$^1H$ COSY NMR spectrum for GG-7-BrW-CONH$_2$ ($D_2O$, 500 MHz).**

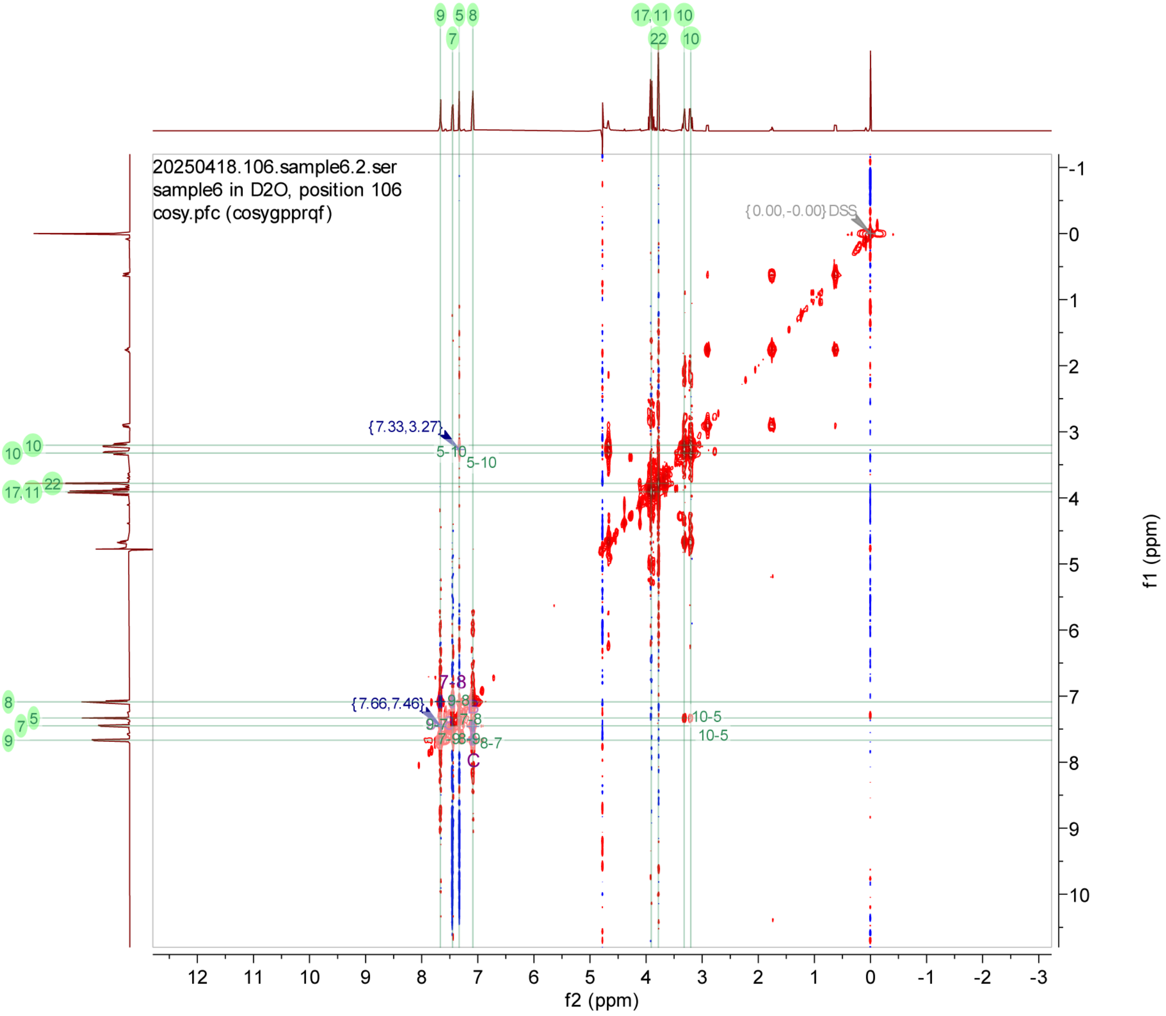

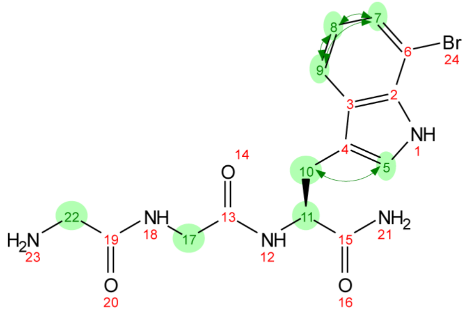

**Supplementary Figure 11. $^1$H-$^1$H TOCSY NMR spectrum for GG-7-BrW-CONH$_2$ ($D_2O$, 500 MHz).**

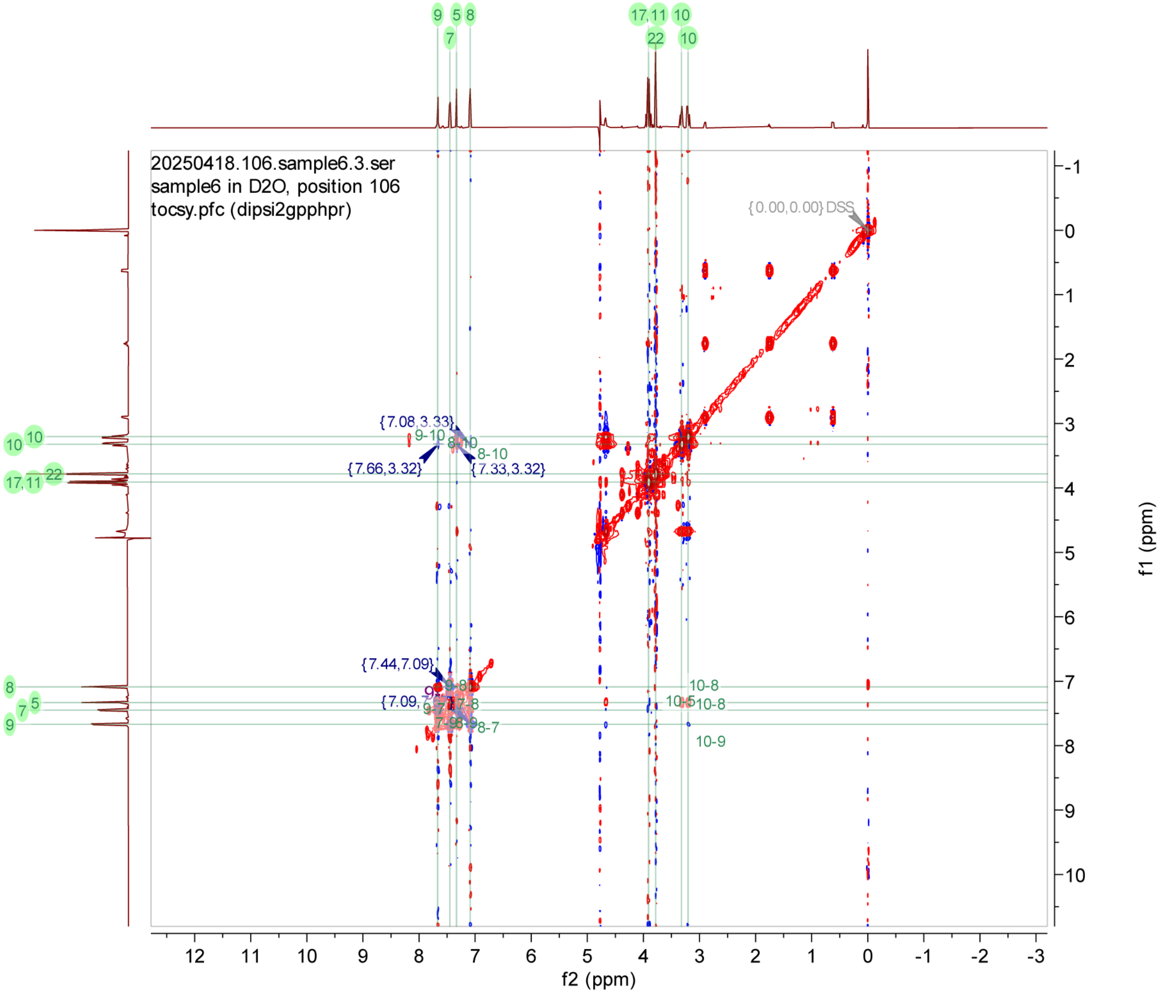

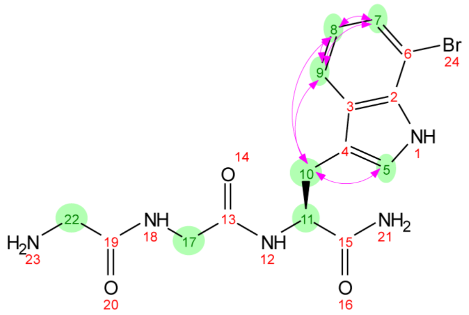

**Supplementary Figure 12. $^1$H-$^1$H NOESY NMR spectrum for GG-7-BrW-$CONH_2$ ($D_2O$, 500 MHz).**

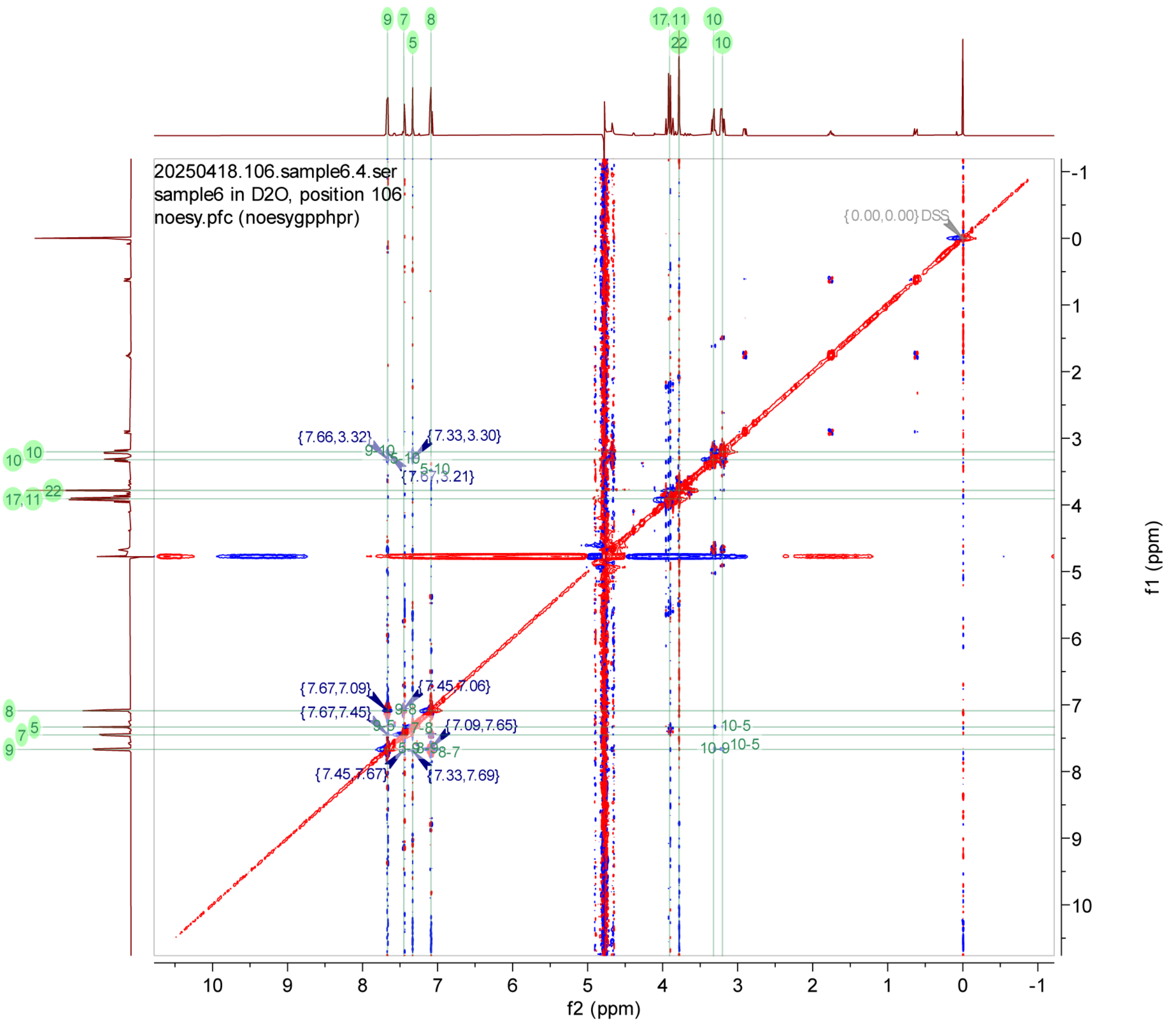

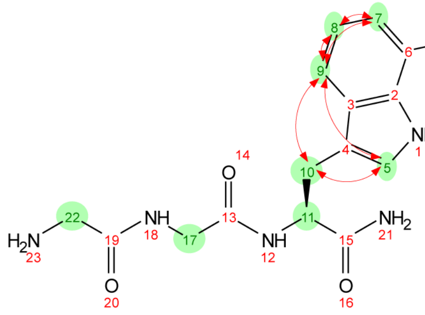

**Supplementary Figure 13. $^{1}H$ NMR spectrum for 7-BrW-GG-CONH$_2$ ($D_2O$, 500 MHz).**

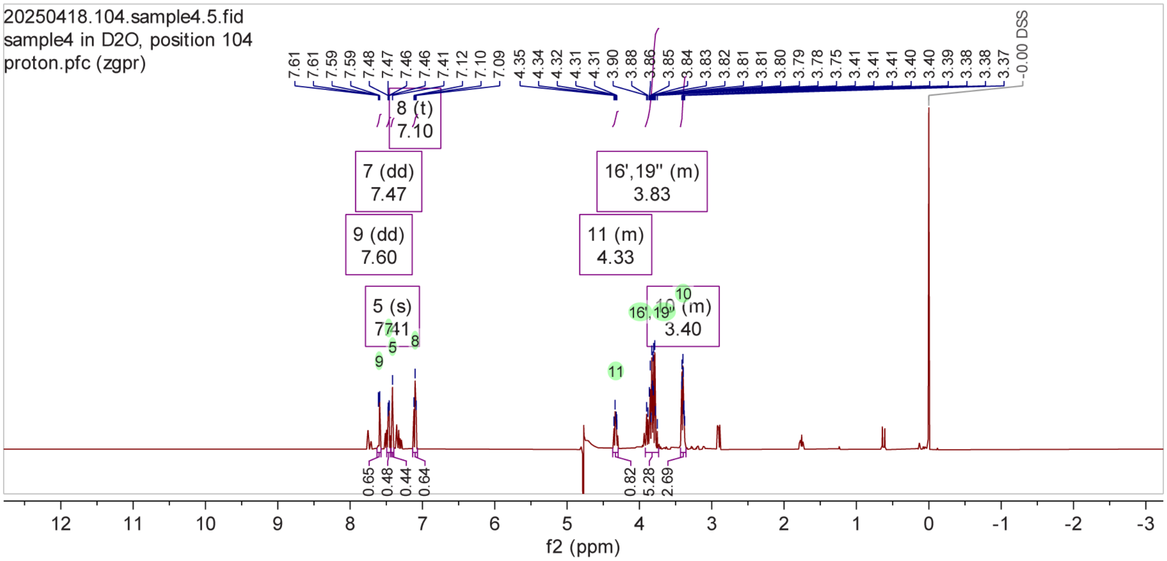

$^{1}H$ NMR (500 MHz, $D_2O$) δ 7.61, 7.61, 7.59, 7.59, 7.48, 7.47, 7.46, 7.46, 7.41, 7.12, 7.10, 7.09, 4.35, 4.34, 4.32, 4.31, 4.31, 3.90, 3.88, 3.86, 3.85, 3.84, 3.83, 3.82, 3.81, 3.81, 3.80, 3.79, 3.78, 3.75, 3.41, 3.41, 3.41, 3.40, 3.40, 3.39, 3.38, 3.38, 3.37.

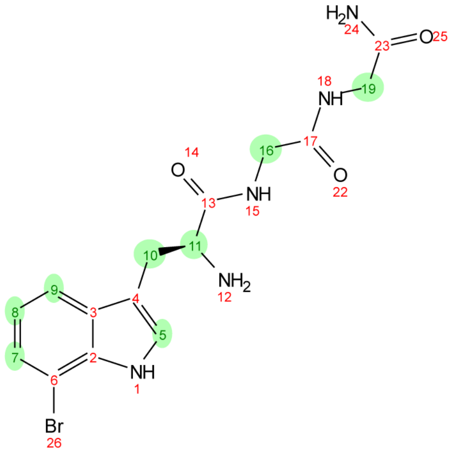

**Supplementary Figure 14. [1]H-[1]H COSY NMR spectrum for 7-BrW-GG-CONH$_2$ ($D_2O$, 500 MHz).**

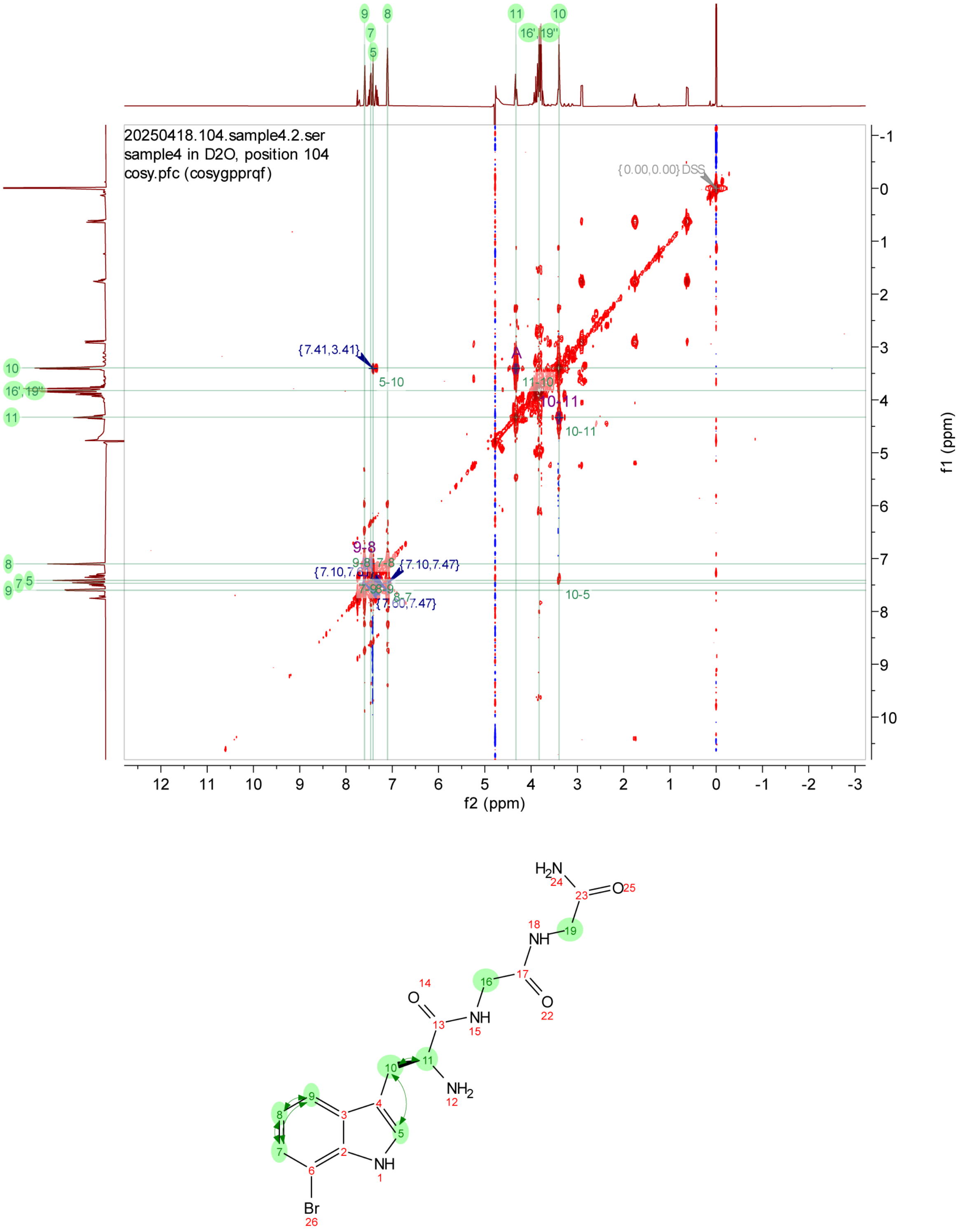

**Supplementary Figure 15. $^{1}H$-$^{1}H$ TOCSY NMR spectrum for 7-BrW-GG-CONH$_2$ ($D_2O$, 500 MHz).**

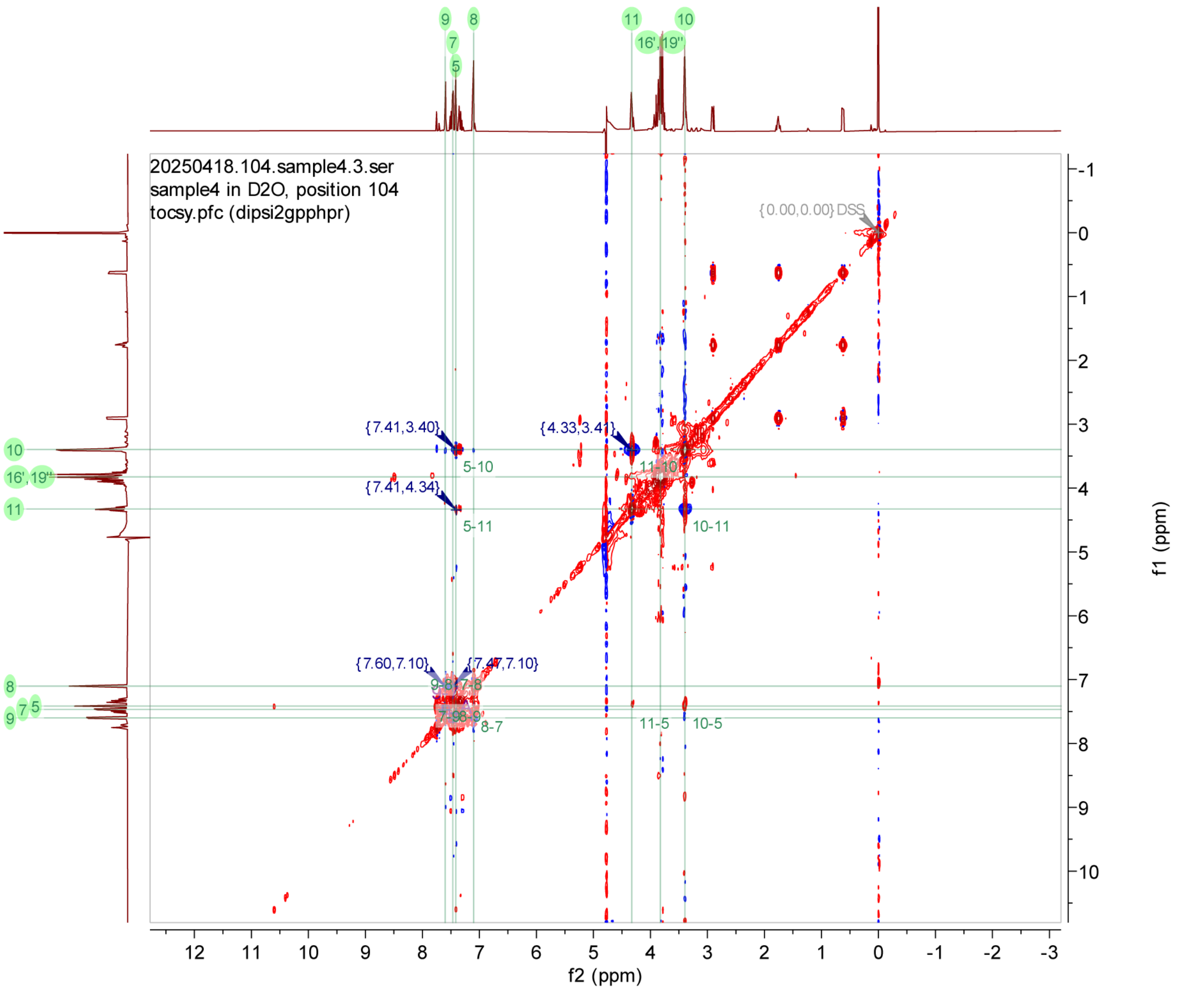

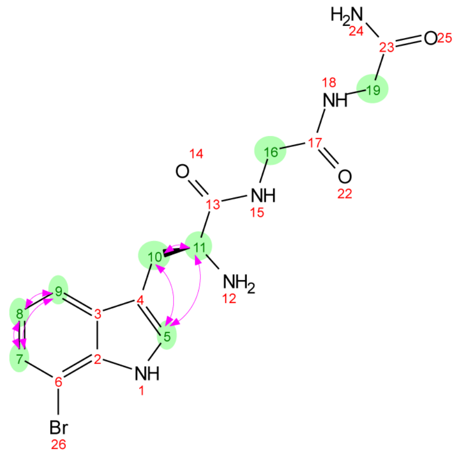

**Supplementary Figure 16. $^1$H-$^1$H NOESY NMR spectrum for 7-BrW-GG-CONH$_2$ ($D_2O$, 500 MHz).**

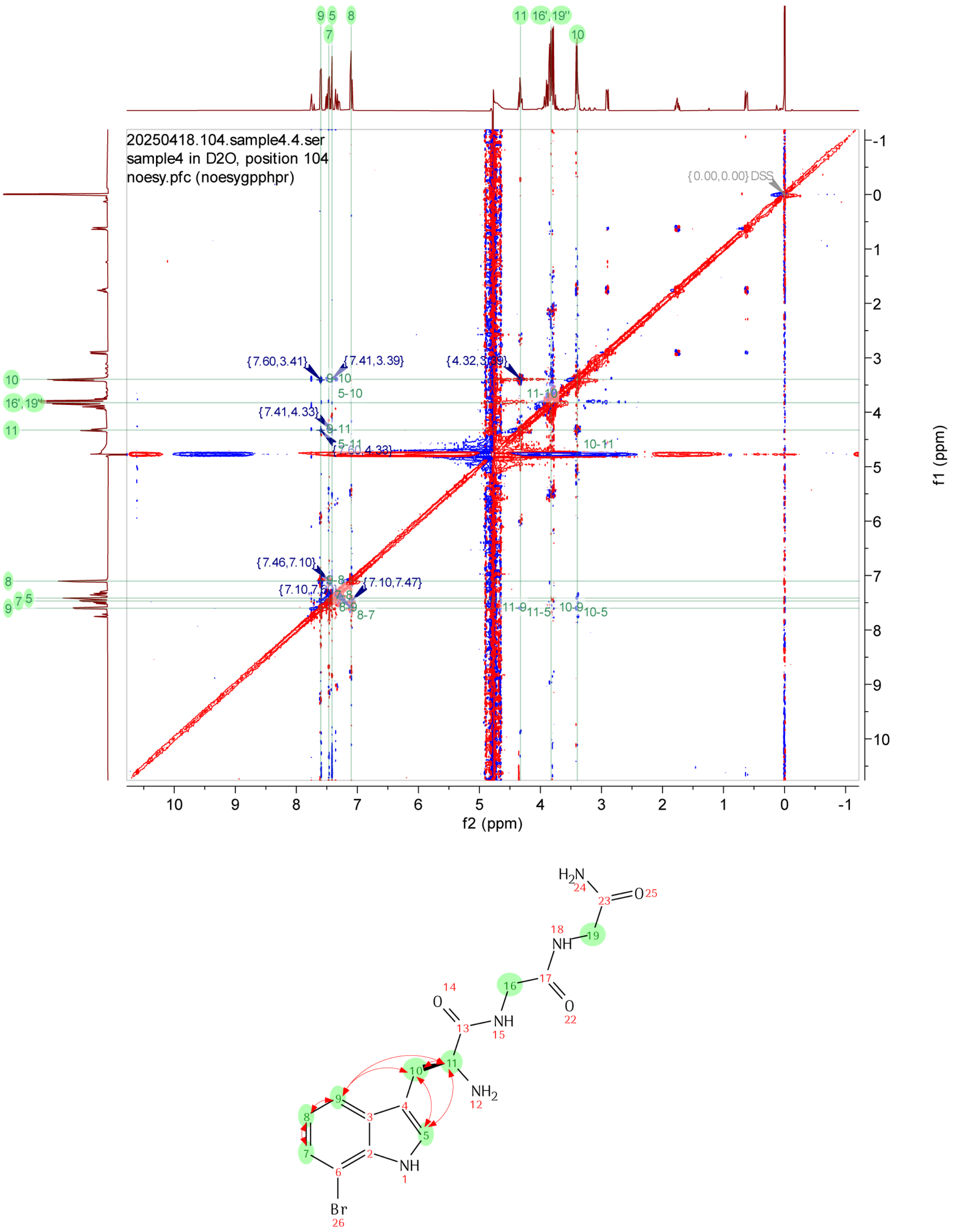

**Supplementary Figure 17. RebH- and 4V-catalyzed mono- and di-bromination of peptides.**
a) RebH-catalyzed mono- (lavender) and debromination (grey) of Trp-containing glycine peptides.
b) 4V-catalyzed mono- (lavender) and debromination (grey) of Trp-containing glycine peptides.

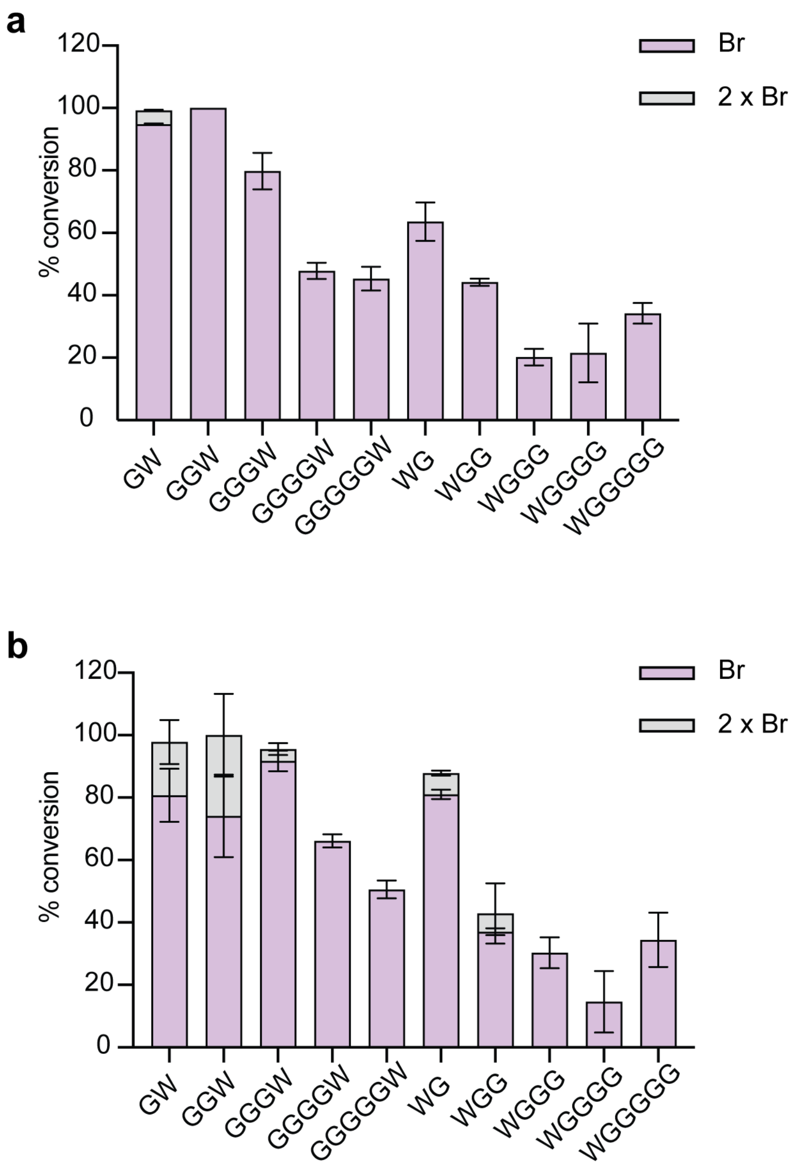

**Supplementary Figure 18. Structural models of RebH-peptide complexes.** Trp peptides modeled into RebH (PDBID 2E4G) using FlexPepDock server as a part of the ROSIE2 commons. The catalytic lysine (K79) is shown as an orange stick for reference. (A) Cartoon view of WGGG modeled into RebH shown as magenta sticks. (B) Surface view of WGGG modeled into RebH shown as magenta sticks. (C) Cartoon view of GGWGG modeled into RebH shown as magenta sticks. (D) Surface view of GGWGG modeled into RebH shown as magenta sticks. (E) Cartoon view of GGGW modeled into RebH shown as magenta sticks. (F) Surface view of GGGW modeled into RebH shown as magenta sticks.

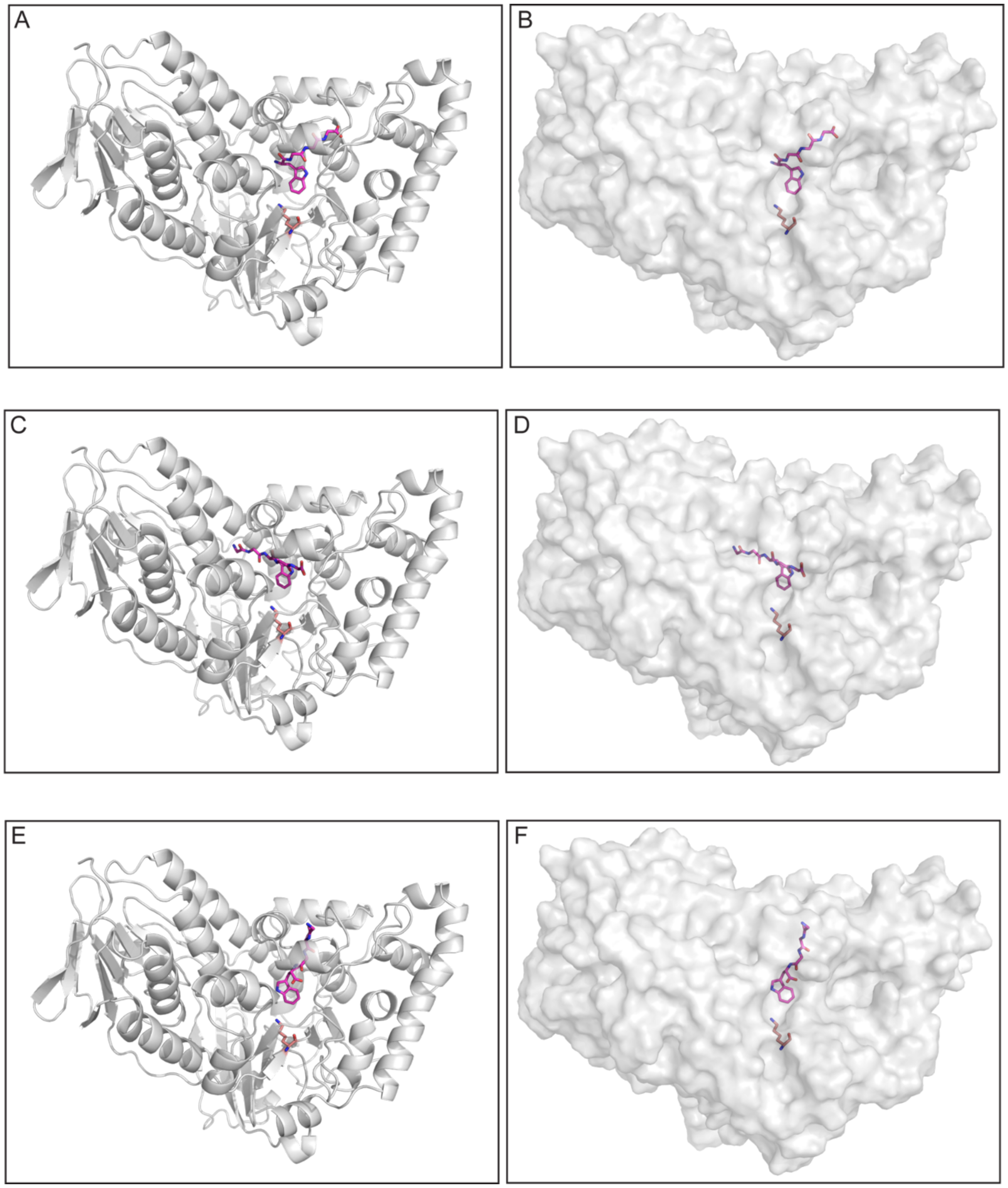

**Supplementary Figure 19. RebH- and 4V-catalyzed mono- and di-bromination of peptides with a central Trp residue.** a) RebH-catalyzed mono- (lavender) and dibromination (grey) of central Trp-containing glycine peptides. b) 4V-catalyzed mono- (lavender) and dibromination (grey) of central Trp-containing glycine peptides.

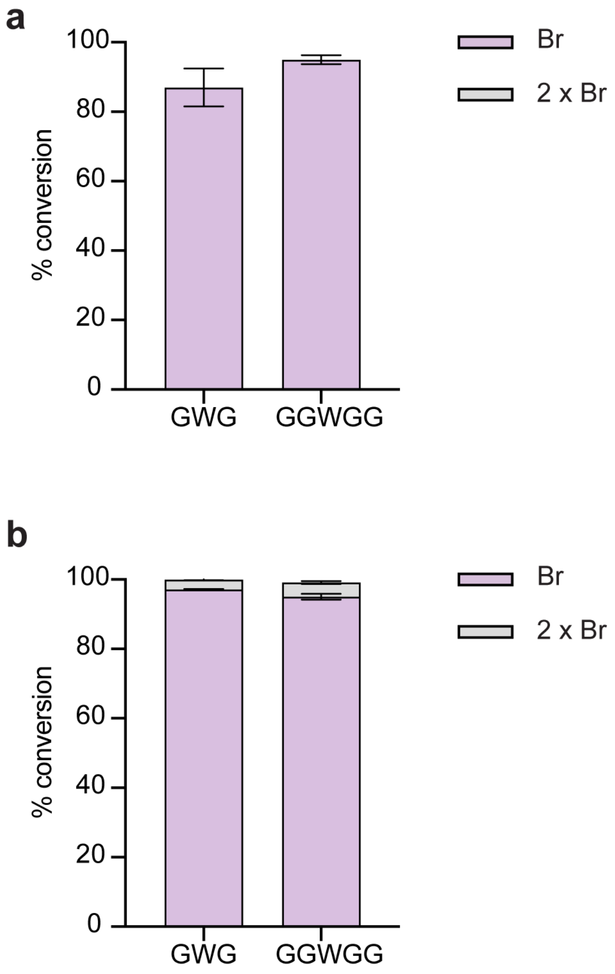

**Supplementary Figure 20. RebH and 4V bromination activity on dipeptides.** (a) Percent conversion for RebH bromination of N-terminal Trp dipeptides. (b) Percent conversion for 4V bromination of N-terminal Trp dipeptides. (c) Percent conversion for RebH bromination of C-terminal Trp dipeptides. (d) Percent conversion for 4V bromination of C-terminal Trp dipeptides. (e) No enzyme control for N-terminal Trp dipeptides. (f) No enzyme control for C-terminal Trp dipeptides.

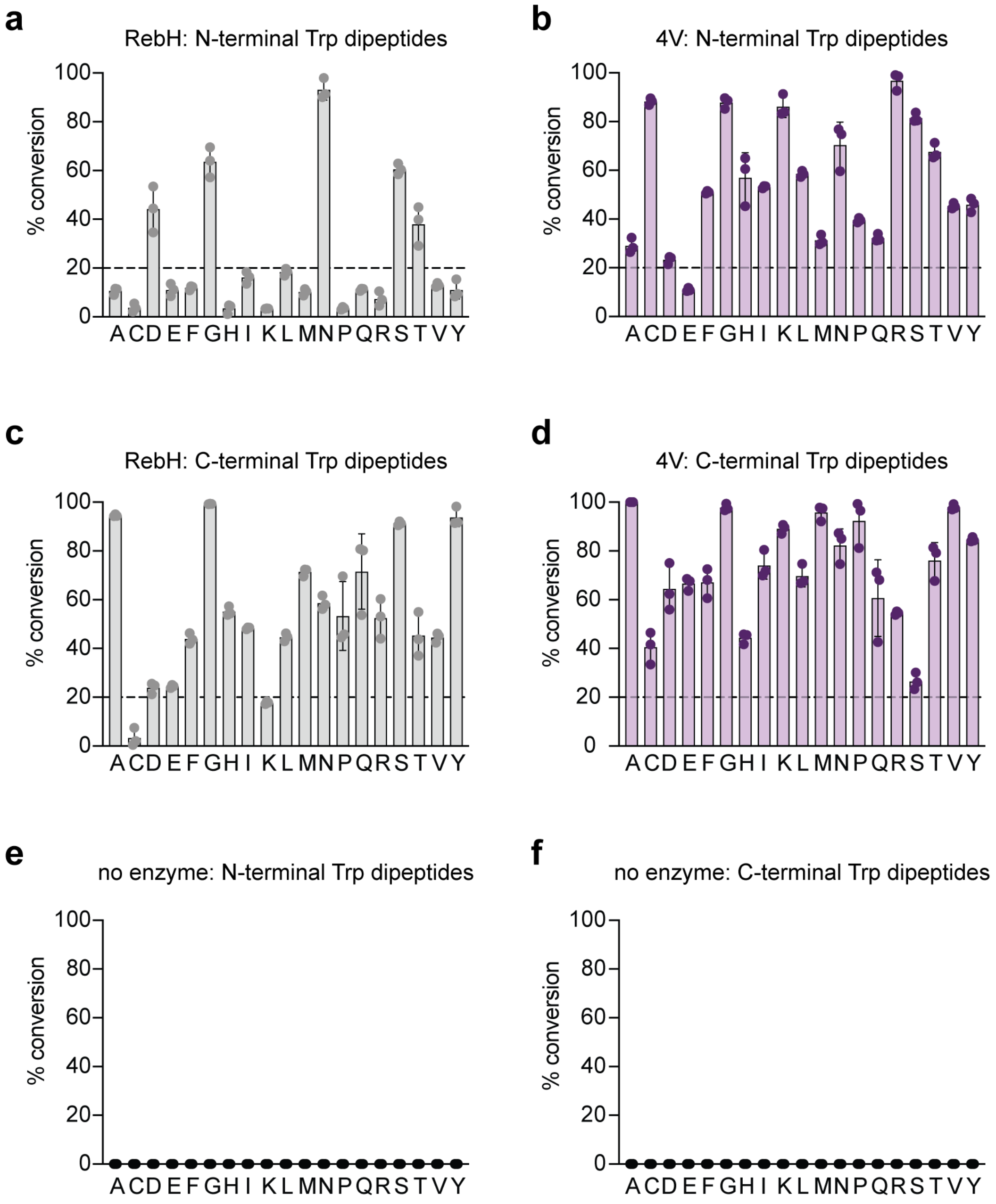

**Supplementary Figure 21. RebH- and 4V-catalyzed mono- and di-bromination of Trp-containing dipeptides.** a) RebH-catalyzed mono- (lavender) and dibromination (grey) of N-terminal Trp dipeptides. b) 4V-catalyzed mono- (lavender) and dibromination (grey) of N-terminal Trp dipeptides. c) RebH-catalyzed mono- (lavender) and dibromination (grey) of C-terminal Trp dipeptides. d) 4V-catalyzed mono- (lavender) and dibromination (grey) of C-terminal Trp dipeptides.

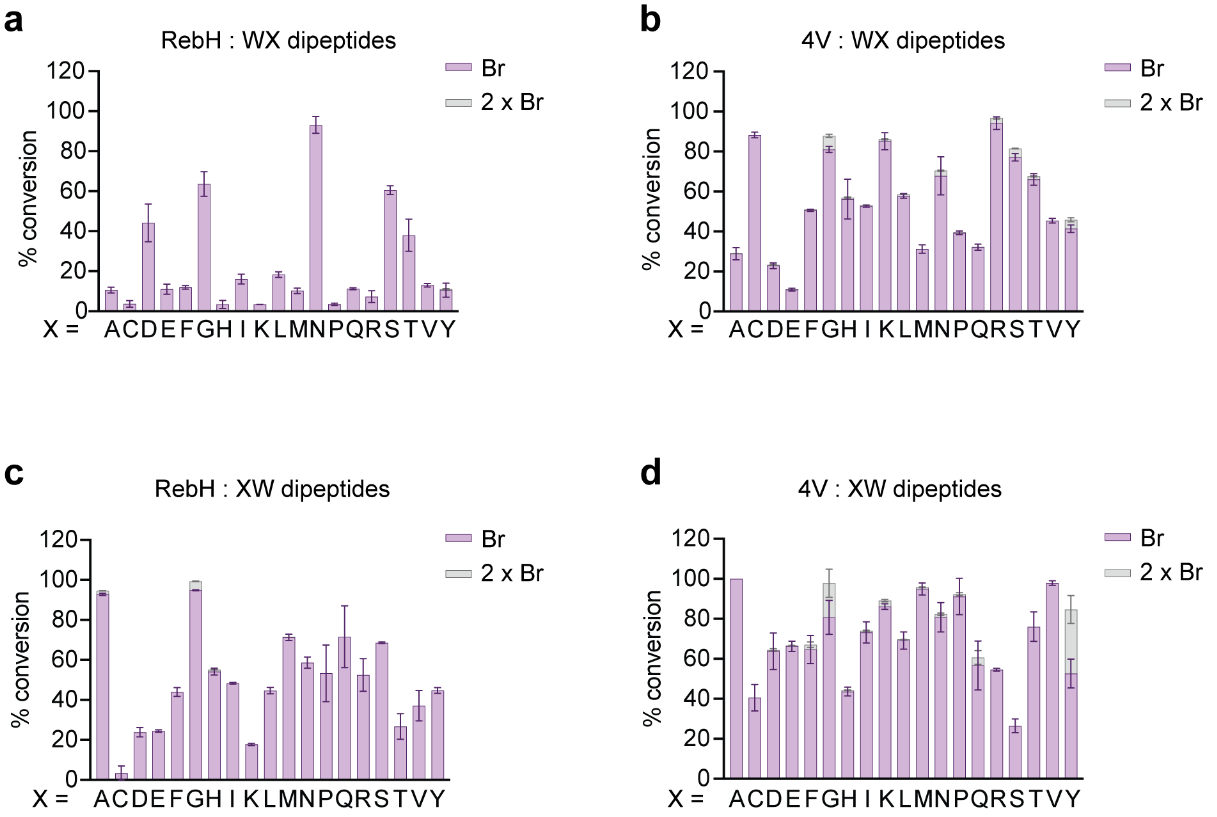

**Supplementary Figure 22. Distances between L-Trp and residues 467 and 470 in RebH and 4V.** (a) In RebH, N467 and N470 are outside hydrogen bonding distance from the l-Trp substrate but are ideally positioned to engage an extended C terminus. (b) In 4V, T467 and S470 are outside hydrogen bonding distance from the l-Trp substrate but are ideally positioned to engage an extended C terminus. Figures were generated from PDB ID 24EG. For (b), N467 and N470 were changed to T and S, respectively, using PyMOL’s mutagenesis tool.

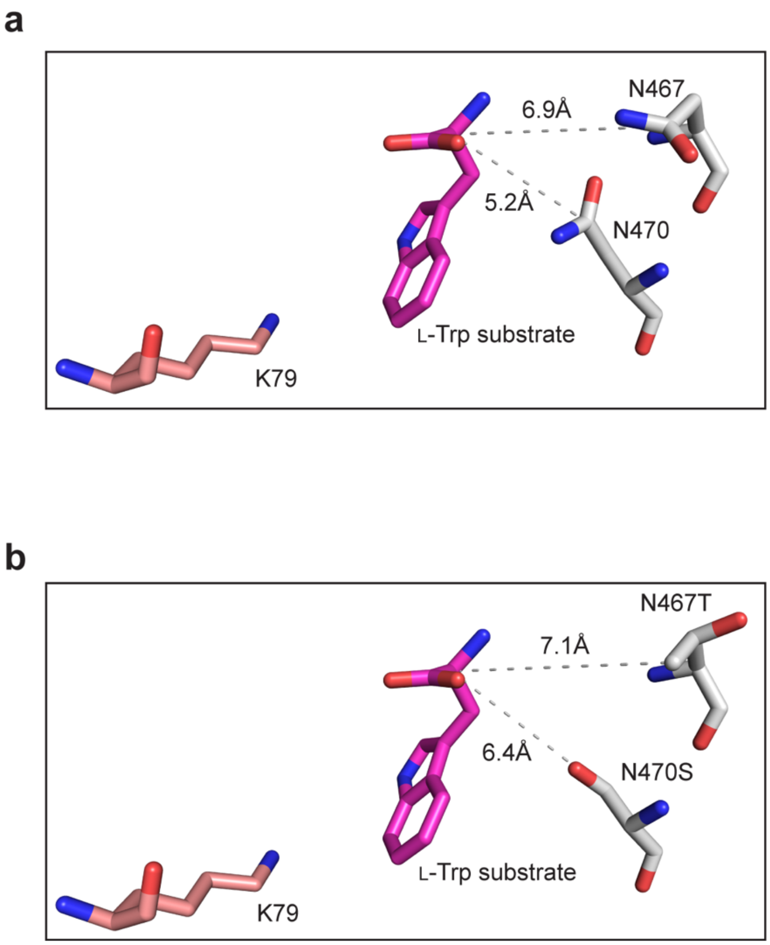

**Supplementary Figure 23. RebH- and 4V-catalyzed bromination of N-terminal Trp tripeptides.** (a) Percent conversion for RebH- and 4V-catalyzed bromination of WGX tripeptides. (b) Percent conversion for RebH- and 4V-catalyzed bromination of WXG tripeptides.

a

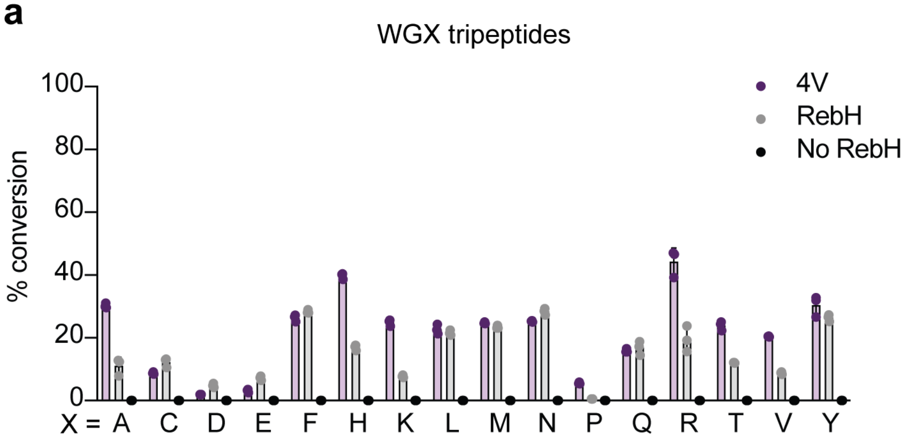

b

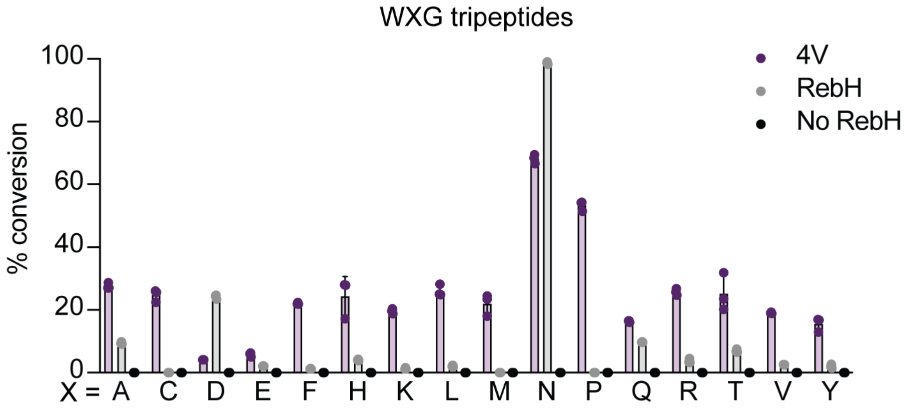

**Supplementary Figure 24. RebH- and 4V-catalyzed bromination of central Trp tripeptides.** (a) Percent conversion for RebH- and 4V-catalyzed bromination of GWX tripeptides. (b) Percent conversion for RebH- and 4V-catalyzed bromination of XWG tripeptides.

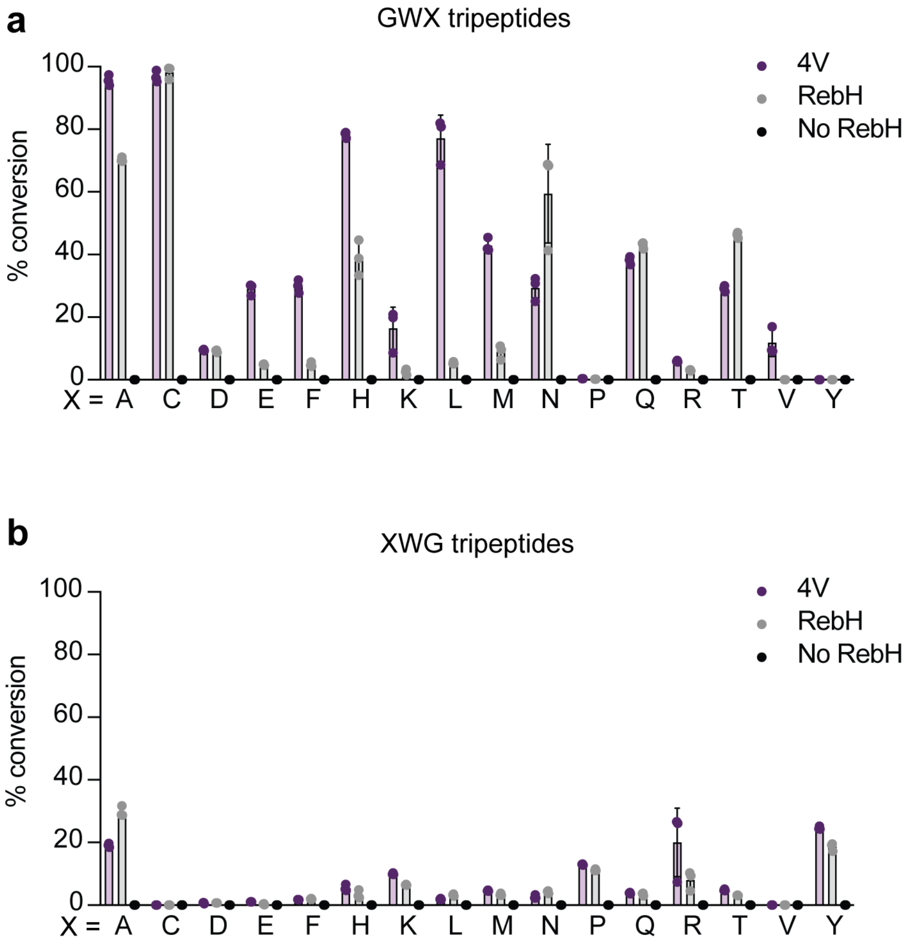

**Supplementary Figure 25. RebH- and 4V-catalyzed bromination of central Trp tripeptides.** (a) Percent conversion for RebH- and 4V-catalyzed bromination of GXW tripeptides. (b) Percent conversion for RebH- and 4V-catalyzed bromination of XGW tripeptides.

**a**

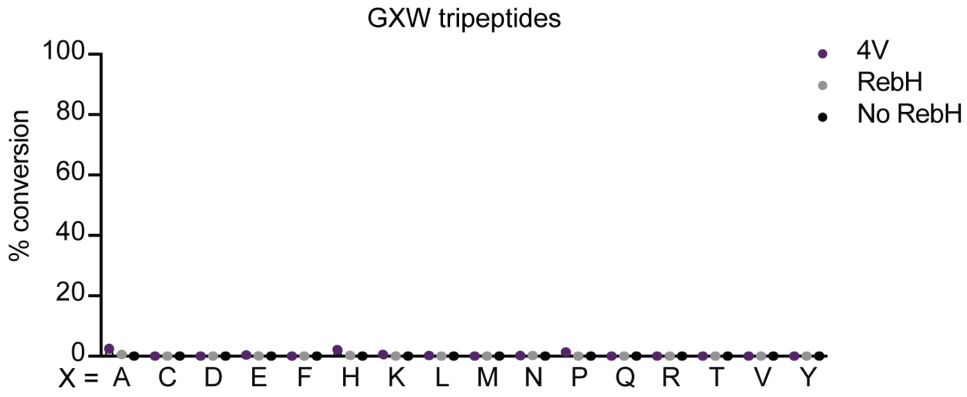

**b**

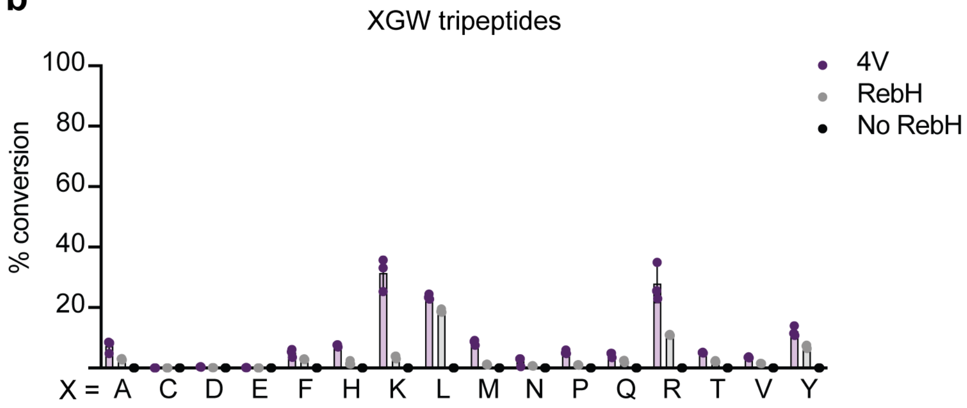

**Supplementary Figure 26. RebH- and 4V-catalyzed bromination of peptides with carboxamide or carboxylate C termini.** (a) Percent conversion for RebH- and 4V-catalyzed bromination of carboxamide (lavender) or carboxylate (grey) version of peptides with C-terminal Trp. (b) Percent conversion for RebH- and 4V-catalyzed bromination of carboxamide (lavender) or carboxylate (grey) version of peptides with N-terminal Trp.

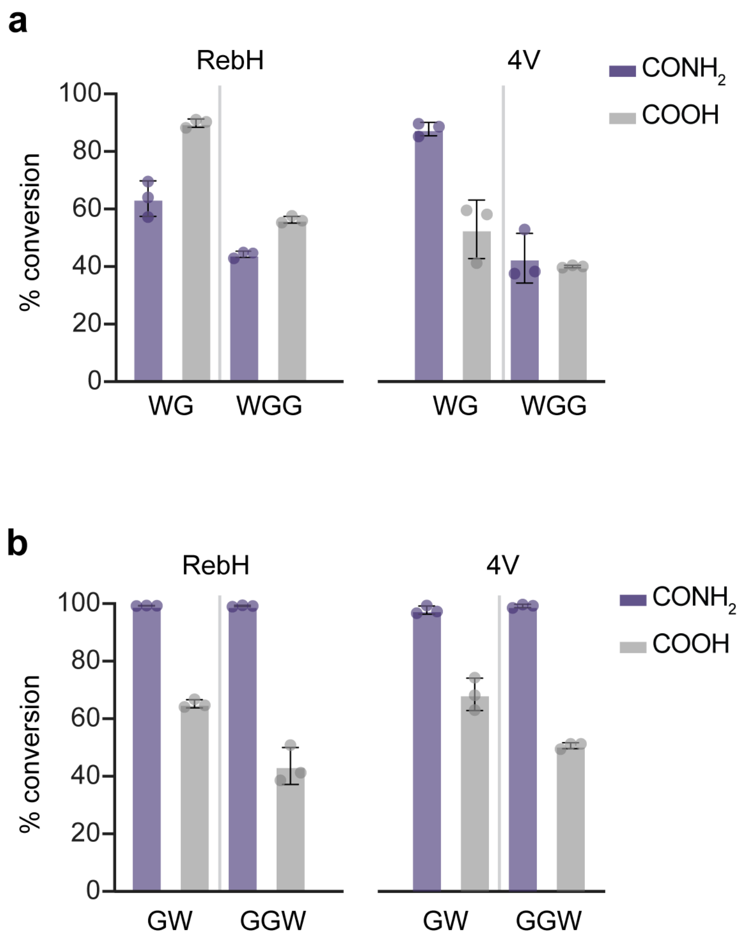

**Supplementary Figure 27. Trypsin digest analysis of 4V bromination reaction with (RW)₃.** (a) Chemical schematic of $(RW)_3$ (**1**) and fragments (**2-8**) resulting from trypsin digestion of $(RW)_3$. (b) LC-HRMS spectrum of trypsin digested 4V halogenation reaction with $(RW)_3$. (c) C-terminal brominated tryptophanamide fragment mass spectrum.

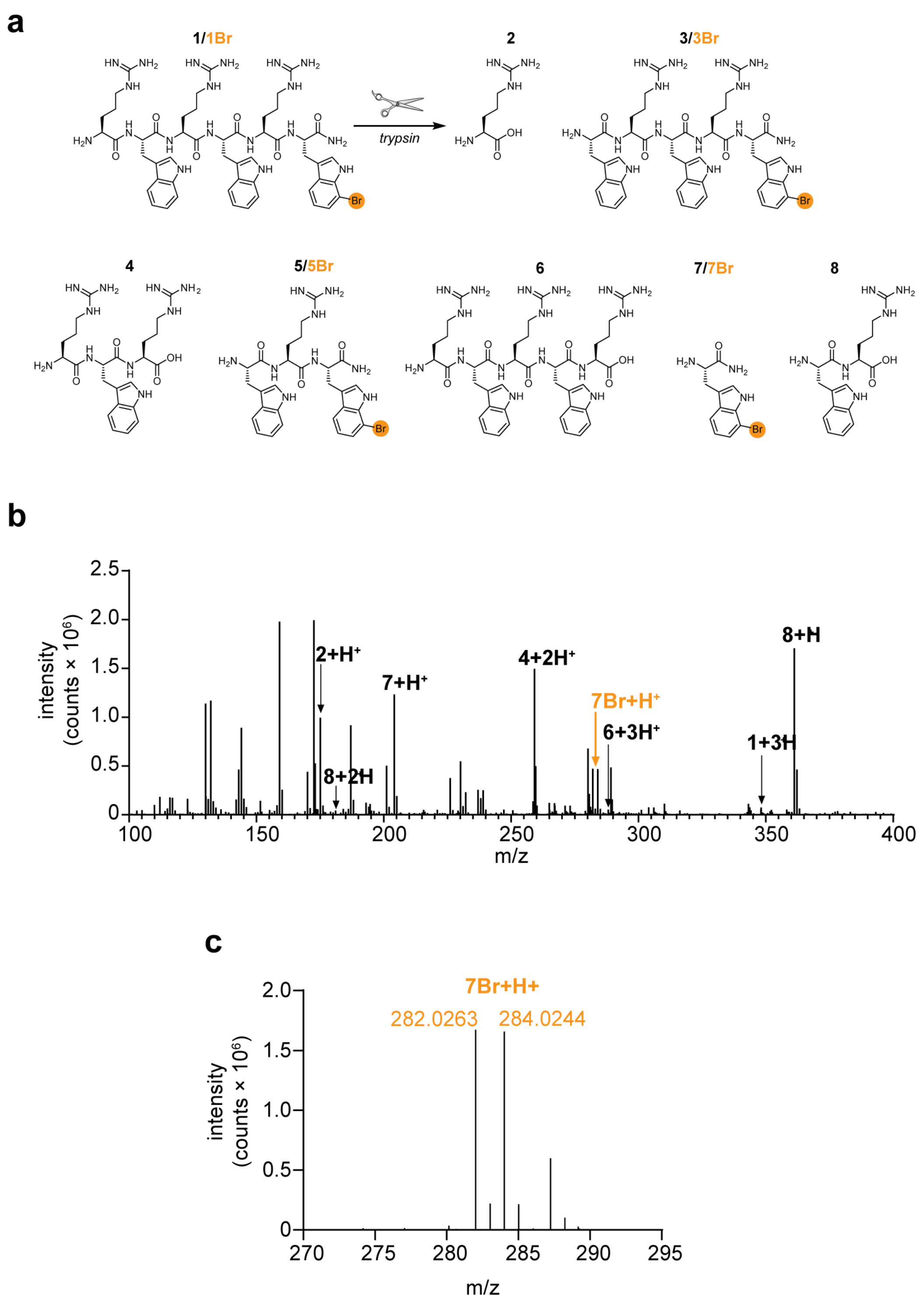

**Supplementary Figure 28. Chemical structures of $(RW)_3$ and $(RW)_4$ derivatives.** (a) $(RW)_3$. (c) Br-$(RW)_3$. (c) 3-Trifluoromethylphenyl-$(RW)_3$. (d) $(RW)_4$. (e) Br-$(RW)_4$.

**a**

$(RW)_3$

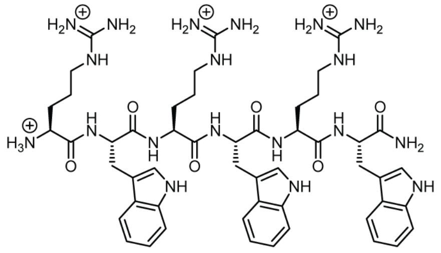

**b**

Br-$(RW)_3$

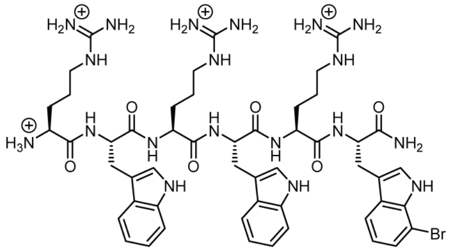

**c**

3-trifluoromethylphenyl-$(RW)_3$

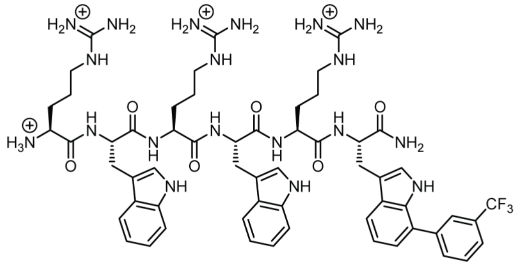

**d**

$(RW)_4$

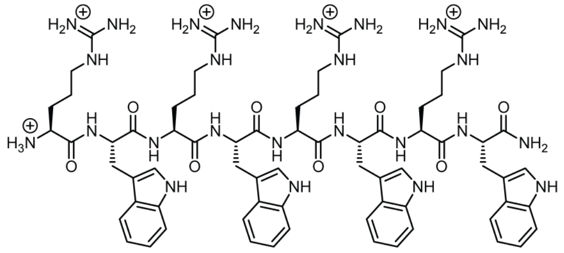

**e**

Br-$(RW)_4$

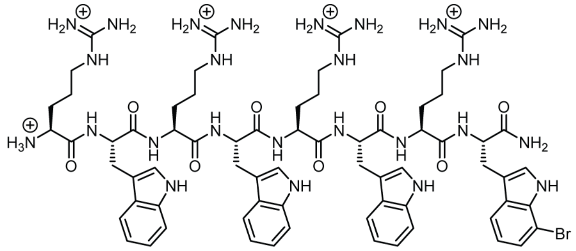

**Supplementary Figure 29. Analytical HPLC traces of $(RW)_3$ and $(RW)_4$ derivatives.** (a) $(RW)_3$. (c) Br-$(RW)_3$. (c) 3-Trifluoromethylphenyl-$(RW)_3$. (d) $(RW)_4$. (e) Br-$(RW)_4$.

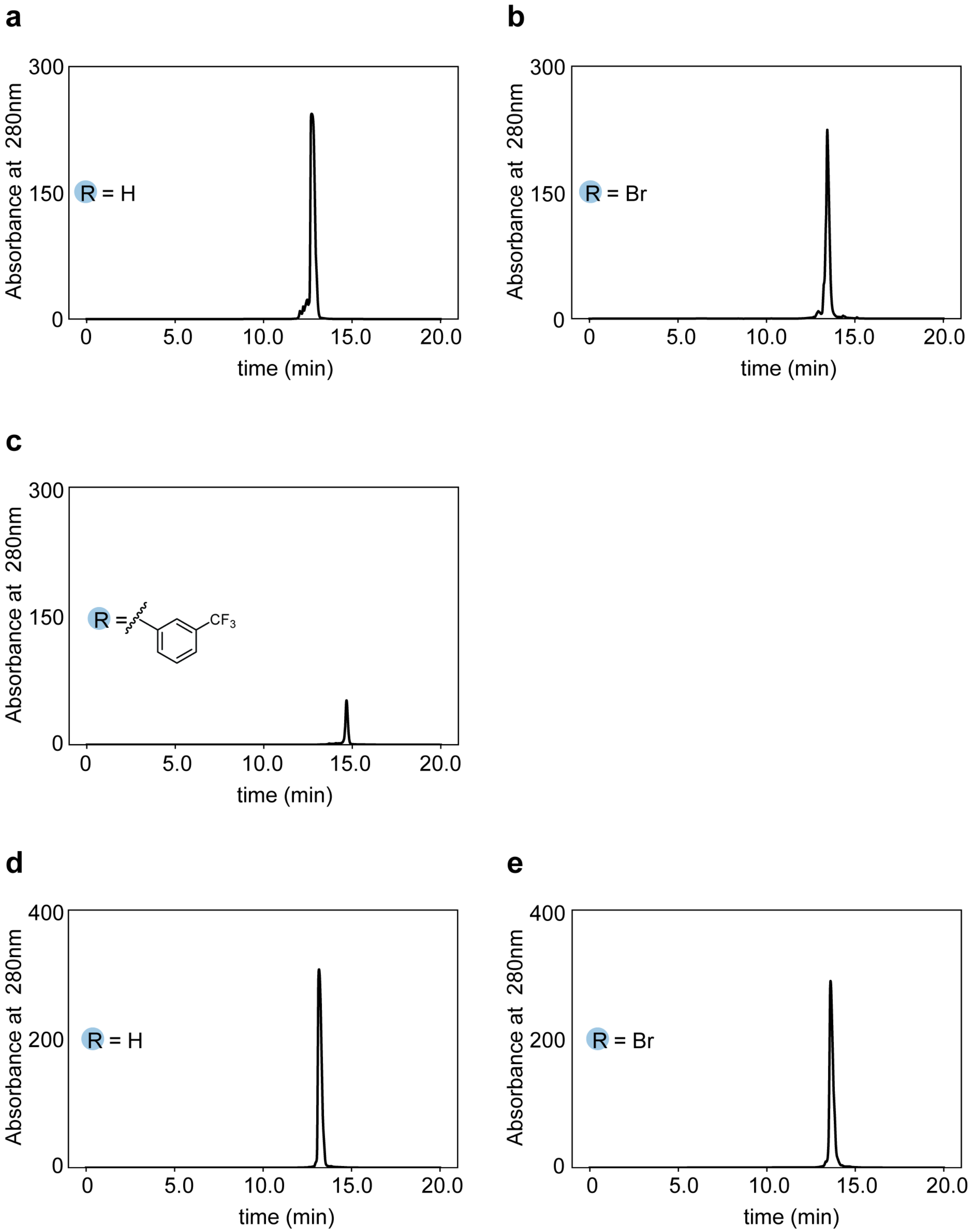

**Supplementary Table 1. Analytical HPLC purity of antimicrobial peptide derivatives.**

| Neuropeptide | A280nm main peak area | A280nm minor peak area(s) | Total A280nm peak area | % purity |
|---|---|---|---|---|
| (RW)3 | 58340.9 | 1852.2 | 60193.1 | 96.92 |
| Br-(RW)3 | 31336.6 | 1149.3 | 32485.9 | 96.46 |
| CF3-(RW)3 | 5252.7 | 204 | 5456.7 | 96.26 |
| (RW)4 | 46845.5 | 176.7 | 47022.2 | 99.62 |
| Br-(RW)4 | 44721 | 177.2 | 44898.2 | 99.61 |

**Supplementary Figure 30. Minimum inhibitory concentration (MIC) and minimum bactericidal concentration (MBC) measurements for $(RW)_3$, Br-$(RW)_3$, and 3-trifluoromethylphenyl-$(RW)_3$.** (A) MIC triplicate data for *E. coli* grown in the presence of 2-fold dilutions of $(RW)_3$ and Br-$(RW)_3$. (B) $OD_{600}$ quantitation of raw MIC triplicate data for *E. coli* grown in the presence of 2-fold dilutions of $(RW)_3$ and Br-$(RW)_3$. (C) MBC spot assay triplicate data for *E. coli* grown in the presence of 2-fold dilutions of $(RW)_3$ and Br-$(RW)_3$. (D) MIC triplicate data for *E. coli* grown in the presence of 2-fold dilutions of Br-$(RW)_3$ and 3-trifluoromethylphenyl-$(RW)_3$. (E) $OD_{600}$ quantitation of raw MIC triplicate data for *E. coli* grown in the presence of 2-fold dilutions of Br-$(RW)_3$ and 3-trifluoromethylphenyl-$(RW)_3$. (F) MBC spot assay triplicate data for *E. coli* grown in the presence of 2-fold dilutions Br-$(RW)_3$ and 3-trifluoromethylphenyl-$(RW)_3$.

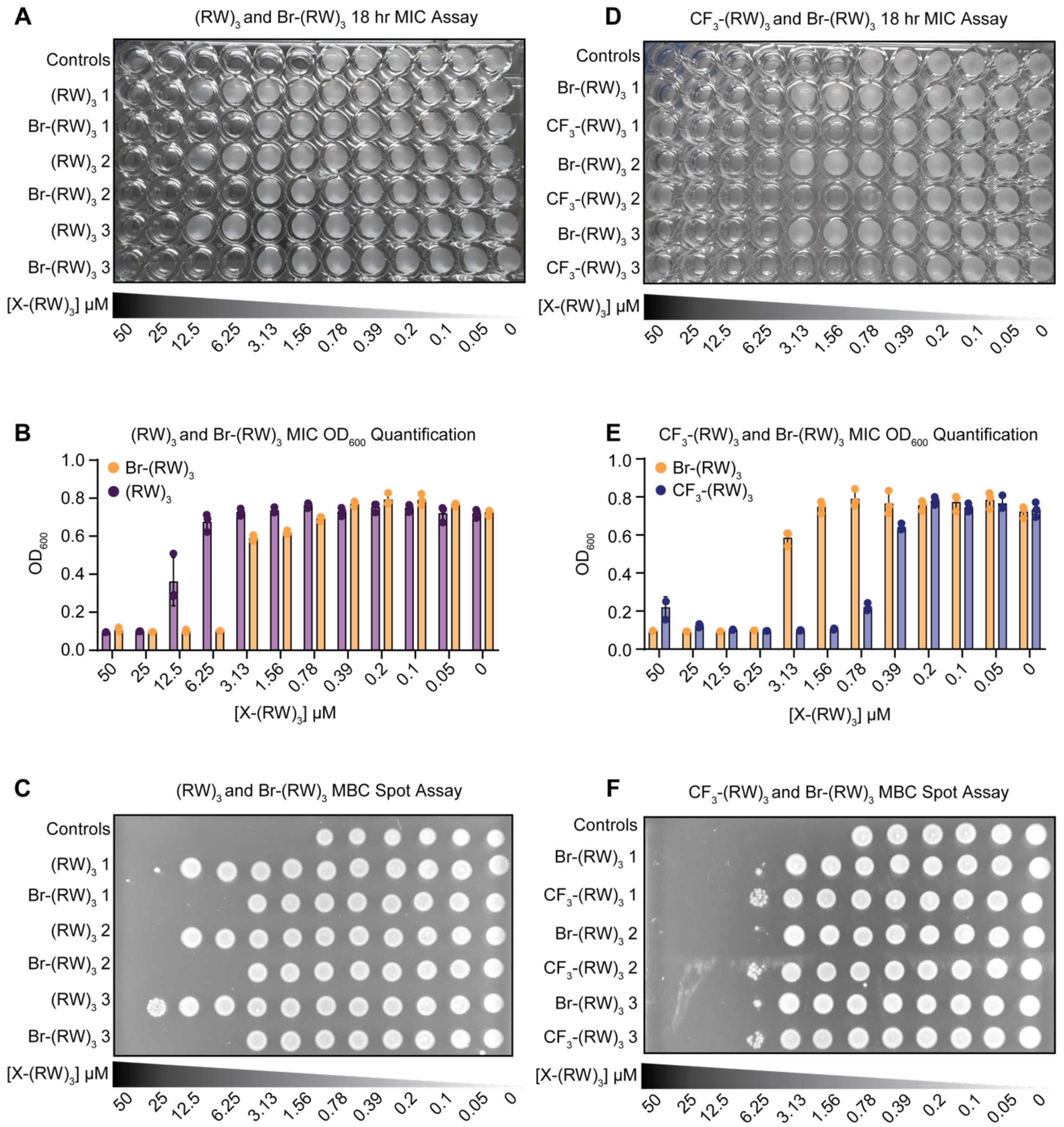

**Supplementary Figure 31. Large-scale MBC assays for $(RW)_3$ and derivatives.** (A) MBC large-scale Petri dish triplicate data for *E. coli* grown in the presence of 2-fold dilutions of $(RW)_3$. (B) MBC large-scale Petri dish triplicate data for *E. coli* grown in the presence of 2-fold dilutions of Br-$(RW)_3$. (C) MBC large-scale Petri dish triplicate data for *E. coli* grown in the presence of 2-fold dilutions of 3-trifluoromethylphenyl-$(RW)_3$. Negative and positive growth controls are shown on the left of each Petri dish figure.

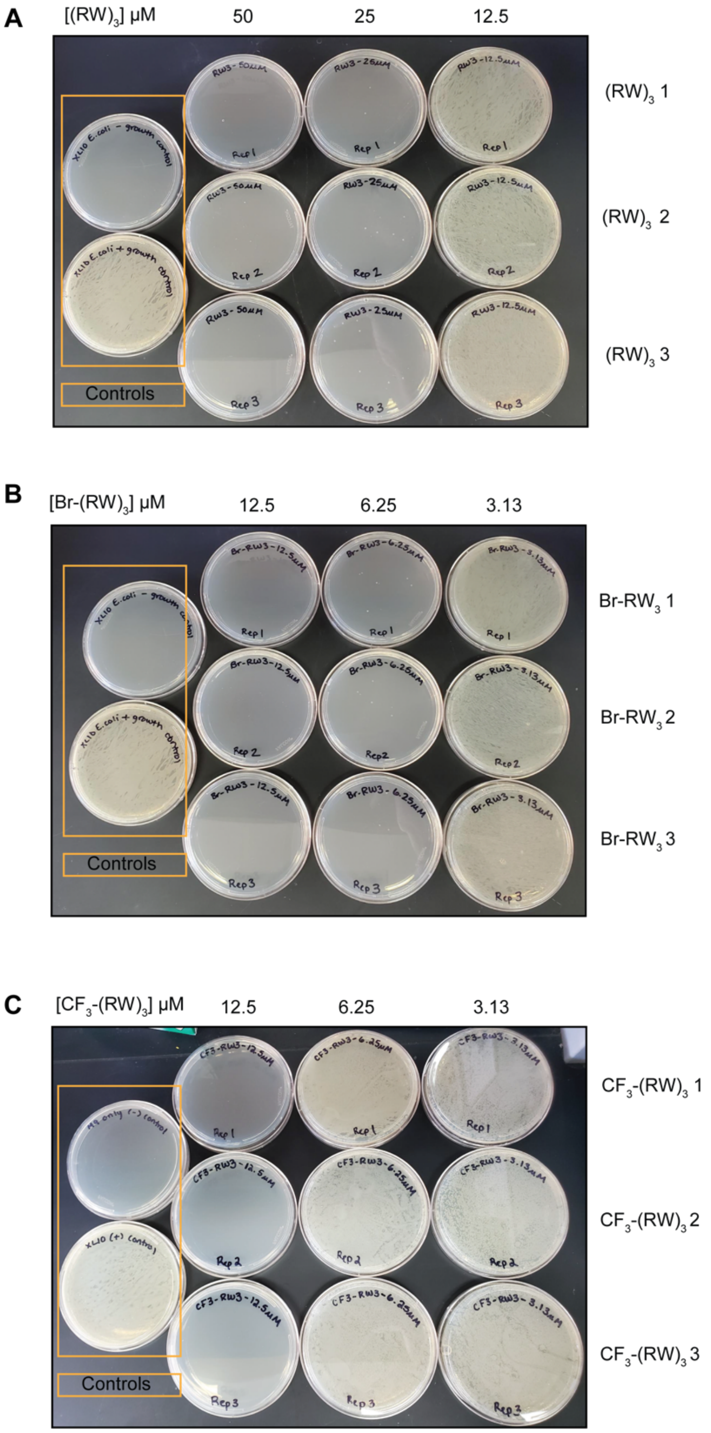

**Supplementary Figure 32. Minimum inhibitory concentration (MIC) and minimum bactericidal concentration (MBC) measurements for $(RW)_3$ and Br-$(RW)_4$.** (A) MIC triplicate data for *E. coli* grown in the presence of 2-fold dilutions of $(RW)_4$ and Br-$(RW)_4$. (B) $OD_{600}$ quantitation of raw MIC triplicate data for *E. coli* grown in the presence of 2-fold dilutions of $(RW)_4$ and Br-$(RW)_4$. (C) MBC spot assay triplicate data for *E. coli* grown in the presence of 2-fold dilutions of $(RW)_4$ and Br-$(RW)_4$. (D) MBC large-scale Petri dish triplicate data for *E. coli* grown in the presence of 2-fold dilutions of $(RW)_4$. (E) MBC large-scale Petri dish triplicate data for *E. coli* grown in the presence of 2-fold dilutions of Br-$(RW)_4$. Negative and positive growth controls are shown on the left of each Petri dish figure.

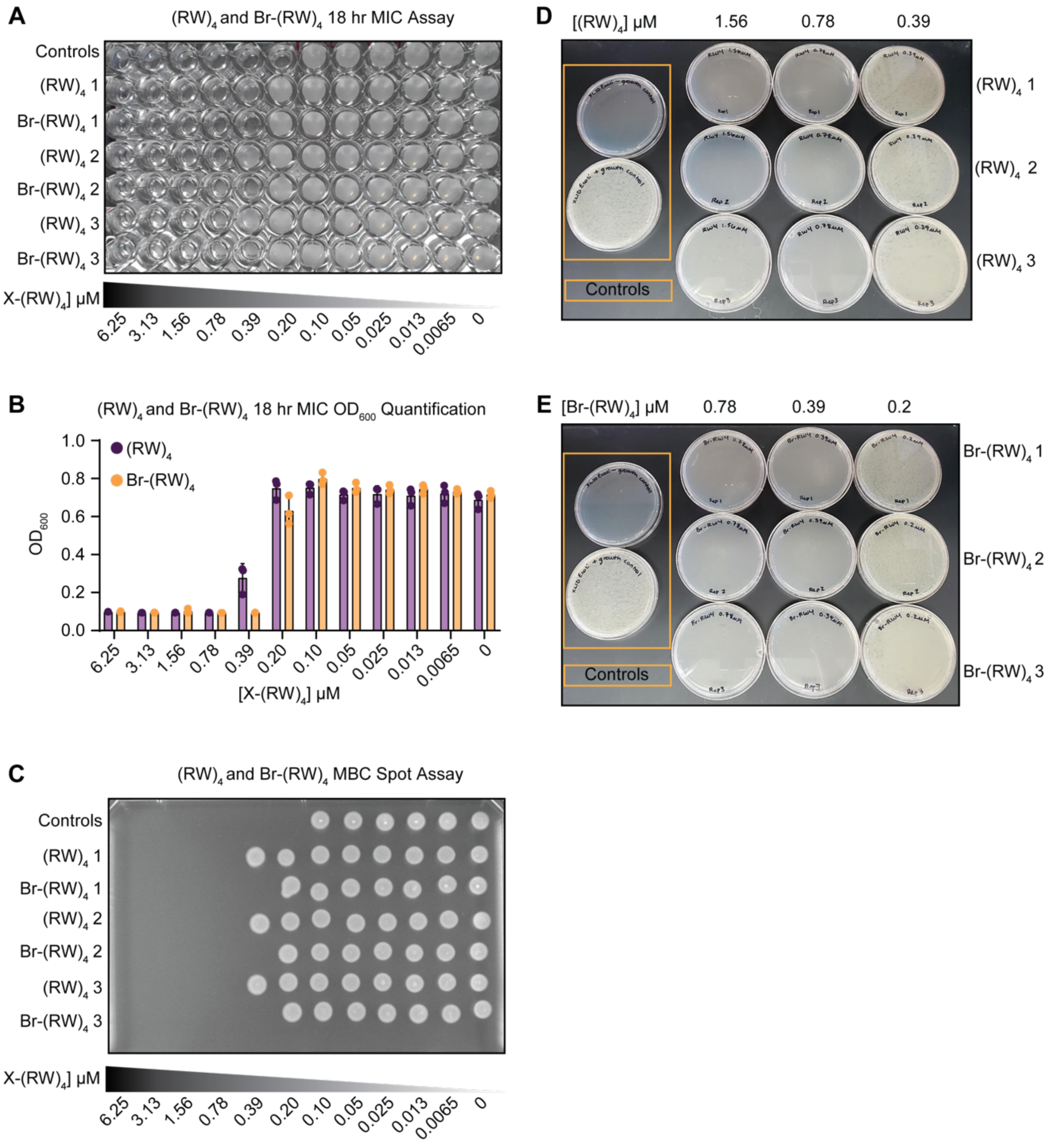

**Supplementary Figure 33. Analytical HPLC traces of transportan derivatives.** (A) Transportan. (B) Br-Transportan. (C) Naphthyl-transportan (Nap-TN).

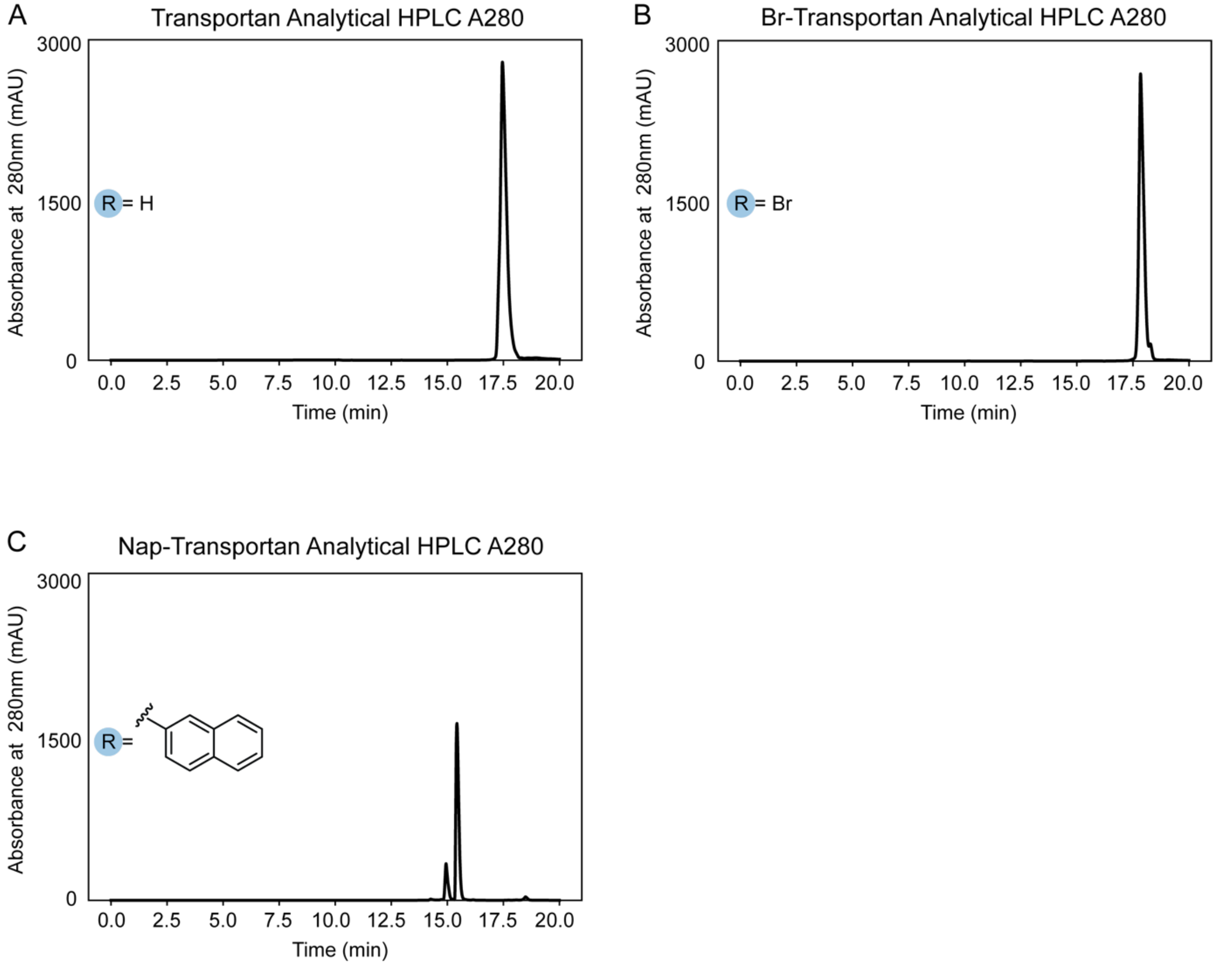

**Supplementary Table 2. Analytical HPLC purity of transportan derivatives.**

| Neuropeptide | A280nm main peak area | A280nm minor peak area(s) | Total A280nm peak area | % purity |
|---|---|---|---|---|
| Transportan | 22304.8 | 15.3 | 22320.1 | 99.93 |
| Br-Transportan | 140019 | 381.4 | 140400.4 | 99.73 |
| Nap-Transportan | 9914.3 | 1984.7 | 11899 | 83.33 |

**Supplementary Figure 34. L-Endomorphin and derivatives synthesized via enzymatic bromination and Suzuki-Miyaura coupling.** (a) [L-Trp 3]endomorphin-1 (EM-1). (b) [Br-L-Trp 3]EM-1. (c) [*p*-tolyl-L-Trp 3]EM-1. (d) [3-trifluoromethylphenyl-L-Trp 3]EM-1. (e) [Naphthyl-L-Trp 3]EM-1.

**a**

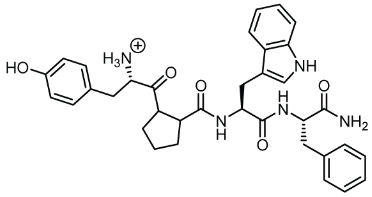

**b**

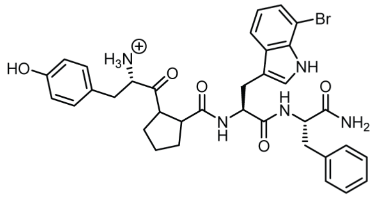

**c**

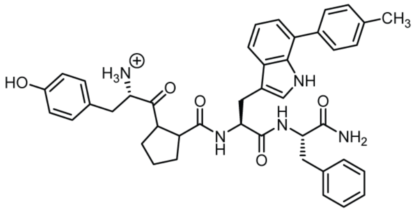

**d**

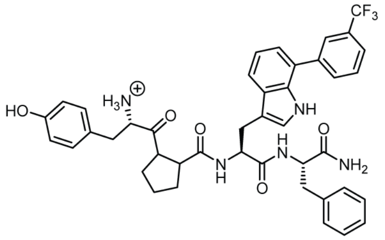

**e**

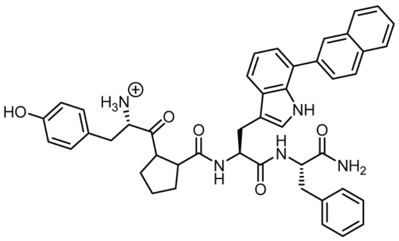

**Supplementary Figure 35. Analytical HPLC traces for L-Endomorphin and derivatives .** (a) [L-Trp 3]endomorphin-1 (EM-1). (b) [Br-L-Trp 3]EM-1. (c) [*p*-tolyl-L-Trp 3]EM-1. (d) [3-trifluoromethylphenyl-L-Trp 3]EM-1. (e) [Naphthyl-L-Trp 3]EM-1.

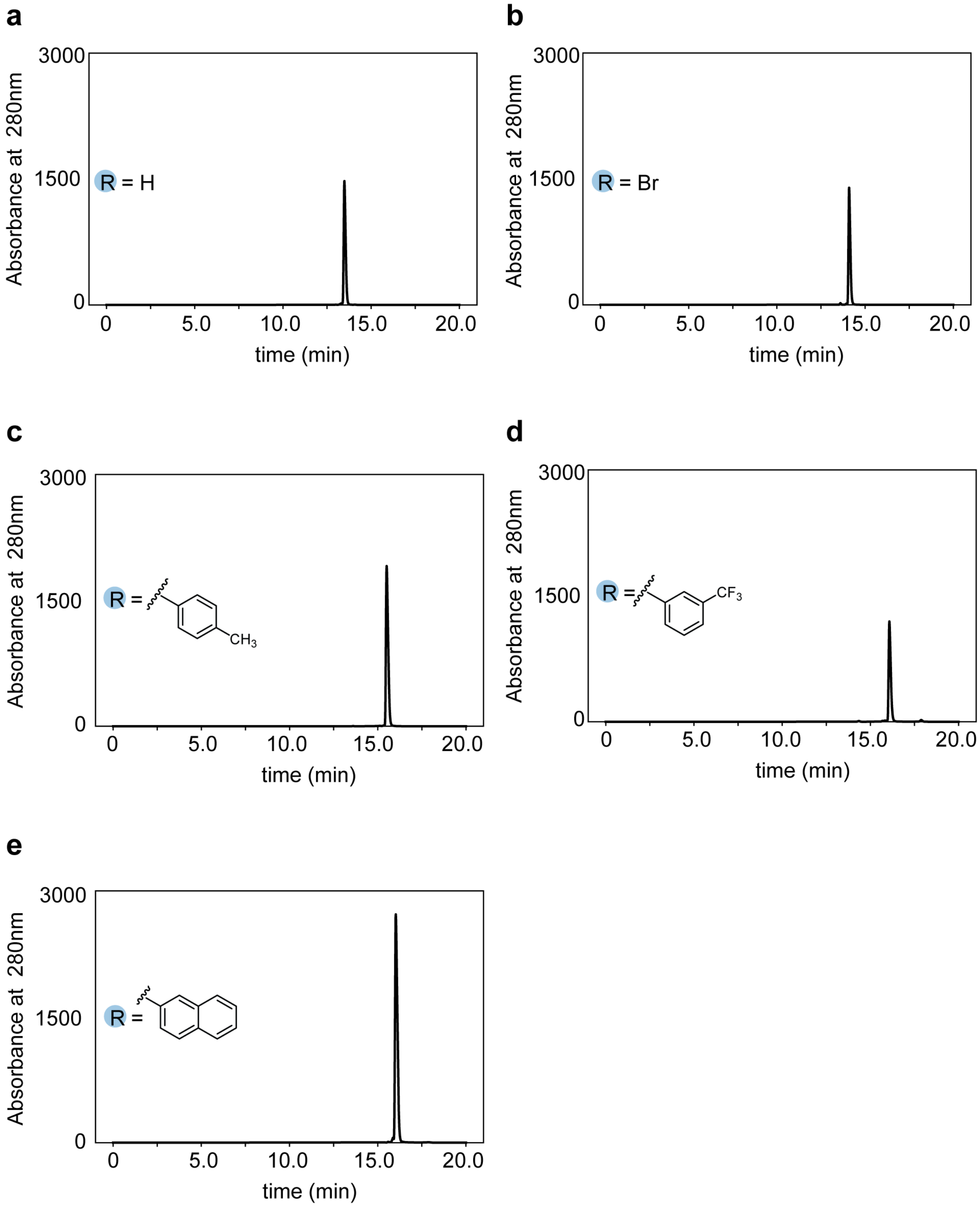

**Supplementary Figure 36. D-Endomorphin and derivatives synthesized via enzymatic bromination and Suzuki-Miyaura coupling.** (a) [D-Trp 3]endomorphin-1 (EM-1). (b) [Br-D-Trp 3]EM-1. (c) [*p*-tolyl-D-Trp 3]EM-1. (d) [3-trifluoromethylphenyl-D-Trp 3]EM-1. (e) [Naphthyl-D-Trp 3]EM-1.

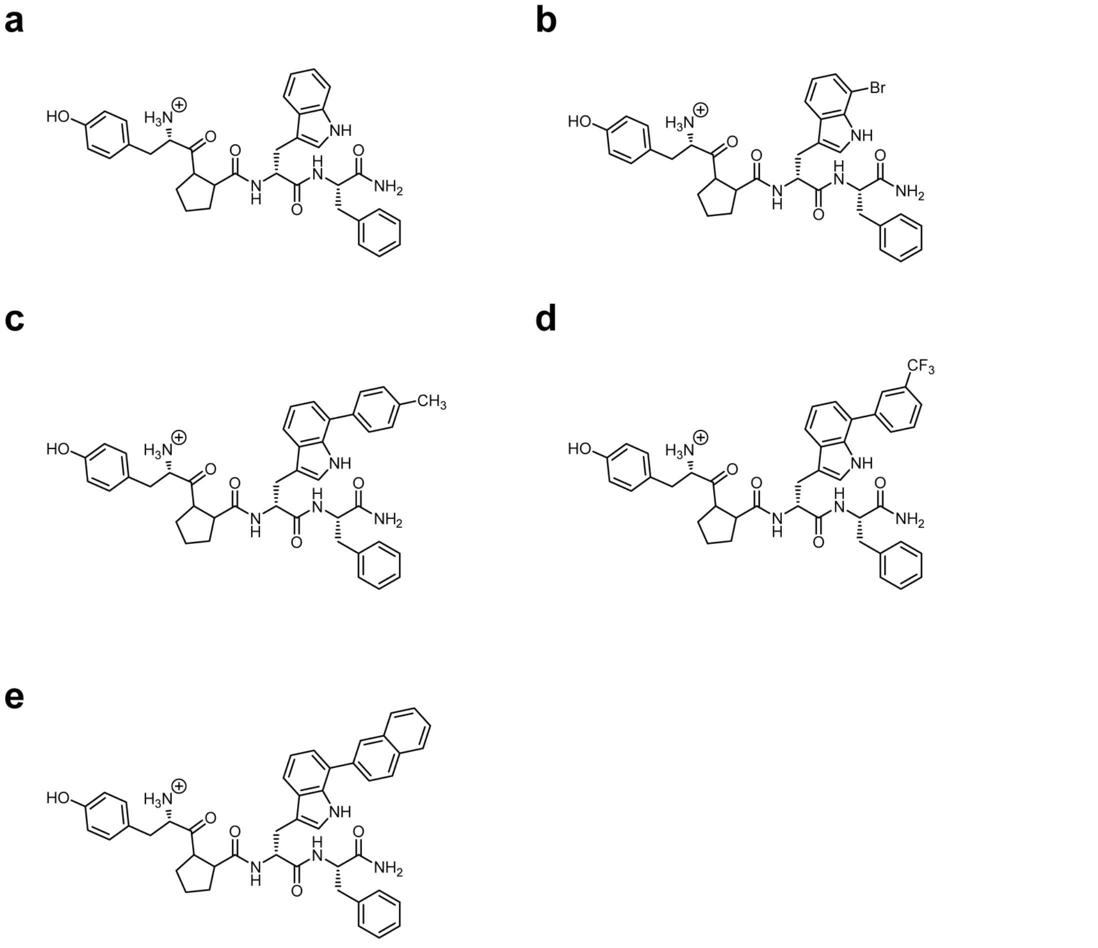

**Supplementary Figure 37. Analytical HPLC traces for D-Endomorphin and derivatives.** (a) [D-Trp 3]endomorphin-1 (EM-1). (b) [Br-D-Trp 3]EM-1. (c) [*p*-tolyl-D-Trp 3]EM-1. (d) [3-trifluoromethylphenyl-D-Trp 3]EM-1. (e) [Naphthyl-D-Trp 3]EM-1.

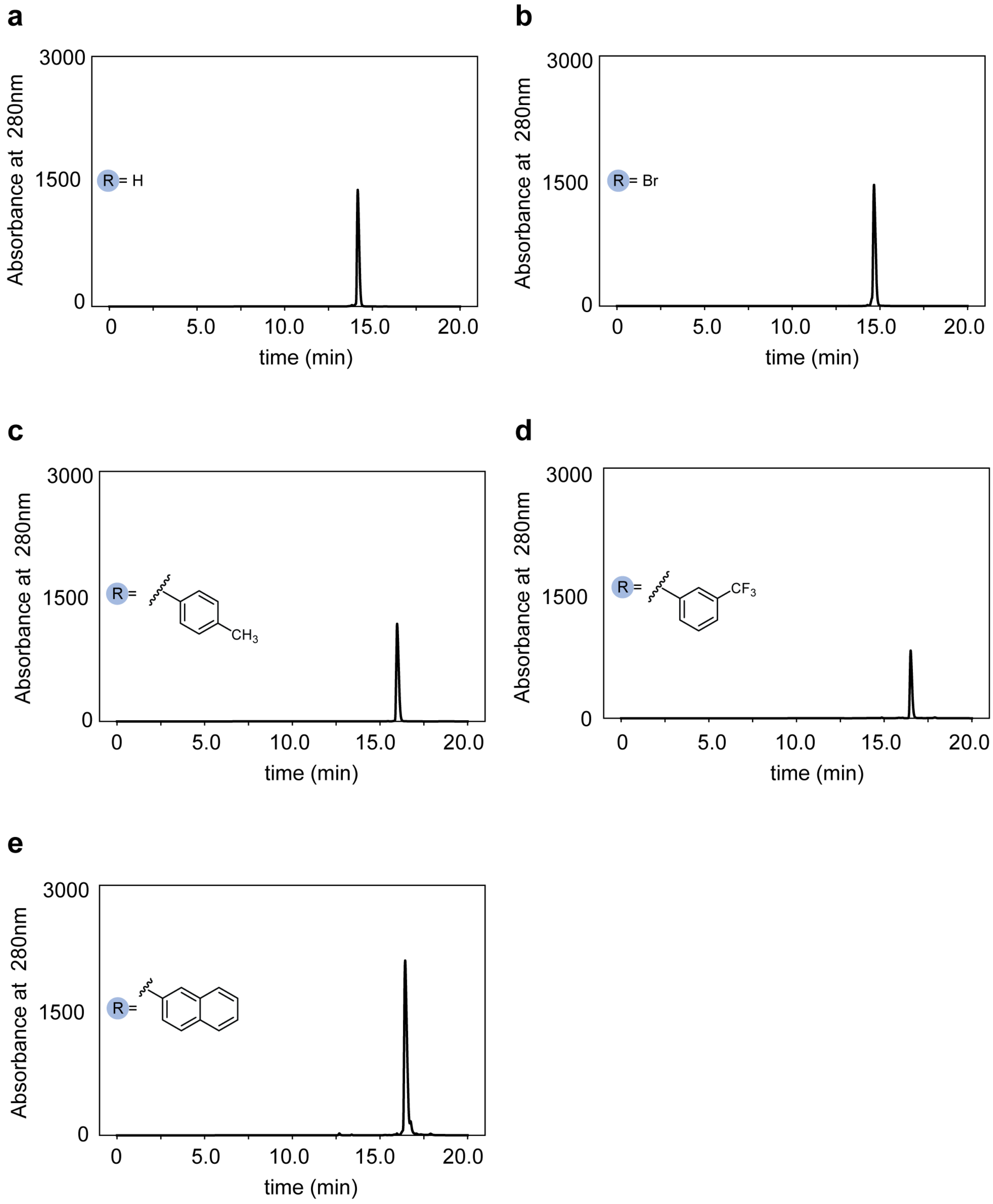

**Supplementary Table 3. Analytical HPLC purity of endomorphin-1 derivatives.**

| Endomorphin-1 derivative | A280nm main peak area | A280nm minor peak(s) area | Total A280nm peak area | % purity |
|---|---|---|---|---|
| L-Endomorphin-1 | 11254.4 | 199.4 | 11453.8 | 98.26 |
| Br-L-Endomorphin-1 | 10094.1 | 209.1 | 10303.2 | 97.97 |
| Br2-L-Endomorphin-1 | 31719.4 | 189.6 | 31909 | 99.41 |
| Ptolyl-L-Endomorphin-1 | 16884.3 | 89 | 16973.3 | 99.48 |
| CF3-L-Endomorphin-1 | 10130.1 | 522.2 | 10652.3 | 95.09 |
| Nap-L-Endomorphin-1 | 25716.6 | 476.7 | 26193.3 | 98.18 |
| D-Endomorphin-1 | 12241.8 | 163.7 | 12405.5 | 98.68 |
| Br-D-Endomorphin-1 | 14170.5 | 84.4 | 14254.9 | 99.41 |
| Br2-D-Endomorphin-1 | 8286.4 | 202.8 | 8489.2 | 97.61 |
| Ptolyl-D-Endomorphin-1 | 11561.4 | 42.2 | 11603.6 | 99.64 |
| CF3-D-Endomorphin-1 | 7027 | 263 | 7290 | 96.39 |
| Nap-D-Endomorphin-1 | 23504.2 | 461.9 | 23966.1 | 98.07 |

**Supplementary Figure 38. Immunostaining of μ opioid receptor-transfected HEK293T cells.** Fixed transfected HEK293T cells were stained with anti-1D4 mouse μOR antibody and secondary goat anti mouse-AlexaFluor 488 antibody (imaged in the FITC channel). Channels (from left) are brightfield, DAPI nuclear stain (blue), FITC (green), and overlay of DAPI and FITC channels.

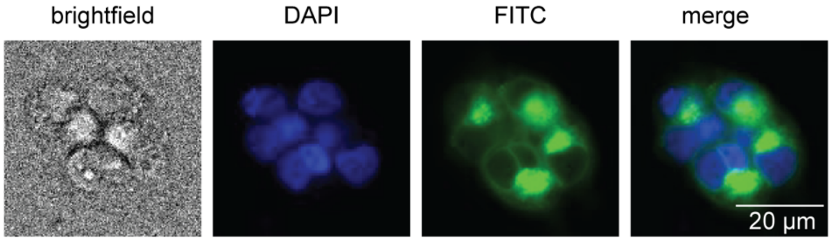

**Supplementary Figure 39. Kinetic traces of μOR GloSensor assay for L-endomorphin-1 derivatives.** HEK293T cells transfected with μOR and GloSensor 22F were treated with GloSensor reagent, stimulated with L-endomorphin-1 derivatives followed by forskolin, and luminescence was monitored for 2 h.( A) Unmodified L-endomorphin-1 (EM-1) stimulation. (B) [Br-L-Trp 3]EM-1 stimulation. (C) [$Br_2$-L-Trp 3]EM-1 stimulation. (D) [*p*-tolyl-L-Trp 3]EM-1 stimulation. (E) [Nap-L-Trp 3]EM-1 stimulation. (F) [3-trifluoromethylphenyl-L-Trp]EM-1 stimulation.

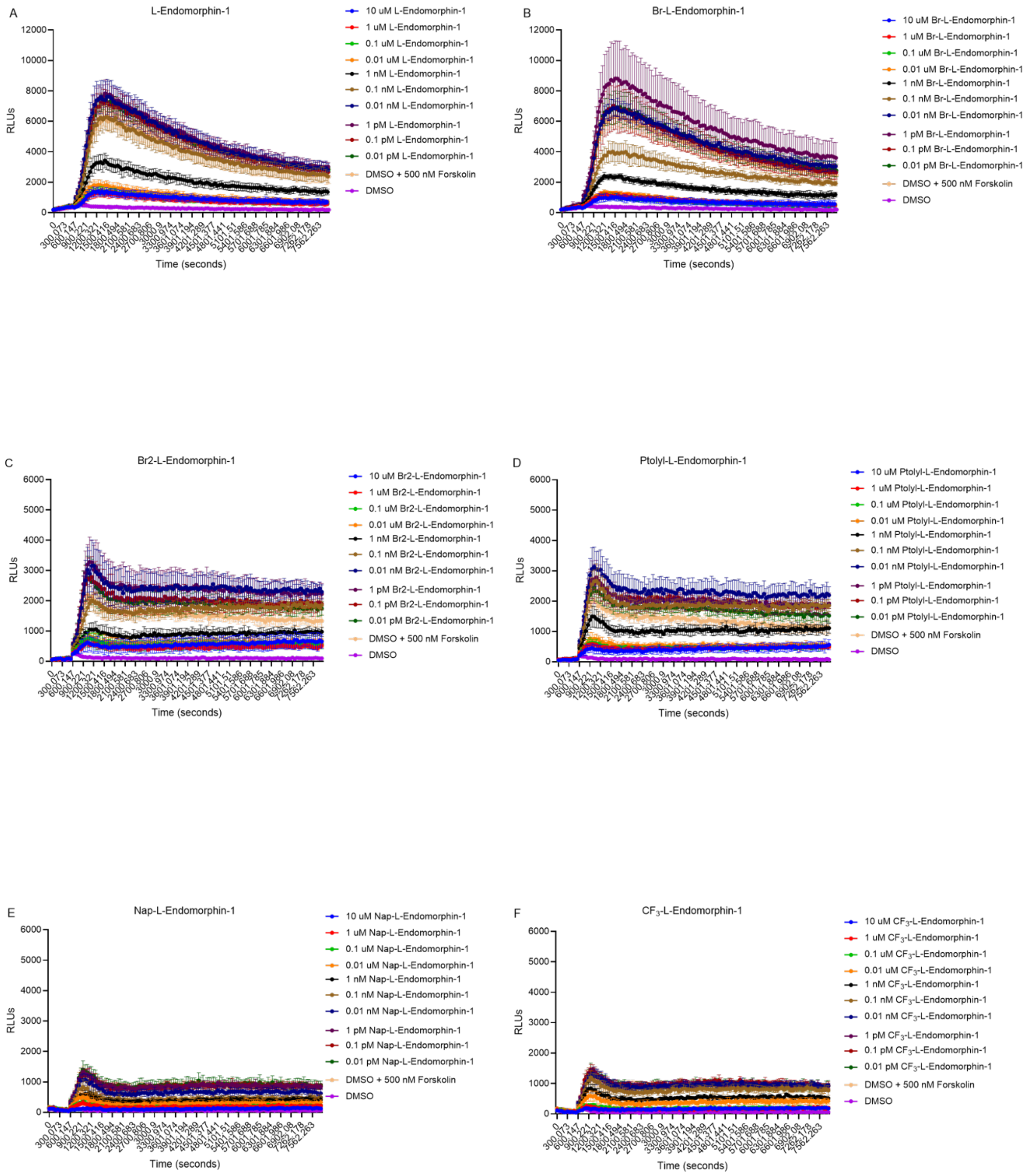

**Supplementary Figure 40. Kinetic traces of μOR GloSensor assay for D-endomorphin-1 derivatives.** HEK293T cells transfected with μOR and GloSensor 22F were treated with GloSensor reagent, stimulated with D-endomorphin-1 derivatives followed by forskolin, and luminescence was monitored for 2 h.( A) Unmodified D-endomorphin-1 (EM-1) stimulation. (B) [Br-D-Trp 3]EM-1 stimulation. (C) [Br$_2$-D-Trp 3]EM-1 stimulation. (D) [*p*-tolyl-D-Trp 3]EM-1 stimulation. (E) [Nap-D-Trp 3]EM-1 stimulation. (F) [3-trifluoromethylphenyl-D-Trp]EM-1 stimulation.

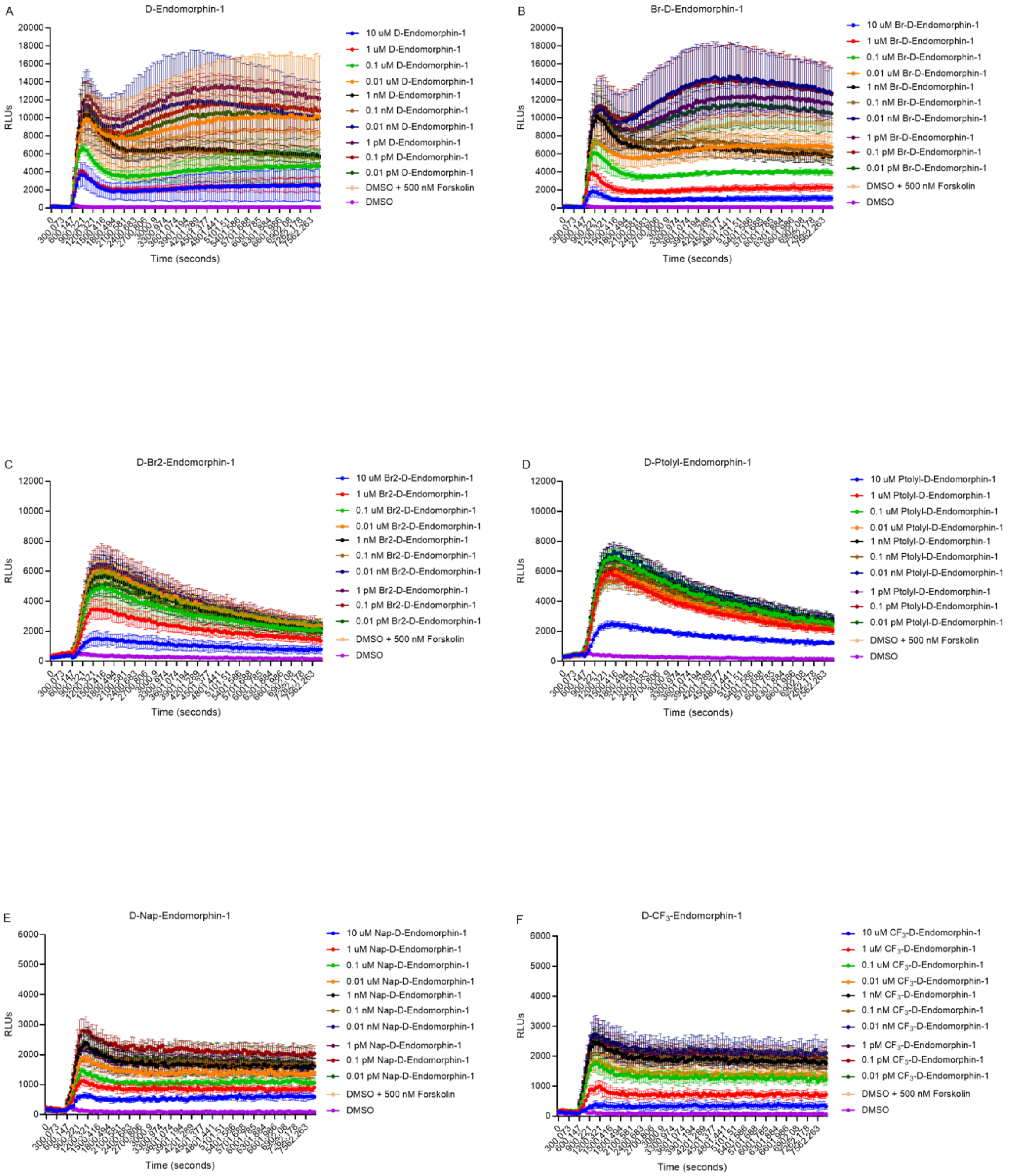

**Supplementary Figure 41. SDS-PAGE gel of Ni-NTA purified RebH and 4V.** PageRuler Plus Prestained 10-250kDa protein ladder (5 µL) was loaded onto the gel. Ni-NTA purified RebH or 4V (20 µg) was mixed with loading dye and loaded onto the gel. Gel was run at 140 volts for 80 minutes.

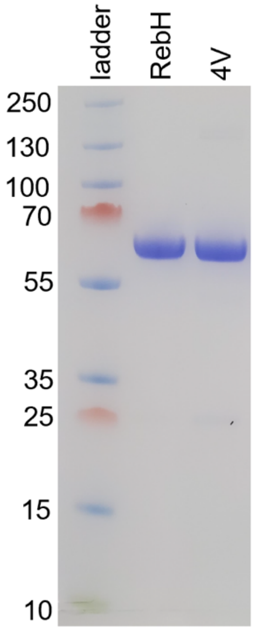
