## Supplementary Appendix for "Enzymatic bromination of native peptides for late-stage structural diversification via Suzuki-Miyaura coupling"

**Affiliation**

#### RebH Brominated Glycine Peptides of Increasing Length Mass Spectra

#### RebH GGW-$CONH_2$

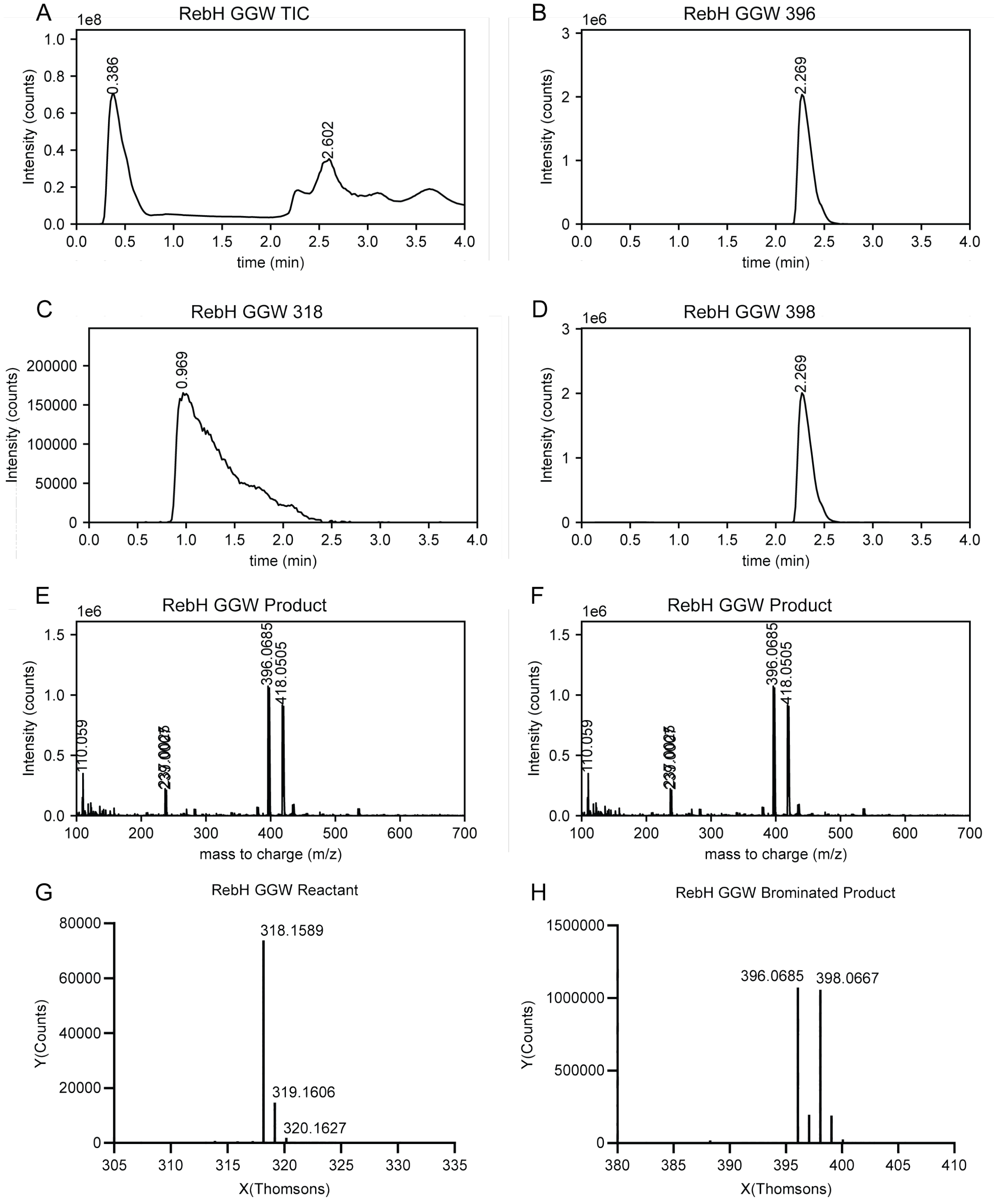

**RebH GGGW-CONH$_2$**

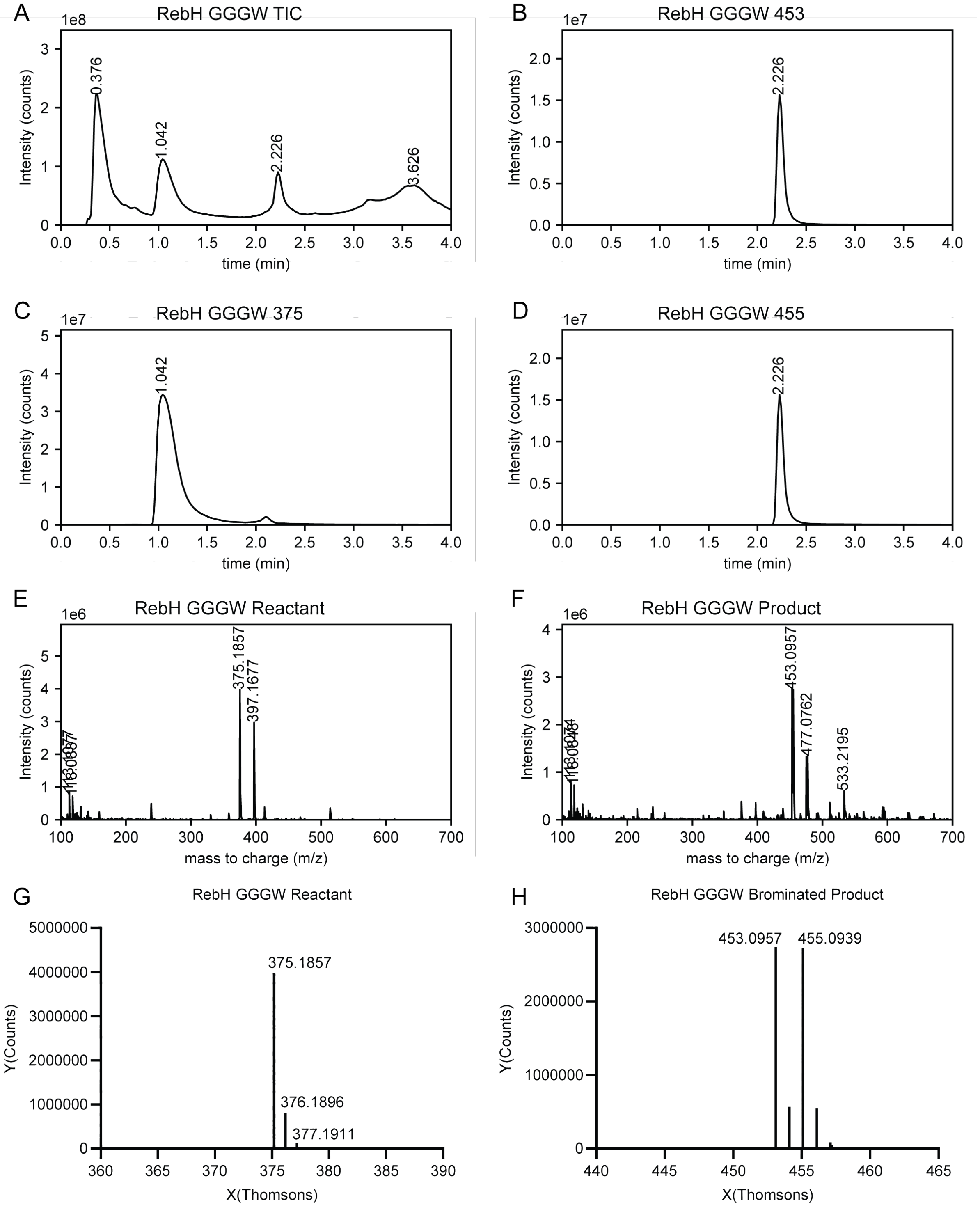

#### RebH GGGGW-CONH$_2$

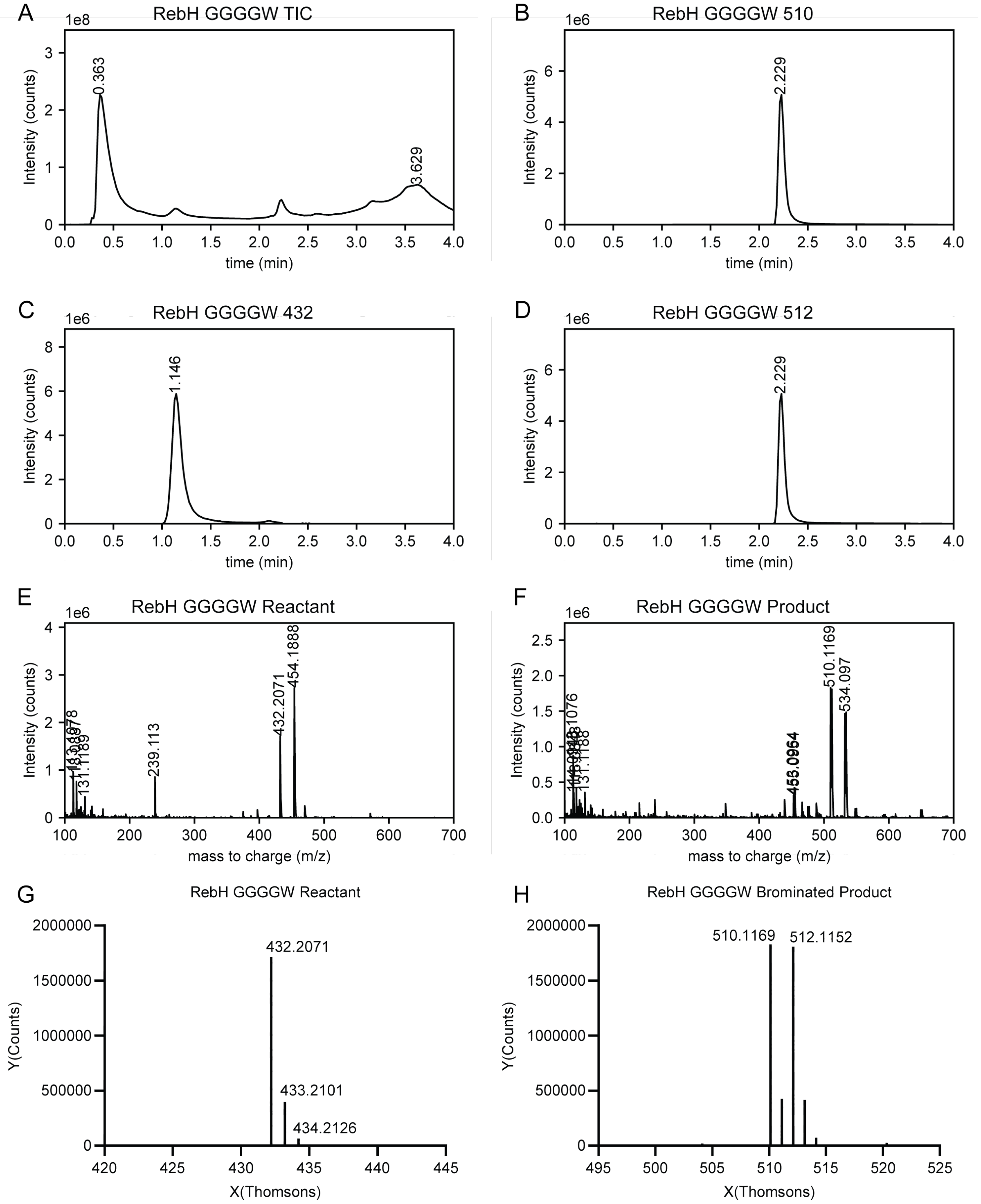

#### RebH GGGGGW-CONH$_2$

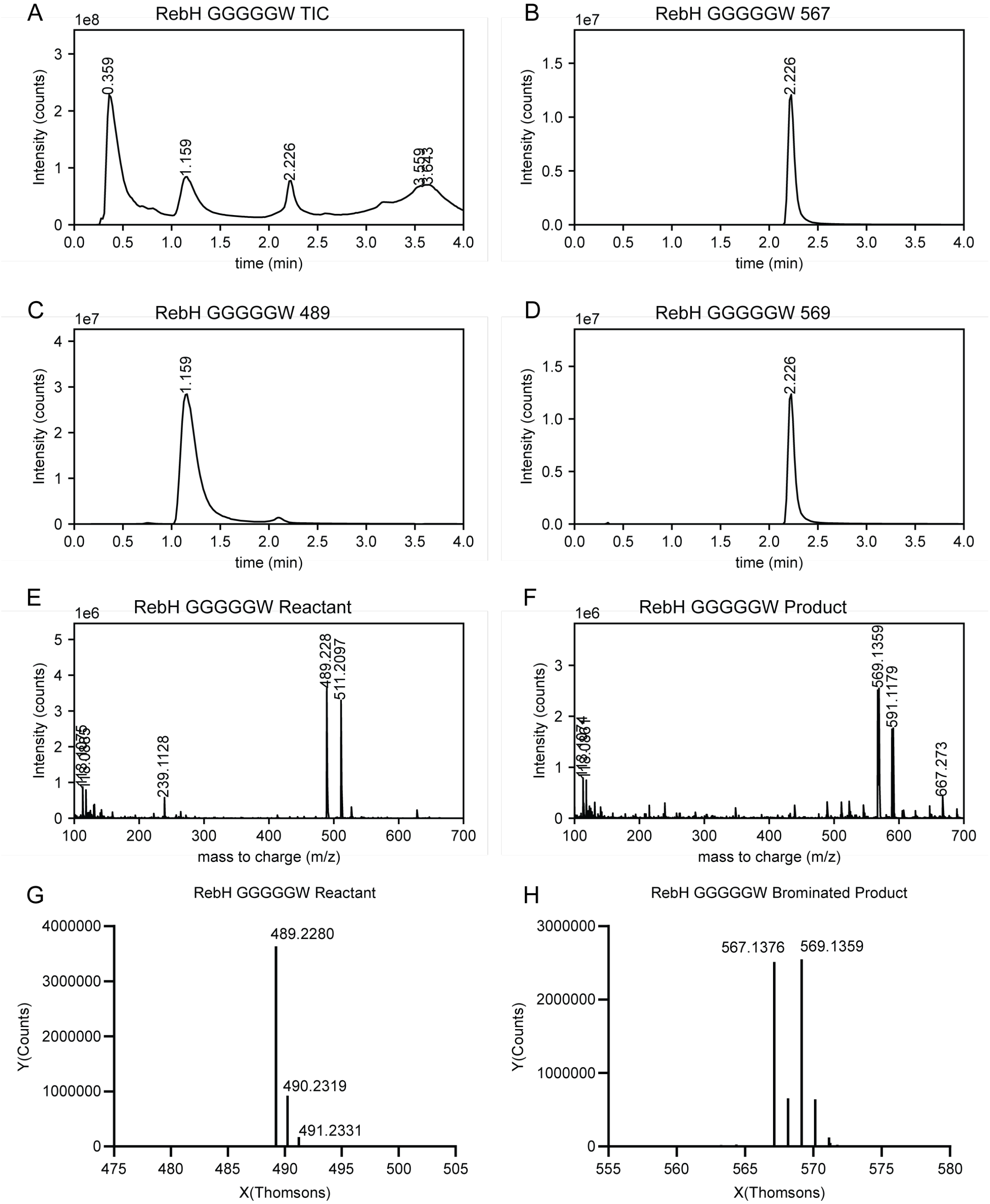

#### RebH WGG-CONH$_2$

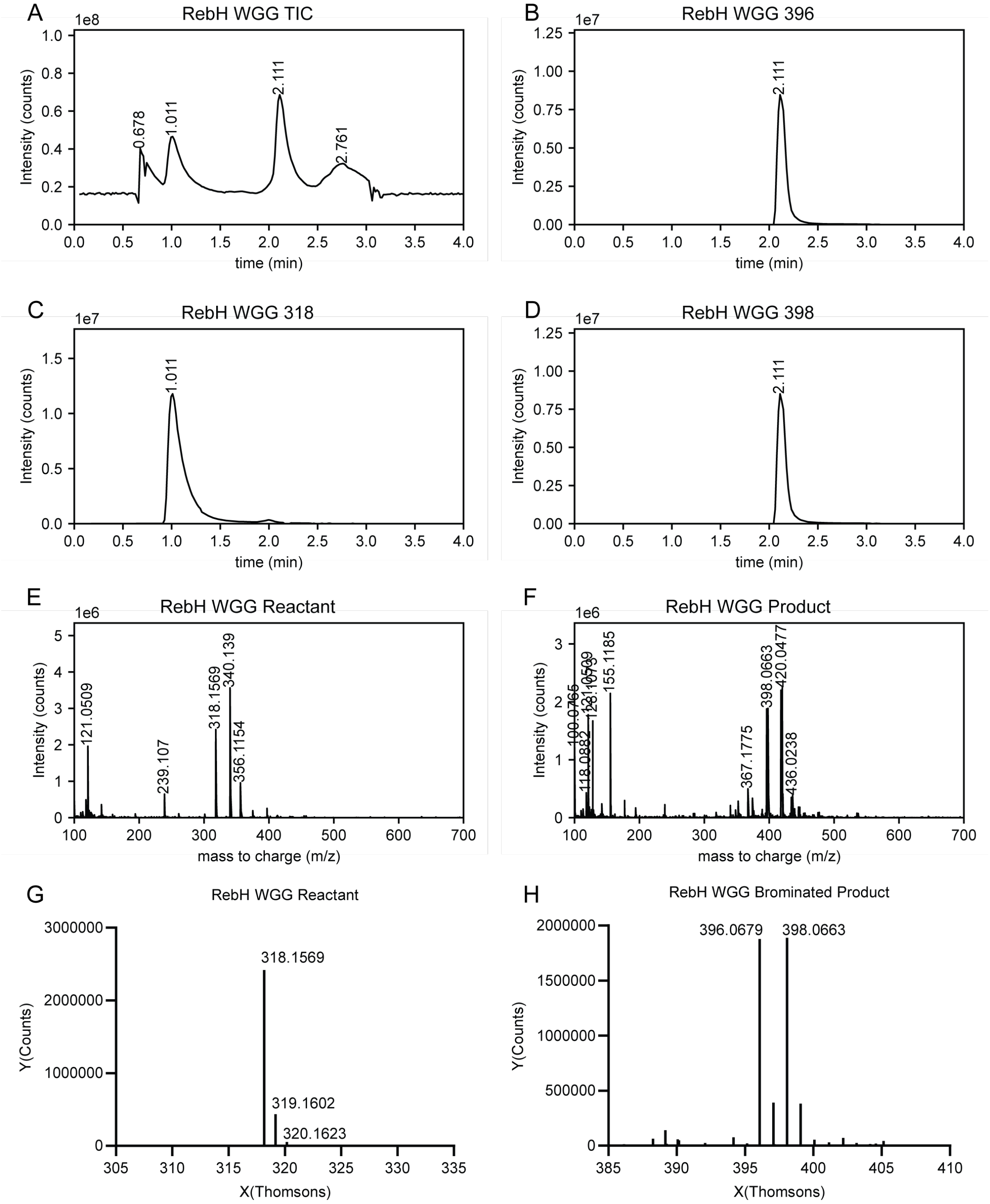

#### RebH WGGG-CONH$_2$

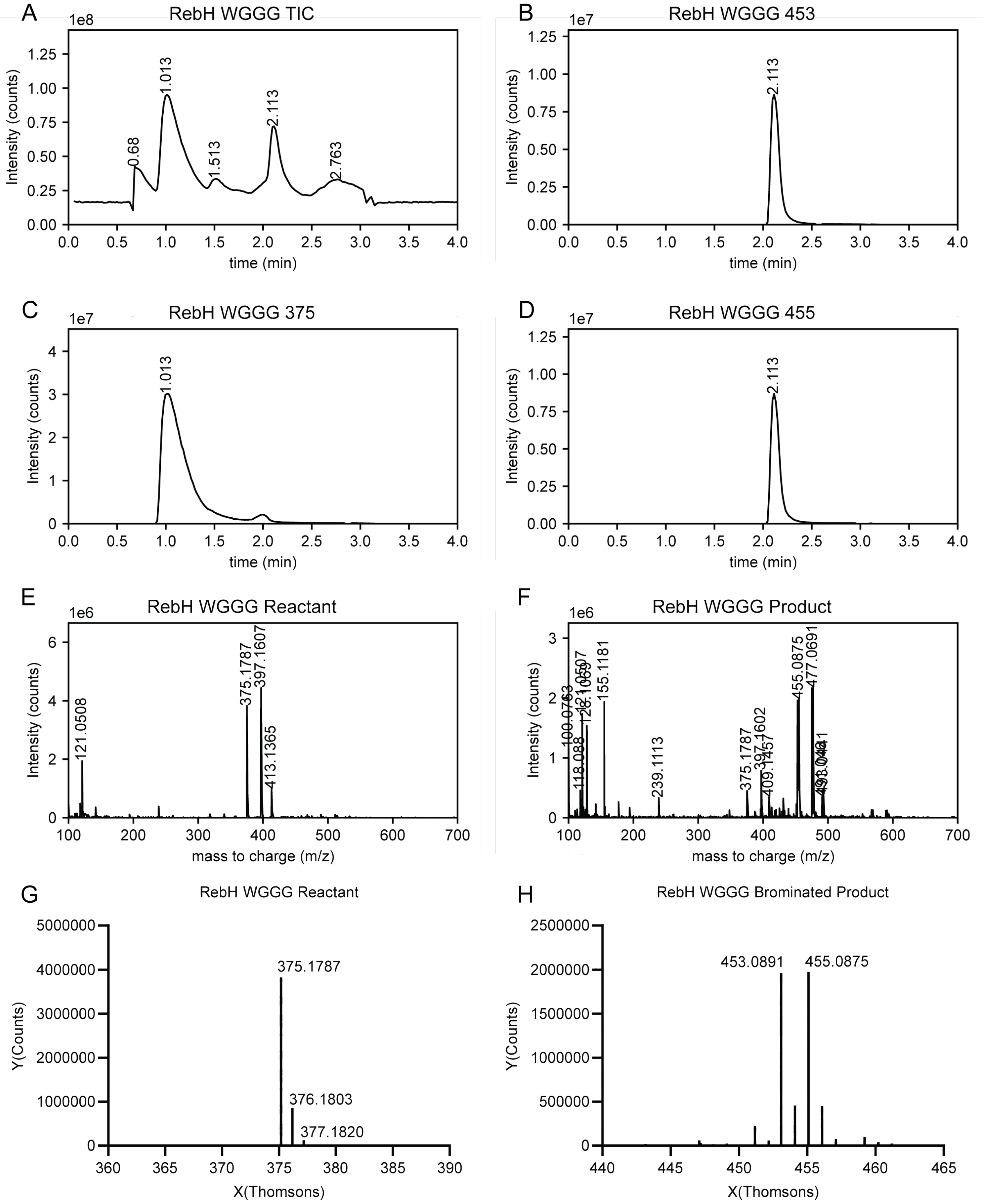

#### RebH WGGGG-CONH$_2$

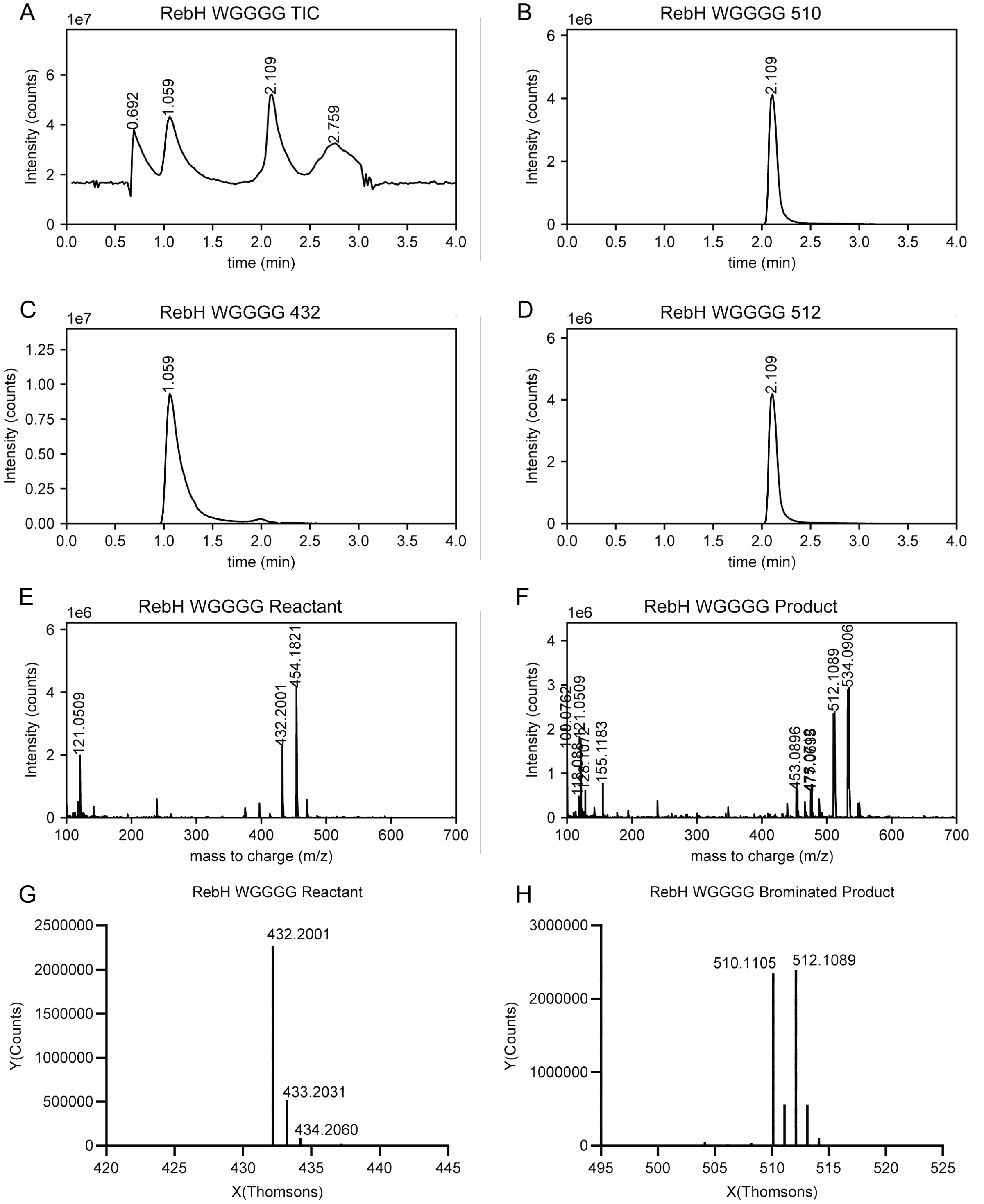

#### RebH WGGGGG-$CONH_2$

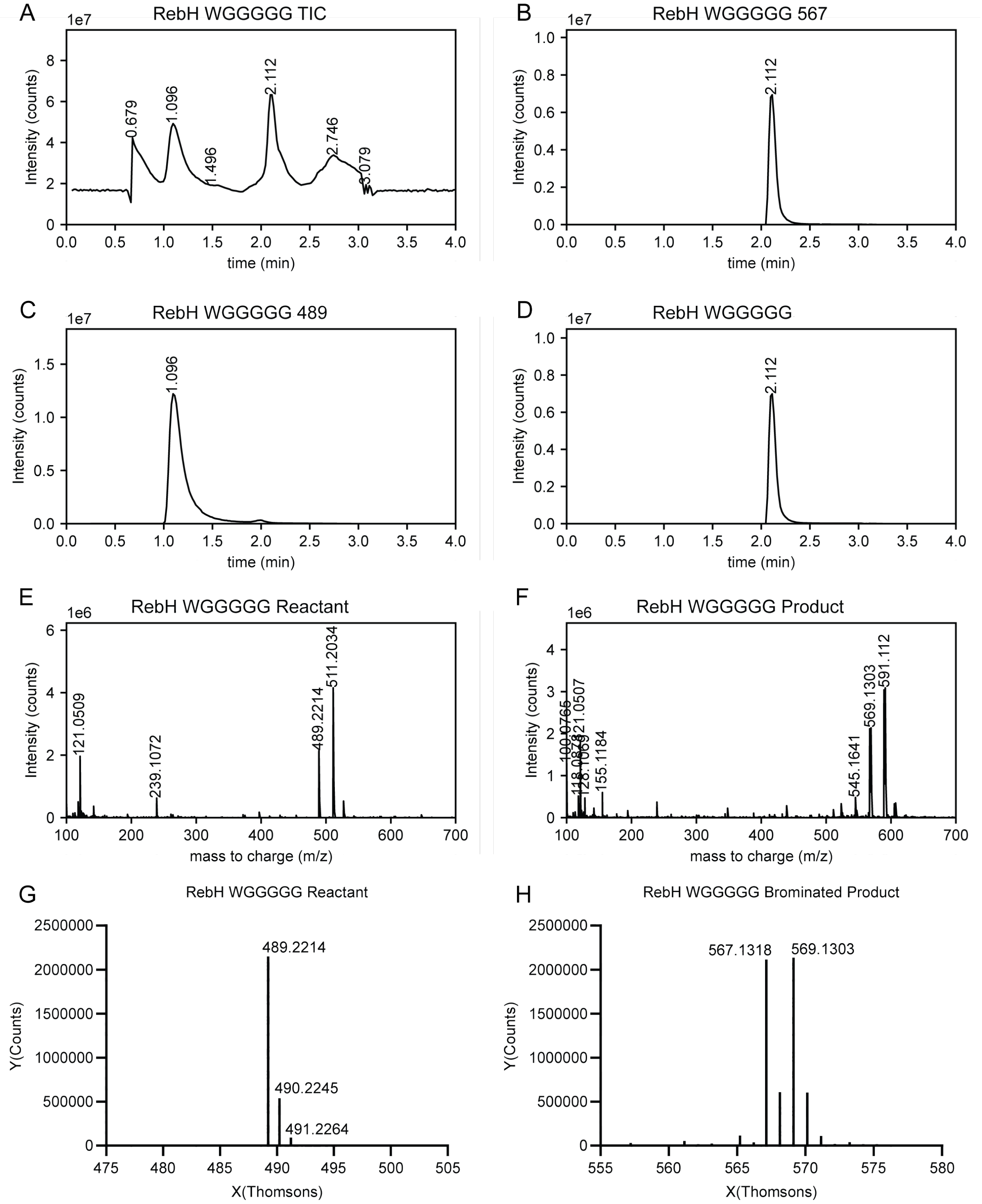

**RebH GWG-CONH$_2$**

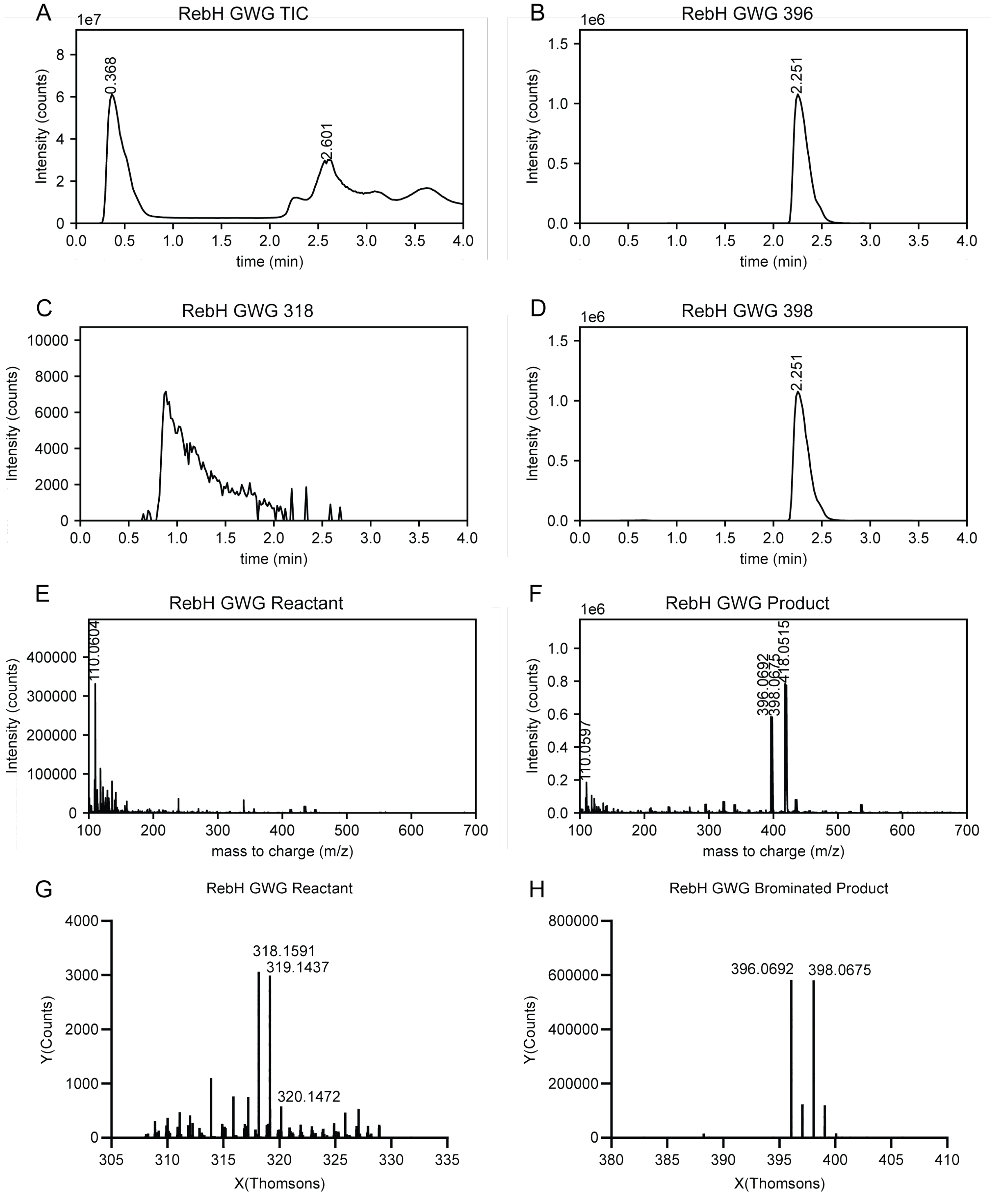

#### RebH GGWGG-CONH$_2$

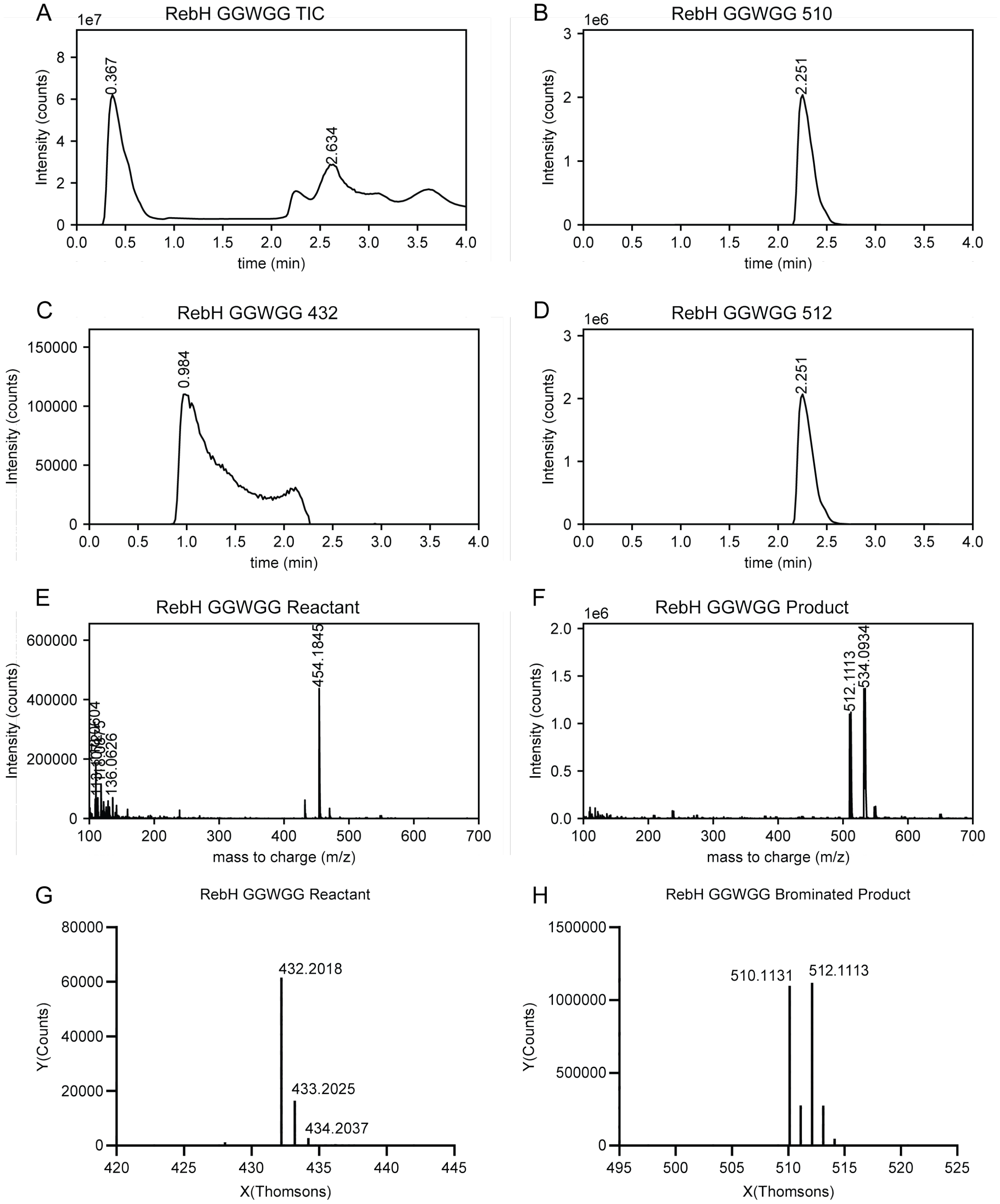

### RebH GW-COOH

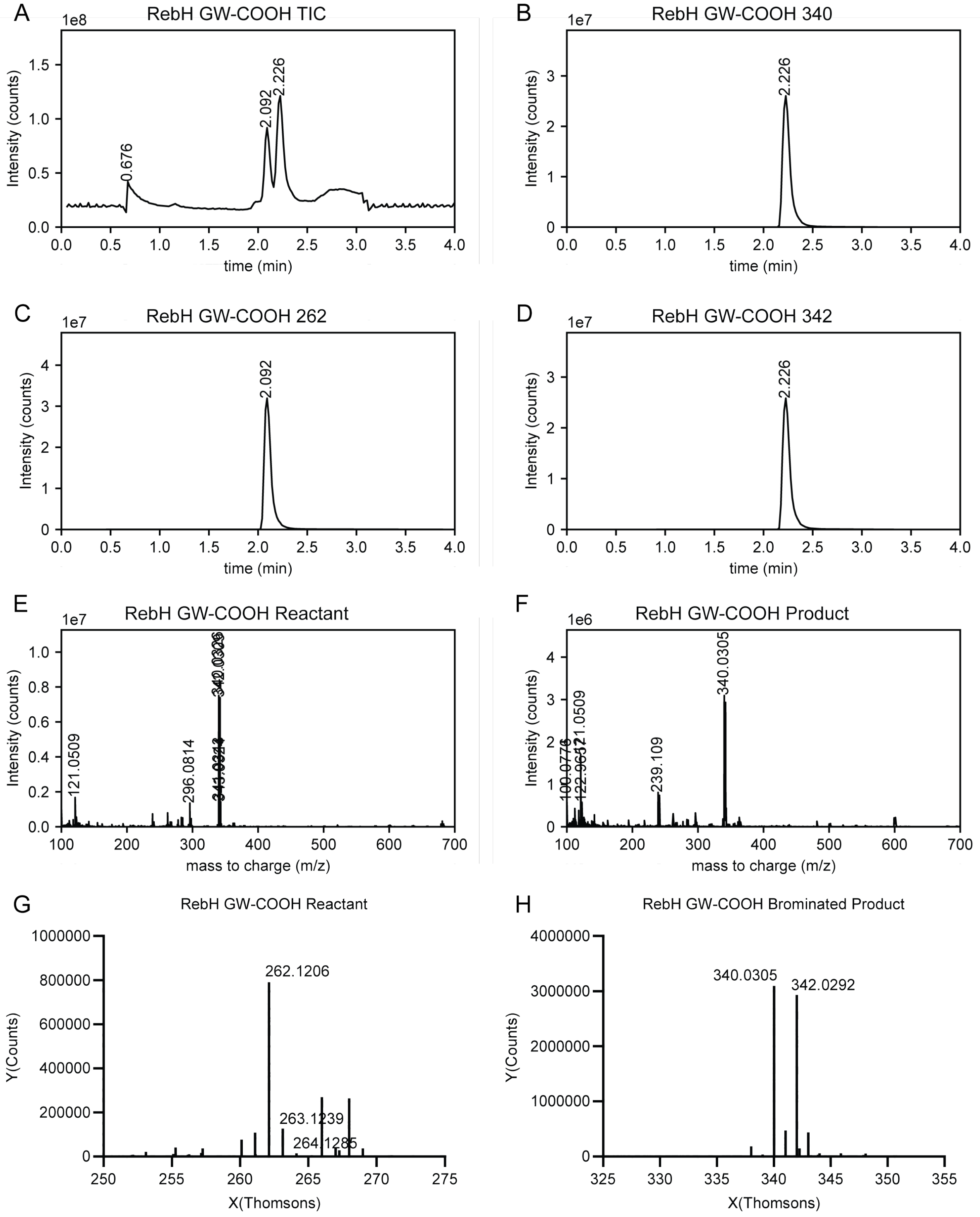
A
RebH GW-COOH TIC
1e8
Intensity (counts)
time (min)
0.676
2.092
2.226
B
RebH GW-COOH 340
1e7
Intensity (counts)
time (min)
2.226
C
RebH GW-COOH 262
1e7
Intensity (counts)
time (min)
2.092
D
RebH GW-COOH 342
1e7
Intensity (counts)
time (min)
2.226
E
RebH GW-COOH Reactant
1e7
Intensity (counts)
mass to charge (m/z)
121.0509
296.0814
F
RebH GW-COOH Product
1e6
Intensity (counts)
mass to charge (m/z)
340.0305
121.0509
239.109
G
RebH GW-COOH Reactant
Y(Counts)
X(Thomsons)
262.1206
263.1239
264.1285
H
RebH GW-COOH Brominated Product
Y(Counts)
X(Thomsons)
340.0305
342.0292

**RebH GGW-COOH**

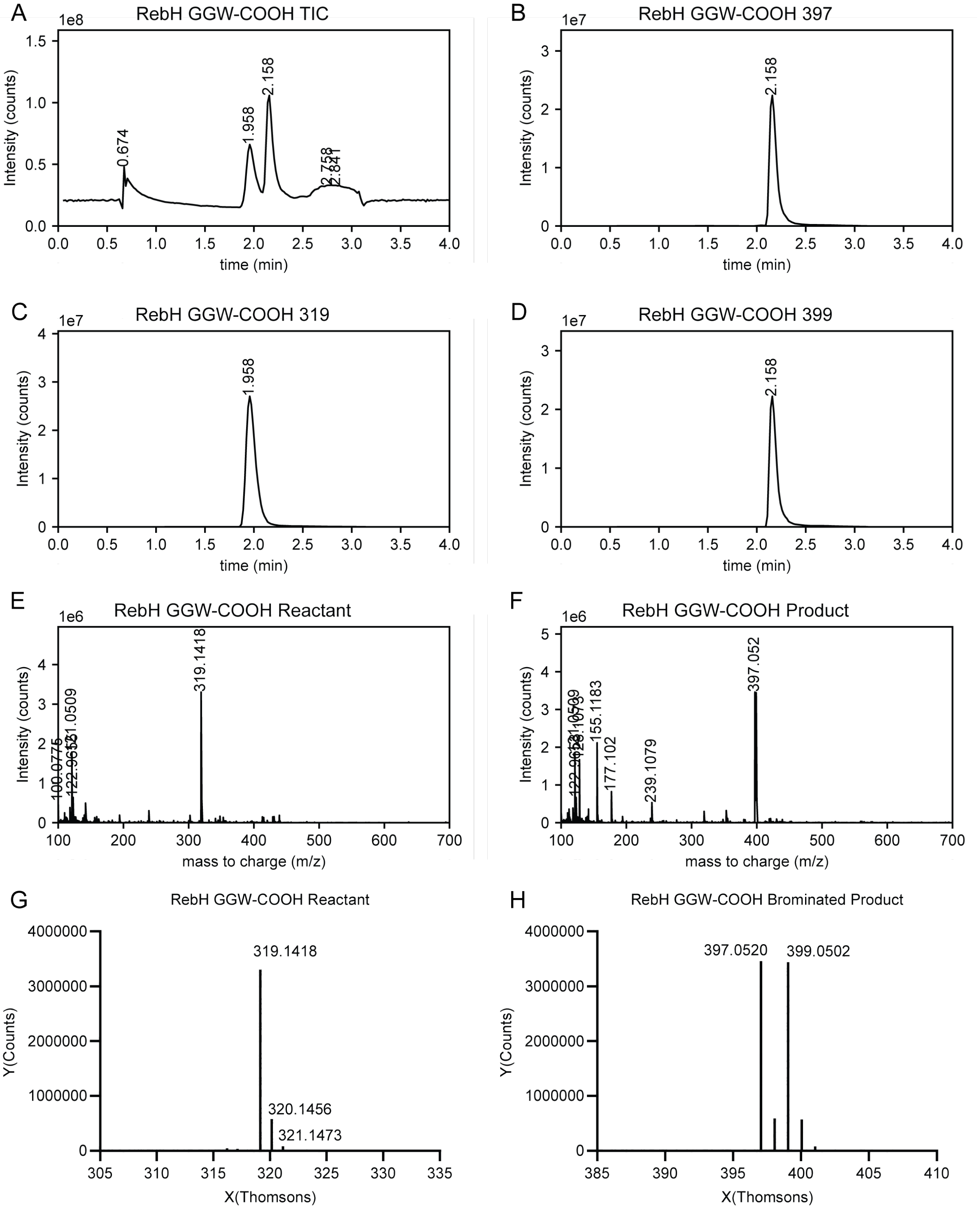

#### RebH WG-COOH

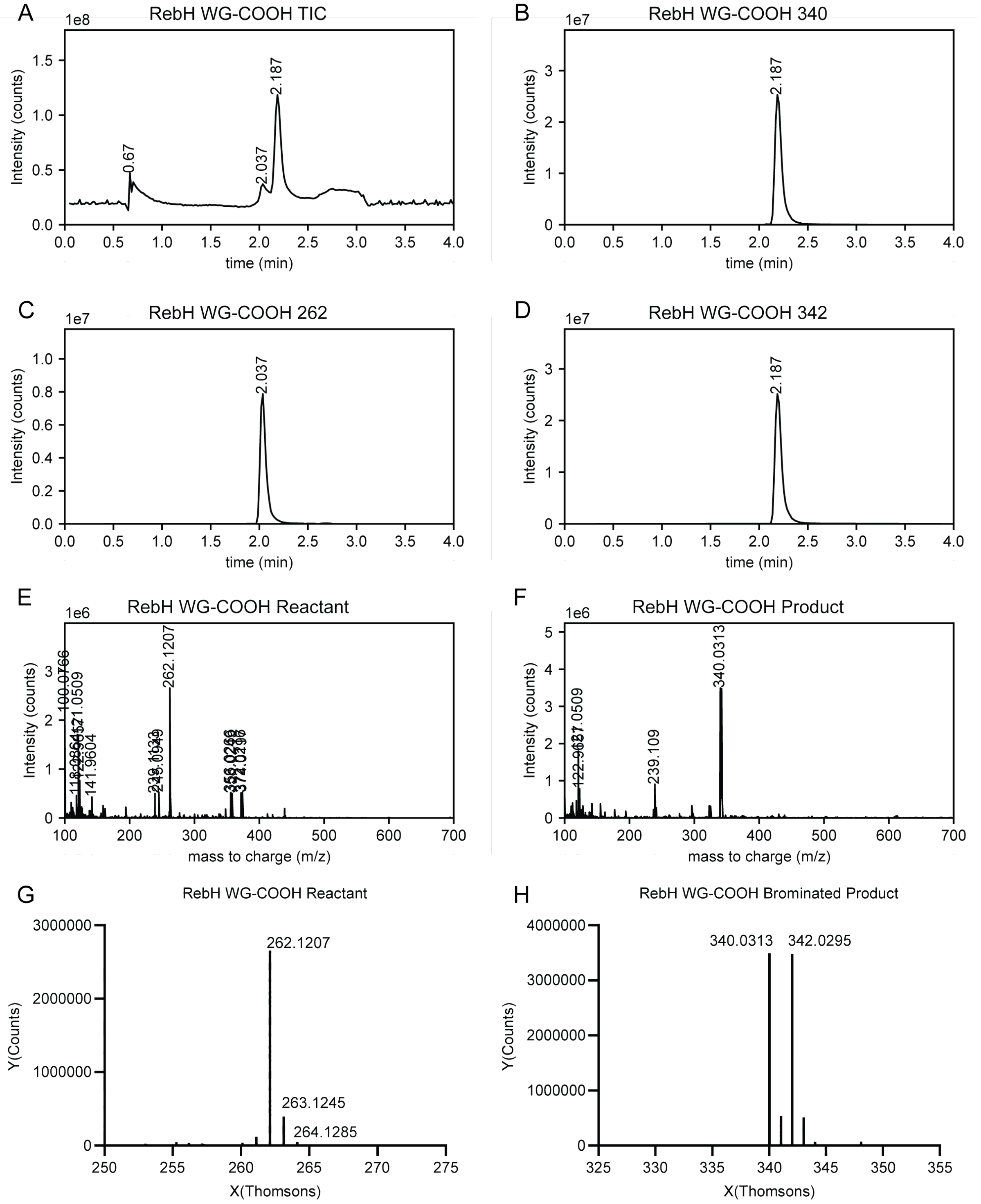
A
RebH WG-COOH TIC
1e8
Intensity (counts)
time (min)
0.67
2.037
2.187
B
RebH WG-COOH 340
1e7
Intensity (counts)
time (min)
2.187
C
RebH WG-COOH 262
1e7
Intensity (counts)
time (min)
2.037
D
RebH WG-COOH 342
1e7
Intensity (counts)
time (min)
2.187
E
RebH WG-COOH Reactant
1e6
Intensity (counts)
mass to charge (m/z)
100.0766
121.0509
141.9604
262.1207
F
RebH WG-COOH Product
1e6
Intensity (counts)
mass to charge (m/z)
121.0509
239.109
340.0313
G
RebH WG-COOH Reactant
Y(Counts)
X(Thomsons)
262.1207
263.1245
264.1285
H
RebH WG-COOH Brominated Product
Y(Counts)
X(Thomsons)
340.0313
342.0295

#### RebH WGG-COOH

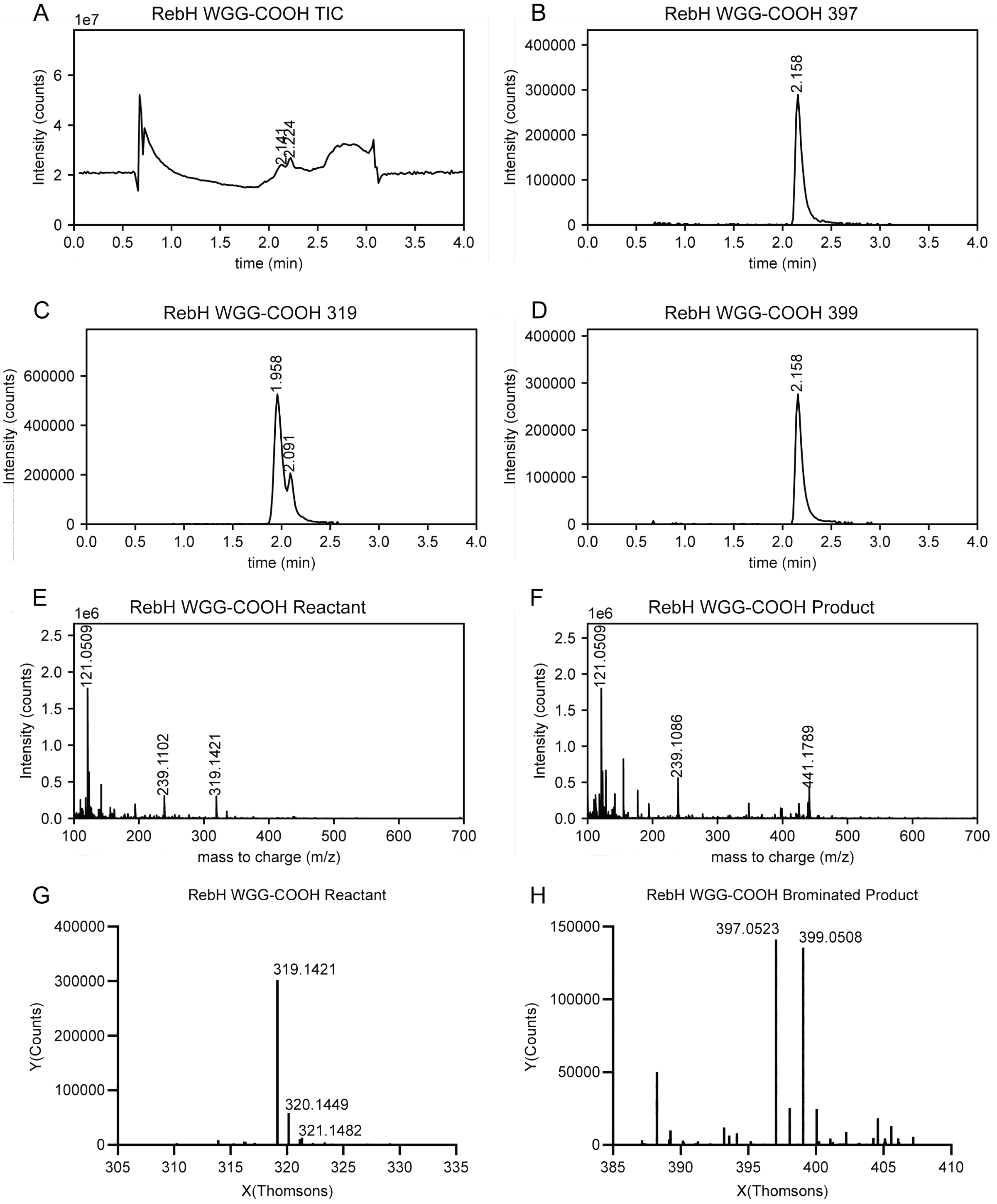

### 4V Brominated Glycine Peptides of Increasing Length Mass Spectra

**4V GGW-$CONH_2$**

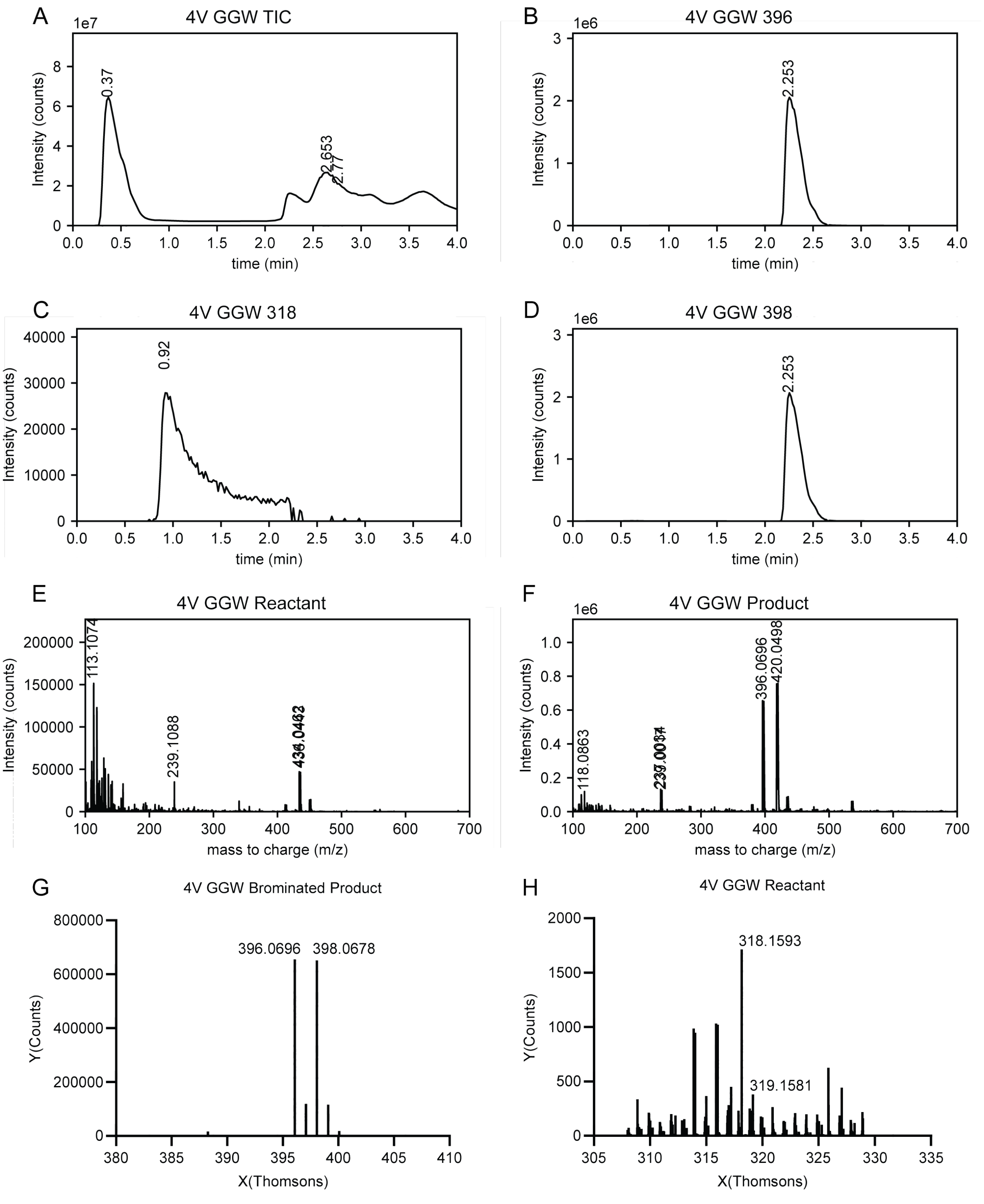
A
4V GGW TIC
1e7
Intensity (counts)
time (min)
0.37
2.653
2.77
B
4V GGW 396
1e6
2.253
C
4V GGW 318
0.92
D
4V GGW 398
2.253
E
4V GGW Reactant
113.1074
239.1088
mass to charge (m/z)
F
4V GGW Product
396.0696
420.0498
118.0863
G
4V GGW Brominated Product
396.0696
398.0678
Y(Counts)
X(Thomsons)
H
4V GGW Reactant
318.1593
319.1581

#### 4V GGGW-CONH$_2$

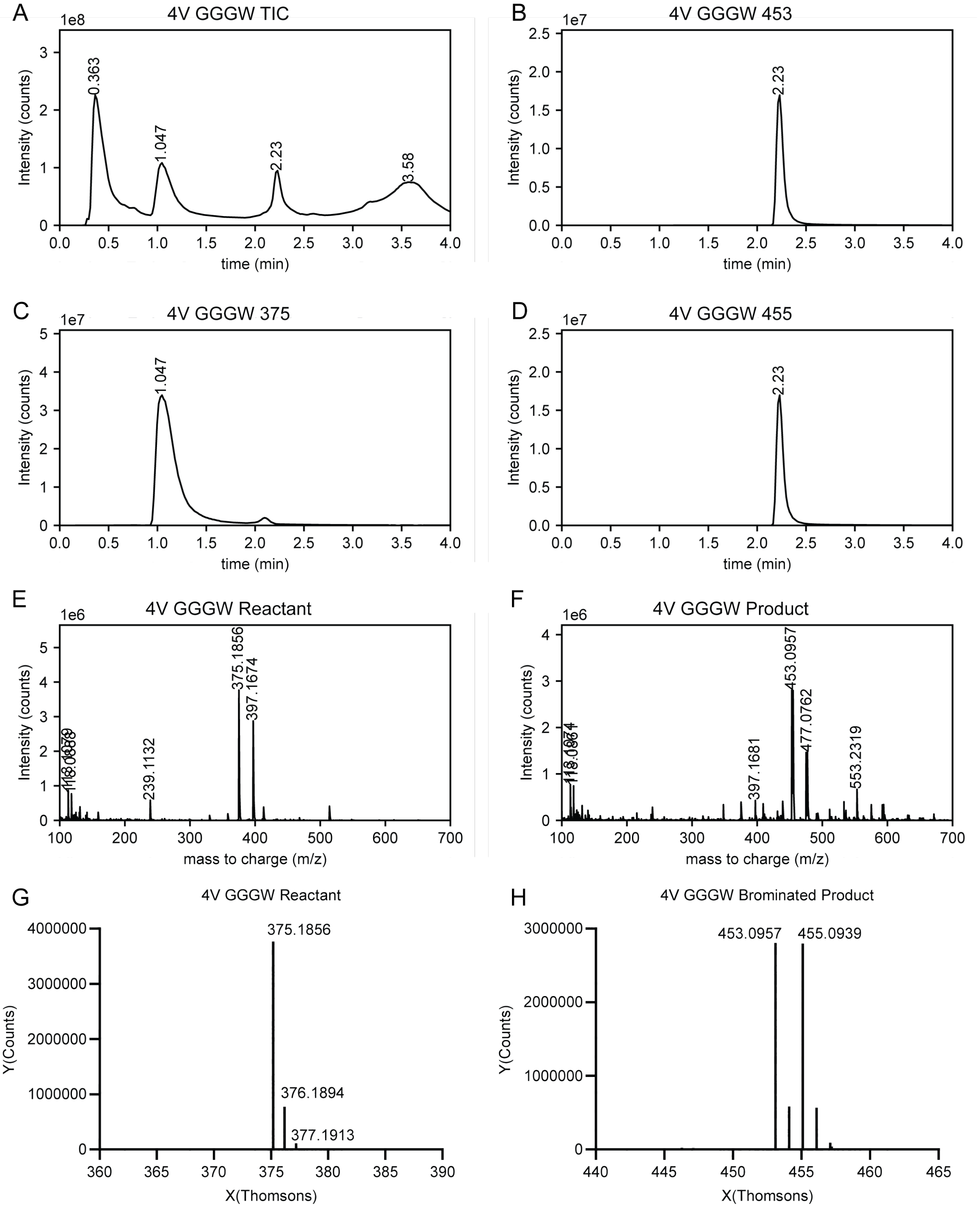

**4V GGGGW-CONH$_2$**

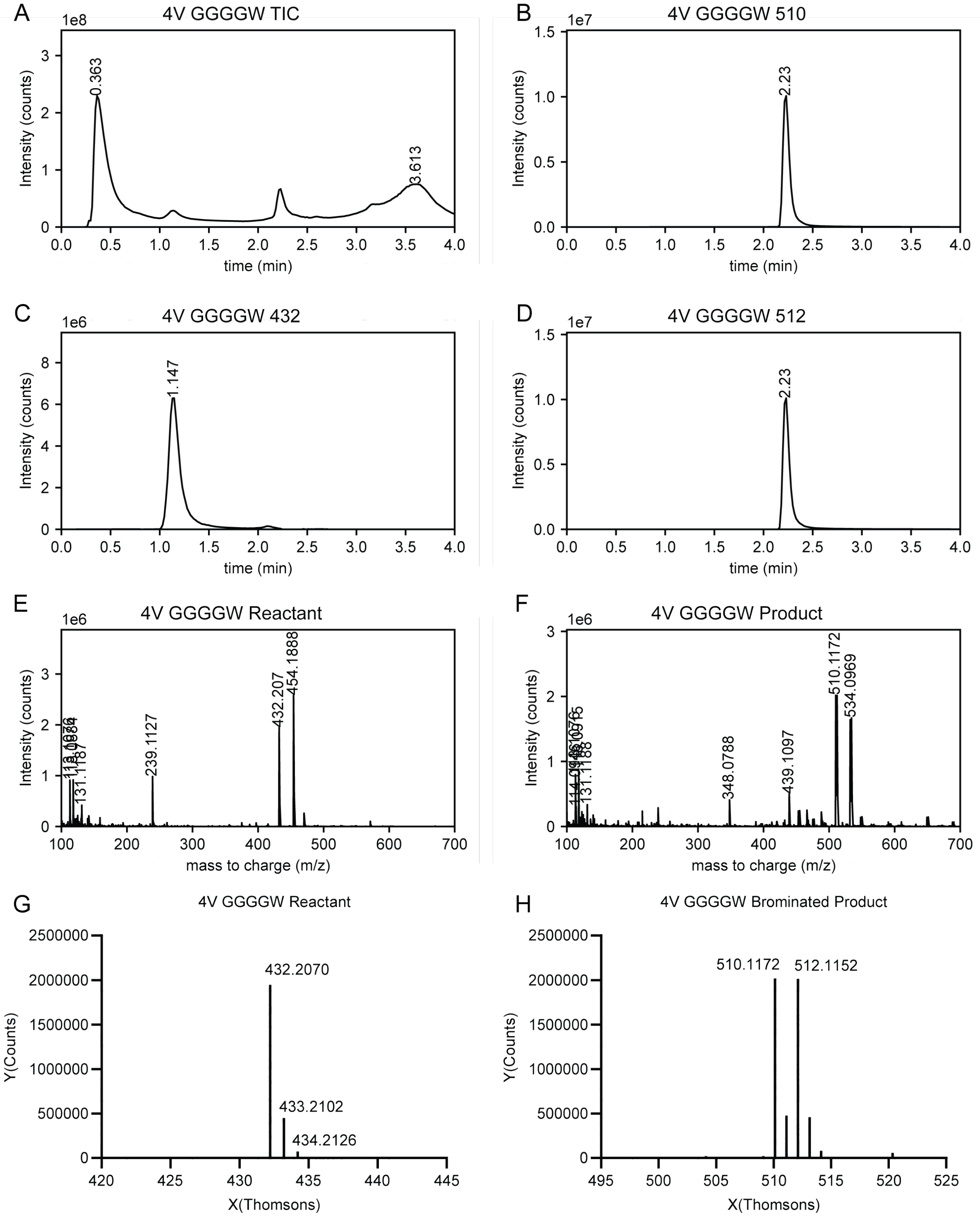

#### 4V GGGGGW-$CONH_2$

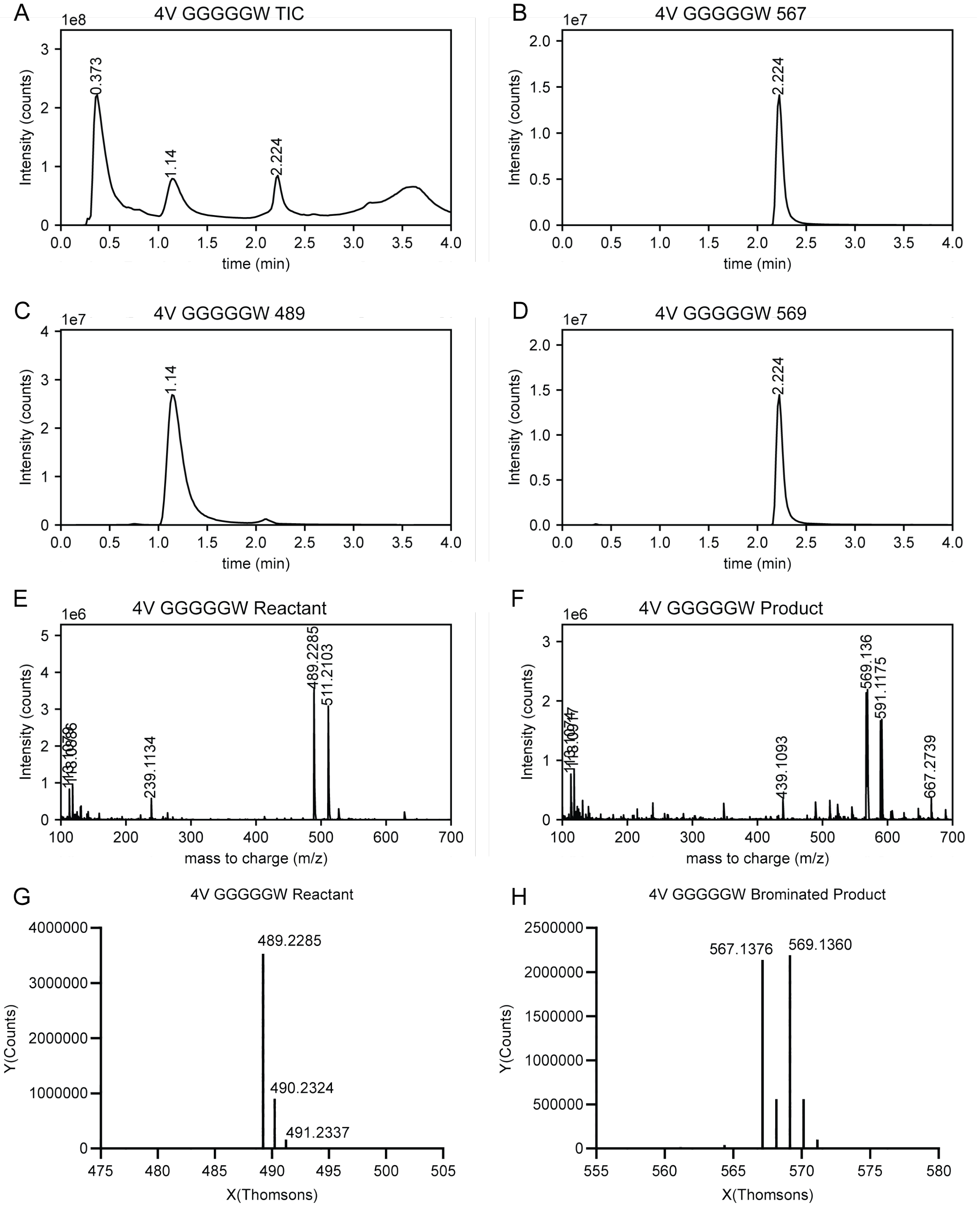

#### 4V WGG-$CONH_2$

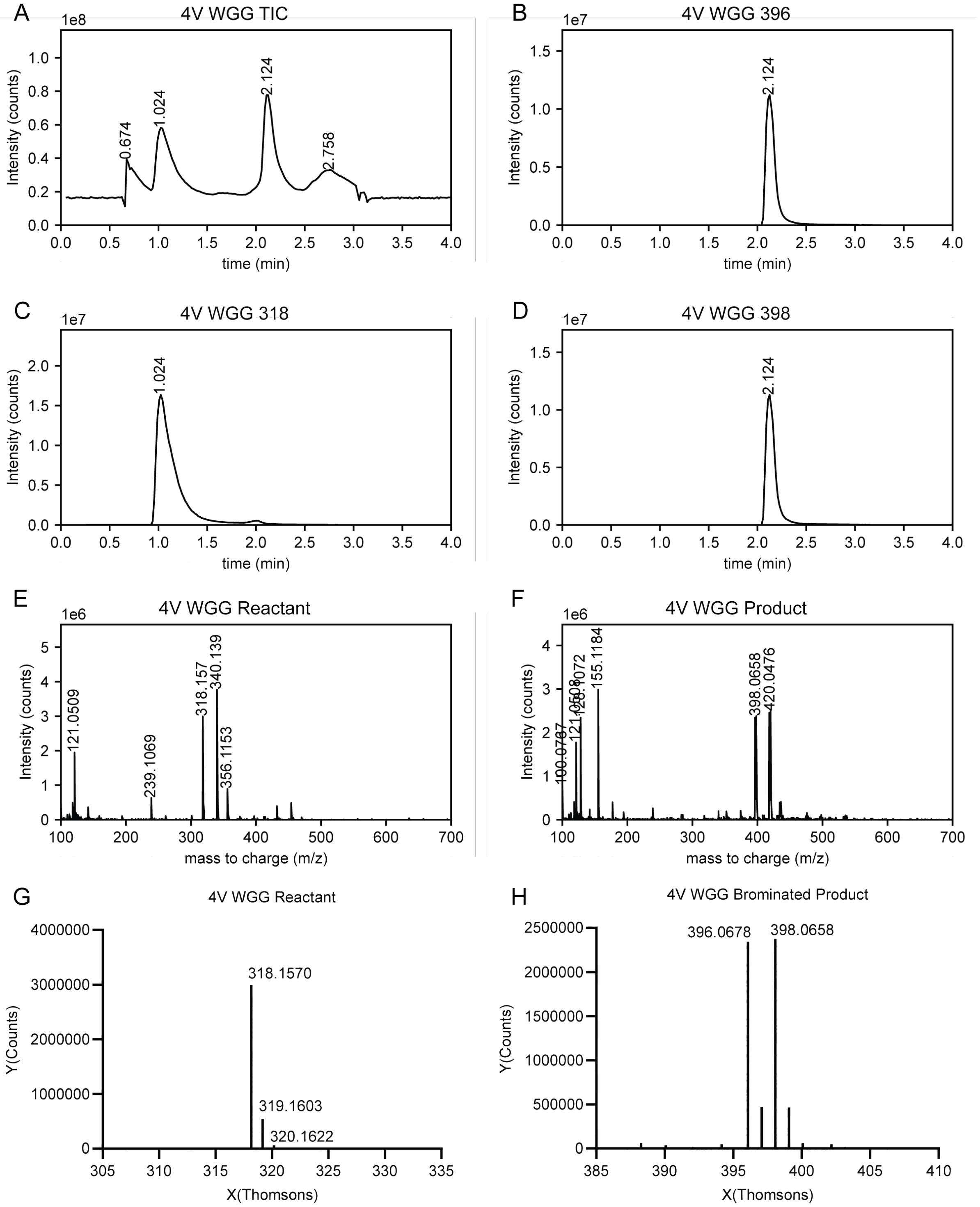
A
4V WGG TIC
1e8
Intensity (counts)
time (min)
0.674
1.024
2.124
2.758
B
4V WGG 396
1e7
2.124
C
4V WGG 318
1.024
D
4V WGG 398
2.124
E
4V WGG Reactant
1e6
mass to charge (m/z)
121.0509
239.1069
318.157
340.139
356.1153
F
4V WGG Product
155.1184
398.0658
420.0476
G
4V WGG Reactant
Y(Counts)
X(Thomsons)
318.1570
319.1603
320.1622
H
4V WGG Brominated Product
396.0678
398.0658

#### 4V WGGG-CONH$_2$

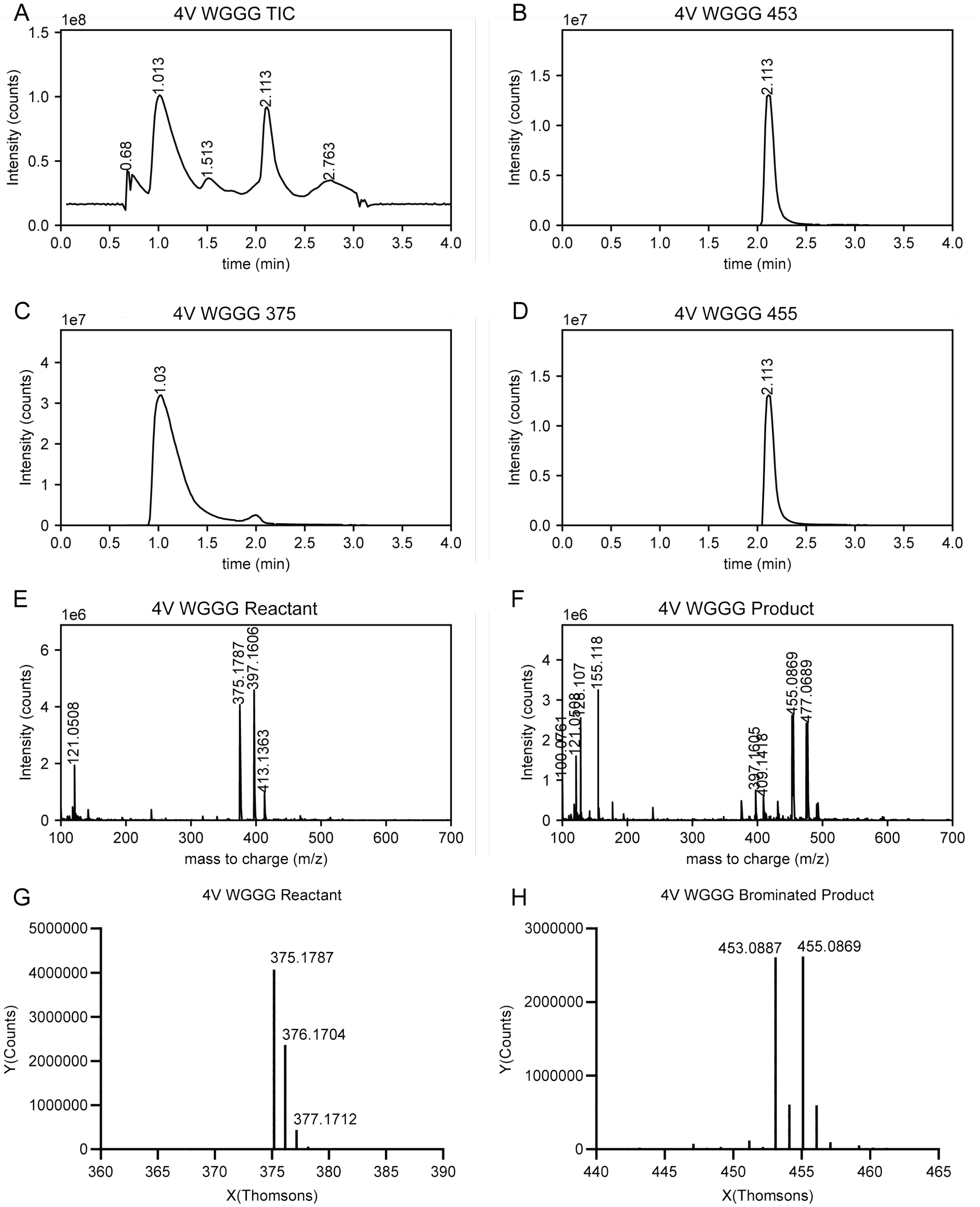

#### 4V WGGGG-CONH$_2$

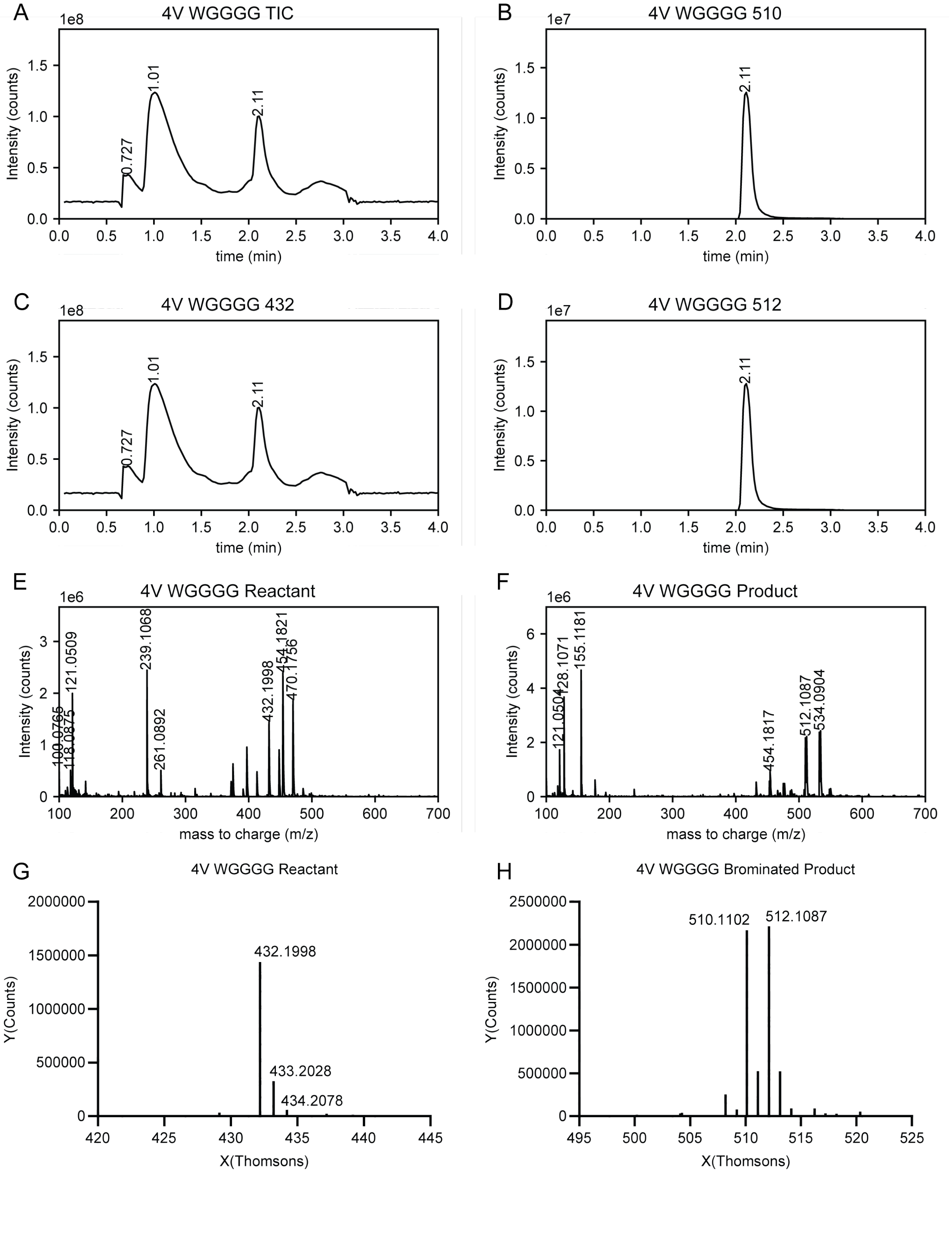

#### 4V WGGGGG-CONH$_2$

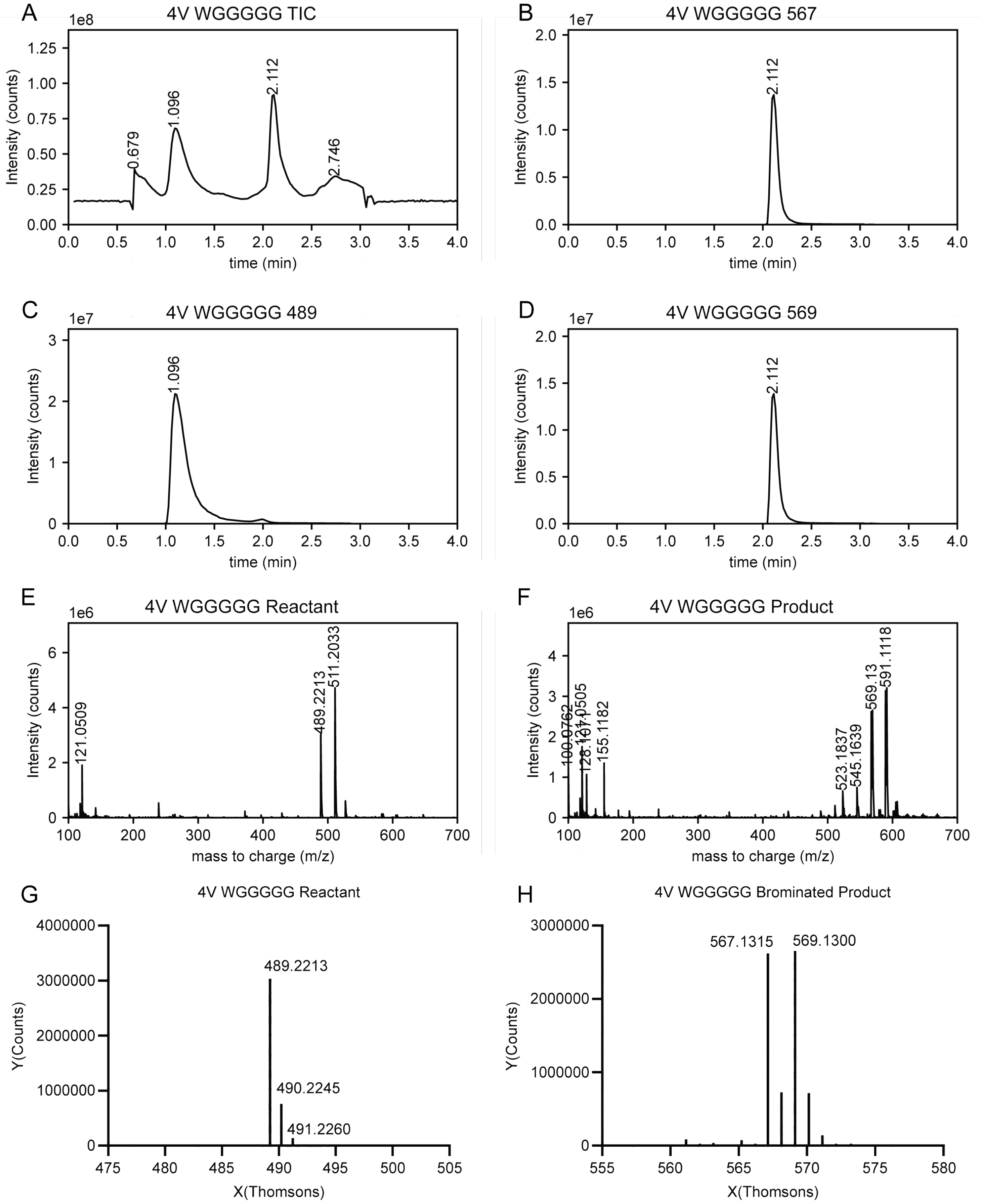

**4V GWG-CONH$_2$**

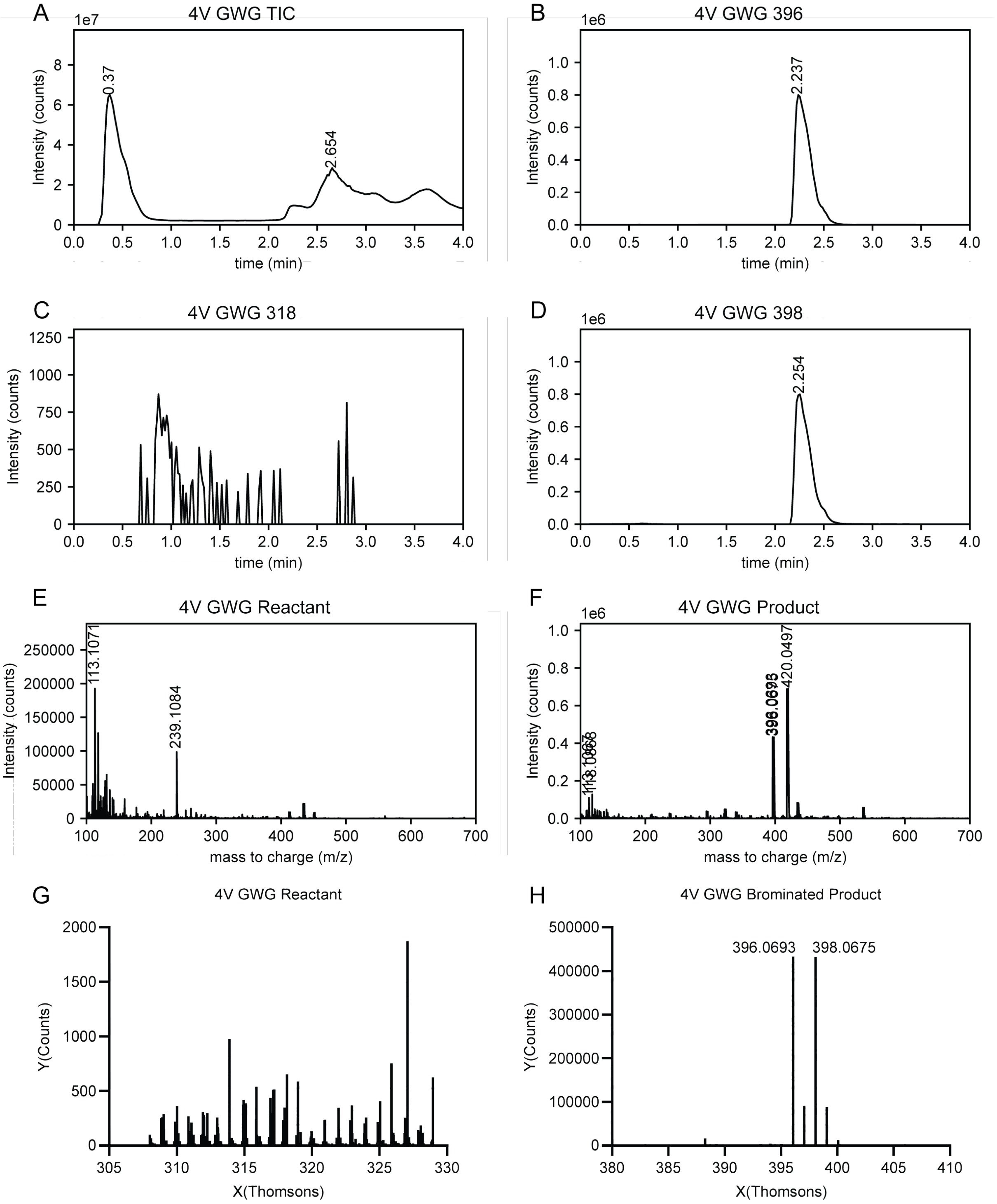
A
4V GWG TIC
1e7
Intensity (counts)
time (min)
0.37
2.654
B
4V GWG 396
1e6
2.237
C
4V GWG 318
D
4V GWG 398
2.254
E
4V GWG Reactant
113.1071
239.1084
mass to charge (m/z)
F
4V GWG Product
420.0497
G
4V GWG Reactant
Y(Counts)
X(Thomsons)
H
4V GWG Brominated Product
396.0693
398.0675

#### 4V GGWGG-$CONH_2$

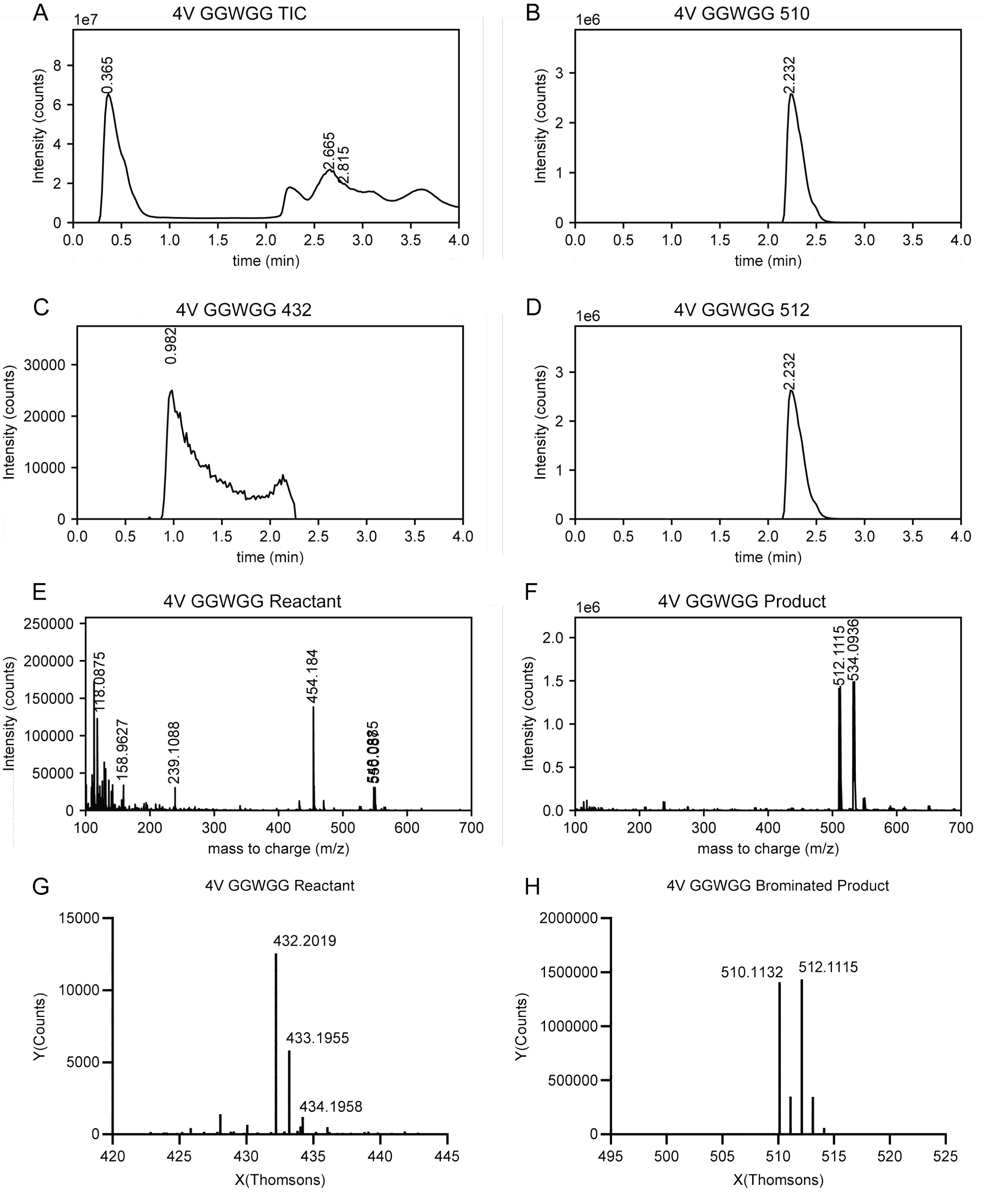

#### 4V GW-COOH

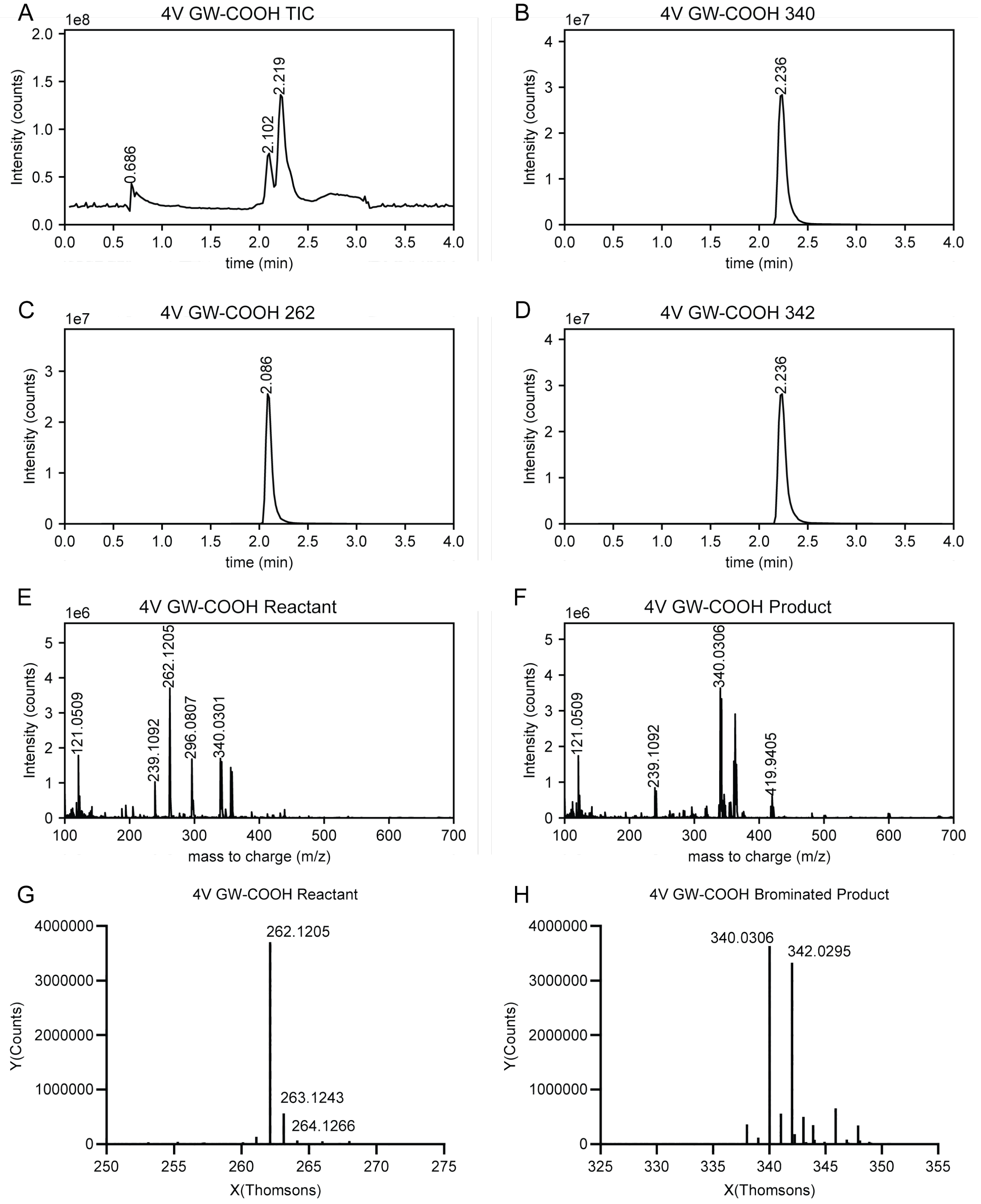
A
4V GW-COOH TIC
1e8
Intensity (counts)
time (min)
0.686
2.102
2.219
B
4V GW-COOH 340
1e7
Intensity (counts)
time (min)
2.236
C
4V GW-COOH 262
1e7
Intensity (counts)
time (min)
2.086
D
4V GW-COOH 342
1e7
Intensity (counts)
time (min)
2.236
E
4V GW-COOH Reactant
1e6
Intensity (counts)
mass to charge (m/z)
121.0509
239.1092
262.1205
296.0807
340.0301
F
4V GW-COOH Product
1e6
Intensity (counts)
mass to charge (m/z)
121.0509
239.1092
340.0306
419.9405
G
4V GW-COOH Reactant
Y(Counts)
X(Thomsons)
262.1205
263.1243
264.1266
H
4V GW-COOH Brominated Product
Y(Counts)
X(Thomsons)
340.0306
342.0295

#### 4V GGW-COOH

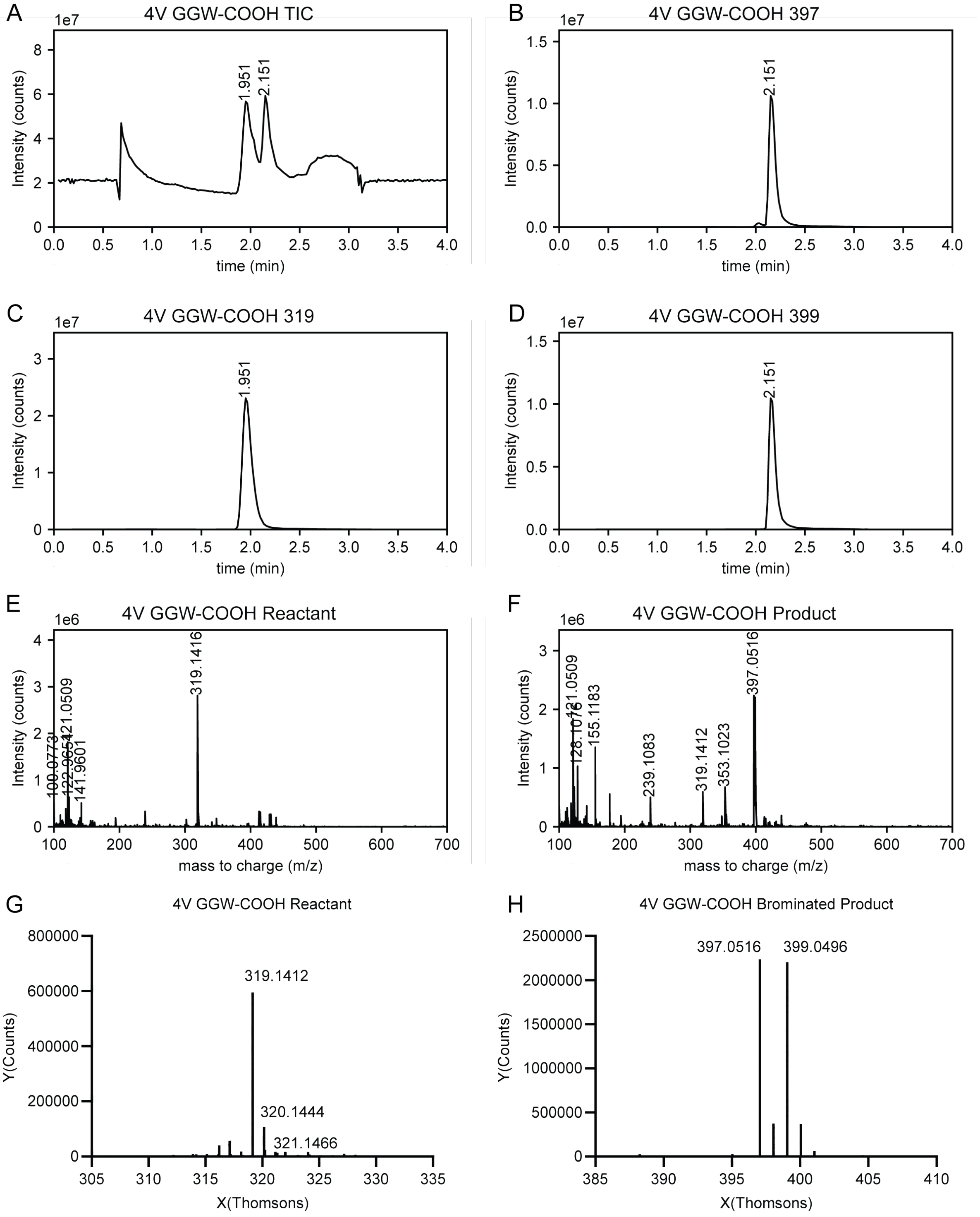
A
4V GGW-COOH TIC
B
4V GGW-COOH 397
C
4V GGW-COOH 319
D
4V GGW-COOH 399
E
4V GGW-COOH Reactant
F
4V GGW-COOH Product
G
4V GGW-COOH Reactant
H
4V GGW-COOH Brominated Product
1e7
1e6
Intensity (counts)
time (min)
mass to charge (m/z)
Y(Counts)
X(Thomsons)
1.951
2.151
319.1416
121.0509
141.9601
397.0516
155.1183
239.1083
319.1412
353.1023
320.1444
321.1466
399.0496

**4V WG-COOH**

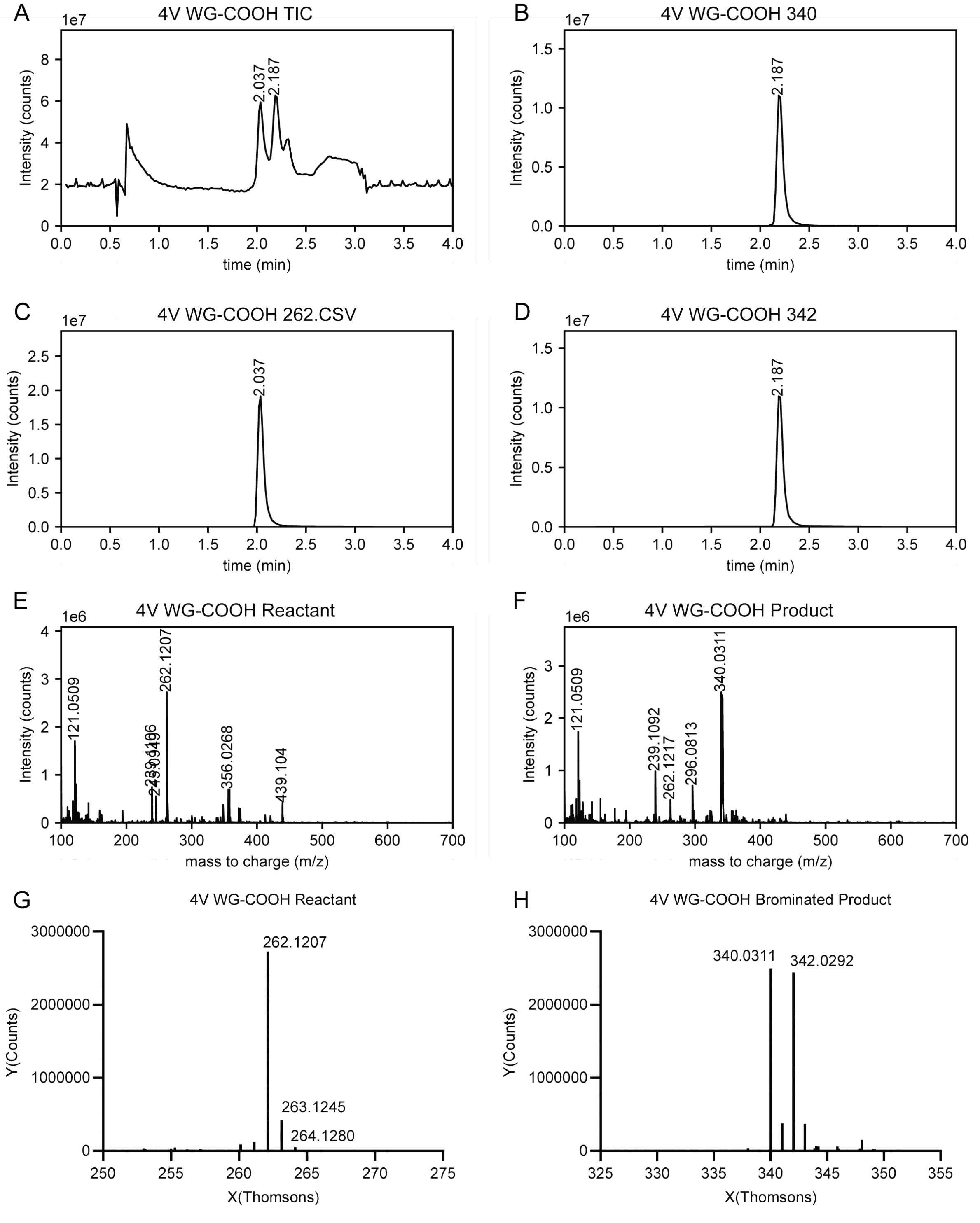
A
4V WG-COOH TIC
1e7
Intensity (counts)
time (min)
2.037
2.187
B
4V WG-COOH 340
1e7
Intensity (counts)
time (min)
2.187
C
4V WG-COOH 262.CSV
1e7
Intensity (counts)
time (min)
2.037
D
4V WG-COOH 342
1e7
Intensity (counts)
time (min)
2.187
E
4V WG-COOH Reactant
1e6
Intensity (counts)
mass to charge (m/z)
121.0509
262.1207
356.0268
439.104
F
4V WG-COOH Product
1e6
Intensity (counts)
mass to charge (m/z)
121.0509
239.1092
262.1217
296.0813
340.0311
G
4V WG-COOH Reactant
Y(Counts)
X(Thomsons)
262.1207
263.1245
264.1280
H
4V WG-COOH Brominated Product
Y(Counts)
X(Thomsons)
340.0311
342.0292

#### 4V WGG-COOH

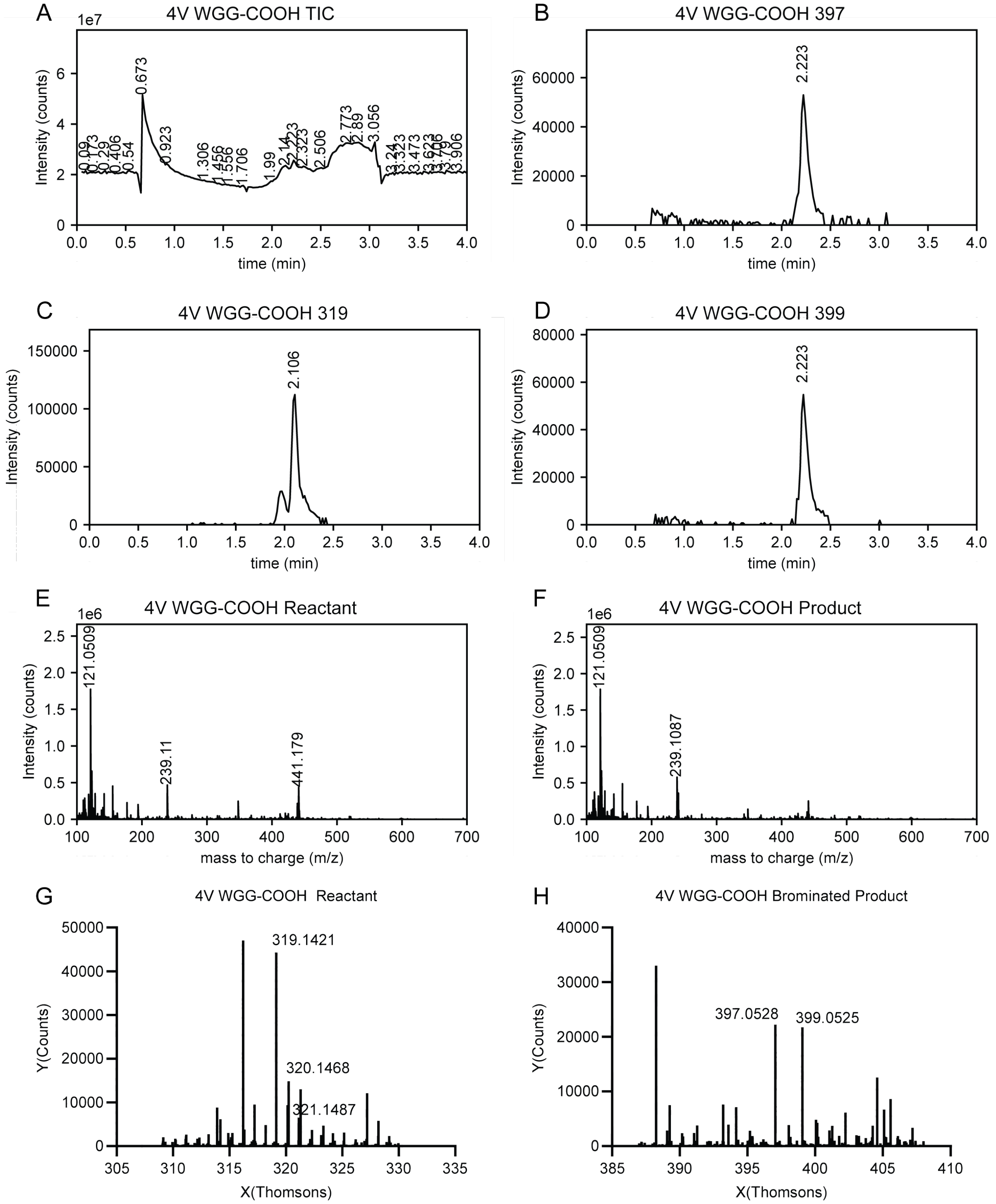
A
4V WGG-COOH TIC
1e7
Intensity (counts)
time (min)
0.673
0.923
1.306
1.99
2.506
3.056
B
4V WGG-COOH 397
Intensity (counts)
time (min)
2.223
C
4V WGG-COOH 319
Intensity (counts)
time (min)
2.106
D
4V WGG-COOH 399
Intensity (counts)
time (min)
2.223
E
4V WGG-COOH Reactant
1e6
Intensity (counts)
mass to charge (m/z)
121.0509
239.11
441.179
F
4V WGG-COOH Product
1e6
Intensity (counts)
mass to charge (m/z)
121.0509
239.1087
G
4V WGG-COOH Reactant
Y(Counts)
X(Thomsons)
319.1421
320.1468
321.1487
H
4V WGG-COOH Brominated Product
Y(Counts)
X(Thomsons)
397.0528
399.0525

#### RebH Brominated Dipeptide Mass Spectra

#### RebH AW

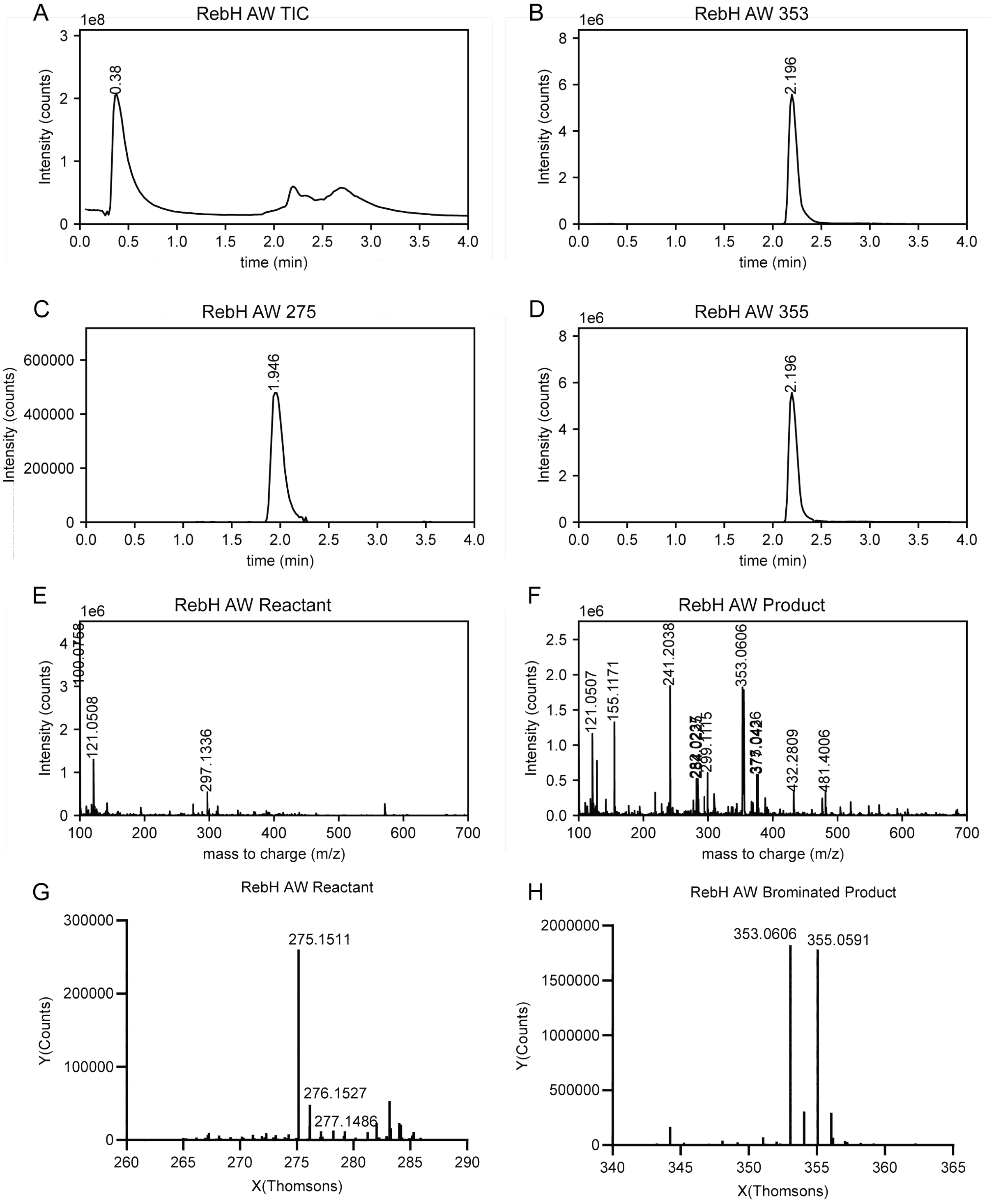
A
RebH AW TIC
1e8
Intensity (counts)
0.38
time (min)
B
RebH AW 353
1e6
Intensity (counts)
2.196
time (min)
C
RebH AW 275
Intensity (counts)
1.946
time (min)
D
RebH AW 355
1e6
Intensity (counts)
2.196
time (min)
E
RebH AW Reactant
1e6
Intensity (counts)
121.0508
297.1336
mass to charge (m/z)
F
RebH AW Product
1e6
Intensity (counts)
121.0507
155.1171
241.2038
299.1115
353.0606
432.2809
481.4006
mass to charge (m/z)
G
RebH AW Reactant
Y(Counts)
275.1511
276.1527
277.1486
X(Thomsons)
H
RebH AW Brominated Product
Y(Counts)
353.0606
355.0591
X(Thomsons)

#### RebH CW

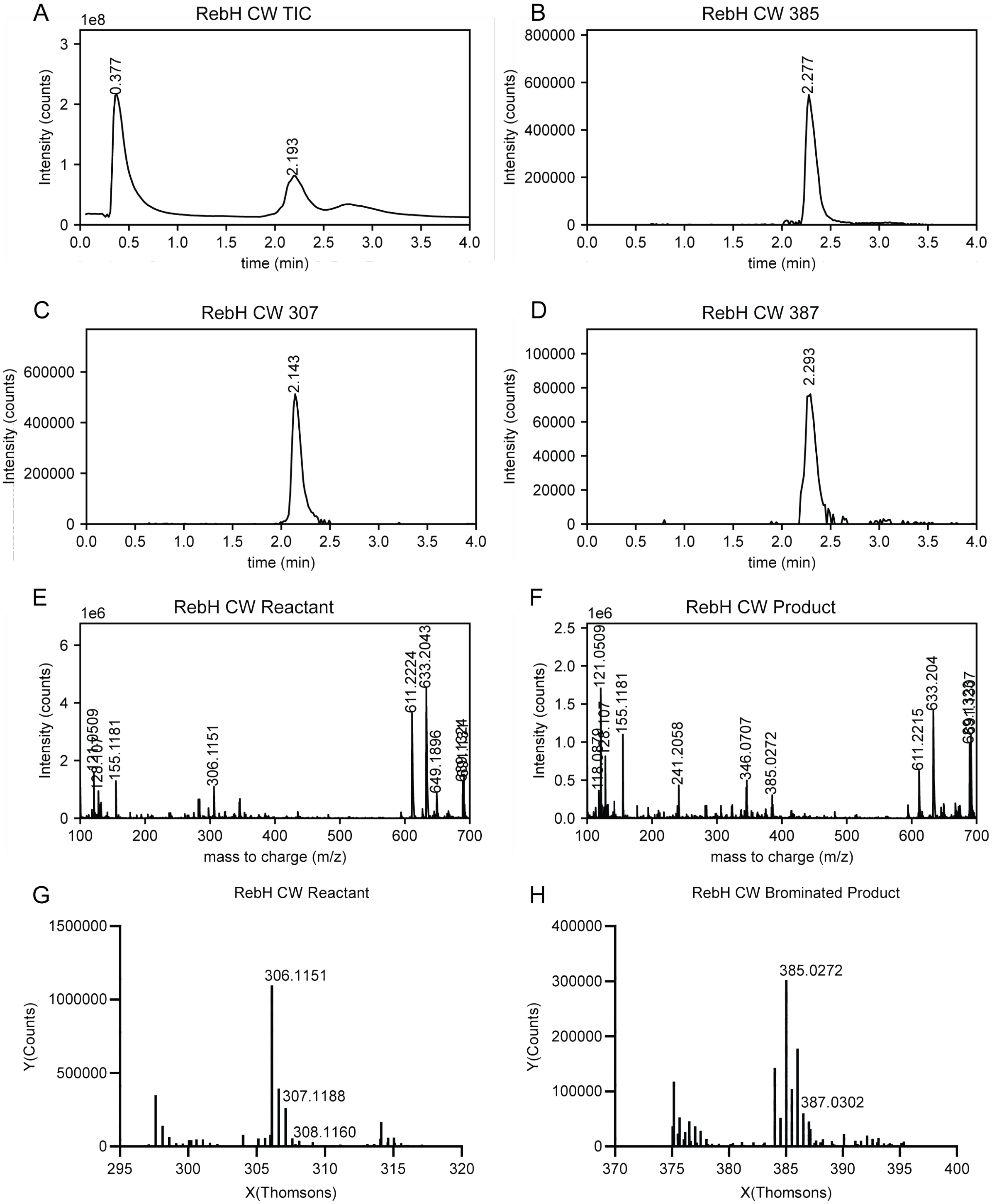
A
RebH CW TIC
1e8
Intensity (counts)
time (min)
0.377
2.193
B
RebH CW 385
2.277
C
RebH CW 307
2.143
D
RebH CW 387
2.293
E
RebH CW Reactant
1e6
mass to charge (m/z)
121.0509
128.107
155.1181
306.1151
611.2224
633.2043
649.1896
F
RebH CW Product
118.0879
121.0509
128.107
155.1181
241.2058
346.0707
385.0272
611.2215
633.204
G
RebH CW Reactant
Y(Counts)
X(Thomsons)
306.1151
307.1188
308.1160
H
RebH CW Brominated Product
385.0272
387.0302

### RebH DW

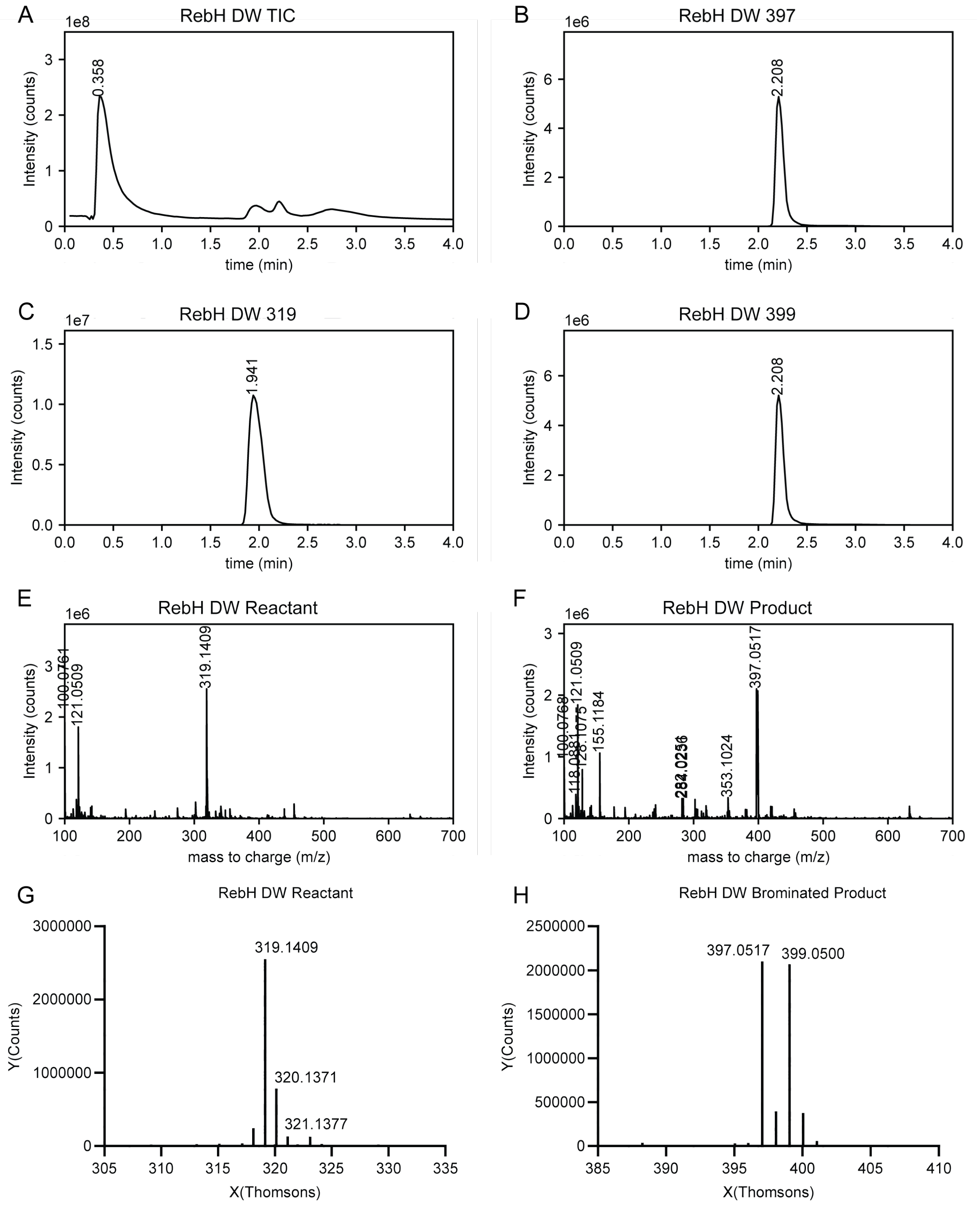
A
RebH DW TIC
1e8
0.358
Intensity (counts)
time (min)
B
RebH DW 397
1e6
2.208
Intensity (counts)
time (min)
C
RebH DW 319
1e7
1.941
Intensity (counts)
time (min)
D
RebH DW 399
1e6
2.208
Intensity (counts)
time (min)
E
RebH DW Reactant
1e6
121.0509
319.1409
Intensity (counts)
mass to charge (m/z)
F
RebH DW Product
1e6
121.0509
155.1184
353.1024
397.0517
Intensity (counts)
mass to charge (m/z)
G
RebH DW Reactant
319.1409
320.1371
321.1377
Y(Counts)
X(Thomsons)
H
RebH DW Brominated Product
397.0517
399.0500
Y(Counts)
X(Thomsons)

#### RebH EW

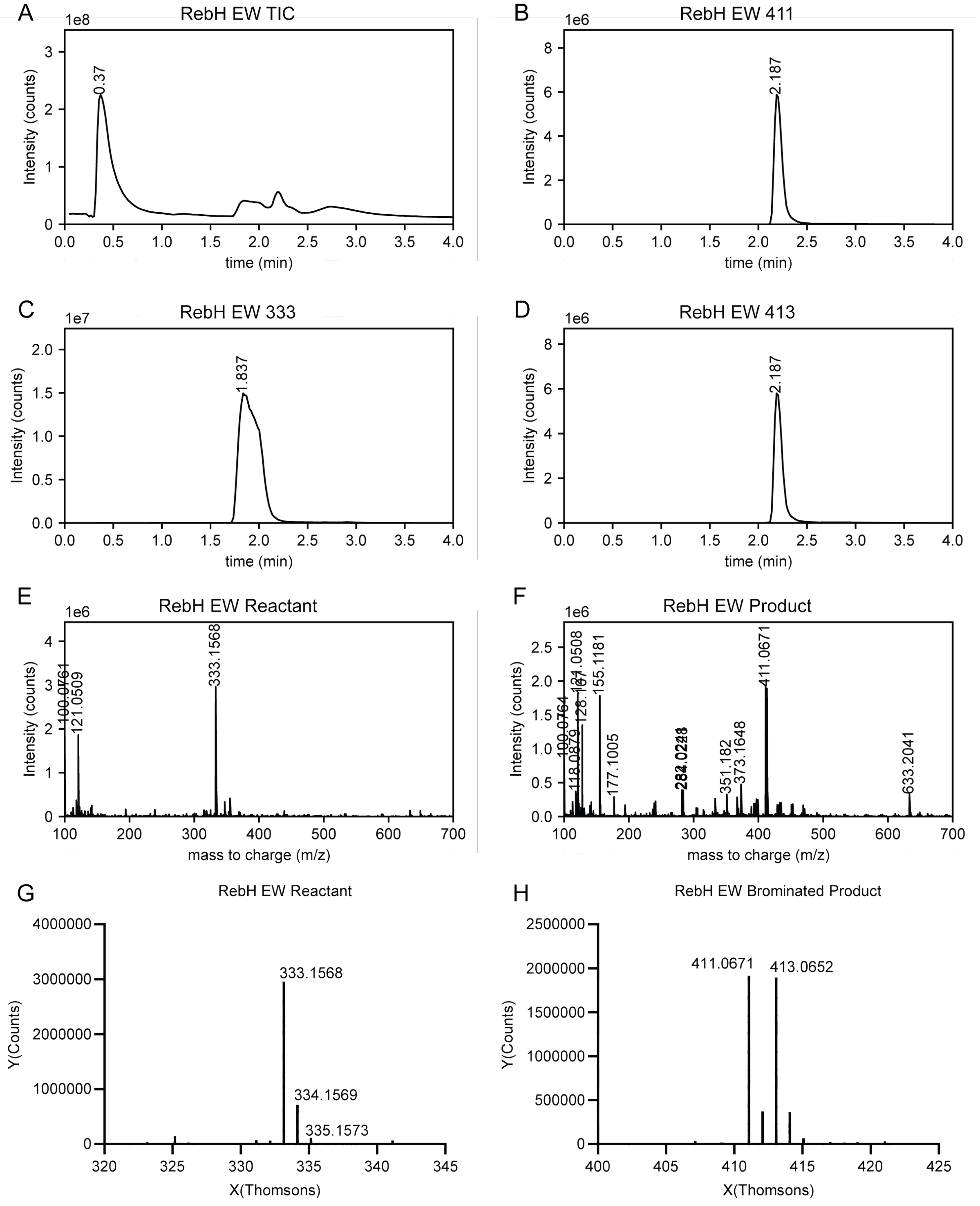
A
RebH EW TIC
1e8
0.37
Intensity (counts)
time (min)
B
RebH EW 411
1e6
2.187
Intensity (counts)
time (min)
C
RebH EW 333
1e7
1.837
Intensity (counts)
time (min)
D
RebH EW 413
1e6
2.187
Intensity (counts)
time (min)
E
RebH EW Reactant
1e6
100.0761
121.0509
333.1568
Intensity (counts)
mass to charge (m/z)
F
RebH EW Product
1e6
100.0764
118.0879
155.1181
177.1005
351.182
373.1648
411.0671
633.2041
Intensity (counts)
mass to charge (m/z)
G
RebH EW Reactant
333.1568
334.1569
335.1573
Y(Counts)
X(Thomsons)
H
RebH EW Brominated Product
411.0671
413.0652
Y(Counts)
X(Thomsons)

#### RebH FW

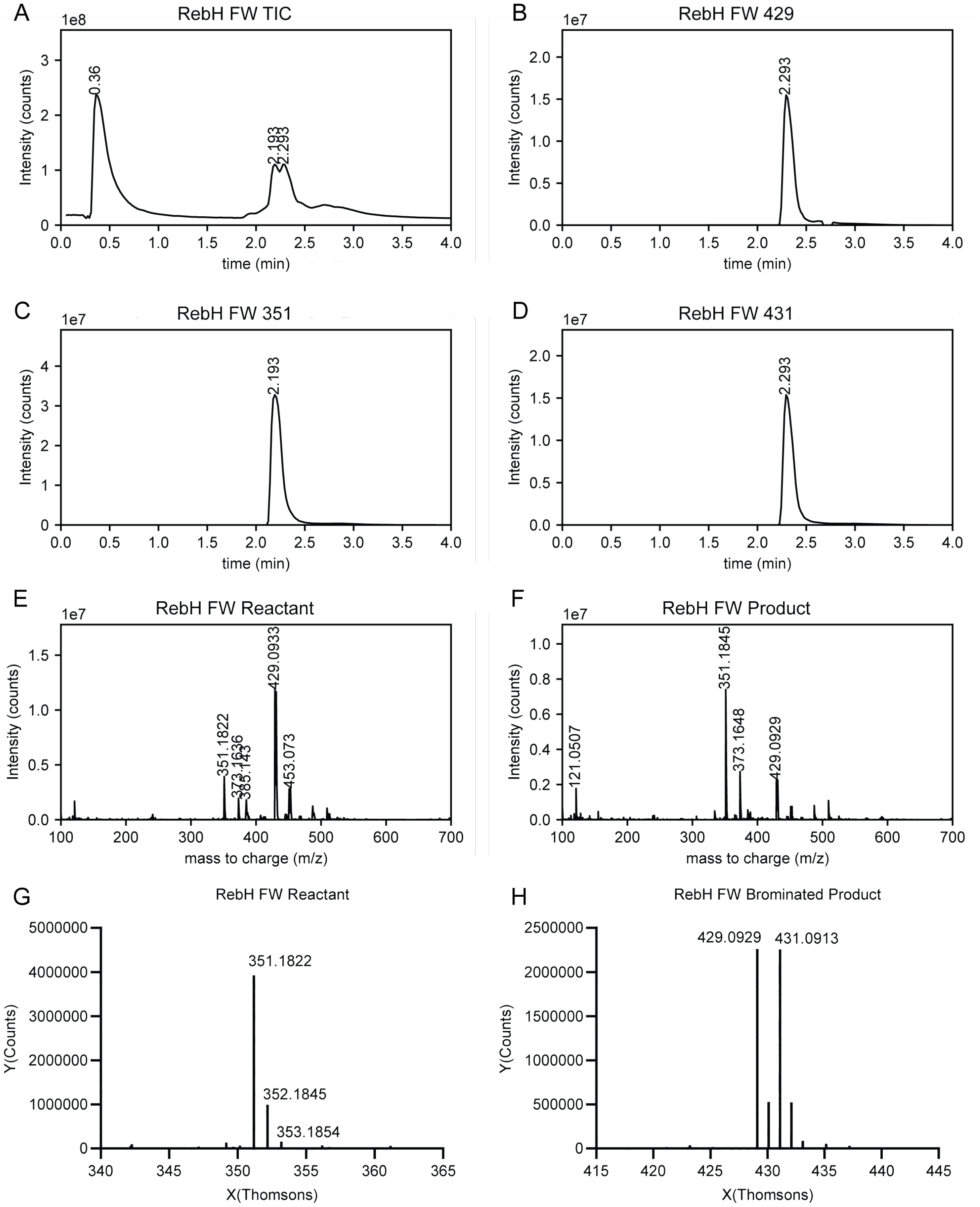

A
RebH FW TIC
1e8
Intensity (counts)
0.36
2.193
2.293
time (min)
B
RebH FW 429
1e7
2.293
time (min)
C
RebH FW 351
1e7
2.193
time (min)
D
RebH FW 431
1e7
2.293
time (min)
E
RebH FW Reactant
1e7
429.0933
351.1822
373.1636
385.143
453.073
mass to charge (m/z)
F
RebH FW Product
1e7
351.1845
373.1648
429.0929
121.0507
mass to charge (m/z)
G
RebH FW Reactant
351.1822
352.1845
353.1854
Y(Counts)
X(Thomsons)
H
RebH FW Brominated Product
429.0929
431.0913
Y(Counts)
X(Thomsons)

#### RebH GW

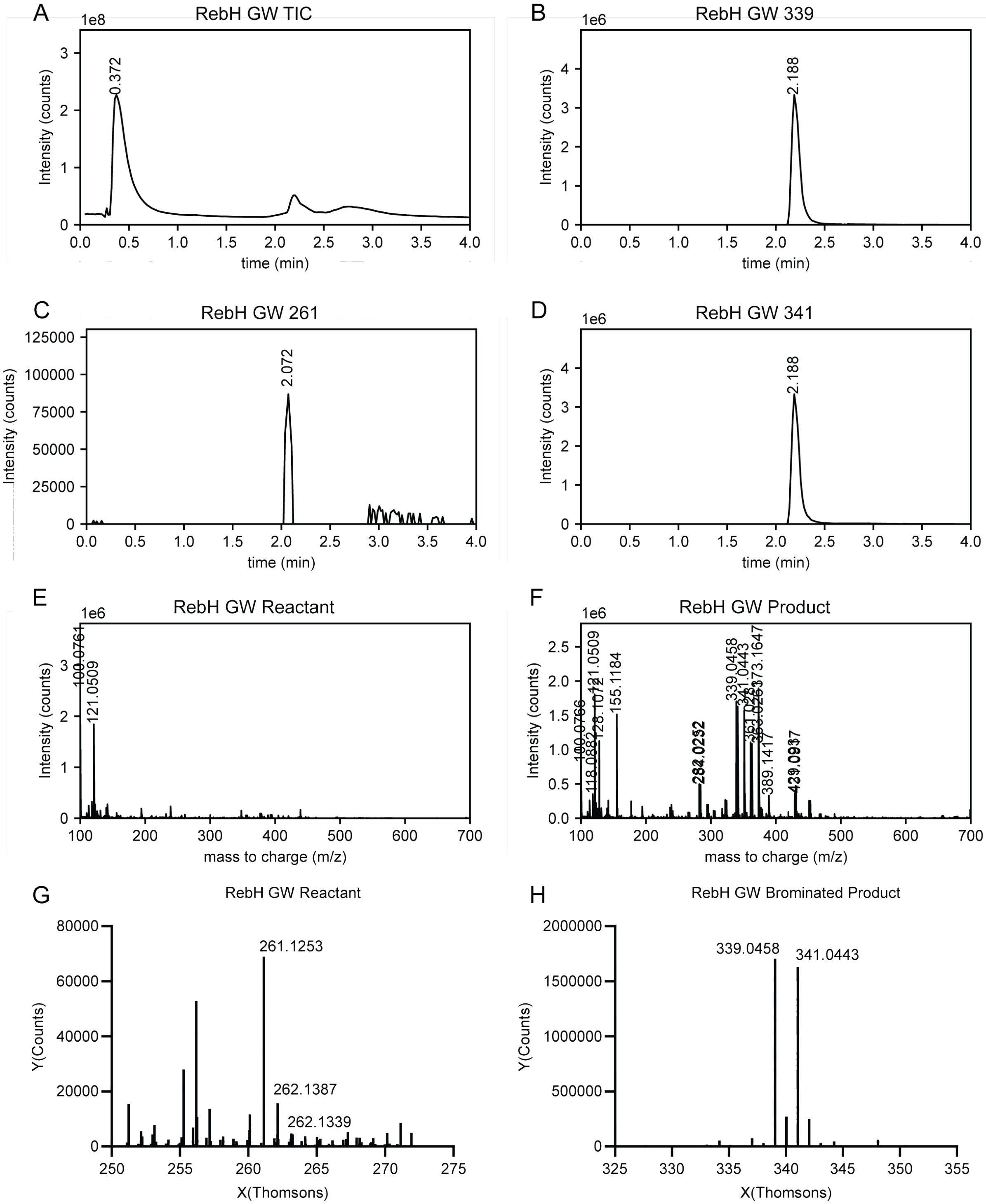

A
RebH GW TIC
1e8
0.372
Intensity (counts)
time (min)
B
RebH GW 339
1e6
2.188
Intensity (counts)
time (min)
C
RebH GW 261
2.072
Intensity (counts)
time (min)
D
RebH GW 341
1e6
2.188
Intensity (counts)
time (min)
E
RebH GW Reactant
1e6
100.0761
121.0509
Intensity (counts)
mass to charge (m/z)
F
RebH GW Product
1e6
121.0509
155.1184
339.0458
341.0443
373.1647
389.1417
Intensity (counts)
mass to charge (m/z)
G
RebH GW Reactant
261.1253
262.1387
262.1339
Y(Counts)
X(Thomsons)
H
RebH GW Brominated Product
339.0458
341.0443
Y(Counts)
X(Thomsons)

#### RebH HW

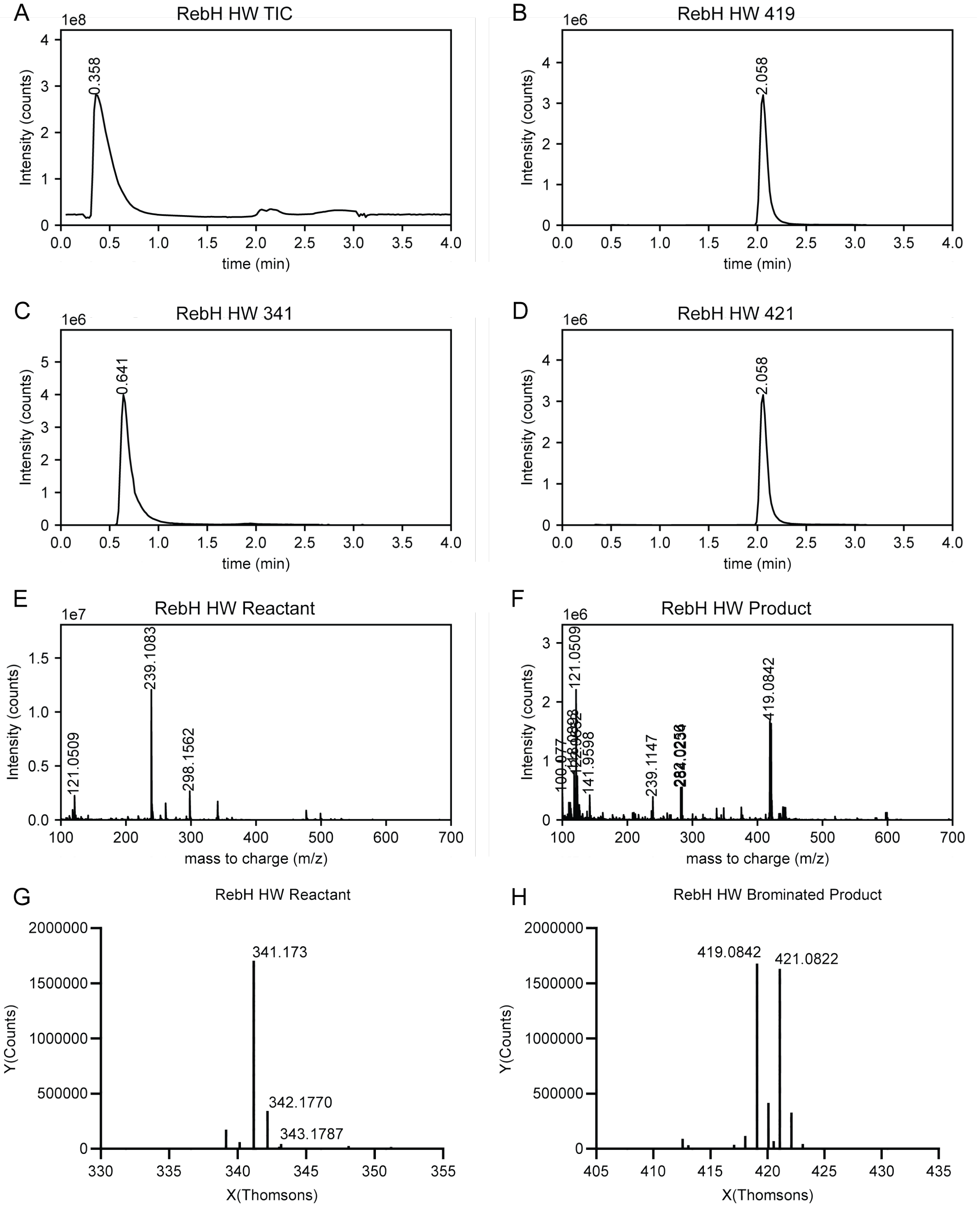
A
RebH HW TIC
1e8
Intensity (counts)
0.358
time (min)
B
RebH HW 419
1e6
2.058
C
RebH HW 341
0.641
D
RebH HW 421
2.058
E
RebH HW Reactant
1e7
121.0509
239.1083
298.1562
mass to charge (m/z)
F
RebH HW Product
121.0509
141.9598
239.1147
419.0842
G
RebH HW Reactant
341.173
342.1770
343.1787
Y(Counts)
X(Thomsons)
H
RebH HW Brominated Product
419.0842
421.0822

#### RebH IW

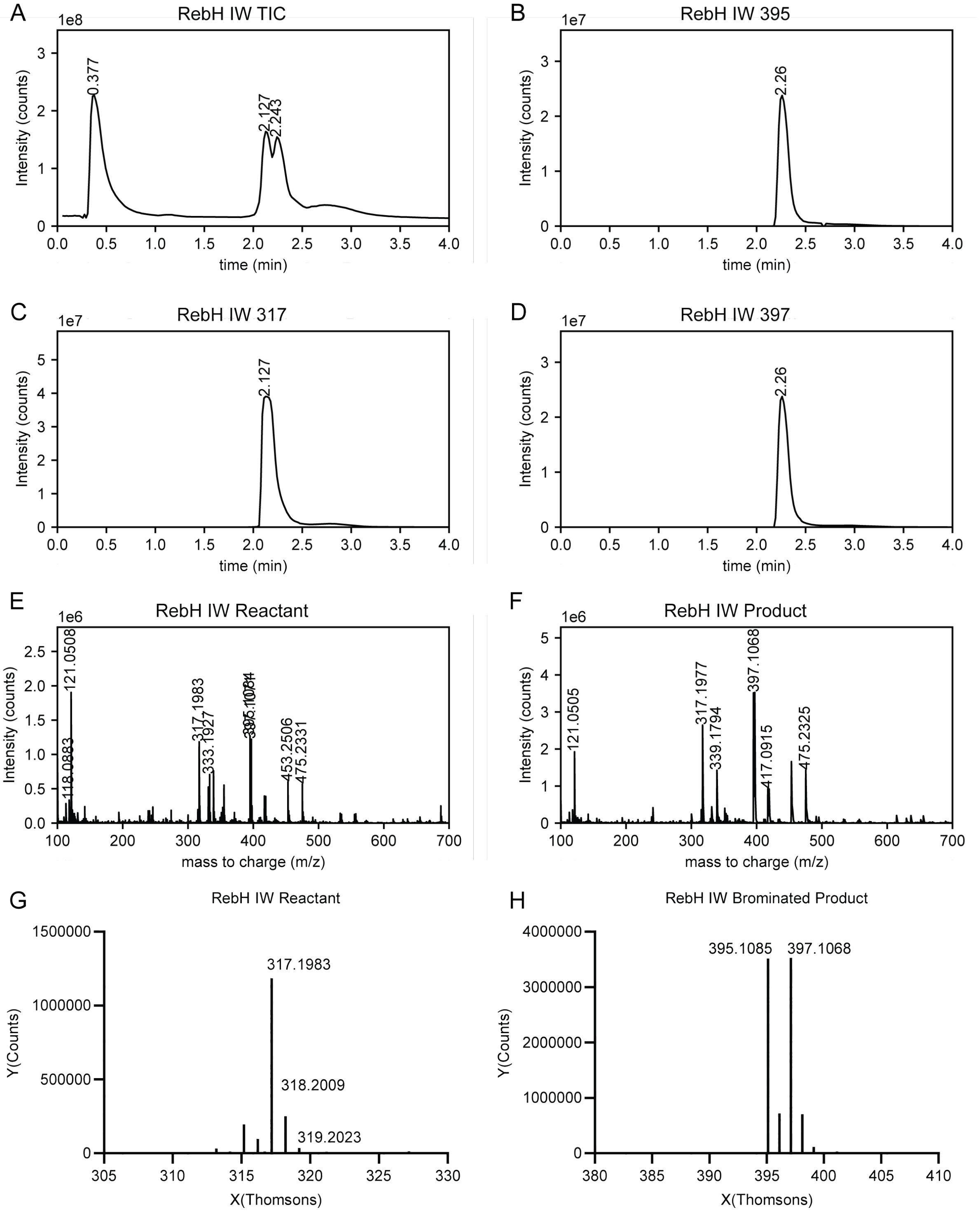
A
RebH IW TIC
1e8
Intensity (counts)
time (min)
0.377
2.127
2.243
B
RebH IW 395
1e7
2.26
C
RebH IW 317
1e7
2.127
D
RebH IW 397
1e7
2.26
E
RebH IW Reactant
1e6
mass to charge (m/z)
121.0508
118.0883
317.1983
333.1927
453.2506
475.2331
F
RebH IW Product
1e6
121.0505
317.1977
339.1794
397.1068
417.0915
475.2325
G
RebH IW Reactant
Y(Counts)
X(Thomsons)
317.1983
318.2009
319.2023
H
RebH IW Brominated Product
395.1085
397.1068

#### RebH KW

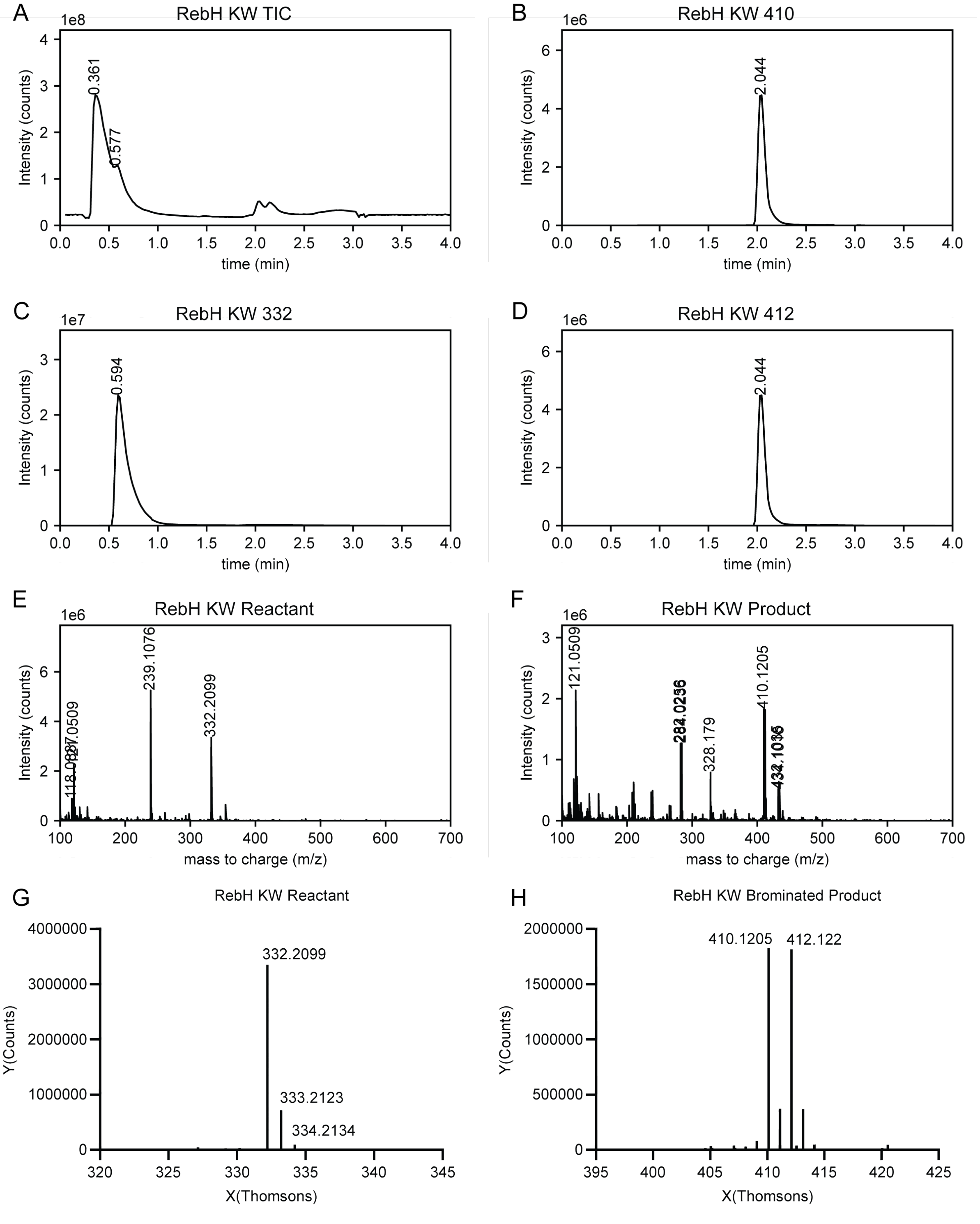

A
RebH KW TIC
1e8
Intensity (counts)
0.361
0.577
time (min)
B
RebH KW 410
1e6
2.044
C
RebH KW 332
1e7
0.594
D
RebH KW 412
1e6
2.044
E
RebH KW Reactant
239.1076
332.2099
121.0509
mass to charge (m/z)
F
RebH KW Product
121.0509
410.1205
328.179
G
RebH KW Reactant
332.2099
333.2123
334.2134
Y(Counts)
X(Thomsons)
H
RebH KW Brominated Product
410.1205
412.122

#### RebH LW

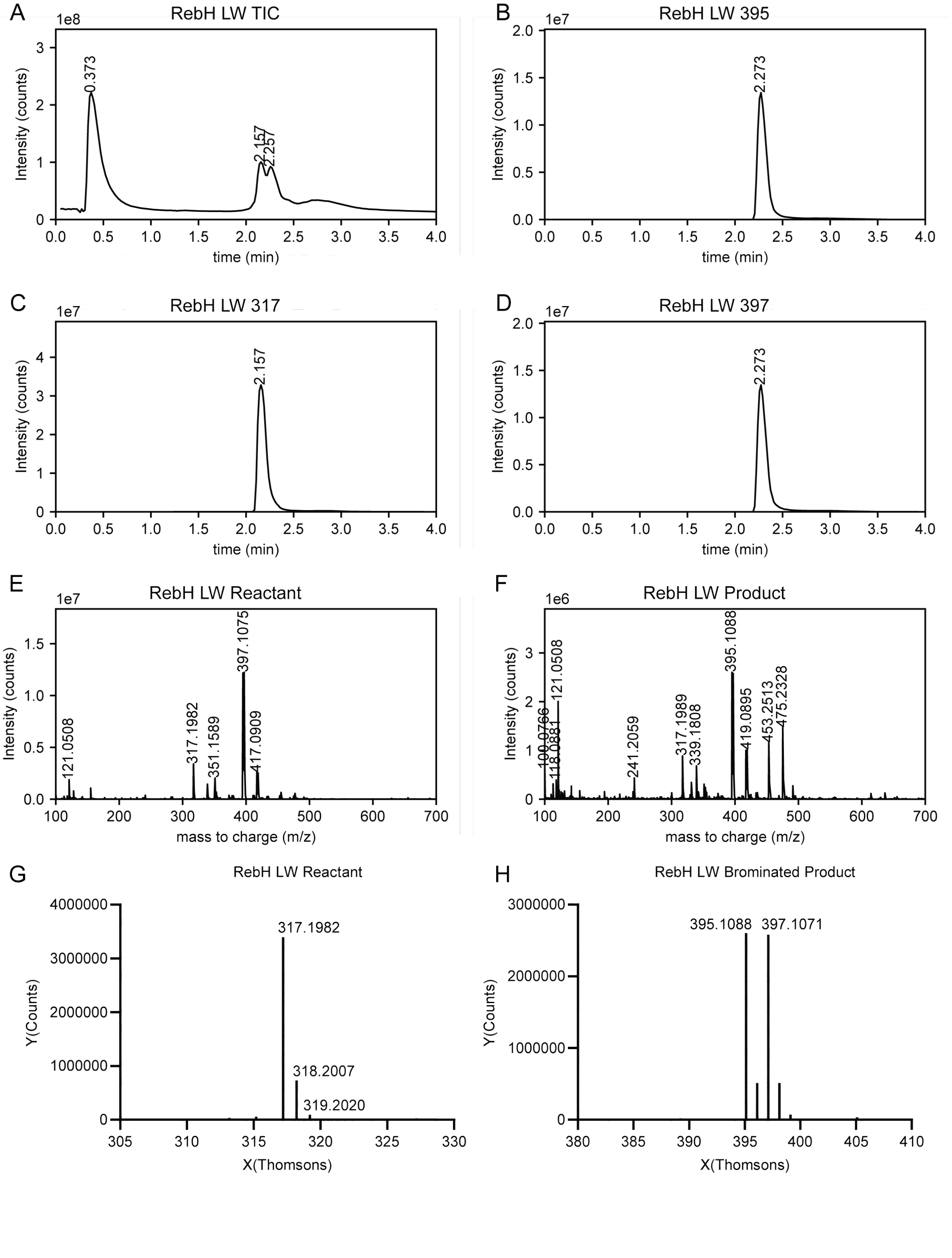
A
RebH LW TIC
1e8
Intensity (counts)
time (min)
0.373
2.157
2.257
B
RebH LW 395
1e7
Intensity (counts)
time (min)
2.273
C
RebH LW 317
1e7
Intensity (counts)
time (min)
2.157
D
RebH LW 397
1e7
Intensity (counts)
time (min)
2.273
E
RebH LW Reactant
1e7
Intensity (counts)
mass to charge (m/z)
121.0508
317.1982
351.1589
397.1075
417.0909
F
RebH LW Product
1e6
Intensity (counts)
mass to charge (m/z)
121.0508
241.2059
317.1989
339.1808
395.1088
419.0895
453.2513
475.2328
G
RebH LW Reactant
Y(Counts)
X(Thomsons)
317.1982
318.2007
319.2020
H
RebH LW Brominated Product
Y(Counts)
X(Thomsons)
395.1088
397.1071

#### RebH MW

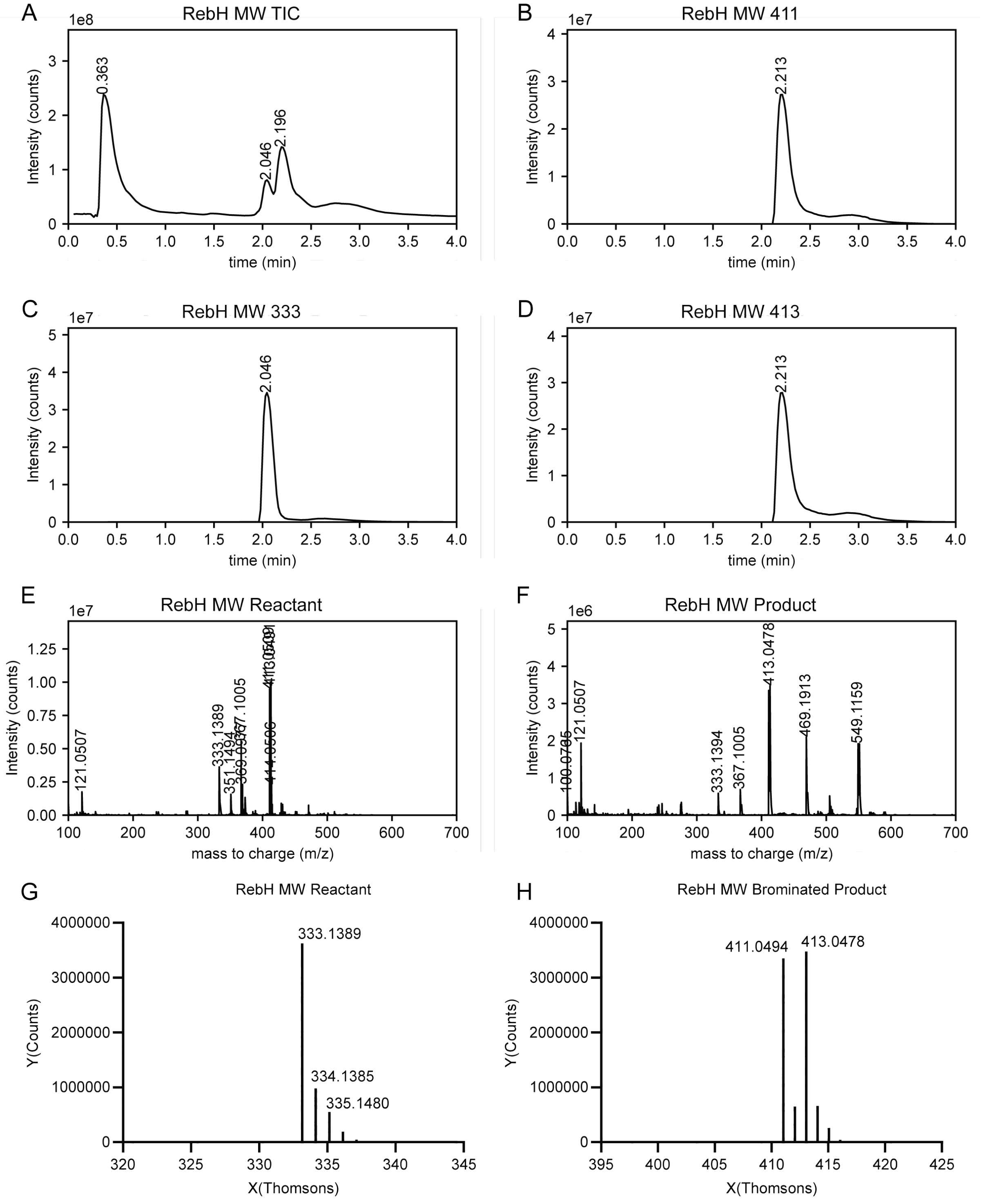
A
RebH MW TIC
1e8
Intensity (counts)
time (min)
0.363
2.046
2.196
B
RebH MW 411
1e7
Intensity (counts)
time (min)
2.213
C
RebH MW 333
1e7
Intensity (counts)
time (min)
2.046
D
RebH MW 413
1e7
Intensity (counts)
time (min)
2.213
E
RebH MW Reactant
1e7
Intensity (counts)
mass to charge (m/z)
121.0507
333.1389
367.1005
F
RebH MW Product
1e6
Intensity (counts)
mass to charge (m/z)
121.0507
333.1394
367.1005
413.0478
469.1913
549.1159
G
RebH MW Reactant
Y(Counts)
X(Thomsons)
333.1389
334.1385
335.1480
H
RebH MW Brominated Product
Y(Counts)
X(Thomsons)
411.0494
413.0478

**RebH NW**

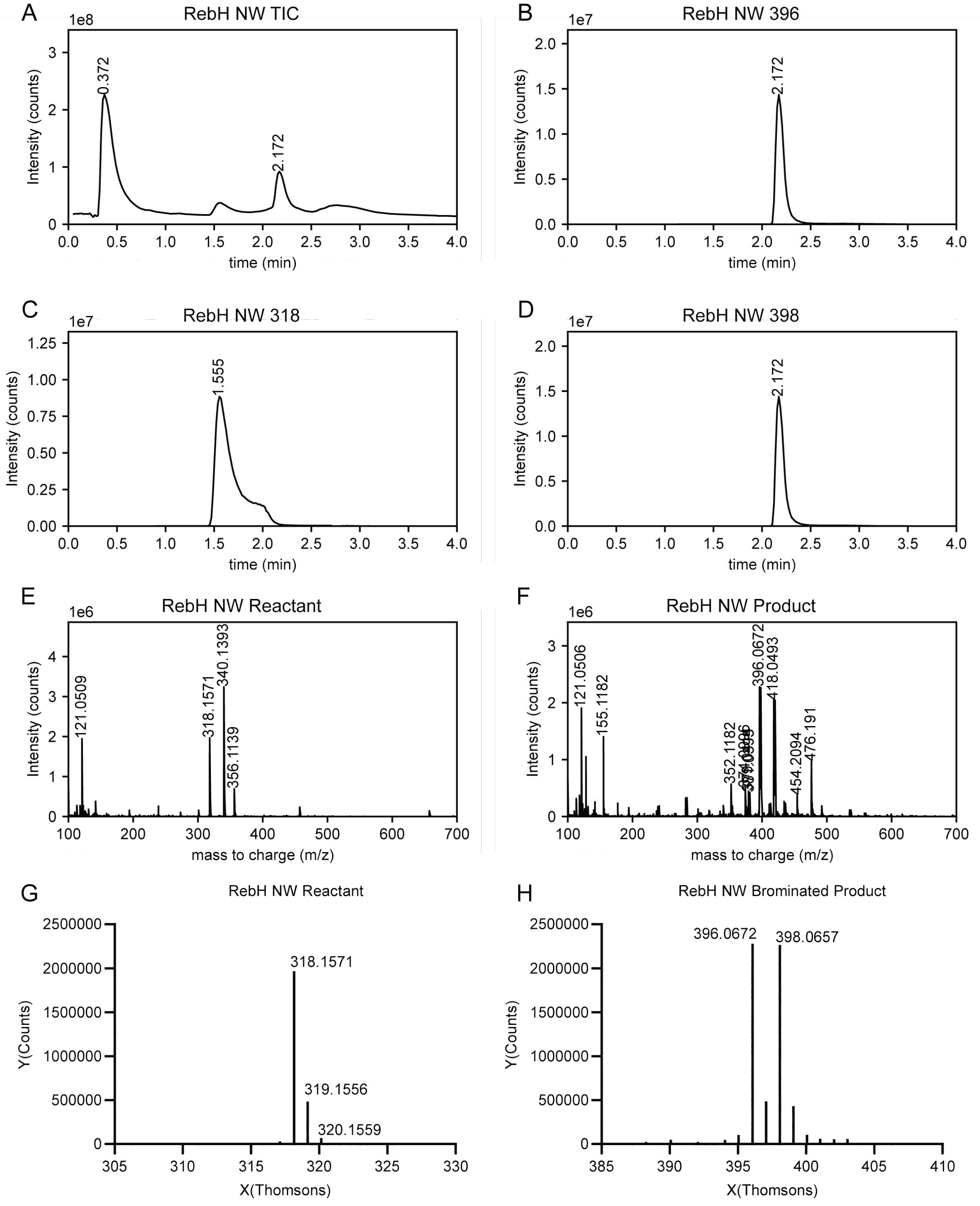
A
RebH NW TIC
1e8
Intensity (counts)
0.372
2.172
time (min)
B
RebH NW 396
1e7
2.172
C
RebH NW 318
1.555
D
RebH NW 398
E
RebH NW Reactant
1e6
121.0509
318.1571
340.1393
356.1139
mass to charge (m/z)
F
RebH NW Product
121.0506
155.1182
352.1182
396.0672
418.0493
454.2094
476.191
G
RebH NW Reactant
Y(Counts)
318.1571
319.1556
320.1559
X(Thomsons)
H
RebH NW Brominated Product
396.0672
398.0657

#### RebH PW

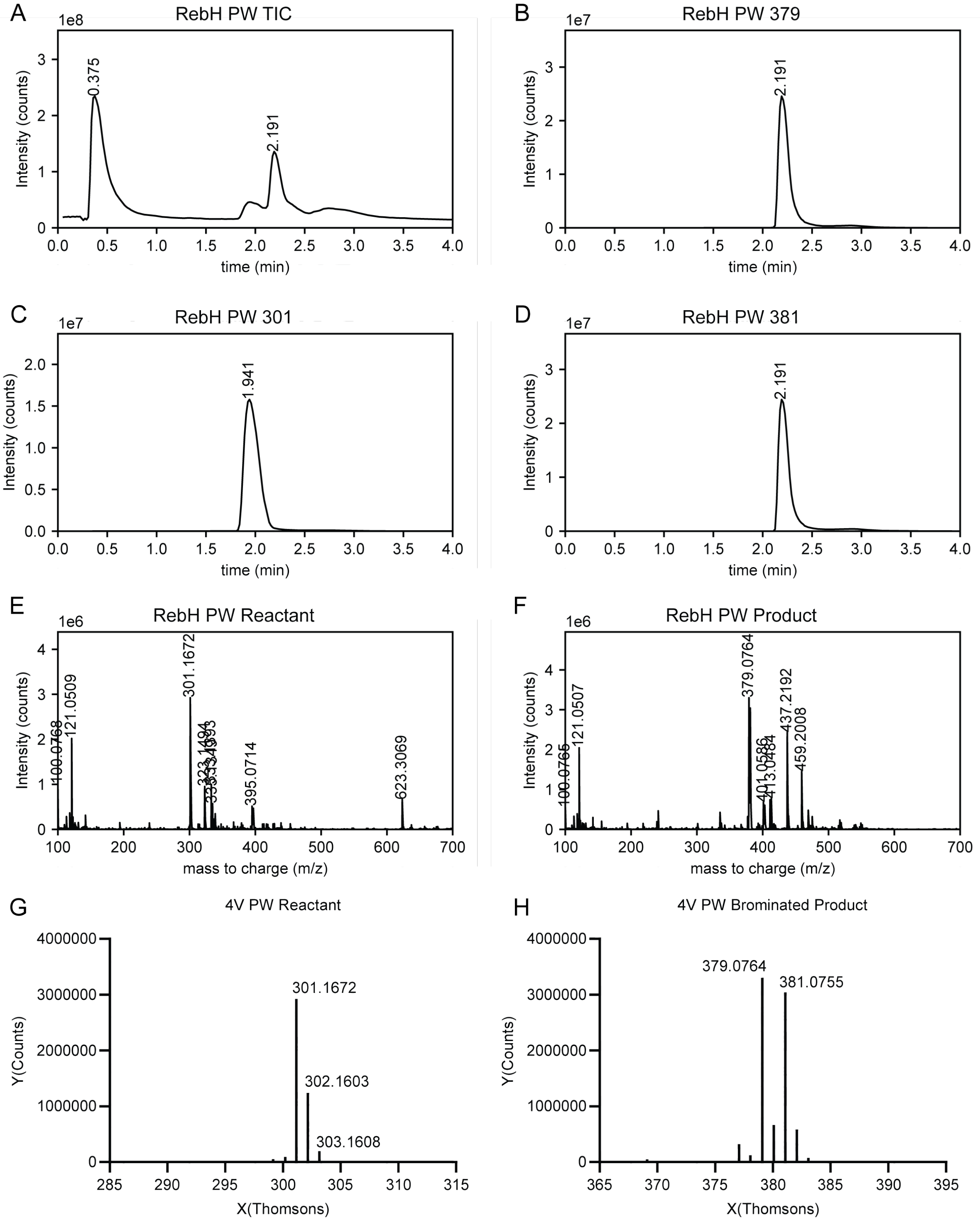

A
RebH PW TIC
1e8
Intensity (counts)
time (min)
0.375
2.191
B
RebH PW 379
1e7
2.191
C
RebH PW 301
1e7
1.941
D
RebH PW 381
1e7
2.191
E
RebH PW Reactant
1e6
mass to charge (m/z)
100.0768
121.0509
301.1672
395.0714
623.3069
F
RebH PW Product
1e6
100.0766
121.0507
379.0764
401.0586
437.2192
459.2008
G
4V PW Reactant
Y(Counts)
X(Thomsons)
301.1672
302.1603
303.1608
H
4V PW Brominated Product
379.0764
381.0755

#### RebH QW

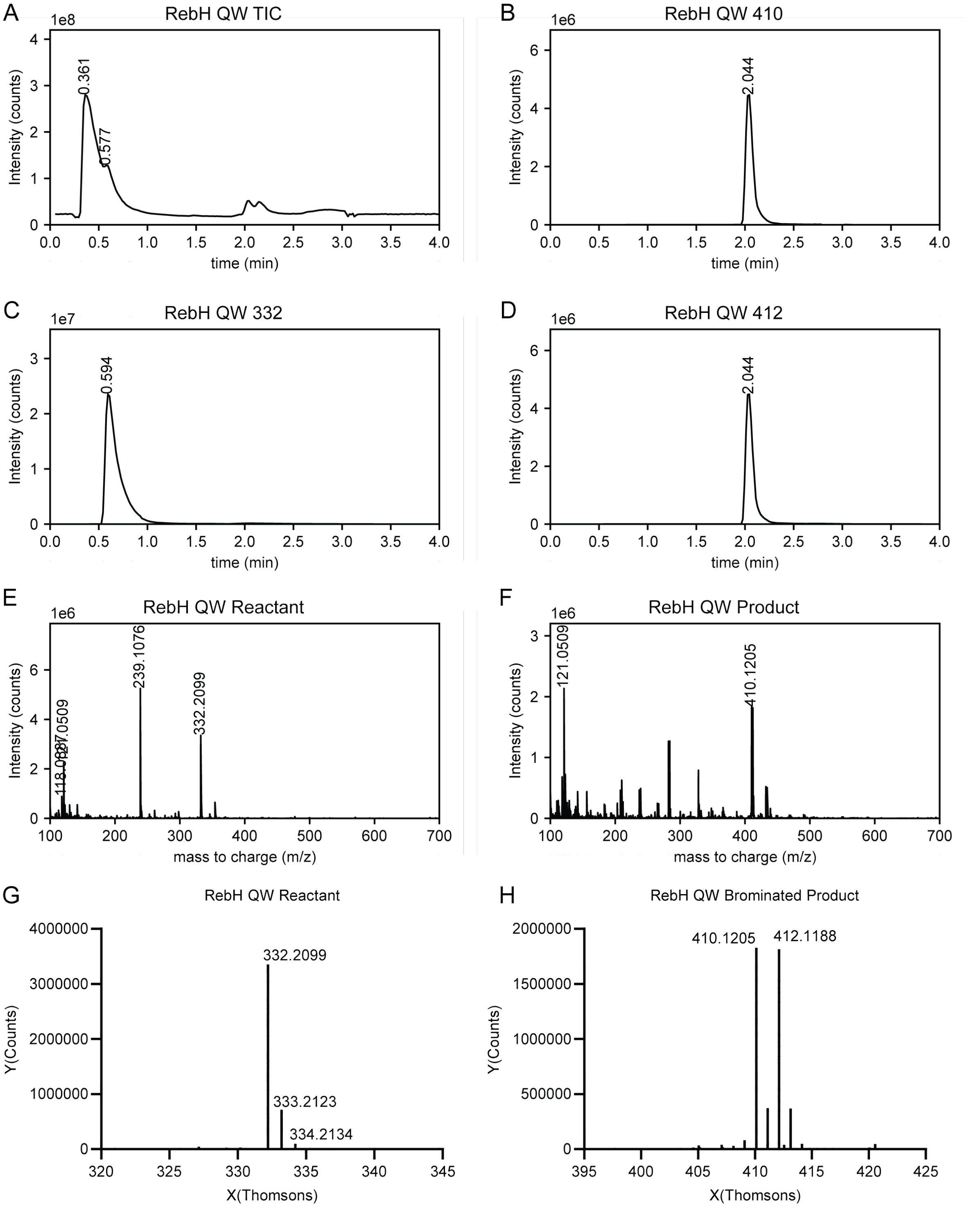

A
RebH QW TIC
1e8
Intensity (counts)
time (min)
0.361
0.577
B
RebH QW 410
1e6
Intensity (counts)
time (min)
2.044
C
RebH QW 332
1e7
Intensity (counts)
time (min)
0.594
D
RebH QW 412
1e6
Intensity (counts)
time (min)
2.044
E
RebH QW Reactant
1e6
Intensity (counts)
mass to charge (m/z)
239.1076
332.2099
F
RebH QW Product
1e6
Intensity (counts)
mass to charge (m/z)
121.0509
410.1205
G
RebH QW Reactant
Y(Counts)
X(Thomsons)
332.2099
333.2123
334.2134
H
RebH QW Brominated Product
Y(Counts)
X(Thomsons)
410.1205
412.1188

#### RebH RW

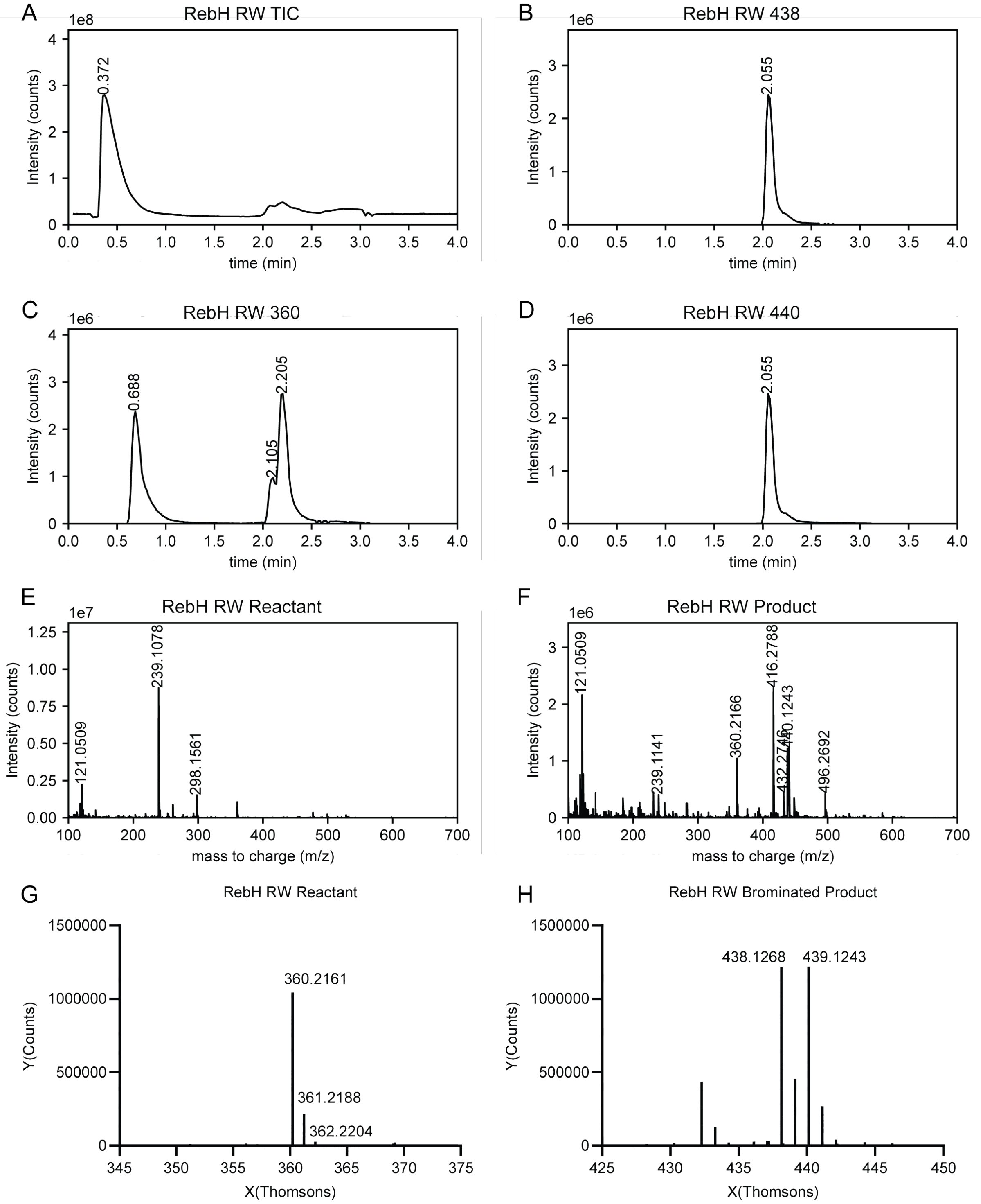

A
RebH RW TIC
1e8
Intensity (counts)
time (min)
0.372
B
RebH RW 438
1e6
2.055
C
RebH RW 360
1e6
0.688
2.105
2.205
D
RebH RW 440
1e6
2.055
E
RebH RW Reactant
1e7
mass to charge (m/z)
121.0509
239.1078
298.1561
F
RebH RW Product
1e6
121.0509
239.1141
360.2166
416.2788
432.2746
440.1243
496.2692
G
RebH RW Reactant
Y(Counts)
X(Thomsons)
360.2161
361.2188
362.2204
H
RebH RW Brominated Product
438.1268
439.1243

**RebH SW**

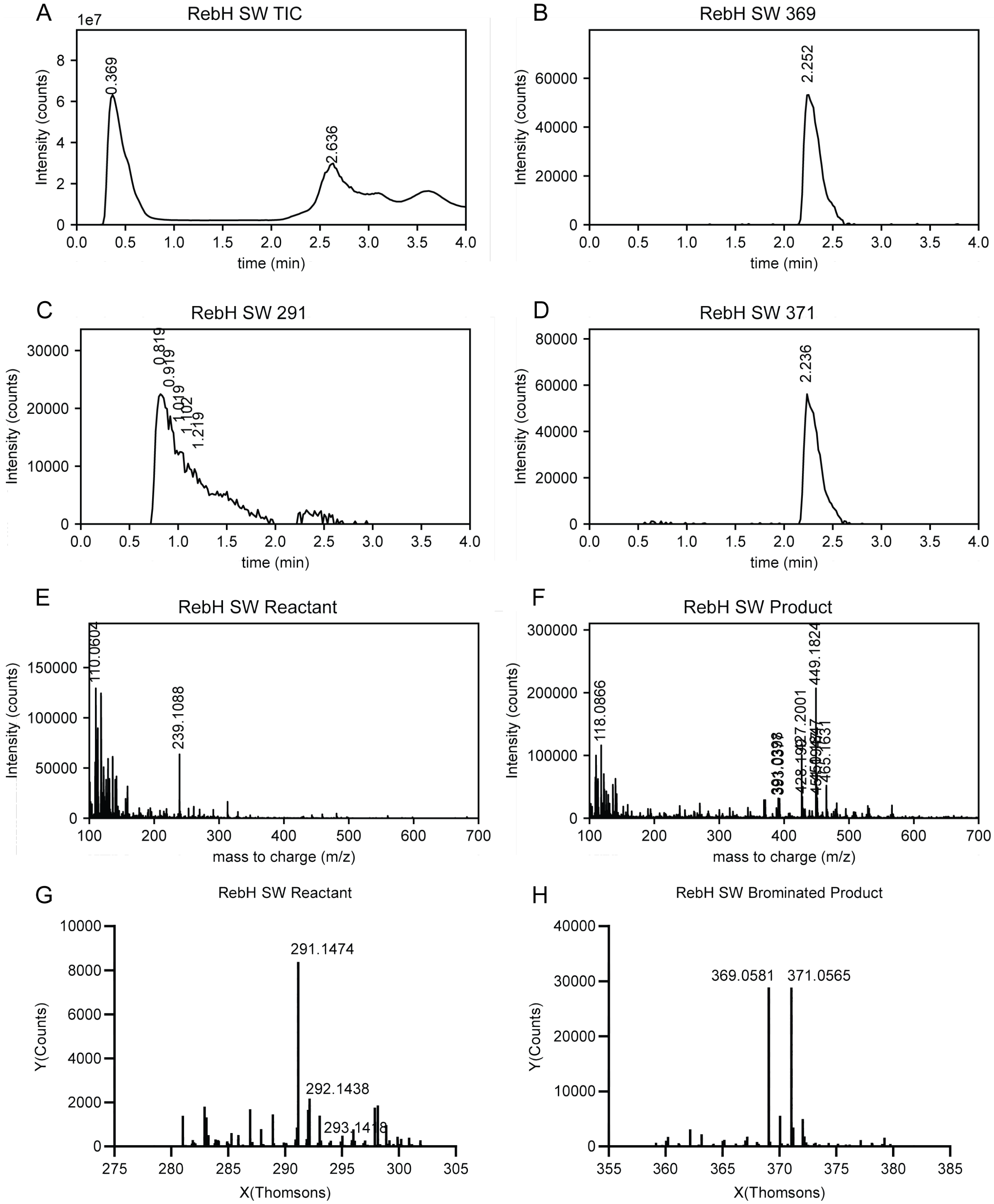
A
RebH SW TIC
1e7
Intensity (counts)
0.369
2.636
time (min)
B
RebH SW 369
2.252
Intensity (counts)
time (min)
C
RebH SW 291
0.819
0.919
1.019
1.102
1.219
Intensity (counts)
time (min)
D
RebH SW 371
2.236
Intensity (counts)
time (min)
E
RebH SW Reactant
110.0604
239.1088
Intensity (counts)
mass to charge (m/z)
F
RebH SW Product
449.1824
118.0866
427.2001
465.1631
Intensity (counts)
mass to charge (m/z)
G
RebH SW Reactant
291.1474
292.1438
293.1418
Y(Counts)
X(Thomsons)
H
RebH SW Brominated Product
369.0581
371.0565
Y(Counts)
X(Thomsons)

#### RebH TW

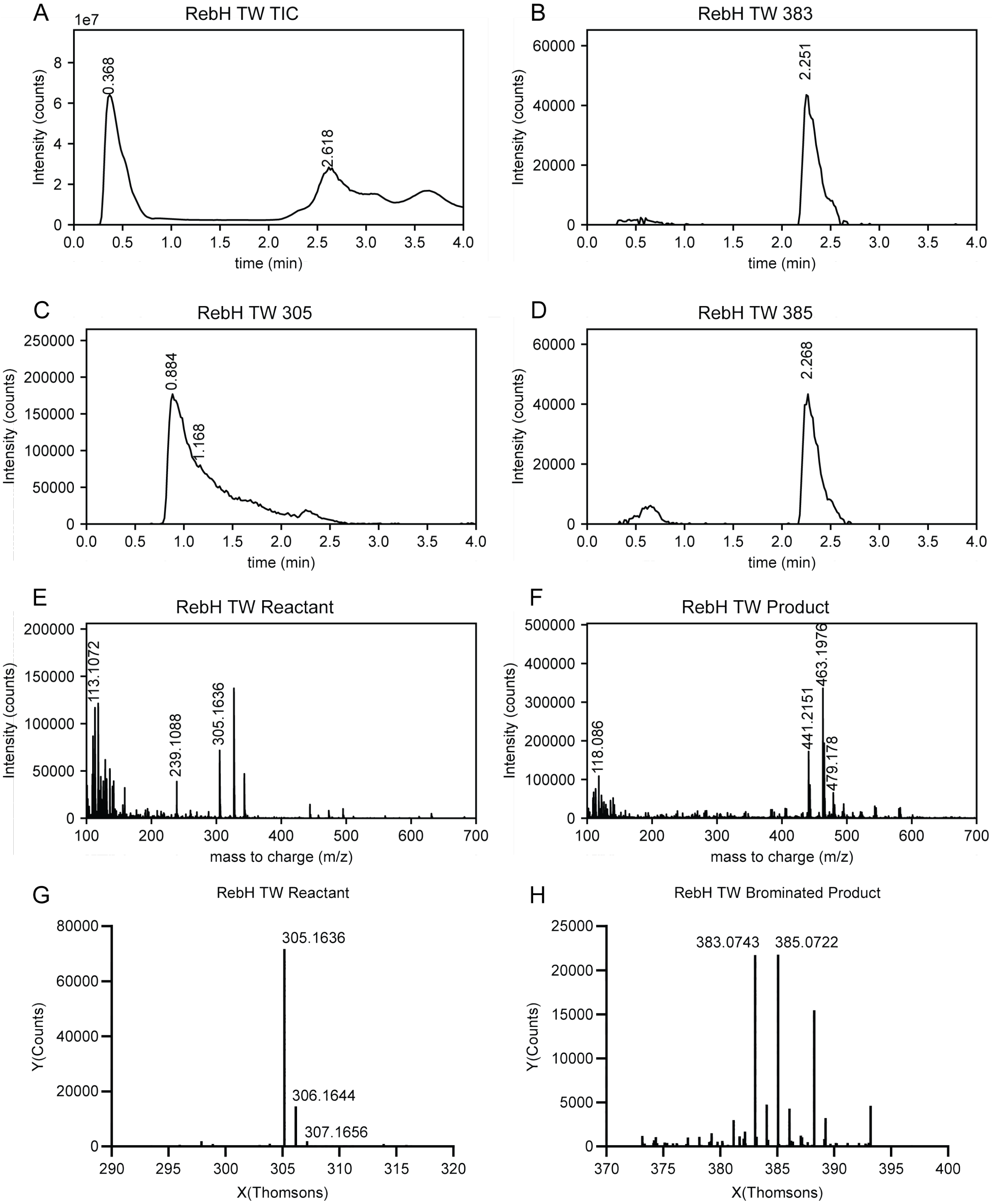
A
RebH TW TIC
1e7
Intensity (counts)
time (min)
0.368
2.618
B
RebH TW 383
2.251
C
RebH TW 305
0.884
1.168
D
RebH TW 385
2.268
E
RebH TW Reactant
mass to charge (m/z)
113.1072
239.1088
305.1636
F
RebH TW Product
118.086
441.2151
463.1976
479.178
G
RebH TW Reactant
Y(Counts)
X(Thomsons)
305.1636
306.1644
307.1656
H
RebH TW Brominated Product
383.0743
385.0722

#### RebH VW

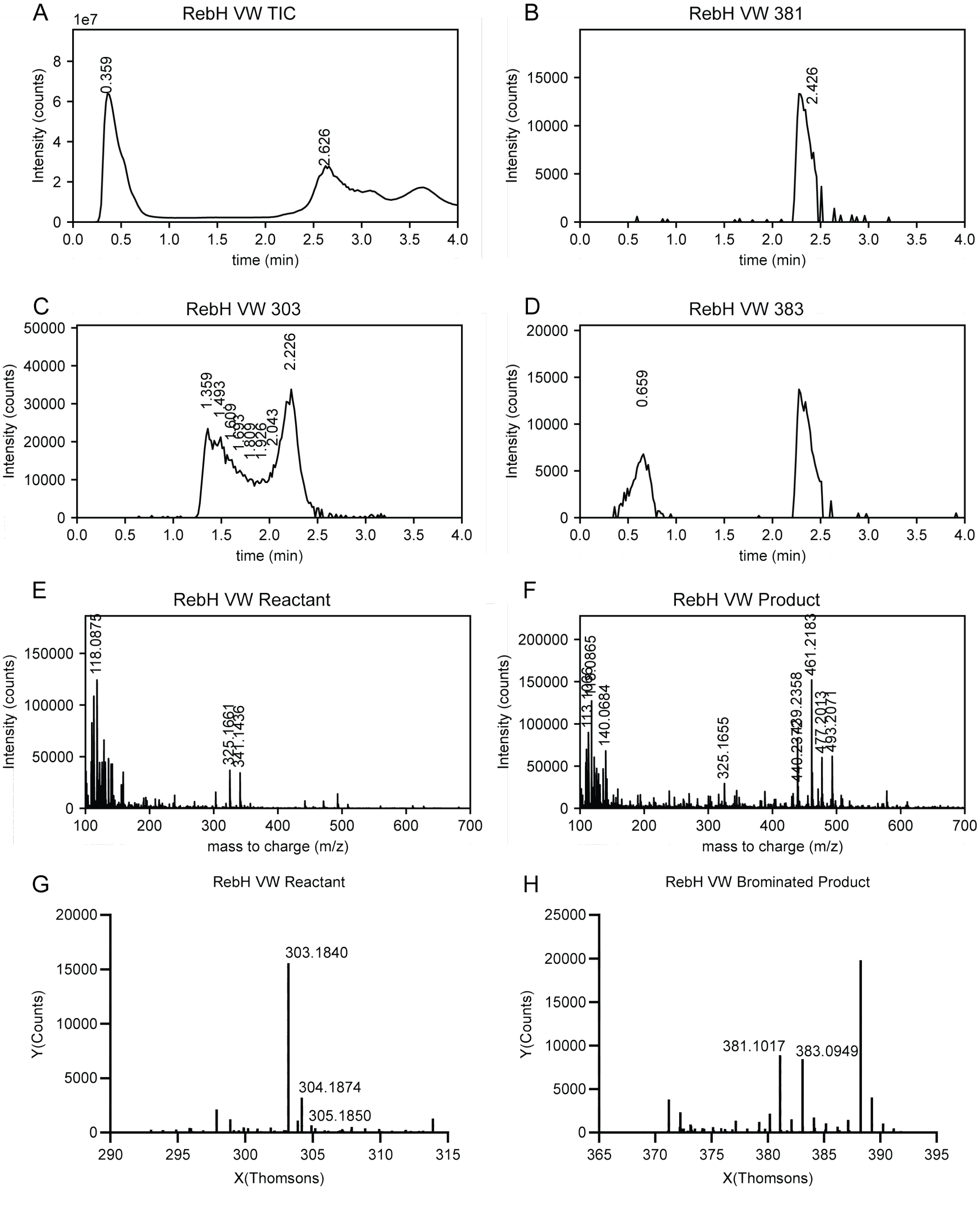
A
RebH VW TIC
1e7
Intensity (counts)
time (min)
0.359
2.626
B
RebH VW 381
Intensity (counts)
time (min)
2.426
C
RebH VW 303
Intensity (counts)
time (min)
1.359
1.493
2.043
2.226
D
RebH VW 383
Intensity (counts)
time (min)
0.659
E
RebH VW Reactant
Intensity (counts)
mass to charge (m/z)
118.0875
325.1661
341.1436
F
RebH VW Product
Intensity (counts)
mass to charge (m/z)
140.0684
325.1655
461.2183
477.2013
493.2071
G
RebH VW Reactant
Y(Counts)
X(Thomsons)
303.1840
304.1874
305.1850
H
RebH VW Brominated Product
Y(Counts)
X(Thomsons)
381.1017
383.0949

#### RebH YW

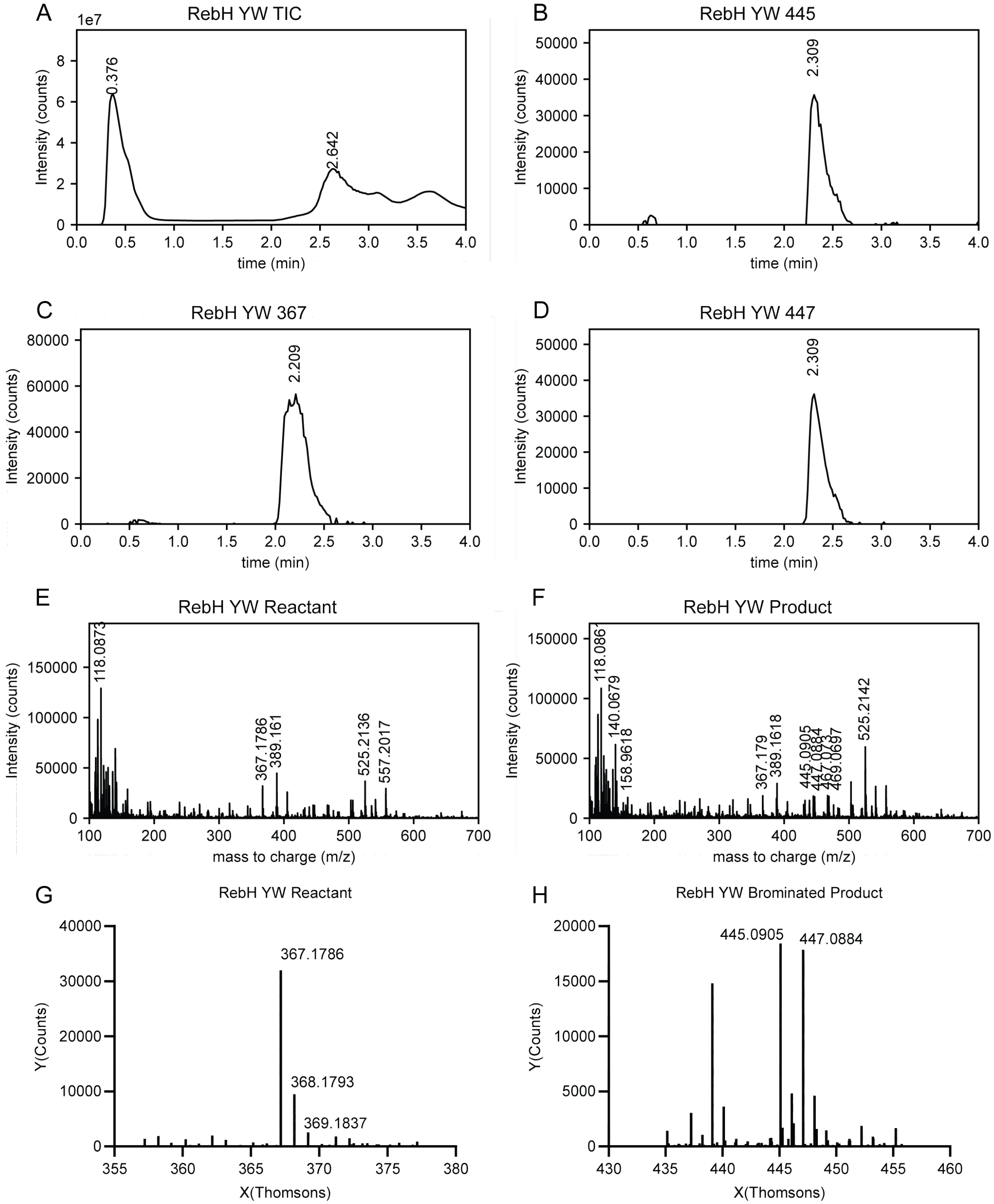

A
RebH YW TIC
1e7
Intensity (counts)
time (min)
0.376
2.642
B
RebH YW 445
Intensity (counts)
time (min)
2.309
C
RebH YW 367
Intensity (counts)
time (min)
2.209
D
RebH YW 447
Intensity (counts)
time (min)
2.309
E
RebH YW Reactant
Intensity (counts)
mass to charge (m/z)
118.0873
367.1786
389.161
525.2136
557.2017
F
RebH YW Product
Intensity (counts)
mass to charge (m/z)
118.0861
140.0679
158.9618
367.179
389.1618
445.0905
447.0884
469.0697
525.2142
G
RebH YW Reactant
Y(Counts)
X(Thomsons)
367.1786
368.1793
369.1837
H
RebH YW Brominated Product
Y(Counts)
X(Thomsons)
445.0905
447.0884

#### RebH WA

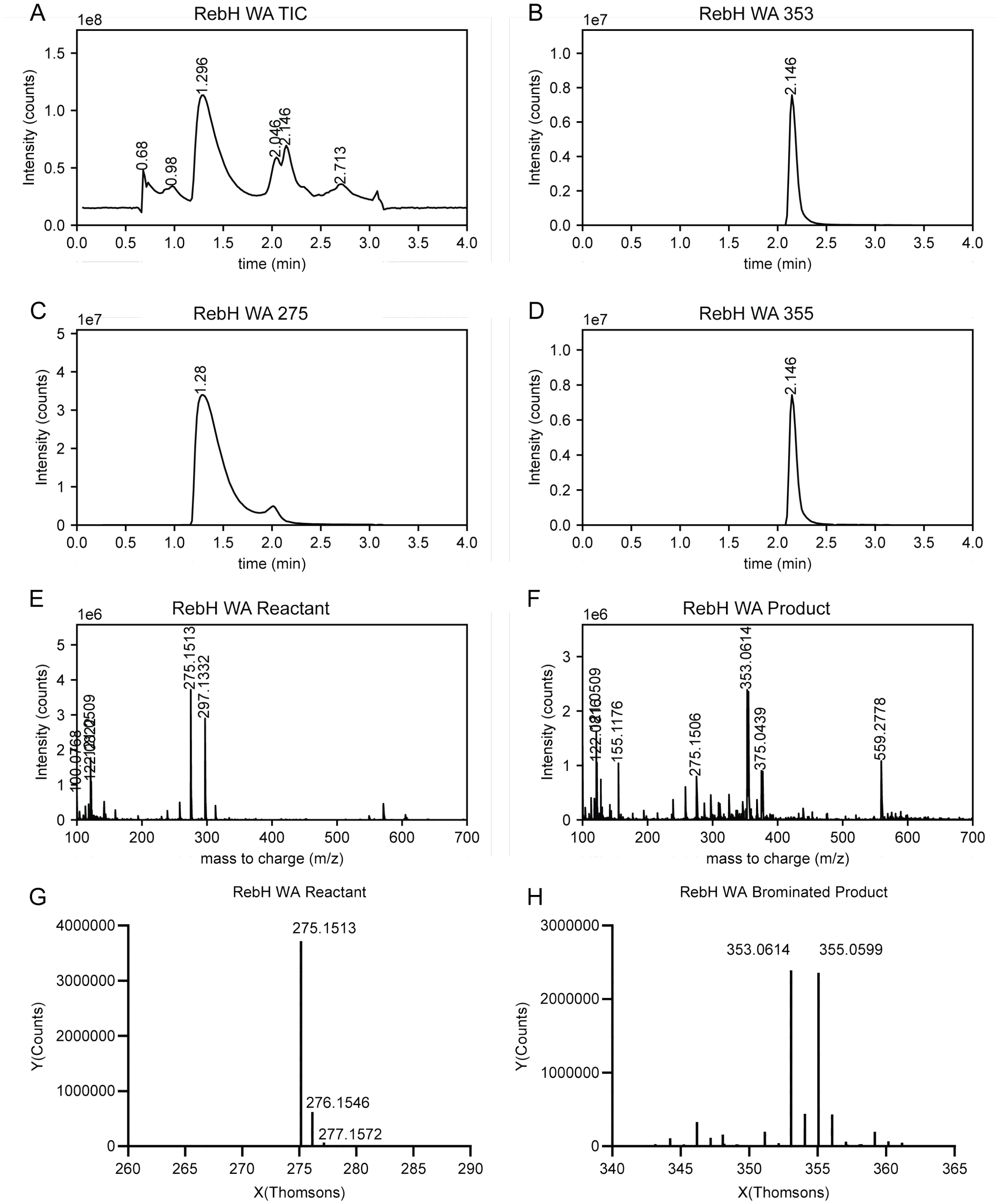
A
RebH WA TIC
1e8
Intensity (counts)
time (min)
0.68
0.98
1.296
2.046
2.146
2.713
B
RebH WA 353
1e7
2.146
C
RebH WA 275
1.28
D
RebH WA 355
2.146
E
RebH WA Reactant
1e6
mass to charge (m/z)
275.1513
297.1332
F
RebH WA Product
353.0614
155.1176
275.1506
375.0439
559.2778
G
RebH WA Reactant
Y(Counts)
X(Thomsons)
275.1513
276.1546
277.1572
H
RebH WA Brominated Product
353.0614
355.0599

#### RebH WC

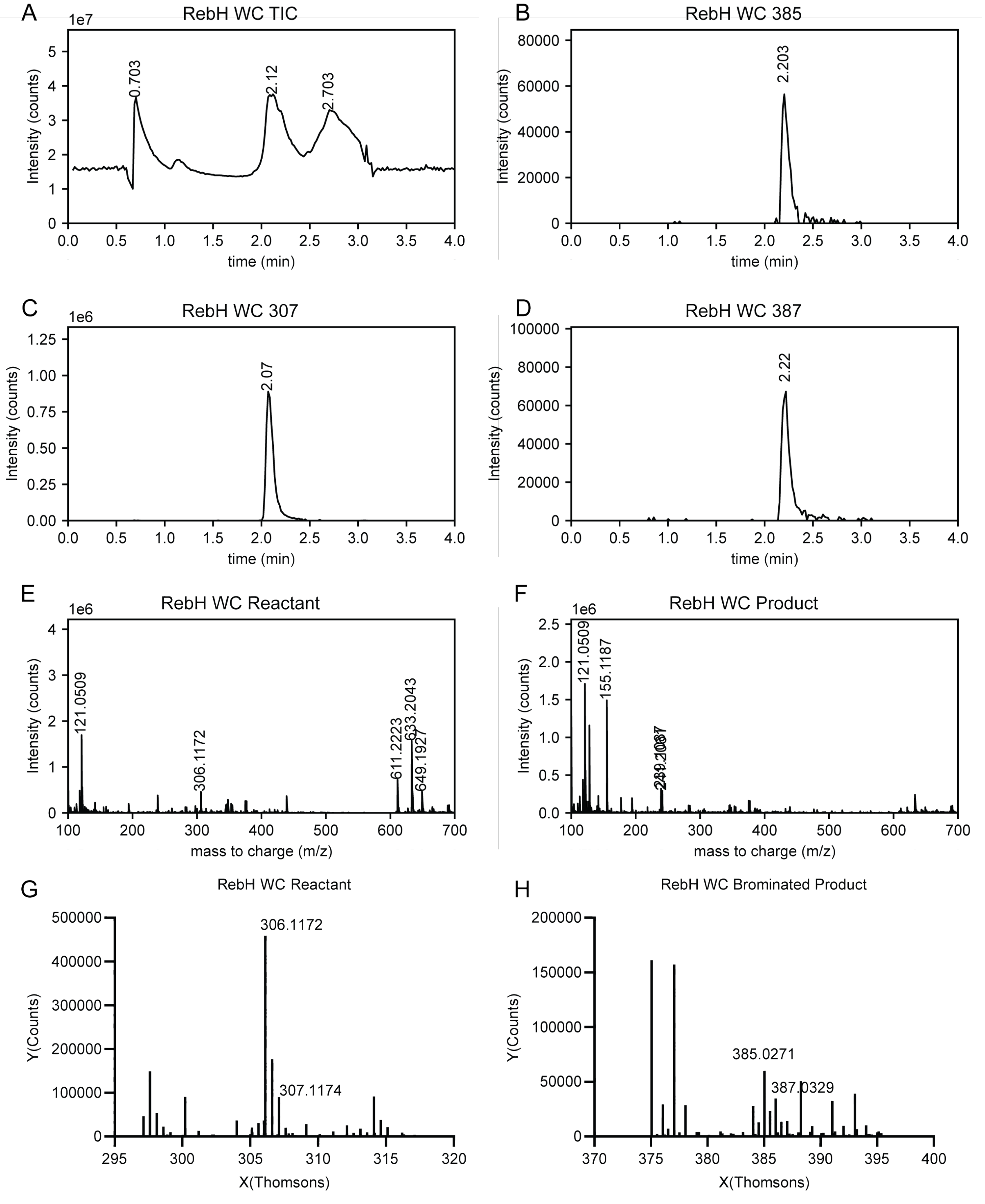
A
RebH WC TIC
1e7
Intensity (counts)
time (min)
0.703
2.12
2.703
B
RebH WC 385
Intensity (counts)
time (min)
2.203
C
RebH WC 307
1e6
Intensity (counts)
time (min)
2.07
D
RebH WC 387
Intensity (counts)
time (min)
2.22
E
RebH WC Reactant
1e6
Intensity (counts)
mass to charge (m/z)
121.0509
306.1172
611.2223
633.2043
649.1927
F
RebH WC Product
1e6
Intensity (counts)
mass to charge (m/z)
121.0509
155.1187
G
RebH WC Reactant
Y(Counts)
X(Thomsons)
306.1172
307.1174
H
RebH WC Brominated Product
Y(Counts)
X(Thomsons)
385.0271
387.0329

#### RebH WD

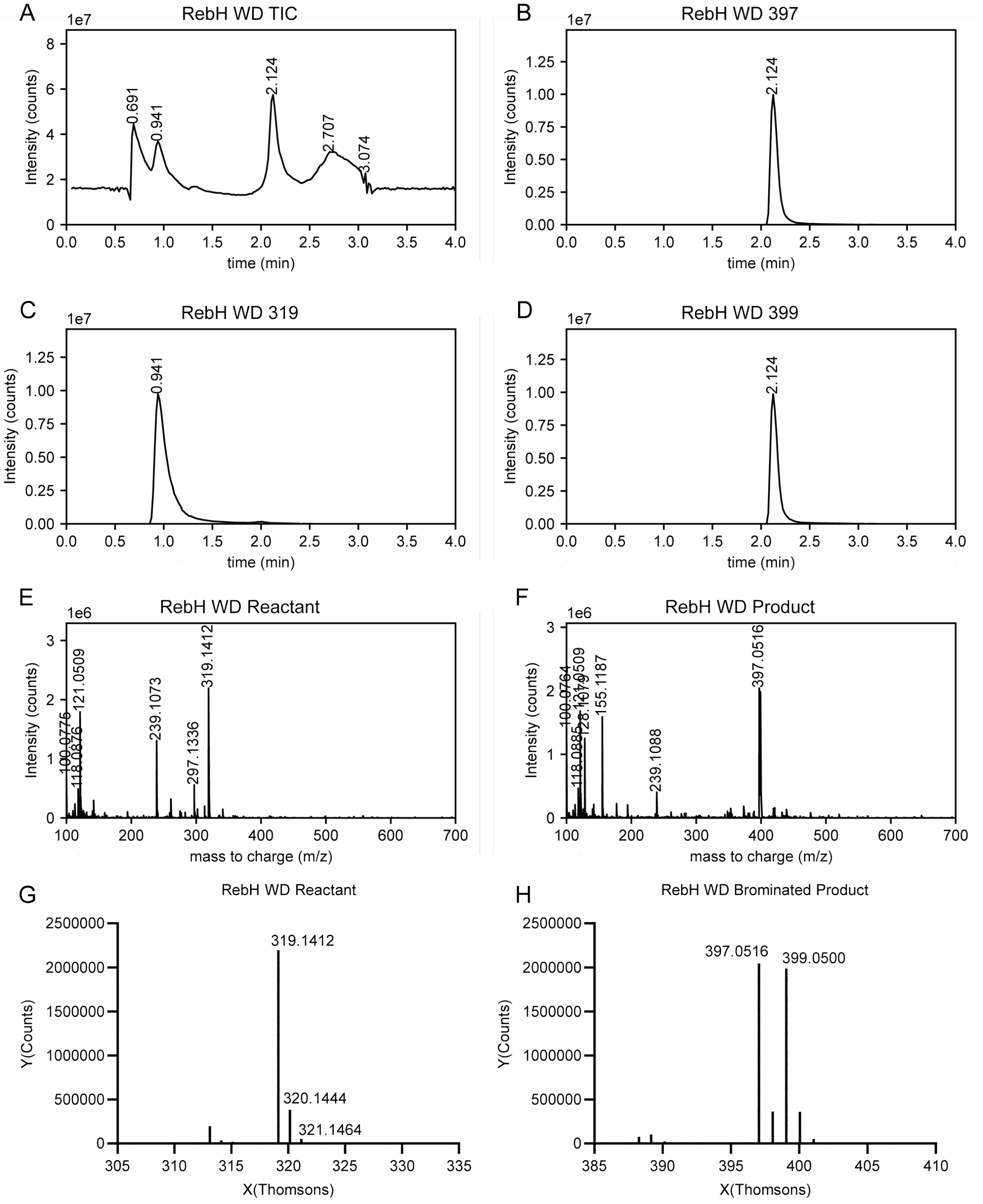

A
RebH WD TIC
1e7
Intensity (counts)
time (min)
0.691
0.941
2.124
2.707
3.074
B
RebH WD 397
1e7
Intensity (counts)
time (min)
2.124
C
RebH WD 319
1e7
Intensity (counts)
time (min)
0.941
D
RebH WD 399
1e7
Intensity (counts)
time (min)
2.124
E
RebH WD Reactant
1e6
Intensity (counts)
mass to charge (m/z)
100.0775
118.0876
121.0509
239.1073
297.1336
319.1412
F
RebH WD Product
1e6
Intensity (counts)
mass to charge (m/z)
100.0764
118.0885
121.0509
155.1187
239.1088
397.0516
G
RebH WD Reactant
Y(Counts)
X(Thomsons)
319.1412
320.1444
321.1464
H
RebH WD Brominated Product
Y(Counts)
X(Thomsons)
397.0516
399.0500

#### RebH WE

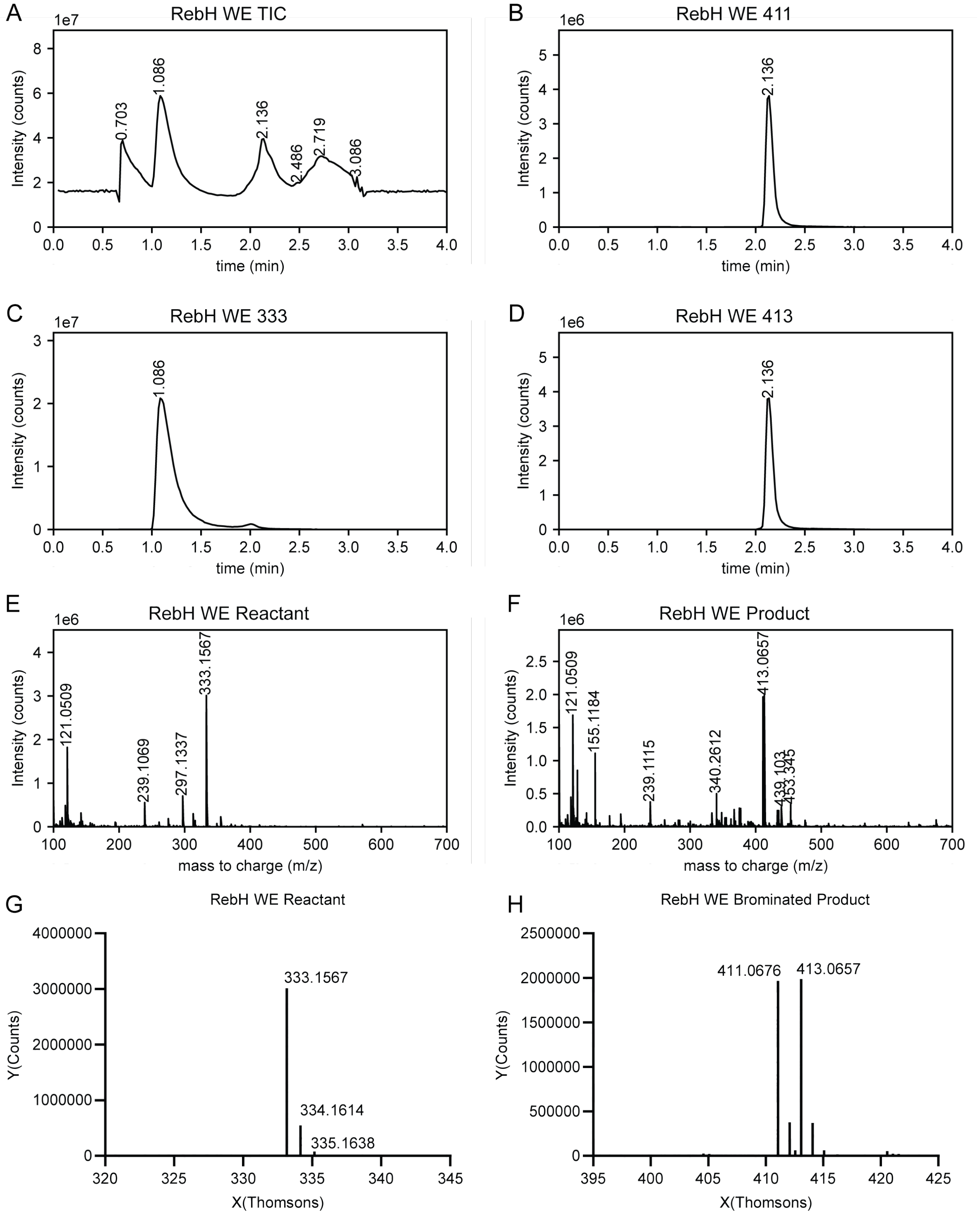
A
RebH WE TIC
1e7
Intensity (counts)
time (min)
0.703
1.086
2.136
2.486
2.719
3.086
B
RebH WE 411
1e6
Intensity (counts)
time (min)
2.136
C
RebH WE 333
1e7
Intensity (counts)
time (min)
1.086
D
RebH WE 413
1e6
Intensity (counts)
time (min)
2.136
E
RebH WE Reactant
1e6
Intensity (counts)
mass to charge (m/z)
121.0509
239.1069
297.1337
333.1567
F
RebH WE Product
1e6
Intensity (counts)
mass to charge (m/z)
121.0509
155.1184
239.1115
340.2612
413.0657
439.103
453.345
G
RebH WE Reactant
Y(Counts)
X(Thomsons)
333.1567
334.1614
335.1638
H
RebH WE Brominated Product
Y(Counts)
X(Thomsons)
411.0676
413.0657

#### RebH WF

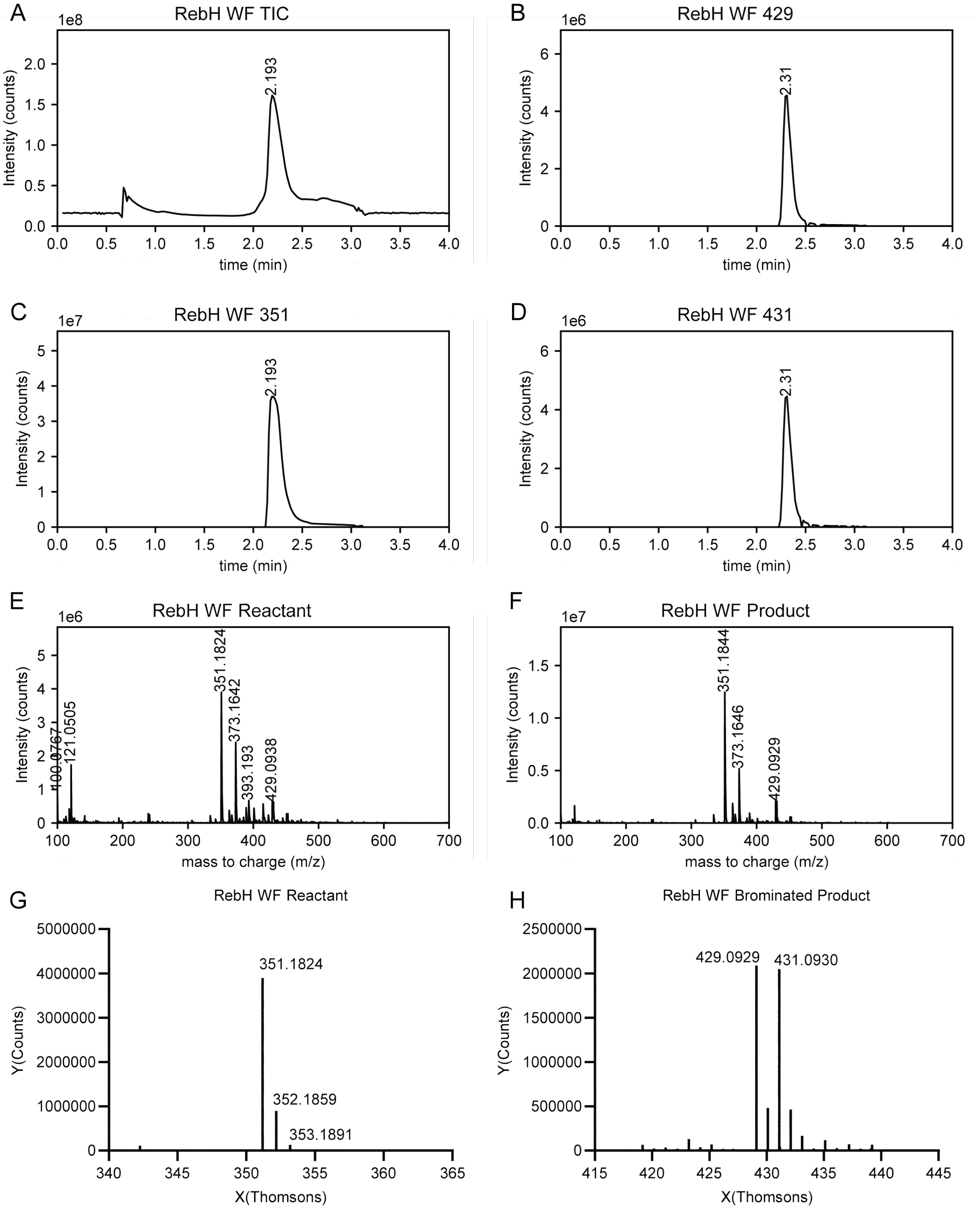

A
RebH WF TIC
1e8
Intensity (counts)
2.193
time (min)
B
RebH WF 429
1e6
2.31
C
RebH WF 351
1e7
2.193
D
RebH WF 431
1e6
2.31
E
RebH WF Reactant
1e6
351.1824
373.1642
121.0505
393.193
429.0938
mass to charge (m/z)
F
RebH WF Product
1e7
351.1844
373.1646
429.0929
G
RebH WF Reactant
Y(Counts)
351.1824
352.1859
353.1891
X(Thomsons)
H
RebH WF Brominated Product
429.0929
431.0930

#### RebH WG

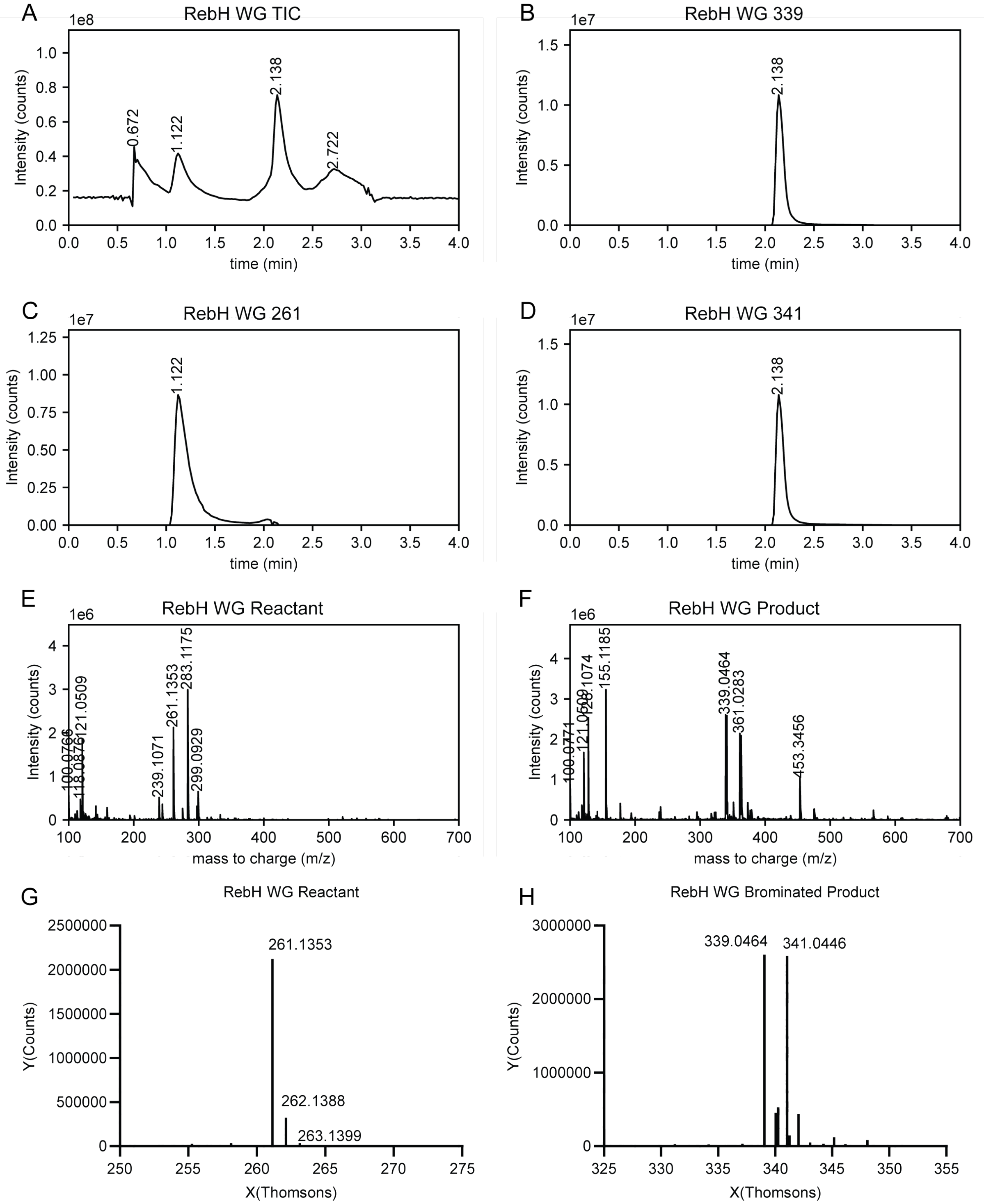
A
RebH WG TIC
1e8
Intensity (counts)
time (min)
0.672
1.122
2.138
2.722
B
RebH WG 339
1e7
Intensity (counts)
time (min)
2.138
C
RebH WG 261
1e7
Intensity (counts)
time (min)
1.122
D
RebH WG 341
1e7
Intensity (counts)
time (min)
2.138
E
RebH WG Reactant
1e6
Intensity (counts)
mass to charge (m/z)
121.0509
239.1071
261.1353
283.1175
299.0929
F
RebH WG Product
1e6
Intensity (counts)
mass to charge (m/z)
155.1185
339.0464
361.0283
453.3456
G
RebH WG Reactant
Y(Counts)
X(Thomsons)
261.1353
262.1388
263.1399
H
RebH WG Brominated Product
Y(Counts)
X(Thomsons)
339.0464
341.0446

**RebH WH**

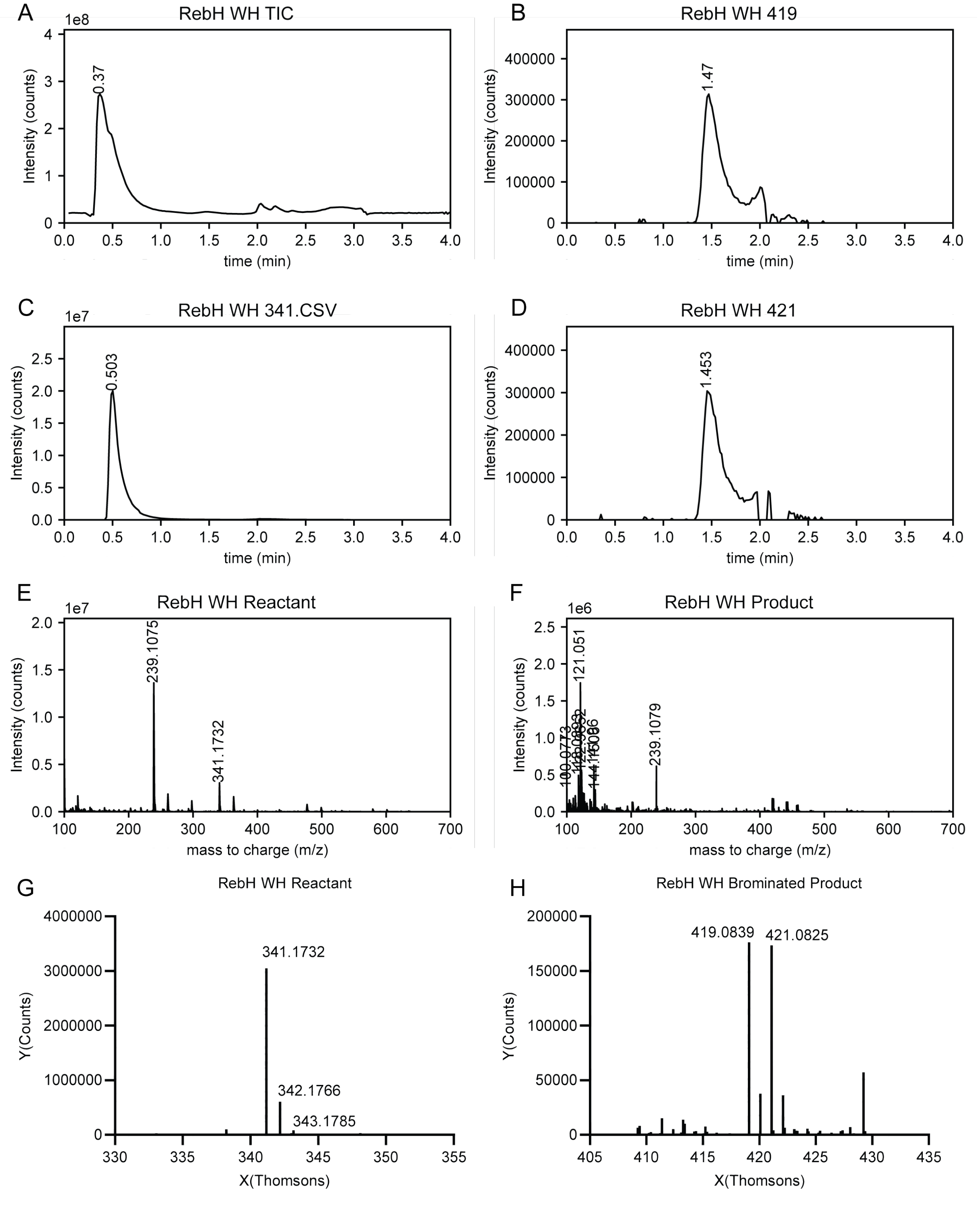

A
RebH WH TIC
1e8
0.37
Intensity (counts)
time (min)
B
RebH WH 419
1.47
Intensity (counts)
time (min)
C
RebH WH 341.CSV
1e7
0.503
Intensity (counts)
time (min)
D
RebH WH 421
1.453
Intensity (counts)
time (min)
E
RebH WH Reactant
1e7
239.1075
341.1732
Intensity (counts)
mass to charge (m/z)
F
RebH WH Product
1e6
121.051
239.1079
Intensity (counts)
mass to charge (m/z)
G
RebH WH Reactant
341.1732
342.1766
343.1785
Y(Counts)
X(Thomsons)
H
RebH WH Brominated Product
419.0839
421.0825
Y(Counts)
X(Thomsons)

#### RebH WI

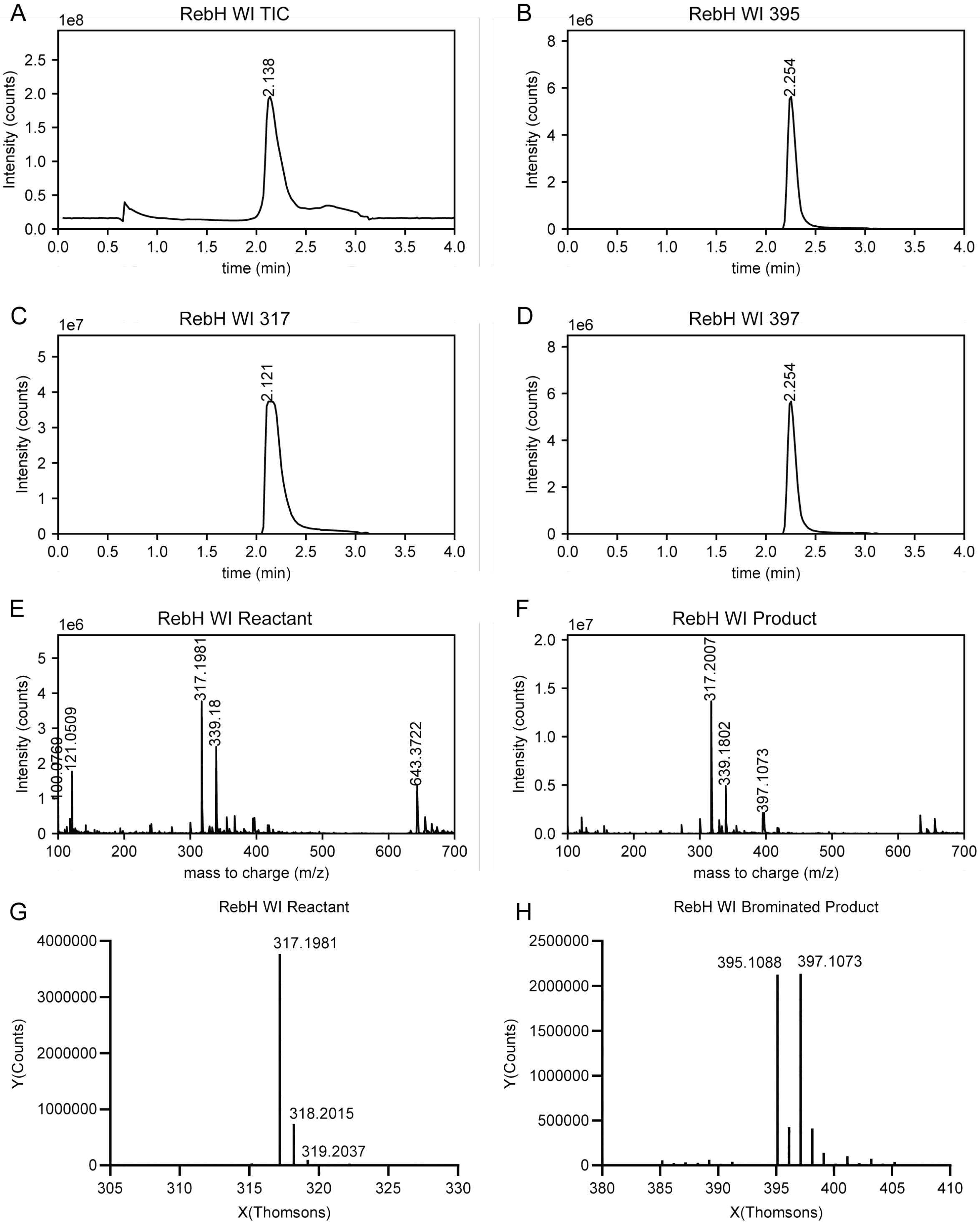
A
RebH WI TIC
1e8
Intensity (counts)
time (min)
2.138
B
RebH WI 395
1e6
time (min)
2.254
C
RebH WI 317
1e7
Intensity (counts)
time (min)
2.121
D
RebH WI 397
1e6
Intensity (counts)
time (min)
2.254
E
RebH WI Reactant
1e6
Intensity (counts)
mass to charge (m/z)
100.0769
121.0509
317.1981
339.18
643.3722
F
RebH WI Product
1e7
Intensity (counts)
mass to charge (m/z)
317.2007
339.1802
397.1073
G
RebH WI Reactant
Y(Counts)
X(Thomsons)
317.1981
318.2015
319.2037
H
RebH WI Brominated Product
Y(Counts)
X(Thomsons)
395.1088
397.1073

#### RebH WK

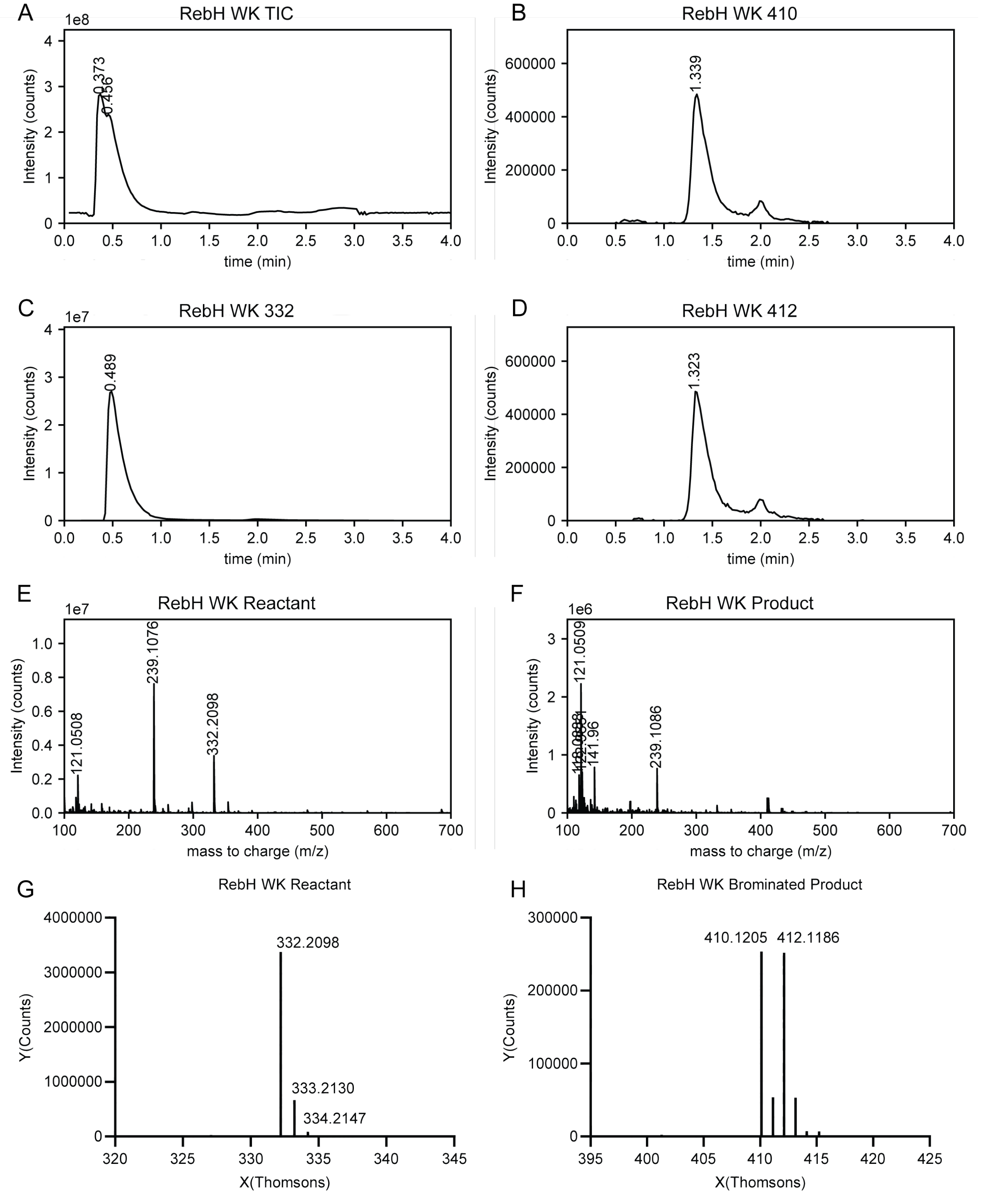
A
RebH WK TIC
1e8
0.373
0.456
Intensity (counts)
time (min)
B
RebH WK 410
1.339
Intensity (counts)
time (min)
C
RebH WK 332
1e7
0.489
Intensity (counts)
time (min)
D
RebH WK 412
1.323
Intensity (counts)
time (min)
E
RebH WK Reactant
1e7
239.1076
332.2098
121.0508
Intensity (counts)
mass to charge (m/z)
F
RebH WK Product
1e6
121.0509
141.96
239.1086
Intensity (counts)
mass to charge (m/z)
G
RebH WK Reactant
332.2098
333.2130
334.2147
Y(Counts)
X(Thomsons)
H
RebH WK Brominated Product
410.1205
412.1186
Y(Counts)
X(Thomsons)

#### RebH WL

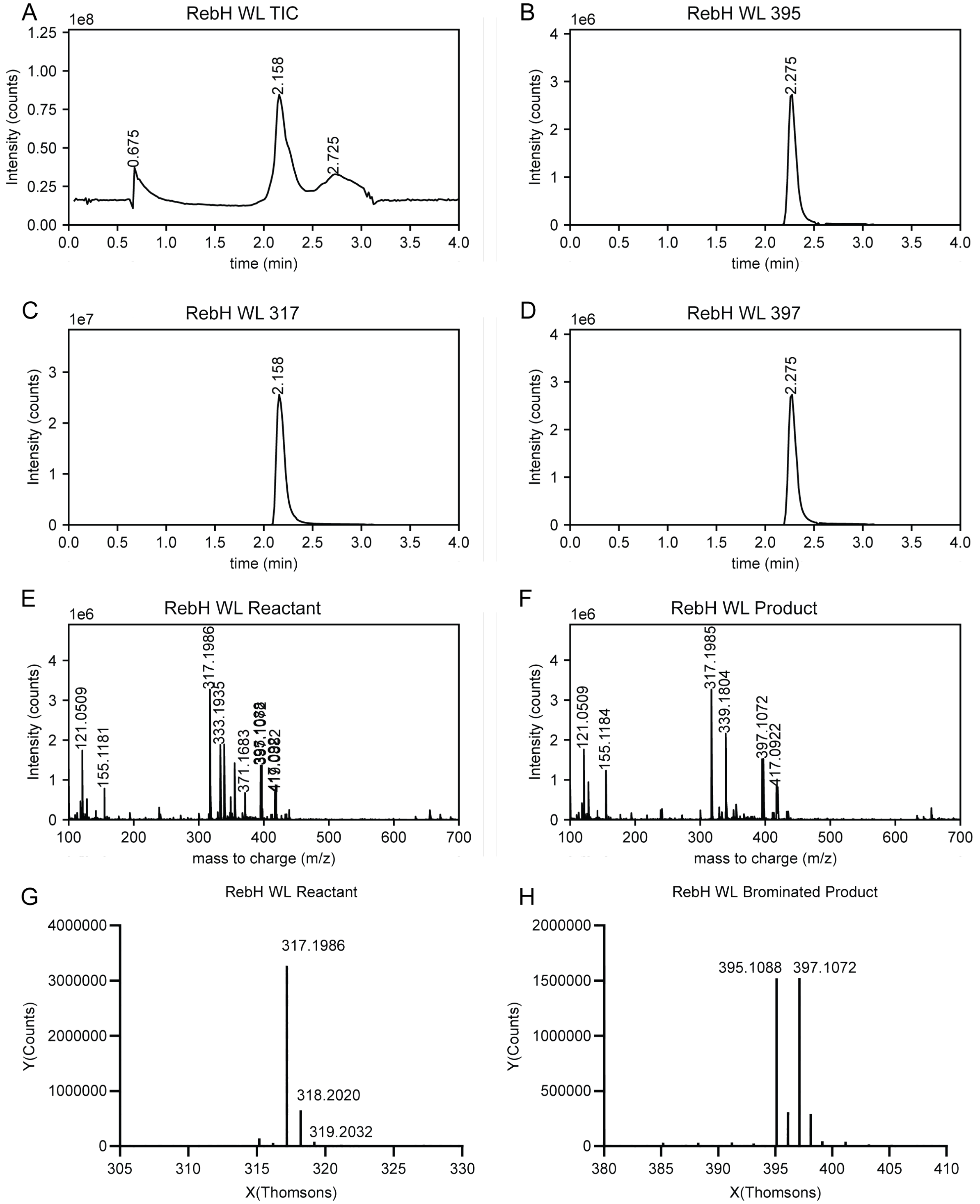
A
RebH WL TIC
1e8
Intensity (counts)
time (min)
0.675
2.158
2.725
B
RebH WL 395
1e6
Intensity (counts)
time (min)
2.275
C
RebH WL 317
1e7
Intensity (counts)
time (min)
2.158
D
RebH WL 397
1e6
Intensity (counts)
time (min)
2.275
E
RebH WL Reactant
1e6
Intensity (counts)
mass to charge (m/z)
121.0509
155.1181
317.1986
333.1935
371.1683
F
RebH WL Product
1e6
Intensity (counts)
mass to charge (m/z)
121.0509
155.1184
317.1985
339.1804
397.1072
417.0922
G
RebH WL Reactant
Y(Counts)
X(Thomsons)
317.1986
318.2020
319.2032
H
RebH WL Brominated Product
Y(Counts)
X(Thomsons)
395.1088
397.1072

#### RebH WM

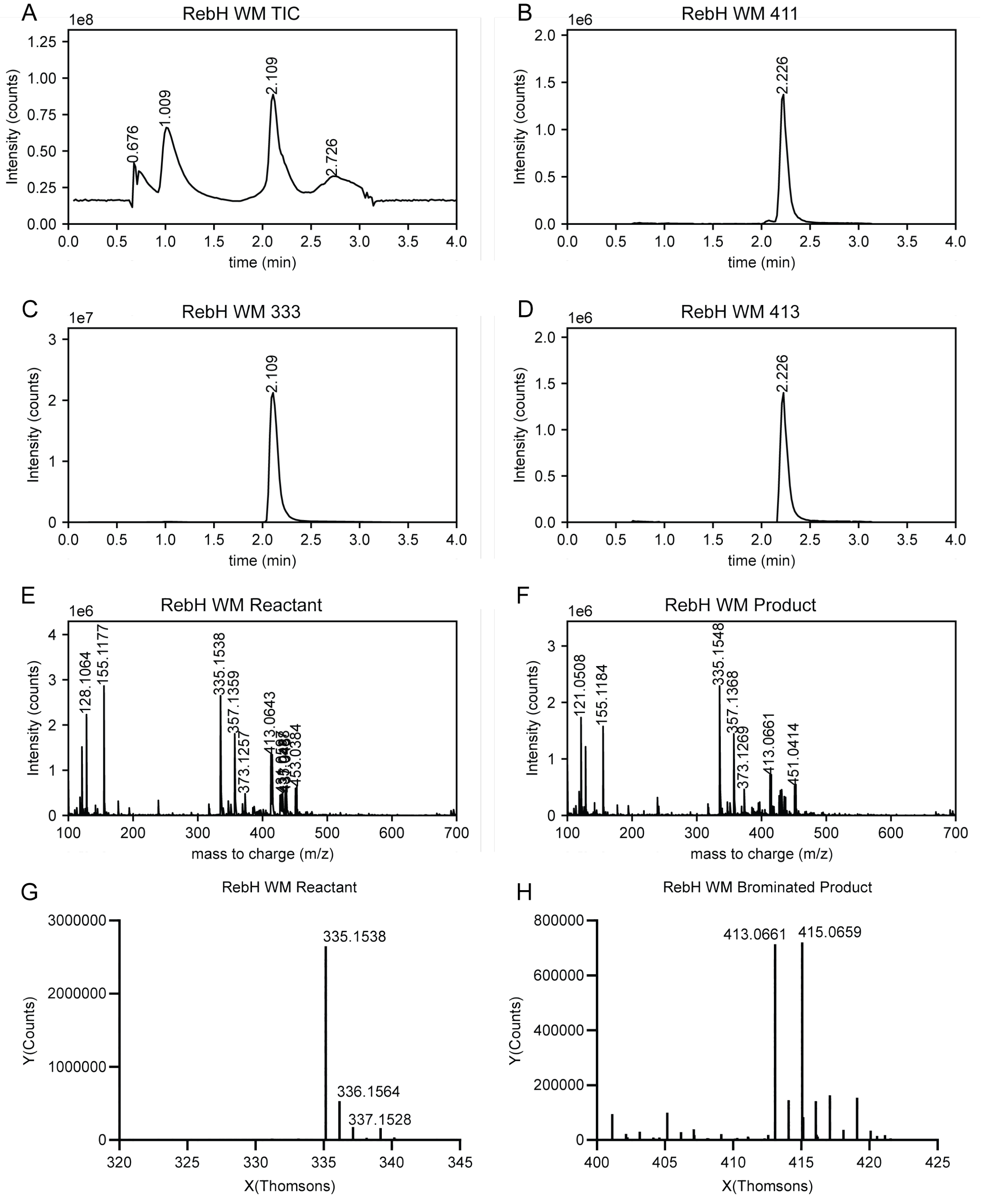
A
RebH WM TIC
1e8
Intensity (counts)
time (min)
0.676
1.009
2.109
2.726
B
RebH WM 411
1e6
Intensity (counts)
time (min)
2.226
C
RebH WM 333
1e7
Intensity (counts)
time (min)
2.109
D
RebH WM 413
1e6
Intensity (counts)
time (min)
2.226
E
RebH WM Reactant
1e6
Intensity (counts)
mass to charge (m/z)
128.1064
155.1177
335.1538
357.1359
373.1257
413.0643
453.0384
F
RebH WM Product
1e6
Intensity (counts)
mass to charge (m/z)
121.0508
155.1184
335.1548
357.1368
373.1269
413.0661
451.0414
G
RebH WM Reactant
Y(Counts)
X(Thomsons)
335.1538
336.1564
337.1528
H
RebH WM Brominated Product
Y(Counts)
X(Thomsons)
413.0661
415.0659

#### RebH WN

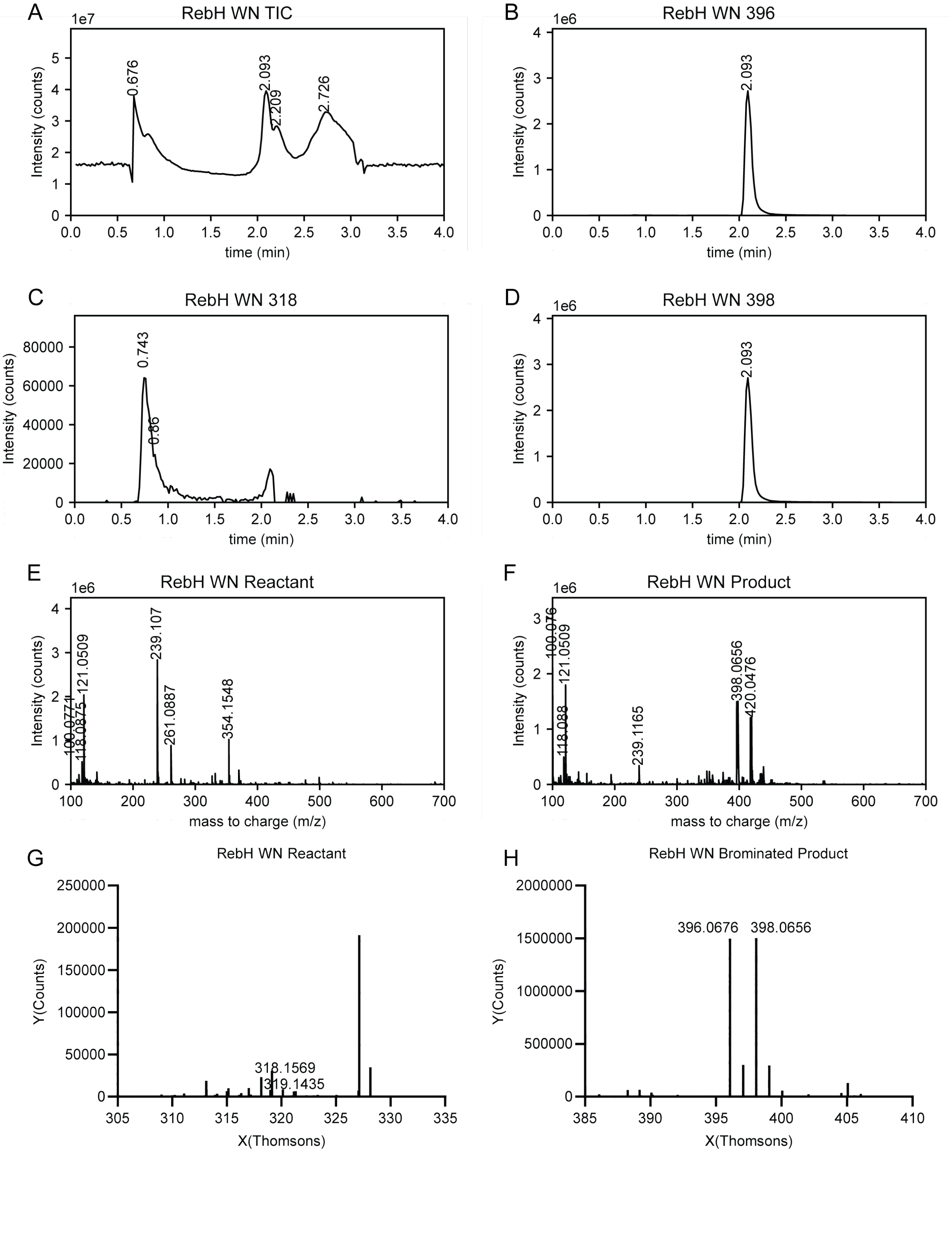

A
RebH WN TIC
1e7
0.676
2.093
2.209
2.726
Intensity (counts)
time (min)
B
RebH WN 396
1e6
2.093
Intensity (counts)
time (min)
C
RebH WN 318
0.743
0.86
Intensity (counts)
time (min)
D
RebH WN 398
1e6
2.093
Intensity (counts)
time (min)
E
RebH WN Reactant
1e6
239.107
121.0509
100.0771
118.0875
261.0887
354.1548
Intensity (counts)
mass to charge (m/z)
F
RebH WN Product
1e6
100.076
121.0509
118.088
239.1165
398.0656
420.0476
Intensity (counts)
mass to charge (m/z)
G
RebH WN Reactant
318.1569
319.1435
Y(Counts)
X(Thomsons)
H
RebH WN Brominated Product
396.0676
398.0656
Y(Counts)
X(Thomsons)

**RebH WP**

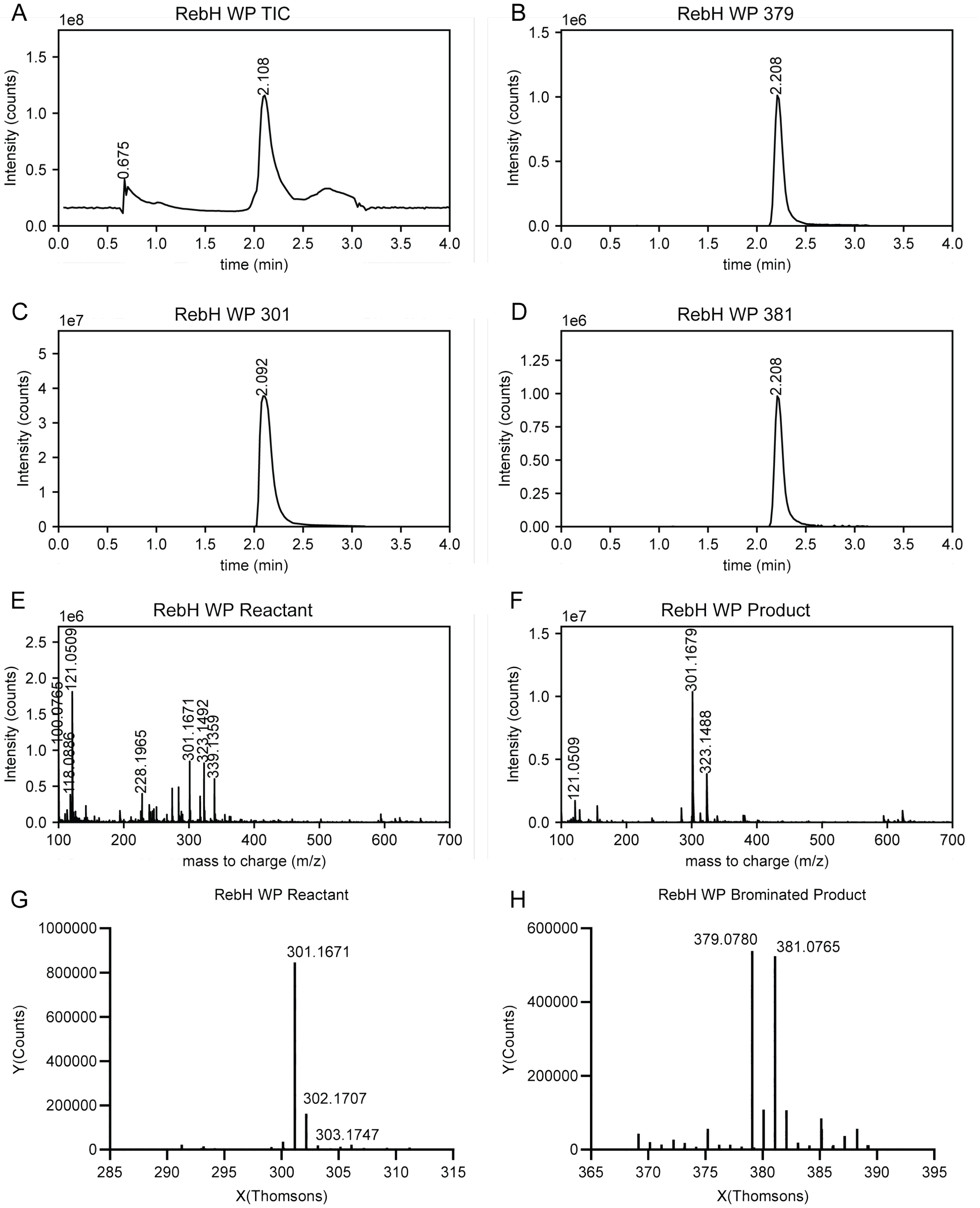

A
RebH WP TIC
1e8
Intensity (counts)
time (min)
0.675
2.108
B
RebH WP 379
1e6
Intensity (counts)
time (min)
2.208
C
RebH WP 301
1e7
Intensity (counts)
time (min)
2.092
D
RebH WP 381
1e6
Intensity (counts)
time (min)
2.208
E
RebH WP Reactant
1e6
Intensity (counts)
mass to charge (m/z)
121.0509
100.0765
118.0886
228.1965
301.1671
323.1492
339.1359
F
RebH WP Product
1e7
Intensity (counts)
mass to charge (m/z)
301.1679
323.1488
121.0509
G
RebH WP Reactant
Y(Counts)
X(Thomsons)
301.1671
302.1707
303.1747
H
RebH WP Brominated Product
Y(Counts)
X(Thomsons)
379.0780
381.0765

#### RebH WQ

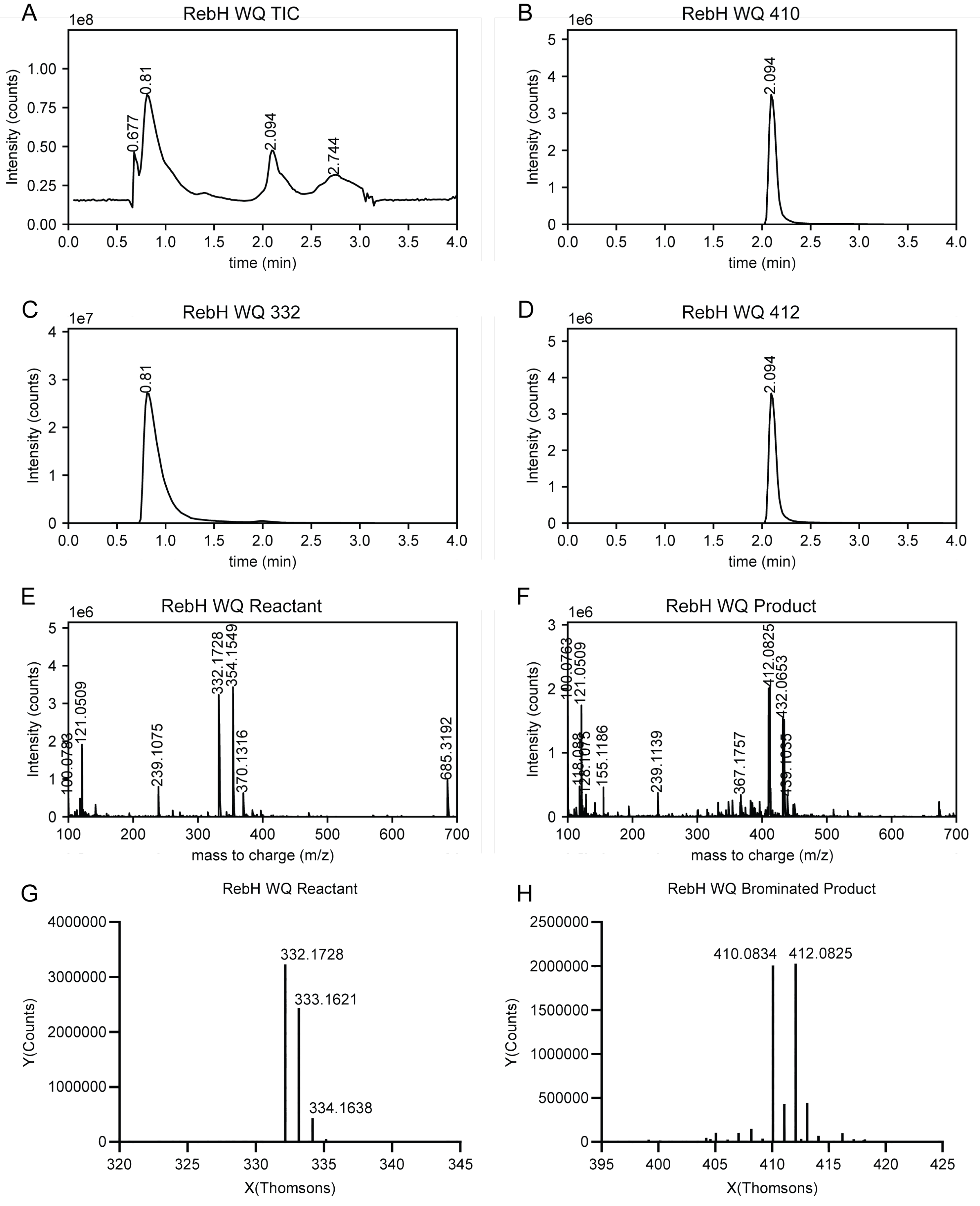
A
RebH WQ TIC
1e8
Intensity (counts)
time (min)
0.677
0.81
2.094
2.744
B
RebH WQ 410
1e6
2.094
C
RebH WQ 332
1e7
0.81
D
RebH WQ 412
1e6
2.094
E
RebH WQ Reactant
121.0509
239.1075
332.1728
354.1549
370.1316
685.3192
mass to charge (m/z)
F
RebH WQ Product
121.0509
155.1186
239.1139
367.1757
412.0825
432.0653
G
RebH WQ Reactant
332.1728
333.1621
334.1638
Y(Counts)
X(Thomsons)
H
RebH WQ Brominated Product
410.0834
412.0825

#### RebH WR

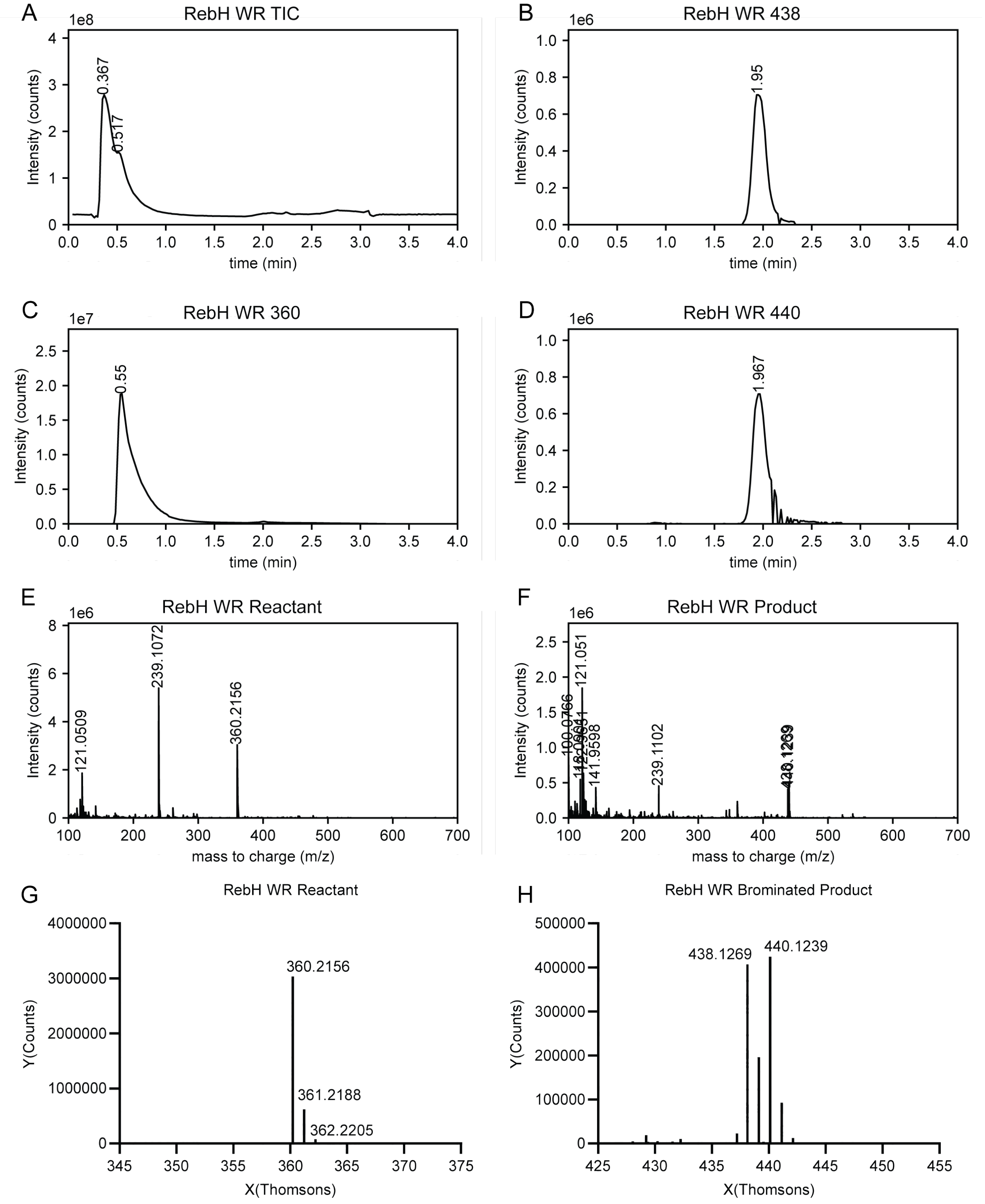
A
RebH WR TIC
1e8
Intensity (counts)
time (min)
0.367
0.517
B
RebH WR 438
1e6
Intensity (counts)
time (min)
1.95
C
RebH WR 360
1e7
Intensity (counts)
time (min)
0.55
D
RebH WR 440
1e6
Intensity (counts)
time (min)
1.967
E
RebH WR Reactant
1e6
Intensity (counts)
mass to charge (m/z)
121.0509
239.1072
360.2156
F
RebH WR Product
1e6
Intensity (counts)
mass to charge (m/z)
121.051
141.9598
239.1102
G
RebH WR Reactant
Y(Counts)
X(Thomsons)
360.2156
361.2188
362.2205
H
RebH WR Brominated Product
Y(Counts)
X(Thomsons)
438.1269
440.1239

#### RebH WS

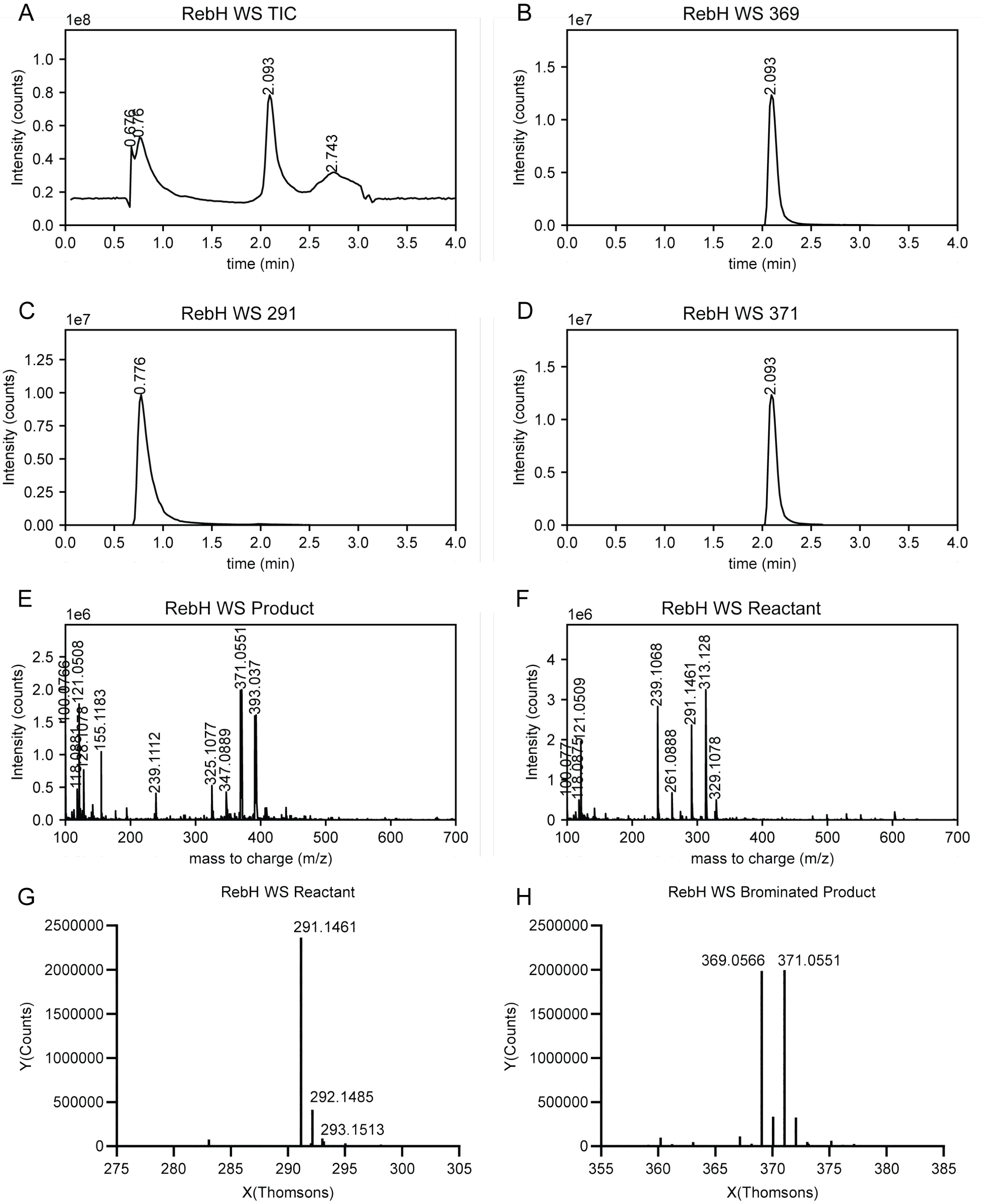

A
RebH WS TIC
1e8
Intensity (counts)
time (min)
0.676
0.76
2.093
2.743
B
RebH WS 369
1e7
2.093
C
RebH WS 291
1e7
0.776
D
RebH WS 371
1e7
2.093
E
RebH WS Product
1e6
mass to charge (m/z)
121.0508
155.1183
239.1112
325.1077
347.0889
371.0551
393.037
F
RebH WS Reactant
1e6
121.0509
239.1068
261.0888
291.1461
313.128
329.1078
G
RebH WS Reactant
Y(Counts)
X(Thomsons)
291.1461
292.1485
293.1513
H
RebH WS Brominated Product
369.0566
371.0551

**RebH WT**

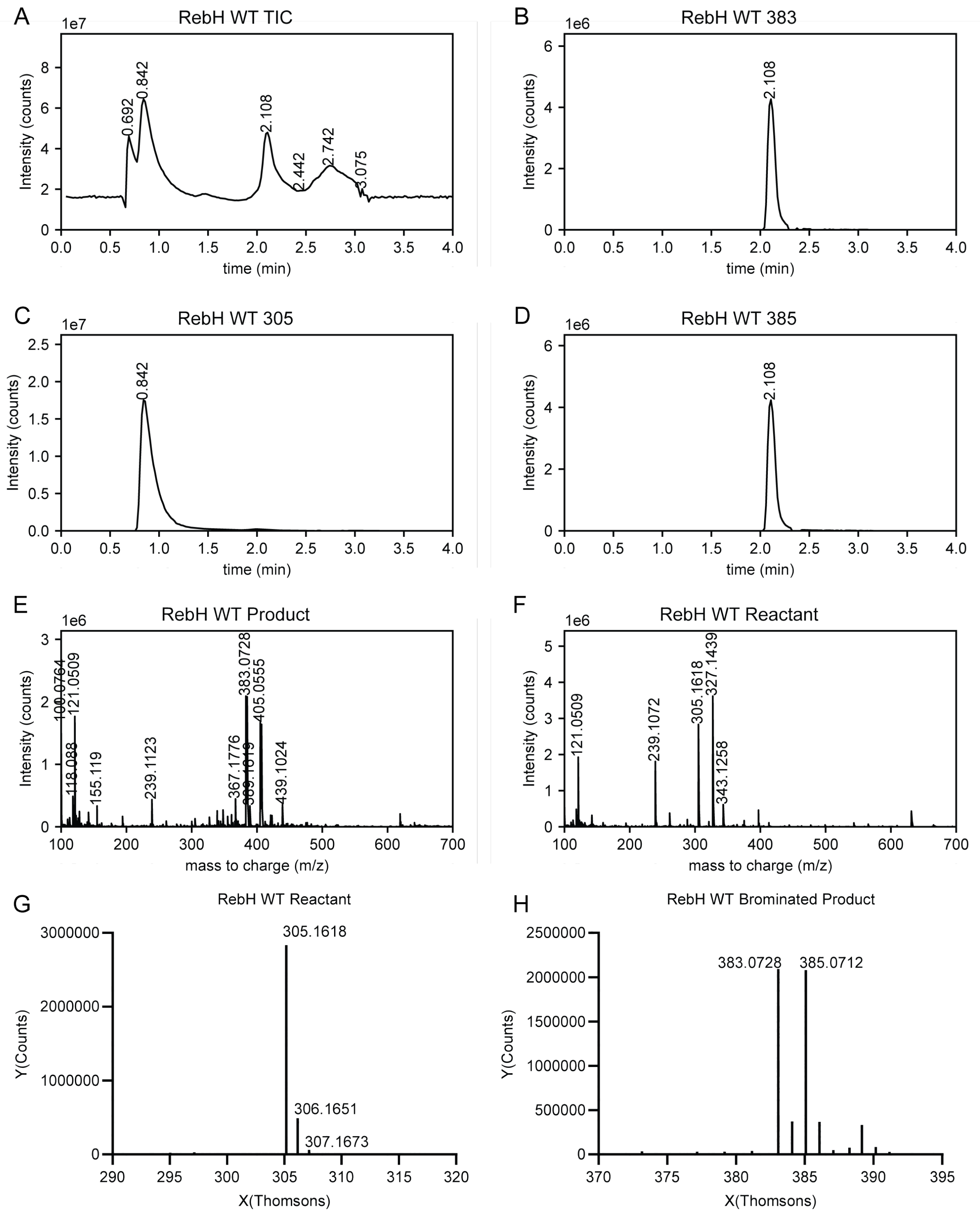

A
RebH WT TIC
1e7
Intensity (counts)
time (min)
0.692
0.842
2.108
2.442
2.742
3.075
B
RebH WT 383
1e6
2.108
C
RebH WT 305
0.842
D
RebH WT 385
2.108
E
RebH WT Product
mass to charge (m/z)
121.0509
118.088
155.119
239.1123
367.1776
383.0728
405.0555
439.1024
F
RebH WT Reactant
121.0509
239.1072
305.1618
327.1439
343.1258
G
RebH WT Reactant
Y(Counts)
X(Thomsons)
305.1618
306.1651
307.1673
H
RebH WT Brominated Product
383.0728
385.0712

**RebH WV**

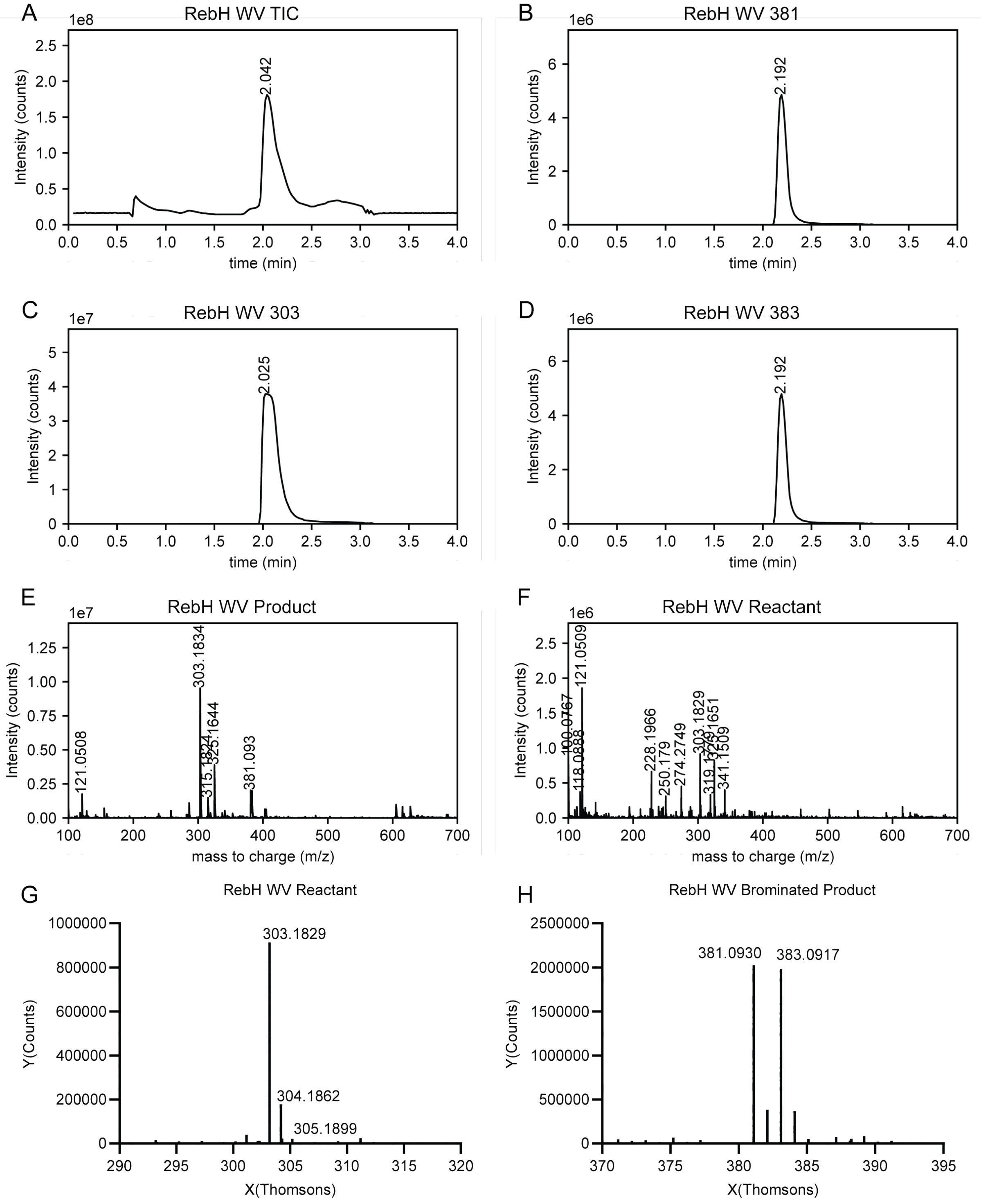

A
RebH WV TIC
B
RebH WV 381
C
RebH WV 303
D
RebH WV 383
E
RebH WV Product
F
RebH WV Reactant
G
RebH WV Reactant
H
RebH WV Brominated Product
Intensity (counts)
time (min)
mass to charge (m/z)
Y(Counts)
X(Thomsons)
2.042
2.192
2.025
2.192
121.0508
303.1834
315.1824
325.1644
381.093
100.0767
118.0888
121.0509
228.1966
250.179
274.2749
303.1829
319.1749
325.1651
341.1509
303.1829
304.1862
305.1899
381.0930
383.0917

#### RebH WY

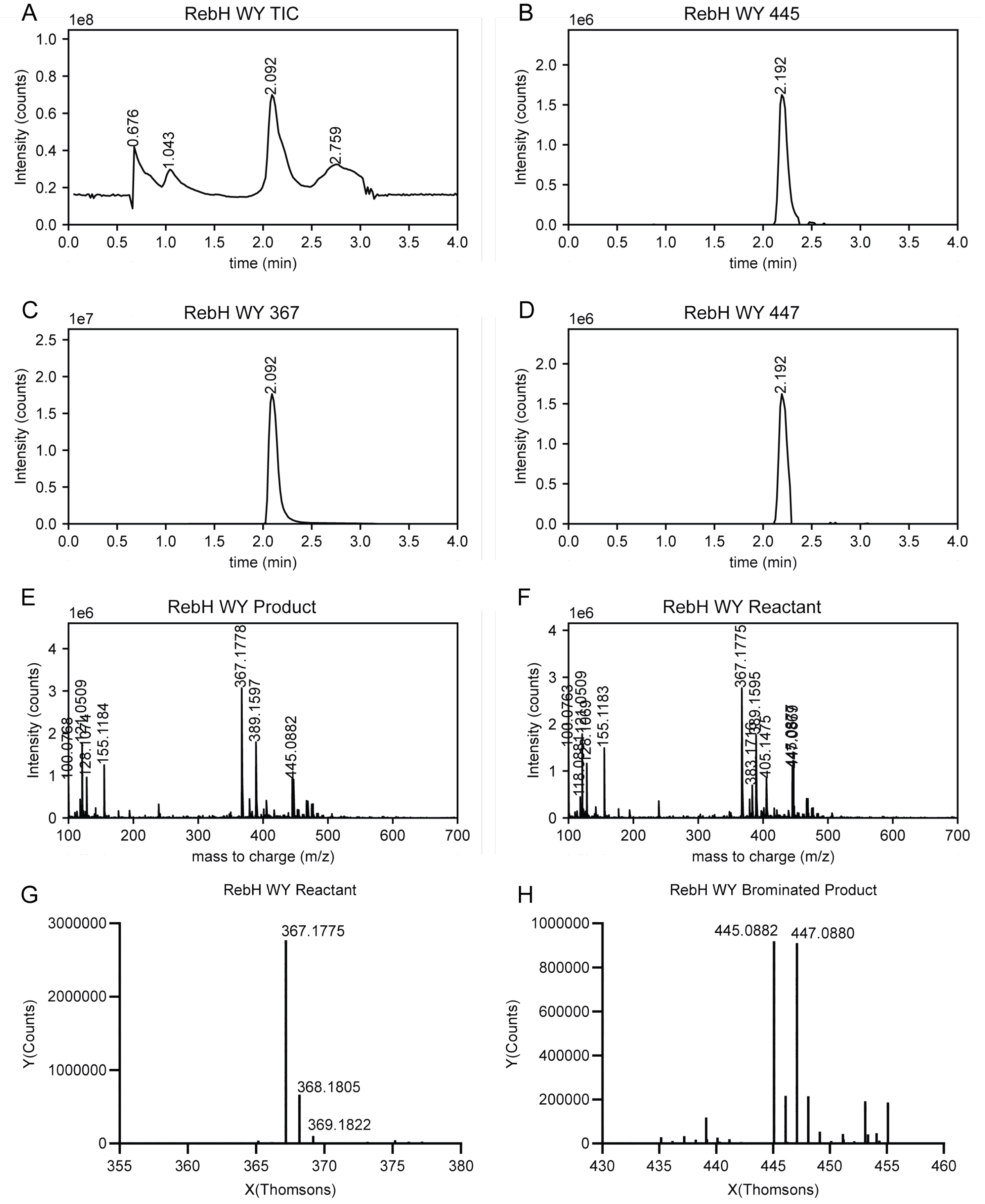
A
RebH WY TIC
1e8
Intensity (counts)
time (min)
0.676
1.043
2.092
2.759
B
RebH WY 445
1e6
Intensity (counts)
time (min)
2.192
C
RebH WY 367
1e7
Intensity (counts)
time (min)
2.092
D
RebH WY 447
1e6
Intensity (counts)
time (min)
2.192
E
RebH WY Product
1e6
Intensity (counts)
mass to charge (m/z)
100.0768
128.1074
155.1184
367.1778
389.1597
445.0882
F
RebH WY Reactant
1e6
Intensity (counts)
mass to charge (m/z)
100.0763
155.1183
367.1775
405.1475
G
RebH WY Reactant
Y(Counts)
X(Thomsons)
367.1775
368.1805
369.1822
H
RebH WY Brominated Product
Y(Counts)
X(Thomsons)
445.0882
447.0880

#### 4V Brominated Dipeptide Mass Spectra

## 4V AW

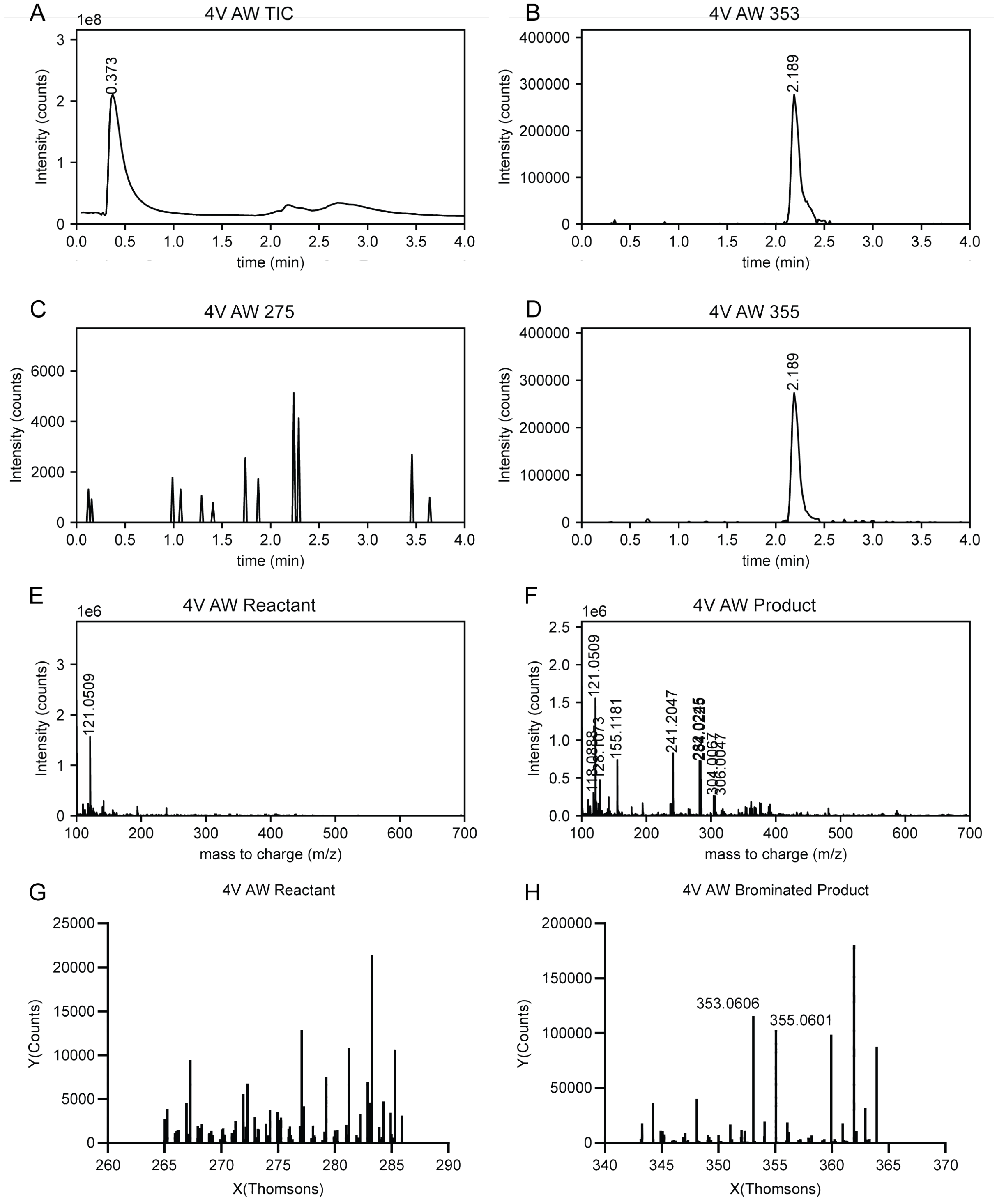

A
4V AW TIC
1e8
Intensity (counts)
0.373
time (min)
B
4V AW 353
Intensity (counts)
2.189
time (min)
C
4V AW 275
Intensity (counts)
time (min)
D
4V AW 355
Intensity (counts)
2.189
time (min)
E
4V AW Reactant
1e6
Intensity (counts)
121.0509
mass to charge (m/z)
F
4V AW Product
1e6
Intensity (counts)
121.0509
118.0888
128.1073
155.1181
241.2047
304.0067
306.0047
mass to charge (m/z)
G
4V AW Reactant
Y(Counts)
X(Thomsons)
H
4V AW Brominated Product
353.0606
355.0601
Y(Counts)
X(Thomsons)

## 4V CW

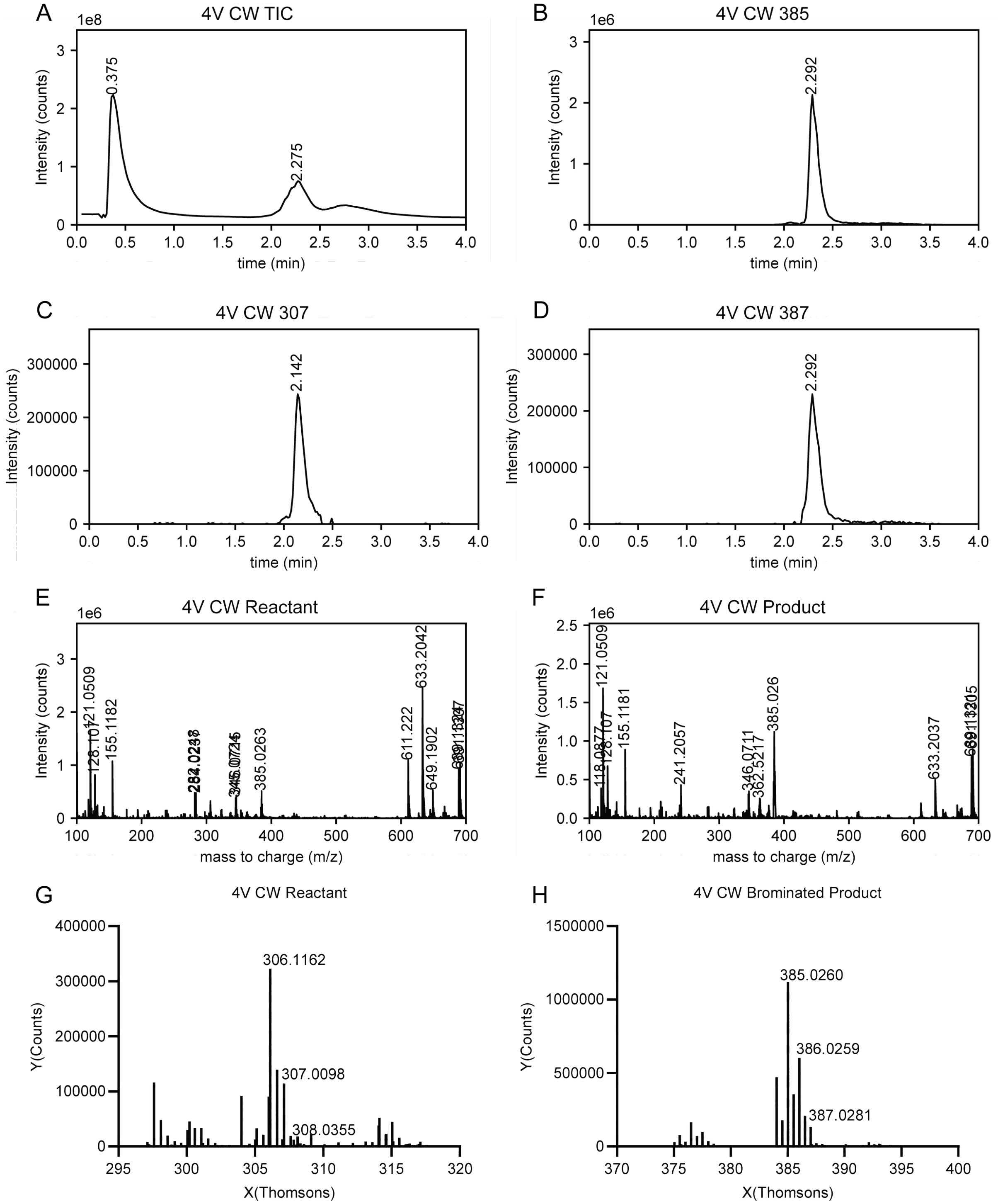

A
4V CW TIC
1e8
Intensity (counts)
time (min)
0.375
2.275
B
4V CW 385
1e6
Intensity (counts)
time (min)
2.292
C
4V CW 307
Intensity (counts)
time (min)
2.142
D
4V CW 387
Intensity (counts)
time (min)
2.292
E
4V CW Reactant
1e6
Intensity (counts)
mass to charge (m/z)
121.0509
128.107
155.1182
385.0263
611.222
633.2042
649.1902
F
4V CW Product
1e6
Intensity (counts)
mass to charge (m/z)
121.0509
118.0877
128.107
155.1181
241.2057
346.0711
362.5217
385.026
633.2037
G
4V CW Reactant
Y(Counts)
X(Thomsons)
306.1162
307.0098
308.0355
H
4V CW Brominated Product
Y(Counts)
X(Thomsons)
385.0260
386.0259
387.0281

**4V DW**

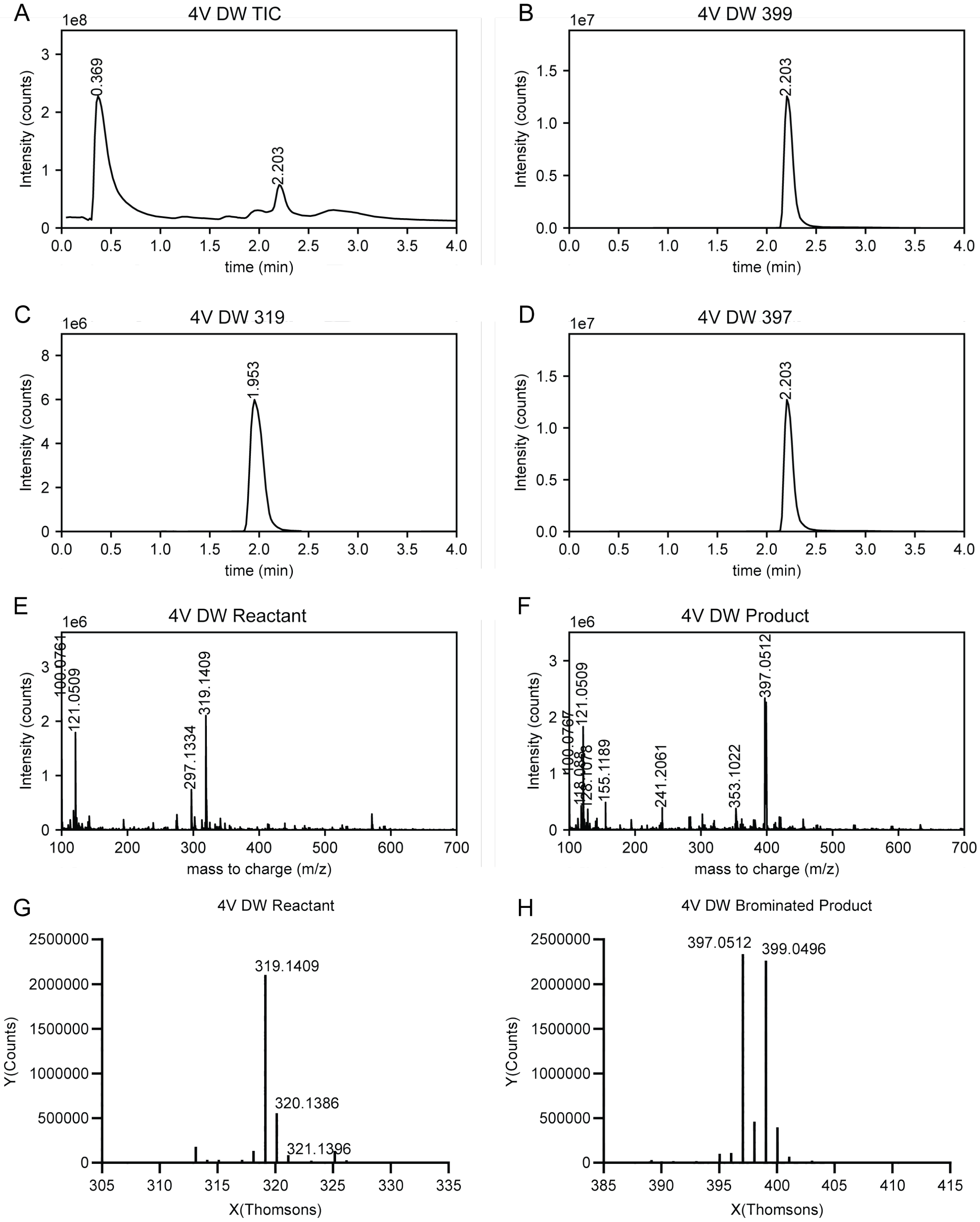

A
4V DW TIC
1e8
Intensity (counts)
time (min)
0.369
2.203
B
4V DW 399
1e7
Intensity (counts)
time (min)
2.203
C
4V DW 319
1e6
Intensity (counts)
time (min)
1.953
D
4V DW 397
1e7
Intensity (counts)
time (min)
2.203
E
4V DW Reactant
1e6
Intensity (counts)
mass to charge (m/z)
121.0509
297.1334
319.1409
F
4V DW Product
1e6
Intensity (counts)
mass to charge (m/z)
121.0509
155.1189
241.2061
353.1022
397.0512
G
4V DW Reactant
Y(Counts)
X(Thomsons)
319.1409
320.1386
321.1396
H
4V DW Brominated Product
Y(Counts)
X(Thomsons)
397.0512
399.0496

## 4V EW

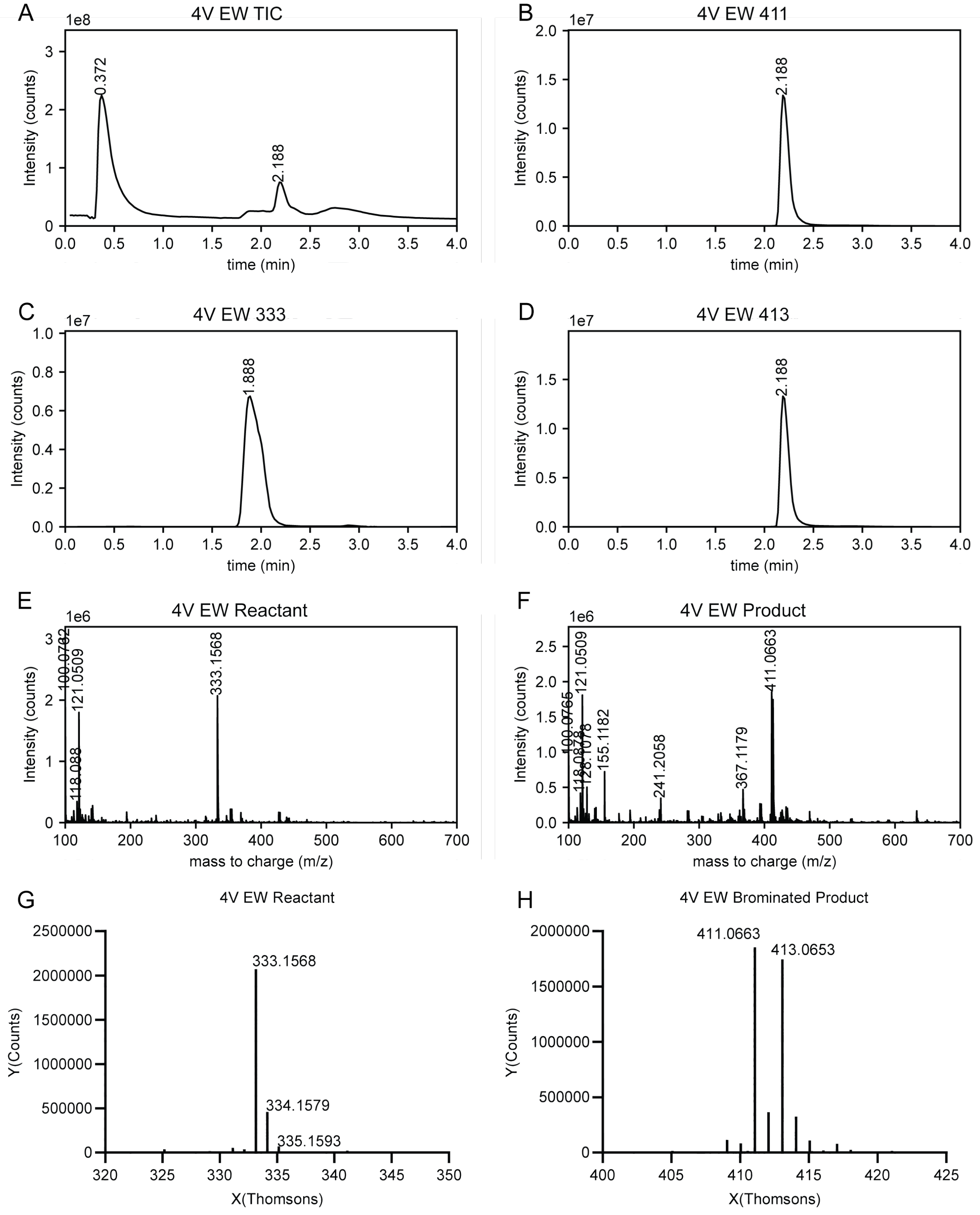

A
4V EW TIC
1e8
Intensity (counts)
time (min)
0.372
2.188
B
4V EW 411
1e7
Intensity (counts)
time (min)
2.188
C
4V EW 333
1e7
Intensity (counts)
time (min)
1.888
D
4V EW 413
1e7
Intensity (counts)
time (min)
2.188
E
4V EW Reactant
1e6
Intensity (counts)
mass to charge (m/z)
100.0782
121.0509
118.088
333.1568
F
4V EW Product
1e6
Intensity (counts)
mass to charge (m/z)
100.0765
118.0873
121.0509
126.1078
155.1182
241.2058
367.1179
411.0663
G
4V EW Reactant
Y(Counts)
X(Thomsons)
333.1568
334.1579
335.1593
H
4V EW Brominated Product
Y(Counts)
X(Thomsons)
411.0663
413.0653

## 4V FW

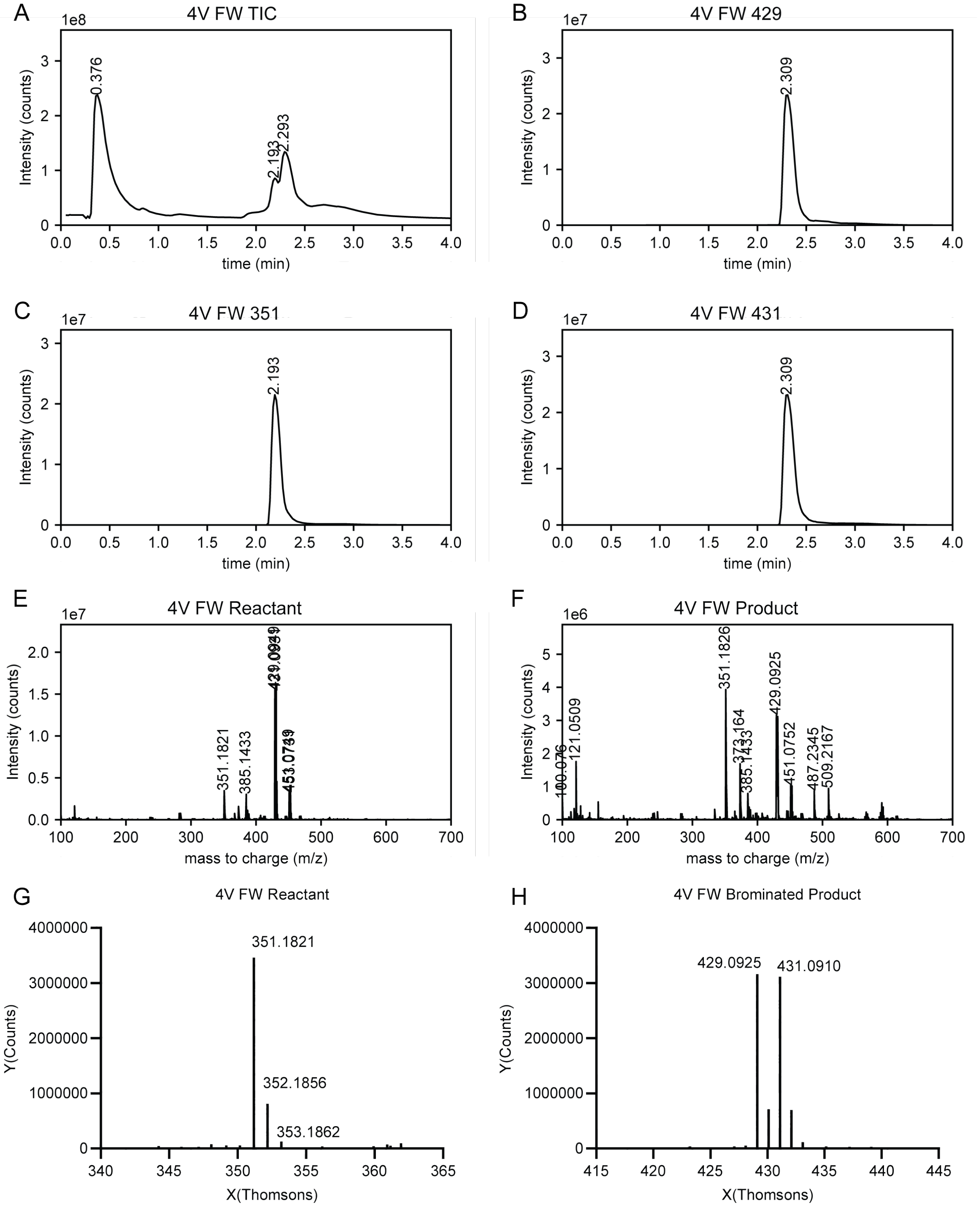
A
4V FW TIC
1e8
Intensity (counts)
time (min)
0.376
2.193
2.293
B
4V FW 429
1e7
2.309
C
4V FW 351
2.193
D
4V FW 431
2.309
E
4V FW Reactant
1e7
mass to charge (m/z)
351.1821
385.1433
F
4V FW Product
1e6
351.1826
429.0925
121.0509
373.164
385.1433
451.0752
487.2345
509.2167
G
4V FW Reactant
Y(Counts)
X(Thomsons)
351.1821
352.1856
353.1862
H
4V FW Brominated Product
429.0925
431.0910

## 4V GW

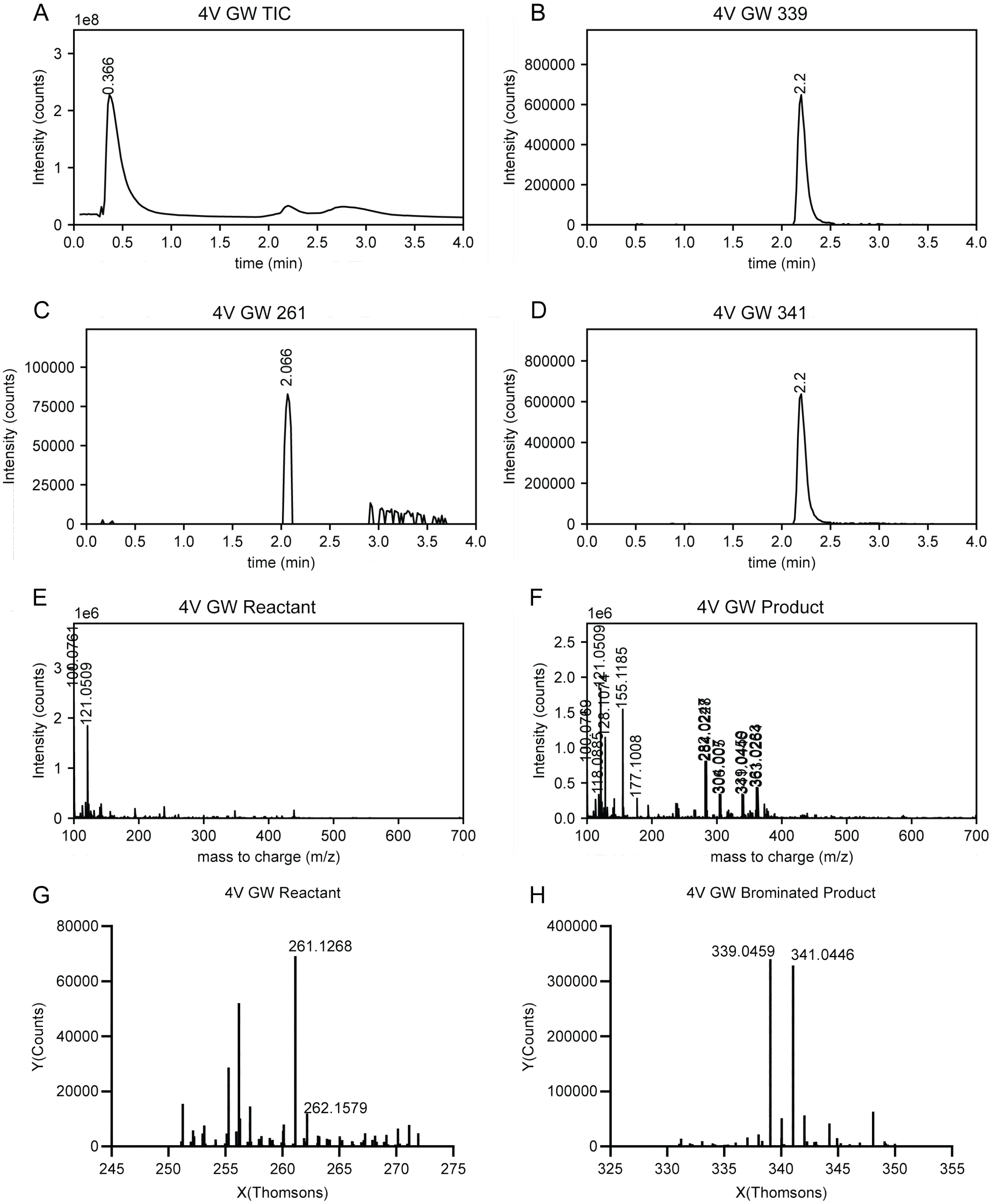

A
4V GW TIC
1e8
Intensity (counts)
0.366
time (min)
B
4V GW 339
Intensity (counts)
2.2
time (min)
C
4V GW 261
Intensity (counts)
2.066
time (min)
D
4V GW 341
Intensity (counts)
2.2
time (min)
E
4V GW Reactant
1e6
Intensity (counts)
100.0761
121.0509
mass to charge (m/z)
F
4V GW Product
1e6
Intensity (counts)
121.0509
155.1185
128.1074
100.0759
118.0885
177.1008
mass to charge (m/z)
G
4V GW Reactant
Y(Counts)
261.1268
262.1579
X(Thomsons)
H
4V GW Brominated Product
Y(Counts)
339.0459
341.0446
X(Thomsons)

## 4V HW

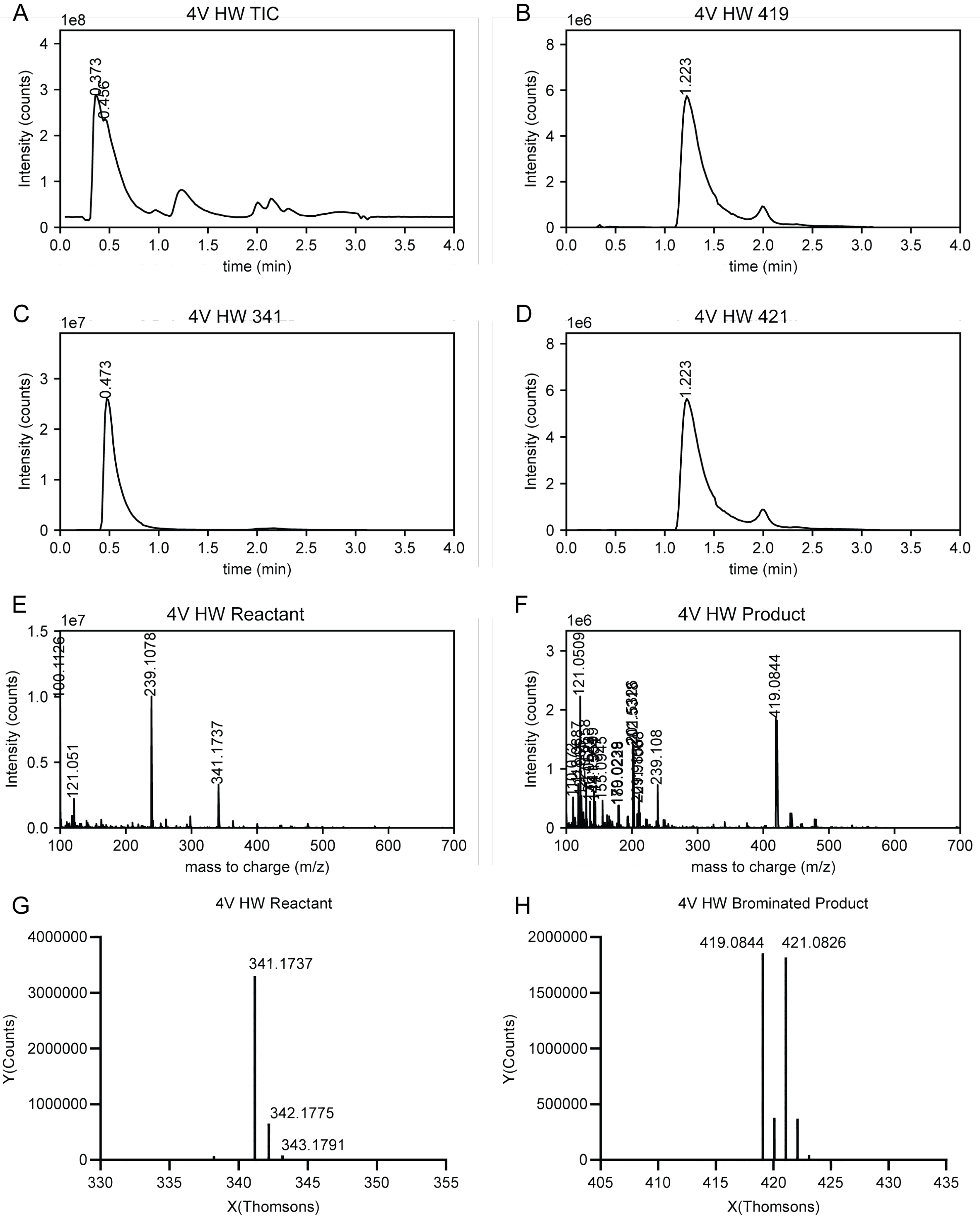
A
4V HW TIC
1e8
Intensity (counts)
time (min)
0.373
0.456
B
4V HW 419
1e6
Intensity (counts)
time (min)
1.223
C
4V HW 341
1e7
Intensity (counts)
time (min)
0.473
D
4V HW 421
1e6
Intensity (counts)
time (min)
1.223
E
4V HW Reactant
1e7
Intensity (counts)
mass to charge (m/z)
100.1126
121.051
239.1078
341.1737
F
4V HW Product
1e6
Intensity (counts)
mass to charge (m/z)
121.0509
239.108
419.0844
G
4V HW Reactant
Y(Counts)
X(Thomsons)
341.1737
342.1775
343.1791
H
4V HW Brominated Product
Y(Counts)
X(Thomsons)
419.0844
421.0826

## 4V IW

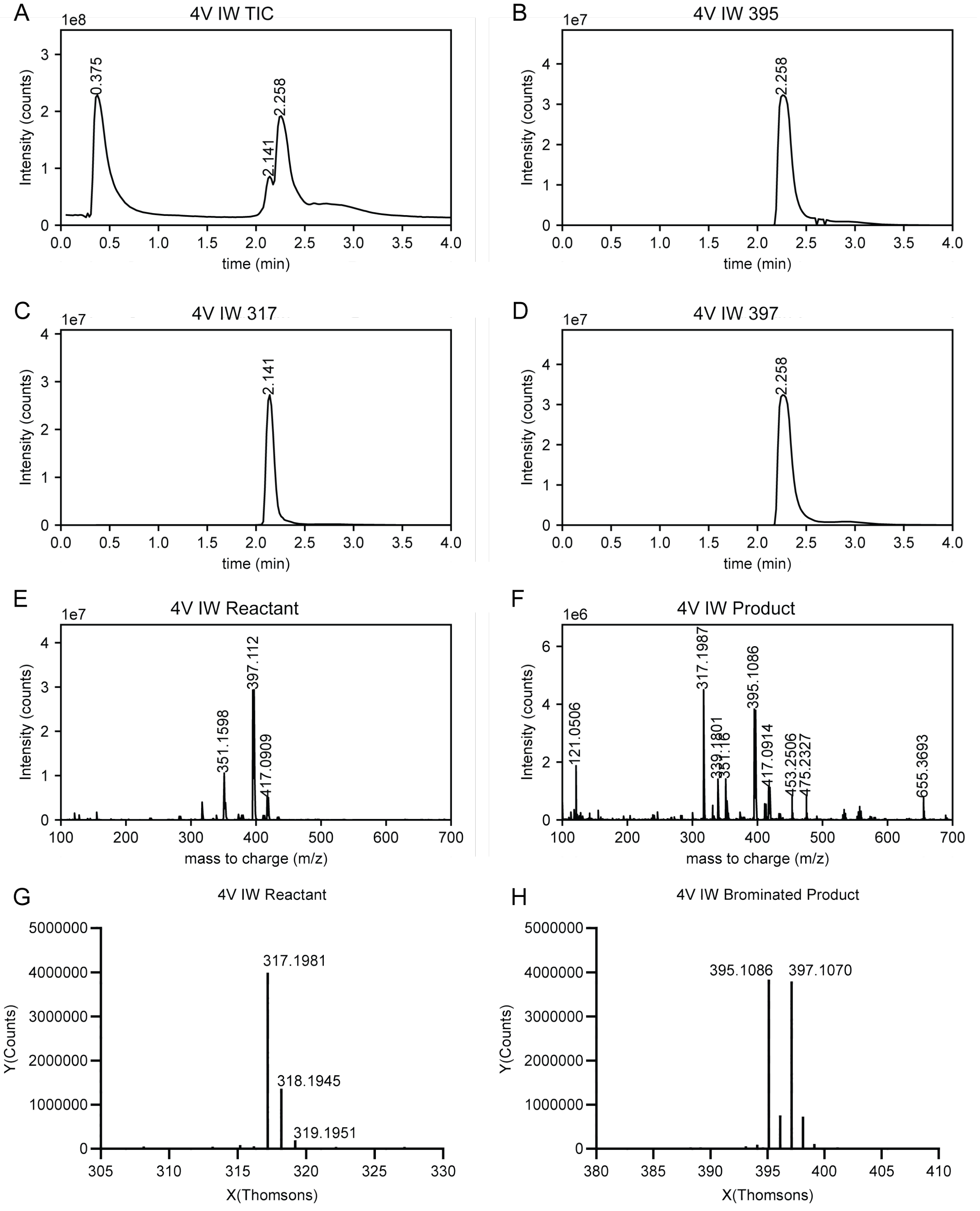
A
4V IW TIC
B
4V IW 395
C
4V IW 317
D
4V IW 397
E
4V IW Reactant
F
4V IW Product
G
4V IW Reactant
H
4V IW Brominated Product
Intensity (counts)
time (min)
mass to charge (m/z)
Y(Counts)
X(Thomsons)
0.375
2.141
2.258
351.1598
397.112
417.0909
121.0506
317.1987
339.1801
351.16
395.1086
417.0914
453.2506
475.2327
655.3693
317.1981
318.1945
319.1951
397.1070

## 4V KW

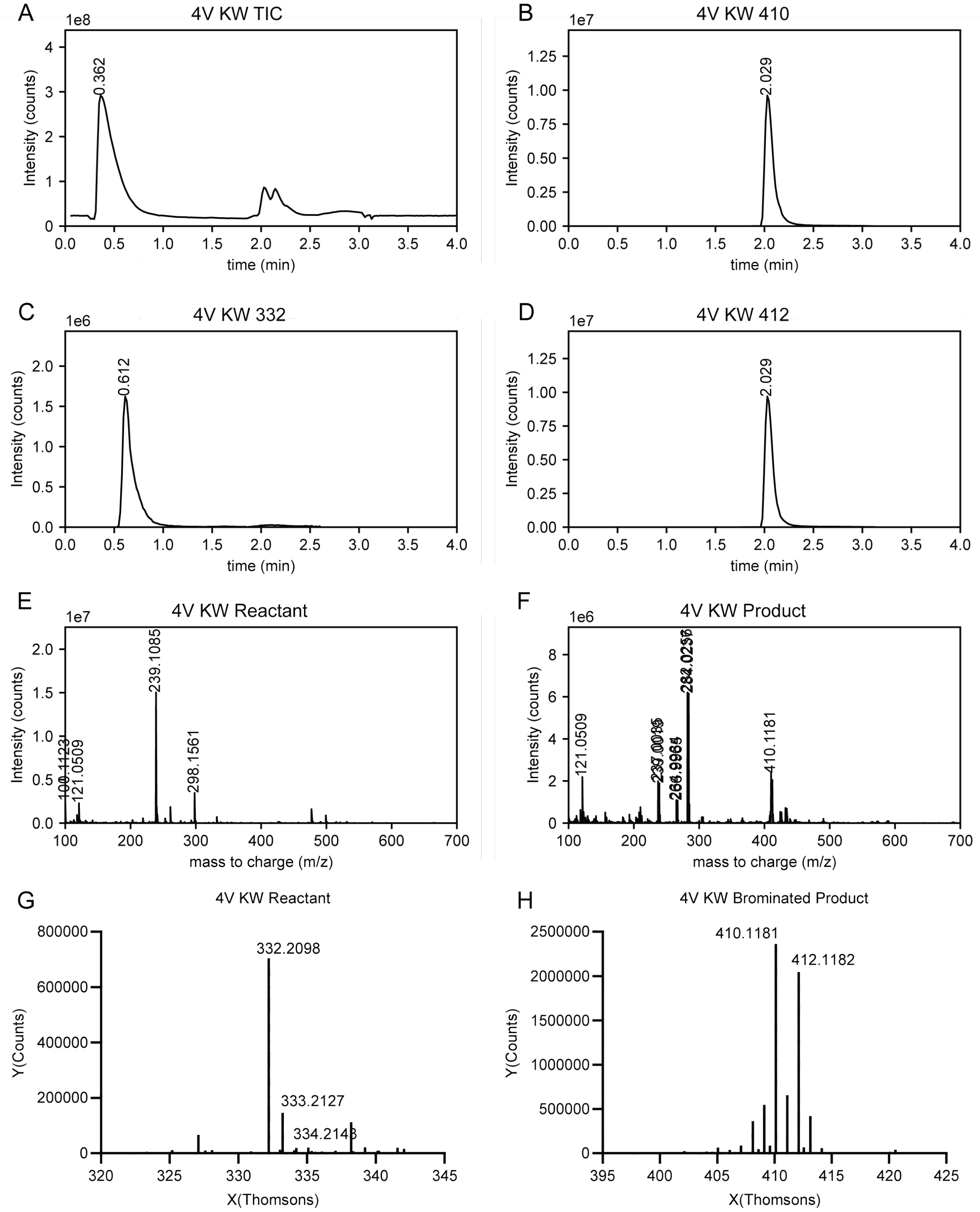
A
4V KW TIC
1e8
0.362
Intensity (counts)
time (min)
B
4V KW 410
1e7
2.029
Intensity (counts)
time (min)
C
4V KW 332
1e6
0.612
Intensity (counts)
time (min)
D
4V KW 412
1e7
2.029
Intensity (counts)
time (min)
E
4V KW Reactant
1e7
239.1085
298.1561
121.0509
Intensity (counts)
mass to charge (m/z)
F
4V KW Product
1e6
121.0509
410.1181
Intensity (counts)
mass to charge (m/z)
G
4V KW Reactant
332.2098
333.2127
334.2143
Y(Counts)
X(Thomsons)
H
4V KW Brominated Product
410.1181
412.1182
Y(Counts)
X(Thomsons)

# 4V LW

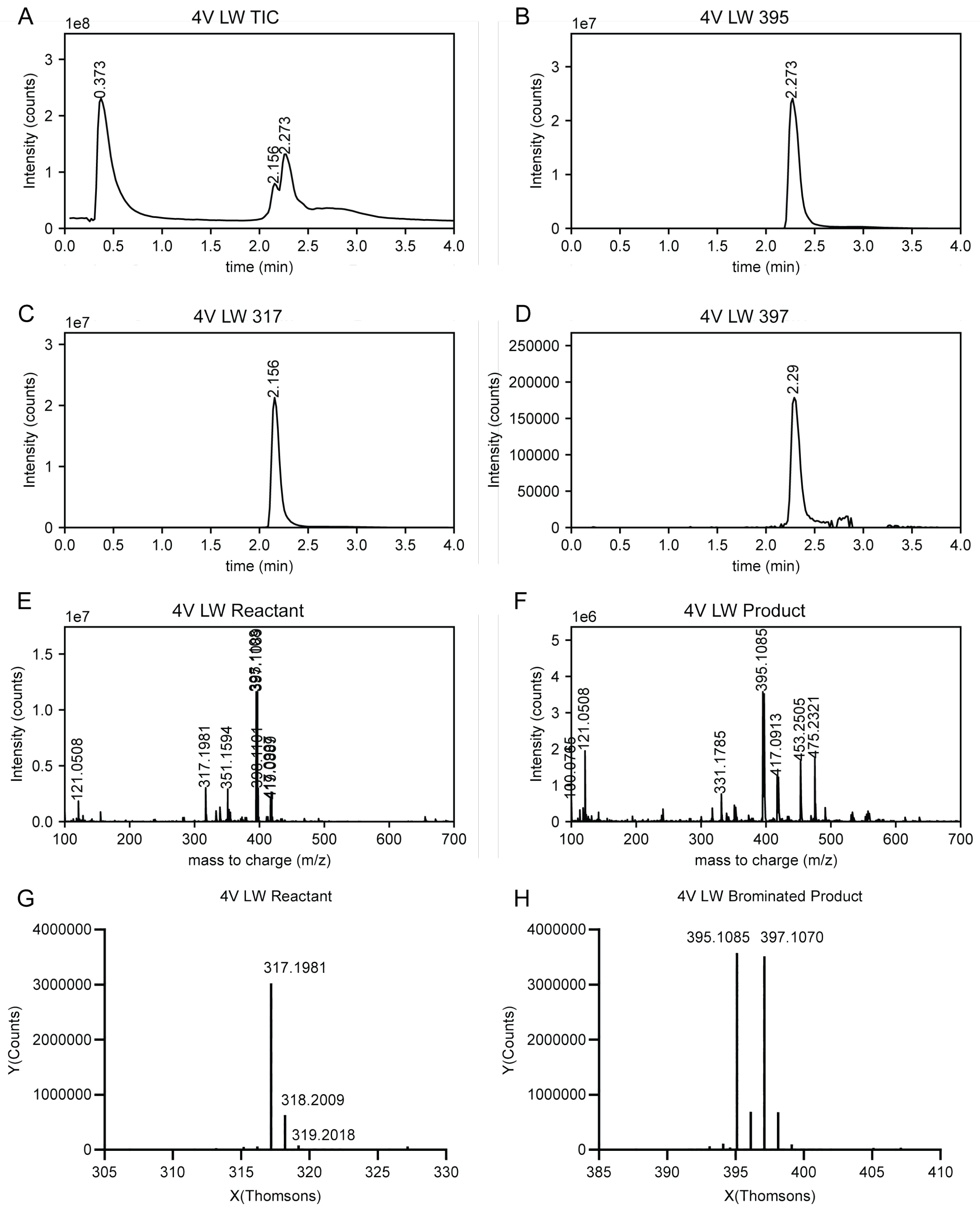

A
4V LW TIC
1e8
Intensity (counts)
time (min)
0.373
2.156
2.273
B
4V LW 395
1e7
Intensity (counts)
time (min)
2.273
C
4V LW 317
1e7
Intensity (counts)
time (min)
2.156
D
4V LW 397
Intensity (counts)
time (min)
2.29
E
4V LW Reactant
1e7
Intensity (counts)
mass to charge (m/z)
121.0508
317.1981
351.1594
F
4V LW Product
1e6
Intensity (counts)
mass to charge (m/z)
121.0508
331.1785
395.1085
417.0913
453.2505
475.2321
G
4V LW Reactant
Y(Counts)
X(Thomsons)
317.1981
318.2009
319.2018
H
4V LW Brominated Product
Y(Counts)
X(Thomsons)
395.1085
397.1070

## 4V MW

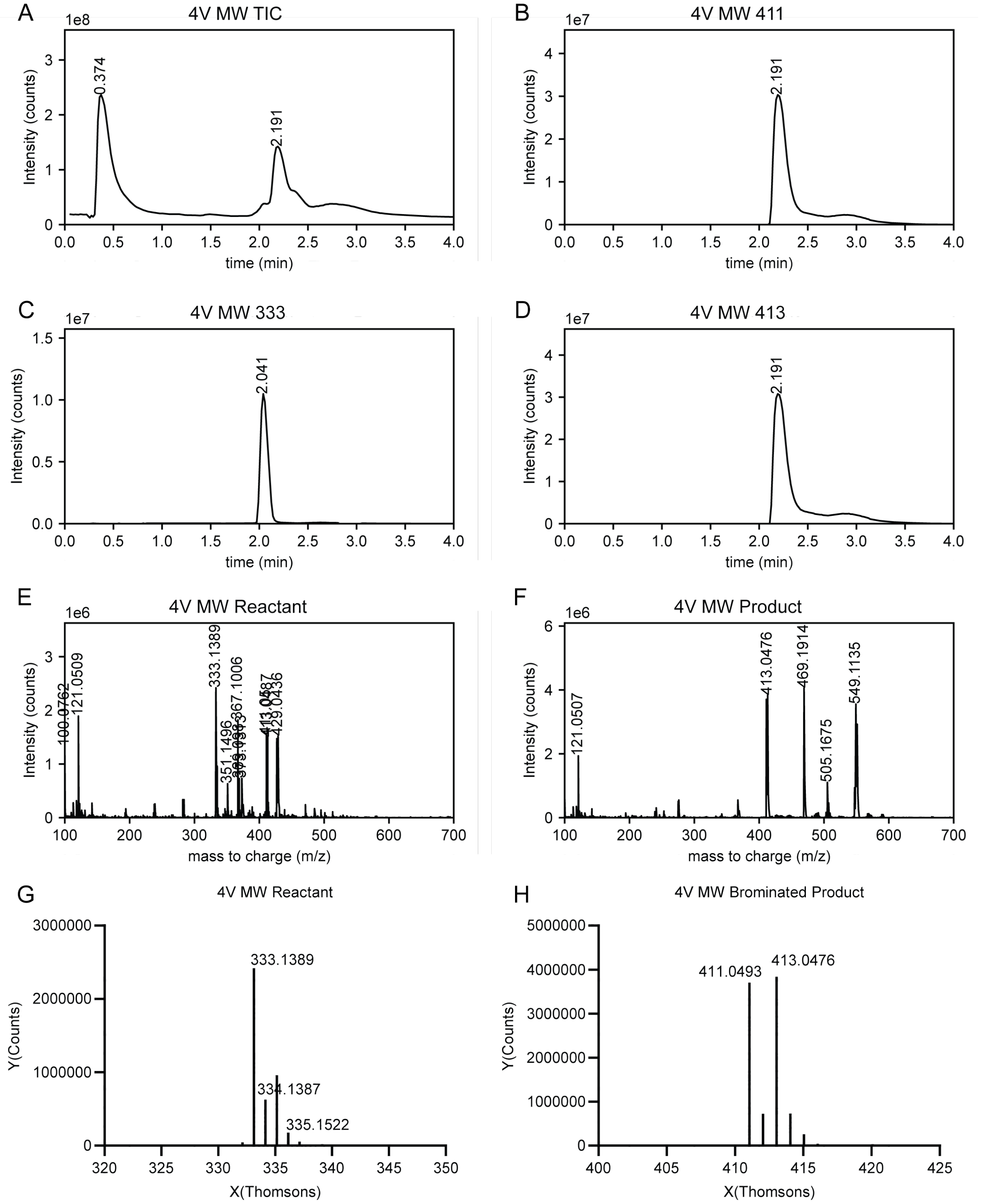

A
4V MW TIC
1e8
Intensity (counts)
0.374
2.191
time (min)
B
4V MW 411
1e7
2.191
C
4V MW 333
1e7
2.041
D
4V MW 413
1e7
2.191
E
4V MW Reactant
1e6
333.1389
121.0509
367.1006
351.1496
429.0436
mass to charge (m/z)
F
4V MW Product
1e6
413.0476
469.1914
549.1135
121.0507
505.1675
G
4V MW Reactant
333.1389
334.1387
335.1522
Y(Counts)
X(Thomsons)
H
4V MW Brominated Product
411.0493
413.0476

**4V NW**

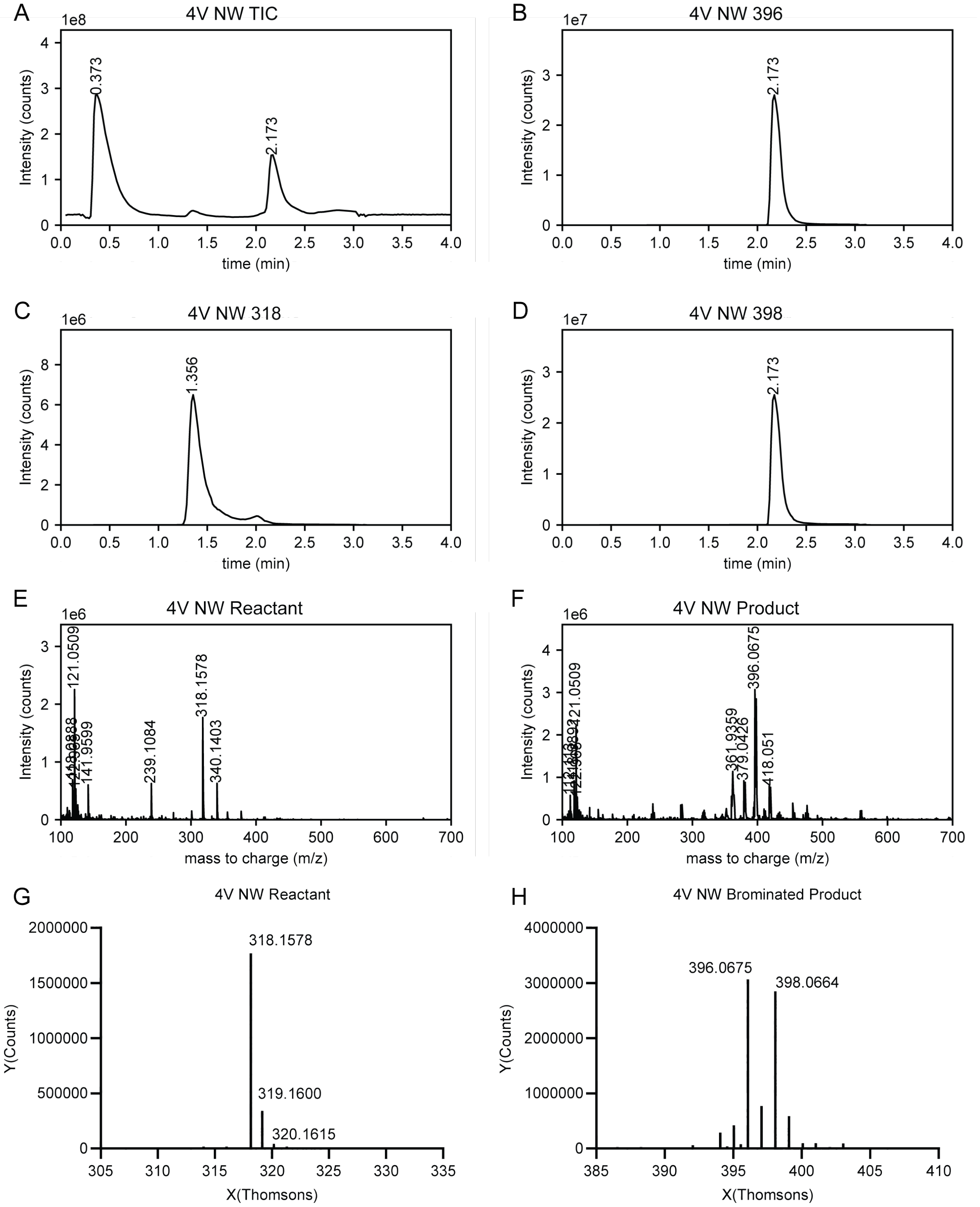
A
4V NW TIC
1e8
Intensity (counts)
time (min)
0.373
2.173
B
4V NW 396
1e7
Intensity (counts)
time (min)
2.173
C
4V NW 318
1e6
Intensity (counts)
time (min)
1.356
D
4V NW 398
1e7
Intensity (counts)
time (min)
2.173
E
4V NW Reactant
1e6
Intensity (counts)
mass to charge (m/z)
121.0509
141.9599
239.1084
318.1578
340.1403
F
4V NW Product
1e6
Intensity (counts)
mass to charge (m/z)
121.0509
361.9359
379.0426
396.0675
418.051
G
4V NW Reactant
Y(Counts)
X(Thomsons)
318.1578
319.1600
320.1615
H
4V NW Brominated Product
Y(Counts)
X(Thomsons)
396.0675
398.0664

## 4V PW

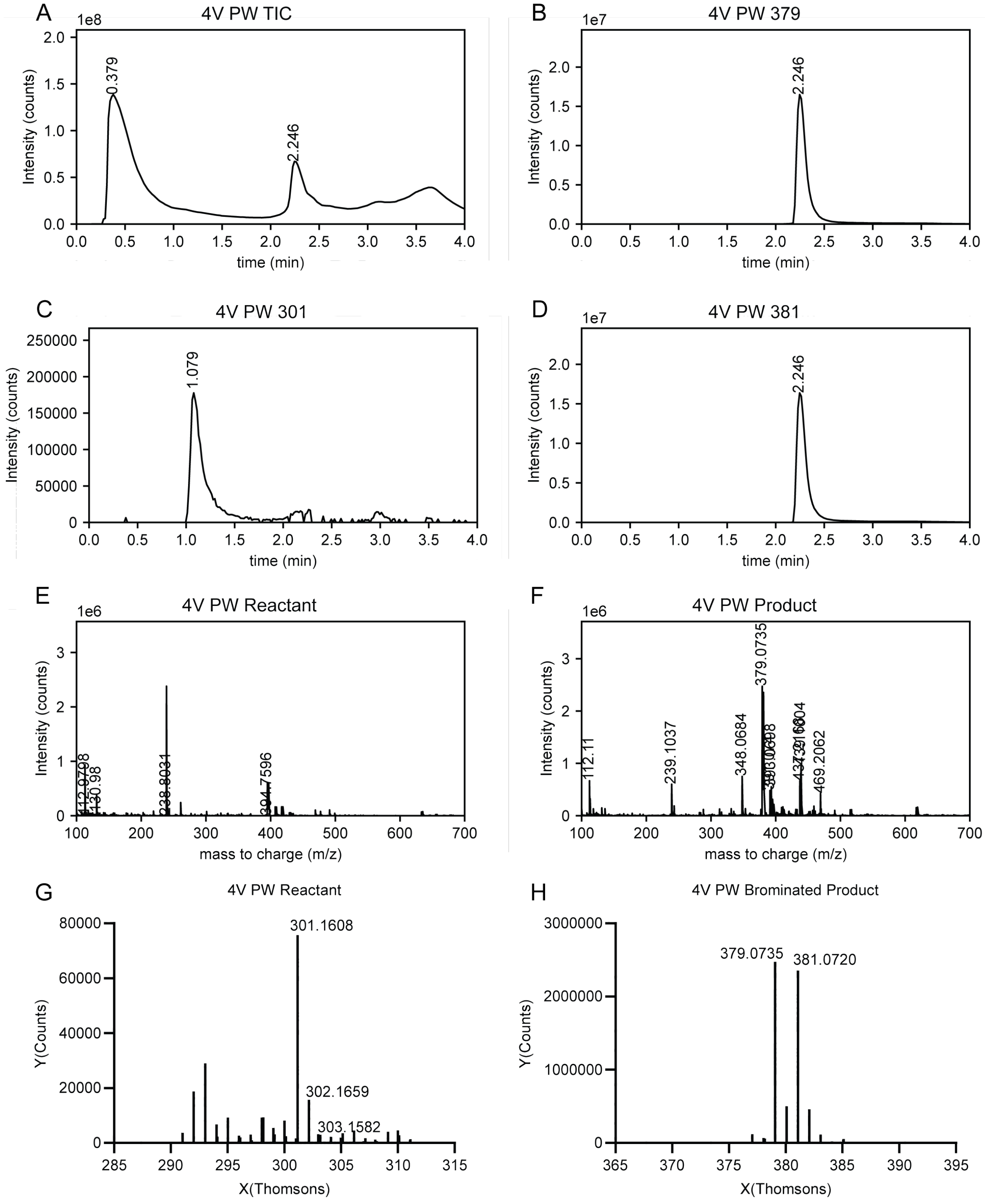

A
4V PW TIC
1e8
Intensity (counts)
time (min)
0.379
2.246
B
4V PW 379
1e7
Intensity (counts)
time (min)
2.246
C
4V PW 301
Intensity (counts)
time (min)
1.079
D
4V PW 381
1e7
Intensity (counts)
time (min)
2.246
E
4V PW Reactant
1e6
Intensity (counts)
mass to charge (m/z)
130.98
238.8031
394.7596
F
4V PW Product
1e6
Intensity (counts)
mass to charge (m/z)
112.11
239.1037
348.0684
379.0735
469.2062
G
4V PW Reactant
Y(Counts)
X(Thomsons)
301.1608
302.1659
303.1582
H
4V PW Brominated Product
Y(Counts)
X(Thomsons)
379.0735
381.0720

## 4V QW

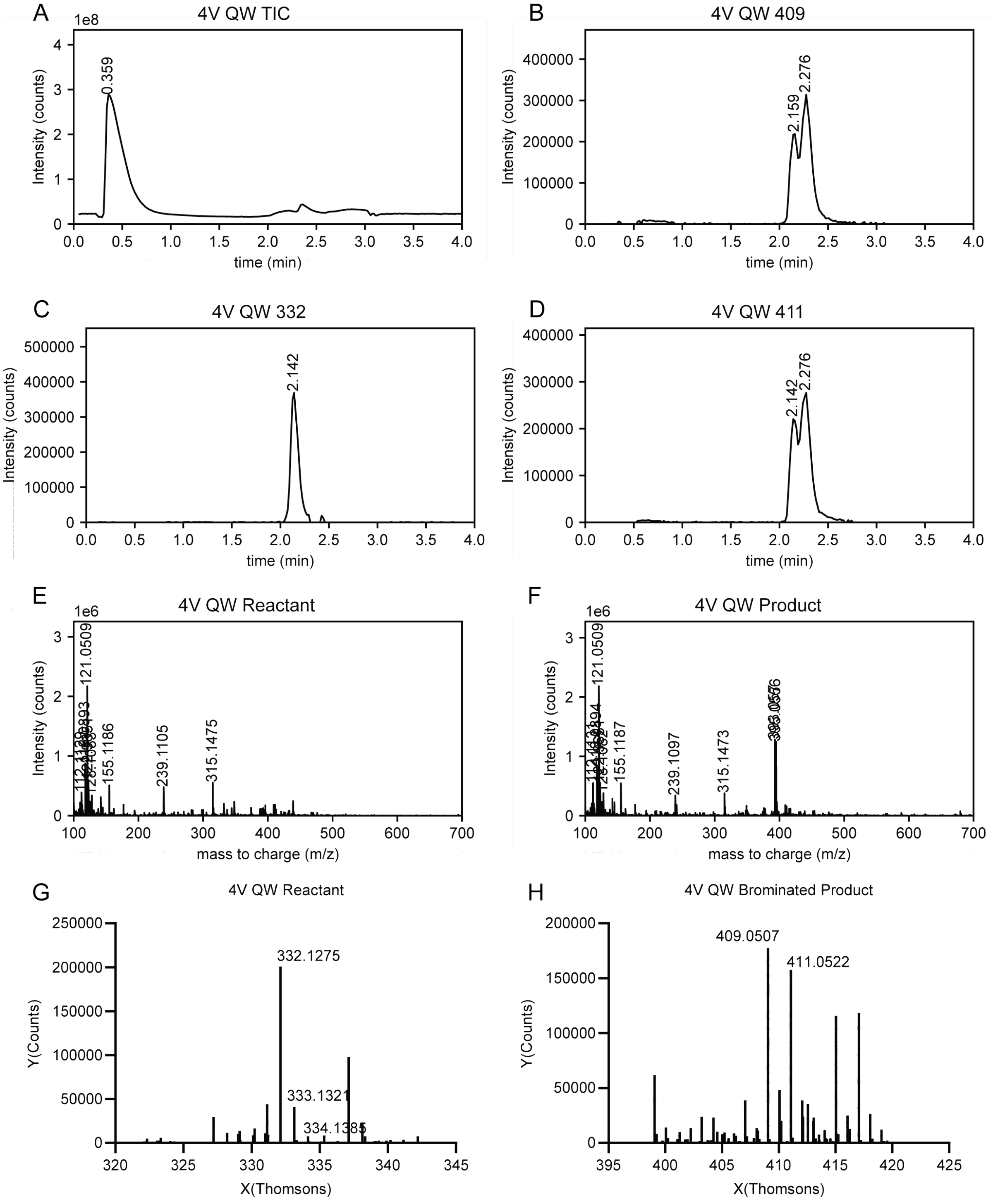
A
4V QW TIC
1e8
0.359
Intensity (counts)
time (min)
B
4V QW 409
2.159
2.276
Intensity (counts)
time (min)
C
4V QW 332
2.142
Intensity (counts)
time (min)
D
4V QW 411
2.142
2.276
Intensity (counts)
time (min)
E
4V QW Reactant
1e6
121.0509
155.1186
239.1105
315.1475
Intensity (counts)
mass to charge (m/z)
F
4V QW Product
1e6
121.0509
155.1187
239.1097
315.1473
Intensity (counts)
mass to charge (m/z)
G
4V QW Reactant
332.1275
333.1321
334.1385
Y(Counts)
X(Thomsons)
H
4V QW Brominated Product
409.0507
411.0522
Y(Counts)
X(Thomsons)

## 4V RW

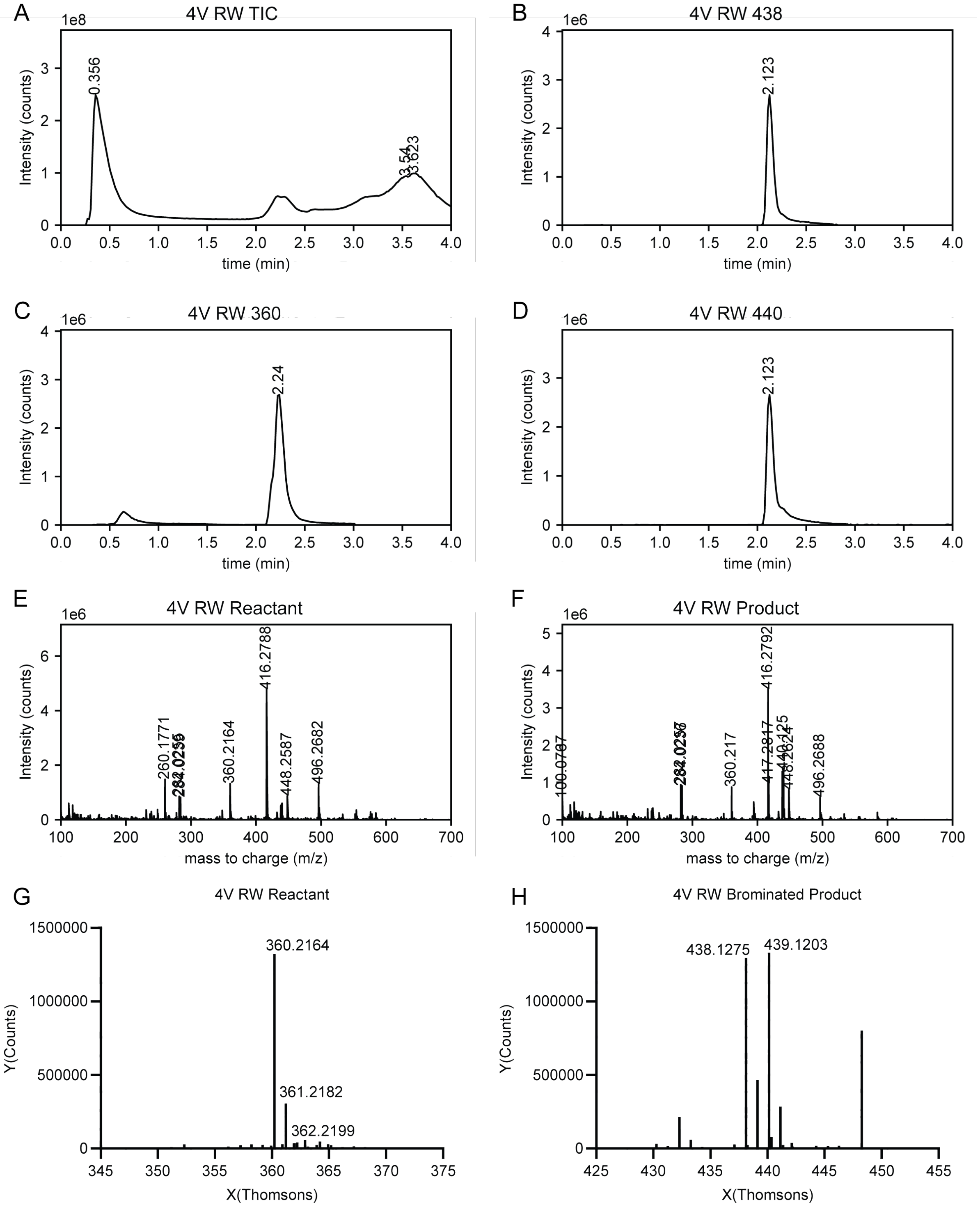

A
4V RW TIC
1e8
Intensity (counts)
0.356
3.54
3.623
time (min)
B
4V RW 438
1e6
Intensity (counts)
2.123
time (min)
C
4V RW 360
1e6
Intensity (counts)
2.24
time (min)
D
4V RW 440
1e6
Intensity (counts)
2.123
time (min)
E
4V RW Reactant
1e6
Intensity (counts)
260.1771
360.2164
416.2788
448.2587
496.2682
mass to charge (m/z)
F
4V RW Product
1e6
Intensity (counts)
100.0787
360.217
416.2792
417.2817
440.125
448.2624
496.2688
mass to charge (m/z)
G
4V RW Reactant
Y(Counts)
360.2164
361.2182
362.2199
X(Thomsons)
H
4V RW Brominated Product
Y(Counts)
438.1275
439.1203
X(Thomsons)

## 4V SW

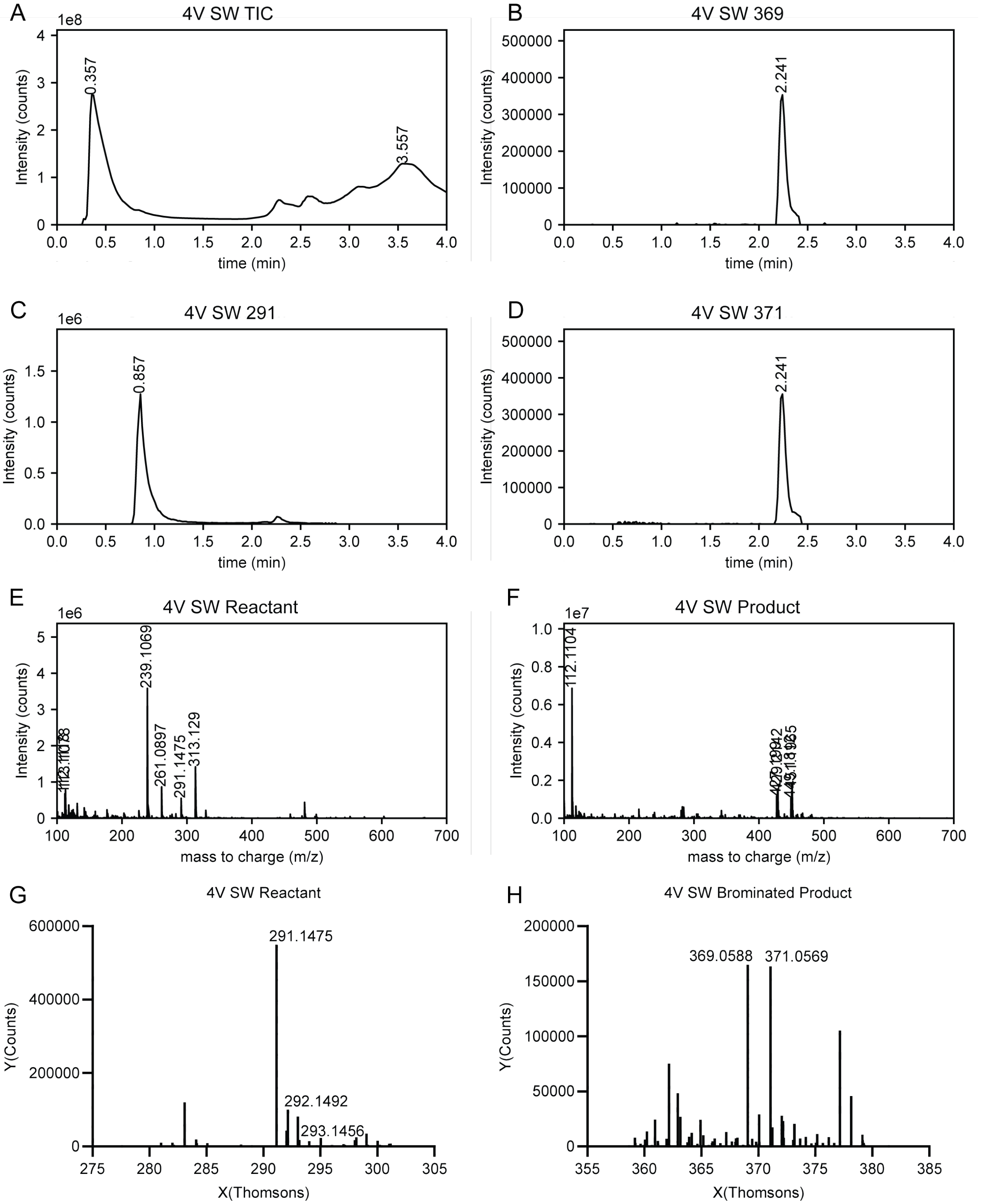
A
4V SW TIC
1e8
Intensity (counts)
time (min)
0.357
3.557
B
4V SW 369
Intensity (counts)
time (min)
2.241
C
4V SW 291
1e6
Intensity (counts)
time (min)
0.857
D
4V SW 371
Intensity (counts)
time (min)
2.241
E
4V SW Reactant
1e6
Intensity (counts)
mass to charge (m/z)
239.1069
261.0897
291.1475
313.129
F
4V SW Product
1e7
Intensity (counts)
mass to charge (m/z)
112.1104
G
4V SW Reactant
Y(Counts)
X(Thomsons)
291.1475
292.1492
293.1456
H
4V SW Brominated Product
Y(Counts)
X(Thomsons)
369.0588
371.0569

## 4V TW

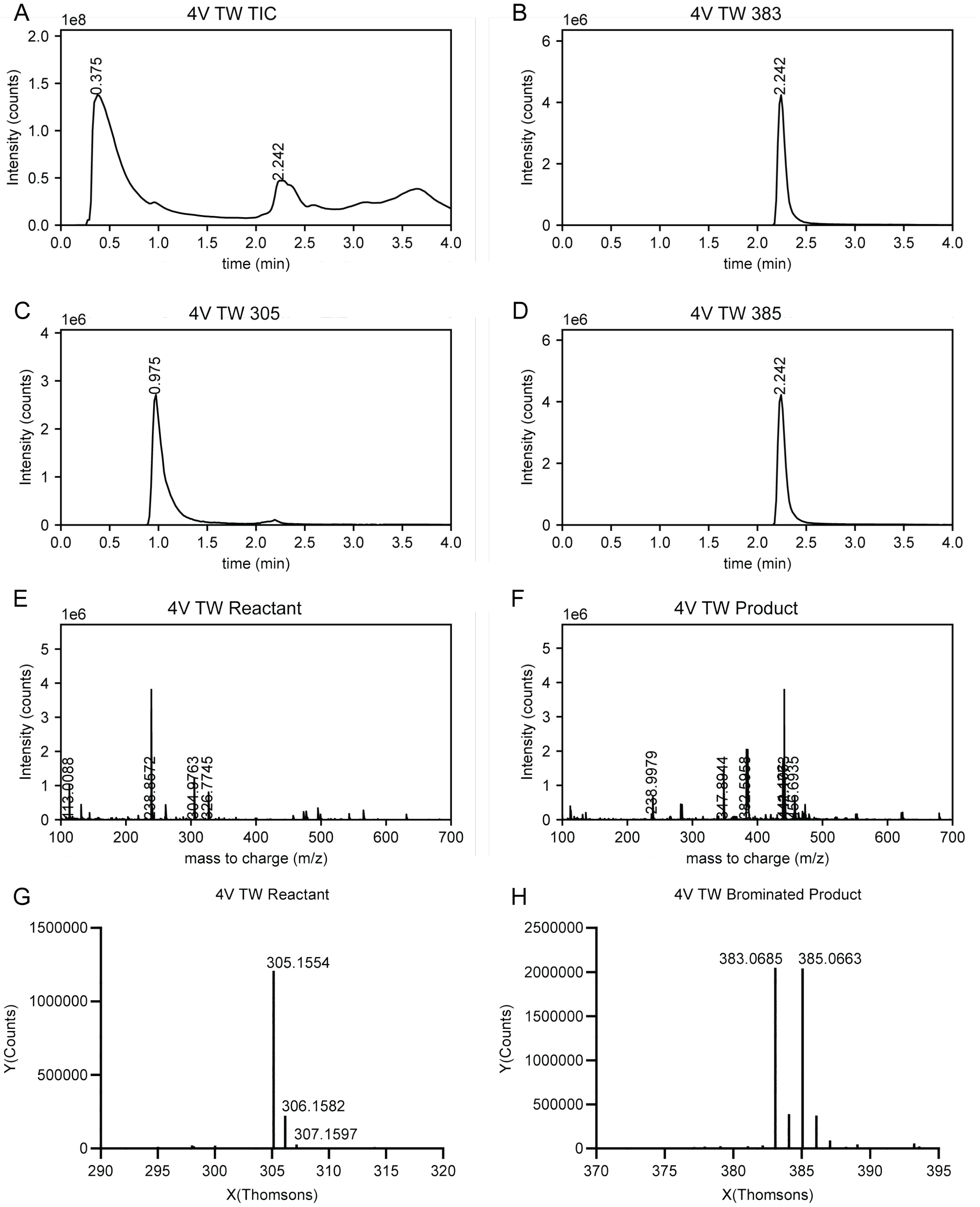
A
4V TW TIC
1e8
Intensity (counts)
time (min)
0.375
2.242
B
4V TW 383
1e6
2.242
C
4V TW 305
0.975
D
4V TW 385
2.242
E
4V TW Reactant
mass to charge (m/z)
238.8572
304.9763
326.7745
F
4V TW Product
238.9979
347.8944
G
4V TW Reactant
Y(Counts)
X(Thomsons)
305.1554
306.1582
307.1597
H
4V TW Brominated Product
383.0685
385.0663

## 4V VW

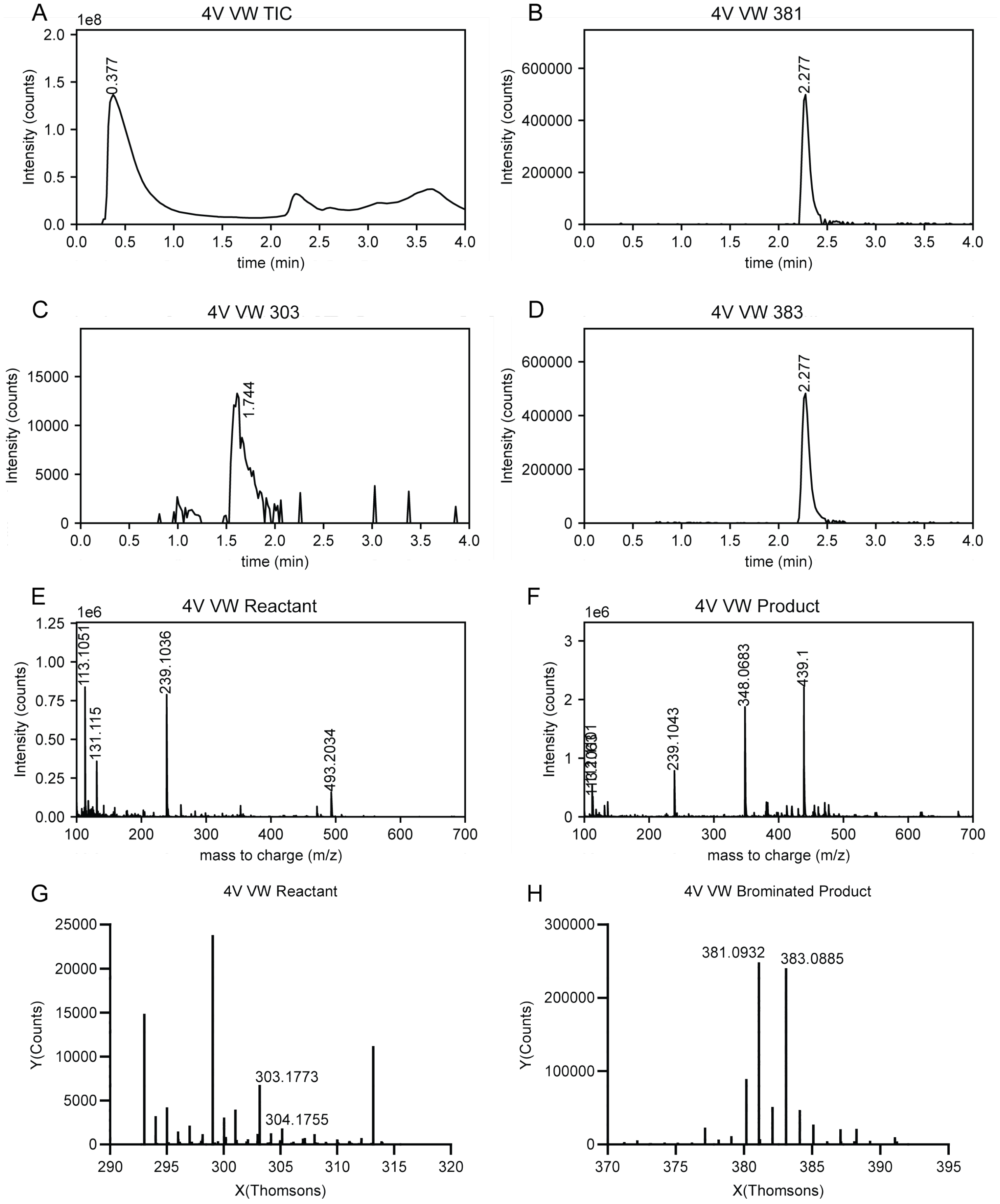

A
4V VW TIC
1e8
Intensity (counts)
time (min)
0.377
B
4V VW 381
Intensity (counts)
time (min)
2.277
C
4V VW 303
Intensity (counts)
time (min)
1.744
D
4V VW 383
Intensity (counts)
time (min)
2.277
E
4V VW Reactant
1e6
Intensity (counts)
mass to charge (m/z)
113.1051
131.115
239.1036
493.2034
F
4V VW Product
1e6
Intensity (counts)
mass to charge (m/z)
239.1043
348.0683
439.1
G
4V VW Reactant
Y(Counts)
X(Thomsons)
303.1773
304.1755
H
4V VW Brominated Product
Y(Counts)
X(Thomsons)
381.0932
383.0885

## 4V YW

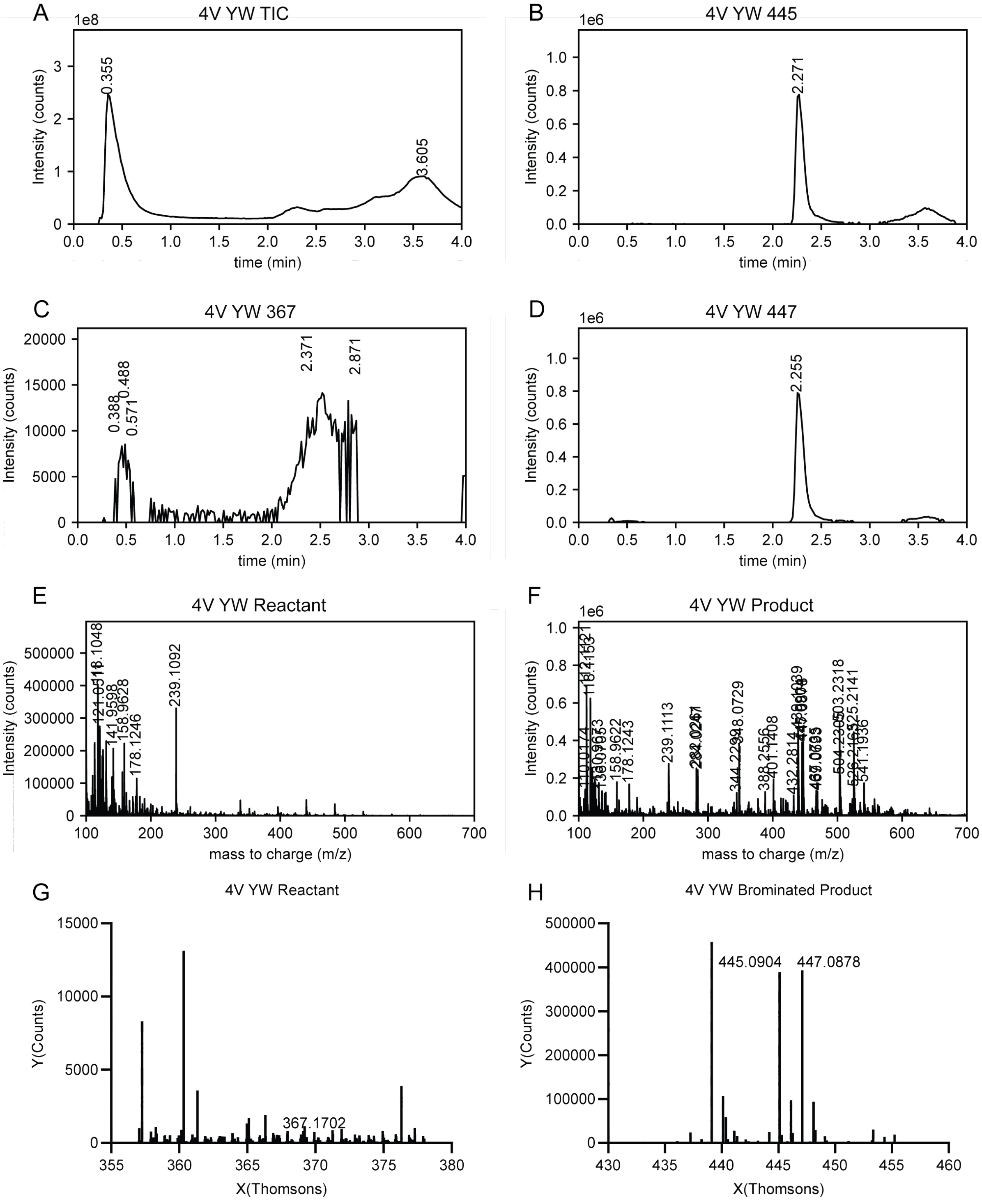
A
4V YW TIC
1e8
Intensity (counts)
time (min)
0.355
3.605
B
4V YW 445
1e6
Intensity (counts)
time (min)
2.271
C
4V YW 367
Intensity (counts)
time (min)
0.388
0.488
0.571
2.371
2.871
D
4V YW 447
1e6
Intensity (counts)
time (min)
2.255
E
4V YW Reactant
Intensity (counts)
mass to charge (m/z)
239.1092
141.9598
158.9628
178.1246
F
4V YW Product
1e6
Intensity (counts)
mass to charge (m/z)
239.1113
348.0729
388.2556
401.1408
432.2814
503.2318
525.2141
541.1936
158.9622
178.1243
G
4V YW Reactant
Y(Counts)
X(Thomsons)
367.1702
H
4V YW Brominated Product
Y(Counts)
X(Thomsons)
445.0904
447.0878

# 4V WA

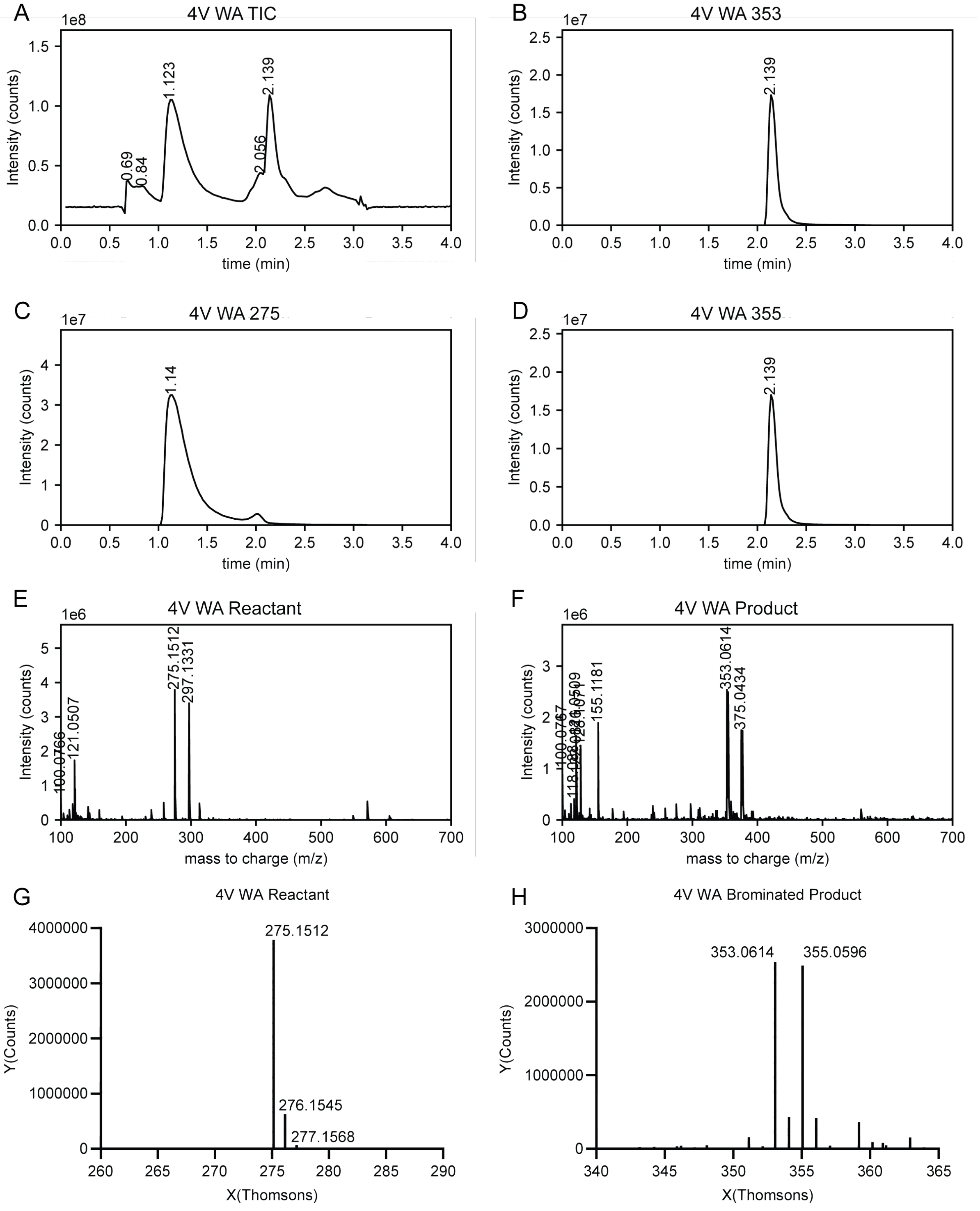

A
4V WA TIC
1e8
Intensity (counts)
time (min)
0.69
0.84
1.123
2.056
2.139
B
4V WA 353
1e7
Intensity (counts)
time (min)
2.139
C
4V WA 275
1e7
Intensity (counts)
time (min)
1.14
D
4V WA 355
1e7
Intensity (counts)
time (min)
2.139
E
4V WA Reactant
1e6
Intensity (counts)
mass to charge (m/z)
100.0766
121.0507
275.1512
297.1331
F
4V WA Product
1e6
Intensity (counts)
mass to charge (m/z)
155.1181
353.0614
375.0434
G
4V WA Reactant
Y(Counts)
X(Thomsons)
275.1512
276.1545
277.1568
H
4V WA Brominated Product
Y(Counts)
X(Thomsons)
353.0614
355.0596

## 4V WC

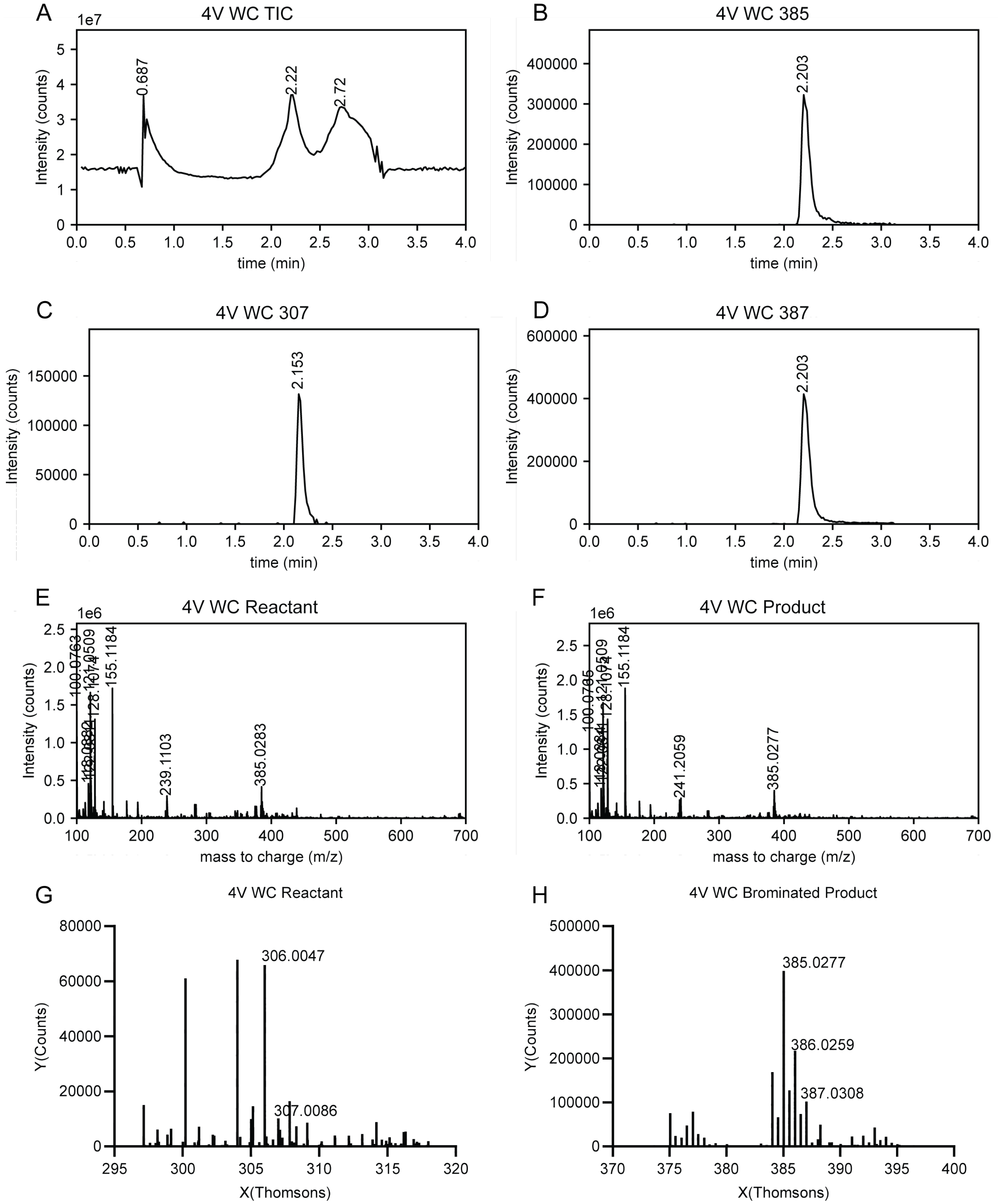
A
4V WC TIC
1e7
Intensity (counts)
time (min)
0.687
2.22
2.72
B
4V WC 385
Intensity (counts)
time (min)
2.203
C
4V WC 307
Intensity (counts)
time (min)
2.153
D
4V WC 387
Intensity (counts)
time (min)
2.203
E
4V WC Reactant
1e6
Intensity (counts)
mass to charge (m/z)
155.1184
239.1103
385.0283
F
4V WC Product
1e6
Intensity (counts)
mass to charge (m/z)
155.1184
241.2059
385.0277
G
4V WC Reactant
Y(Counts)
X(Thomsons)
306.0047
307.0086
H
4V WC Brominated Product
Y(Counts)
X(Thomsons)
385.0277
386.0259
387.0308

## 4V WD

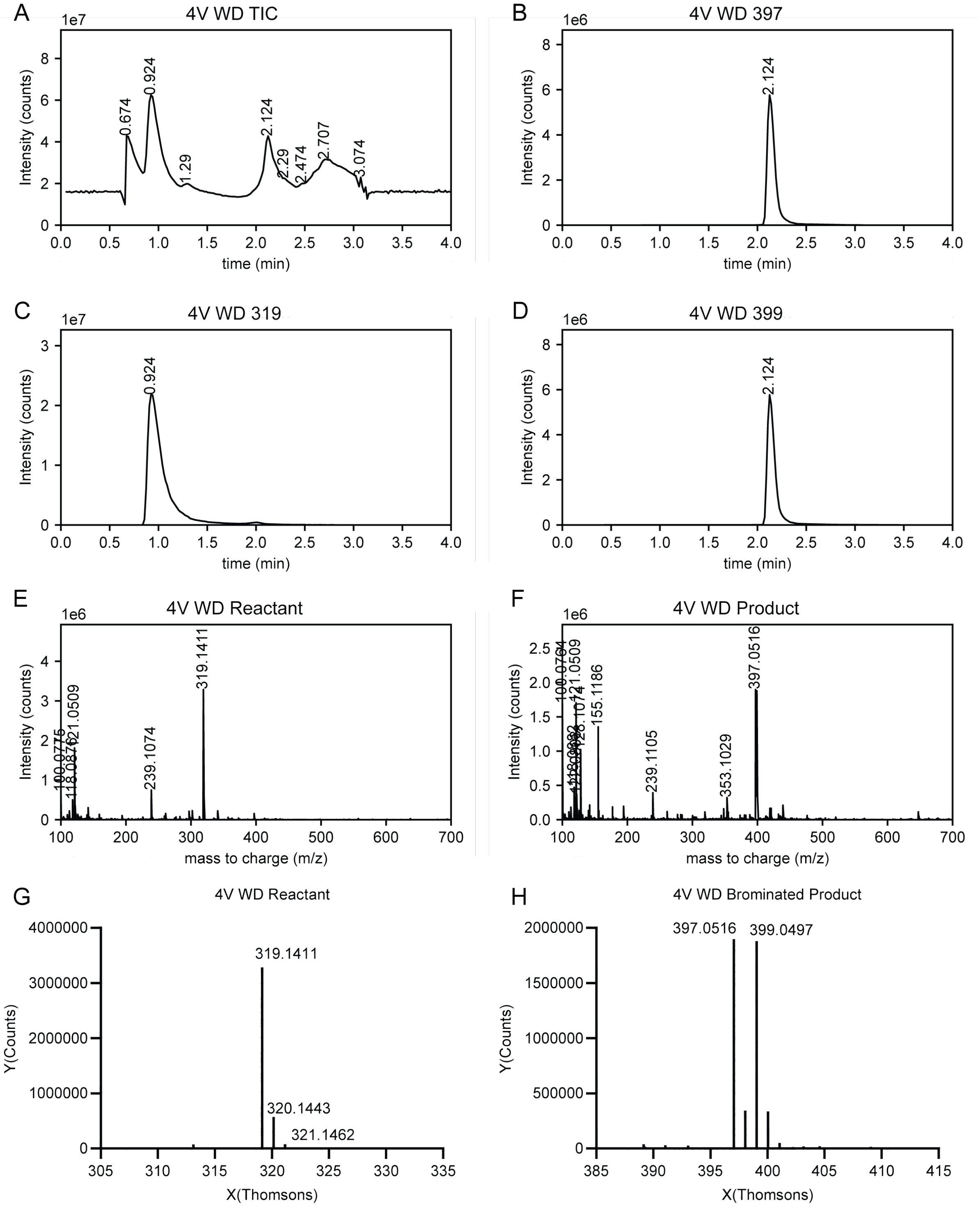

A
4V WD TIC
1e7
Intensity (counts)
time (min)
0.674
0.924
1.29
2.124
2.29
2.474
2.707
3.074
B
4V WD 397
1e6
2.124
C
4V WD 319
1e7
0.924
D
4V WD 399
1e6
2.124
E
4V WD Reactant
1e6
mass to charge (m/z)
121.0509
239.1074
319.1411
F
4V WD Product
1e6
121.0509
155.1186
239.1105
353.1029
397.0516
G
4V WD Reactant
Y(Counts)
X(Thomsons)
319.1411
320.1443
321.1462
H
4V WD Brominated Product
397.0516
399.0497

## 4V WE

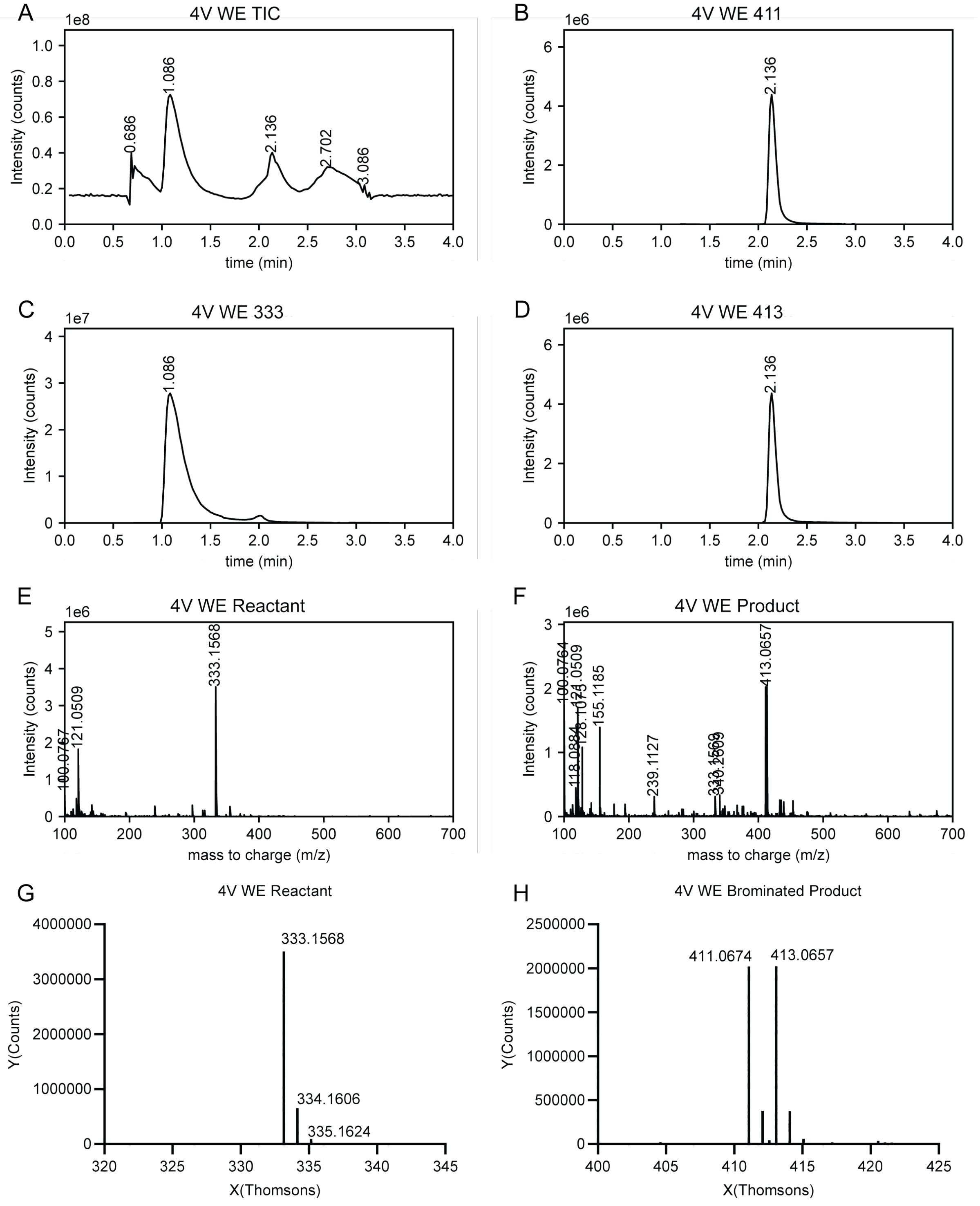
A
4V WE TIC
1e8
Intensity (counts)
time (min)
0.686
1.086
2.136
2.702
3.086
B
4V WE 411
1e6
Intensity (counts)
time (min)
2.136
C
4V WE 333
1e7
Intensity (counts)
time (min)
1.086
D
4V WE 413
1e6
Intensity (counts)
time (min)
2.136
E
4V WE Reactant
1e6
Intensity (counts)
mass to charge (m/z)
100.0767
121.0509
333.1568
F
4V WE Product
1e6
Intensity (counts)
mass to charge (m/z)
100.0764
118.0864
121.0509
126.1073
155.1185
239.1127
333.1569
340.2609
413.0657
G
4V WE Reactant
Y(Counts)
X(Thomsons)
333.1568
334.1606
335.1624
H
4V WE Brominated Product
Y(Counts)
X(Thomsons)
411.0674
413.0657

## 4V WF

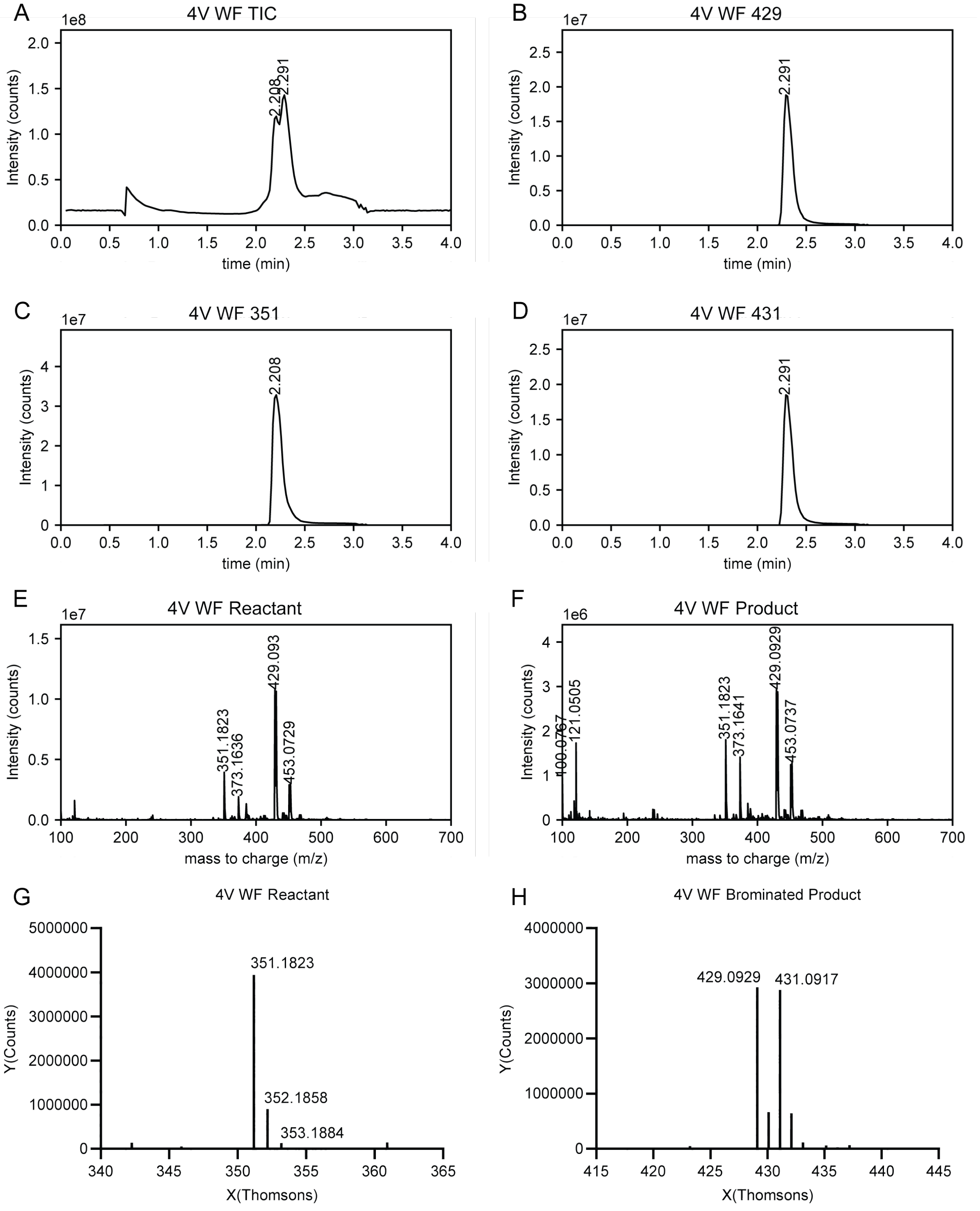

A
4V WF TIC
1e8
Intensity (counts)
time (min)
2.208
2.291
B
4V WF 429
1e7
Intensity (counts)
time (min)
2.291
C
4V WF 351
1e7
Intensity (counts)
time (min)
2.208
D
4V WF 431
1e7
Intensity (counts)
time (min)
2.291
E
4V WF Reactant
1e7
Intensity (counts)
mass to charge (m/z)
351.1823
373.1636
429.093
453.0729
F
4V WF Product
1e6
Intensity (counts)
mass to charge (m/z)
100.0767
121.0505
351.1823
373.1641
429.0929
453.0737
G
4V WF Reactant
Y(Counts)
X(Thomsons)
351.1823
352.1858
353.1884
H
4V WF Brominated Product
Y(Counts)
X(Thomsons)
429.0929
431.0917

## 4V WG

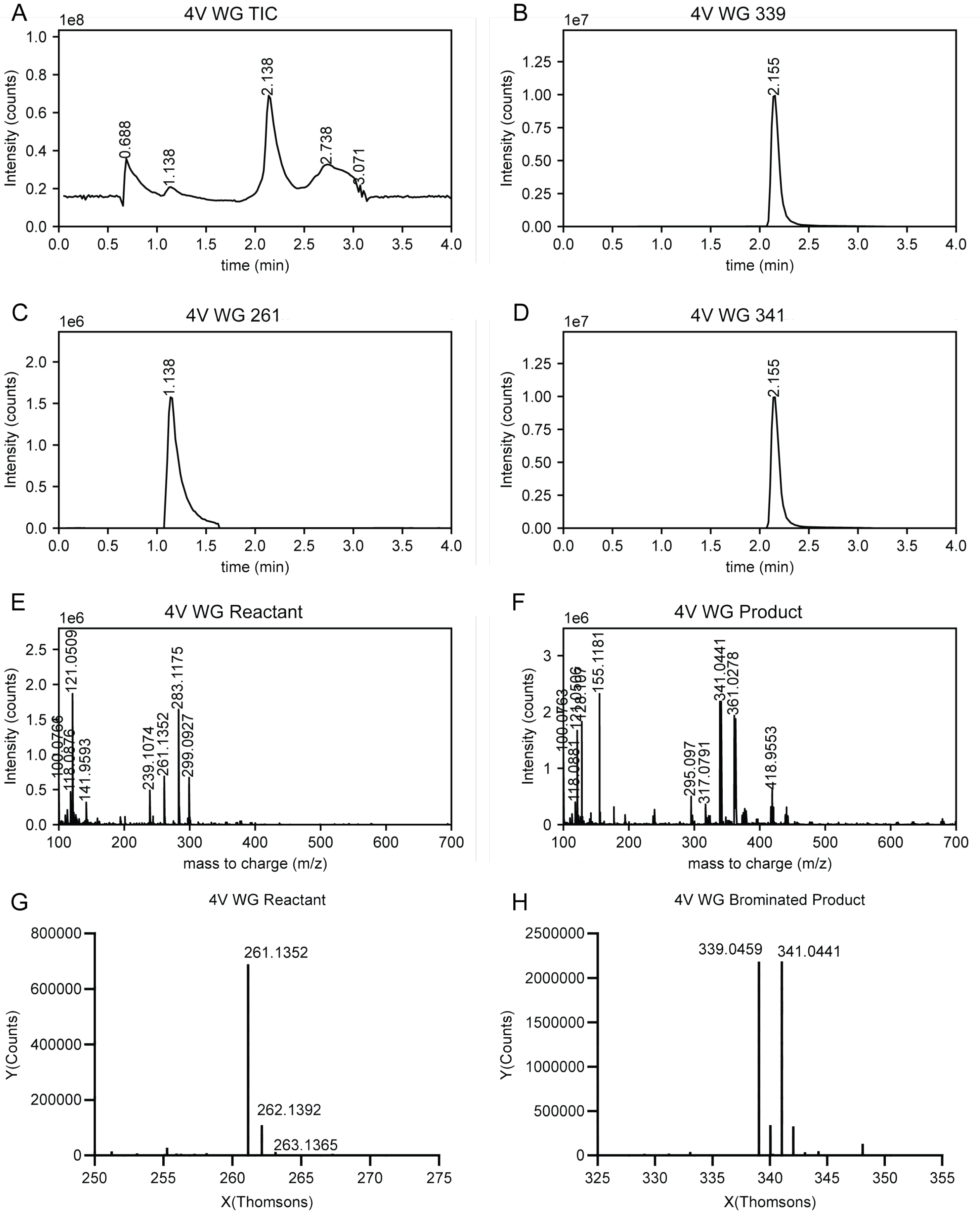

A
4V WG TIC
1e8
0.688
1.138
2.138
2.738
3.071
Intensity (counts)
time (min)
B
4V WG 339
1e7
2.155
Intensity (counts)
time (min)
C
4V WG 261
1e6
1.138
Intensity (counts)
time (min)
D
4V WG 341
1e7
2.155
Intensity (counts)
time (min)
E
4V WG Reactant
1e6
121.0509
141.9593
239.1074
261.1352
283.1175
299.0927
Intensity (counts)
mass to charge (m/z)
F
4V WG Product
1e6
155.1181
341.0441
361.0278
295.097
317.0791
418.9553
Intensity (counts)
mass to charge (m/z)
G
4V WG Reactant
261.1352
262.1392
263.1365
Y(Counts)
X(Thomsons)
H
4V WG Brominated Product
339.0459
341.0441
Y(Counts)
X(Thomsons)

## 4V WH

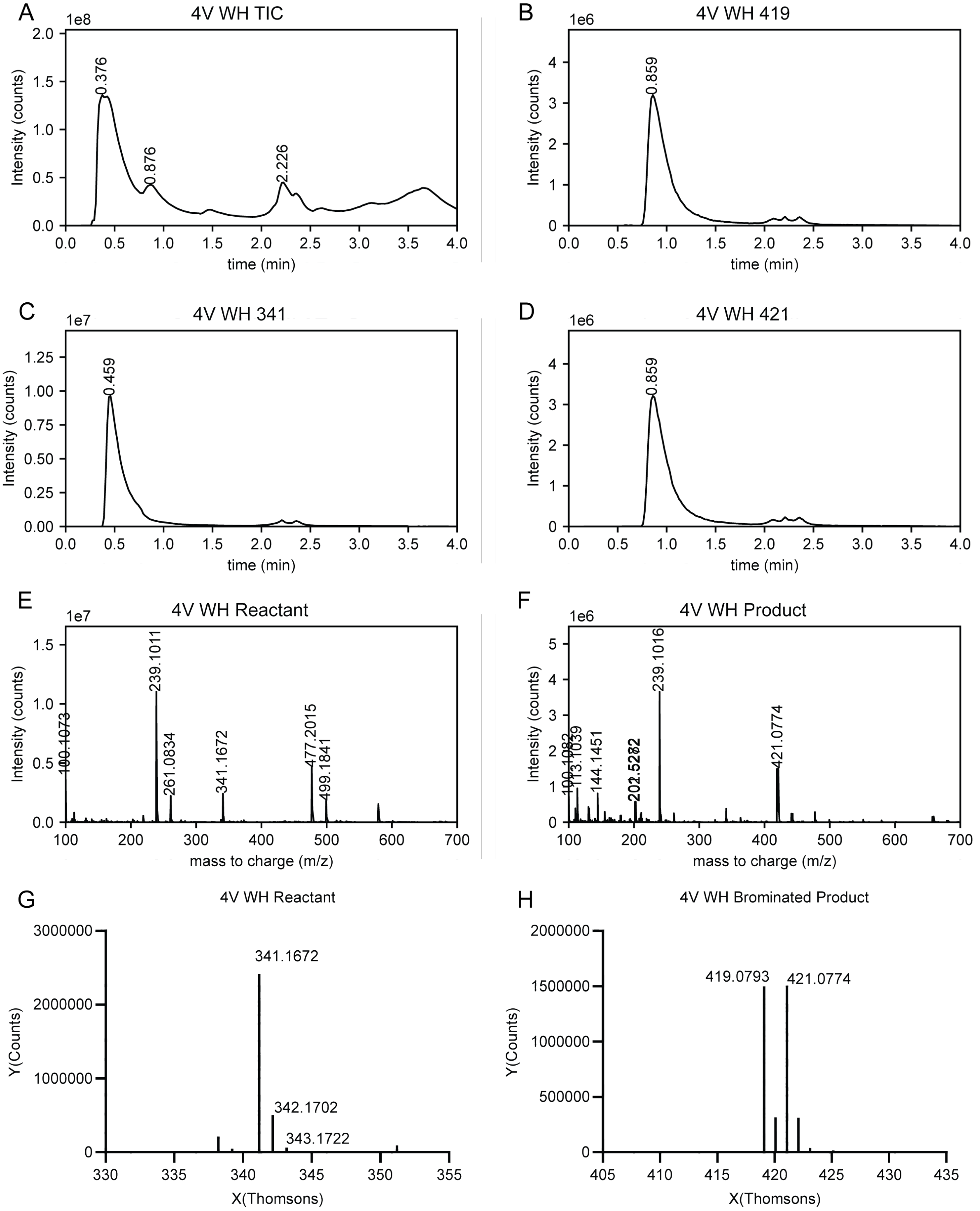

A
4V WH TIC
1e8
Intensity (counts)
time (min)
0.376
0.876
2.226
B
4V WH 419
1e6
Intensity (counts)
time (min)
0.859
C
4V WH 341
1e7
Intensity (counts)
time (min)
0.459
D
4V WH 421
1e6
Intensity (counts)
time (min)
0.859
E
4V WH Reactant
1e7
Intensity (counts)
mass to charge (m/z)
239.1011
261.0834
341.1672
477.2015
499.1841
F
4V WH Product
1e6
Intensity (counts)
mass to charge (m/z)
113.1039
144.1451
239.1016
421.0774
G
4V WH Reactant
Y(Counts)
X(Thomsons)
341.1672
342.1702
343.1722
H
4V WH Brominated Product
Y(Counts)
X(Thomsons)
419.0793
421.0774

## 4V WI

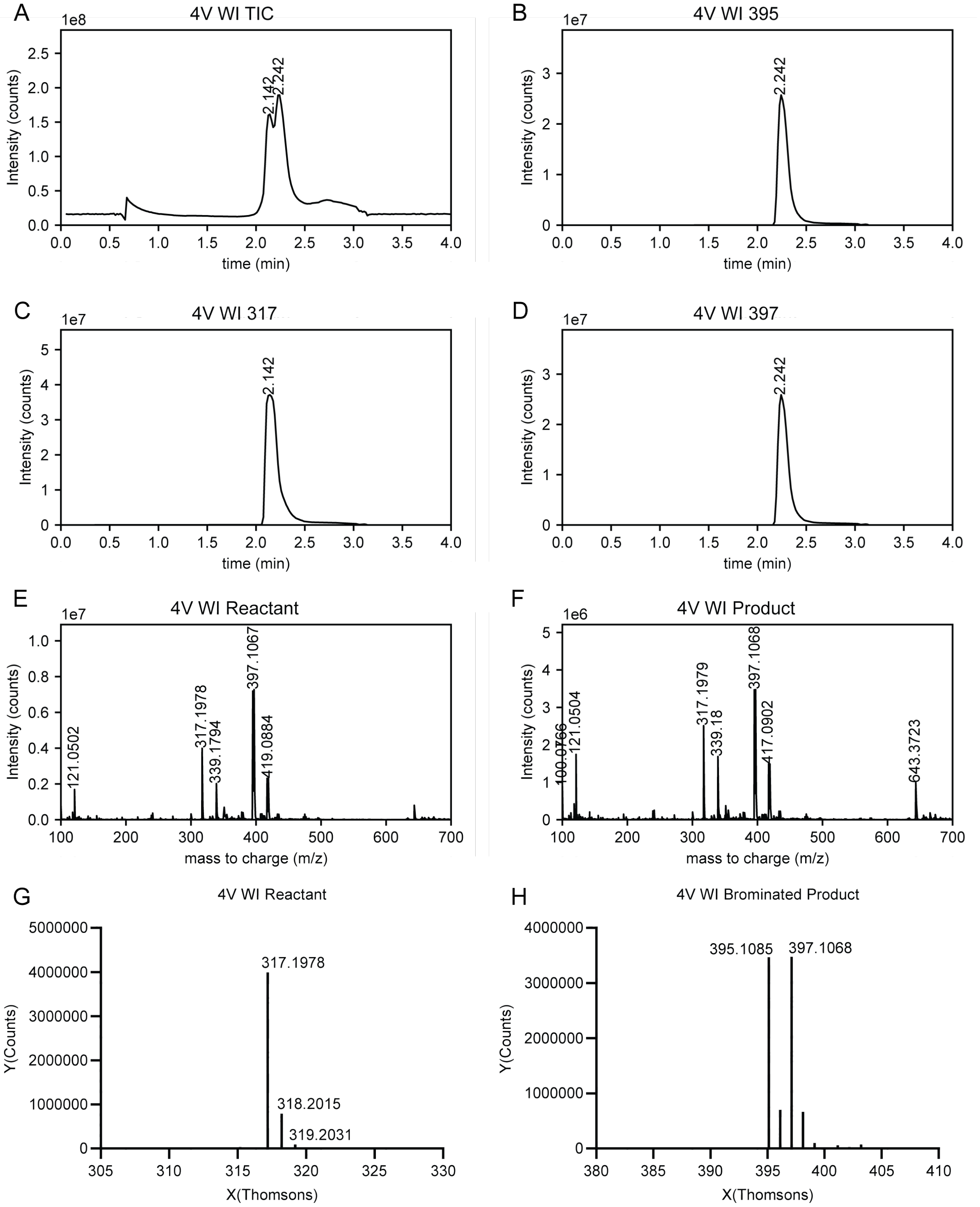

A
4V WI TIC
1e8
Intensity (counts)
time (min)
2.142
2.242
B
4V WI 395
1e7
Intensity (counts)
time (min)
2.242
C
4V WI 317
1e7
Intensity (counts)
time (min)
2.142
D
4V WI 397
1e7
Intensity (counts)
time (min)
2.242
E
4V WI Reactant
1e7
Intensity (counts)
mass to charge (m/z)
121.0502
317.1978
339.1794
397.1067
419.0884
F
4V WI Product
1e6
Intensity (counts)
mass to charge (m/z)
121.0504
317.1979
339.18
397.1068
417.0902
643.3723
G
4V WI Reactant
Y(Counts)
X(Thomsons)
317.1978
318.2015
319.2031
H
4V WI Brominated Product
Y(Counts)
X(Thomsons)
395.1085
397.1068

## 4V WK

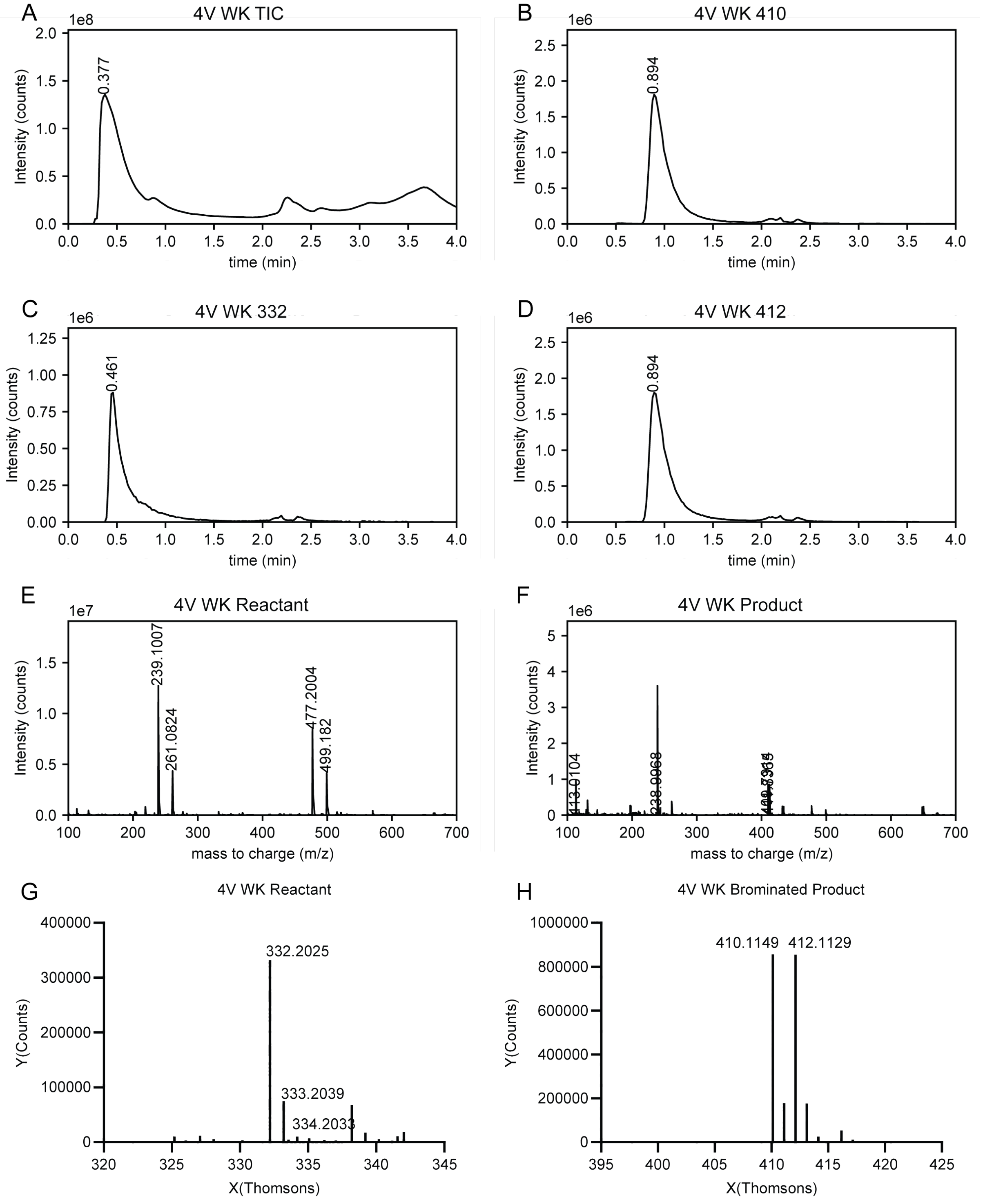

A
4V WK TIC
1e8
Intensity (counts)
time (min)
0.377
B
4V WK 410
1e6
Intensity (counts)
time (min)
0.894
C
4V WK 332
1e6
Intensity (counts)
time (min)
0.461
D
4V WK 412
1e6
Intensity (counts)
time (min)
0.894
E
4V WK Reactant
1e7
Intensity (counts)
mass to charge (m/z)
239.1007
261.0824
477.2004
499.182
F
4V WK Product
1e6
Intensity (counts)
mass to charge (m/z)
113.0104
G
4V WK Reactant
Y(Counts)
X(Thomsons)
332.2025
333.2039
334.2033
H
4V WK Brominated Product
Y(Counts)
X(Thomsons)
410.1149
412.1129

**4V WL**

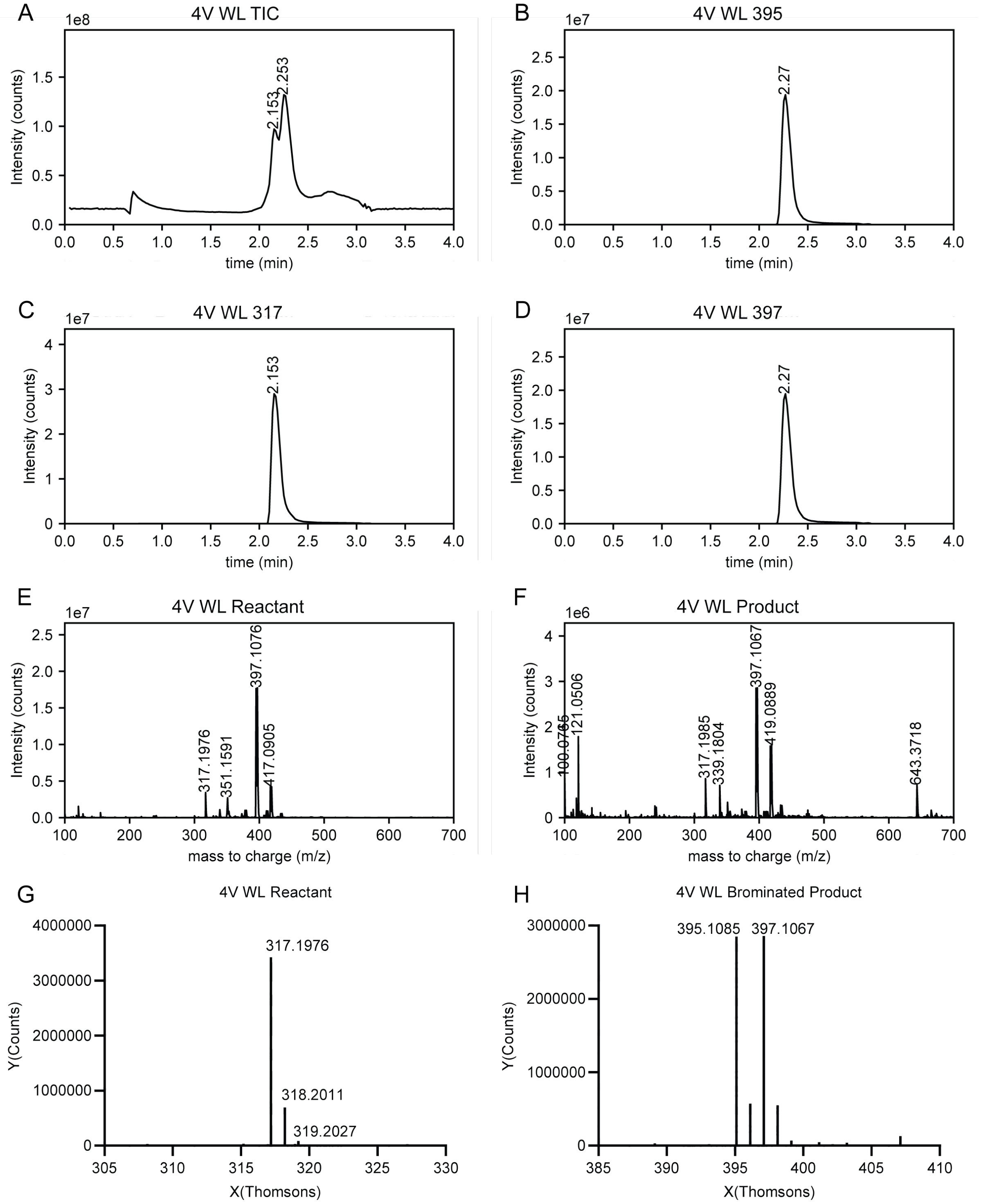

A
4V WL TIC
1e8
Intensity (counts)
2.153
2.253
time (min)
B
4V WL 395
1e7
2.27
C
4V WL 317
2.153
D
4V WL 397
2.27
E
4V WL Reactant
397.1076
317.1976
351.1591
417.0905
mass to charge (m/z)
F
4V WL Product
1e6
397.1067
121.0506
100.0765
419.0889
317.1985
339.1804
643.3718
G
4V WL Reactant
317.1976
318.2011
319.2027
Y(Counts)
X(Thomsons)
H
4V WL Brominated Product
395.1085
397.1067

## 4V WM

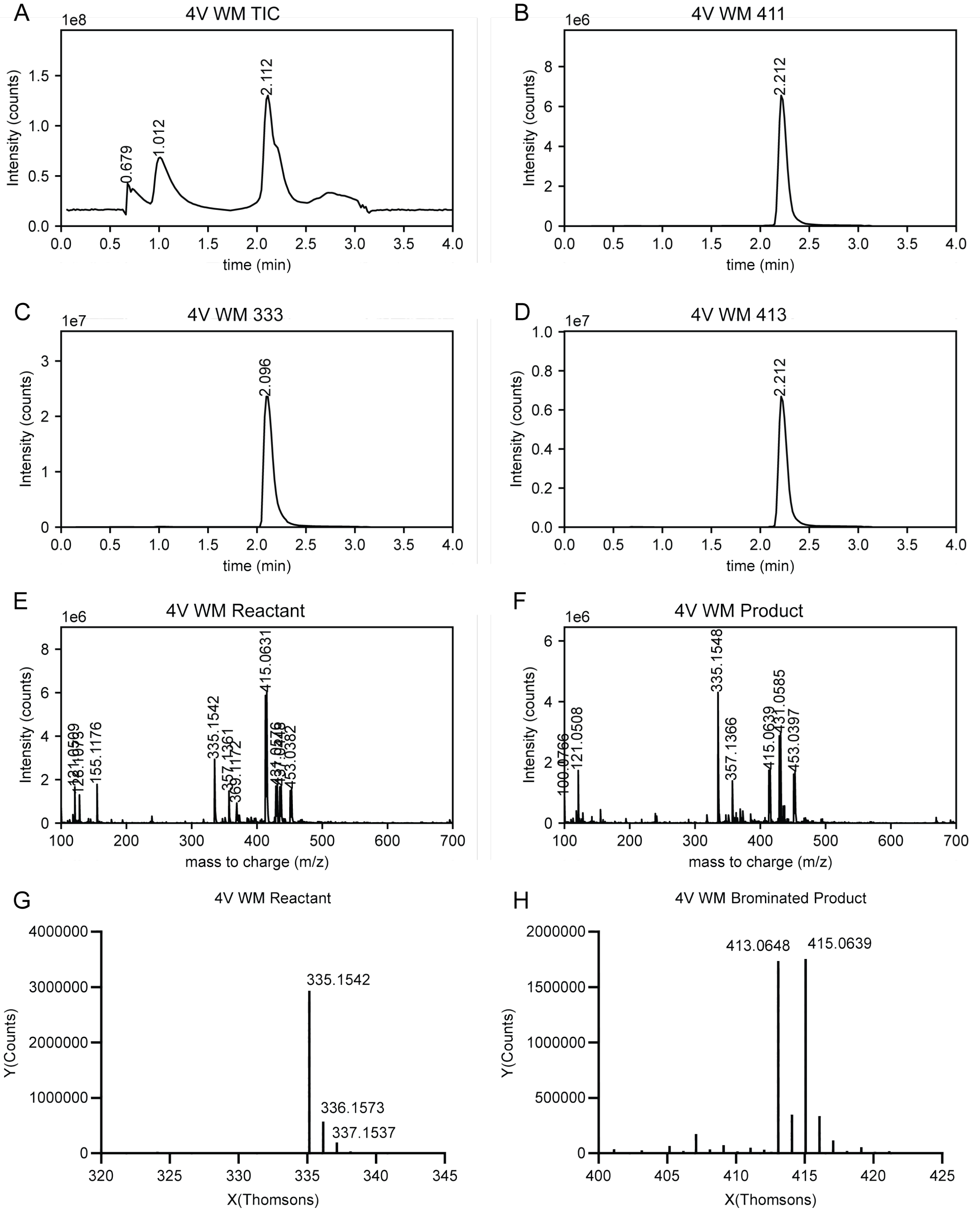

A
4V WM TIC
1e8
Intensity (counts)
time (min)
0.679
1.012
2.112
B
4V WM 411
1e6
Intensity (counts)
time (min)
2.212
C
4V WM 333
1e7
Intensity (counts)
time (min)
2.096
D
4V WM 413
1e7
Intensity (counts)
time (min)
2.212
E
4V WM Reactant
1e6
Intensity (counts)
mass to charge (m/z)
155.1176
335.1542
415.0631
453.0382
F
4V WM Product
1e6
Intensity (counts)
mass to charge (m/z)
100.0766
121.0508
335.1548
357.1366
415.0639
431.0585
453.0397
G
4V WM Reactant
Y(Counts)
X(Thomsons)
335.1542
336.1573
337.1537
H
4V WM Brominated Product
Y(Counts)
X(Thomsons)
413.0648
415.0639

**4V WN**

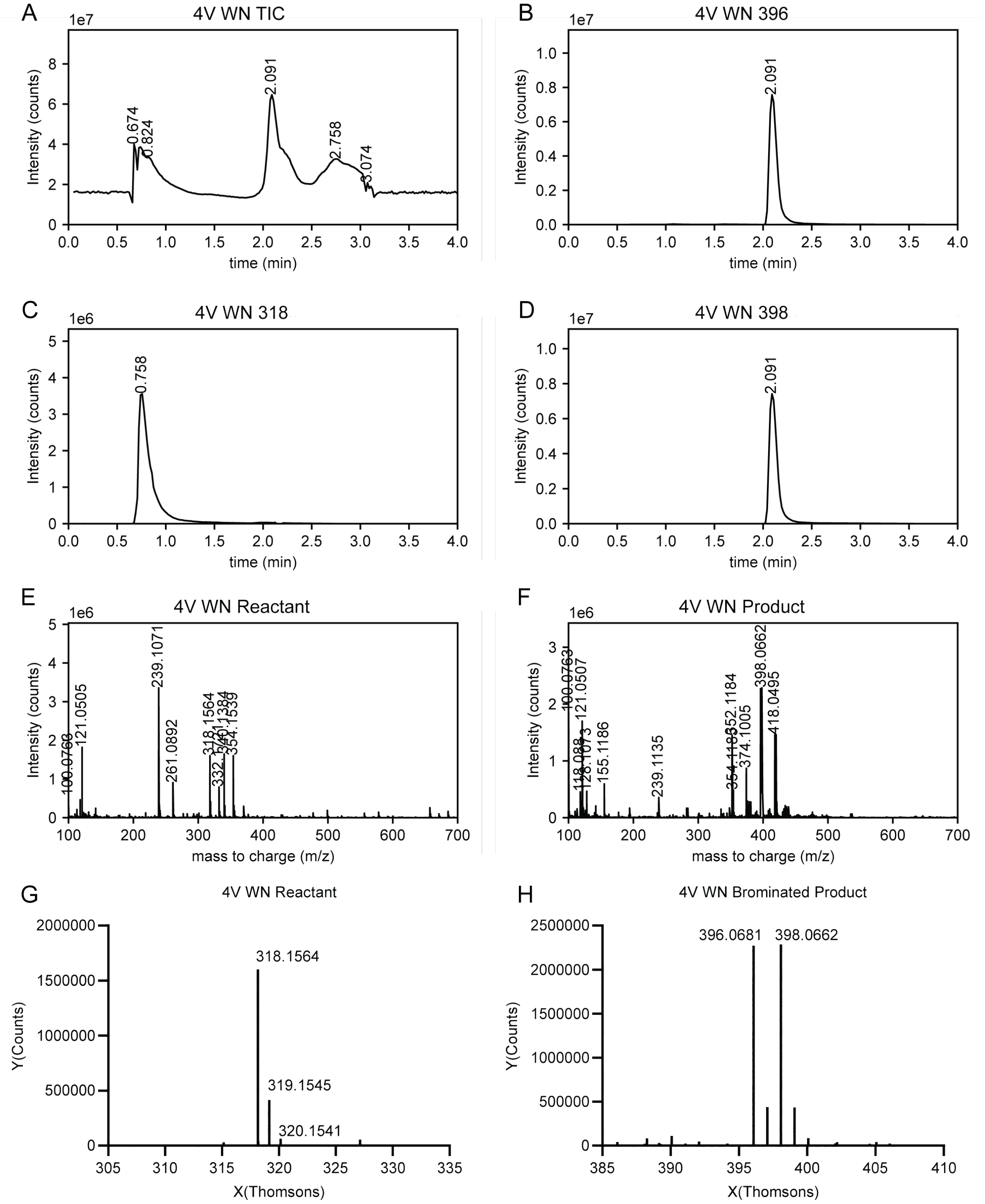
A
4V WN TIC
1e7
Intensity (counts)
time (min)
0.674
0.824
2.091
2.758
3.074
B
4V WN 396
1e7
Intensity (counts)
time (min)
2.091
C
4V WN 318
1e6
Intensity (counts)
time (min)
0.758
D
4V WN 398
1e7
Intensity (counts)
time (min)
2.091
E
4V WN Reactant
1e6
Intensity (counts)
mass to charge (m/z)
100.0763
121.0505
239.1071
261.0892
318.1564
354.1539
F
4V WN Product
1e6
Intensity (counts)
mass to charge (m/z)
100.0763
121.0507
155.1186
239.1135
352.1184
374.1005
398.0662
418.0495
G
4V WN Reactant
Y(Counts)
X(Thomsons)
318.1564
319.1545
320.1541
H
4V WN Brominated Product
Y(Counts)
X(Thomsons)
396.0681
398.0662

## 4V WP

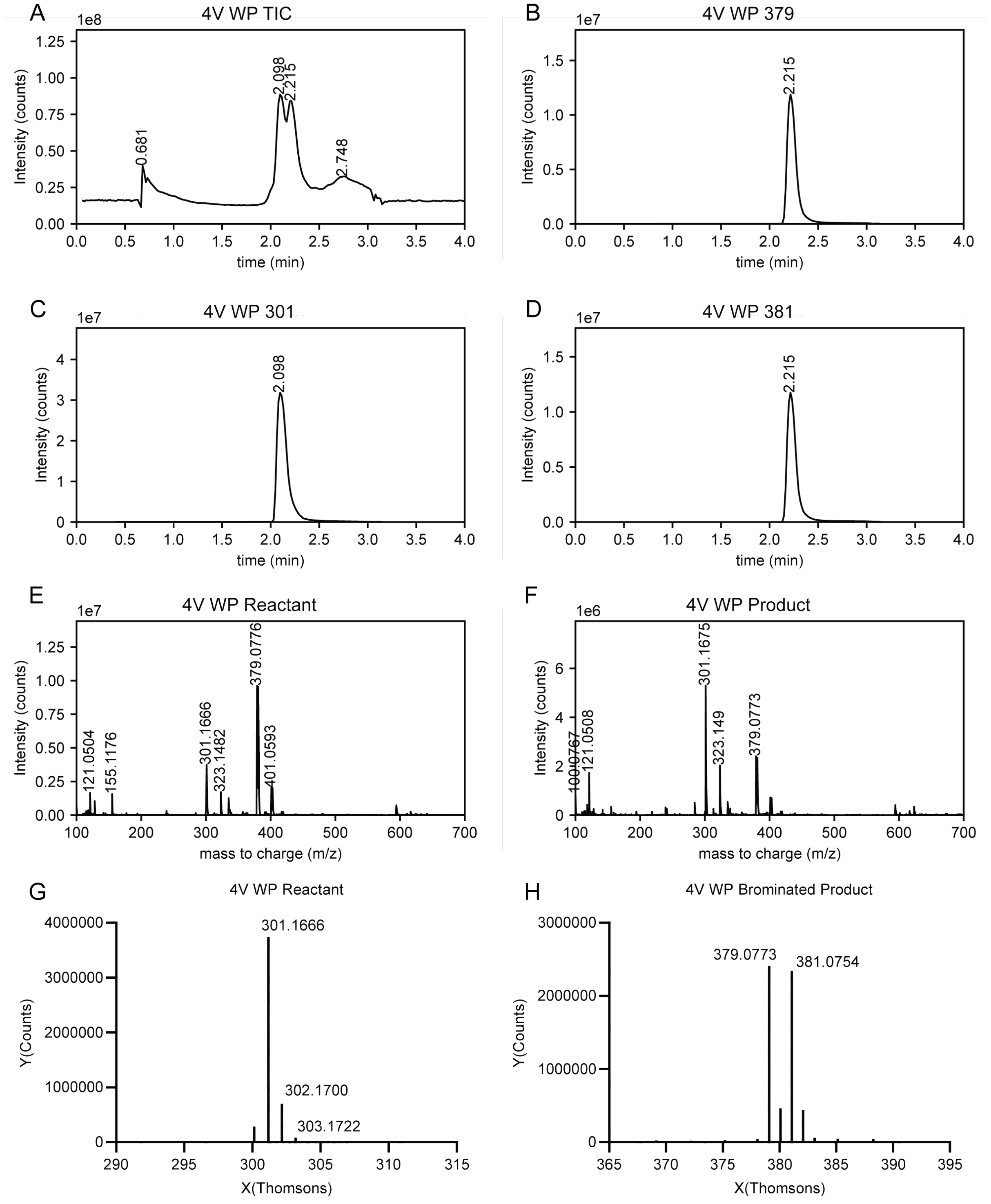

A
4V WP TIC
1e8
Intensity (counts)
time (min)
0.681
2.098
2.215
2.748
B
4V WP 379
1e7
Intensity (counts)
time (min)
2.215
C
4V WP 301
1e7
Intensity (counts)
time (min)
2.098
D
4V WP 381
1e7
Intensity (counts)
time (min)
2.215
E
4V WP Reactant
1e7
Intensity (counts)
mass to charge (m/z)
121.0504
155.1176
301.1666
323.1482
379.0776
401.0593
F
4V WP Product
1e6
Intensity (counts)
mass to charge (m/z)
121.0508
301.1675
323.149
379.0773
G
4V WP Reactant
Y(Counts)
X(Thomsons)
301.1666
302.1700
303.1722
H
4V WP Brominated Product
Y(Counts)
X(Thomsons)
379.0773
381.0754

## 4V WQ

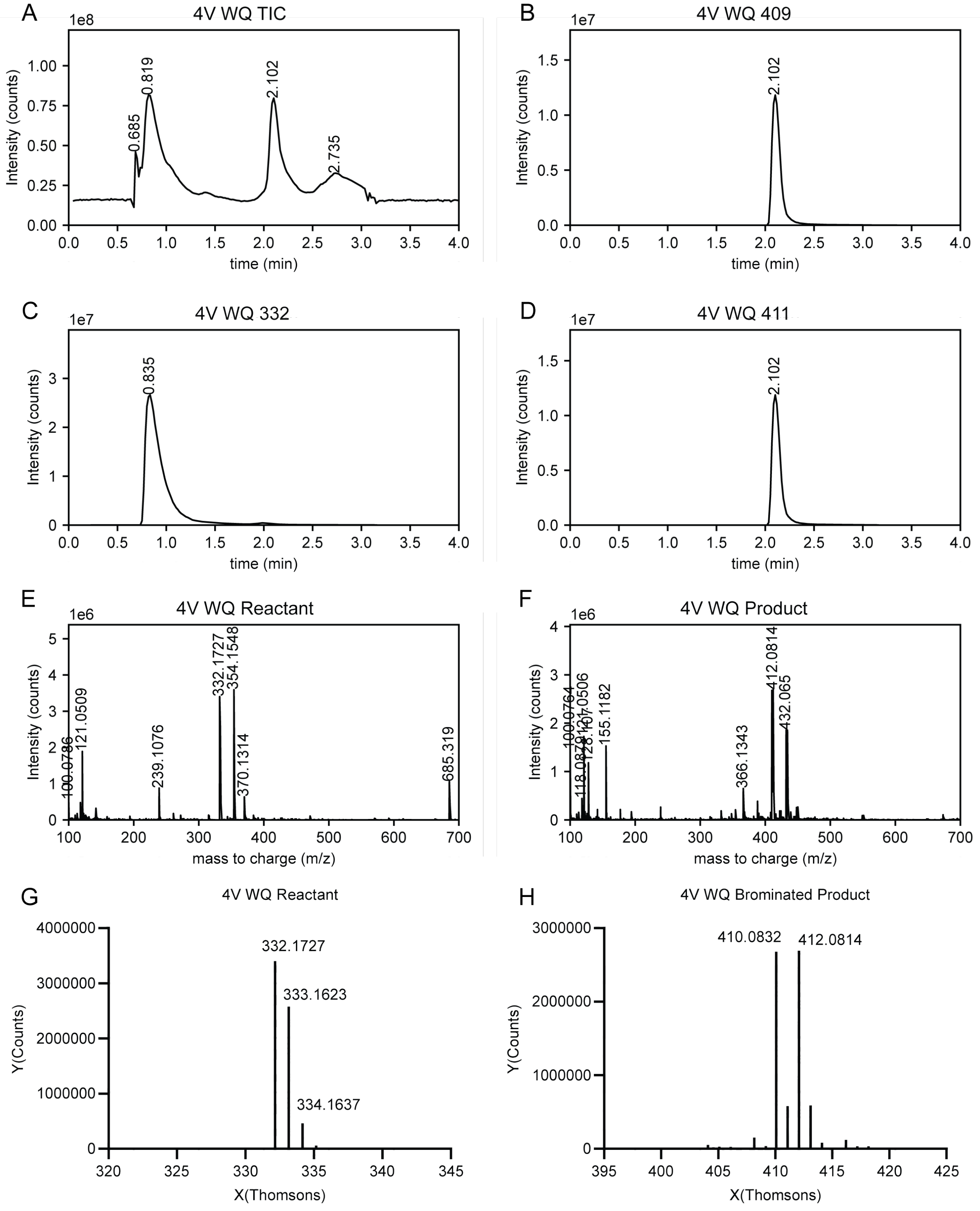

A
4V WQ TIC
1e8
Intensity (counts)
time (min)
0.685
0.819
2.102
2.735
B
4V WQ 409
1e7
Intensity (counts)
time (min)
2.102
C
4V WQ 332
1e7
Intensity (counts)
time (min)
0.835
D
4V WQ 411
1e7
Intensity (counts)
time (min)
2.102
E
4V WQ Reactant
1e6
Intensity (counts)
mass to charge (m/z)
121.0509
239.1076
332.1727
354.1548
370.1314
685.319
F
4V WQ Product
1e6
Intensity (counts)
mass to charge (m/z)
155.1182
366.1343
412.0814
432.065
G
4V WQ Reactant
Y(Counts)
X(Thomsons)
332.1727
333.1623
334.1637
H
4V WQ Brominated Product
Y(Counts)
X(Thomsons)
410.0832
412.0814

## 4V WR

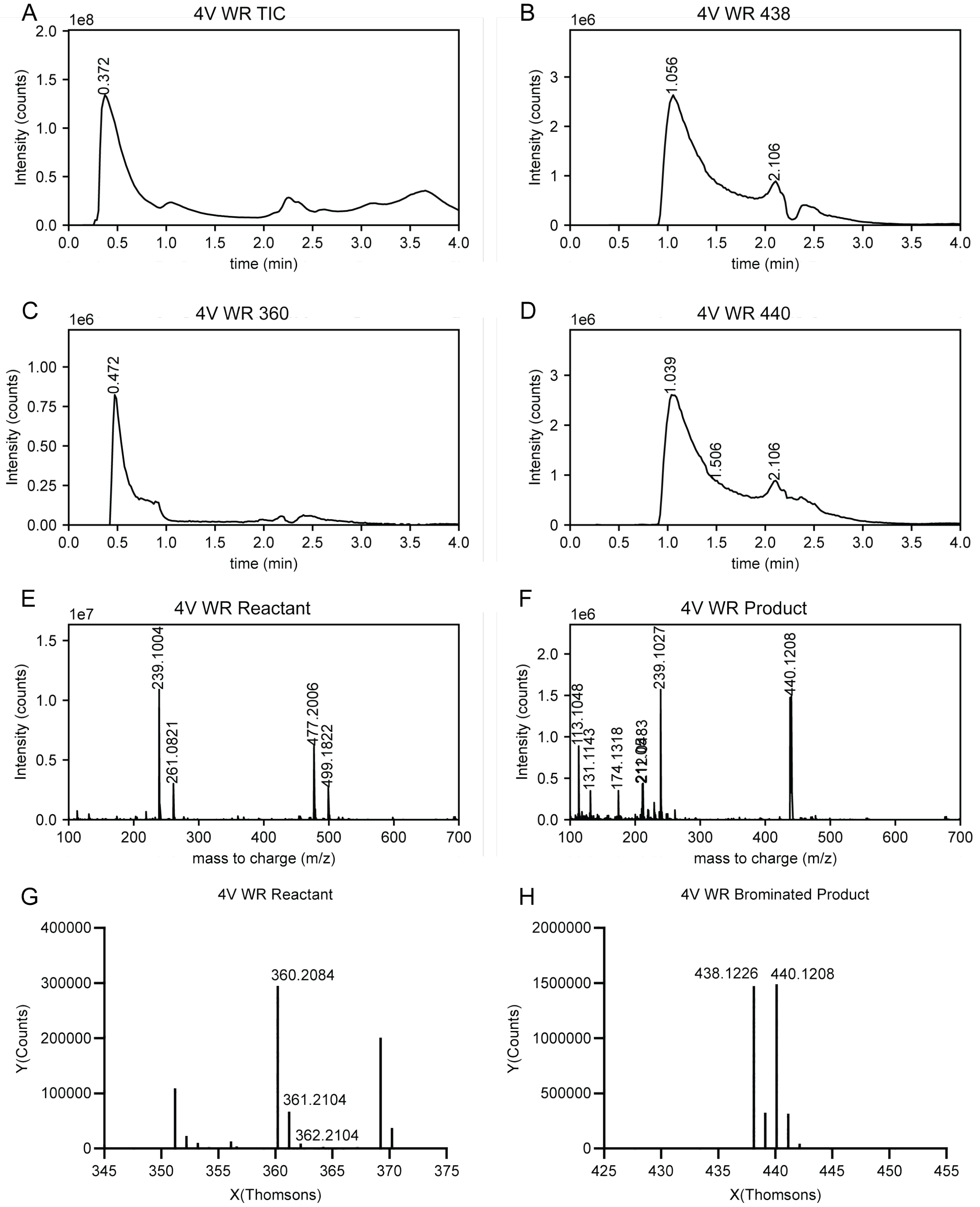
A
4V WR TIC
1e8
0.372
Intensity (counts)
time (min)
B
4V WR 438
1e6
1.056
2.106
Intensity (counts)
time (min)
C
4V WR 360
1e6
0.472
Intensity (counts)
time (min)
D
4V WR 440
1e6
1.039
1.506
2.106
Intensity (counts)
time (min)
E
4V WR Reactant
1e7
239.1004
261.0821
477.2006
499.1822
Intensity (counts)
mass to charge (m/z)
F
4V WR Product
1e6
113.1048
131.1143
174.1318
212.0483
239.1027
440.1208
Intensity (counts)
mass to charge (m/z)
G
4V WR Reactant
360.2084
361.2104
362.2104
Y(Counts)
X(Thomsons)
H
4V WR Brominated Product
438.1226
440.1208
Y(Counts)
X(Thomsons)

## 4V WS

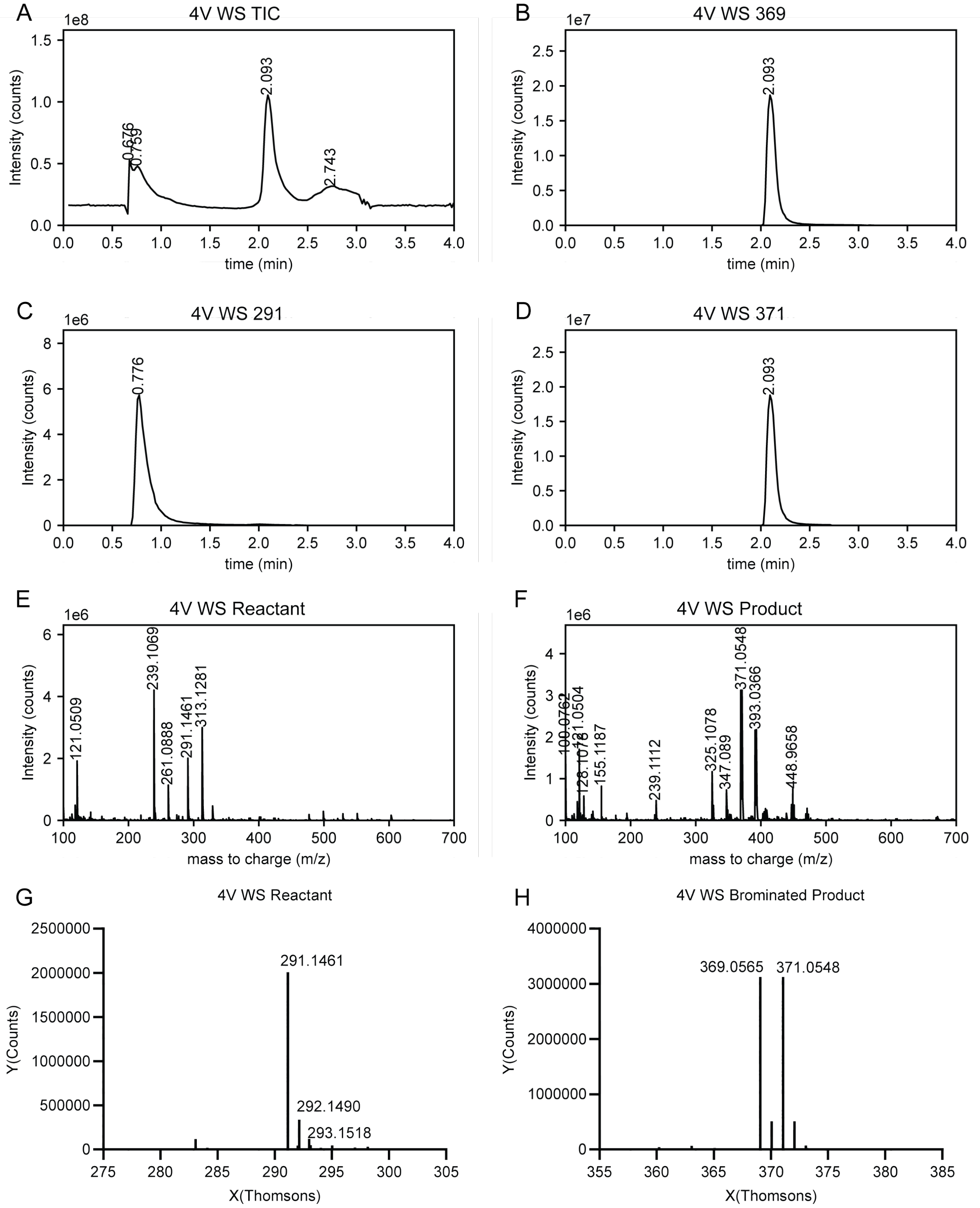

A
4V WS TIC
1e8
Intensity (counts)
time (min)
0.676
0.759
2.093
2.743
B
4V WS 369
1e7
Intensity (counts)
time (min)
2.093
C
4V WS 291
1e6
Intensity (counts)
time (min)
0.776
D
4V WS 371
1e7
Intensity (counts)
time (min)
2.093
E
4V WS Reactant
1e6
Intensity (counts)
mass to charge (m/z)
121.0509
239.1069
261.0888
291.1461
313.1281
F
4V WS Product
1e6
Intensity (counts)
mass to charge (m/z)
100.0762
121.0504
128.1078
155.1187
239.1112
325.1078
347.089
371.0548
393.0366
448.9658
G
4V WS Reactant
Y(Counts)
X(Thomsons)
291.1461
292.1490
293.1518
H
4V WS Brominated Product
Y(Counts)
X(Thomsons)
369.0565
371.0548

## 4V WT

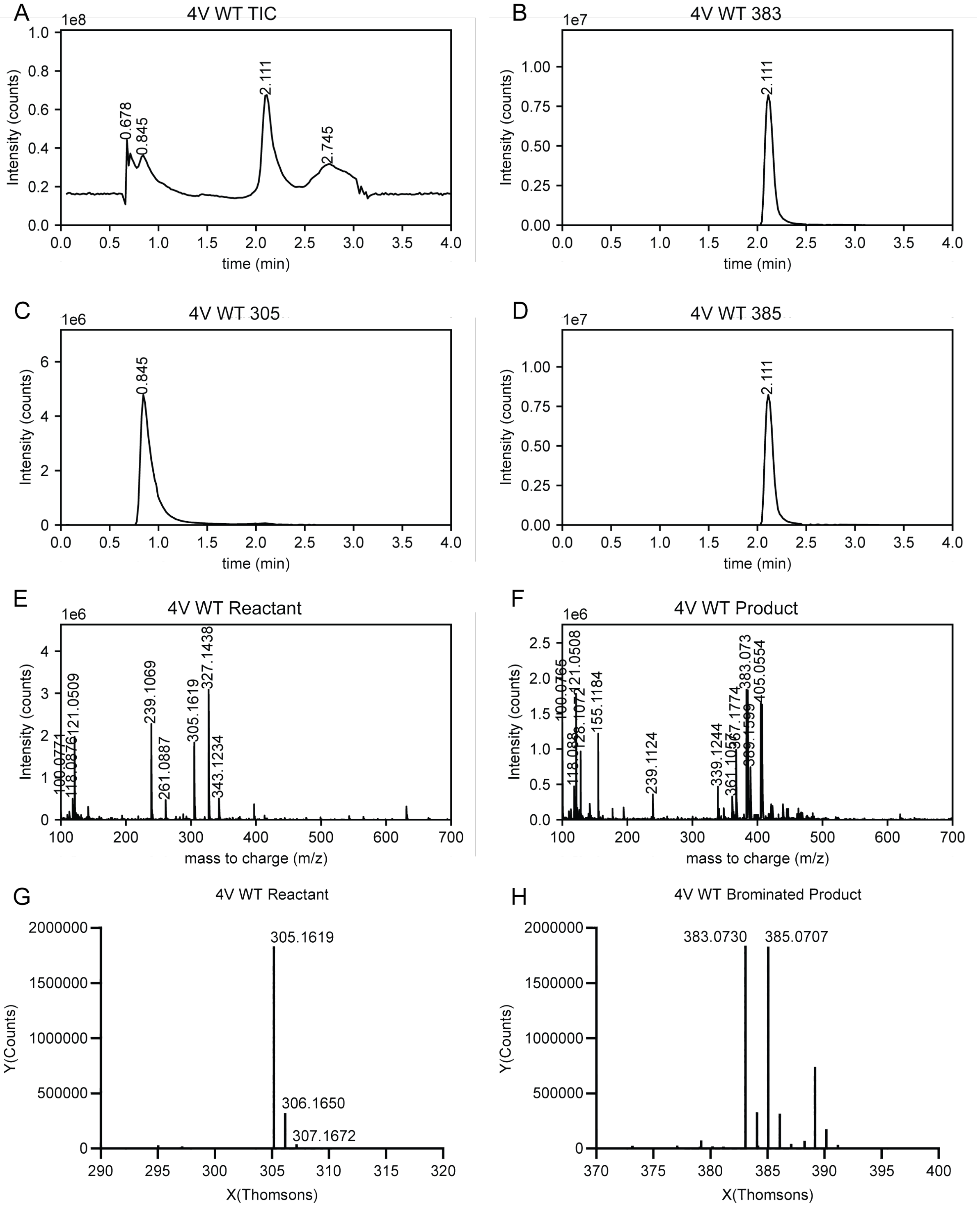
A
4V WT TIC
1e8
Intensity (counts)
time (min)
0.678
0.845
2.111
2.745
B
4V WT 383
1e7
Intensity (counts)
time (min)
2.111
C
4V WT 305
1e6
Intensity (counts)
time (min)
0.845
D
4V WT 385
1e7
Intensity (counts)
time (min)
2.111
E
4V WT Reactant
1e6
Intensity (counts)
mass to charge (m/z)
100.0771
118.0876
121.0509
239.1069
261.0887
305.1619
327.1438
343.1234
F
4V WT Product
1e6
Intensity (counts)
mass to charge (m/z)
100.0765
118.088
121.0508
128.1072
155.1184
239.1124
339.1244
361.1056
367.1774
383.073
405.0554
G
4V WT Reactant
Y(Counts)
X(Thomsons)
305.1619
306.1650
307.1672
H
4V WT Brominated Product
Y(Counts)
X(Thomsons)
383.0730
385.0707

## 4V WV

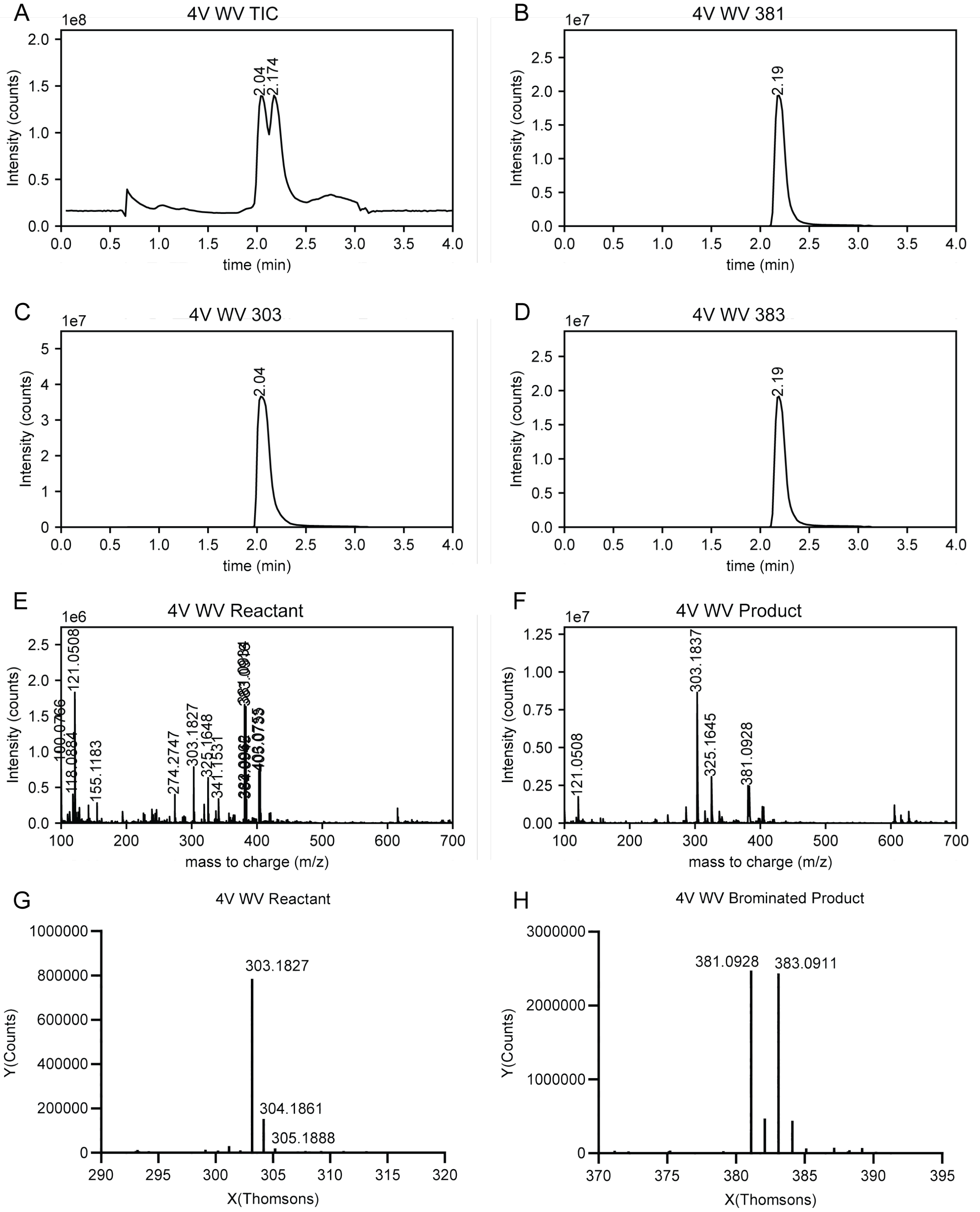
A
4V WV TIC
1e8
Intensity (counts)
time (min)
2.04
2.174
B
4V WV 381
1e7
Intensity (counts)
time (min)
2.19
C
4V WV 303
1e7
Intensity (counts)
time (min)
2.04
D
4V WV 383
1e7
Intensity (counts)
time (min)
2.19
E
4V WV Reactant
1e6
Intensity (counts)
mass to charge (m/z)
121.0508
100.0766
118.0884
155.1183
274.2747
303.1827
325.1648
341.1531
F
4V WV Product
1e7
Intensity (counts)
mass to charge (m/z)
303.1837
325.1645
381.0928
121.0508
G
4V WV Reactant
Y(Counts)
X(Thomsons)
303.1827
304.1861
305.1888
H
4V WV Brominated Product
Y(Counts)
X(Thomsons)
381.0928
383.0911

# 4V WY

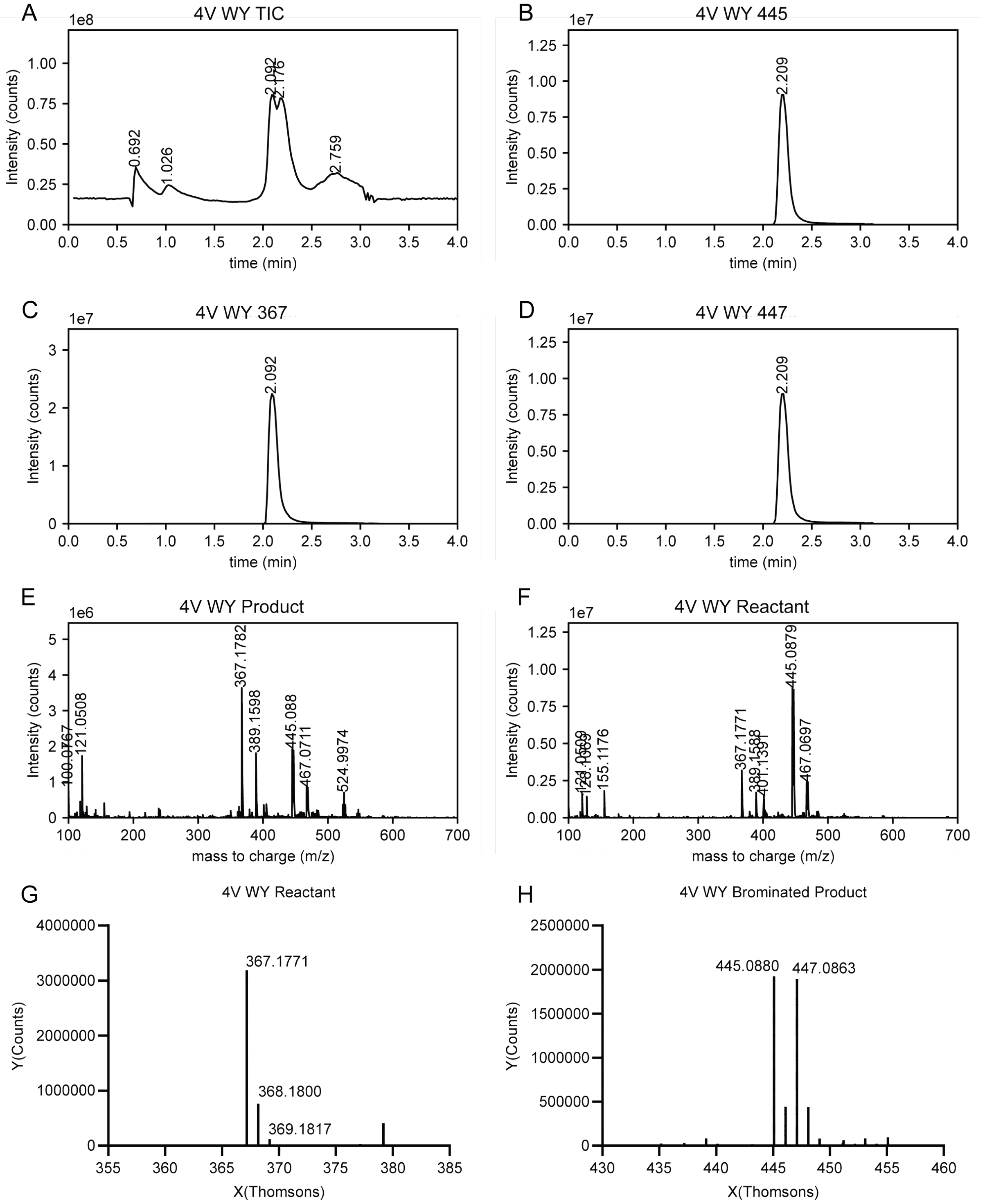
A
4V WY TIC
1e8
Intensity (counts)
time (min)
0.692
1.026
2.092
2.176
2.759
B
4V WY 445
1e7
2.209
C
4V WY 367
2.092
D
4V WY 447
2.209
E
4V WY Product
1e6
mass to charge (m/z)
100.0767
121.0508
367.1782
389.1598
445.088
467.0711
524.9974
F
4V WY Reactant
155.1176
367.1771
401.1391
445.0879
467.0697
G
4V WY Reactant
Y(Counts)
X(Thomsons)
367.1771
368.1800
369.1817
H
4V WY Brominated Product
445.0880
447.0863

#### RebH Brominated Tripeptide Mass Spectra

### RebH WAG

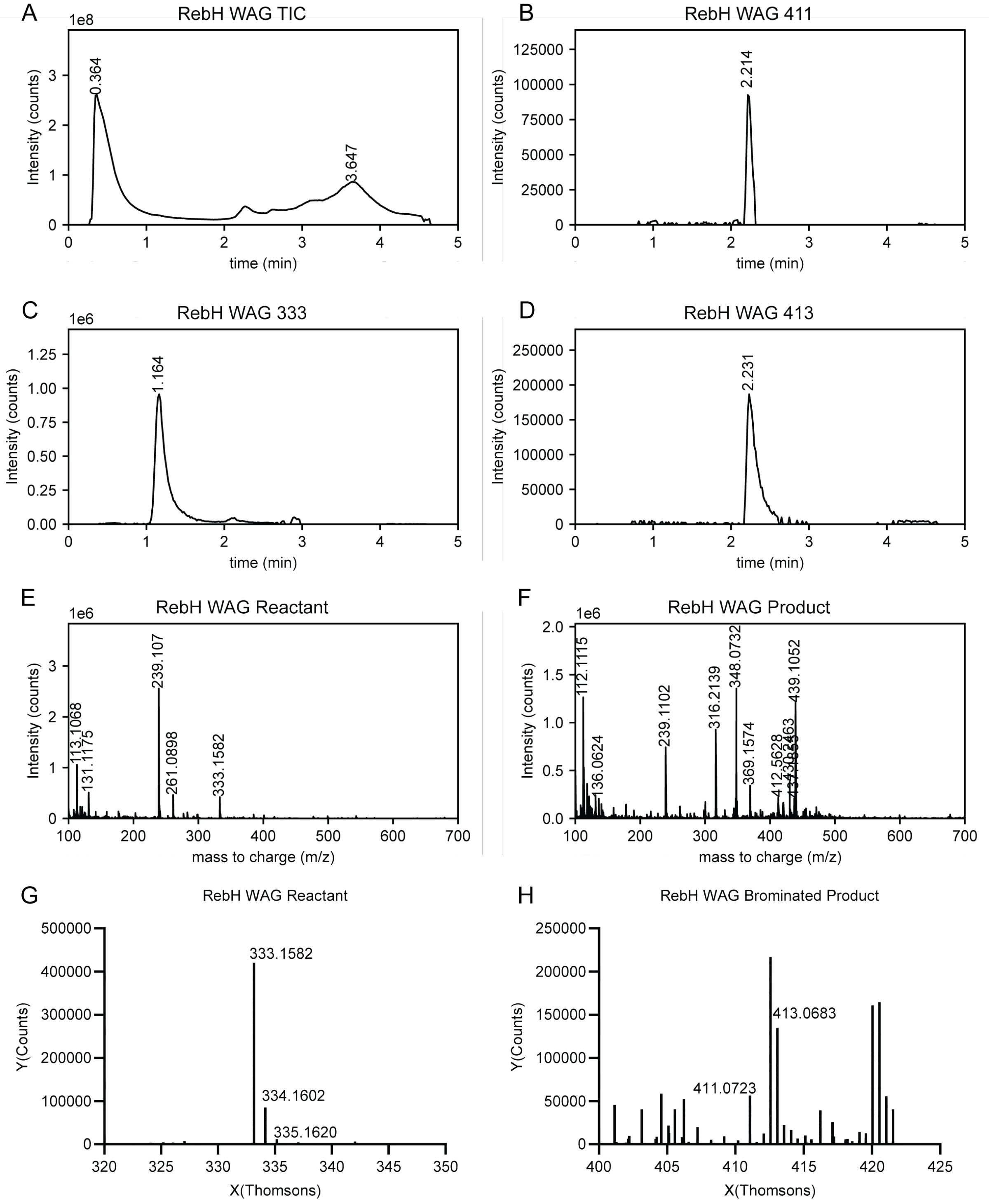

A
RebH WAG TIC
1e8
Intensity (counts)
time (min)
0.364
3.647
B
RebH WAG 411
Intensity (counts)
time (min)
2.214
C
RebH WAG 333
1e6
Intensity (counts)
time (min)
1.164
D
RebH WAG 413
Intensity (counts)
time (min)
2.231
E
RebH WAG Reactant
1e6
Intensity (counts)
mass to charge (m/z)
113.1068
131.1175
239.107
261.0898
333.1582
F
RebH WAG Product
1e6
Intensity (counts)
mass to charge (m/z)
112.1115
136.0624
239.1102
316.2139
348.0732
369.1574
412.5628
439.1052
G
RebH WAG Reactant
Y(Counts)
X(Thomsons)
333.1582
334.1602
335.1620
H
RebH WAG Brominated Product
Y(Counts)
X(Thomsons)
411.0723
413.0683

#### RebH WCG

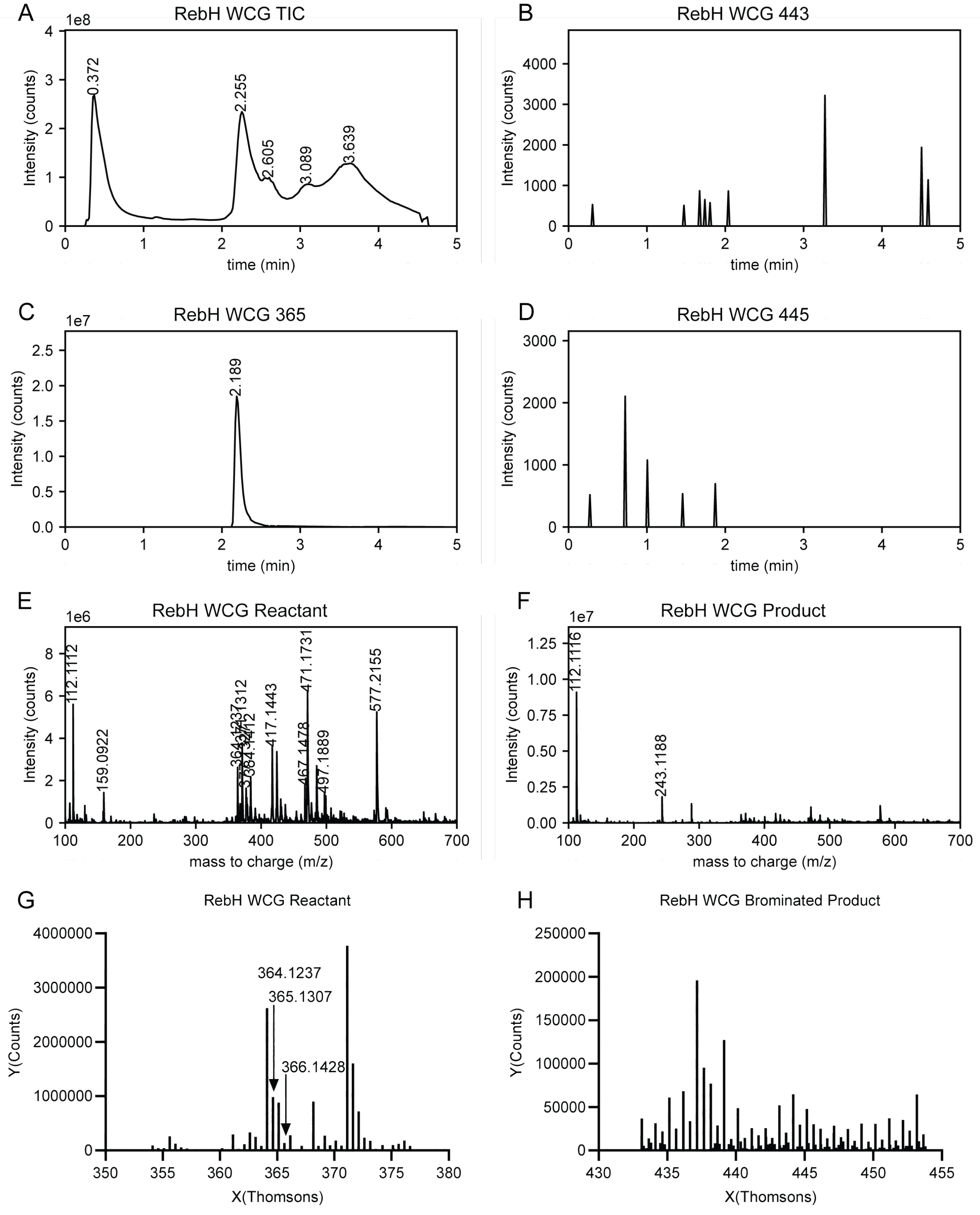
A
RebH WCG TIC
1e8
Intensity (counts)
time (min)
0.372
2.255
2.605
3.089
3.639
B
RebH WCG 443
Intensity (counts)
time (min)
C
RebH WCG 365
1e7
Intensity (counts)
time (min)
2.189
D
RebH WCG 445
Intensity (counts)
time (min)
E
RebH WCG Reactant
1e6
Intensity (counts)
mass to charge (m/z)
112.1112
159.0922
417.1443
471.1731
467.1478
497.1889
577.2155
F
RebH WCG Product
1e7
Intensity (counts)
mass to charge (m/z)
112.1116
243.1188
G
RebH WCG Reactant
Y(Counts)
X(Thomsons)
364.1237
365.1307
366.1428
H
RebH WCG Brominated Product
Y(Counts)
X(Thomsons)

#### RebH WDG

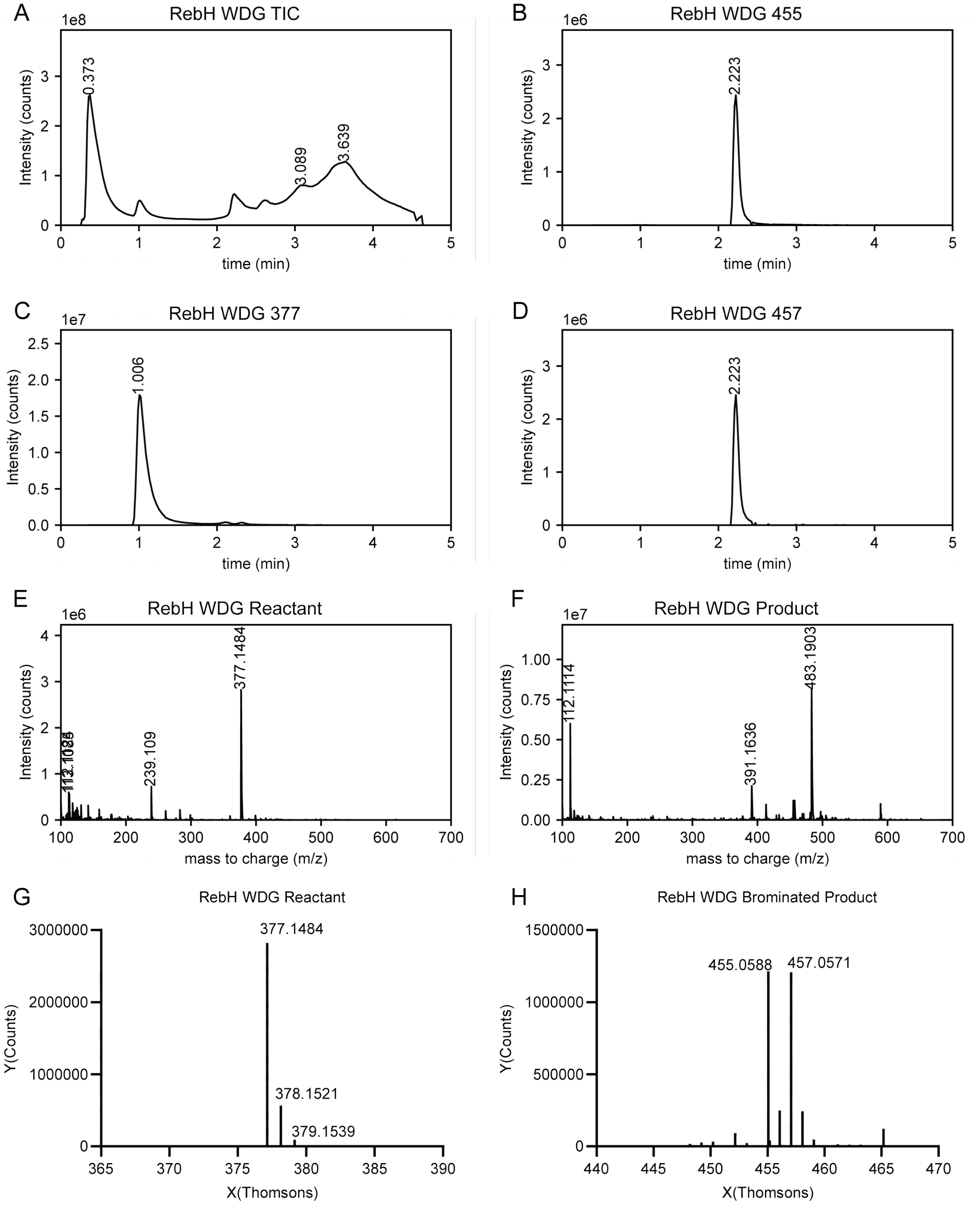

A
RebH WDG TIC
1e8
Intensity (counts)
time (min)
0.373
3.089
3.639
B
RebH WDG 455
1e6
Intensity (counts)
time (min)
2.223
C
RebH WDG 377
1e7
Intensity (counts)
time (min)
1.006
D
RebH WDG 457
1e6
Intensity (counts)
time (min)
2.223
E
RebH WDG Reactant
1e6
Intensity (counts)
mass to charge (m/z)
239.109
377.1484
F
RebH WDG Product
1e7
Intensity (counts)
mass to charge (m/z)
112.1114
391.1636
483.1903
G
RebH WDG Reactant
Y(Counts)
X(Thomsons)
377.1484
378.1521
379.1539
H
RebH WDG Brominated Product
Y(Counts)
X(Thomsons)
455.0588
457.0571

#### RebH WEG

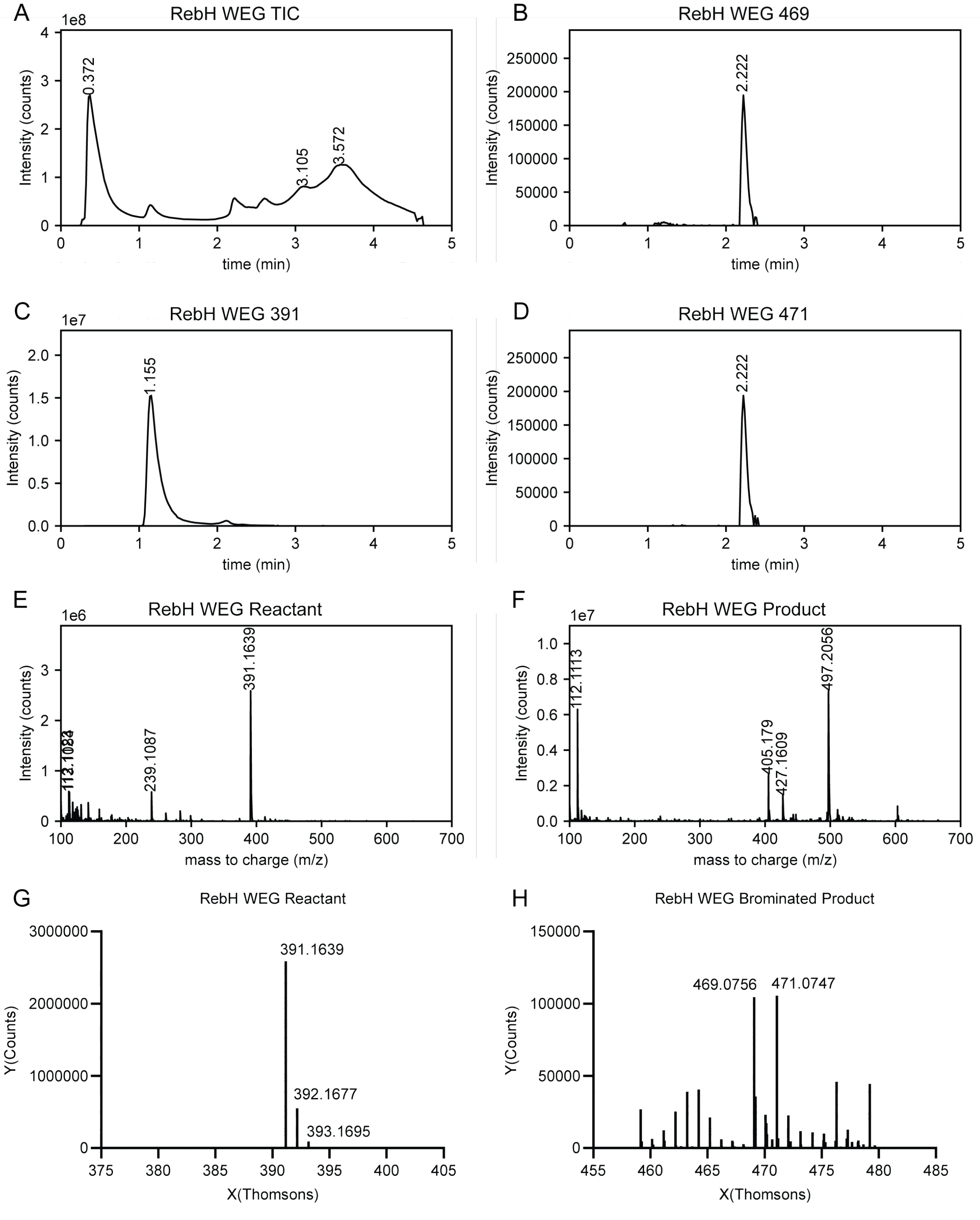

A
RebH WEG TIC
1e8
Intensity (counts)
time (min)
0.372
3.105
3.572
B
RebH WEG 469
Intensity (counts)
time (min)
2.222
C
RebH WEG 391
1e7
Intensity (counts)
time (min)
1.155
D
RebH WEG 471
Intensity (counts)
time (min)
2.222
E
RebH WEG Reactant
1e6
Intensity (counts)
mass to charge (m/z)
391.1639
239.1087
F
RebH WEG Product
1e7
Intensity (counts)
mass to charge (m/z)
112.1113
497.2056
405.179
427.1609
G
RebH WEG Reactant
Y(Counts)
X(Thomsons)
391.1639
392.1677
393.1695
H
RebH WEG Brominated Product
Y(Counts)
X(Thomsons)
469.0756
471.0747

#### RebH WFG

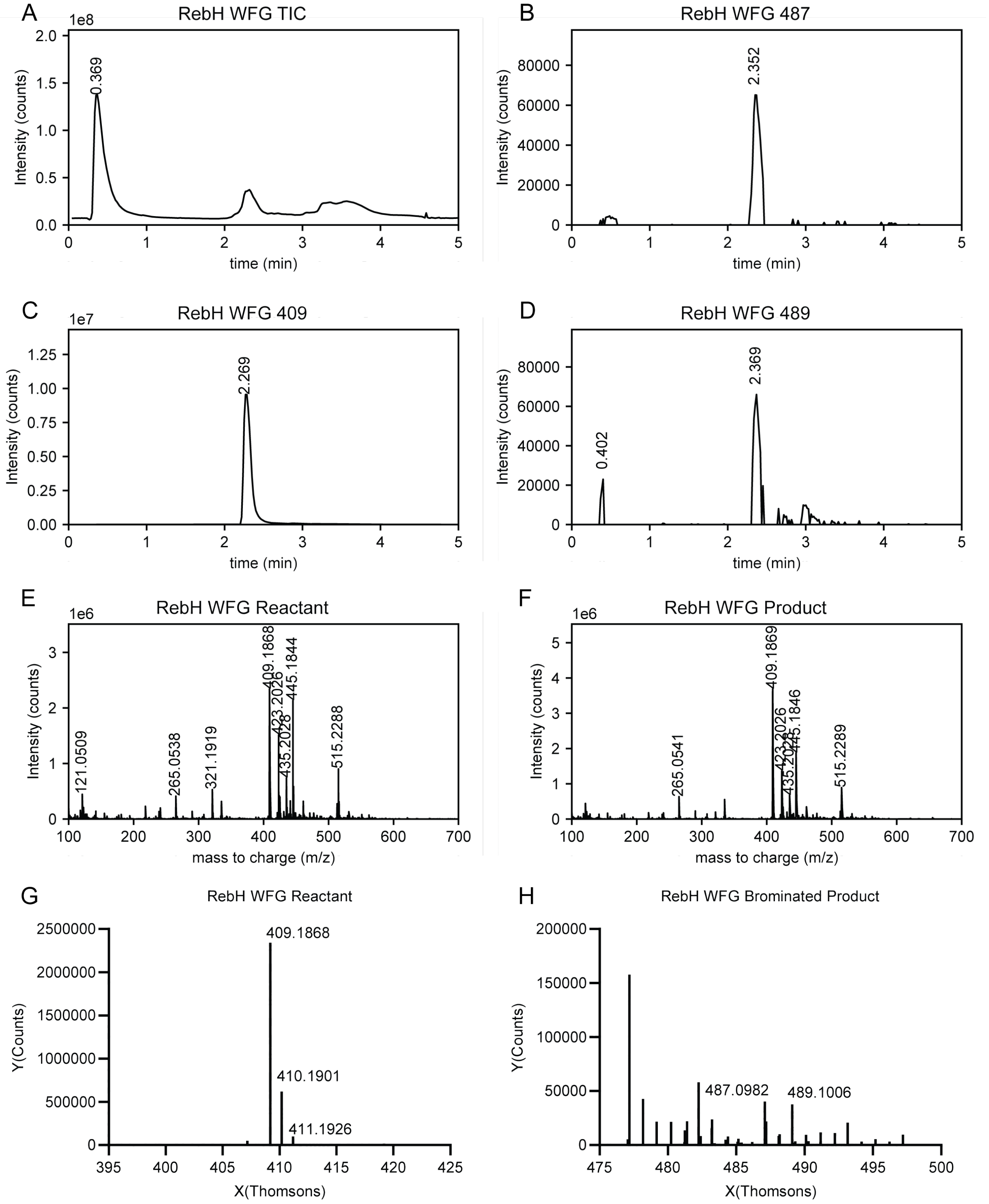

A
RebH WFG TIC
1e8
2.0
1.5
1.0
0.5
0.0
Intensity (counts)
0.369
time (min)
B
RebH WFG 487
80000
60000
40000
20000
0
Intensity (counts)
2.352
time (min)
C
RebH WFG 409
1e7
1.25
1.00
0.75
0.50
0.25
0.00
Intensity (counts)
2.269
time (min)
D
RebH WFG 489
80000
60000
40000
20000
0
Intensity (counts)
2.369
0.402
time (min)
0
1
2
3
4
5
E
RebH WFG Reactant
1e6
Intensity (counts)
121.0509
265.0538
321.1919
409.1868
423.2026
435.2028
445.1844
515.2288
mass to charge (m/z)
F
RebH WFG Product
1e6
Intensity (counts)
265.0541
409.1869
423.2026
435.2028
445.1846
515.2289
mass to charge (m/z)
100
200
300
400
500
600
700
G
RebH WFG Reactant
2500000
2000000
1500000
1000000
500000
0
Y(Counts)
409.1868
410.1901
411.1926
395
400
405
410
415
420
425
X(Thomsons)
H
RebH WFG Brominated Product
200000
150000
100000
50000
0
Y(Counts)
487.0982
489.1006
475
480
485
490
495
500
X(Thomsons)

#### RebH WHG

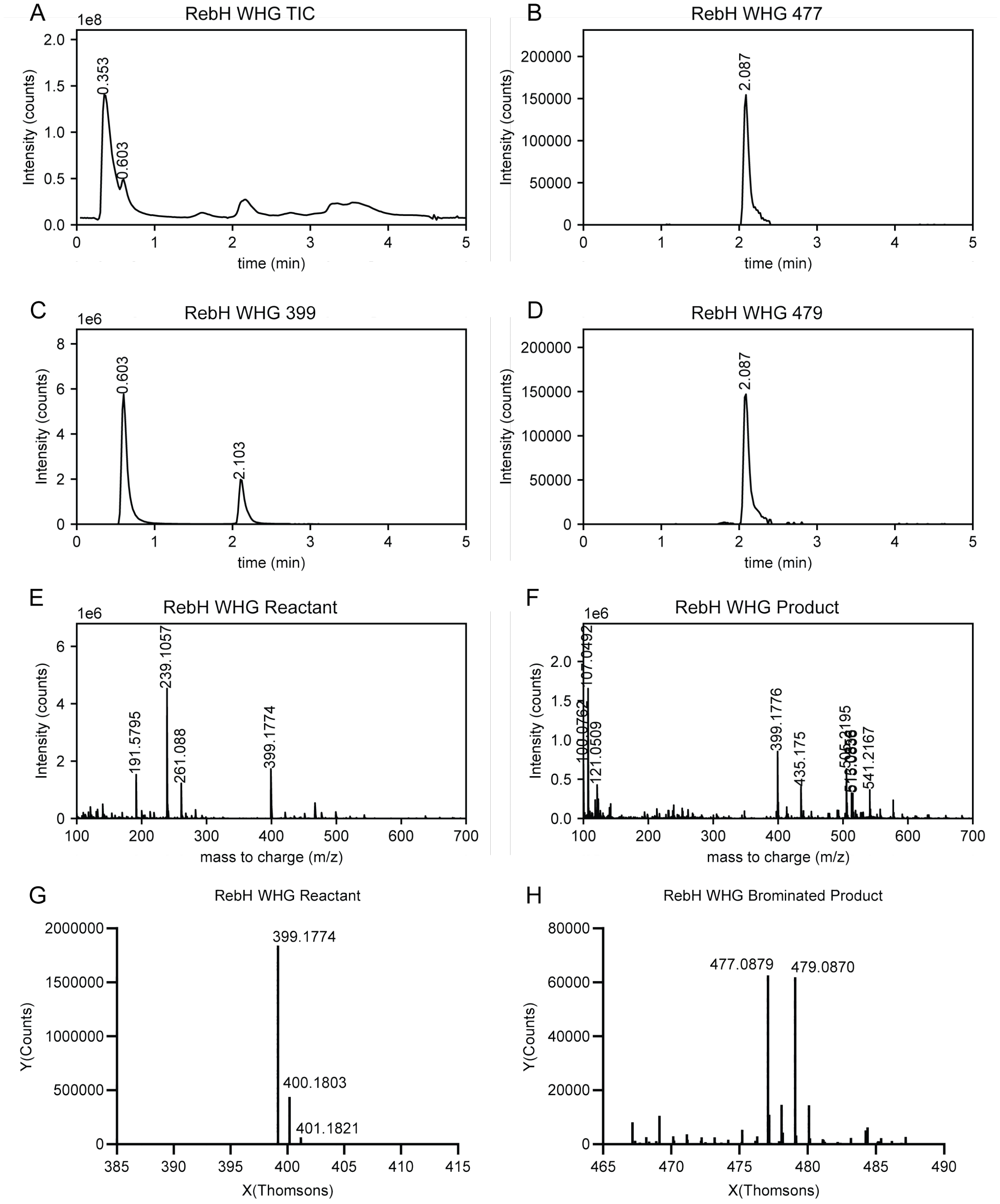

A
RebH WHG TIC
1e8
Intensity (counts)
time (min)
0.353
0.603
B
RebH WHG 477
Intensity (counts)
time (min)
2.087
C
RebH WHG 399
1e6
Intensity (counts)
time (min)
0.603
2.103
D
RebH WHG 479
Intensity (counts)
time (min)
2.087
E
RebH WHG Reactant
1e6
Intensity (counts)
mass to charge (m/z)
191.5795
239.1057
261.088
399.1774
F
RebH WHG Product
1e6
Intensity (counts)
mass to charge (m/z)
121.0509
399.1776
435.175
541.2167
G
RebH WHG Reactant
Y(Counts)
X(Thomsons)
399.1774
400.1803
401.1821
H
RebH WHG Brominated Product
Y(Counts)
X(Thomsons)
477.0879
479.0870

#### RebH WKG

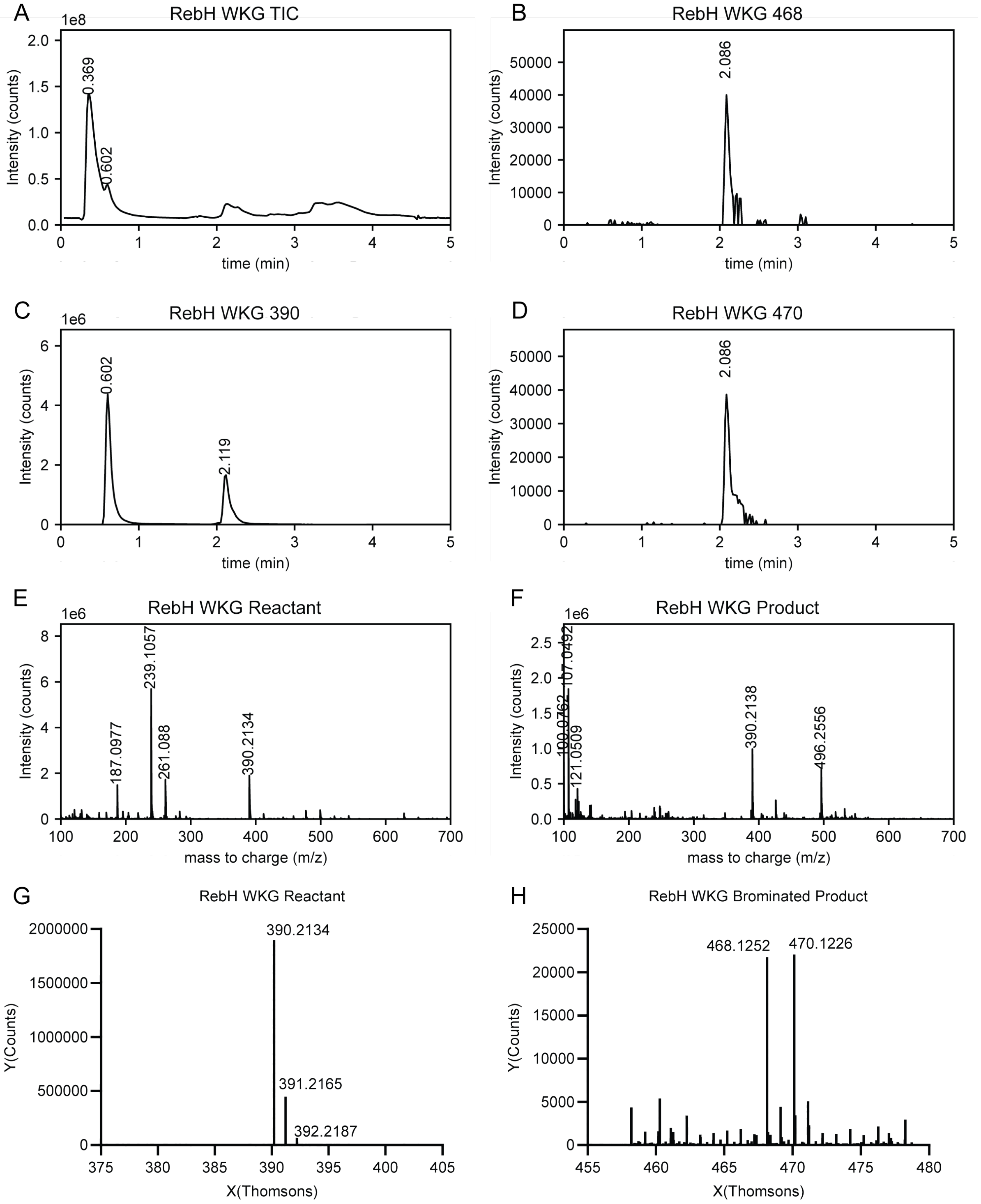

A
RebH WKG TIC
1e8
Intensity (counts)
time (min)
0.369
0.602
B
RebH WKG 468
Intensity (counts)
time (min)
2.086
C
RebH WKG 390
1e6
Intensity (counts)
time (min)
0.602
2.119
D
RebH WKG 470
Intensity (counts)
time (min)
2.086
E
RebH WKG Reactant
1e6
Intensity (counts)
mass to charge (m/z)
187.0977
239.1057
261.088
390.2134
F
RebH WKG Product
1e6
Intensity (counts)
mass to charge (m/z)
107.0492
121.0509
390.2138
496.2556
G
RebH WKG Reactant
Y(Counts)
X(Thomsons)
390.2134
391.2165
392.2187
H
RebH WKG Brominated Product
Y(Counts)
X(Thomsons)
468.1252
470.1226

#### RebH WLG

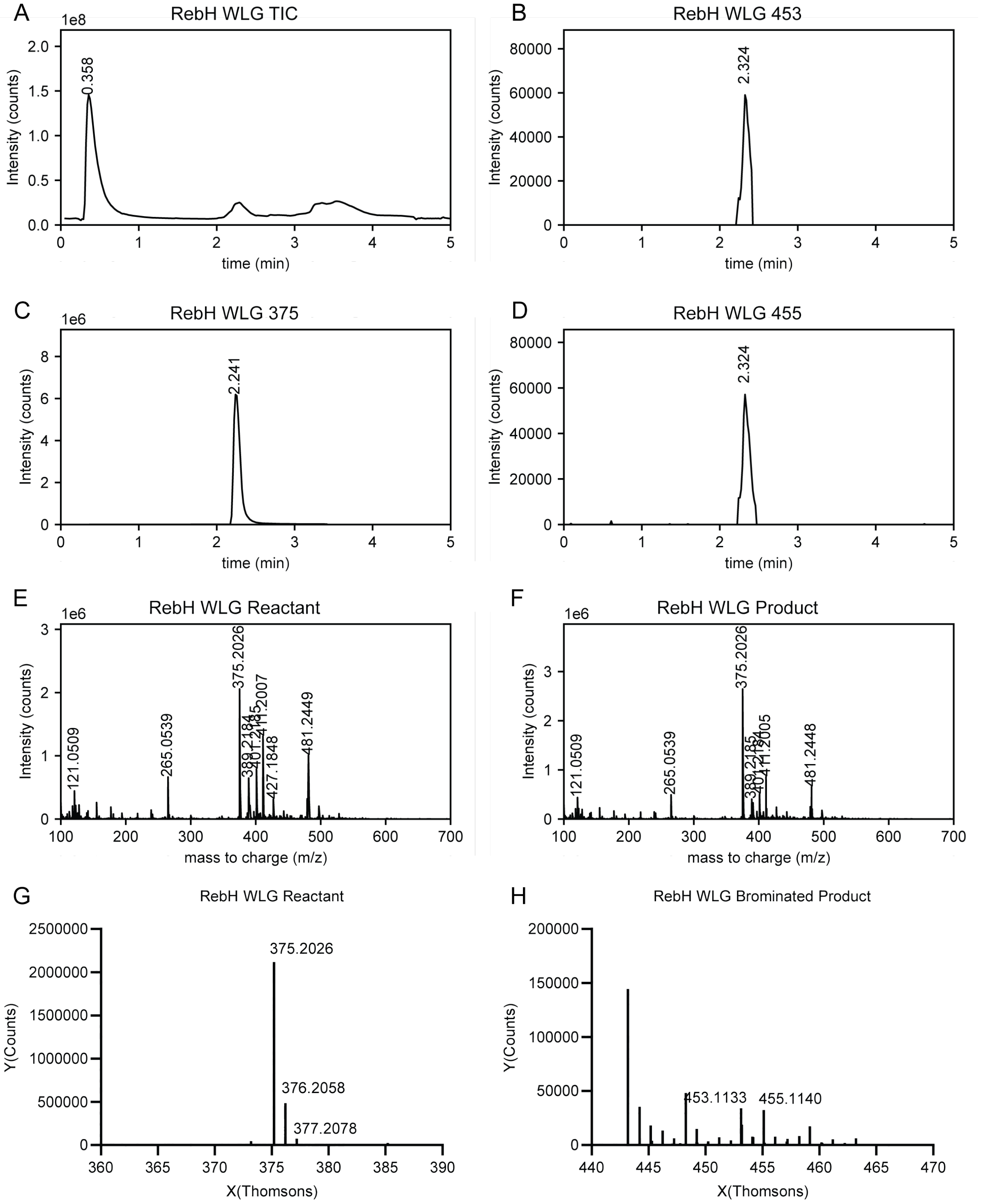

A
RebH WLG TIC
1e8
0.358
Intensity (counts)
time (min)
B
RebH WLG 453
2.324
Intensity (counts)
time (min)
C
RebH WLG 375
1e6
2.241
Intensity (counts)
time (min)
D
RebH WLG 455
2.324
Intensity (counts)
time (min)
E
RebH WLG Reactant
1e6
121.0509
265.0539
375.2026
389.2184
427.1848
481.2449
Intensity (counts)
mass to charge (m/z)
F
RebH WLG Product
1e6
121.0509
265.0539
375.2026
481.2448
Intensity (counts)
mass to charge (m/z)
G
RebH WLG Reactant
375.2026
376.2058
377.2078
Y(Counts)
X(Thomsons)
H
RebH WLG Brominated Product
453.1133
455.1140
Y(Counts)
X(Thomsons)

#### RebH WMG

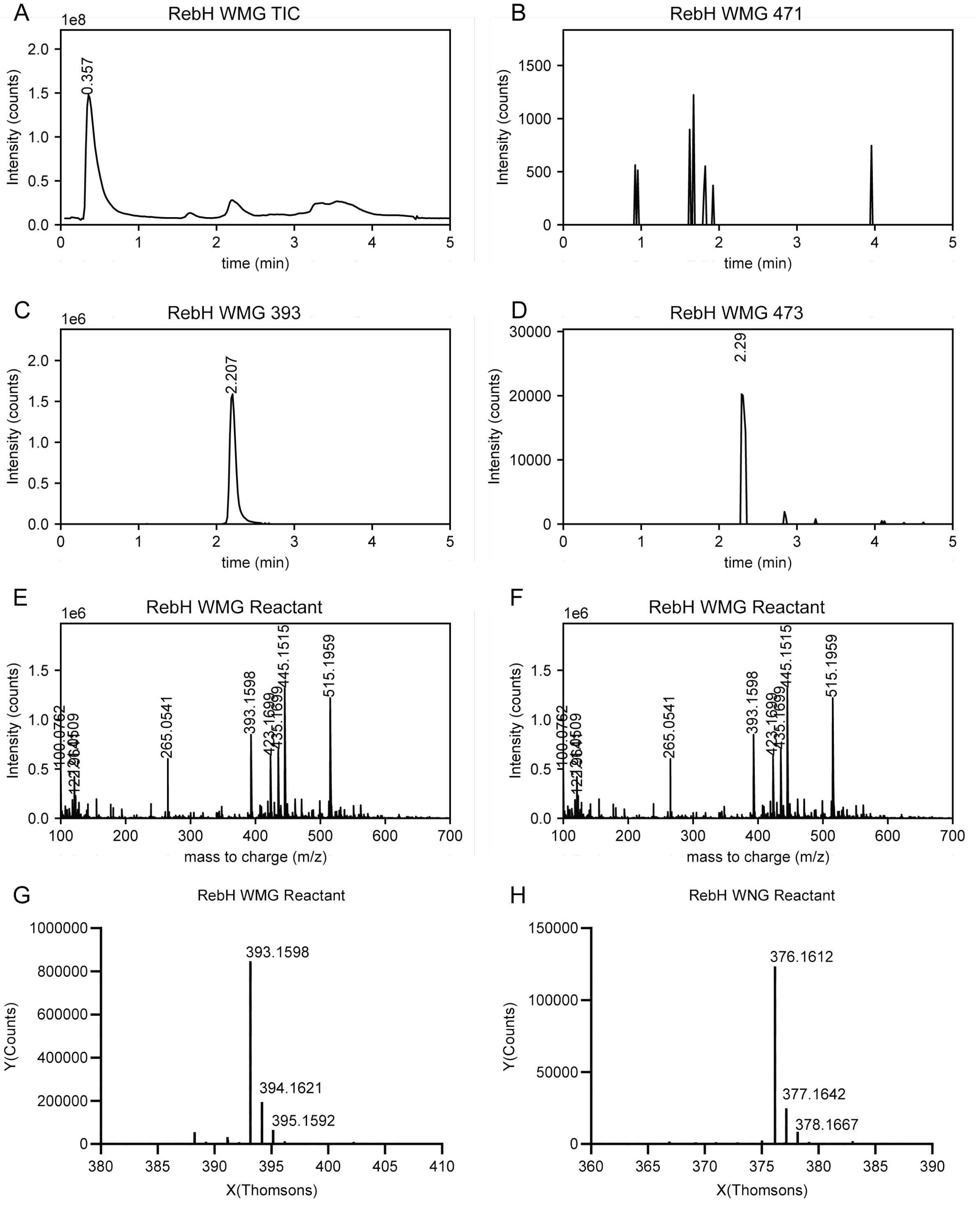

A
RebH WMG TIC
1e8
0.357
Intensity (counts)
time (min)
B
RebH WMG 471
Intensity (counts)
time (min)
C
RebH WMG 393
1e6
2.207
Intensity (counts)
time (min)
D
RebH WMG 473
2.29
Intensity (counts)
time (min)
E
RebH WMG Reactant
1e6
100.0762
265.0541
393.1598
423.1699
435.1699
445.1515
515.1959
Intensity (counts)
mass to charge (m/z)
F
RebH WMG Reactant
1e6
100.0762
265.0541
393.1598
423.1699
435.1699
445.1515
515.1959
Intensity (counts)
mass to charge (m/z)
G
RebH WMG Reactant
393.1598
394.1621
395.1592
Y(Counts)
X(Thomsons)
H
RebH WNG Reactant
376.1612
377.1642
378.1667
Y(Counts)
X(Thomsons)

#### RebH WNG

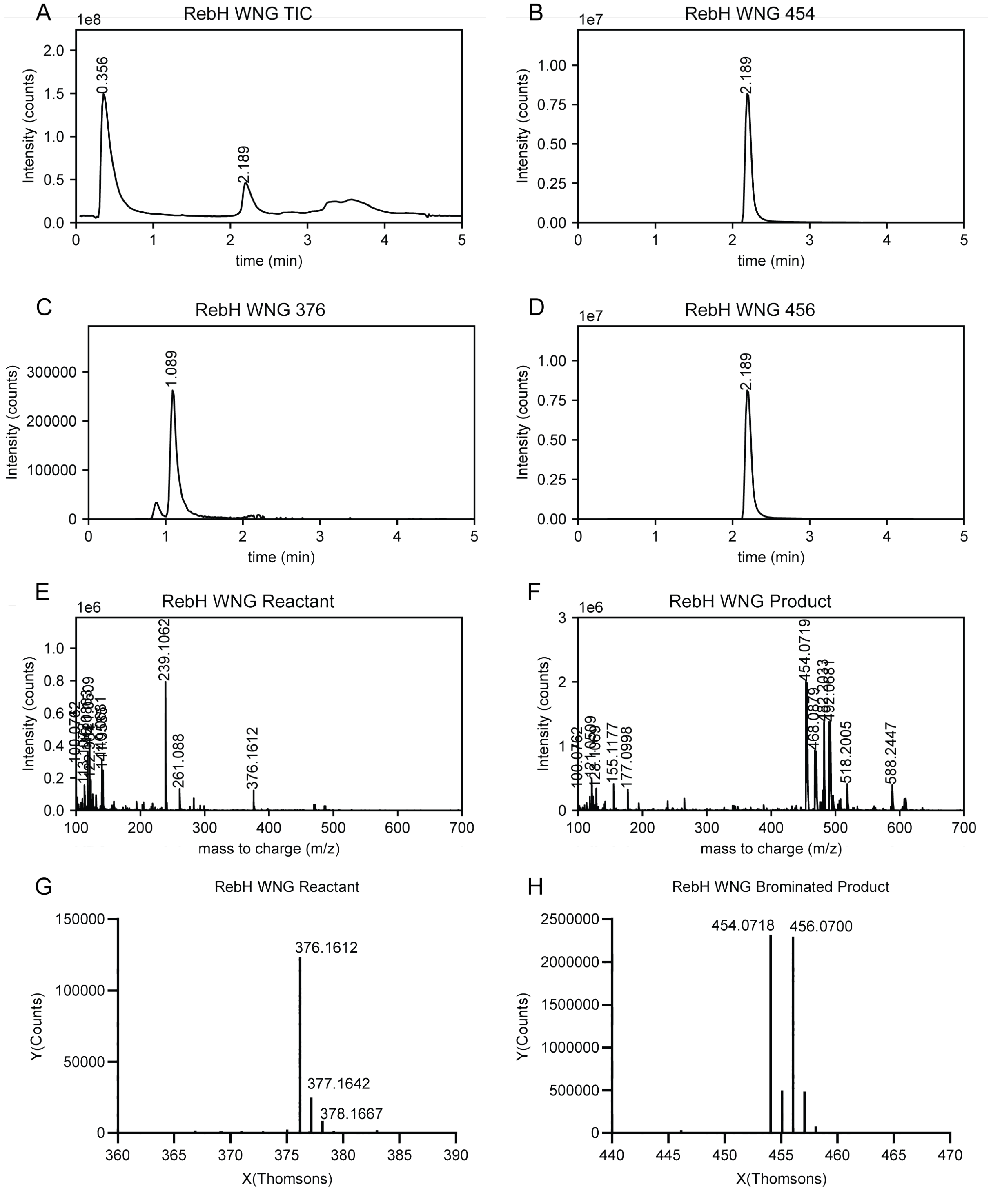

A
RebH WNG TIC
1e8
Intensity (counts)
time (min)
0.356
2.189
B
RebH WNG 454
1e7
Intensity (counts)
time (min)
2.189
C
RebH WNG 376
Intensity (counts)
time (min)
1.089
D
RebH WNG 456
1e7
Intensity (counts)
time (min)
2.189
E
RebH WNG Reactant
1e6
Intensity (counts)
mass to charge (m/z)
239.1062
261.088
376.1612
F
RebH WNG Product
1e6
Intensity (counts)
mass to charge (m/z)
454.0719
155.1177
177.0998
518.2005
588.2447
G
RebH WNG Reactant
Y(Counts)
X(Thomsons)
376.1612
377.1642
378.1667
H
RebH WNG Brominated Product
Y(Counts)
X(Thomsons)
454.0718
456.0700

#### RebH WPG

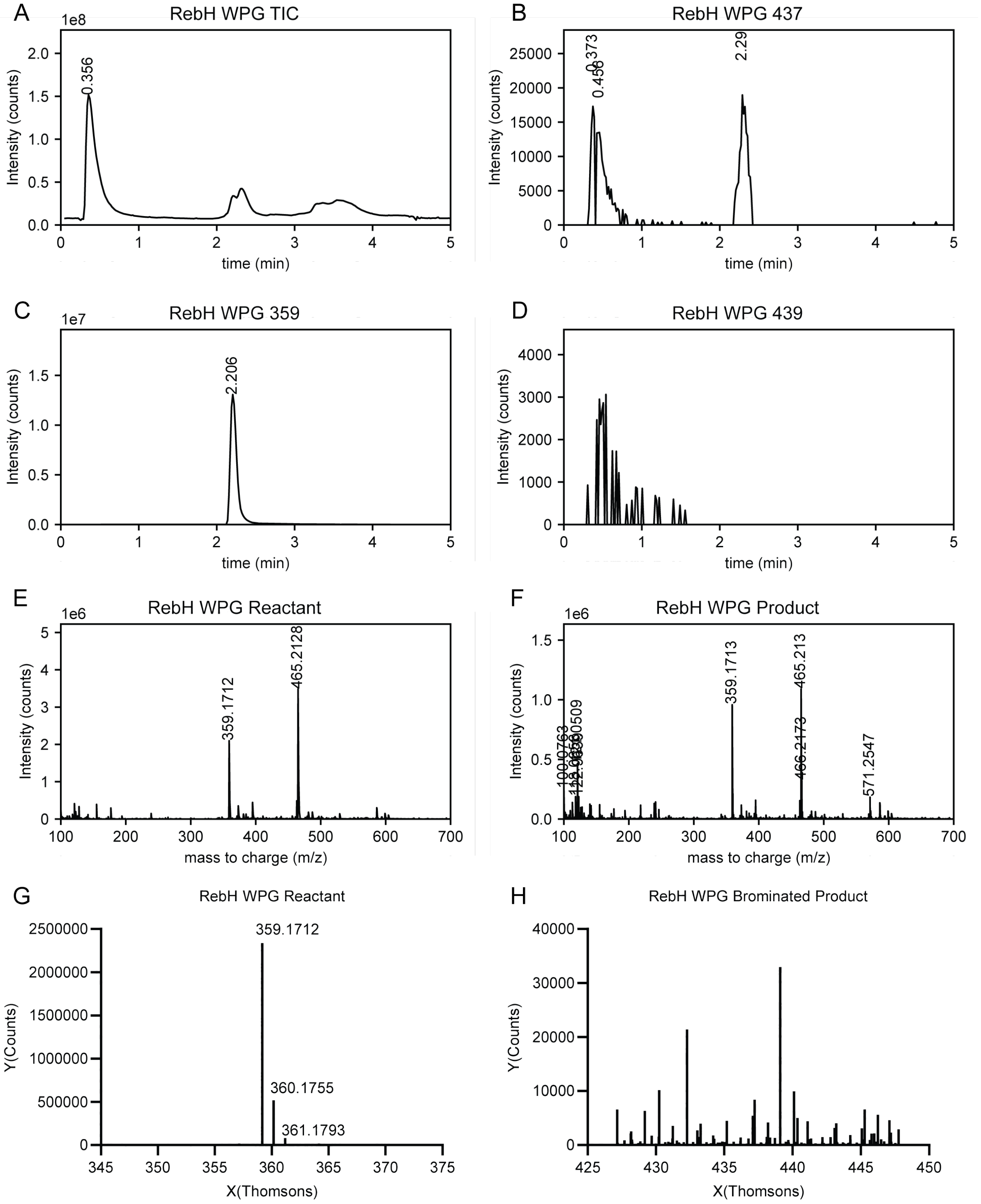

A
RebH WPG TIC
1e8
Intensity (counts)
time (min)
0.356
B
RebH WPG 437
Intensity (counts)
time (min)
0.373
0.456
2.29
C
RebH WPG 359
1e7
Intensity (counts)
time (min)
2.206
D
RebH WPG 439
Intensity (counts)
time (min)
E
RebH WPG Reactant
1e6
Intensity (counts)
mass to charge (m/z)
359.1712
465.2128
F
RebH WPG Product
1e6
Intensity (counts)
mass to charge (m/z)
359.1713
465.213
466.2173
571.2547
G
RebH WPG Reactant
Y(Counts)
X(Thomsons)
359.1712
360.1755
361.1793
H
RebH WPG Brominated Product
Y(Counts)
X(Thomsons)

#### RebH WQG

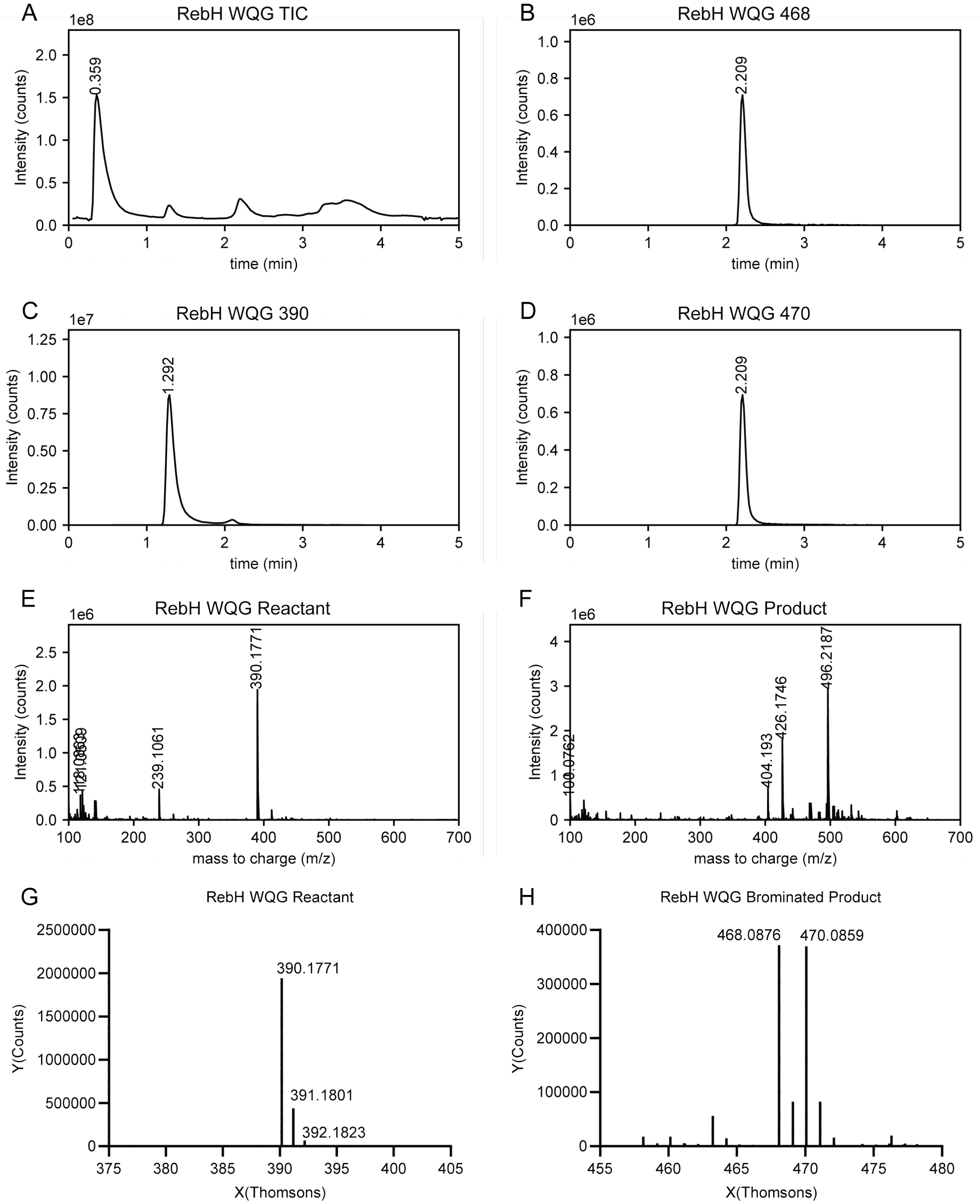
A
RebH WQG TIC
1e8
0.359
Intensity (counts)
time (min)
B
RebH WQG 468
1e6
2.209
Intensity (counts)
time (min)
C
RebH WQG 390
1e7
1.292
Intensity (counts)
time (min)
D
RebH WQG 470
1e6
2.209
Intensity (counts)
time (min)
E
RebH WQG Reactant
1e6
390.1771
239.1061
Intensity (counts)
mass to charge (m/z)
F
RebH WQG Product
1e6
496.2187
426.1746
404.193
Intensity (counts)
mass to charge (m/z)
G
RebH WQG Reactant
390.1771
391.1801
392.1823
Y(Counts)
X(Thomsons)
H
RebH WQG Brominated Product
468.0876
470.0859
Y(Counts)
X(Thomsons)

#### RebH WRG

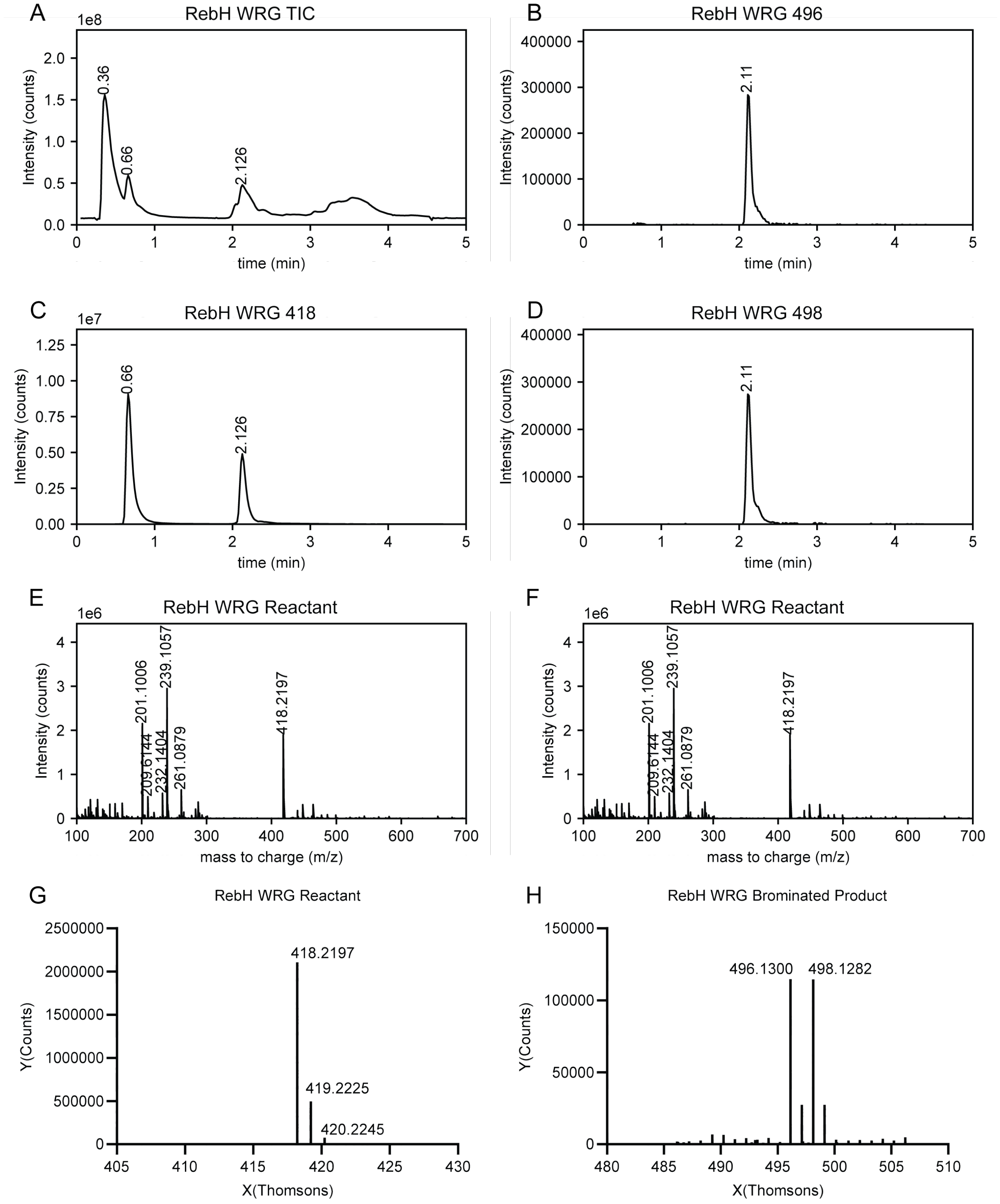

A
RebH WRG TIC
1e8
Intensity (counts)
time (min)
0.36
0.66
2.126
B
RebH WRG 496
Intensity (counts)
time (min)
2.11
C
RebH WRG 418
1e7
Intensity (counts)
time (min)
0.66
2.126
D
RebH WRG 498
Intensity (counts)
time (min)
2.11
E
RebH WRG Reactant
1e6
Intensity (counts)
mass to charge (m/z)
201.1006
209.6144
232.1404
239.1057
261.0879
418.2197
F
RebH WRG Reactant
1e6
Intensity (counts)
mass to charge (m/z)
201.1006
209.6144
232.1404
239.1057
261.0879
418.2197
G
RebH WRG Reactant
Y(Counts)
X(Thomsons)
418.2197
419.2225
420.2245
H
RebH WRG Brominated Product
Y(Counts)
X(Thomsons)
496.1300
498.1282

#### RebH WTG

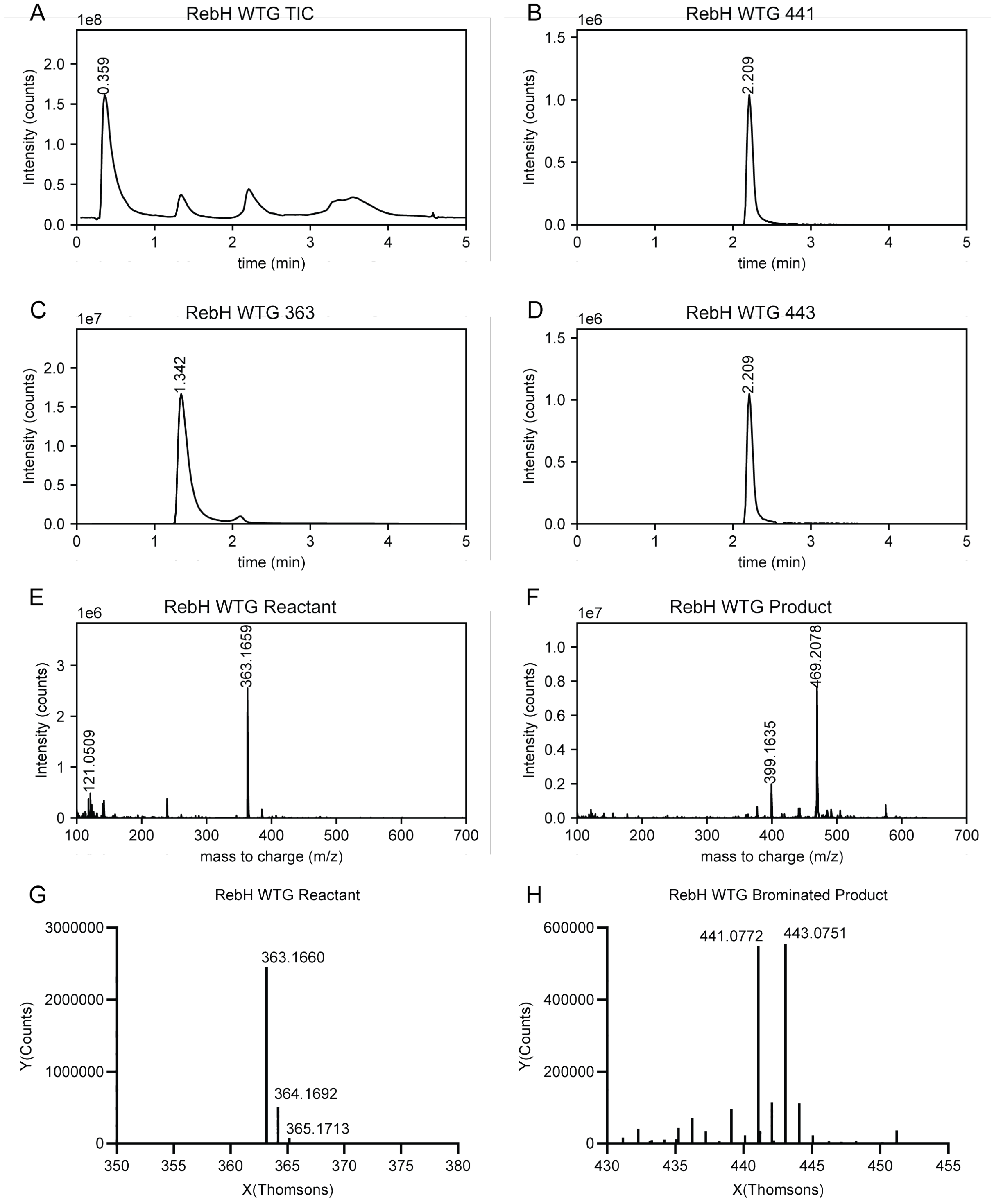
A
RebH WTG TIC
1e8
0.359
Intensity (counts)
time (min)
B
RebH WTG 441
1e6
2.209
Intensity (counts)
time (min)
C
RebH WTG 363
1e7
1.342
Intensity (counts)
time (min)
D
RebH WTG 443
1e6
2.209
Intensity (counts)
time (min)
E
RebH WTG Reactant
1e6
363.1659
121.0509
Intensity (counts)
mass to charge (m/z)
F
RebH WTG Product
1e7
469.2078
399.1635
Intensity (counts)
mass to charge (m/z)
G
RebH WTG Reactant
363.1660
364.1692
365.1713
Y(Counts)
X(Thomsons)
H
RebH WTG Brominated Product
441.0772
443.0751
Y(Counts)
X(Thomsons)

#### RebH WVG

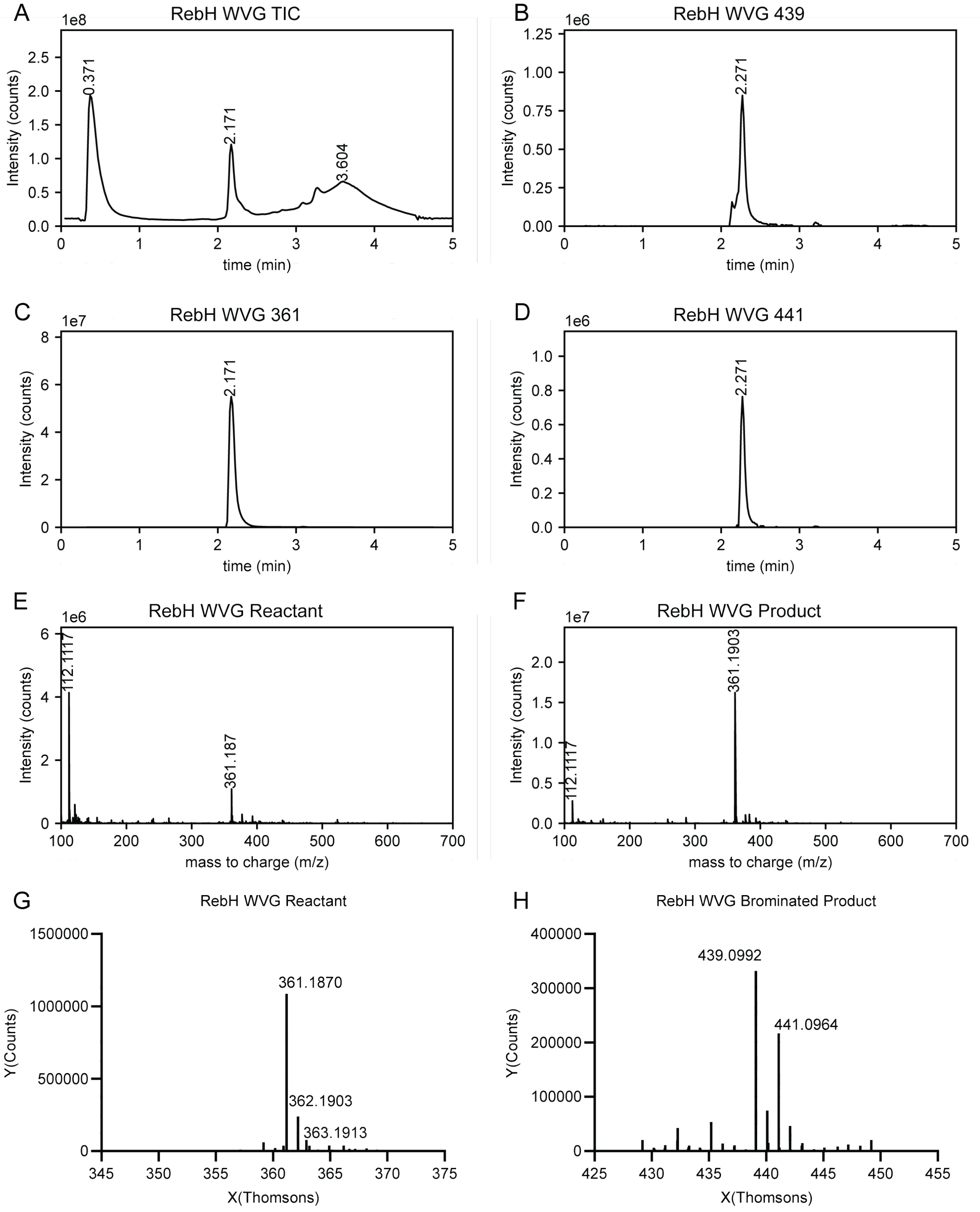
A
RebH WVG TIC
1e8
Intensity (counts)
time (min)
0.371
2.171
3.604
B
RebH WVG 439
1e6
Intensity (counts)
time (min)
2.271
C
RebH WVG 361
1e7
Intensity (counts)
time (min)
2.171
D
RebH WVG 441
1e6
Intensity (counts)
time (min)
2.271
E
RebH WVG Reactant
1e6
Intensity (counts)
mass to charge (m/z)
112.1117
361.187
F
RebH WVG Product
1e7
Intensity (counts)
mass to charge (m/z)
361.1903
112.1117
G
RebH WVG Reactant
Y(Counts)
X(Thomsons)
361.1870
362.1903
363.1913
H
RebH WVG Brominated Product
Y(Counts)
X(Thomsons)
439.0992
441.0964

#### RebH WYG

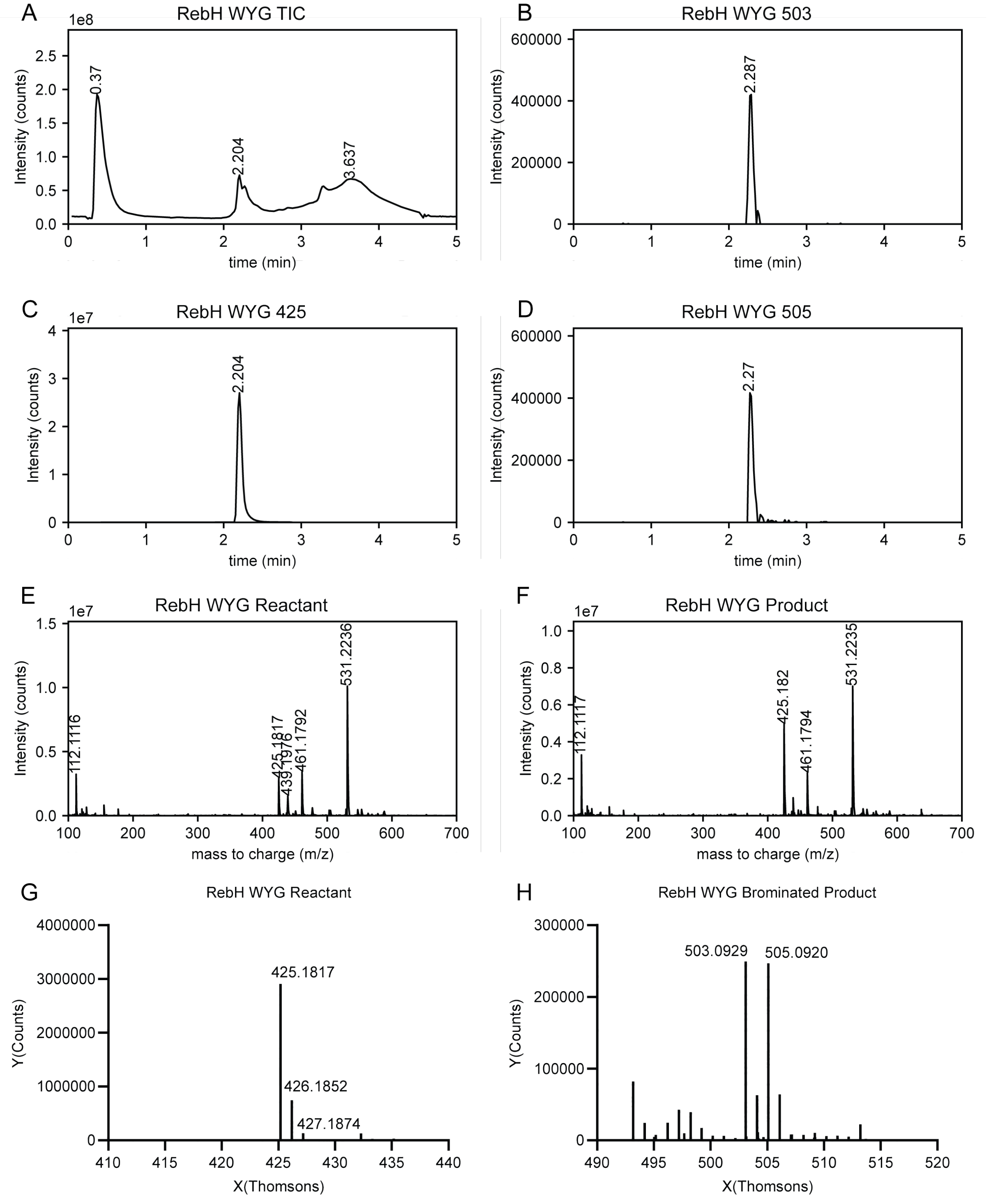

A
RebH WYG TIC
1e8
Intensity (counts)
time (min)
0.37
2.204
3.637
B
RebH WYG 503
Intensity (counts)
time (min)
2.287
C
RebH WYG 425
1e7
Intensity (counts)
time (min)
2.204
D
RebH WYG 505
Intensity (counts)
time (min)
2.27
E
RebH WYG Reactant
1e7
Intensity (counts)
mass to charge (m/z)
112.1116
425.1817
439.1976
461.1792
531.2236
F
RebH WYG Product
1e7
Intensity (counts)
mass to charge (m/z)
112.1117
425.182
461.1794
531.2235
G
RebH WYG Reactant
Y(Counts)
X(Thomsons)
425.1817
426.1852
427.1874
H
RebH WYG Brominated Product
Y(Counts)
X(Thomsons)
503.0929
505.0920

#### RebH WGA

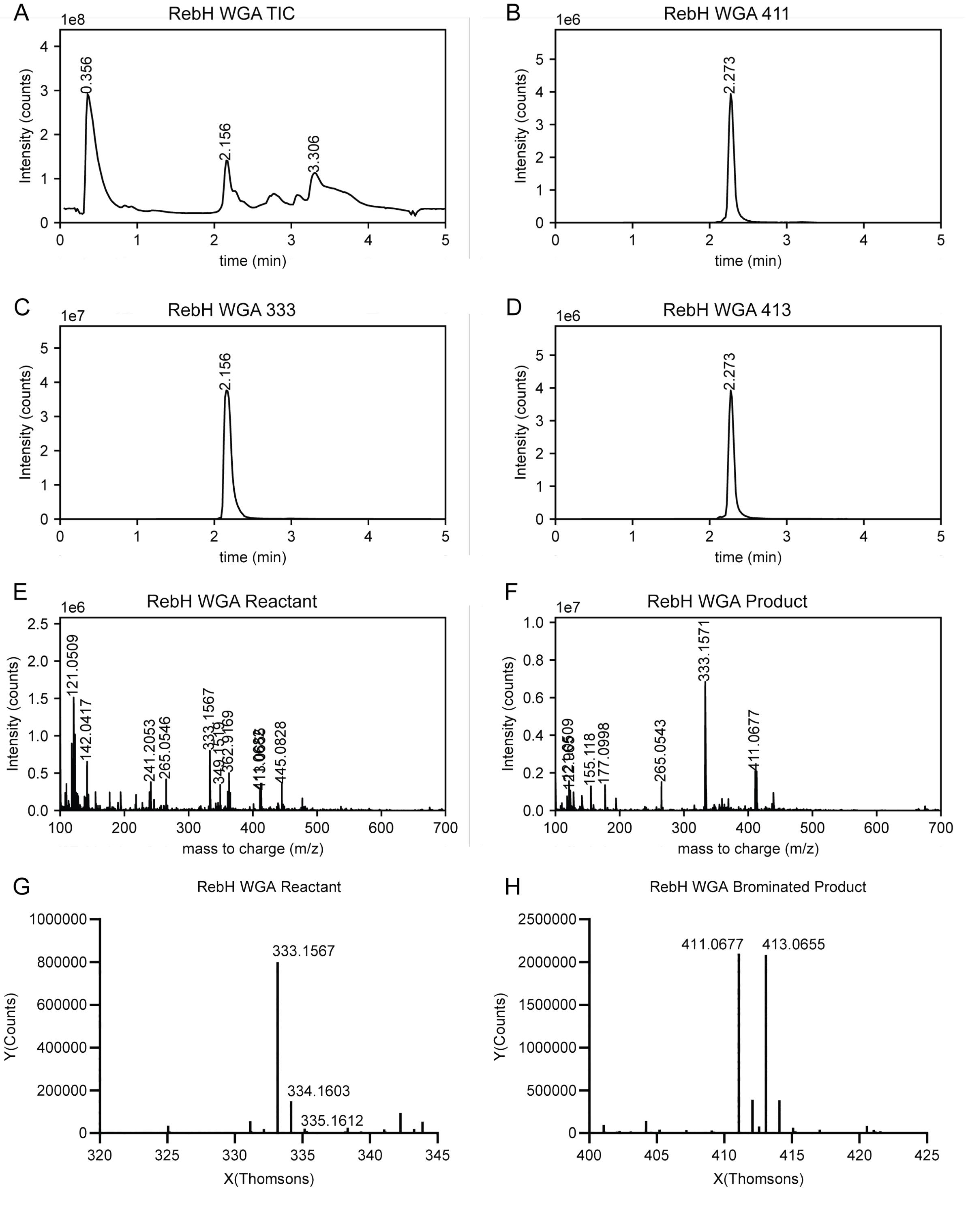

A
RebH WGA TIC
1e8
Intensity (counts)
time (min)
0.356
2.156
3.306
B
RebH WGA 411
1e6
2.273
C
RebH WGA 333
1e7
2.156
D
RebH WGA 413
1e6
2.273
E
RebH WGA Reactant
1e6
mass to charge (m/z)
121.0509
142.0417
241.2053
265.0546
333.1567
349.1519
362.9169
445.0828
F
RebH WGA Product
1e7
333.1571
411.0677
265.0543
155.118
177.0998
G
RebH WGA Reactant
Y(Counts)
X(Thomsons)
333.1567
334.1603
335.1612
H
RebH WGA Brominated Product
411.0677
413.0655

#### RebH WGC

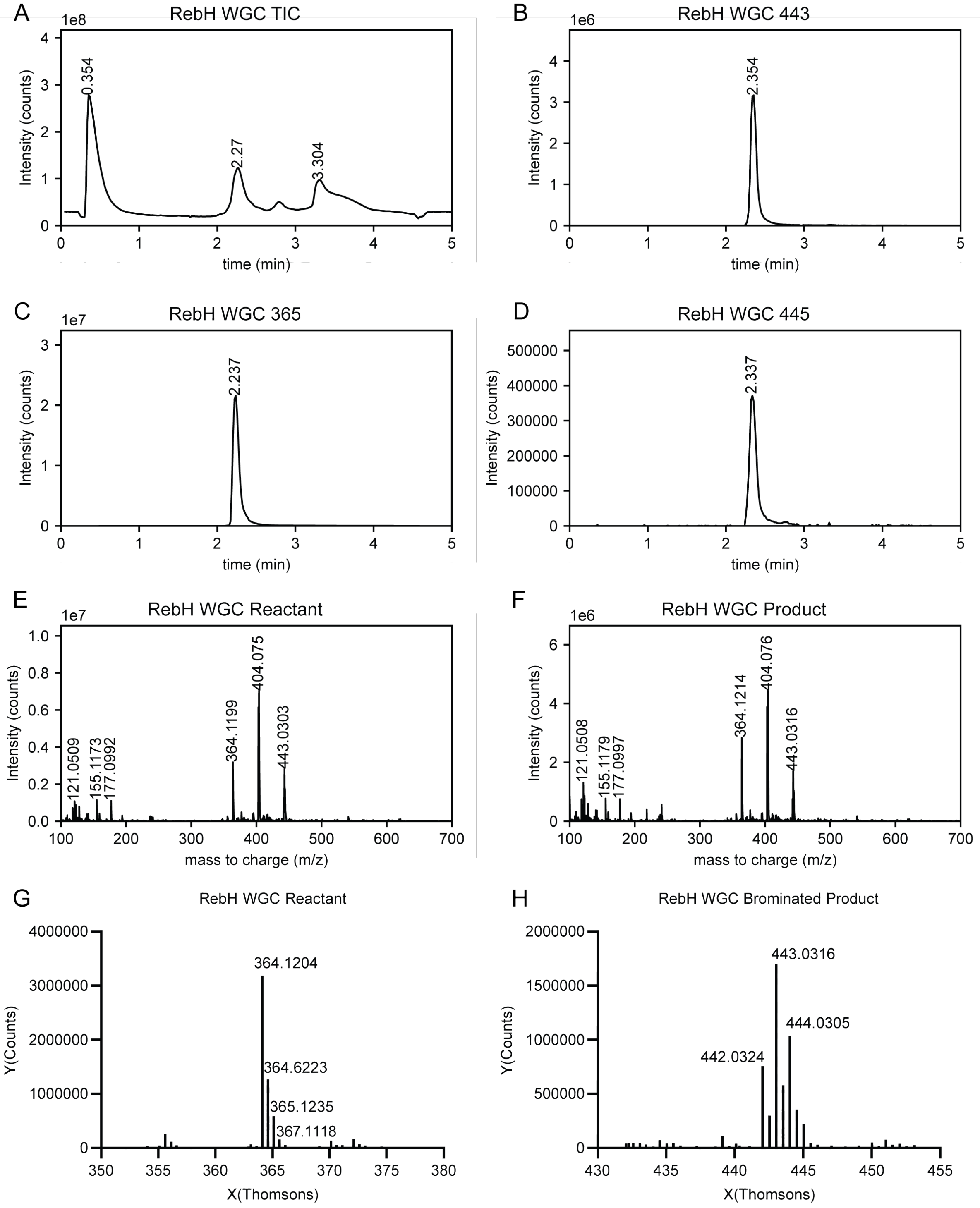
A
RebH WGC TIC
1e8
0.354
2.27
3.304
Intensity (counts)
time (min)
B
RebH WGC 443
1e6
2.354
Intensity (counts)
time (min)
C
RebH WGC 365
1e7
2.237
Intensity (counts)
time (min)
D
RebH WGC 445
2.337
Intensity (counts)
time (min)
E
RebH WGC Reactant
1e7
121.0509
155.1173
177.0992
364.1199
404.075
443.0303
Intensity (counts)
mass to charge (m/z)
F
RebH WGC Product
1e6
121.0508
155.1179
177.0997
364.1214
404.076
443.0316
Intensity (counts)
mass to charge (m/z)
G
RebH WGC Reactant
364.1204
364.6223
365.1235
367.1118
Y(Counts)
X(Thomsons)
H
RebH WGC Brominated Product
443.0316
444.0305
442.0324
Y(Counts)
X(Thomsons)

**RebH WGD**

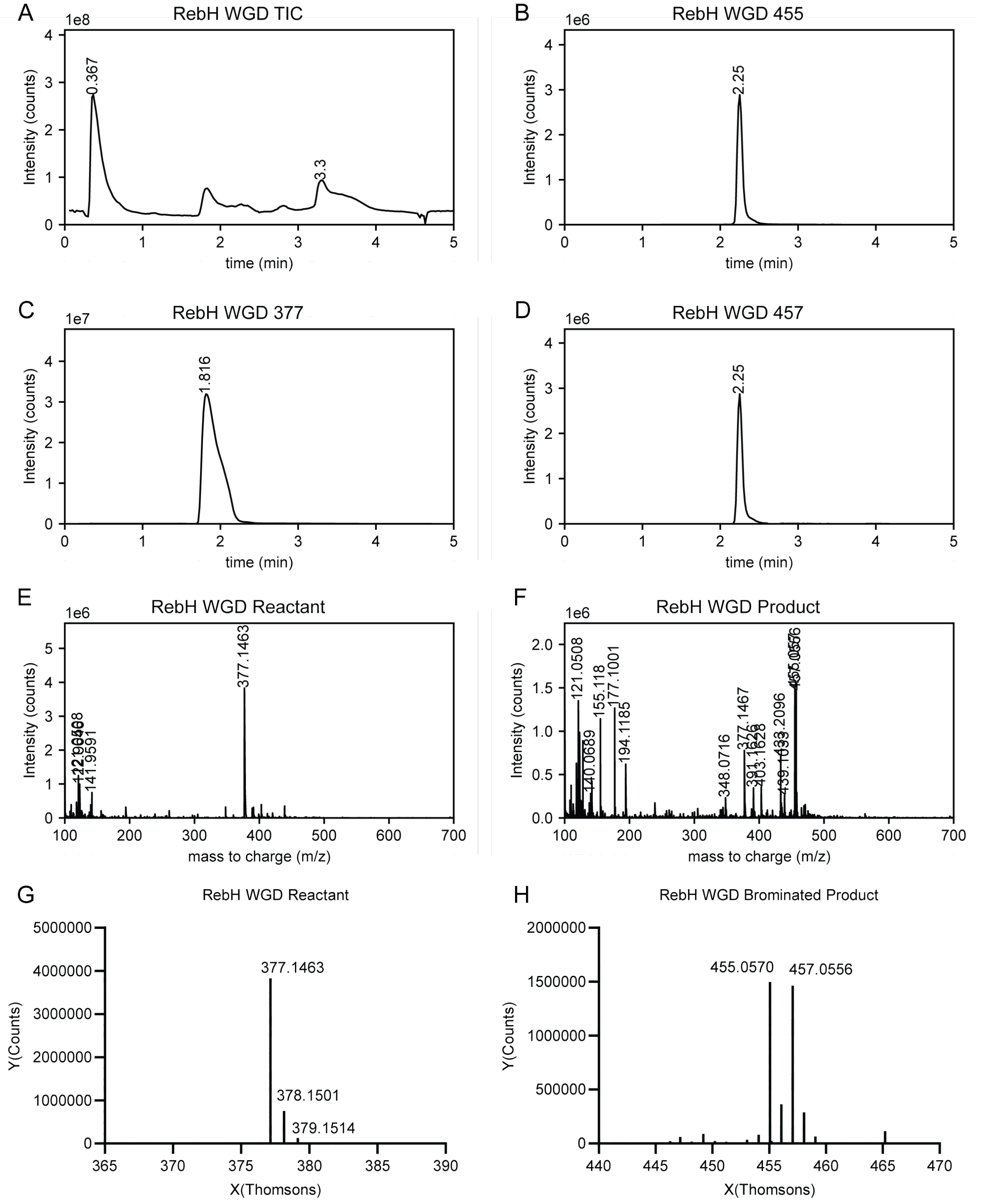
A
RebH WGD TIC
1e8
Intensity (counts)
time (min)
0.367
3.3
B
RebH WGD 455
1e6
Intensity (counts)
time (min)
2.25
C
RebH WGD 377
1e7
Intensity (counts)
time (min)
1.816
D
RebH WGD 457
1e6
Intensity (counts)
time (min)
2.25
E
RebH WGD Reactant
1e6
Intensity (counts)
mass to charge (m/z)
377.1463
141.9591
F
RebH WGD Product
1e6
Intensity (counts)
mass to charge (m/z)
121.0508
155.118
177.1001
194.1185
140.0689
348.0716
377.1467
433.2096
G
RebH WGD Reactant
Y(Counts)
X(Thomsons)
377.1463
378.1501
379.1514
H
RebH WGD Brominated Product
Y(Counts)
X(Thomsons)
455.0570
457.0556

#### RebH WGE

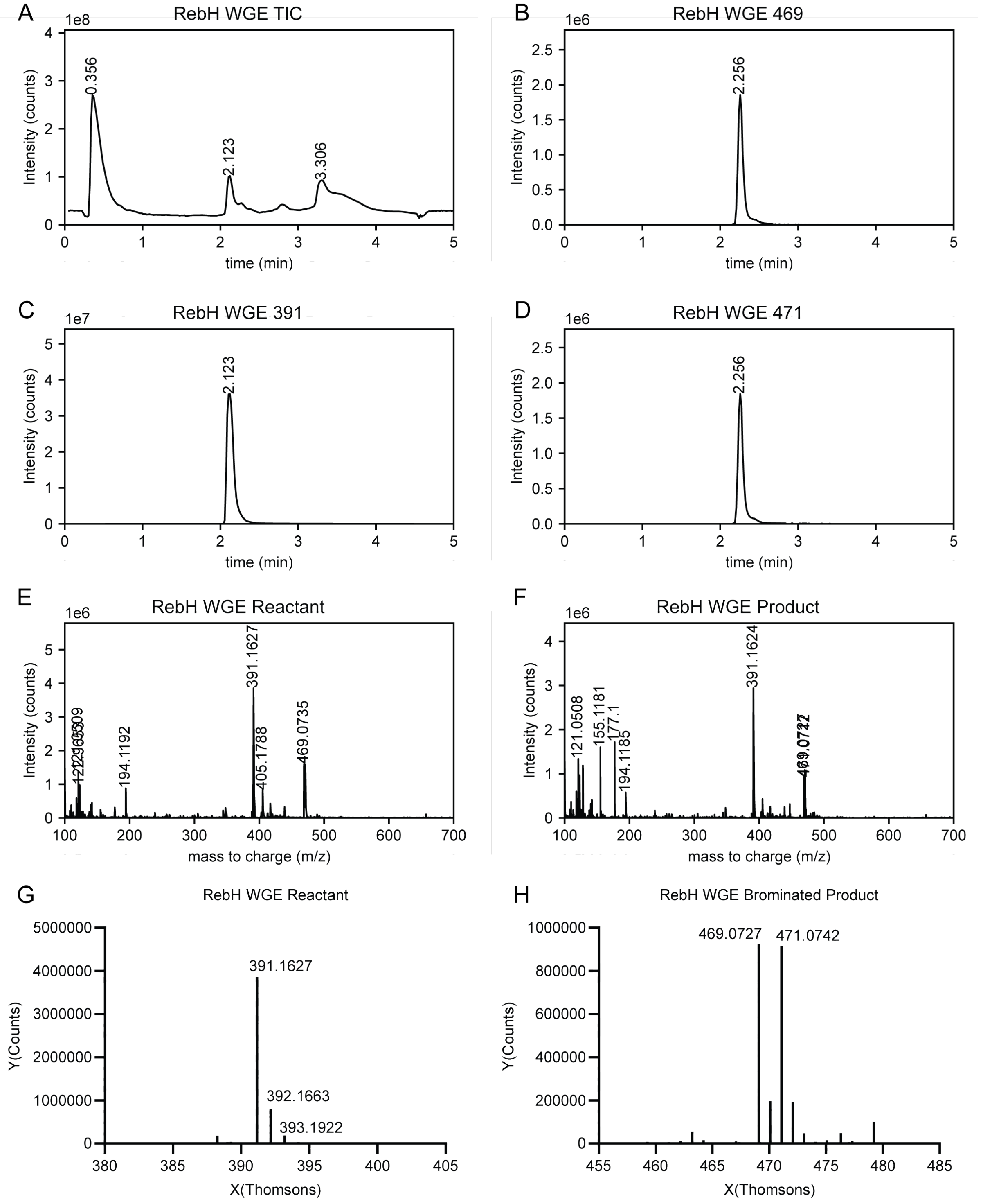
A
RebH WGE TIC
1e8
0.356
2.123
3.306
Intensity (counts)
time (min)
B
RebH WGE 469
1e6
2.256
Intensity (counts)
time (min)
C
RebH WGE 391
1e7
2.123
Intensity (counts)
time (min)
D
RebH WGE 471
1e6
2.256
Intensity (counts)
time (min)
E
RebH WGE Reactant
1e6
391.1627
469.0735
405.1788
194.1192
Intensity (counts)
mass to charge (m/z)
F
RebH WGE Product
1e6
391.1624
121.0508
155.1181
177.1
194.1185
Intensity (counts)
mass to charge (m/z)
G
RebH WGE Reactant
391.1627
392.1663
393.1922
Y(Counts)
X(Thomsons)
H
RebH WGE Brominated Product
469.0727
471.0742
Y(Counts)
X(Thomsons)

#### RebH WGF

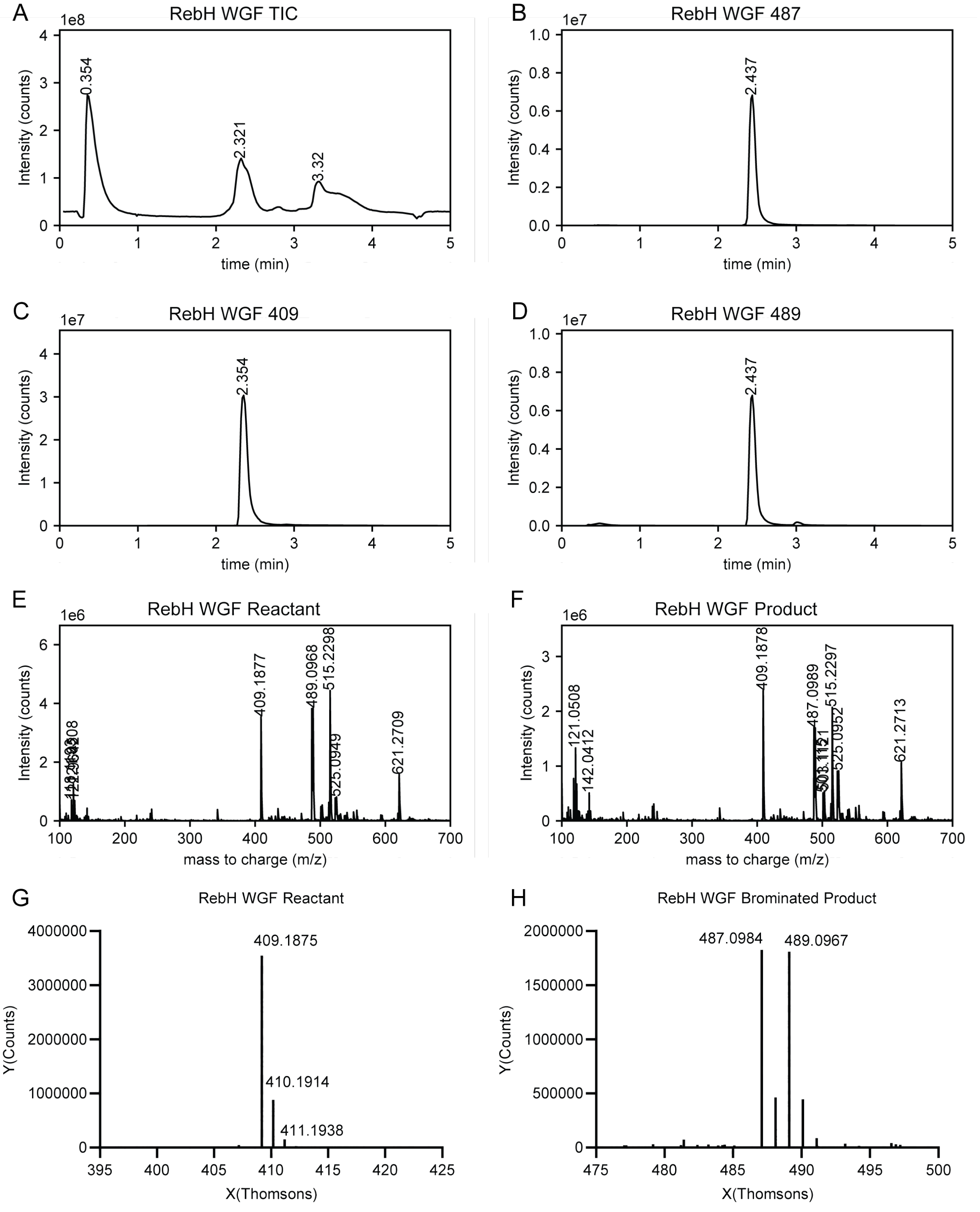
A
RebH WGF TIC
1e8
Intensity (counts)
time (min)
0.354
2.321
3.32
B
RebH WGF 487
1e7
Intensity (counts)
time (min)
2.437
C
RebH WGF 409
1e7
Intensity (counts)
time (min)
2.354
D
RebH WGF 489
1e7
Intensity (counts)
time (min)
2.437
E
RebH WGF Reactant
1e6
Intensity (counts)
mass to charge (m/z)
409.1877
489.0968
515.2298
525.0949
621.2709
F
RebH WGF Product
1e6
Intensity (counts)
mass to charge (m/z)
121.0508
142.0412
409.1878
487.0989
515.2297
525.0952
621.2713
G
RebH WGF Reactant
Y(Counts)
X(Thomsons)
409.1875
410.1914
411.1938
H
RebH WGF Brominated Product
Y(Counts)
X(Thomsons)
487.0984
489.0967

#### RebH WGH

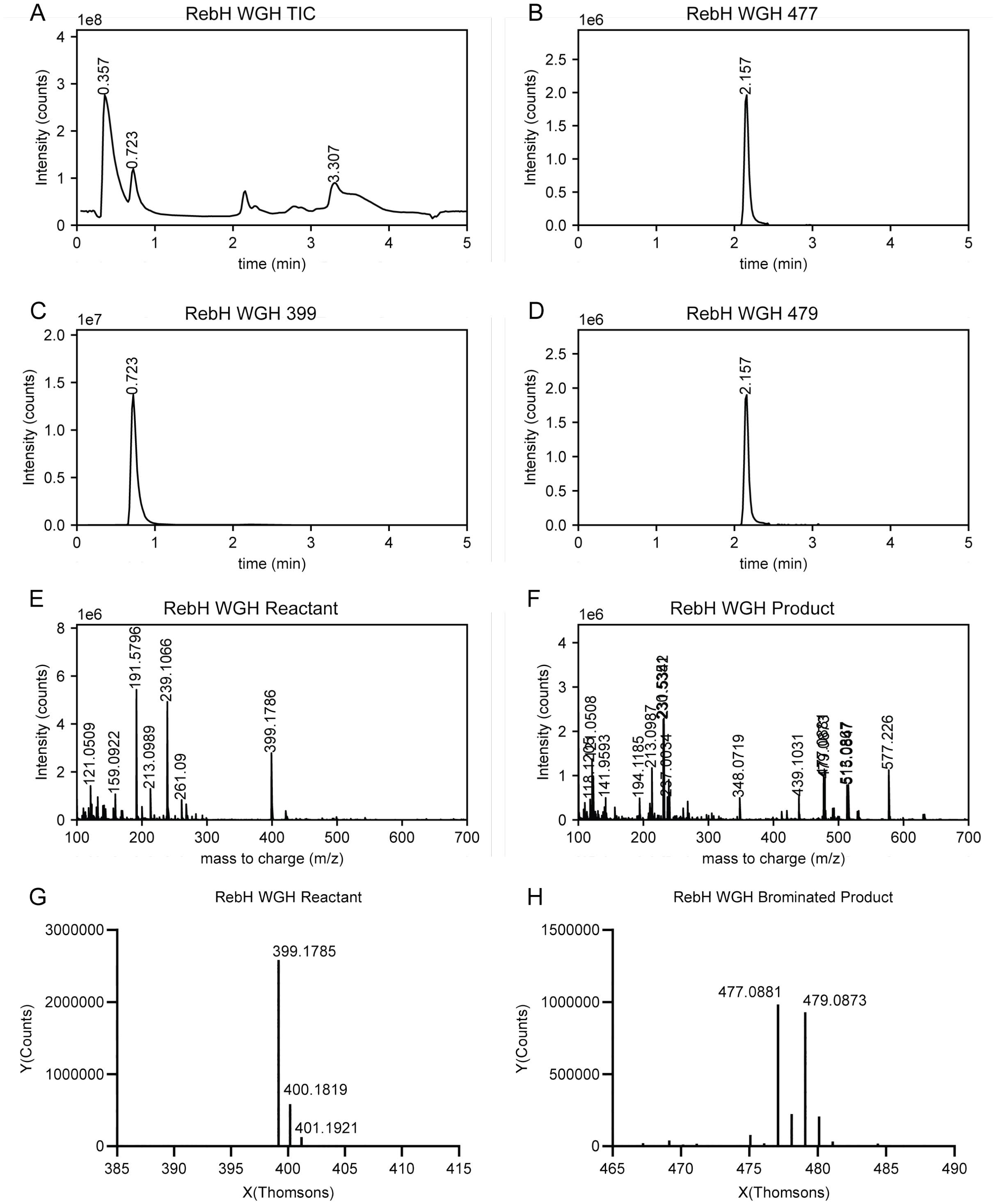
A
RebH WGH TIC
1e8
Intensity (counts)
time (min)
0.357
0.723
3.307
B
RebH WGH 477
1e6
2.157
C
RebH WGH 399
1e7
0.723
D
RebH WGH 479
1e6
2.157
E
RebH WGH Reactant
mass to charge (m/z)
121.0509
159.0922
191.5796
213.0989
239.1066
261.09
399.1786
F
RebH WGH Product
118.1205
121.0508
141.9593
194.1185
213.0987
237.0034
348.0719
439.1031
577.226
G
RebH WGH Reactant
Y(Counts)
X(Thomsons)
399.1785
400.1819
401.1921
H
RebH WGH Brominated Product
477.0881
479.0873

#### RebH WGK

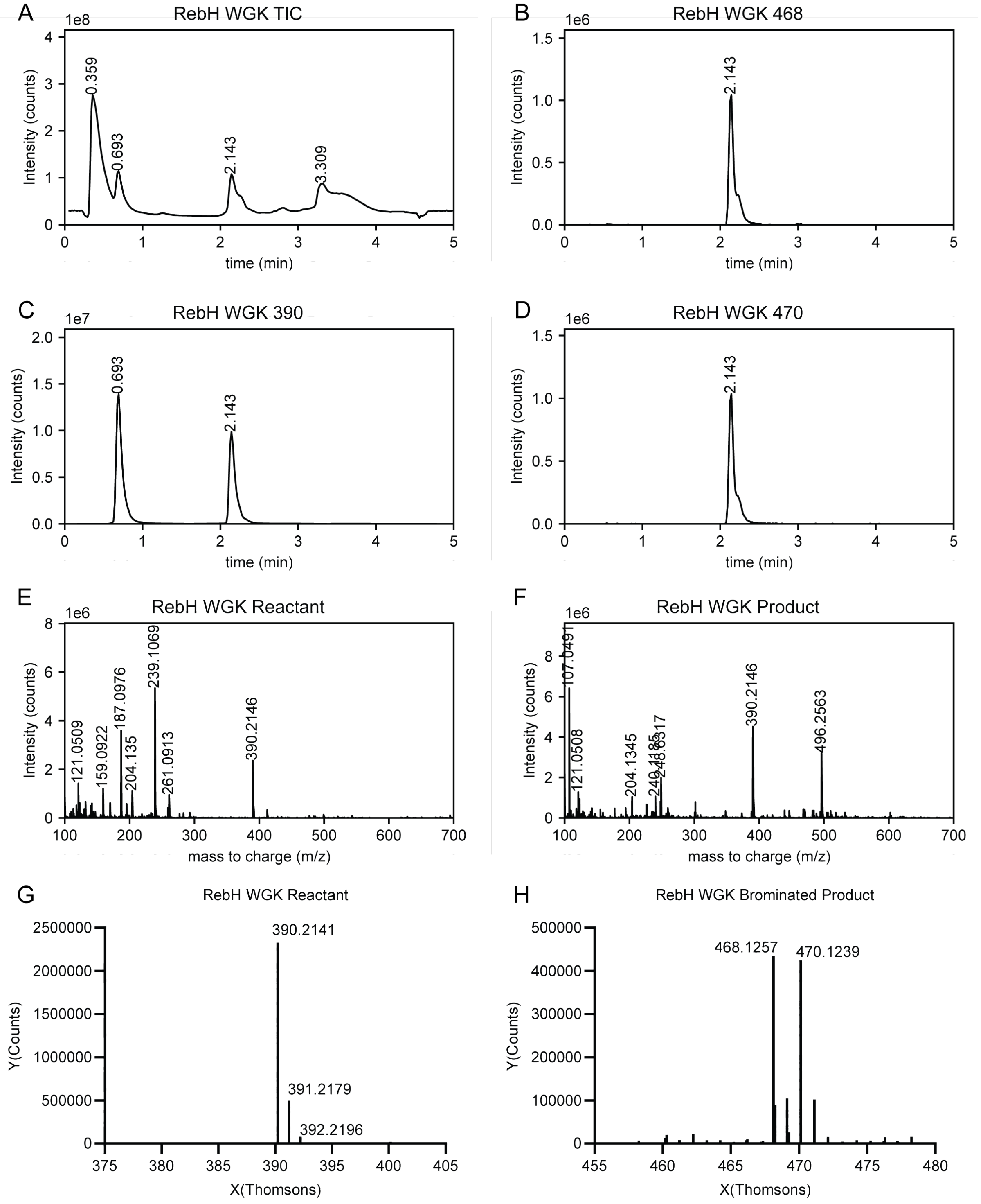

A
RebH WGK TIC
1e8
Intensity (counts)
time (min)
0.359
0.693
2.143
3.309
B
RebH WGK 468
1e6
Intensity (counts)
time (min)
2.143
C
RebH WGK 390
1e7
Intensity (counts)
time (min)
0.693
2.143
D
RebH WGK 470
1e6
Intensity (counts)
time (min)
2.143
E
RebH WGK Reactant
1e6
Intensity (counts)
mass to charge (m/z)
121.0509
159.0922
187.0976
204.135
239.1069
261.0913
390.2146
F
RebH WGK Product
1e6
Intensity (counts)
mass to charge (m/z)
121.0508
204.1345
390.2146
496.2563
G
RebH WGK Reactant
Y(Counts)
X(Thomsons)
390.2141
391.2179
392.2196
H
RebH WGK Brominated Product
Y(Counts)
X(Thomsons)
468.1257
470.1239

#### RebH WGL

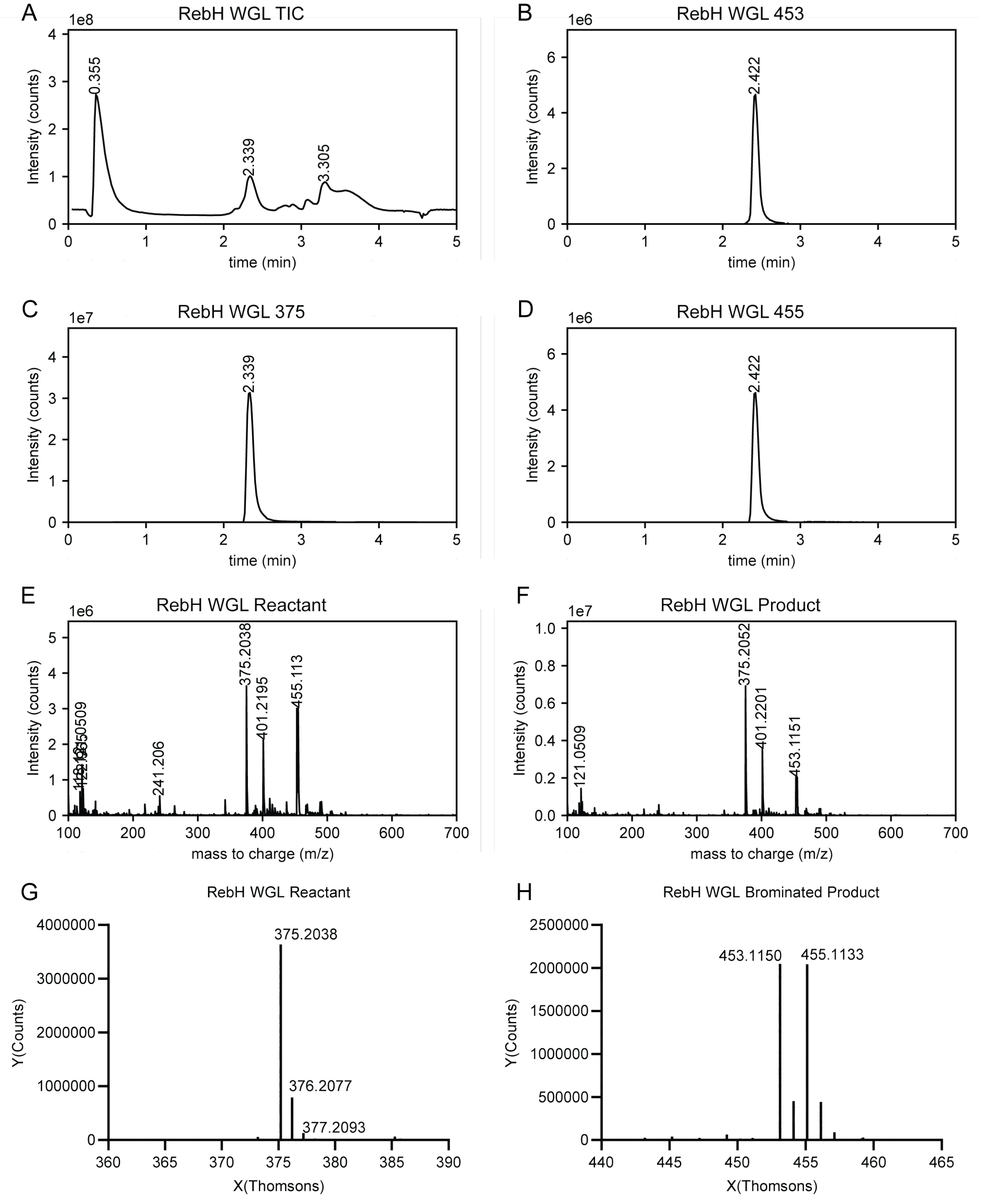

A
RebH WGL TIC
1e8
0.355
2.339
3.305
Intensity (counts)
time (min)
B
RebH WGL 453
1e6
2.422
Intensity (counts)
time (min)
C
RebH WGL 375
1e7
2.339
Intensity (counts)
time (min)
D
RebH WGL 455
1e6
2.422
Intensity (counts)
time (min)
E
RebH WGL Reactant
1e6
241.206
375.2038
401.2195
455.113
Intensity (counts)
mass to charge (m/z)
F
RebH WGL Product
1e7
121.0509
375.2052
401.2201
453.1151
Intensity (counts)
mass to charge (m/z)
G
RebH WGL Reactant
375.2038
376.2077
377.2093
Y(Counts)
X(Thomsons)
H
RebH WGL Brominated Product
453.1150
455.1133
Y(Counts)
X(Thomsons)

#### RebH WGM

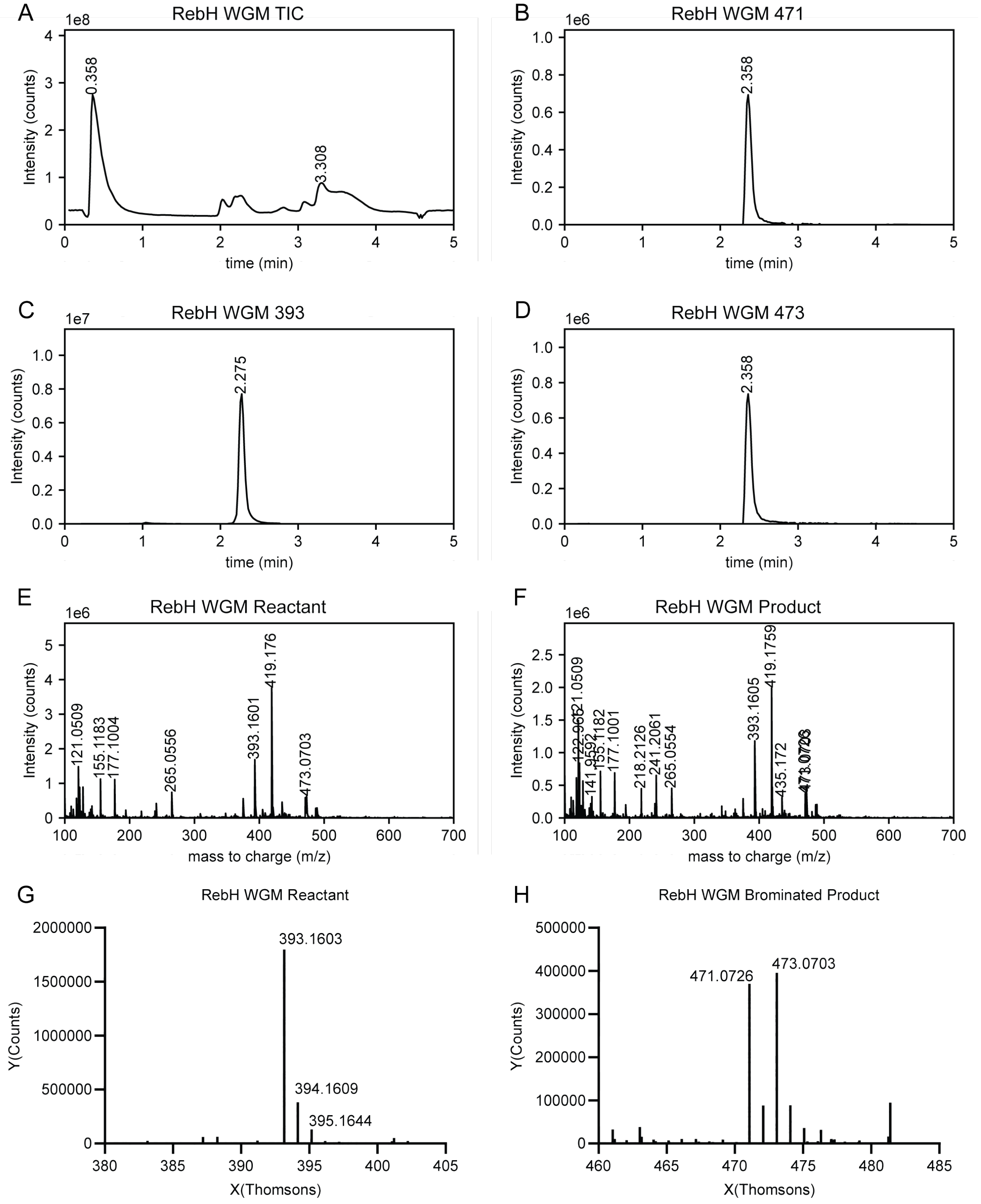
A
RebH WGM TIC
1e8
Intensity (counts)
time (min)
0.358
3.308
B
RebH WGM 471
1e6
Intensity (counts)
time (min)
2.358
C
RebH WGM 393
1e7
Intensity (counts)
time (min)
2.275
D
RebH WGM 473
1e6
Intensity (counts)
time (min)
2.358
E
RebH WGM Reactant
1e6
Intensity (counts)
mass to charge (m/z)
121.0509
155.1183
177.1004
265.0556
393.1601
419.176
473.0703
F
RebH WGM Product
1e6
Intensity (counts)
mass to charge (m/z)
141.9592
155.1182
177.1001
218.2126
241.2061
265.0554
393.1605
419.1759
435.172
G
RebH WGM Reactant
Y(Counts)
X(Thomsons)
393.1603
394.1609
395.1644
H
RebH WGM Brominated Product
Y(Counts)
X(Thomsons)
471.0726
473.0703

#### RebH WGN

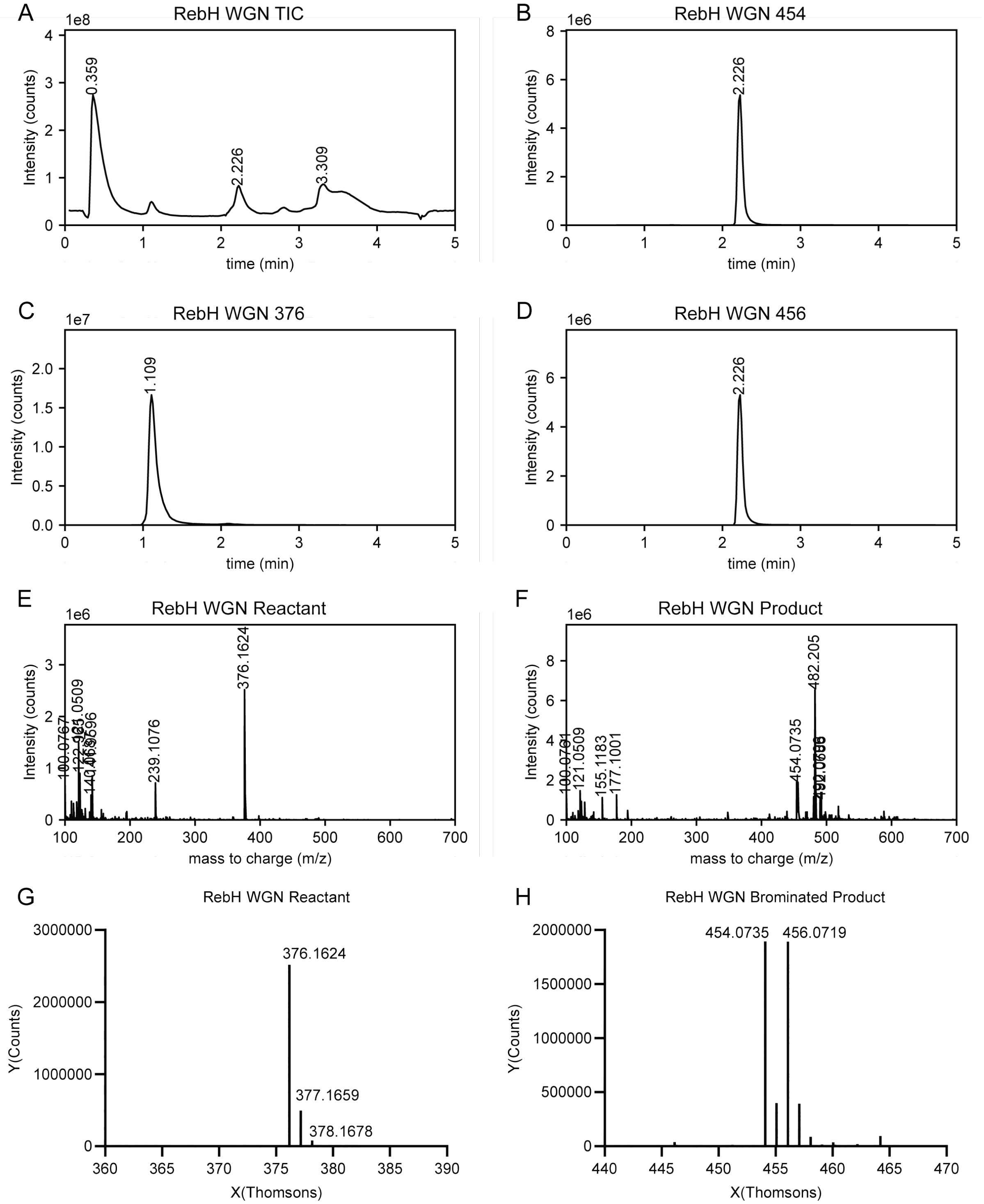
A
RebH WGN TIC
1e8
0.359
2.226
3.309
Intensity (counts)
time (min)
B
RebH WGN 454
1e6
2.226
Intensity (counts)
time (min)
C
RebH WGN 376
1e7
1.109
Intensity (counts)
time (min)
D
RebH WGN 456
1e6
2.226
Intensity (counts)
time (min)
E
RebH WGN Reactant
1e6
376.1624
239.1076
Intensity (counts)
mass to charge (m/z)
F
RebH WGN Product
1e6
482.205
454.0735
121.0509
155.1183
177.1001
Intensity (counts)
mass to charge (m/z)
G
RebH WGN Reactant
376.1624
377.1659
378.1678
Y(Counts)
X(Thomsons)
H
RebH WGN Brominated Product
454.0735
456.0719
Y(Counts)
X(Thomsons)

#### RebH WGP

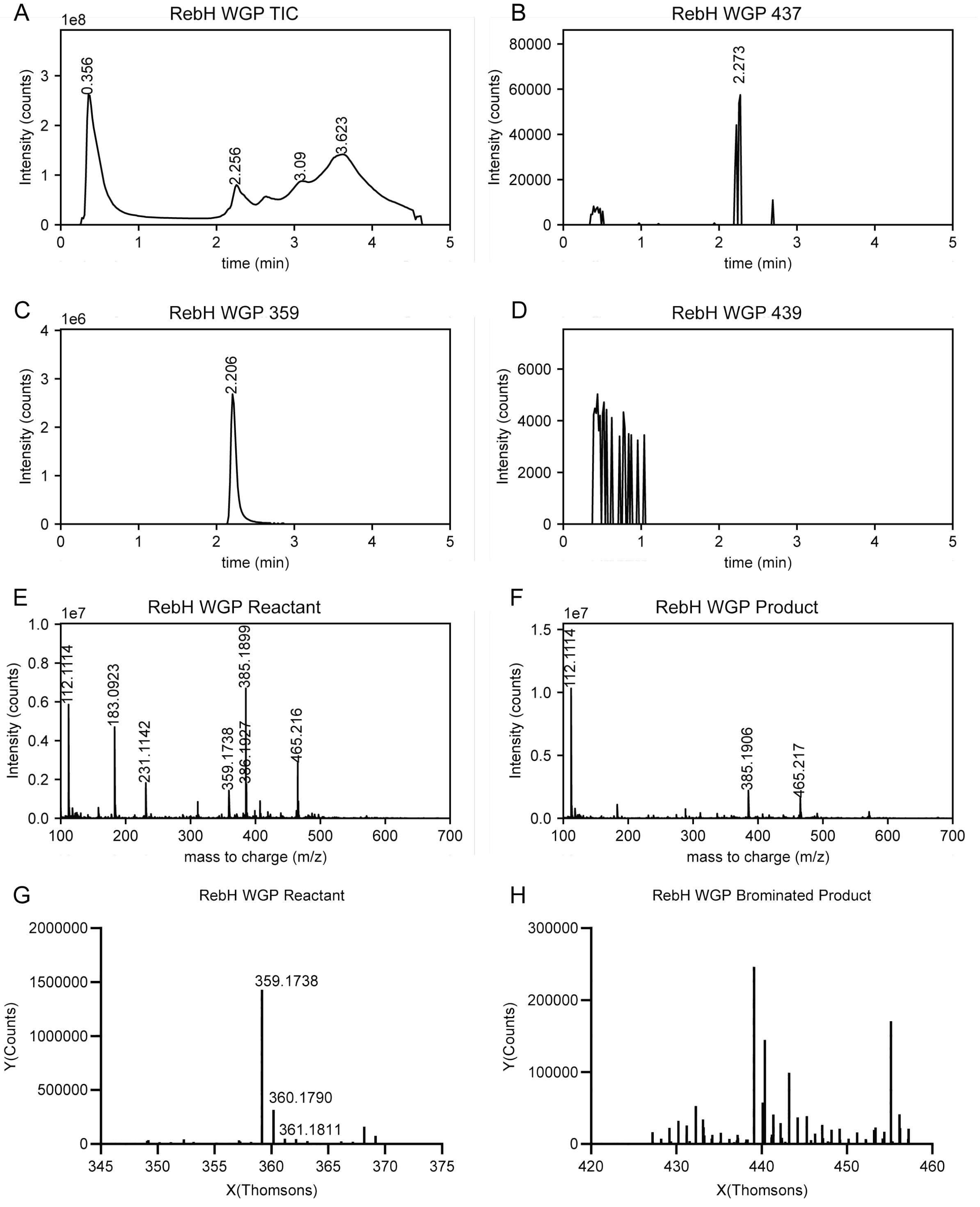
A
RebH WGP TIC
1e8
Intensity (counts)
time (min)
0.356
2.256
3.09
3.623
B
RebH WGP 437
2.273
Intensity (counts)
time (min)
C
RebH WGP 359
1e6
2.206
Intensity (counts)
time (min)
D
RebH WGP 439
Intensity (counts)
time (min)
E
RebH WGP Reactant
1e7
112.1114
183.0923
231.1142
359.1738
386.1927
385.1899
465.216
Intensity (counts)
mass to charge (m/z)
F
RebH WGP Product
1e7
112.1114
385.1906
465.217
Intensity (counts)
mass to charge (m/z)
G
RebH WGP Reactant
359.1738
360.1790
361.1811
Y(Counts)
X(Thomsons)
H
RebH WGP Brominated Product
Y(Counts)
X(Thomsons)

#### RebH WGQ

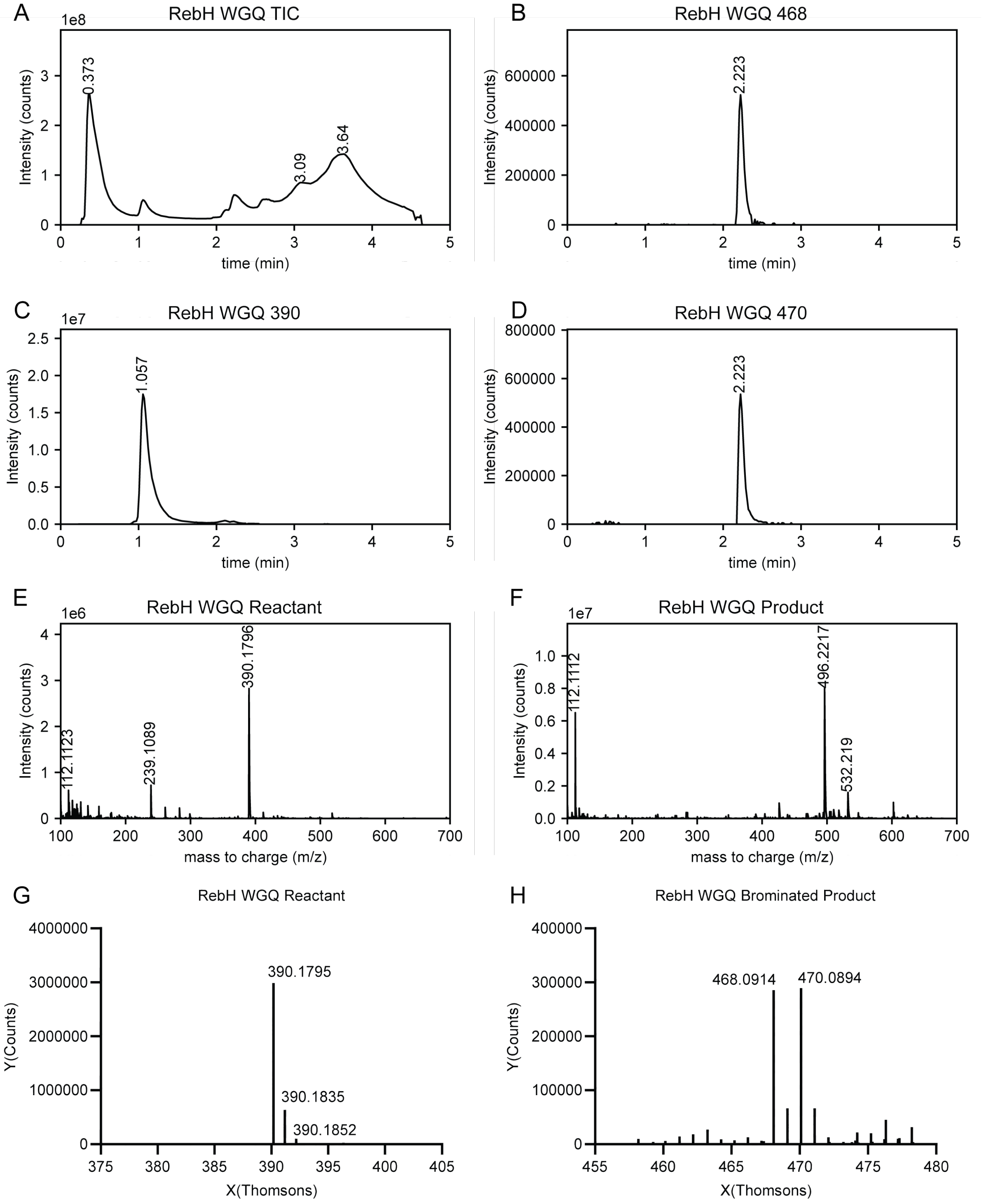
A
RebH WGQ TIC
1e8
Intensity (counts)
0.373
3.09
3.64
time (min)
B
RebH WGQ 468
Intensity (counts)
2.223
time (min)
C
RebH WGQ 390
1e7
Intensity (counts)
1.057
time (min)
D
RebH WGQ 470
Intensity (counts)
2.223
time (min)
E
RebH WGQ Reactant
1e6
Intensity (counts)
112.1123
239.1089
390.1796
mass to charge (m/z)
F
RebH WGQ Product
1e7
Intensity (counts)
112.1112
496.2217
532.219
mass to charge (m/z)
G
RebH WGQ Reactant
Y(Counts)
390.1795
390.1835
390.1852
X(Thomsons)
H
RebH WGQ Brominated Product
Y(Counts)
468.0914
470.0894
X(Thomsons)

#### RebH WGR

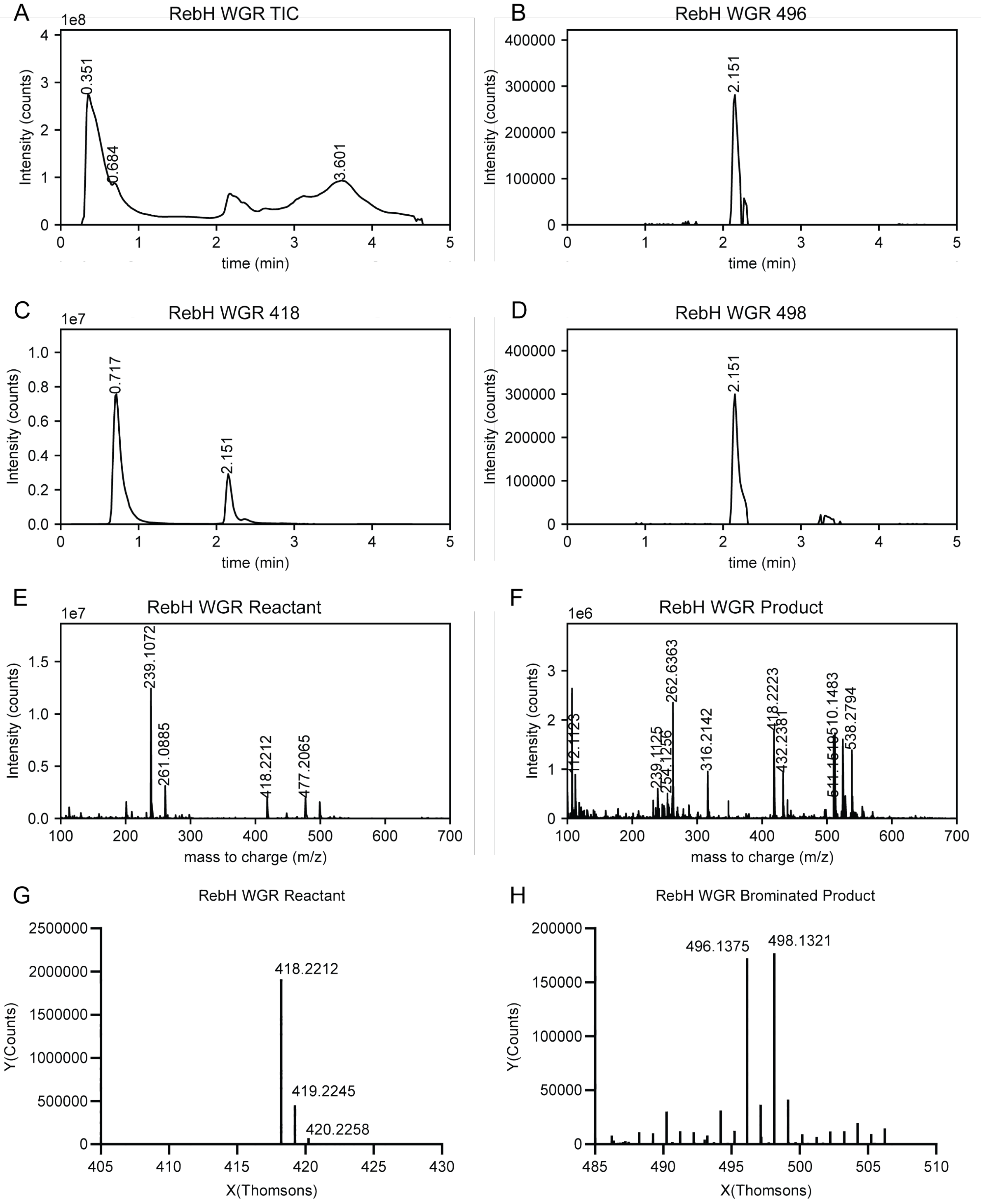

A
RebH WGR TIC
1e8
Intensity (counts)
time (min)
0.351
0.684
3.601
B
RebH WGR 496
Intensity (counts)
time (min)
2.151
C
RebH WGR 418
1e7
Intensity (counts)
time (min)
0.717
2.151
D
RebH WGR 498
Intensity (counts)
time (min)
2.151
E
RebH WGR Reactant
1e7
Intensity (counts)
mass to charge (m/z)
239.1072
261.0885
418.2212
477.2065
F
RebH WGR Product
1e6
Intensity (counts)
mass to charge (m/z)
112.1123
239.1125
254.1256
262.6363
316.2142
418.2223
432.2381
510.1483
538.2794
G
RebH WGR Reactant
Y(Counts)
X(Thomsons)
418.2212
419.2245
420.2258
H
RebH WGR Brominated Product
Y(Counts)
X(Thomsons)
496.1375
498.1321

#### RebH WGT

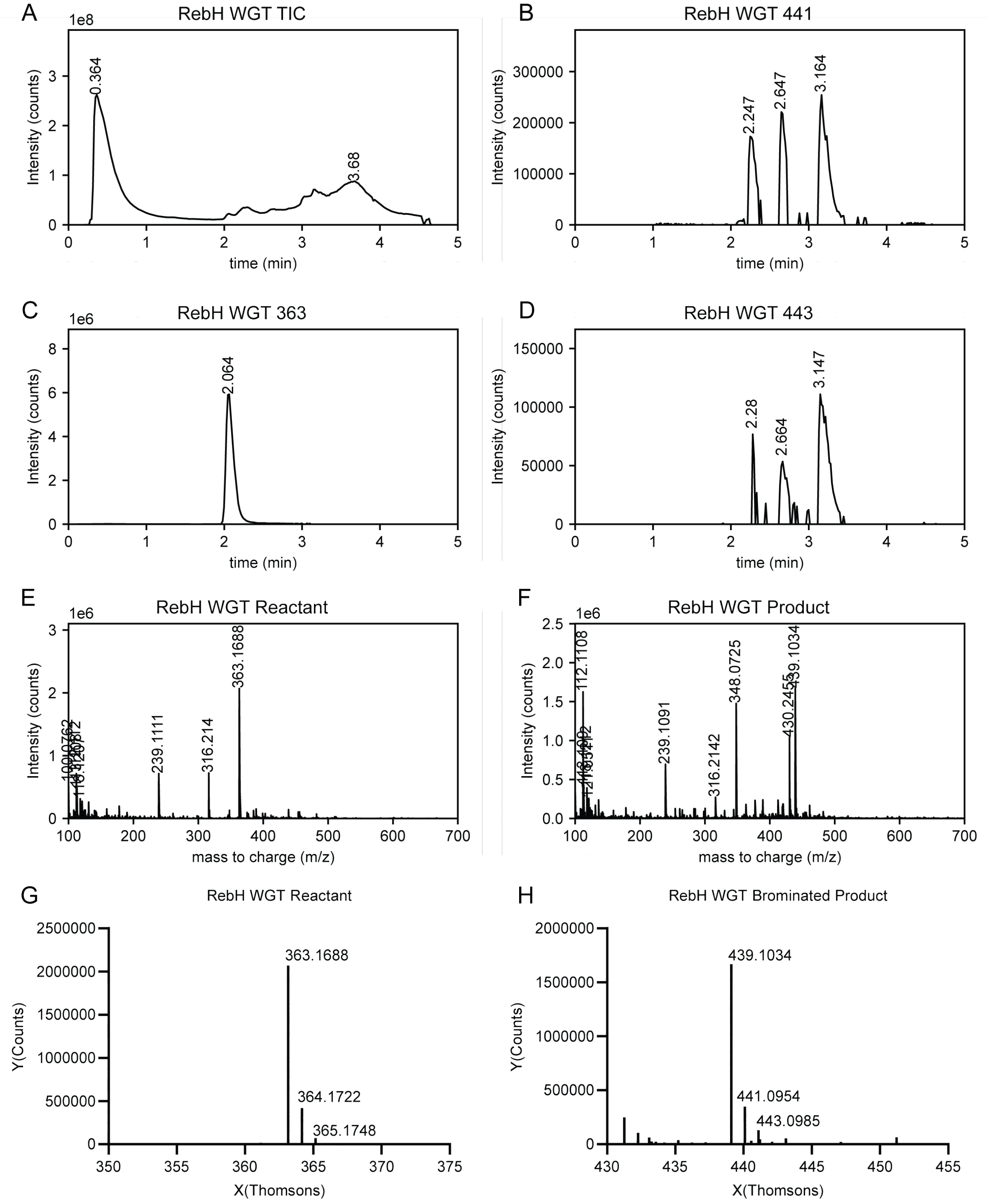
A
RebH WGT TIC
1e8
Intensity (counts)
time (min)
0.364
3.68
B
RebH WGT 441
Intensity (counts)
time (min)
2.247
2.647
3.164
C
RebH WGT 363
1e6
Intensity (counts)
time (min)
2.064
D
RebH WGT 443
Intensity (counts)
time (min)
2.28
2.664
3.147
E
RebH WGT Reactant
1e6
Intensity (counts)
mass to charge (m/z)
239.1111
316.214
363.1688
F
RebH WGT Product
1e6
Intensity (counts)
mass to charge (m/z)
112.1108
239.1091
316.2142
348.0725
439.1034
G
RebH WGT Reactant
Y(Counts)
X(Thomsons)
363.1688
364.1722
365.1748
H
RebH WGT Brominated Product
Y(Counts)
X(Thomsons)
439.1034
441.0954
443.0985

#### RebH WGV

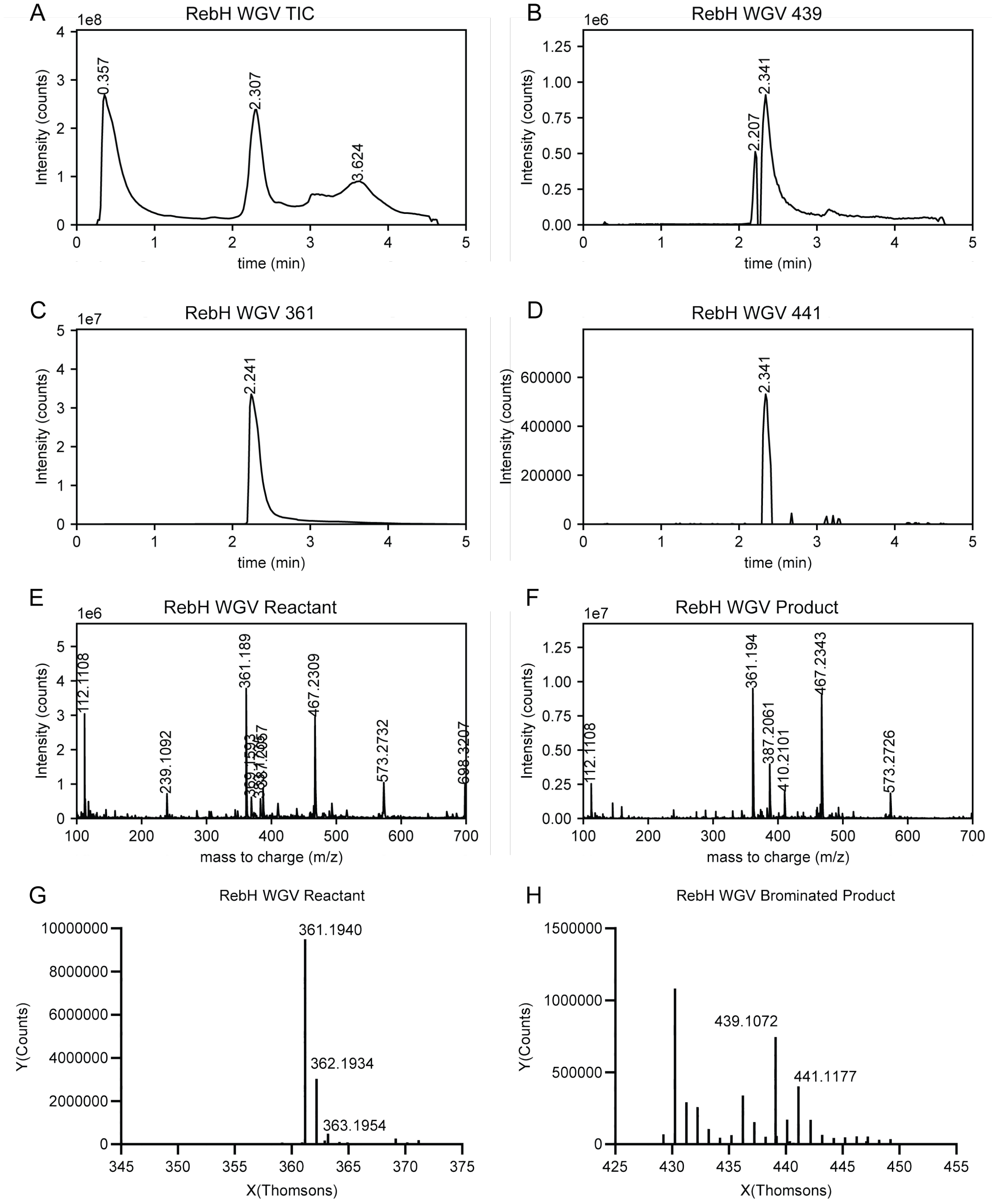

A
RebH WGV TIC
1e8
Intensity (counts)
time (min)
0.357
2.307
3.624
B
RebH WGV 439
1e6
Intensity (counts)
time (min)
2.207
2.341
C
RebH WGV 361
1e7
Intensity (counts)
time (min)
2.241
D
RebH WGV 441
Intensity (counts)
time (min)
2.341
E
RebH WGV Reactant
1e6
Intensity (counts)
mass to charge (m/z)
112.1108
239.1092
361.189
467.2309
573.2732
698.3207
F
RebH WGV Product
1e7
Intensity (counts)
mass to charge (m/z)
112.1108
361.194
387.2061
410.2101
467.2343
573.2726
G
RebH WGV Reactant
Y(Counts)
X(Thomsons)
361.1940
362.1934
363.1954
H
RebH WGV Brominated Product
Y(Counts)
X(Thomsons)
439.1072
441.1177

#### RebH WGY

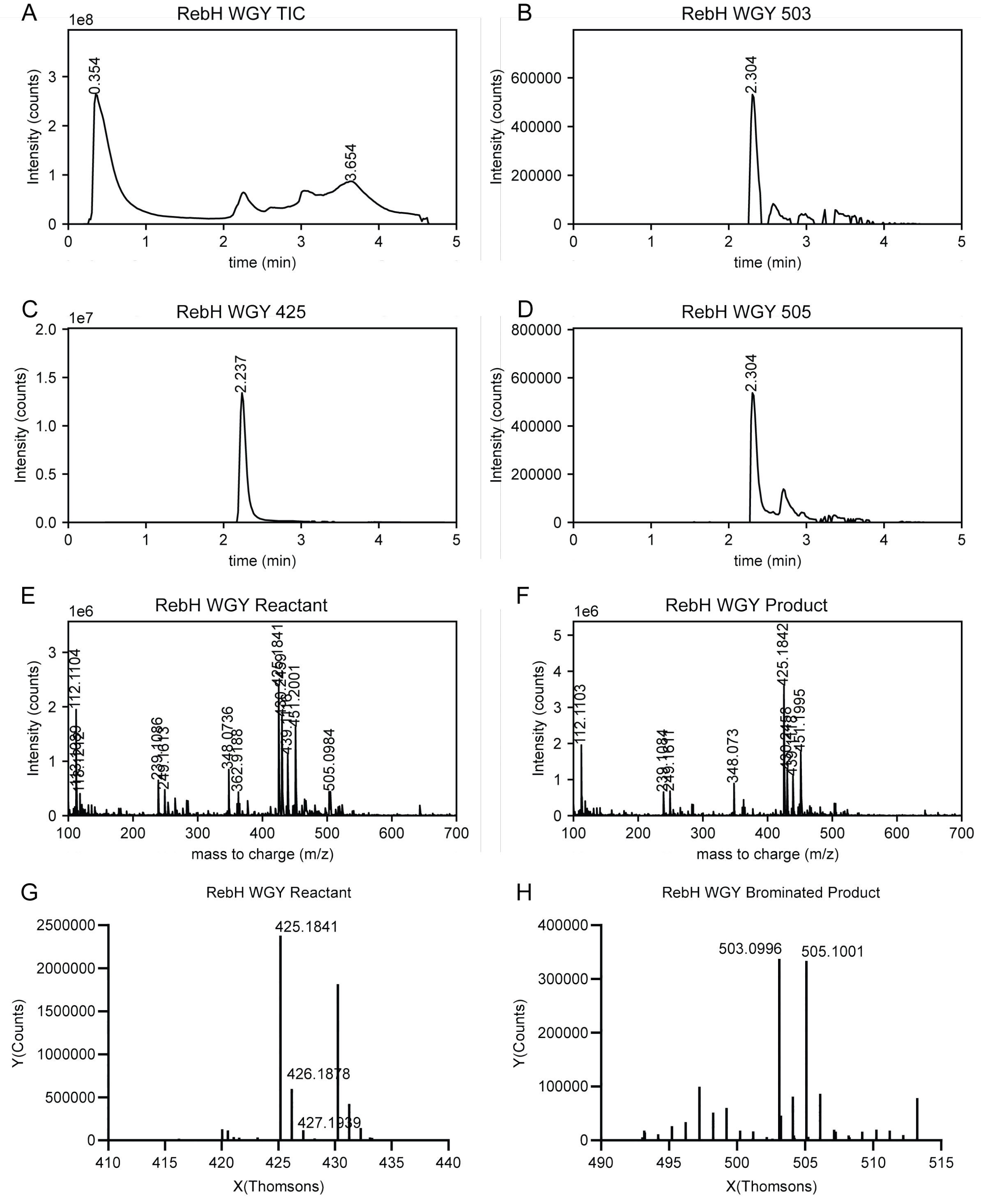

A
RebH WGY TIC
1e8
Intensity (counts)
0.354
3.654
time (min)
B
RebH WGY 503
2.304
Intensity (counts)
time (min)
C
RebH WGY 425
1e7
2.237
Intensity (counts)
time (min)
D
RebH WGY 505
2.304
Intensity (counts)
time (min)
E
RebH WGY Reactant
1e6
112.1104
239.1086
249.1613
348.0736
362.9188
451.2001
505.0984
Intensity (counts)
mass to charge (m/z)
F
RebH WGY Product
1e6
112.1103
348.073
425.1842
451.1995
Intensity (counts)
mass to charge (m/z)
G
RebH WGY Reactant
425.1841
426.1878
427.1939
Y(Counts)
X(Thomsons)
H
RebH WGY Brominated Product
503.0996
505.1001
Y(Counts)
X(Thomsons)

#### RebH AWG

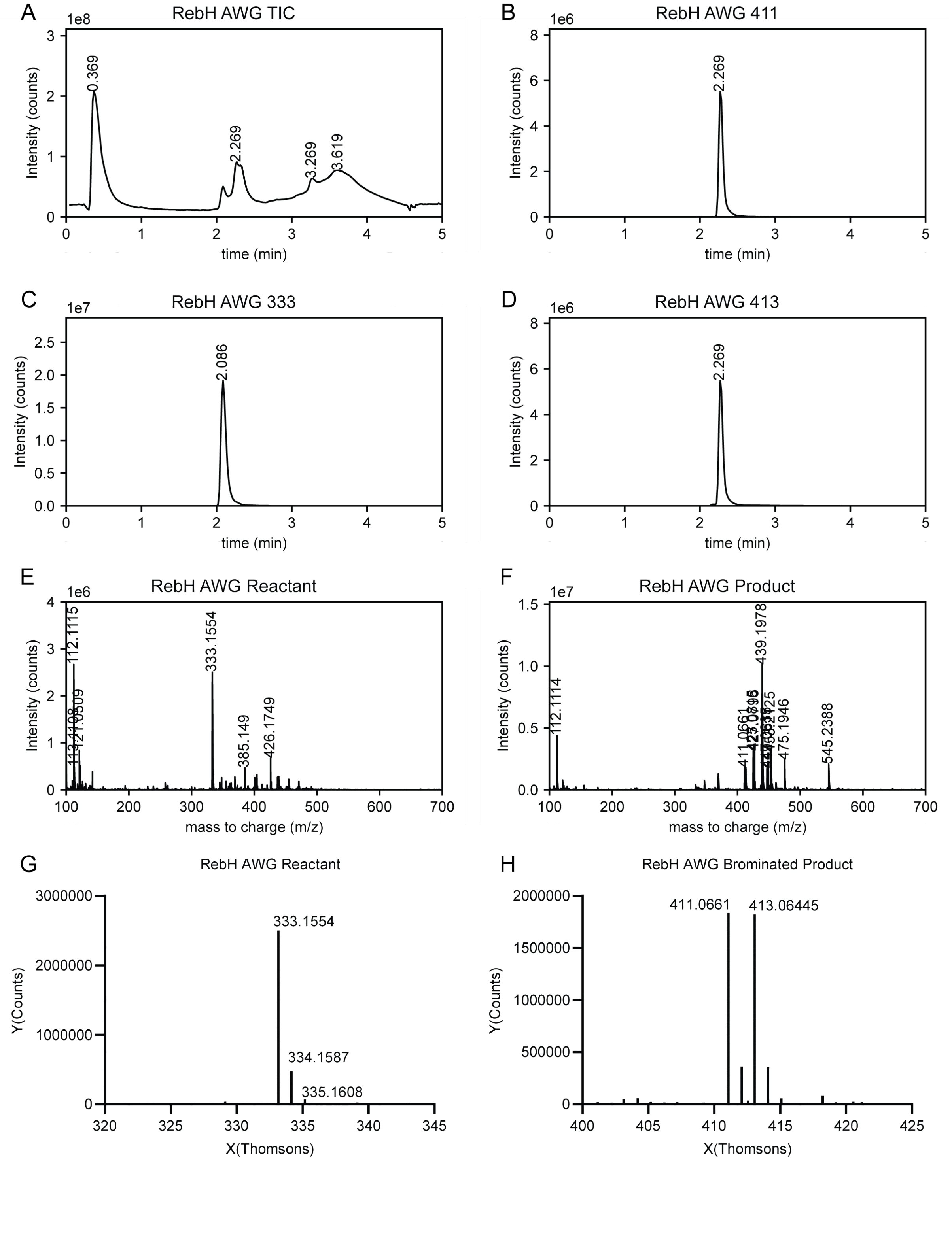
A
RebH AWG TIC
1e8
Intensity (counts)
time (min)
0.369
2.269
3.269
3.619
B
RebH AWG 411
1e6
2.269
C
RebH AWG 333
1e7
2.086
D
RebH AWG 413
1e6
2.269
E
RebH AWG Reactant
1e6
mass to charge (m/z)
112.1115
121.0509
333.1554
385.149
426.1749
F
RebH AWG Product
1e7
112.1114
411.0661
439.1978
475.1946
545.2388
G
RebH AWG Reactant
Y(Counts)
X(Thomsons)
333.1554
334.1587
335.1608
H
RebH AWG Brominated Product
411.0661
413.06445

#### RebH CWG

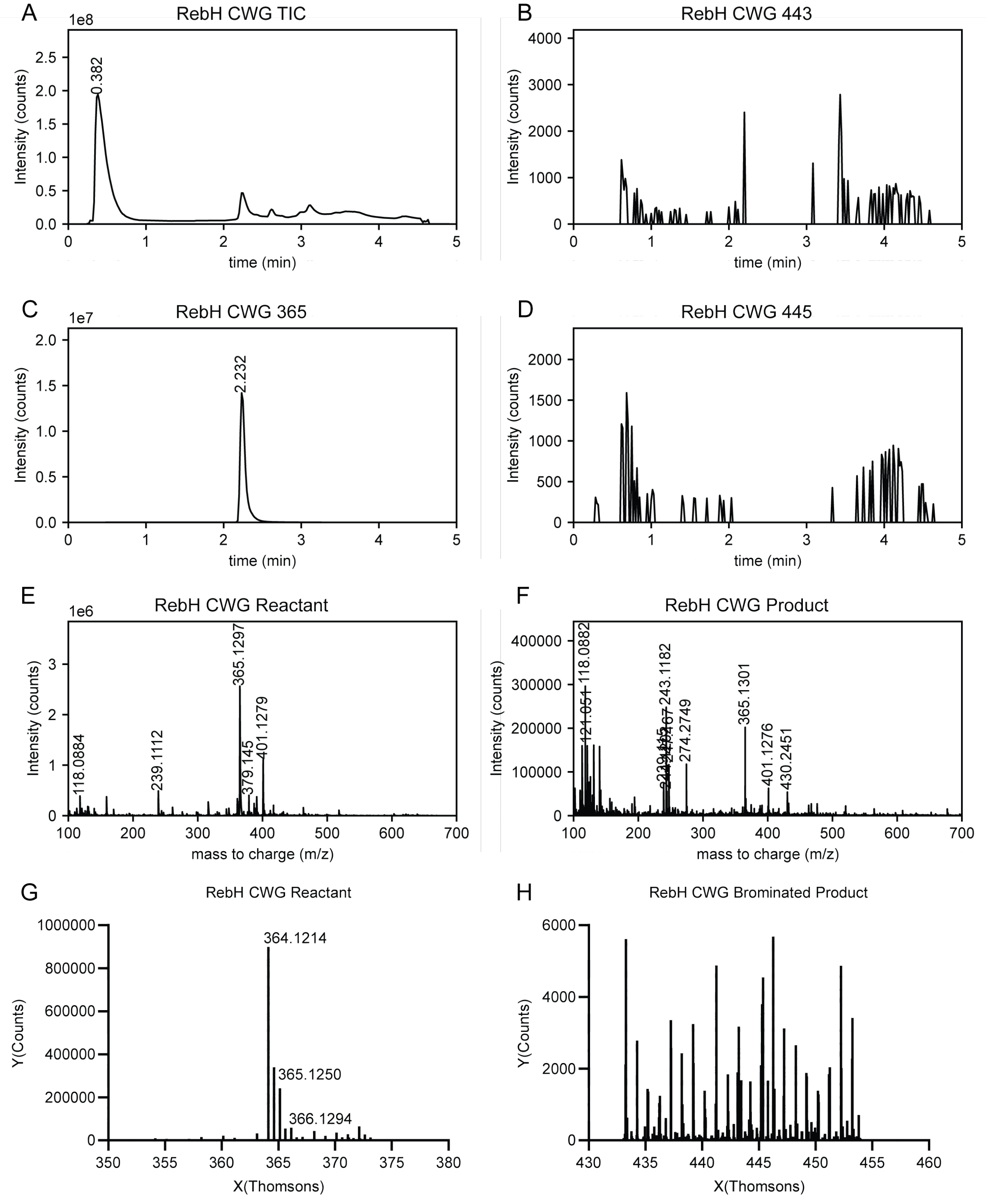

A
RebH CWG TIC
1e8
Intensity (counts)
time (min)
0.382
B
RebH CWG 443
Intensity (counts)
time (min)
C
RebH CWG 365
1e7
Intensity (counts)
time (min)
2.232
D
RebH CWG 445
Intensity (counts)
time (min)
E
RebH CWG Reactant
1e6
Intensity (counts)
mass to charge (m/z)
118.0884
239.1112
365.1297
379.145
401.1279
F
RebH CWG Product
Intensity (counts)
mass to charge (m/z)
118.0882
121.054
243.1182
274.2749
365.1301
401.1276
430.2451
G
RebH CWG Reactant
Y(Counts)
X(Thomsons)
364.1214
365.1250
366.1294
H
RebH CWG Brominated Product
Y(Counts)
X(Thomsons)

#### RebH DWG

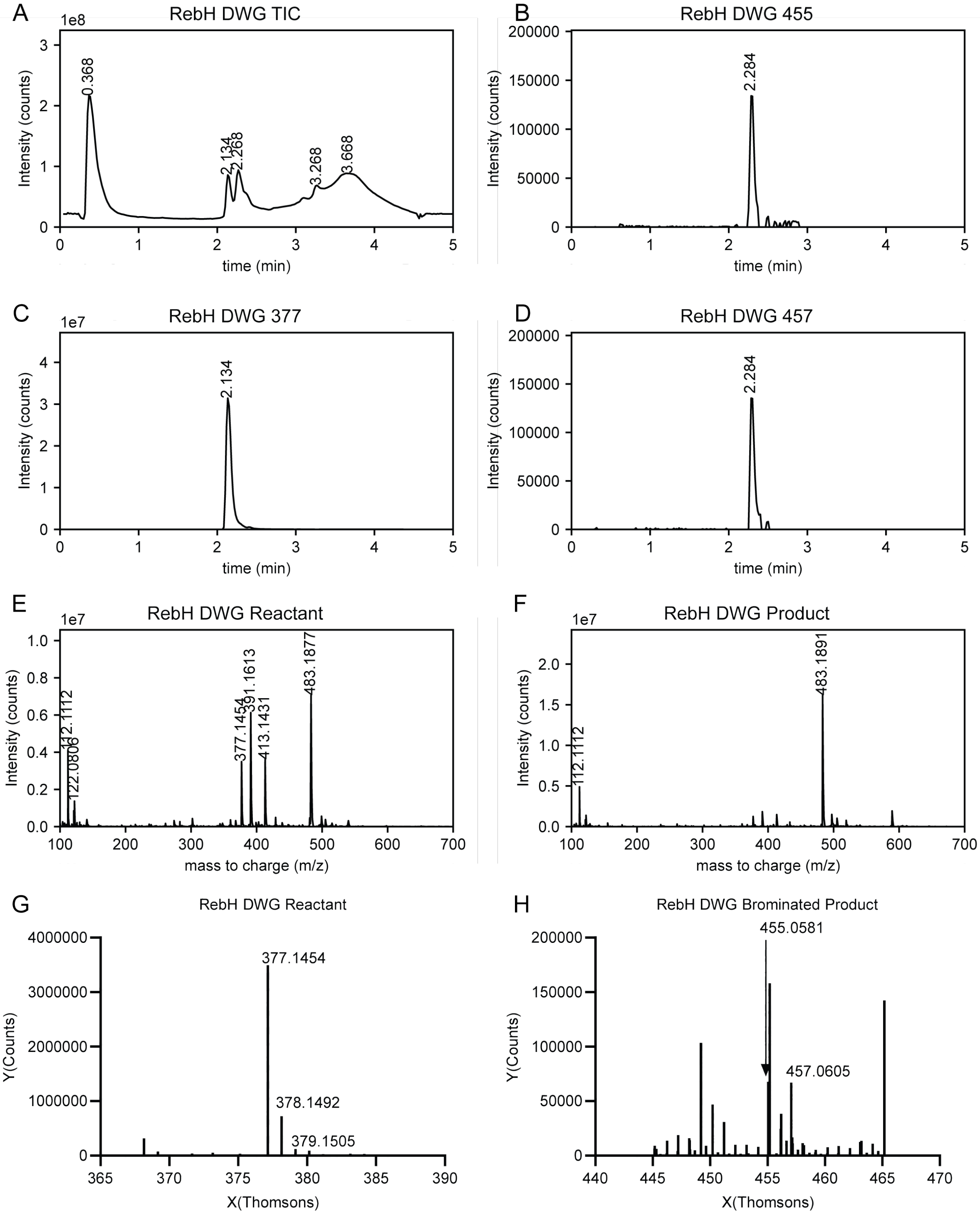
A
RebH DWG TIC
1e8
Intensity (counts)
time (min)
0.368
2.134
2.268
3.268
3.668
B
RebH DWG 455
2.284
C
RebH DWG 377
1e7
2.134
D
RebH DWG 457
2.284
E
RebH DWG Reactant
mass to charge (m/z)
112.1112
122.0806
377.1454
391.1613
413.1431
483.1877
F
RebH DWG Product
483.1891
112.1112
G
RebH DWG Reactant
Y(Counts)
X(Thomsons)
377.1454
378.1492
379.1505
H
RebH DWG Brominated Product
455.0581
457.0605

#### RebH EWG

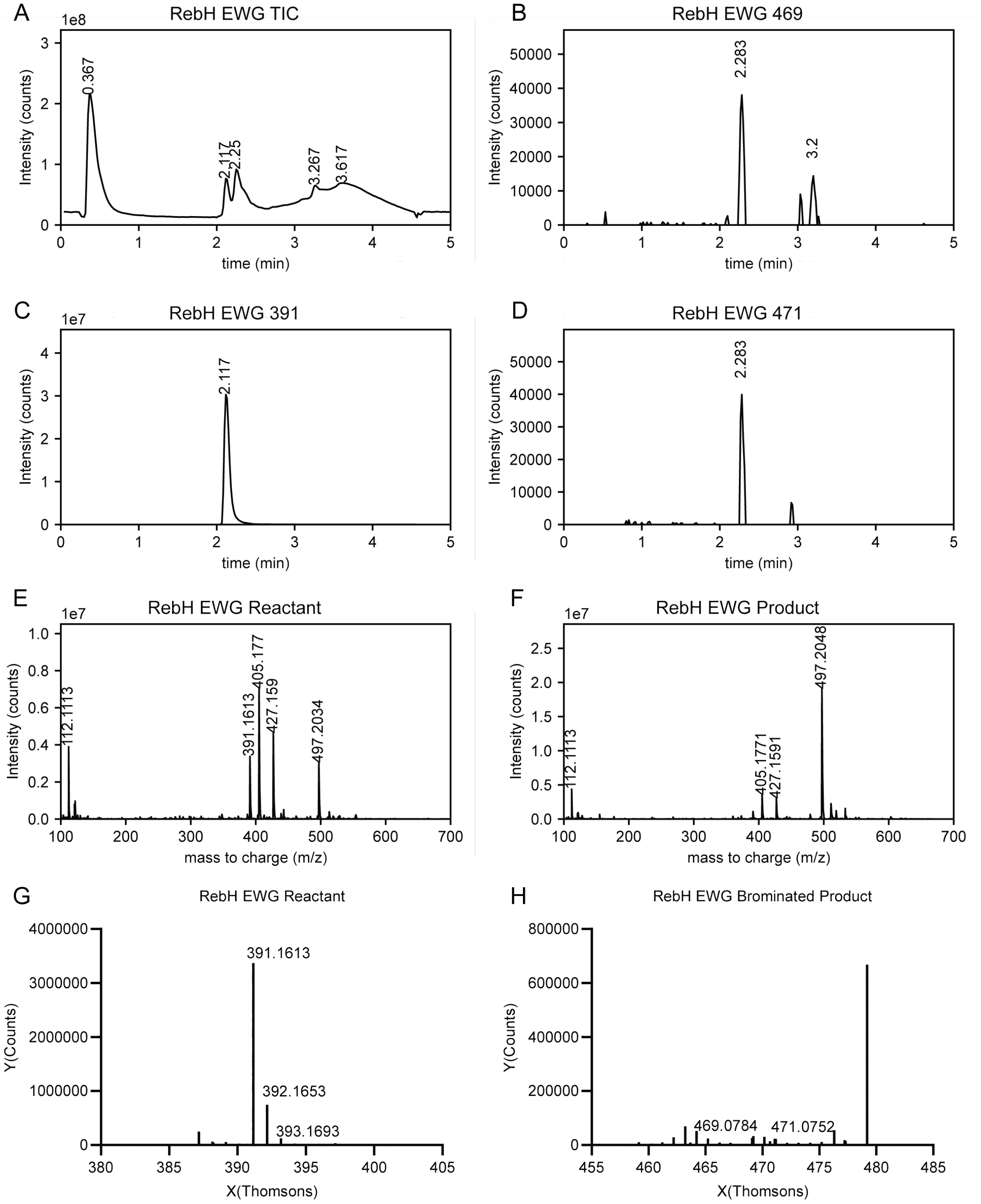
A
RebH EWG TIC
1e8
Intensity (counts)
time (min)
0.367
2.117
2.25
3.267
3.617
B
RebH EWG 469
Intensity (counts)
time (min)
2.283
3.2
C
RebH EWG 391
1e7
Intensity (counts)
time (min)
2.117
D
RebH EWG 471
Intensity (counts)
time (min)
2.283
E
RebH EWG Reactant
1e7
Intensity (counts)
mass to charge (m/z)
112.1113
391.1613
405.177
427.159
497.2034
F
RebH EWG Product
1e7
Intensity (counts)
mass to charge (m/z)
112.1113
405.1771
427.1591
497.2048
G
RebH EWG Reactant
Y(Counts)
X(Thomsons)
391.1613
392.1653
393.1693
H
RebH EWG Brominated Product
Y(Counts)
X(Thomsons)
469.0784
471.0752

#### RebH FWG

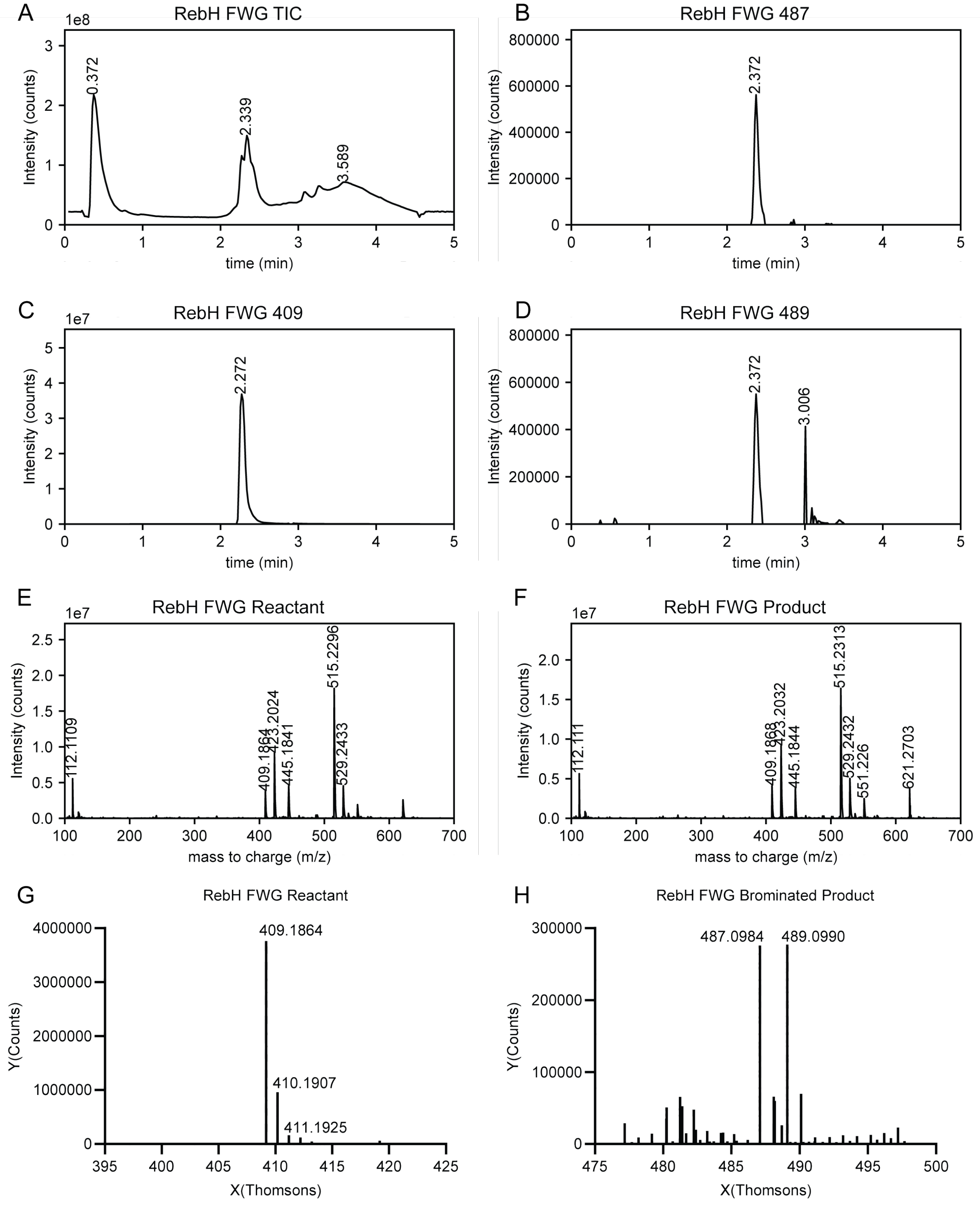

A
RebH FWG TIC
1e8
Intensity (counts)
time (min)
0.372
2.339
3.589
B
RebH FWG 487
Intensity (counts)
time (min)
2.372
C
RebH FWG 409
1e7
Intensity (counts)
time (min)
2.272
D
RebH FWG 489
Intensity (counts)
time (min)
2.372
3.006
E
RebH FWG Reactant
1e7
Intensity (counts)
mass to charge (m/z)
112.1109
409.1864
423.2024
445.1841
515.2296
529.2433
F
RebH FWG Product
1e7
Intensity (counts)
mass to charge (m/z)
112.111
409.1868
423.2032
445.1844
515.2313
529.2432
551.226
621.2703
G
RebH FWG Reactant
Y(Counts)
X(Thomsons)
409.1864
410.1907
411.1925
H
RebH FWG Brominated Product
Y(Counts)
X(Thomsons)
487.0984
489.0990

#### RebH HWG

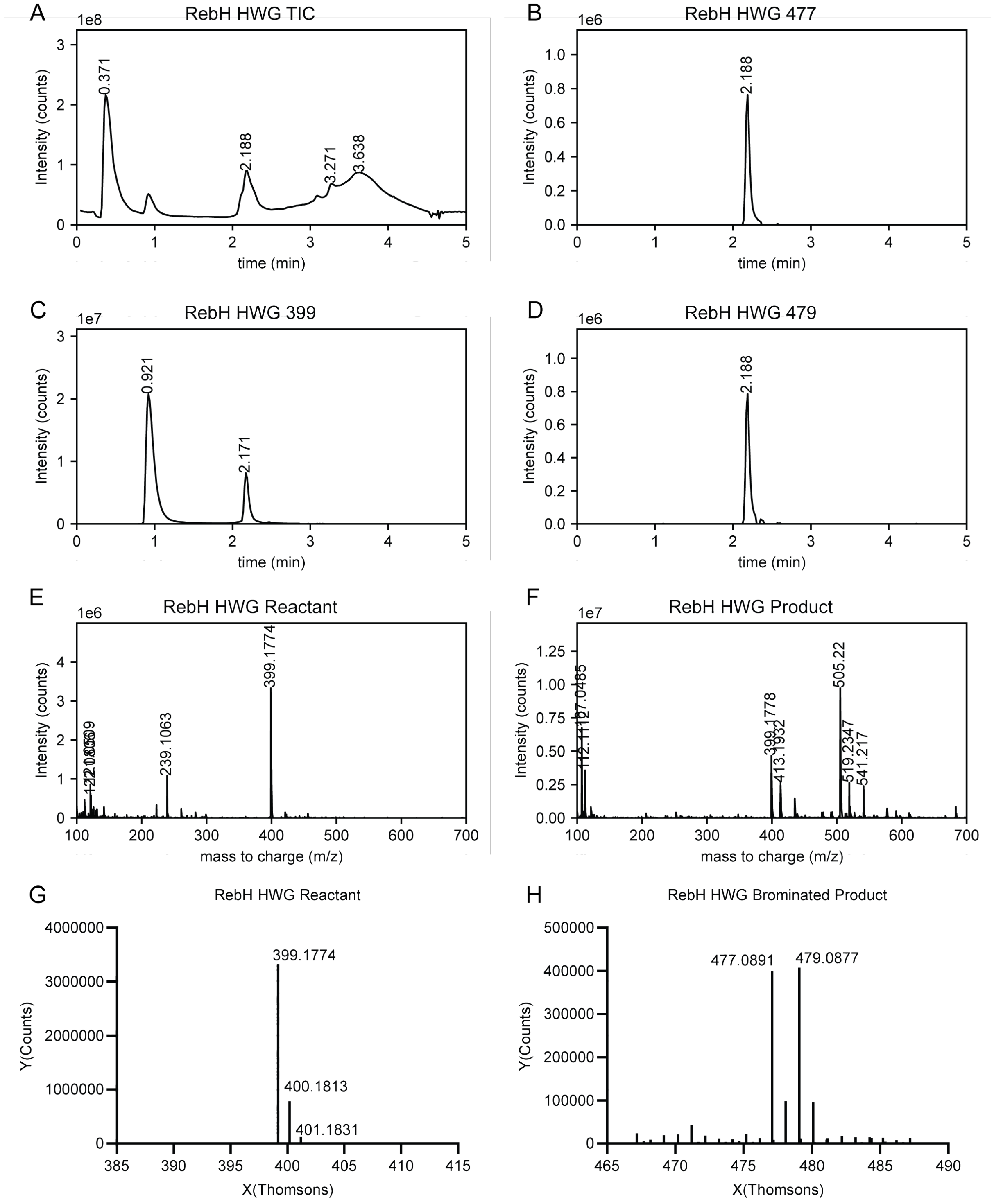

A
RebH HWG TIC
1e8
Intensity (counts)
time (min)
0.371
2.188
3.271
3.638
B
RebH HWG 477
1e6
Intensity (counts)
time (min)
2.188
C
RebH HWG 399
1e7
Intensity (counts)
time (min)
0.921
2.171
D
RebH HWG 479
1e6
Intensity (counts)
time (min)
2.188
E
RebH HWG Reactant
1e6
Intensity (counts)
mass to charge (m/z)
399.1774
239.1063
F
RebH HWG Product
1e7
Intensity (counts)
mass to charge (m/z)
505.22
399.1778
413.1932
519.2347
541.217
G
RebH HWG Reactant
Y(Counts)
X(Thomsons)
399.1774
400.1813
401.1831
H
RebH HWG Brominated Product
Y(Counts)
X(Thomsons)
477.0891
479.0877

#### RebH KWG

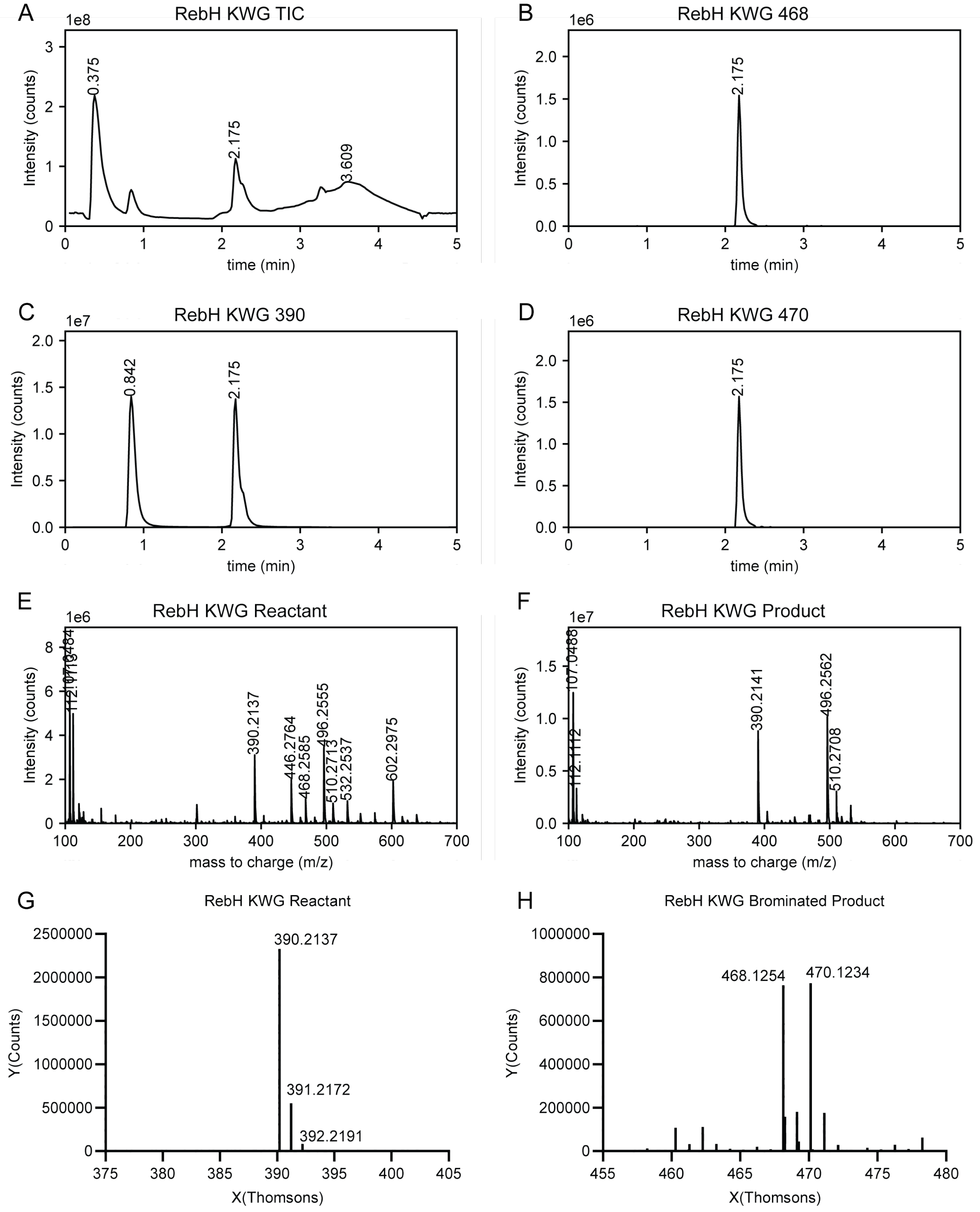
A
RebH KWG TIC
1e8
Intensity (counts)
time (min)
0.375
2.175
3.609
B
RebH KWG 468
1e6
Intensity (counts)
time (min)
2.175
C
RebH KWG 390
1e7
Intensity (counts)
time (min)
0.842
2.175
D
RebH KWG 470
1e6
Intensity (counts)
time (min)
2.175
E
RebH KWG Reactant
1e6
Intensity (counts)
mass to charge (m/z)
390.2137
446.2764
468.2585
496.2555
510.2713
532.2537
602.2975
F
RebH KWG Product
1e7
Intensity (counts)
mass to charge (m/z)
107.0488
112.1112
390.2141
496.2562
510.2708
G
RebH KWG Reactant
Y(Counts)
X(Thomsons)
390.2137
391.2172
392.2191
H
RebH KWG Brominated Product
Y(Counts)
X(Thomsons)
468.1254
470.1234

#### RebH LWG

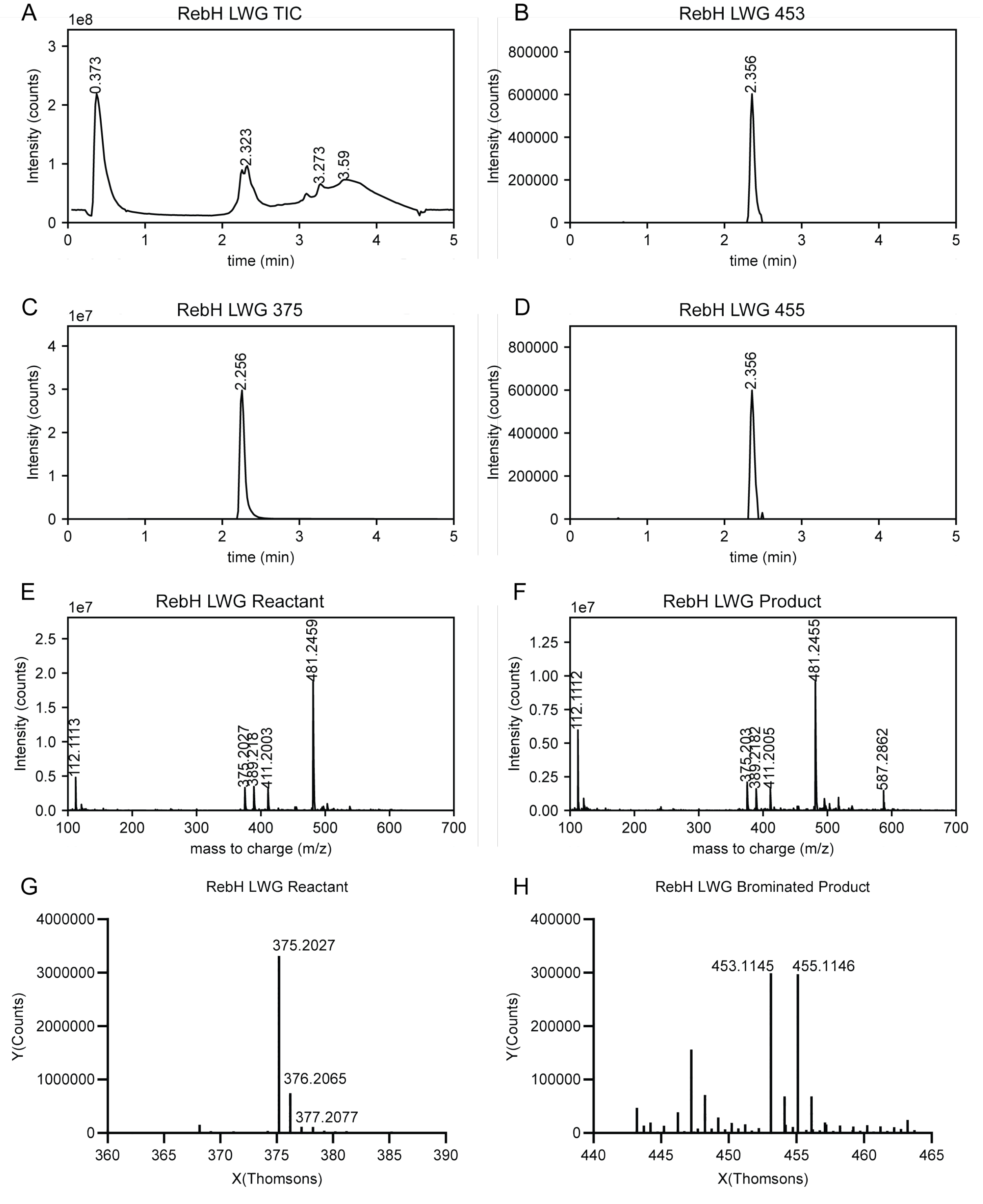
A
RebH LWG TIC
1e8
Intensity (counts)
time (min)
0.373
2.323
3.273
3.59
B
RebH LWG 453
2.356
C
RebH LWG 375
1e7
2.256
D
RebH LWG 455
2.356
E
RebH LWG Reactant
mass to charge (m/z)
112.1113
375.2027
389.218
411.2003
481.2459
F
RebH LWG Product
112.1112
375.203
389.2182
411.2005
481.2455
587.2862
G
RebH LWG Reactant
Y(Counts)
X(Thomsons)
375.2027
376.2065
377.2077
H
RebH LWG Brominated Product
453.1145
455.1146

#### RebH MWG

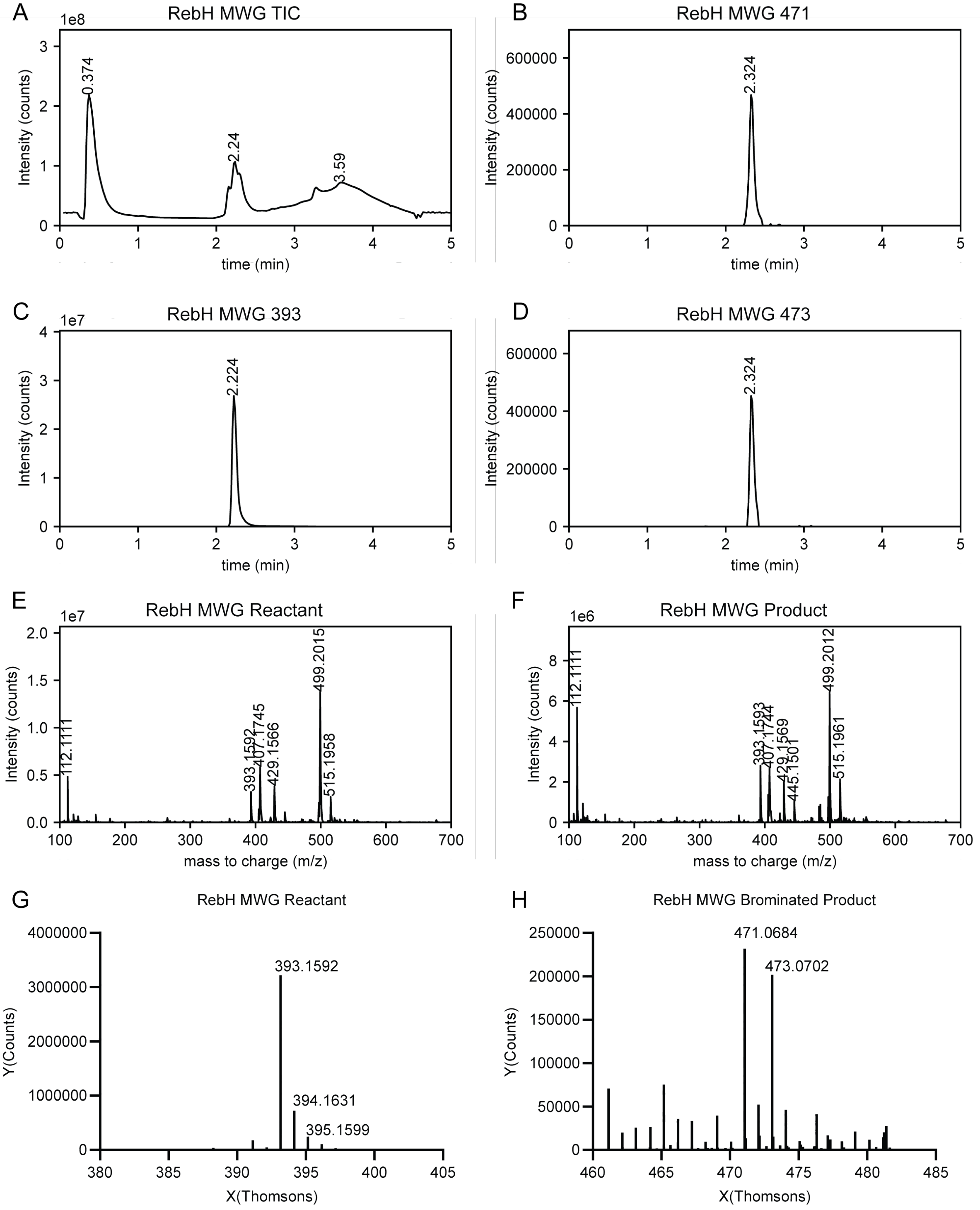

A
RebH MWG TIC
1e8
Intensity (counts)
time (min)
0.374
2.24
3.59
B
RebH MWG 471
Intensity (counts)
time (min)
2.324
C
RebH MWG 393
1e7
Intensity (counts)
time (min)
2.224
D
RebH MWG 473
Intensity (counts)
time (min)
2.324
E
RebH MWG Reactant
1e7
Intensity (counts)
mass to charge (m/z)
112.1111
393.1592
407.1745
429.1566
499.2015
515.1958
F
RebH MWG Product
1e6
Intensity (counts)
mass to charge (m/z)
112.1111
393.1593
407.1744
429.1569
445.1501
499.2012
515.1961
G
RebH MWG Reactant
Y(Counts)
X(Thomsons)
393.1592
394.1631
395.1599
H
RebH MWG Brominated Product
Y(Counts)
X(Thomsons)
471.0684
473.0702

#### RebH NWG

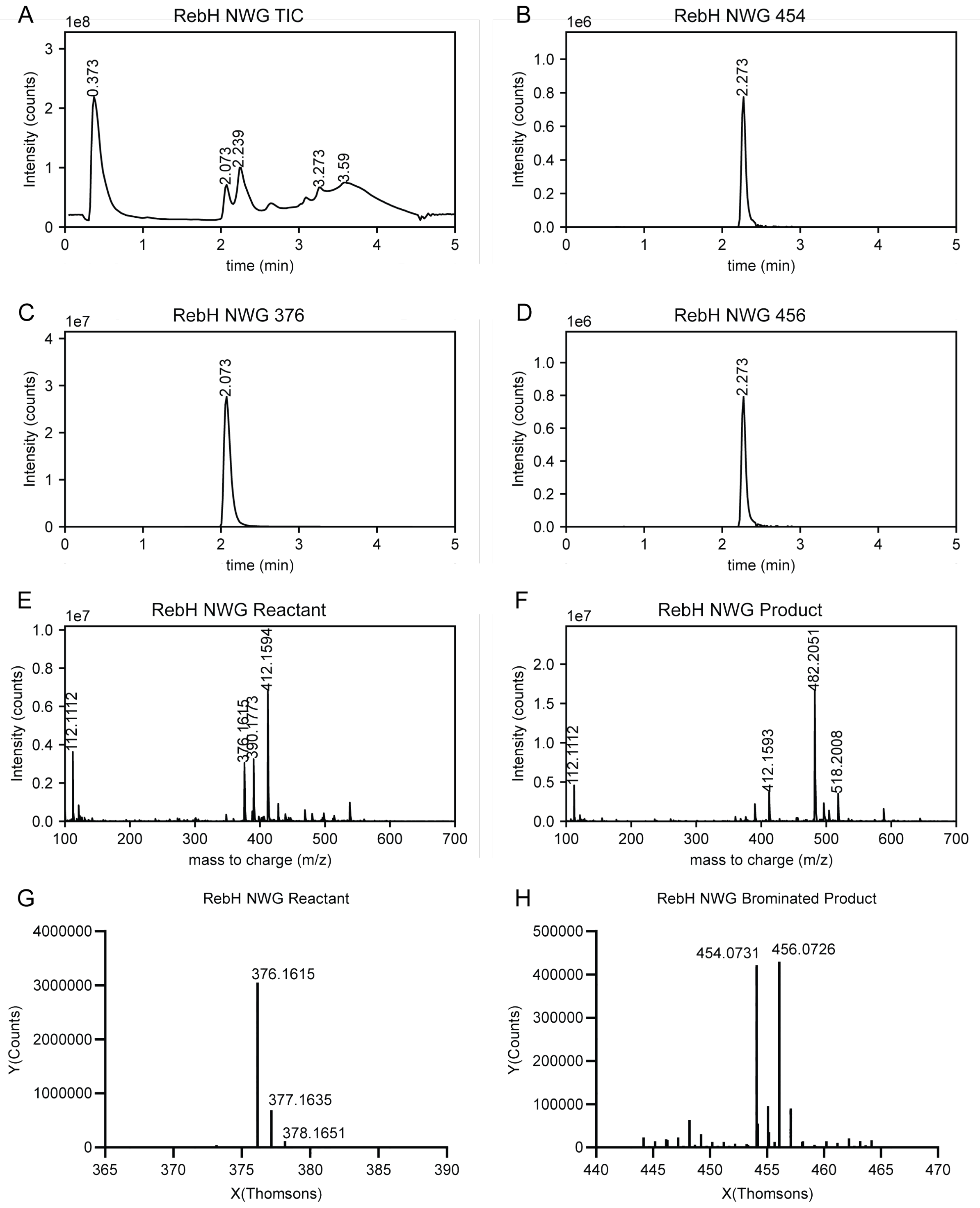
A
RebH NWG TIC
1e8
Intensity (counts)
time (min)
0.373
2.073
2.239
3.273
3.59
B
RebH NWG 454
1e6
Intensity (counts)
time (min)
2.273
C
RebH NWG 376
1e7
Intensity (counts)
time (min)
2.073
D
RebH NWG 456
1e6
Intensity (counts)
time (min)
2.273
E
RebH NWG Reactant
1e7
Intensity (counts)
mass to charge (m/z)
112.1112
376.1615
390.1773
412.1594
F
RebH NWG Product
1e7
Intensity (counts)
mass to charge (m/z)
112.1112
412.1593
482.2051
518.2008
G
RebH NWG Reactant
Y(Counts)
X(Thomsons)
376.1615
377.1635
378.1651
H
RebH NWG Brominated Product
Y(Counts)
X(Thomsons)
454.0731
456.0726

#### RebH PWG

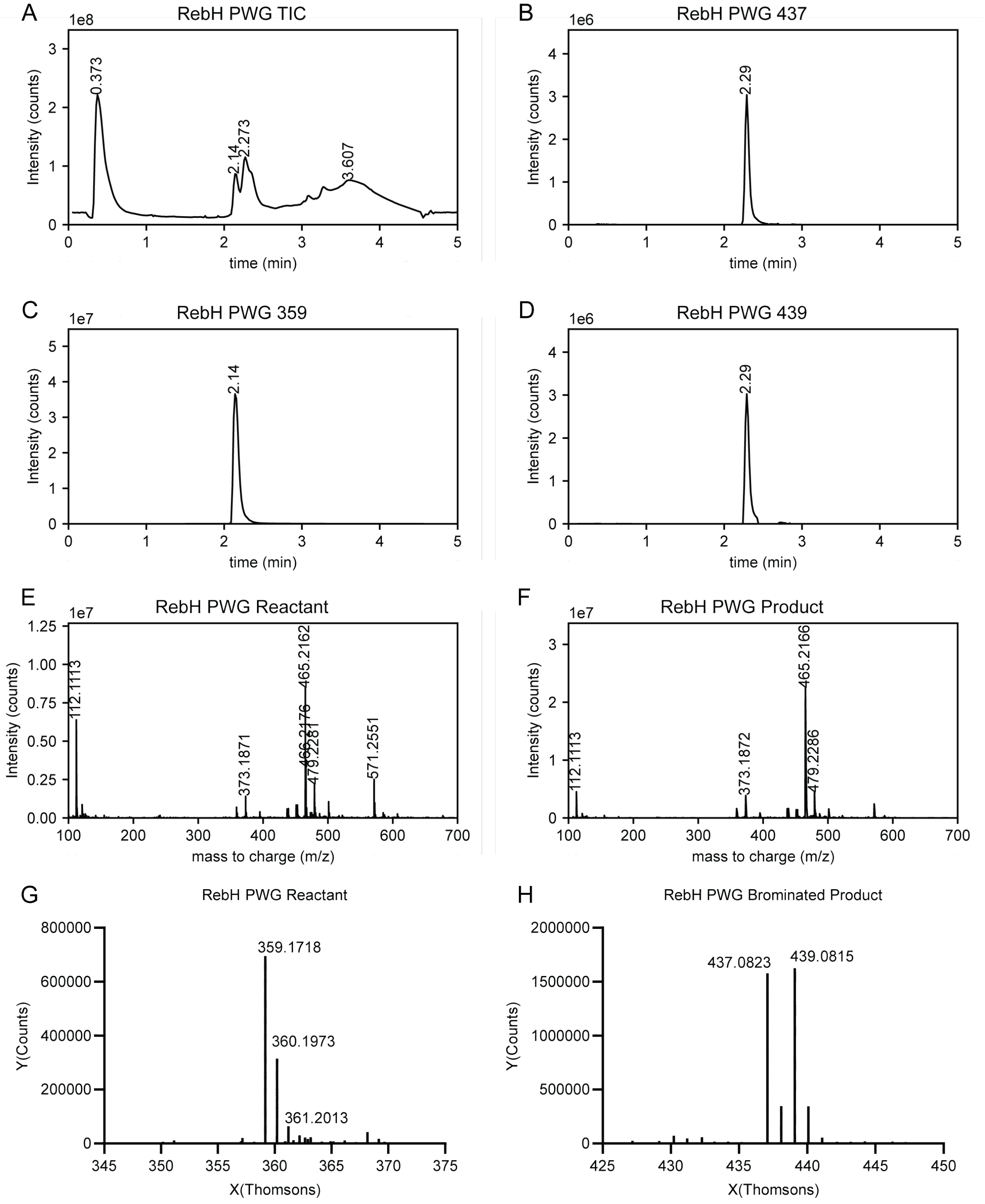
A
RebH PWG TIC
1e8
Intensity (counts)
time (min)
0.373
2.14
2.273
3.607
B
RebH PWG 437
1e6
2.29
C
RebH PWG 359
1e7
2.14
D
RebH PWG 439
1e6
2.29
E
RebH PWG Reactant
1e7
mass to charge (m/z)
112.1113
373.1871
465.2162
466.2176
479.2281
571.2551
F
RebH PWG Product
1e7
112.1113
373.1872
465.2166
479.2286
G
RebH PWG Reactant
Y(Counts)
X(Thomsons)
359.1718
360.1973
361.2013
H
RebH PWG Brominated Product
437.0823
439.0815

#### RebH QWG

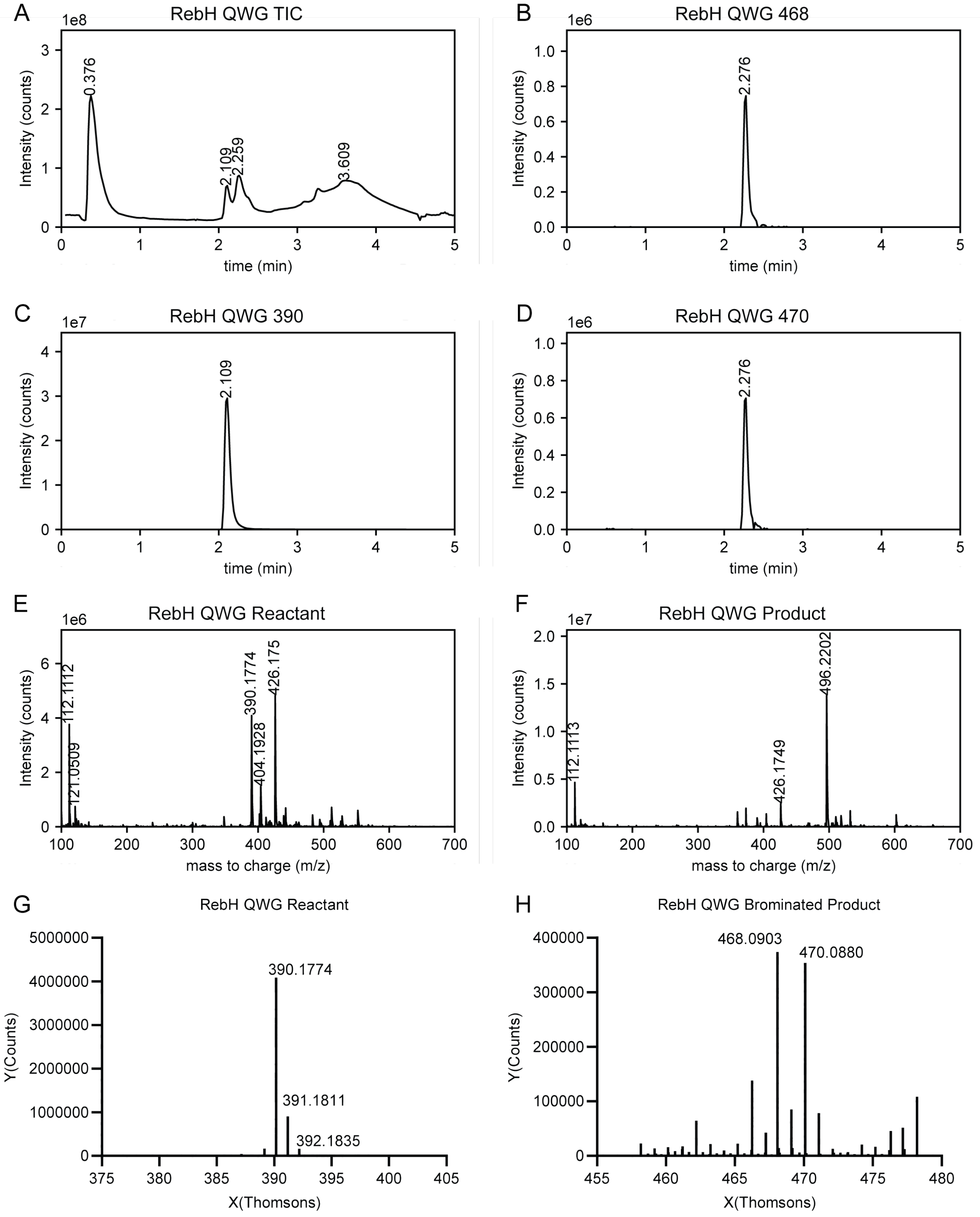
A
RebH QWG TIC
1e8
Intensity (counts)
time (min)
0.376
2.109
2.259
3.609
B
RebH QWG 468
1e6
Intensity (counts)
time (min)
2.276
C
RebH QWG 390
1e7
Intensity (counts)
time (min)
2.109
D
RebH QWG 470
1e6
Intensity (counts)
time (min)
2.276
E
RebH QWG Reactant
1e6
Intensity (counts)
mass to charge (m/z)
112.1112
121.0509
390.1774
404.1928
426.175
F
RebH QWG Product
1e7
Intensity (counts)
mass to charge (m/z)
112.1113
426.1749
496.2202
G
RebH QWG Reactant
Y(Counts)
X(Thomsons)
390.1774
391.1811
392.1835
H
RebH QWG Brominated Product
Y(Counts)
X(Thomsons)
468.0903
470.0880

**RebH RWG**

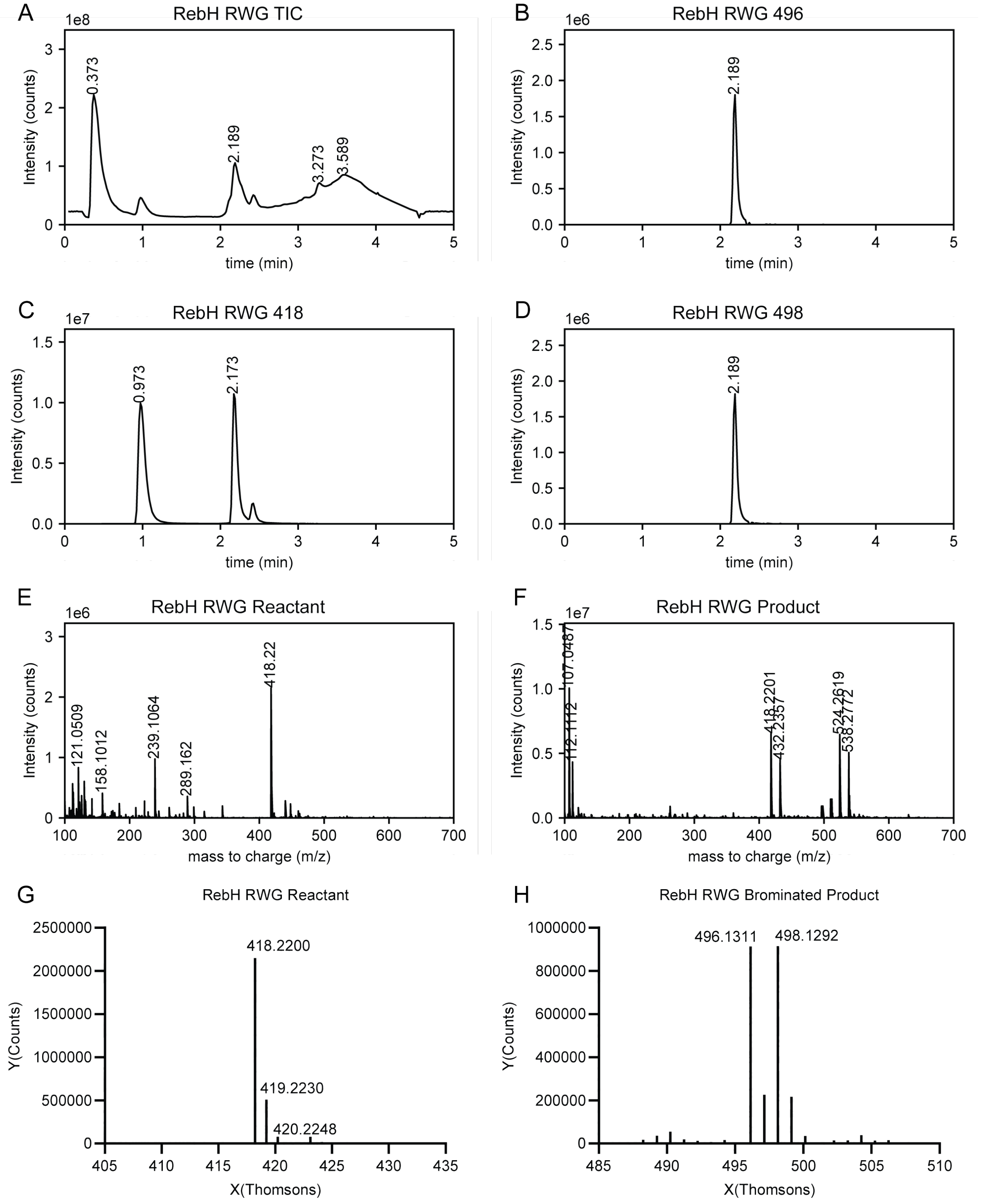

A
RebH RWG TIC
1e8
Intensity (counts)
time (min)
0.373
2.189
3.273
3.589
B
RebH RWG 496
1e6
2.189
C
RebH RWG 418
1e7
0.973
2.173
D
RebH RWG 498
1e6
2.189
E
RebH RWG Reactant
1e6
mass to charge (m/z)
121.0509
158.1012
239.1064
289.162
418.22
F
RebH RWG Product
1e7
107.0487
112.1112
418.2201
432.2357
524.2619
538.2772
G
RebH RWG Reactant
Y(Counts)
X(Thomsons)
418.2200
419.2230
420.2248
H
RebH RWG Brominated Product
496.1311
498.1292

#### RebH TWG

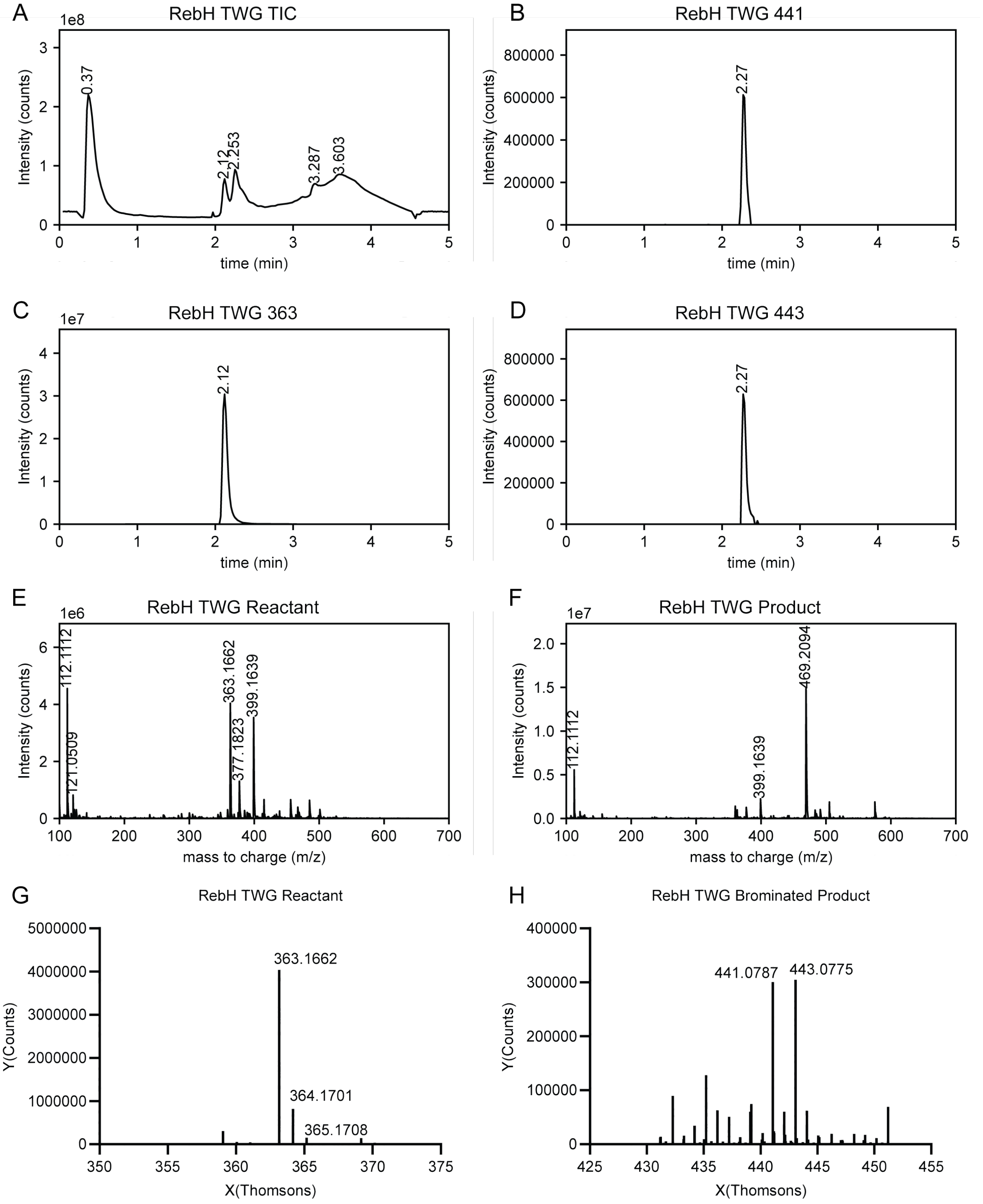
A
RebH TWG TIC
1e8
Intensity (counts)
time (min)
0.37
2.12
2.253
3.287
3.603
B
RebH TWG 441
Intensity (counts)
time (min)
2.27
C
RebH TWG 363
1e7
Intensity (counts)
time (min)
2.12
D
RebH TWG 443
Intensity (counts)
time (min)
2.27
E
RebH TWG Reactant
1e6
Intensity (counts)
mass to charge (m/z)
112.1112
121.0509
363.1662
377.1823
399.1639
F
RebH TWG Product
1e7
Intensity (counts)
mass to charge (m/z)
112.1112
399.1639
469.2094
G
RebH TWG Reactant
Y(Counts)
X(Thomsons)
363.1662
364.1701
365.1708
H
RebH TWG Brominated Product
Y(Counts)
X(Thomsons)
441.0787
443.0775

#### RebH VWG

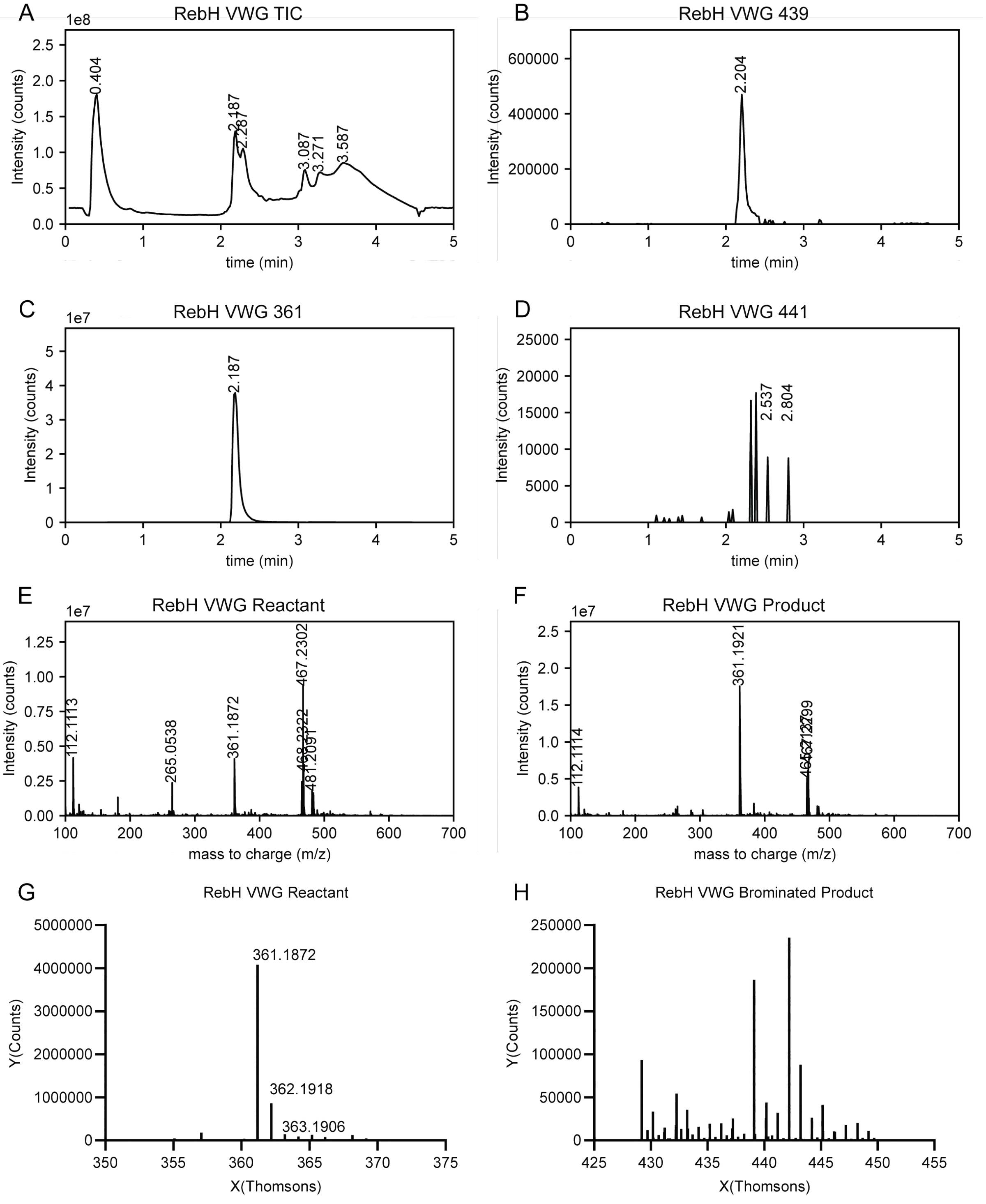
A
RebH VWG TIC
1e8
Intensity (counts)
time (min)
0.404
2.187
2.287
3.087
3.271
3.587
B
RebH VWG 439
Intensity (counts)
time (min)
2.204
C
RebH VWG 361
1e7
Intensity (counts)
time (min)
2.187
D
RebH VWG 441
Intensity (counts)
time (min)
2.537
2.804
E
RebH VWG Reactant
1e7
Intensity (counts)
mass to charge (m/z)
112.1113
265.0538
361.1872
467.2302
468.2322
481.2091
F
RebH VWG Product
1e7
Intensity (counts)
mass to charge (m/z)
112.1114
361.1921
G
RebH VWG Reactant
Y(Counts)
X(Thomsons)
361.1872
362.1918
363.1906
H
RebH VWG Brominated Product
Y(Counts)
X(Thomsons)

#### RebH YWG

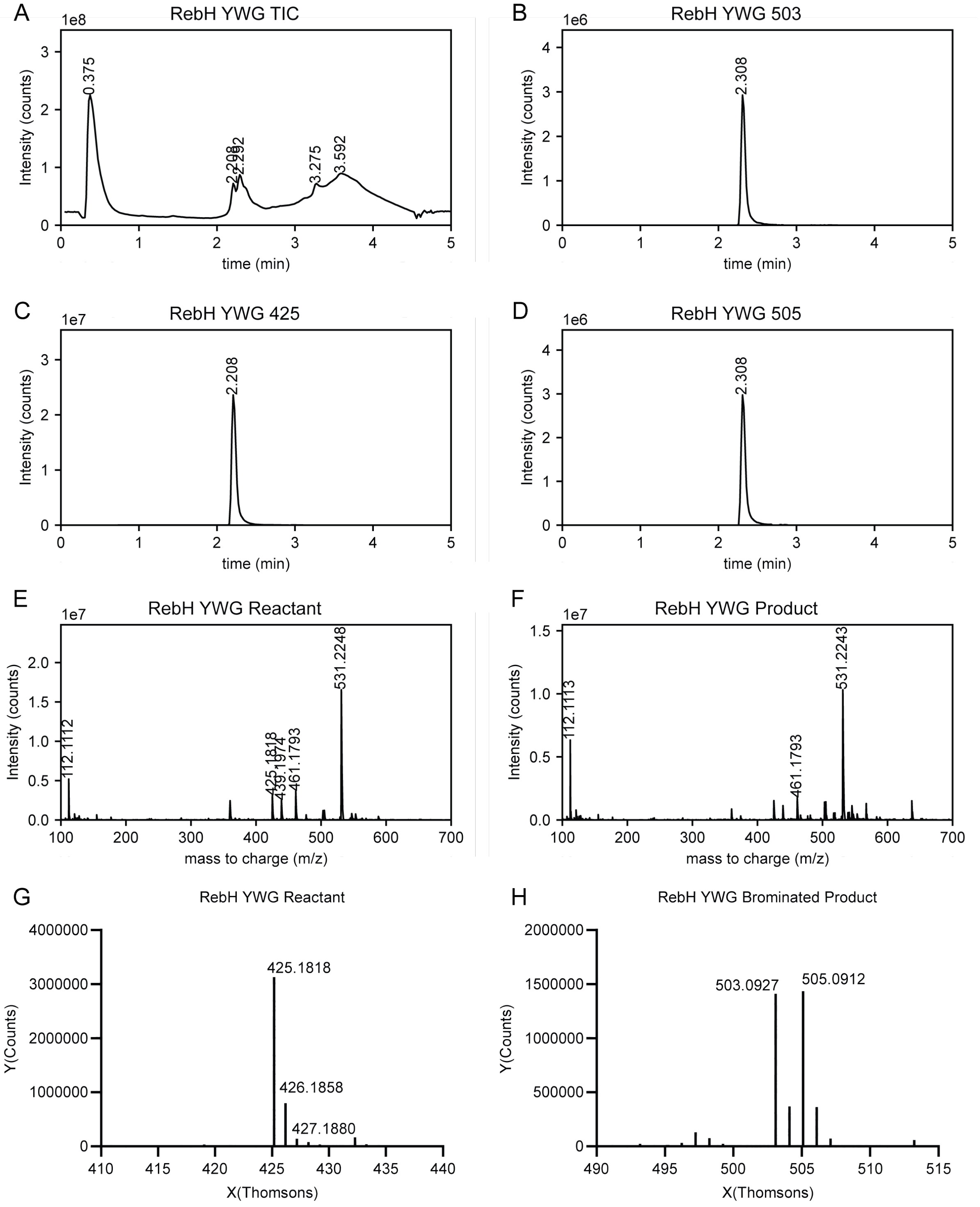

A
RebH YWG TIC
1e8
Intensity (counts)
0.375
2.208
2.292
3.275
3.592
time (min)
B
RebH YWG 503
1e6
2.308
C
RebH YWG 425
1e7
2.208
D
RebH YWG 505
1e6
2.308
E
RebH YWG Reactant
1e7
112.1112
425.1818
439.1974
461.1793
531.2248
mass to charge (m/z)
F
RebH YWG Product
1e7
112.1113
461.1793
531.2243
G
RebH YWG Reactant
Y(Counts)
425.1818
426.1858
427.1880
X(Thomsons)
H
RebH YWG Brominated Product
503.0927
505.0912

#### RebH GWA

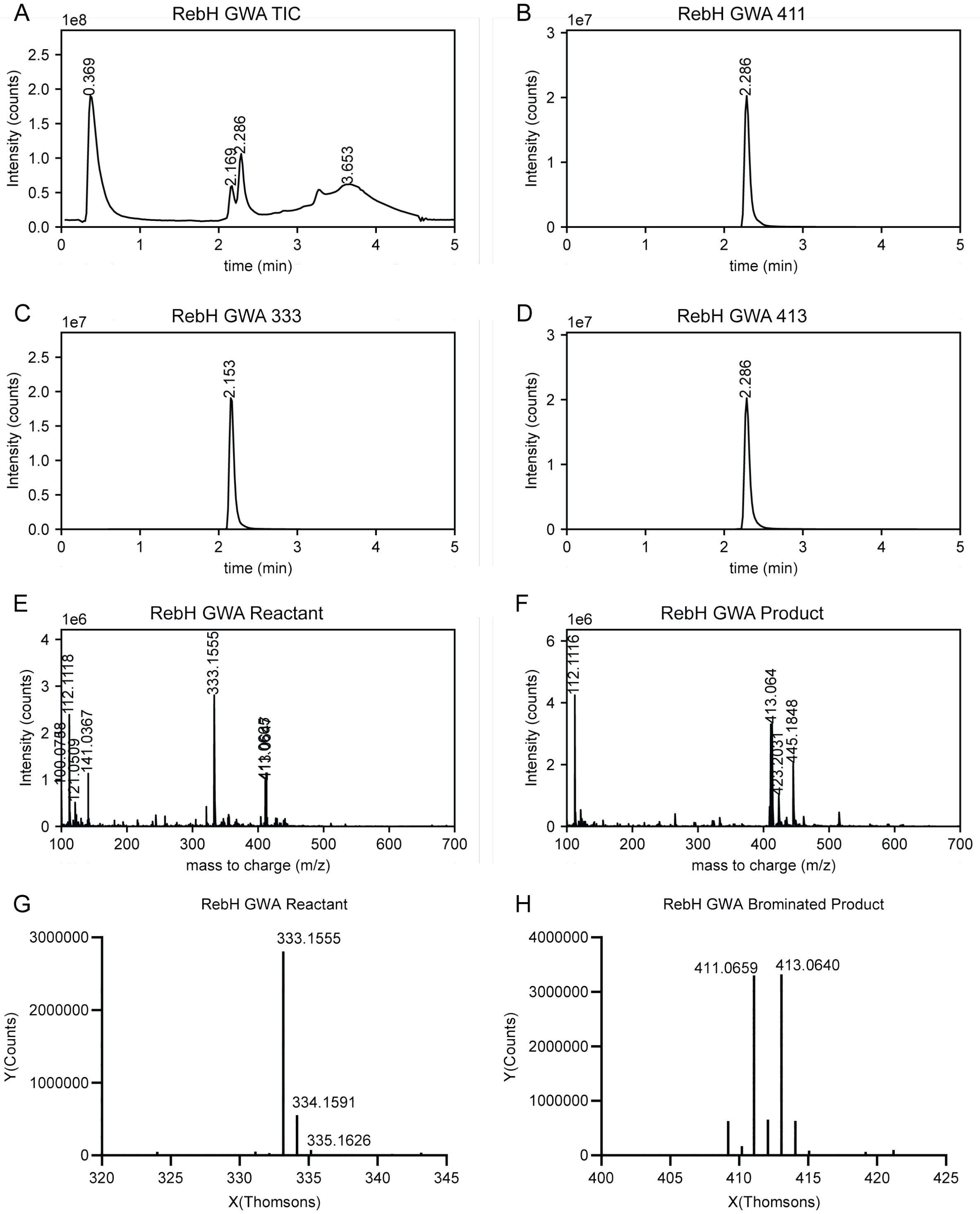

A
RebH GWA TIC
1e8
Intensity (counts)
time (min)
0.369
2.169
2.286
3.653
B
RebH GWA 411
1e7
Intensity (counts)
time (min)
2.286
C
RebH GWA 333
1e7
Intensity (counts)
time (min)
2.153
D
RebH GWA 413
1e7
Intensity (counts)
time (min)
2.286
E
RebH GWA Reactant
1e6
Intensity (counts)
mass to charge (m/z)
333.1555
112.1118
100.0758
121.0509
141.0367
F
RebH GWA Product
1e6
Intensity (counts)
mass to charge (m/z)
112.1116
413.064
445.1848
423.2031
G
RebH GWA Reactant
Y(Counts)
X(Thomsons)
333.1555
334.1591
335.1626
H
RebH GWA Brominated Product
Y(Counts)
X(Thomsons)
411.0659
413.0640

#### RebH GWC

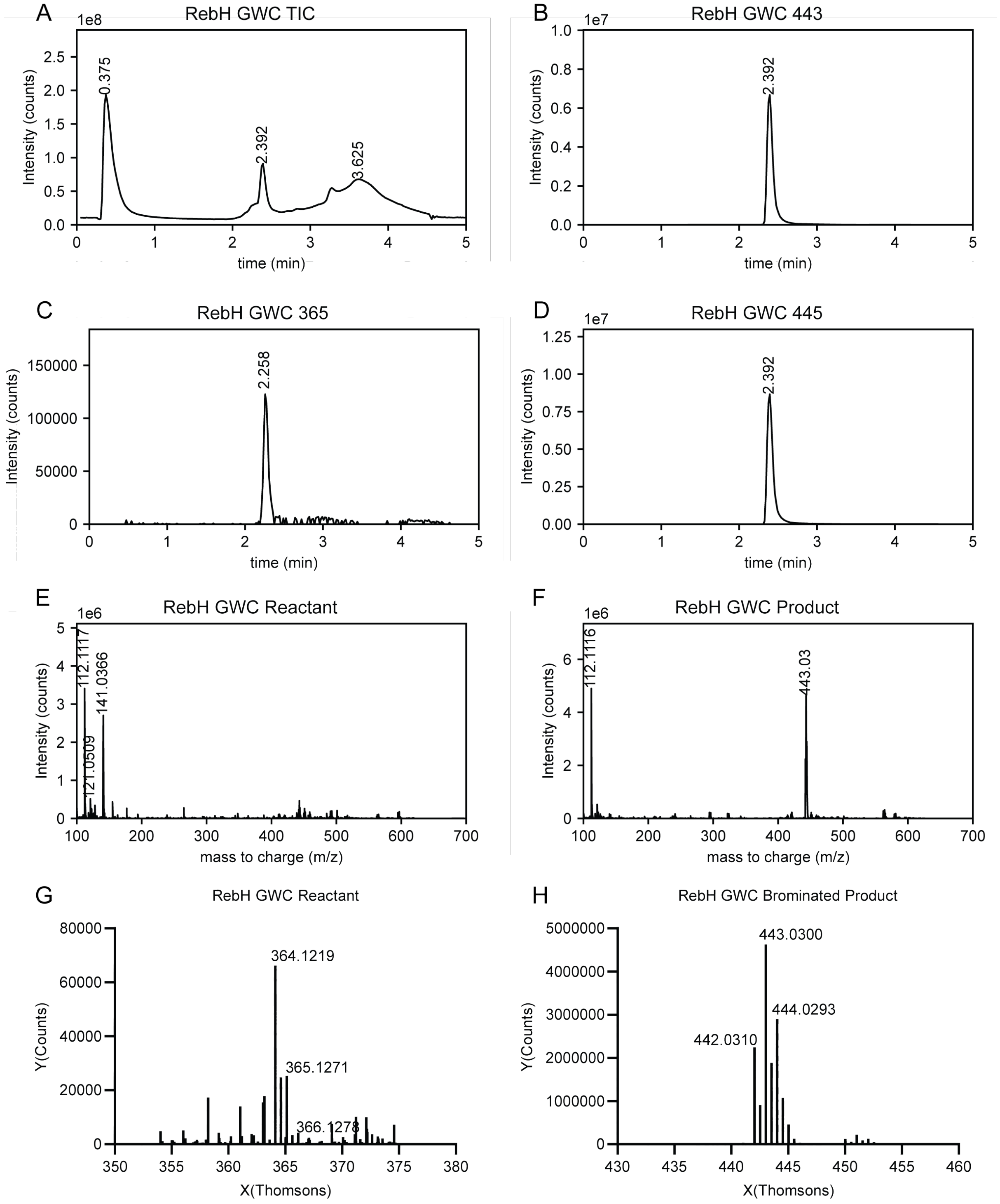
A
RebH GWC TIC
1e8
Intensity (counts)
time (min)
0.375
2.392
3.625
B
RebH GWC 443
1e7
2.392
C
RebH GWC 365
2.258
D
RebH GWC 445
1e7
2.392
E
RebH GWC Reactant
1e6
112.1117
141.0366
121.0509
mass to charge (m/z)
F
RebH GWC Product
1e6
112.1116
443.03
G
RebH GWC Reactant
364.1219
365.1271
366.1278
Y(Counts)
X(Thomsons)
H
RebH GWC Brominated Product
443.0300
444.0293
442.0310

#### RebH GWD

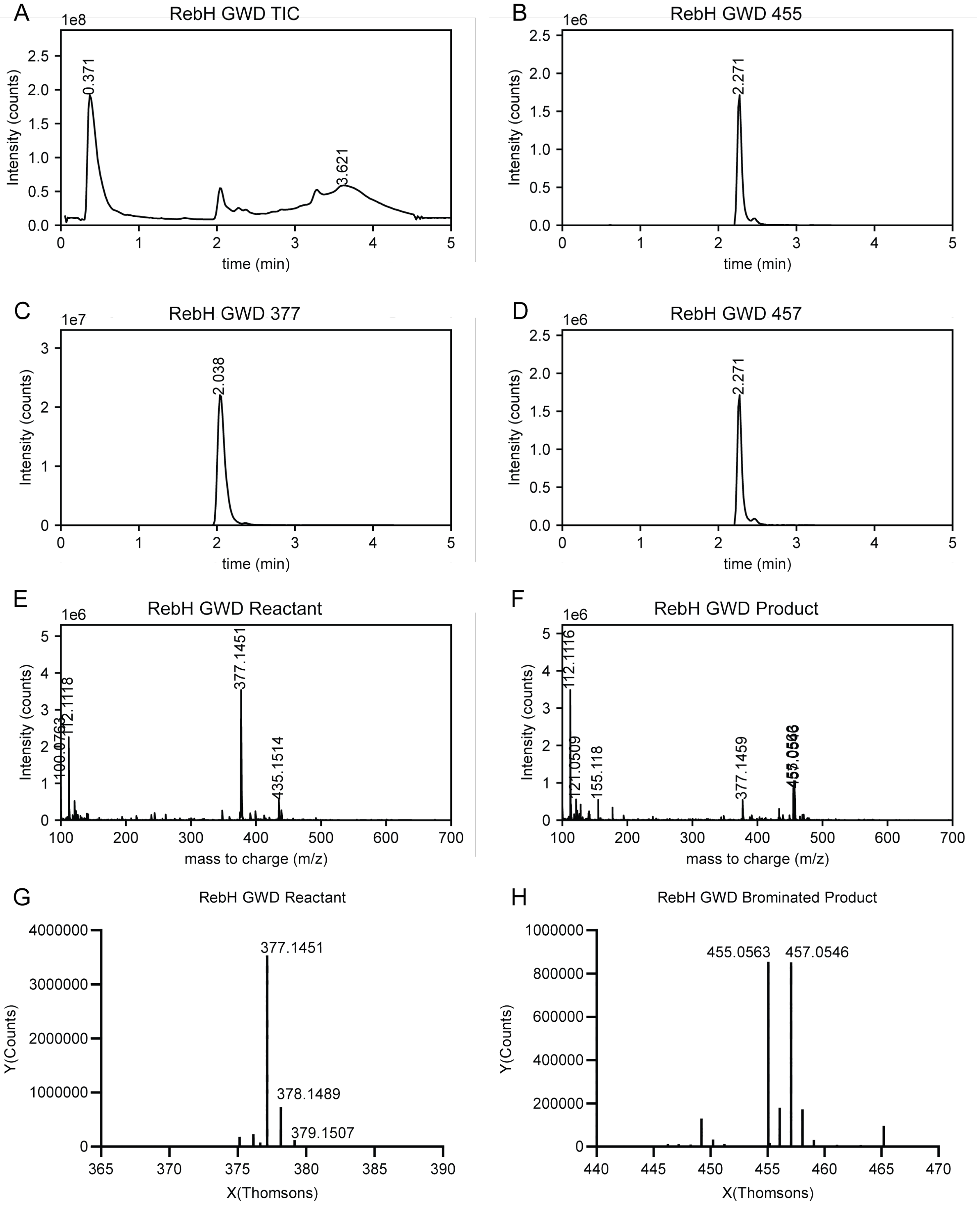

A
RebH GWD TIC
1e8
Intensity (counts)
time (min)
0.371
3.621
B
RebH GWD 455
1e6
Intensity (counts)
time (min)
2.271
C
RebH GWD 377
1e7
Intensity (counts)
time (min)
2.038
D
RebH GWD 457
1e6
Intensity (counts)
time (min)
2.271
E
RebH GWD Reactant
1e6
Intensity (counts)
mass to charge (m/z)
100.0763
112.1118
377.1451
435.1514
F
RebH GWD Product
1e6
Intensity (counts)
mass to charge (m/z)
112.1116
121.0509
155.118
377.1459
G
RebH GWD Reactant
Y(Counts)
X(Thomsons)
377.1451
378.1489
379.1507
H
RebH GWD Brominated Product
Y(Counts)
X(Thomsons)
455.0563
457.0546

#### RebH GWE

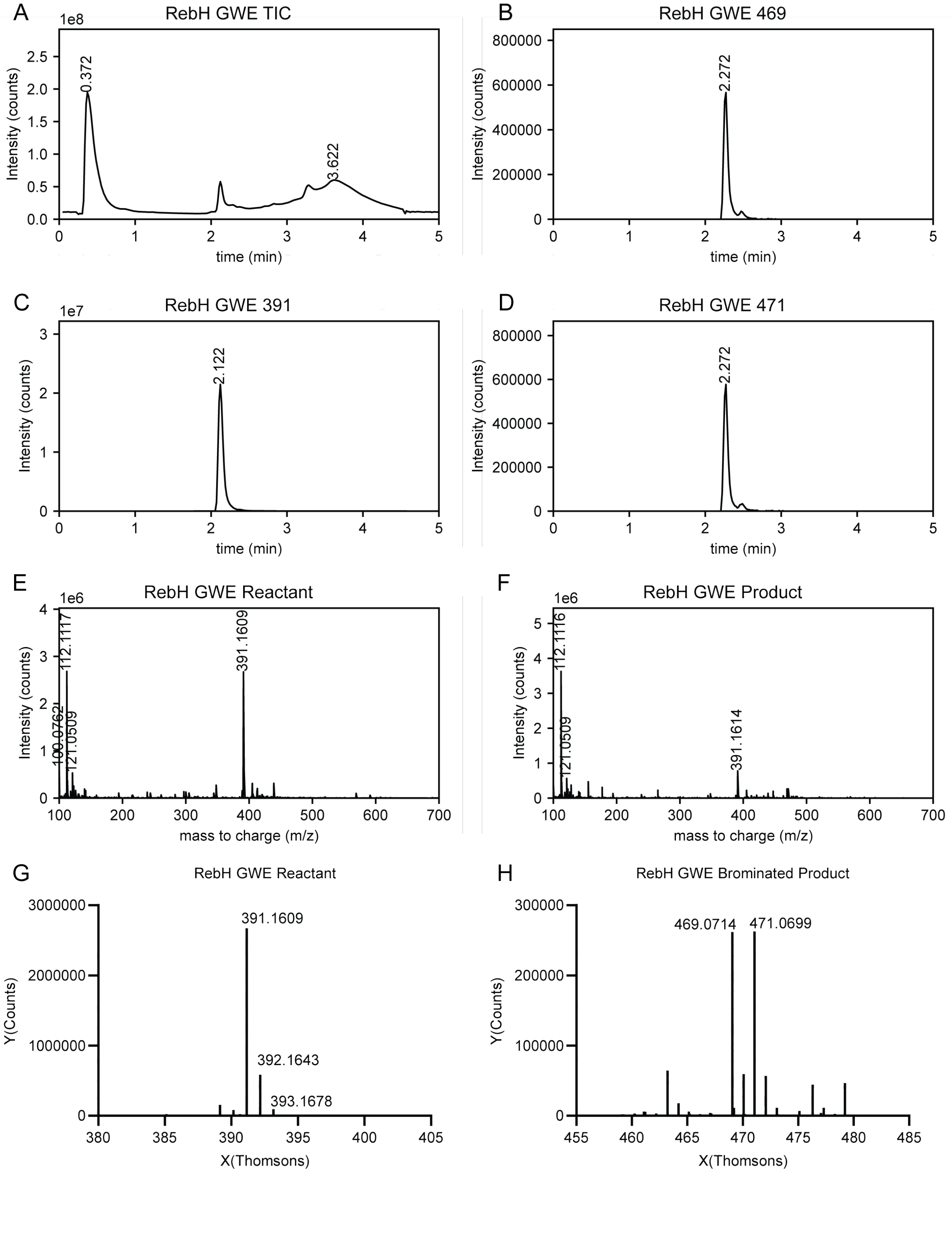

A
RebH GWE TIC
1e8
Intensity (counts)
time (min)
0.372
3.622
B
RebH GWE 469
Intensity (counts)
time (min)
2.272
C
RebH GWE 391
1e7
Intensity (counts)
time (min)
2.122
D
RebH GWE 471
Intensity (counts)
time (min)
2.272
E
RebH GWE Reactant
1e6
Intensity (counts)
mass to charge (m/z)
112.1117
100.0762
121.0509
391.1609
F
RebH GWE Product
1e6
Intensity (counts)
mass to charge (m/z)
112.1116
121.0509
391.1614
G
RebH GWE Reactant
Y(Counts)
X(Thomsons)
391.1609
392.1643
393.1678
H
RebH GWE Brominated Product
Y(Counts)
X(Thomsons)
469.0714
471.0699

#### RebH GWF

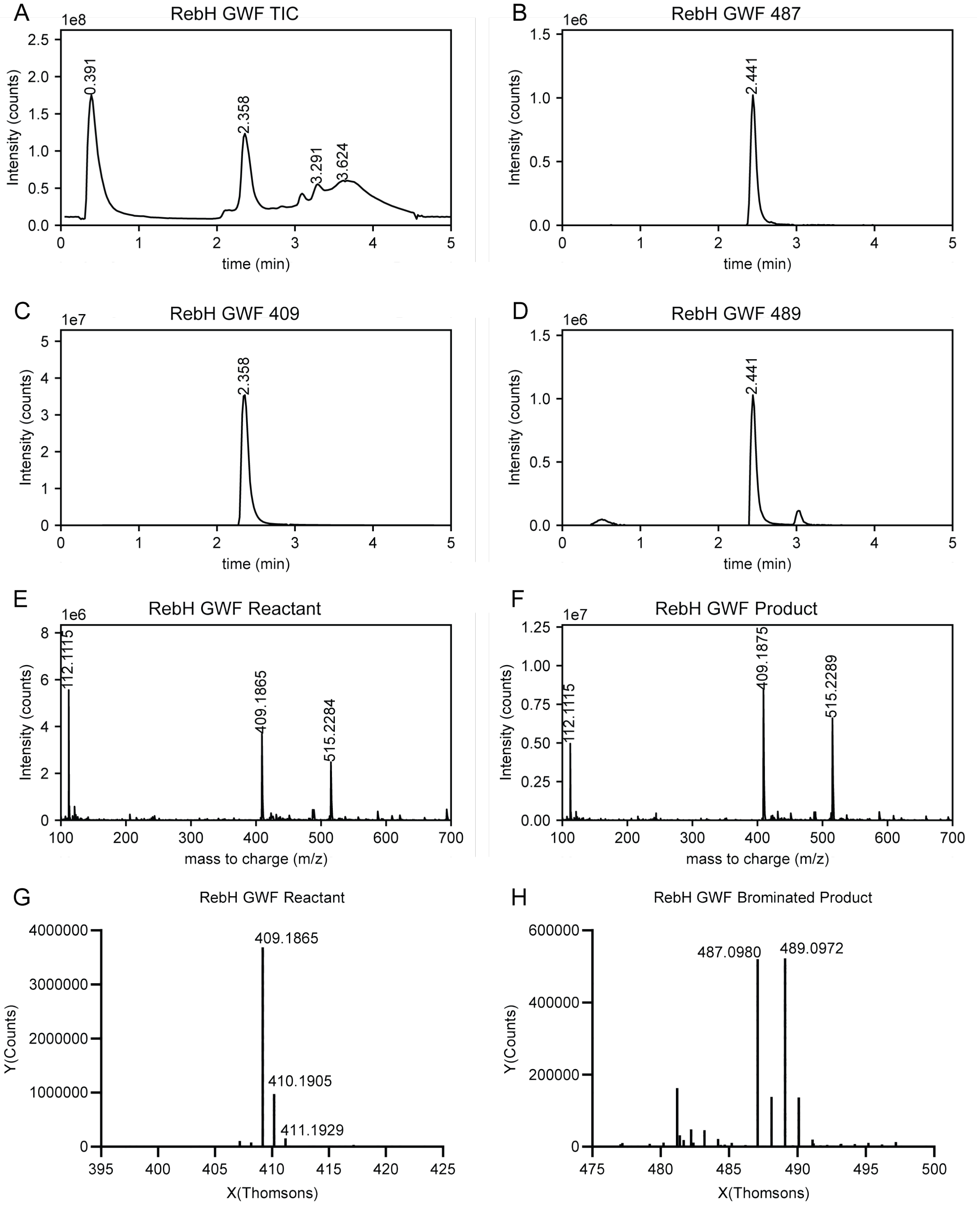
A
RebH GWF TIC
1e8
Intensity (counts)
time (min)
0.391
2.358
3.291
3.624
B
RebH GWF 487
1e6
2.441
C
RebH GWF 409
1e7
2.358
D
RebH GWF 489
1e6
2.441
E
RebH GWF Reactant
1e6
112.1115
409.1865
515.2284
mass to charge (m/z)
F
RebH GWF Product
1e7
112.1115
409.1875
515.2289
G
RebH GWF Reactant
409.1865
410.1905
411.1929
Y(Counts)
X(Thomsons)
H
RebH GWF Brominated Product
487.0980
489.0972

#### RebH GWH

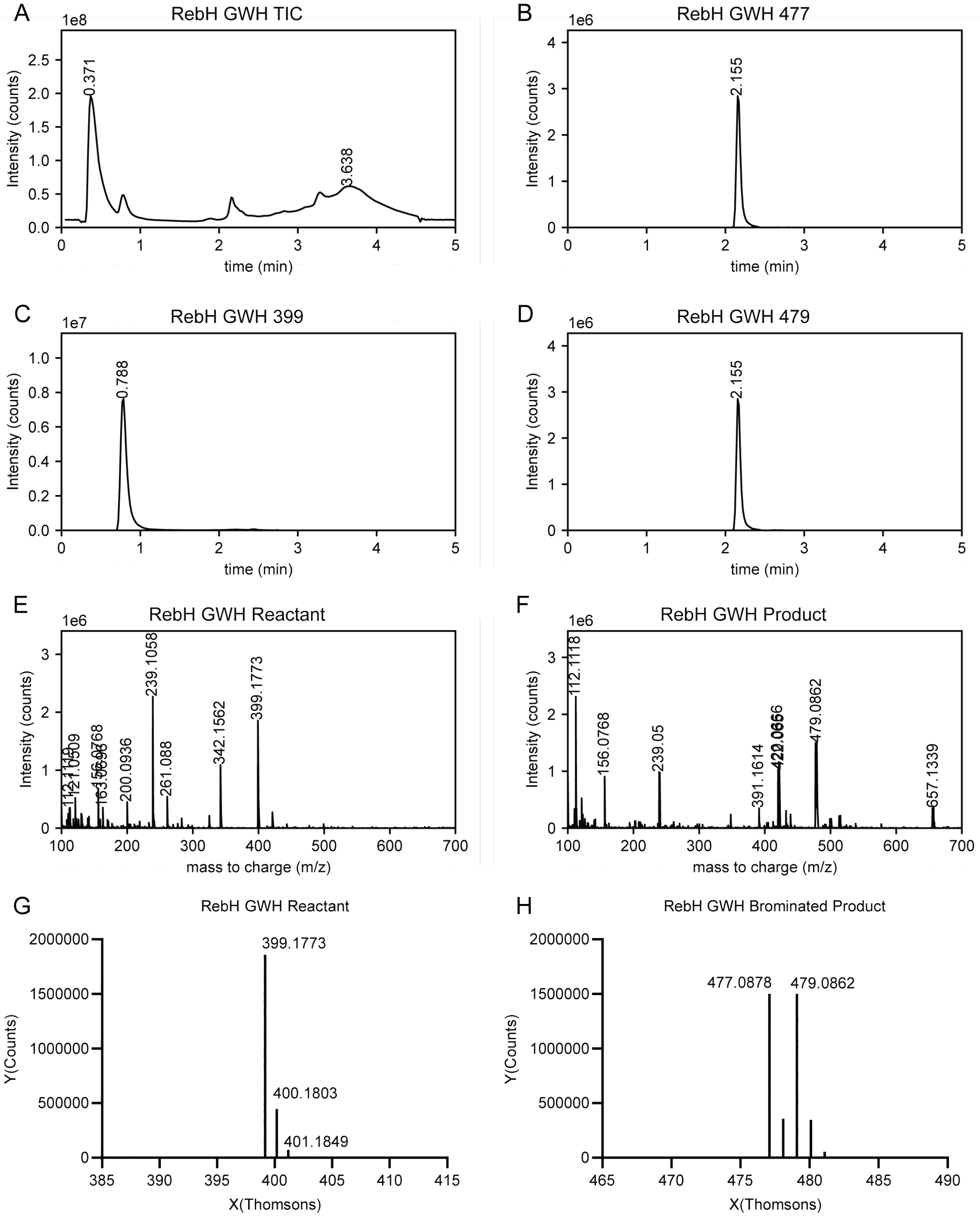
A
RebH GWH TIC
1e8
Intensity (counts)
time (min)
0.371
3.638
B
RebH GWH 477
1e6
Intensity (counts)
time (min)
2.155
C
RebH GWH 399
1e7
Intensity (counts)
time (min)
0.788
D
RebH GWH 479
1e6
Intensity (counts)
time (min)
2.155
E
RebH GWH Reactant
1e6
Intensity (counts)
mass to charge (m/z)
239.1058
399.1773
342.1562
200.0936
261.088
F
RebH GWH Product
1e6
Intensity (counts)
mass to charge (m/z)
112.1118
156.0768
239.05
391.1614
479.0862
657.1339
G
RebH GWH Reactant
Y(Counts)
X(Thomsons)
399.1773
400.1803
401.1849
H
RebH GWH Brominated Product
Y(Counts)
X(Thomsons)
477.0878
479.0862

#### RebH GWK

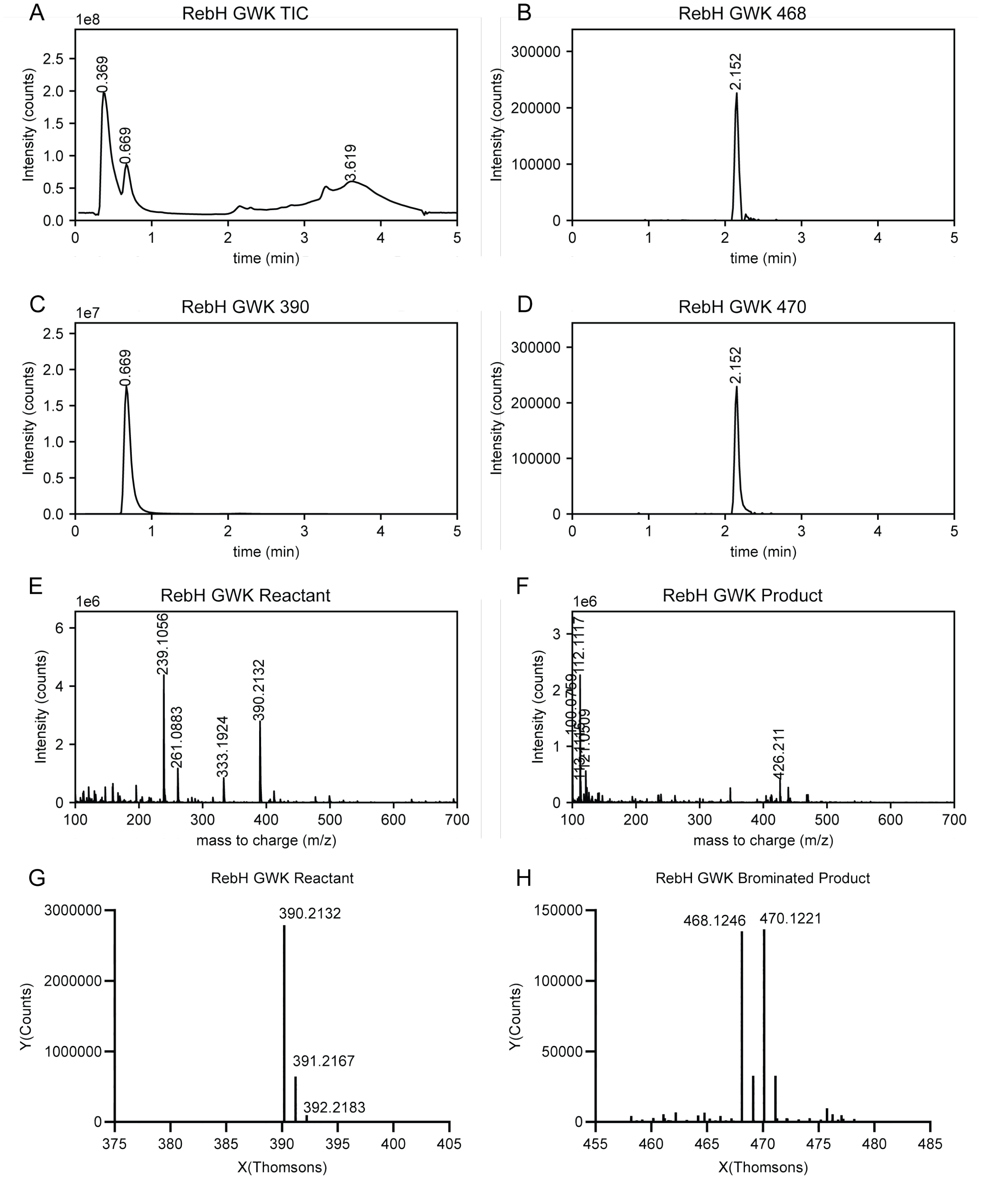

A
RebH GWK TIC
1e8
Intensity (counts)
time (min)
0.369
0.669
3.619
B
RebH GWK 468
2.152
C
RebH GWK 390
1e7
0.669
D
RebH GWK 470
2.152
E
RebH GWK Reactant
1e6
mass to charge (m/z)
239.1056
261.0883
333.1924
390.2132
F
RebH GWK Product
1e6
112.1117
426.211
G
RebH GWK Reactant
Y(Counts)
X(Thomsons)
390.2132
391.2167
392.2183
H
RebH GWK Brominated Product
468.1246
470.1221

#### RebH GWL

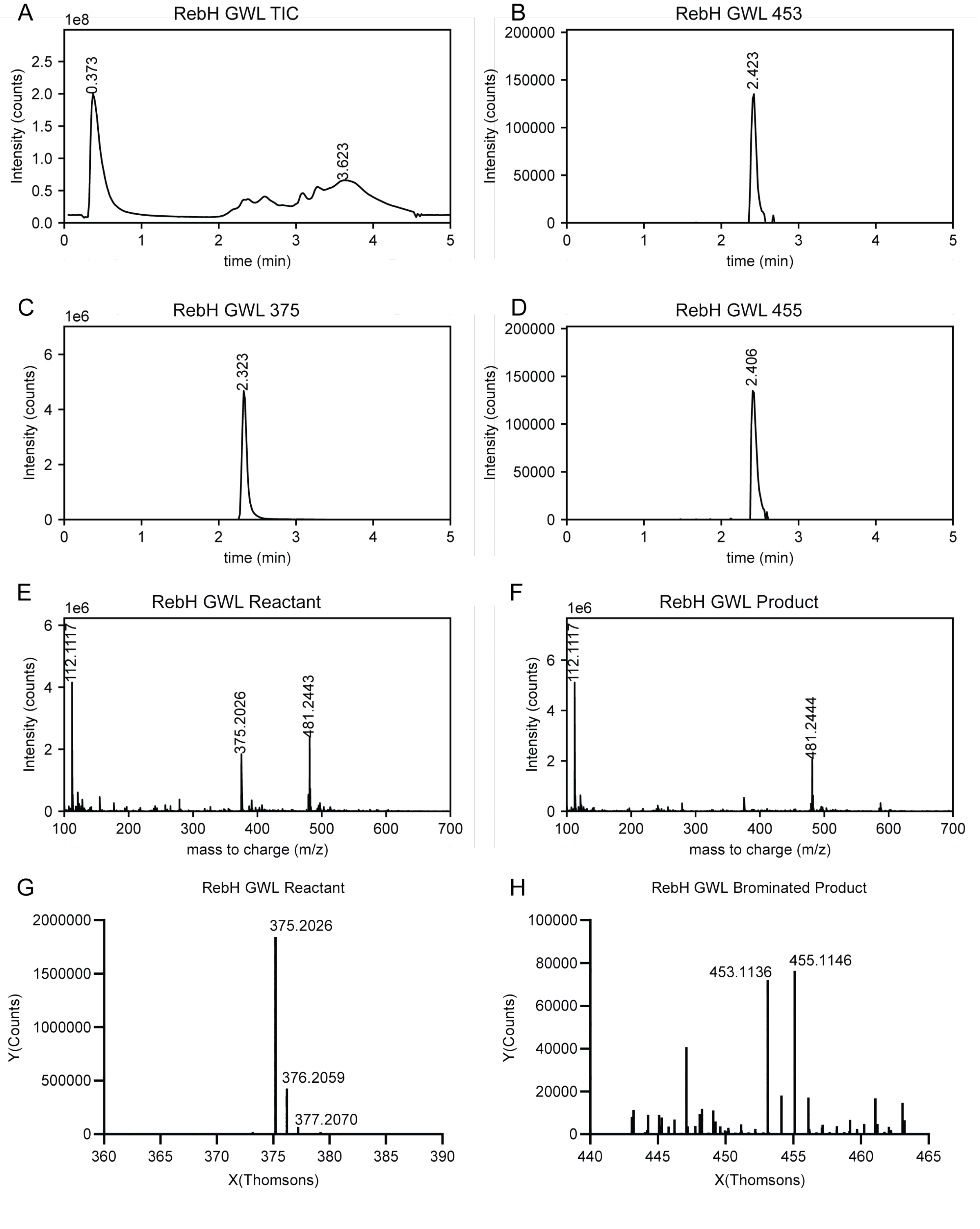

A
RebH GWL TIC
1e8
Intensity (counts)
time (min)
0.373
3.623
B
RebH GWL 453
Intensity (counts)
time (min)
2.423
C
RebH GWL 375
1e6
Intensity (counts)
time (min)
2.323
D
RebH GWL 455
Intensity (counts)
time (min)
2.406
E
RebH GWL Reactant
1e6
Intensity (counts)
mass to charge (m/z)
112.1117
375.2026
481.2443
F
RebH GWL Product
1e6
Intensity (counts)
mass to charge (m/z)
112.1117
481.2444
G
RebH GWL Reactant
Y(Counts)
X(Thomsons)
375.2026
376.2059
377.2070
H
RebH GWL Brominated Product
Y(Counts)
X(Thomsons)
453.1136
455.1146

#### RebH GWM

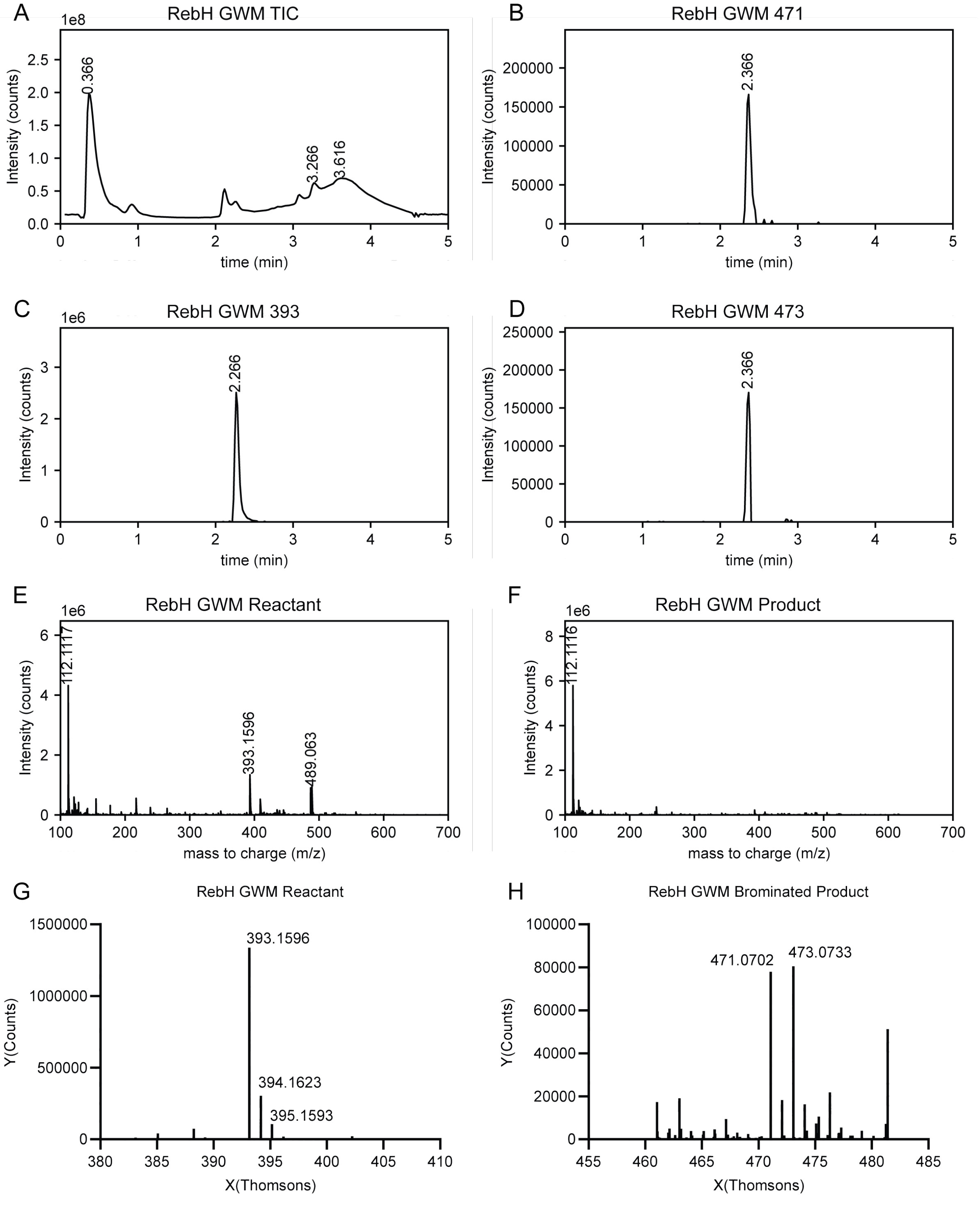
A
RebH GWM TIC
1e8
Intensity (counts)
time (min)
0.366
3.266
3.616
B
RebH GWM 471
Intensity (counts)
time (min)
2.366
C
RebH GWM 393
1e6
Intensity (counts)
time (min)
2.266
D
RebH GWM 473
Intensity (counts)
time (min)
2.366
E
RebH GWM Reactant
1e6
Intensity (counts)
mass to charge (m/z)
112.1117
393.1596
489.063
F
RebH GWM Product
1e6
Intensity (counts)
mass to charge (m/z)
112.1116
G
RebH GWM Reactant
Y(Counts)
X(Thomsons)
393.1596
394.1623
395.1593
H
RebH GWM Brominated Product
Y(Counts)
X(Thomsons)
471.0702
473.0733

#### RebH GWN

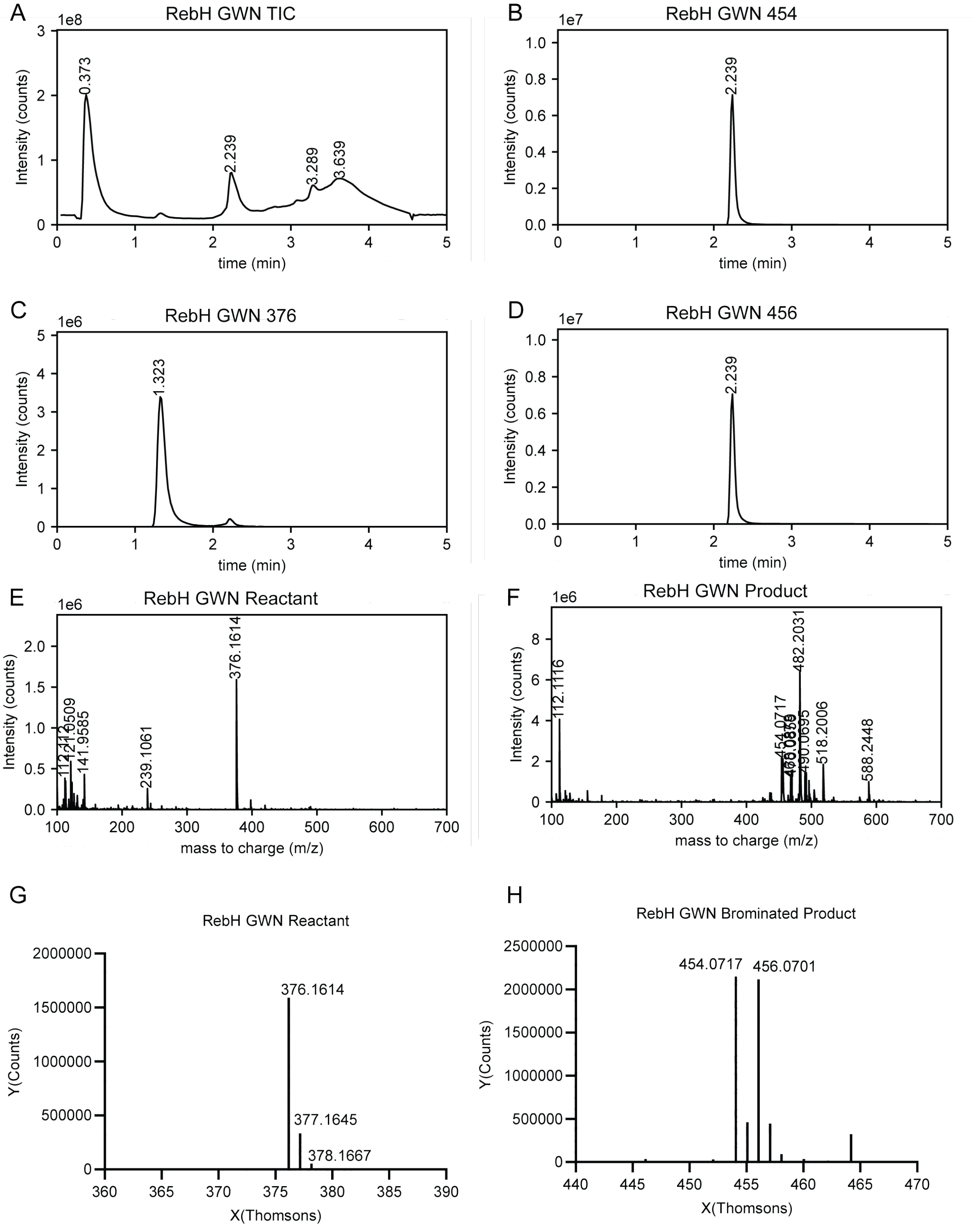
A
RebH GWN TIC
1e8
Intensity (counts)
time (min)
0.373
2.239
3.289
3.639
B
RebH GWN 454
1e7
Intensity (counts)
time (min)
2.239
C
RebH GWN 376
1e6
Intensity (counts)
time (min)
1.323
D
RebH GWN 456
1e7
Intensity (counts)
time (min)
2.239
E
RebH GWN Reactant
1e6
Intensity (counts)
mass to charge (m/z)
121.0509
141.9585
239.1061
376.1614
F
RebH GWN Product
1e6
Intensity (counts)
mass to charge (m/z)
112.1116
454.0717
482.2031
490.0695
518.2006
588.2448
G
RebH GWN Reactant
Y(Counts)
X(Thomsons)
376.1614
377.1645
378.1667
H
RebH GWN Brominated Product
Y(Counts)
X(Thomsons)
454.0717
456.0701

#### RebH GWP

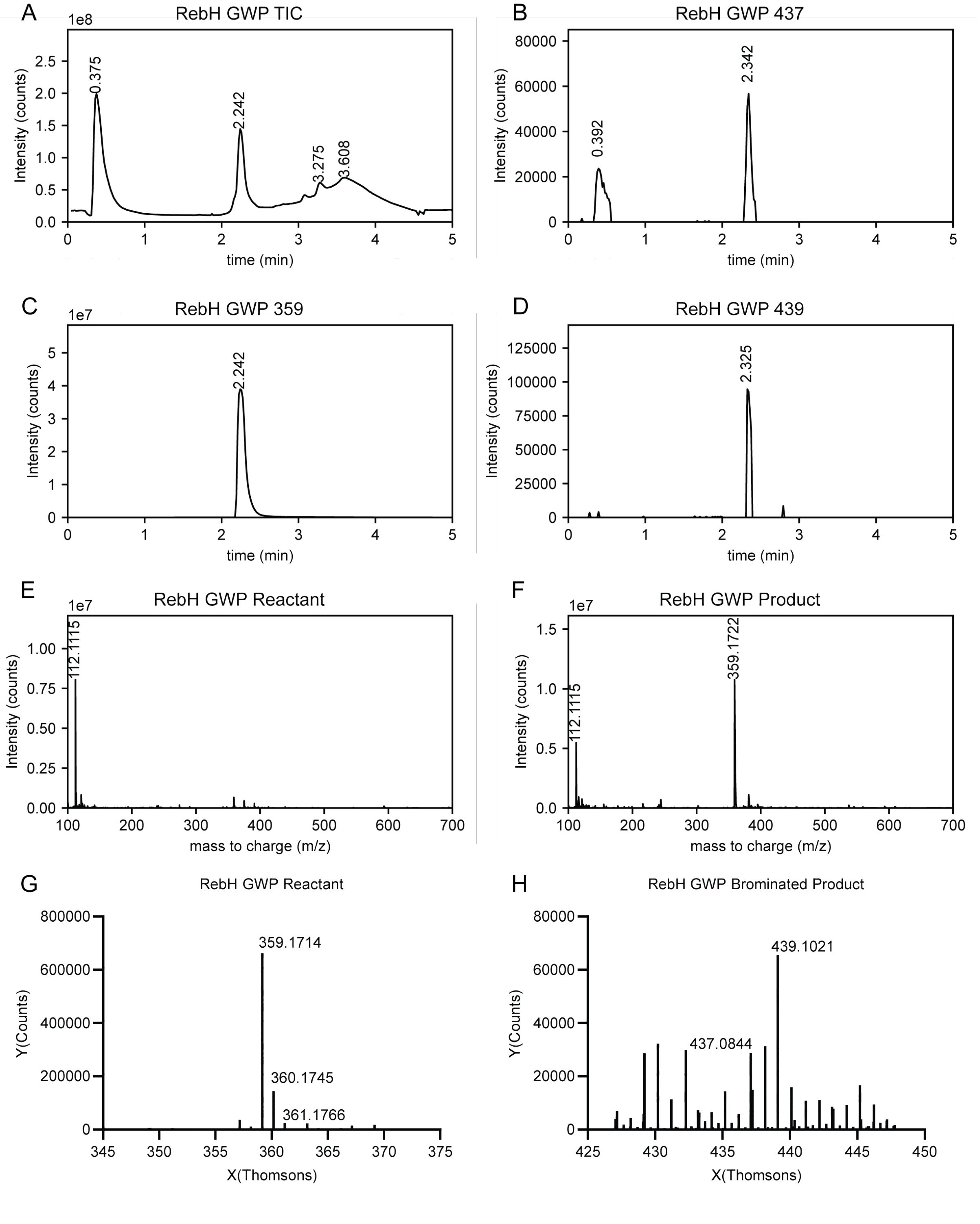
A
RebH GWP TIC
1e8
Intensity (counts)
time (min)
0.375
2.242
3.275
3.608
B
RebH GWP 437
Intensity (counts)
time (min)
0.392
2.342
C
RebH GWP 359
1e7
Intensity (counts)
time (min)
2.242
D
RebH GWP 439
Intensity (counts)
time (min)
2.325
E
RebH GWP Reactant
1e7
Intensity (counts)
mass to charge (m/z)
112.1115
F
RebH GWP Product
1e7
Intensity (counts)
mass to charge (m/z)
112.1115
359.1722
G
RebH GWP Reactant
Y(Counts)
X(Thomsons)
359.1714
360.1745
361.1766
H
RebH GWP Brominated Product
Y(Counts)
X(Thomsons)
439.1021
437.0844

#### RebH GWQ

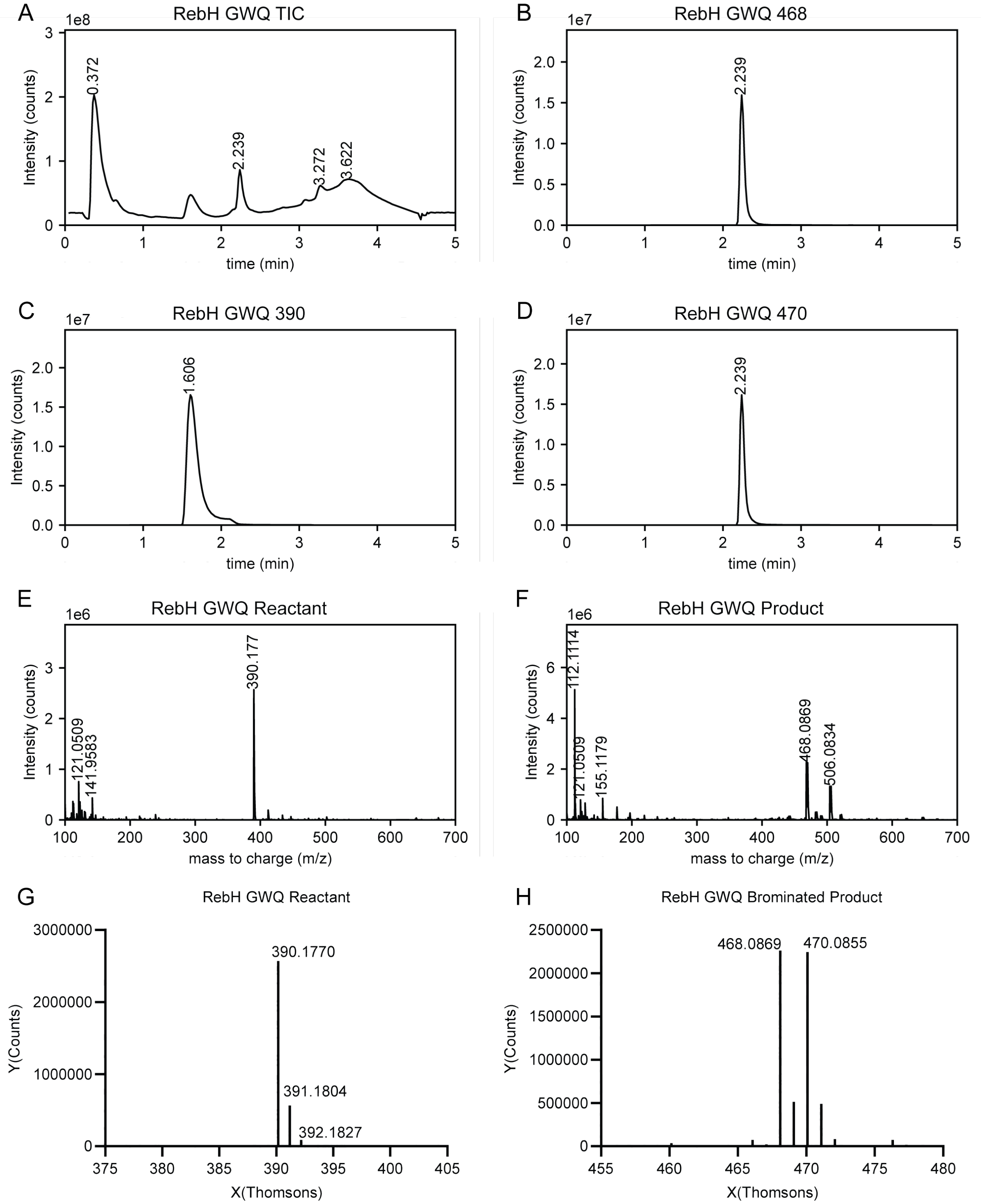

A
RebH GWQ TIC
1e8
Intensity (counts)
time (min)
0.372
2.239
3.272
3.622
B
RebH GWQ 468
1e7
Intensity (counts)
time (min)
2.239
C
RebH GWQ 390
1e7
Intensity (counts)
time (min)
1.606
D
RebH GWQ 470
1e7
Intensity (counts)
time (min)
2.239
E
RebH GWQ Reactant
1e6
Intensity (counts)
mass to charge (m/z)
390.177
121.0509
141.9583
F
RebH GWQ Product
1e6
Intensity (counts)
mass to charge (m/z)
112.1114
121.0509
155.1179
468.0869
506.0834
G
RebH GWQ Reactant
Y(Counts)
X(Thomsons)
390.1770
391.1804
392.1827
H
RebH GWQ Brominated Product
Y(Counts)
X(Thomsons)
468.0869
470.0855

#### RebH GWR

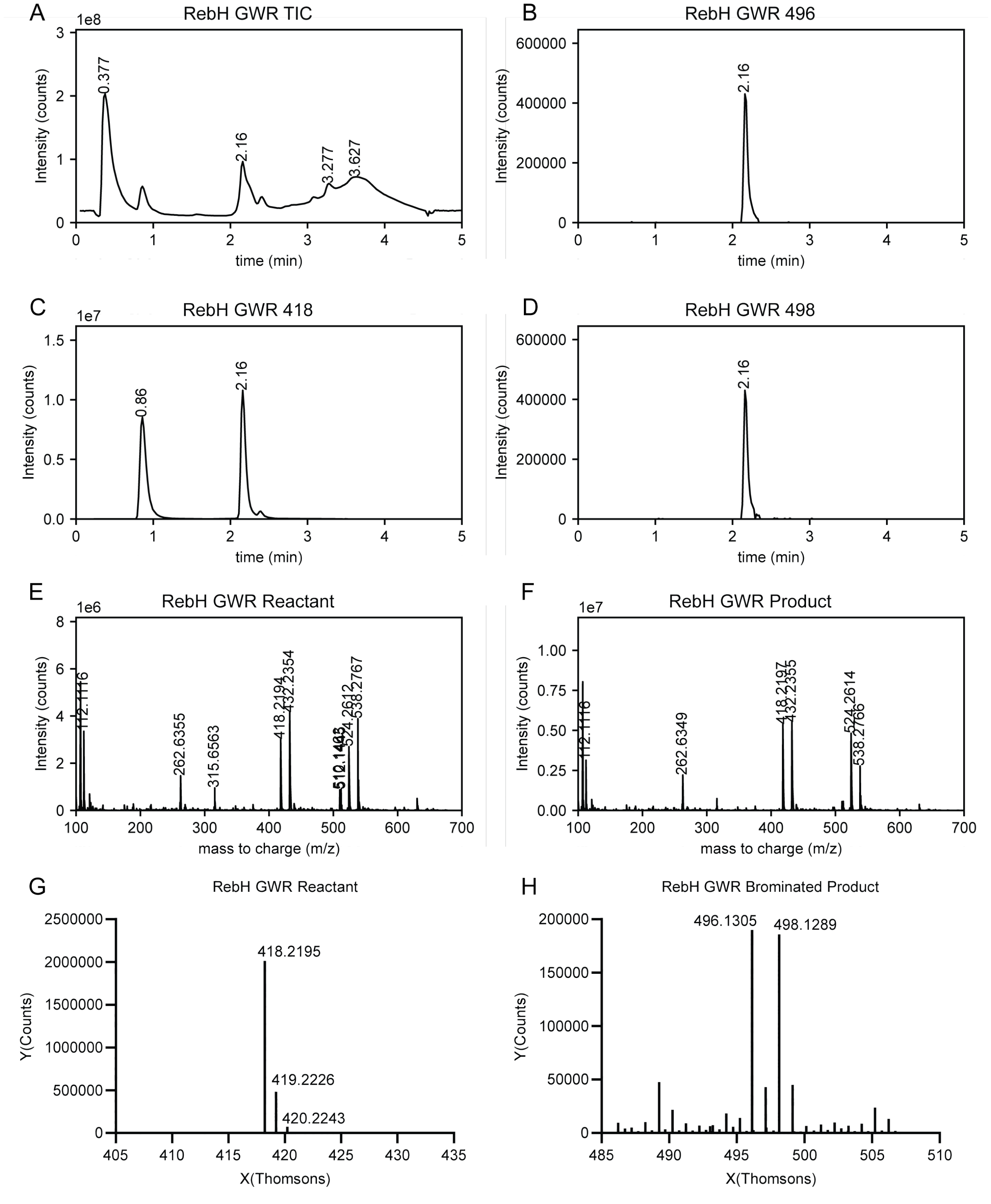

A
RebH GWR TIC
1e8
Intensity (counts)
0.377
2.16
3.277
3.627
time (min)
B
RebH GWR 496
Intensity (counts)
2.16
time (min)
C
RebH GWR 418
1e7
Intensity (counts)
0.86
2.16
time (min)
D
RebH GWR 498
Intensity (counts)
2.16
time (min)
E
RebH GWR Reactant
1e6
Intensity (counts)
112.1116
262.6355
315.6563
418.2194
432.2354
524.2612
538.2767
mass to charge (m/z)
F
RebH GWR Product
1e7
Intensity (counts)
112.1116
262.6349
418.2197
432.2355
524.2614
538.2766
mass to charge (m/z)
G
RebH GWR Reactant
Y(Counts)
418.2195
419.2226
420.2243
X(Thomsons)
H
RebH GWR Brominated Product
Y(Counts)
496.1305
498.1289
X(Thomsons)

#### RebH GWT

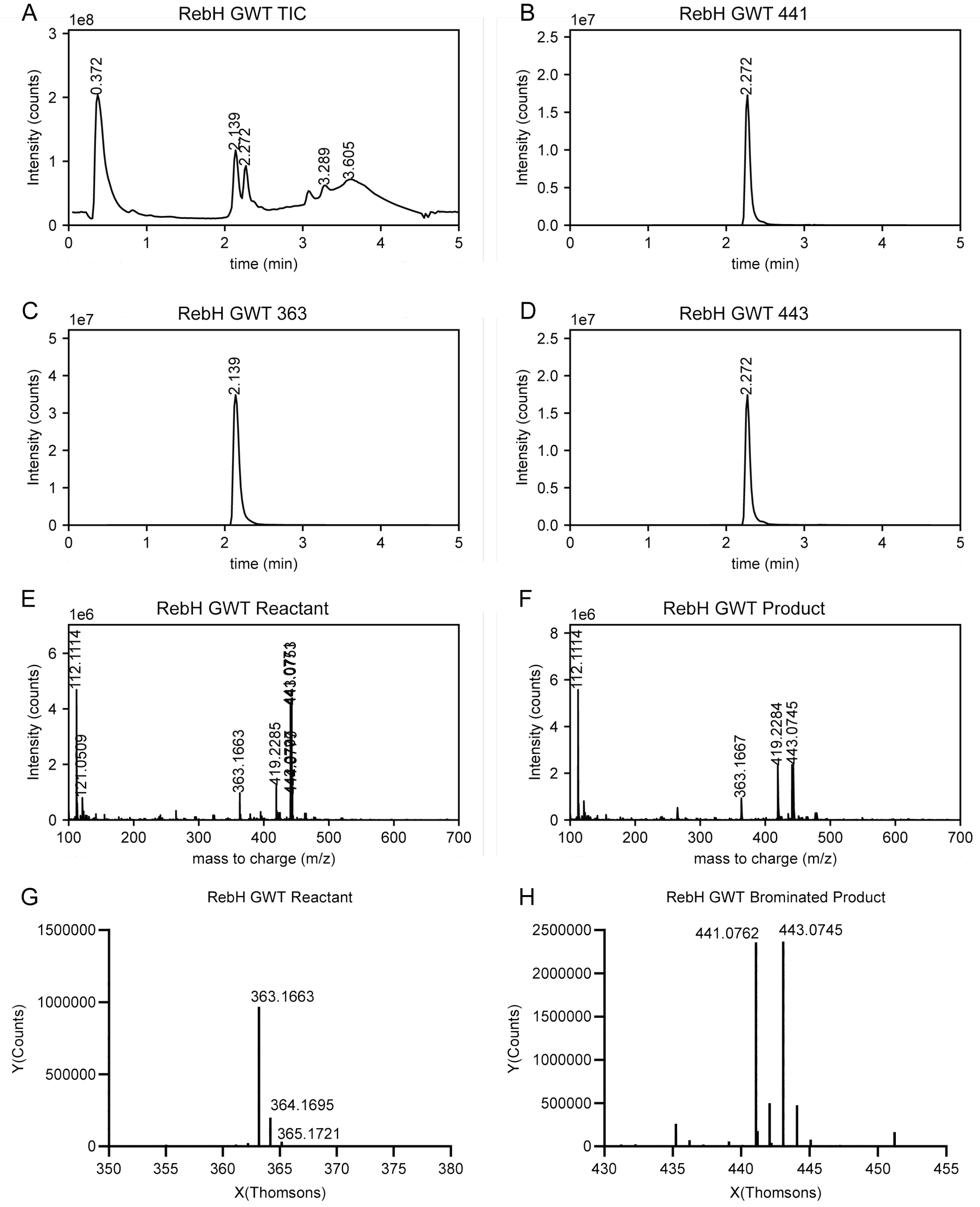
A
RebH GWT TIC
1e8
Intensity (counts)
time (min)
0.372
2.139
2.272
3.289
3.605
B
RebH GWT 441
1e7
Intensity (counts)
time (min)
2.272
C
RebH GWT 363
1e7
Intensity (counts)
time (min)
2.139
D
RebH GWT 443
1e7
Intensity (counts)
time (min)
2.272
E
RebH GWT Reactant
1e6
Intensity (counts)
mass to charge (m/z)
112.1114
121.0509
363.1663
419.2285
F
RebH GWT Product
1e6
Intensity (counts)
mass to charge (m/z)
112.1114
363.1667
419.2284
443.0745
G
RebH GWT Reactant
Y(Counts)
X(Thomsons)
363.1663
364.1695
365.1721
H
RebH GWT Brominated Product
Y(Counts)
X(Thomsons)
441.0762
443.0745

#### RebH GWV

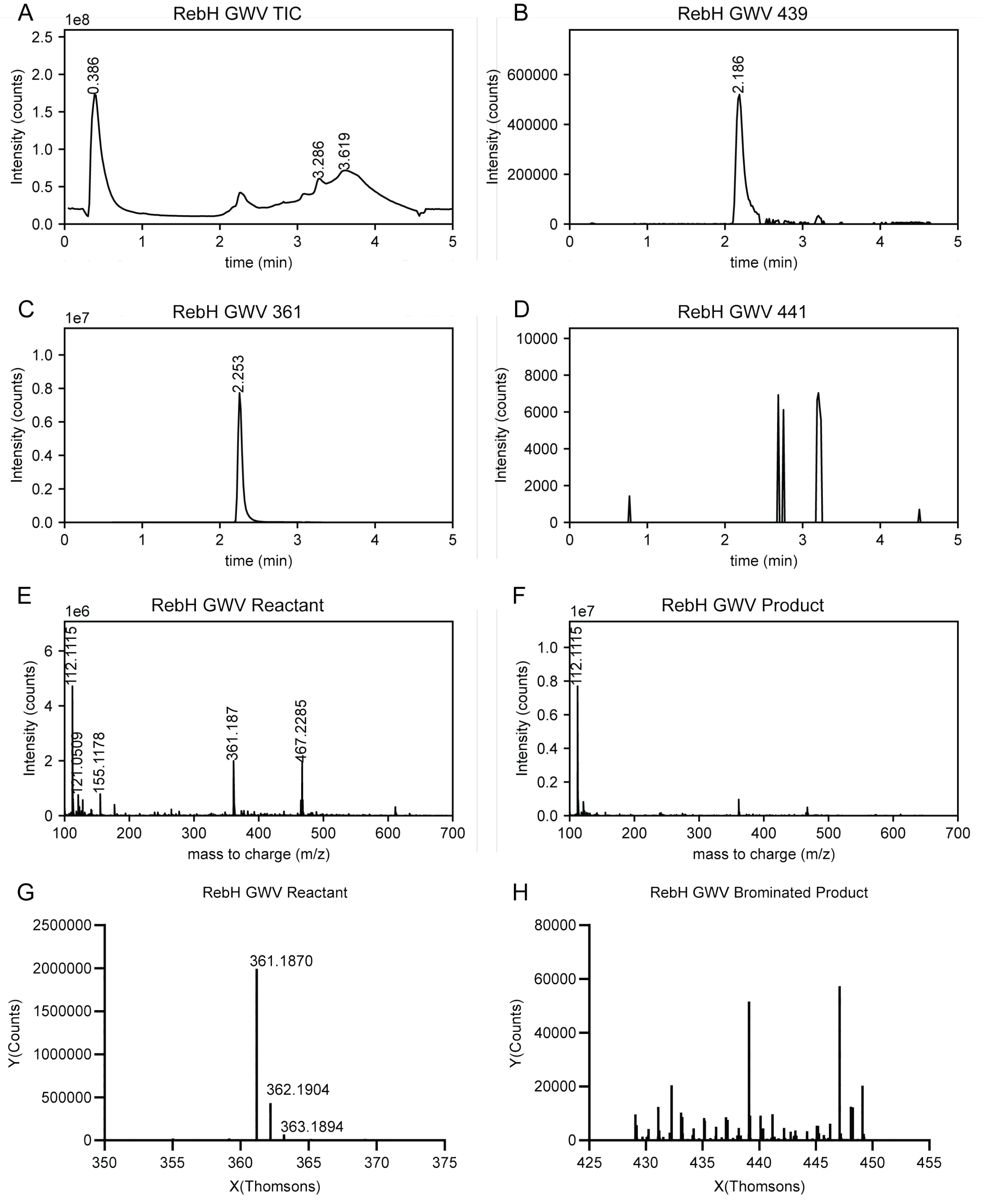

A
RebH GWV TIC
1e8
Intensity (counts)
time (min)
0.386
3.286
3.619
B
RebH GWV 439
2.186
C
RebH GWV 361
1e7
2.253
D
RebH GWV 441
E
RebH GWV Reactant
1e6
mass to charge (m/z)
112.1115
121.0509
155.1178
361.187
467.2285
F
RebH GWV Product
1e7
112.1115
G
RebH GWV Reactant
Y(Counts)
X(Thomsons)
361.1870
362.1904
363.1894
H
RebH GWV Brominated Product

#### RebH GWY

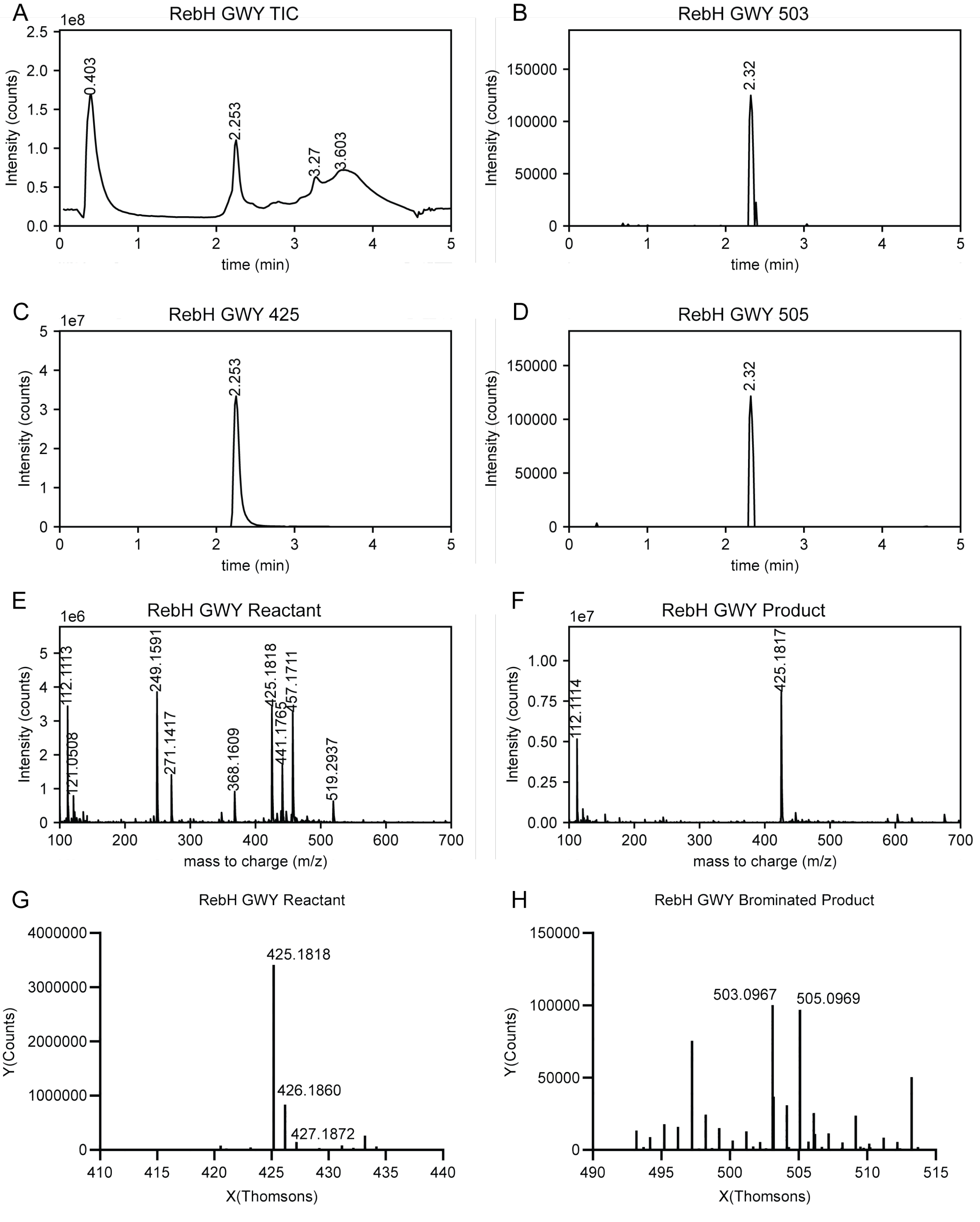

A
RebH GWY TIC
1e8
Intensity (counts)
time (min)
0.403
2.253
3.27
3.603
B
RebH GWY 503
Intensity (counts)
time (min)
2.32
C
RebH GWY 425
1e7
Intensity (counts)
time (min)
2.253
D
RebH GWY 505
Intensity (counts)
time (min)
2.32
E
RebH GWY Reactant
1e6
Intensity (counts)
mass to charge (m/z)
112.1113
121.0508
249.1591
271.1417
368.1609
425.1818
441.1765
457.1711
519.2937
F
RebH GWY Product
1e7
Intensity (counts)
mass to charge (m/z)
112.1114
425.1817
G
RebH GWY Reactant
Y(Counts)
X(Thomsons)
425.1818
426.1860
427.1872
H
RebH GWY Brominated Product
Y(Counts)
X(Thomsons)
503.0967
505.0969

#### RebH AGW

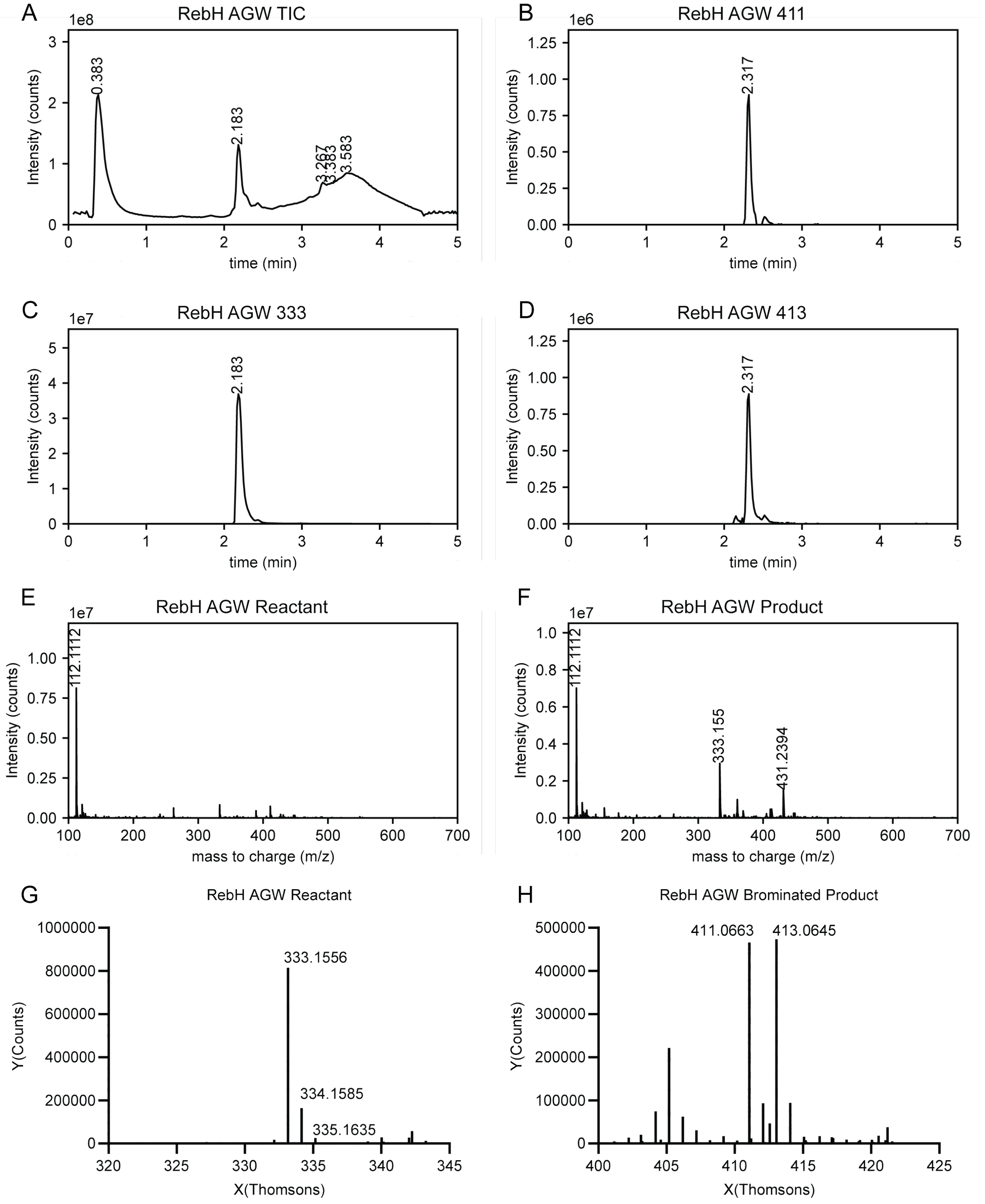

A
RebH AGW TIC
1e8
Intensity (counts)
time (min)
0.383
2.183
3.267
3.383
3.583
B
RebH AGW 411
1e6
Intensity (counts)
time (min)
2.317
C
RebH AGW 333
1e7
Intensity (counts)
time (min)
2.183
D
RebH AGW 413
1e6
Intensity (counts)
time (min)
2.317
E
RebH AGW Reactant
1e7
Intensity (counts)
mass to charge (m/z)
112.1112
F
RebH AGW Product
1e7
Intensity (counts)
mass to charge (m/z)
112.1112
333.155
431.2394
G
RebH AGW Reactant
Y(Counts)
X(Thomsons)
333.1556
334.1585
335.1635
H
RebH AGW Brominated Product
Y(Counts)
X(Thomsons)
411.0663
413.0645

#### RebH CGW

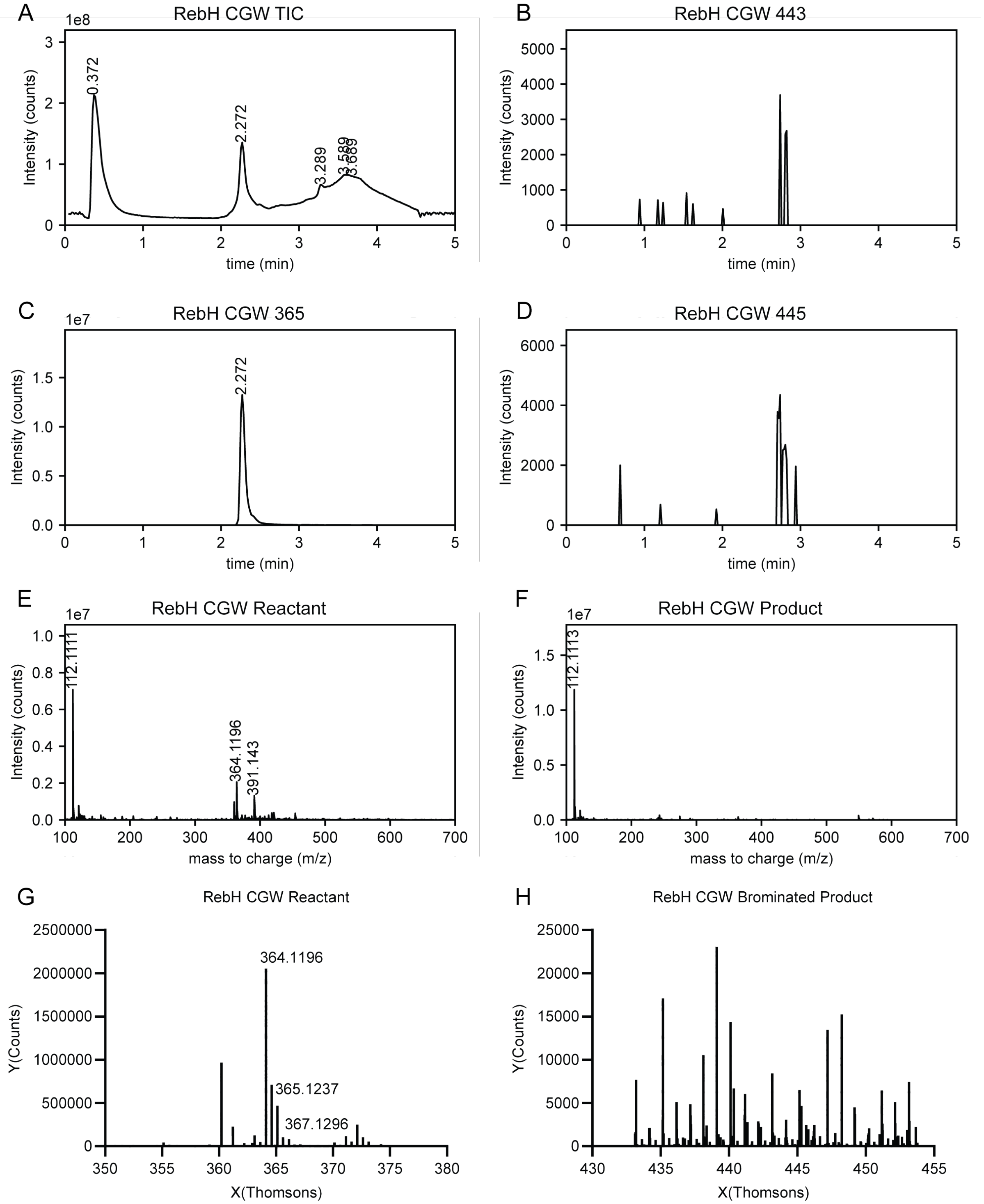

A
RebH CGW TIC
1e8
Intensity (counts)
time (min)
0.372
2.272
3.289
3.589
3.689
B
RebH CGW 443
Intensity (counts)
time (min)
C
RebH CGW 365
1e7
Intensity (counts)
time (min)
2.272
D
RebH CGW 445
Intensity (counts)
time (min)
E
RebH CGW Reactant
1e7
Intensity (counts)
mass to charge (m/z)
112.1111
364.1196
391.143
F
RebH CGW Product
1e7
Intensity (counts)
mass to charge (m/z)
112.1113
G
RebH CGW Reactant
Y(Counts)
X(Thomsons)
364.1196
365.1237
367.1296
H
RebH CGW Brominated Product
Y(Counts)
X(Thomsons)

#### RebH DGW

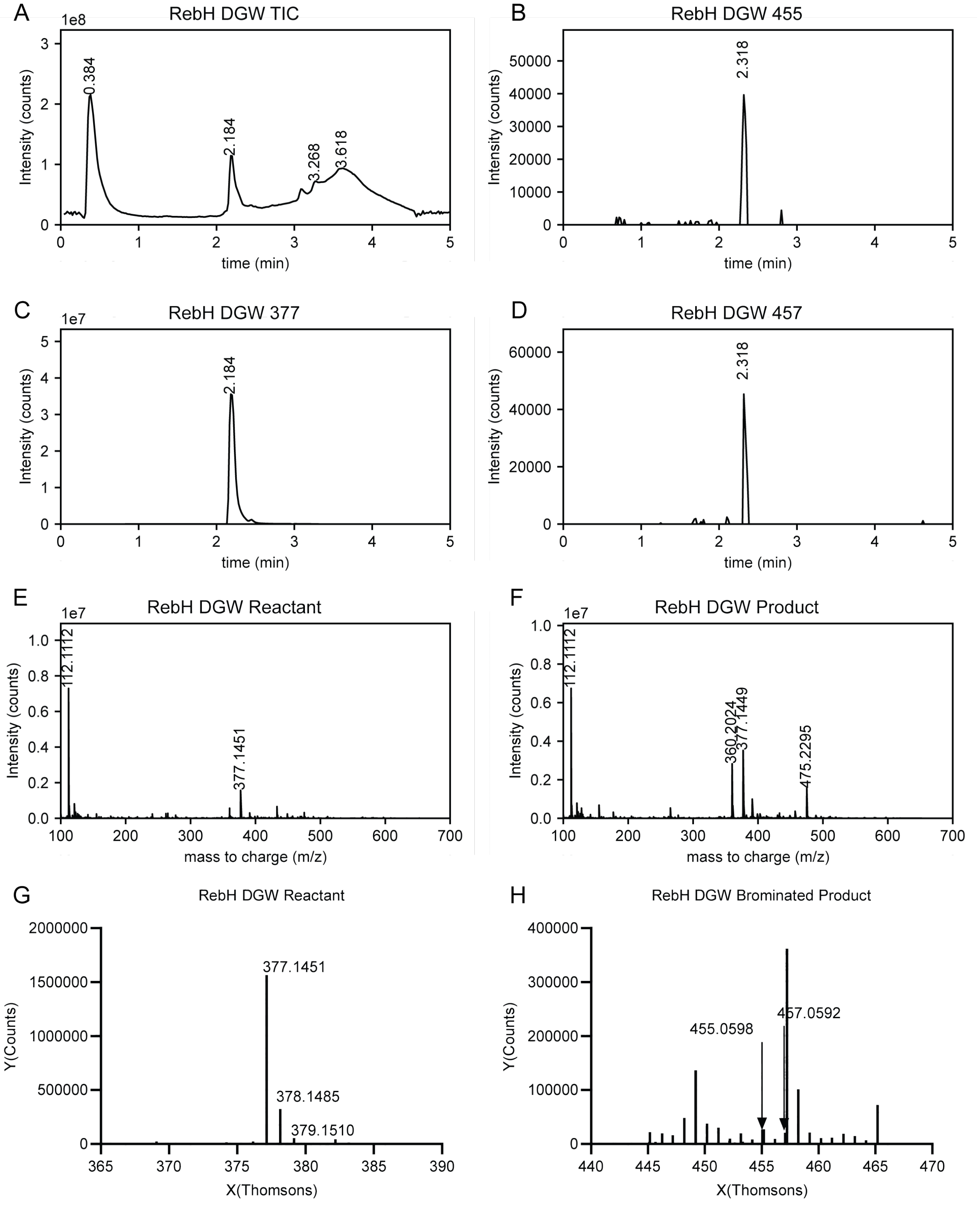
A
RebH DGW TIC
1e8
Intensity (counts)
time (min)
0.384
2.184
3.268
3.618
B
RebH DGW 455
Intensity (counts)
time (min)
2.318
C
RebH DGW 377
1e7
Intensity (counts)
time (min)
2.184
D
RebH DGW 457
Intensity (counts)
time (min)
2.318
E
RebH DGW Reactant
1e7
Intensity (counts)
mass to charge (m/z)
112.1112
377.1451
F
RebH DGW Product
1e7
Intensity (counts)
mass to charge (m/z)
112.1112
360.2024
377.1449
475.2295
G
RebH DGW Reactant
Y(Counts)
X(Thomsons)
377.1451
378.1485
379.1510
H
RebH DGW Brominated Product
Y(Counts)
X(Thomsons)
455.0598
457.0592

#### RebH EGW

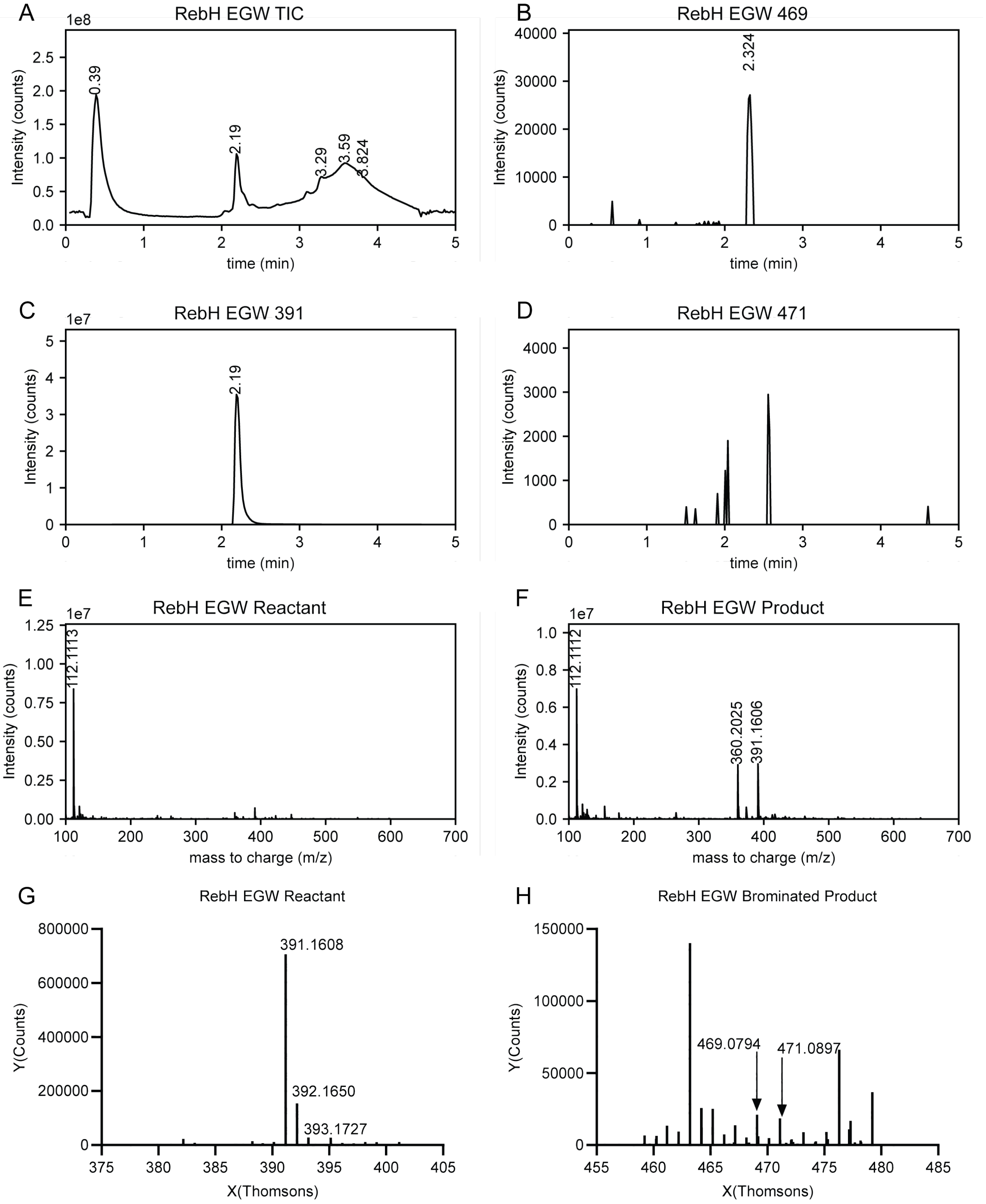
A
RebH EGW TIC
1e8
Intensity (counts)
time (min)
0.39
2.19
3.29
3.59
3.824
B
RebH EGW 469
Intensity (counts)
time (min)
2.324
C
RebH EGW 391
1e7
Intensity (counts)
time (min)
2.19
D
RebH EGW 471
Intensity (counts)
time (min)
E
RebH EGW Reactant
1e7
Intensity (counts)
mass to charge (m/z)
112.1113
F
RebH EGW Product
1e7
Intensity (counts)
mass to charge (m/z)
112.1112
360.2025
391.1606
G
RebH EGW Reactant
Y(Counts)
X(Thomsons)
391.1608
392.1650
393.1727
H
RebH EGW Brominated Product
Y(Counts)
X(Thomsons)
469.0794
471.0897

#### RebH FGW

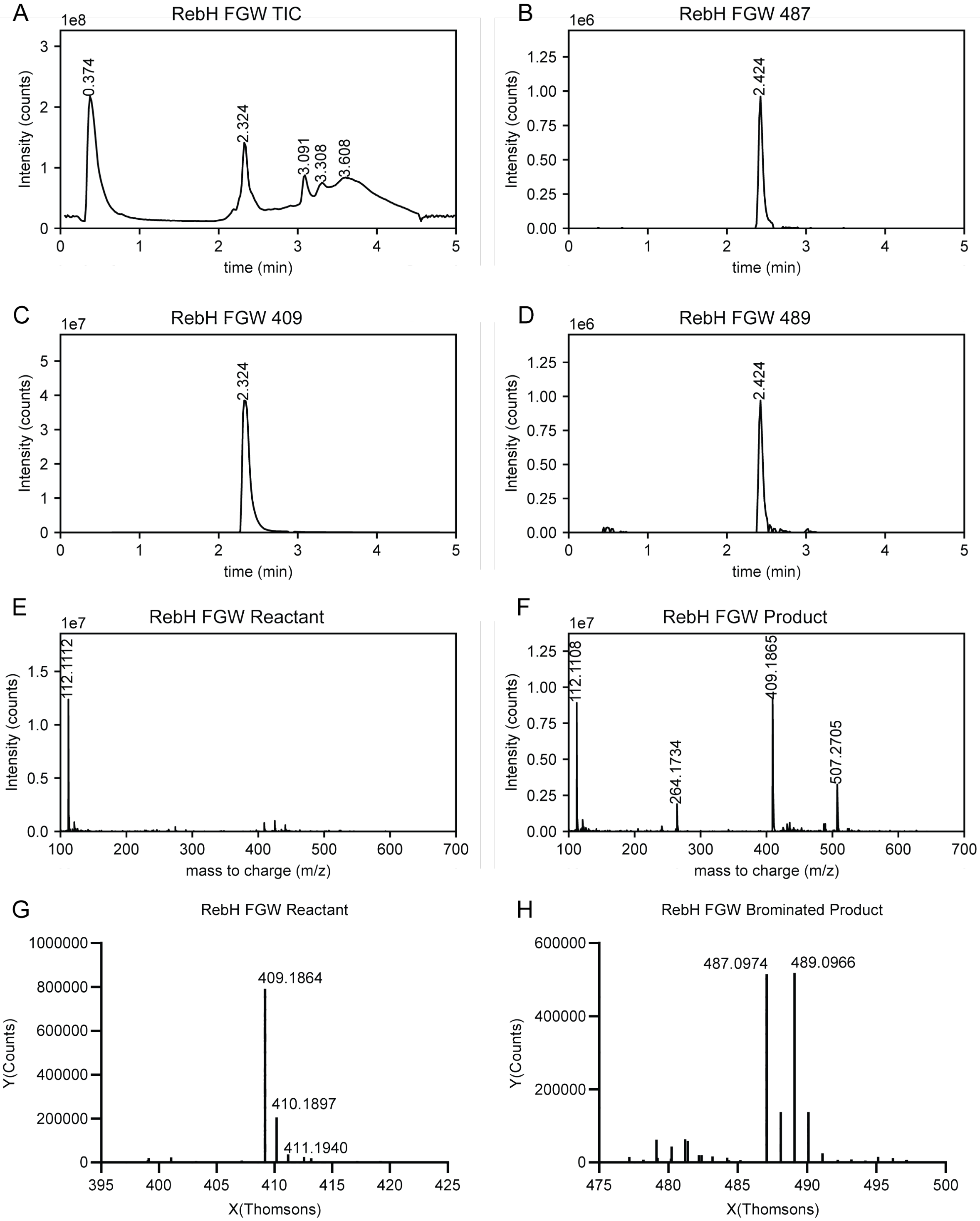

A
RebH FGW TIC
1e8
0.374
2.324
3.091
3.308
3.608
Intensity (counts)
time (min)
B
RebH FGW 487
1e6
2.424
Intensity (counts)
time (min)
C
RebH FGW 409
1e7
2.324
Intensity (counts)
time (min)
D
RebH FGW 489
1e6
2.424
Intensity (counts)
time (min)
E
RebH FGW Reactant
1e7
112.1112
Intensity (counts)
mass to charge (m/z)
F
RebH FGW Product
1e7
112.1108
409.1865
264.1734
507.2705
Intensity (counts)
mass to charge (m/z)
G
RebH FGW Reactant
409.1864
410.1897
411.1940
Y(Counts)
X(Thomsons)
H
RebH FGW Brominated Product
487.0974
489.0966
Y(Counts)
X(Thomsons)

#### RebH HGW

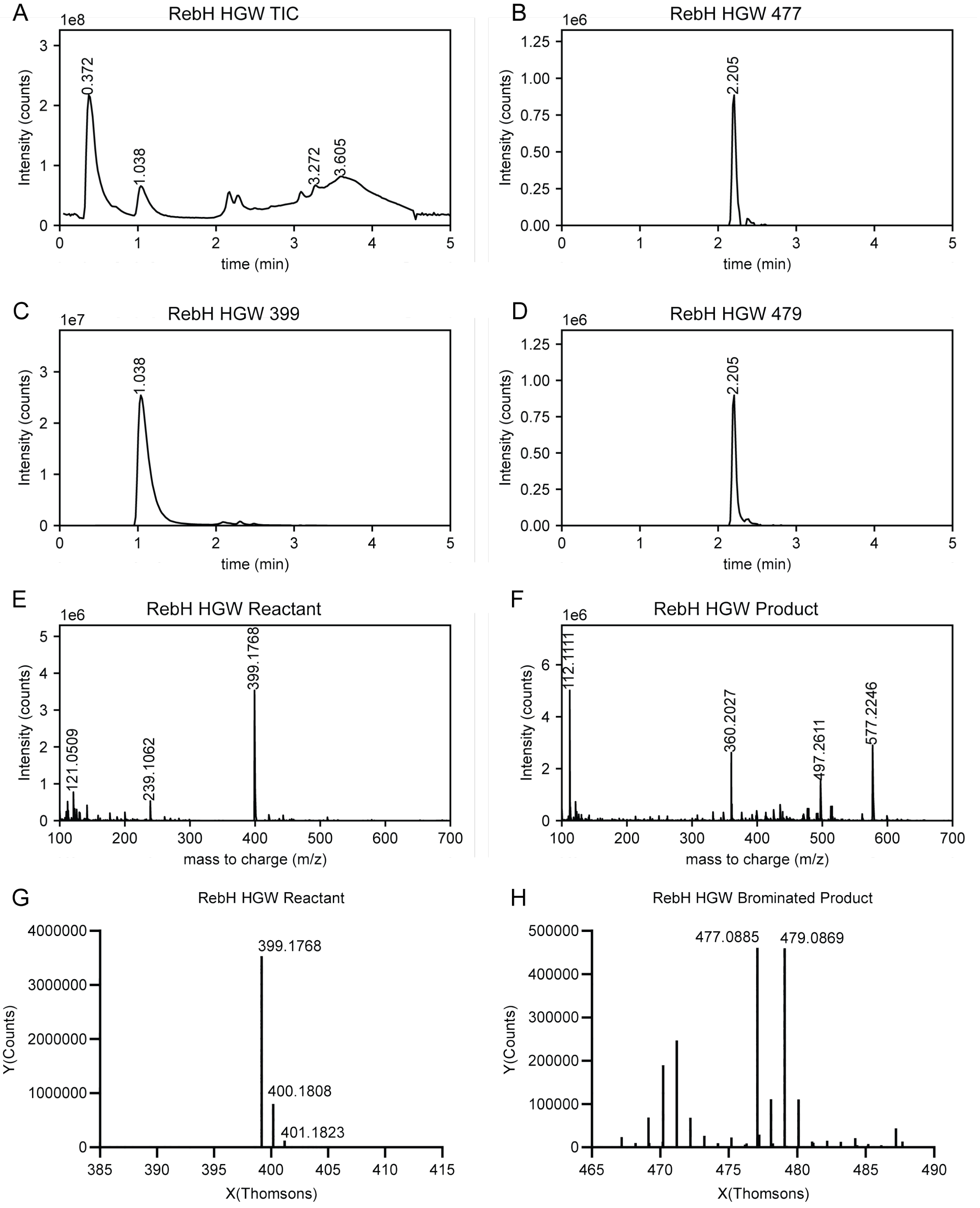

A
RebH HGW TIC
1e8
Intensity (counts)
time (min)
0.372
1.038
3.272
3.605
B
RebH HGW 477
1e6
Intensity (counts)
time (min)
2.205
C
RebH HGW 399
1e7
Intensity (counts)
time (min)
1.038
D
RebH HGW 479
1e6
Intensity (counts)
time (min)
2.205
E
RebH HGW Reactant
1e6
Intensity (counts)
mass to charge (m/z)
121.0509
239.1062
399.1768
F
RebH HGW Product
1e6
Intensity (counts)
mass to charge (m/z)
112.1111
360.2027
497.2611
577.2246
G
RebH HGW Reactant
Y(Counts)
X(Thomsons)
399.1768
400.1808
401.1823
H
RebH HGW Brominated Product
Y(Counts)
X(Thomsons)
477.0885
479.0869

#### RebH KGW

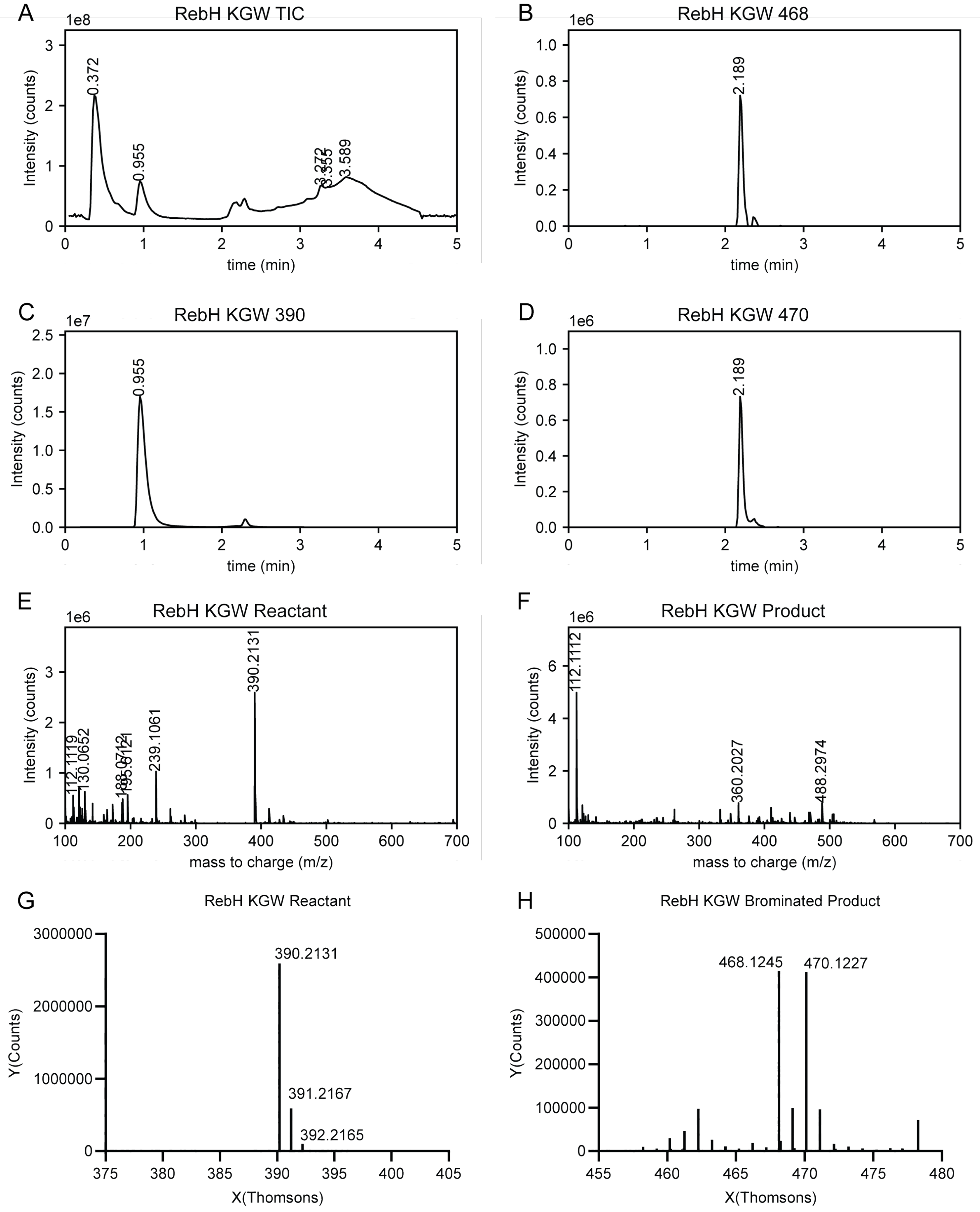
A
RebH KGW TIC
B
RebH KGW 468
C
RebH KGW 390
D
RebH KGW 470
E
RebH KGW Reactant
F
RebH KGW Product
G
RebH KGW Reactant
H
RebH KGW Brominated Product
Intensity (counts)
time (min)
mass to charge (m/z)
Y(Counts)
X(Thomsons)
0.372
0.955
3.589
2.189
390.2131
239.1061
130.0652
112.1119
112.1112
360.2027
488.2974
391.2167
392.2165
468.1245
470.1227

#### RebH LGW

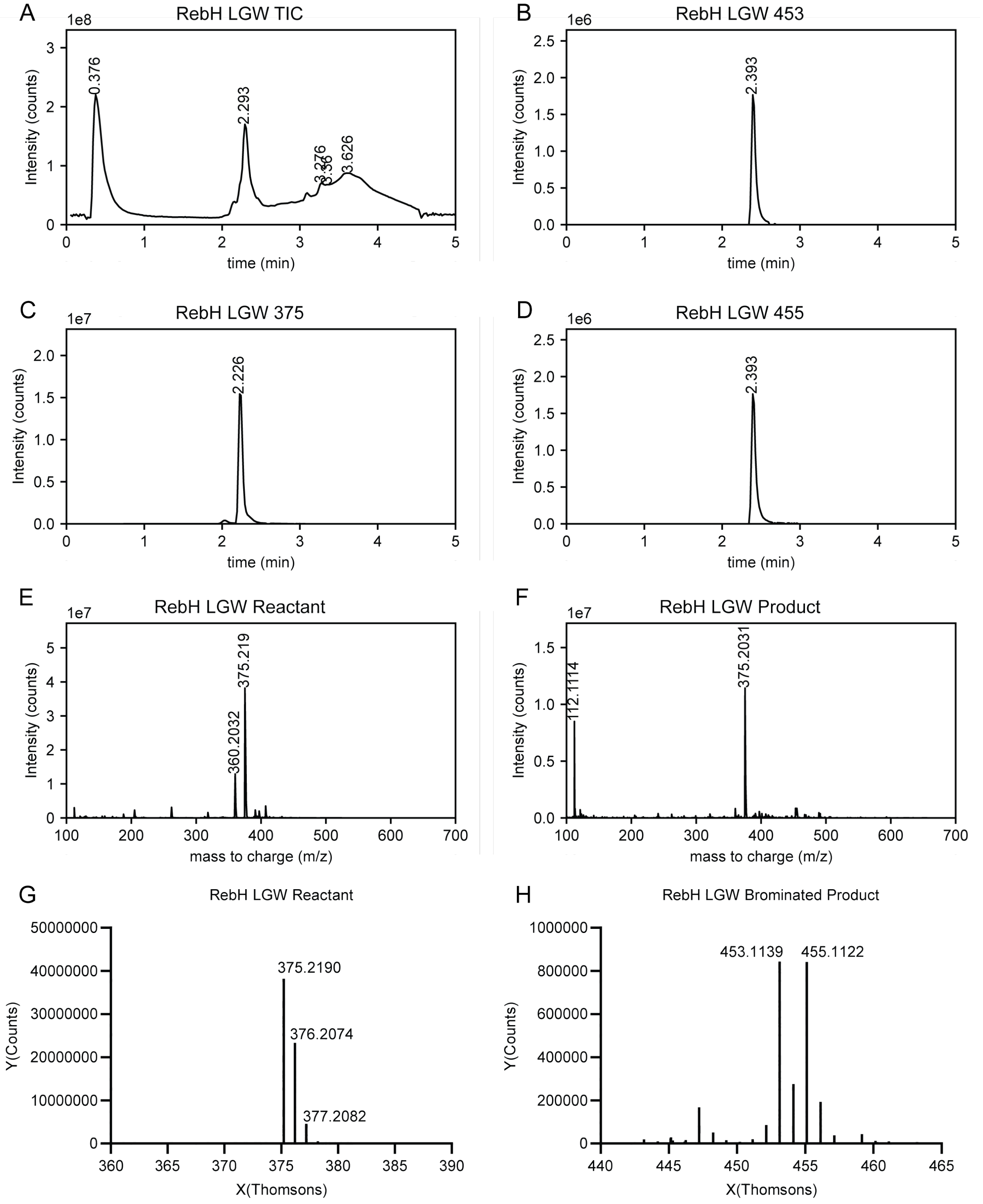

A
RebH LGW TIC
1e8
Intensity (counts)
time (min)
0.376
2.293
3.276
3.36
3.626
B
RebH LGW 453
1e6
Intensity (counts)
time (min)
2.393
C
RebH LGW 375
1e7
Intensity (counts)
time (min)
2.226
D
RebH LGW 455
1e6
Intensity (counts)
time (min)
2.393
E
RebH LGW Reactant
1e7
Intensity (counts)
mass to charge (m/z)
360.2032
375.219
F
RebH LGW Product
1e7
Intensity (counts)
mass to charge (m/z)
112.1114
375.2031
G
RebH LGW Reactant
Y(Counts)
X(Thomsons)
375.2190
376.2074
377.2082
H
RebH LGW Brominated Product
Y(Counts)
X(Thomsons)
453.1139
455.1122

#### RebH MGW

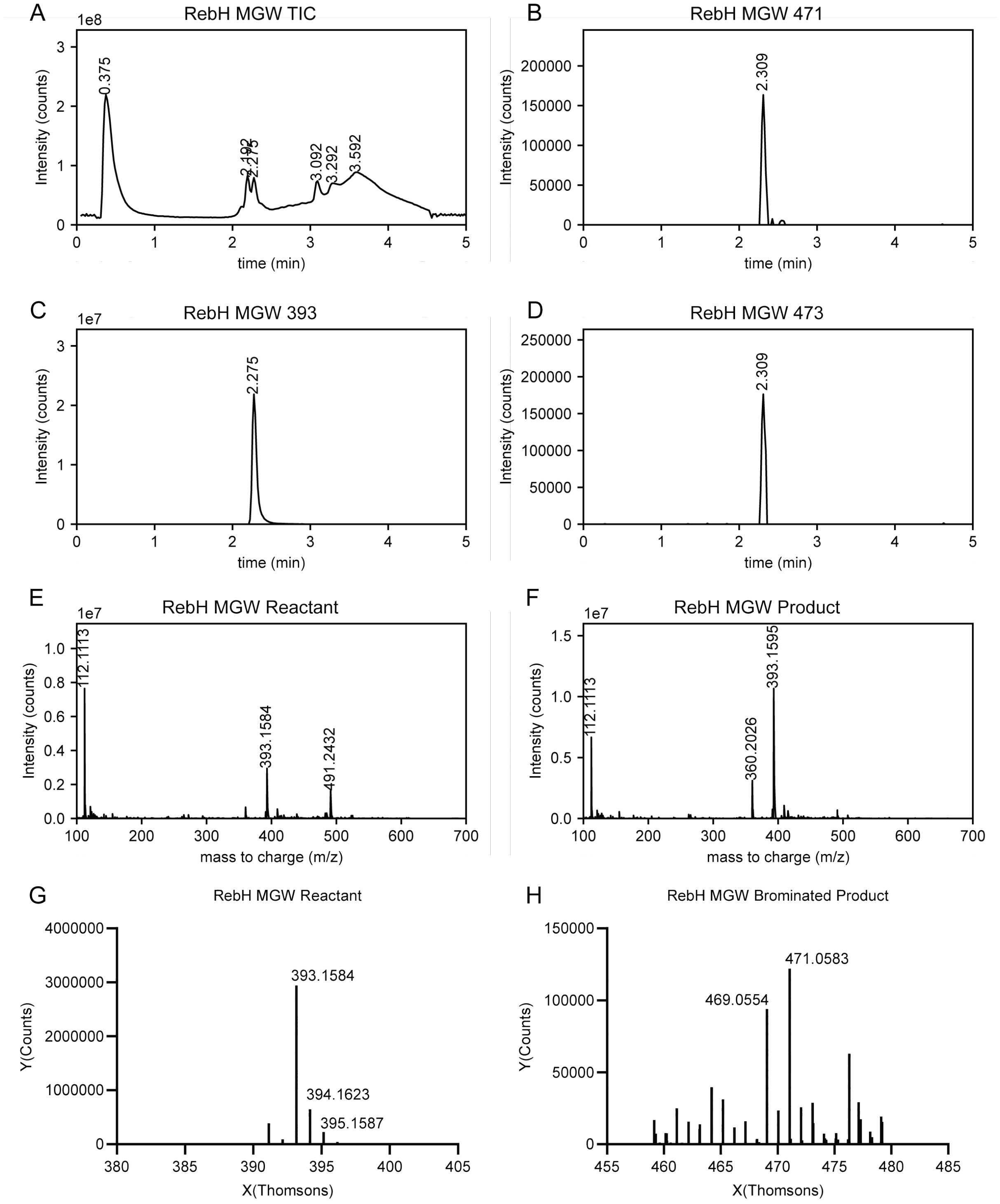

A
RebH MGW TIC
1e8
Intensity (counts)
time (min)
0.375
2.192
2.275
3.092
3.292
3.592
B
RebH MGW 471
Intensity (counts)
time (min)
2.309
C
RebH MGW 393
1e7
Intensity (counts)
time (min)
2.275
D
RebH MGW 473
Intensity (counts)
time (min)
2.309
E
RebH MGW Reactant
1e7
Intensity (counts)
mass to charge (m/z)
112.1113
393.1584
491.2432
F
RebH MGW Product
1e7
Intensity (counts)
mass to charge (m/z)
112.1113
360.2026
393.1595
G
RebH MGW Reactant
Y(Counts)
X(Thomsons)
393.1584
394.1623
395.1587
H
RebH MGW Brominated Product
Y(Counts)
X(Thomsons)
471.0583
469.0554

#### RebH NGW

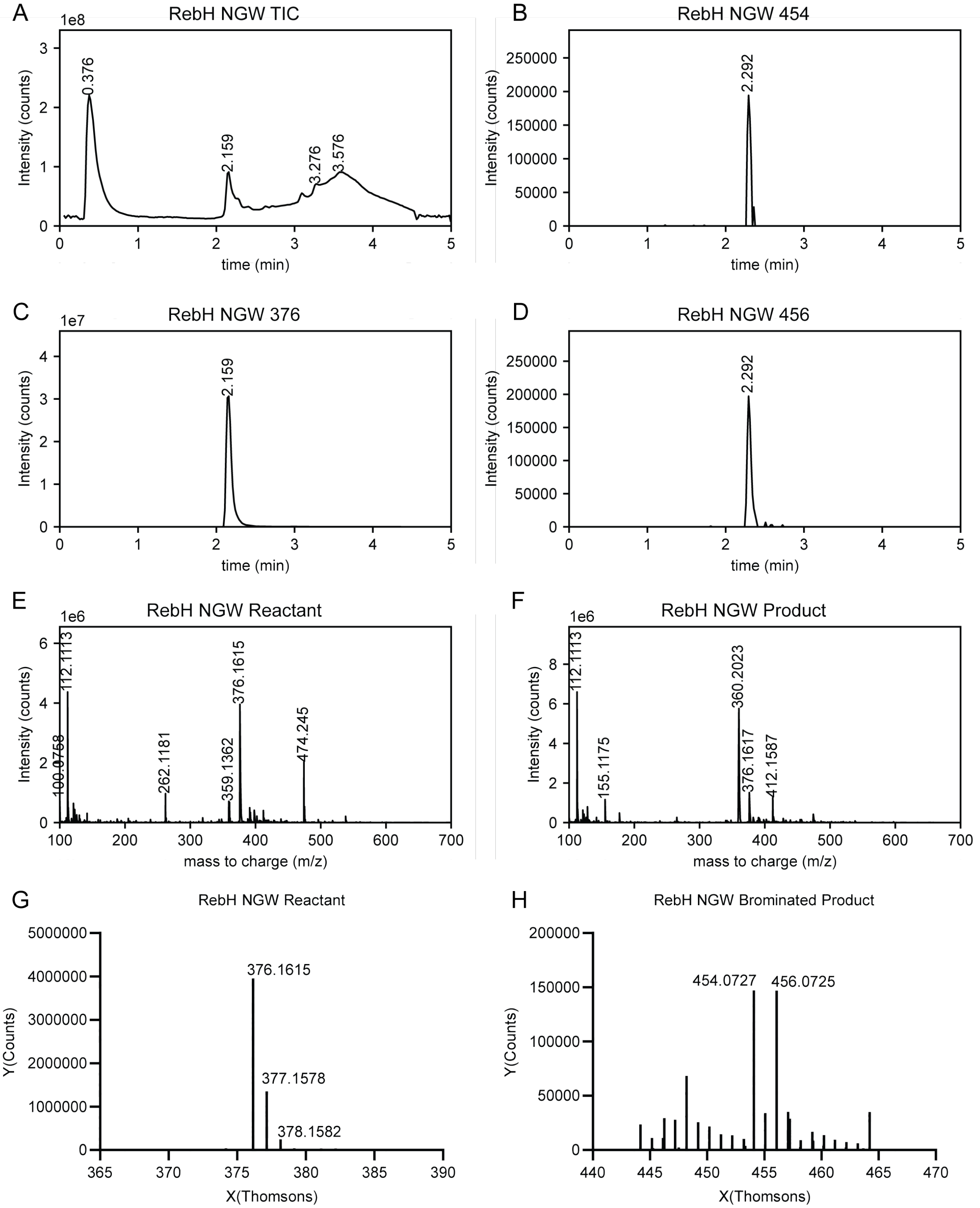
A
RebH NGW TIC
1e8
Intensity (counts)
time (min)
0.376
2.159
3.276
3.576
B
RebH NGW 454
Intensity (counts)
time (min)
2.292
C
RebH NGW 376
1e7
Intensity (counts)
time (min)
2.159
D
RebH NGW 456
Intensity (counts)
time (min)
2.292
E
RebH NGW Reactant
1e6
Intensity (counts)
mass to charge (m/z)
100.0758
112.1113
262.1181
359.1362
376.1615
474.245
F
RebH NGW Product
1e6
Intensity (counts)
mass to charge (m/z)
112.1113
155.1175
360.2023
376.1617
412.1587
G
RebH NGW Reactant
Y(Counts)
X(Thomsons)
376.1615
377.1578
378.1582
H
RebH NGW Brominated Product
Y(Counts)
X(Thomsons)
454.0727
456.0725

**RebH PGW**

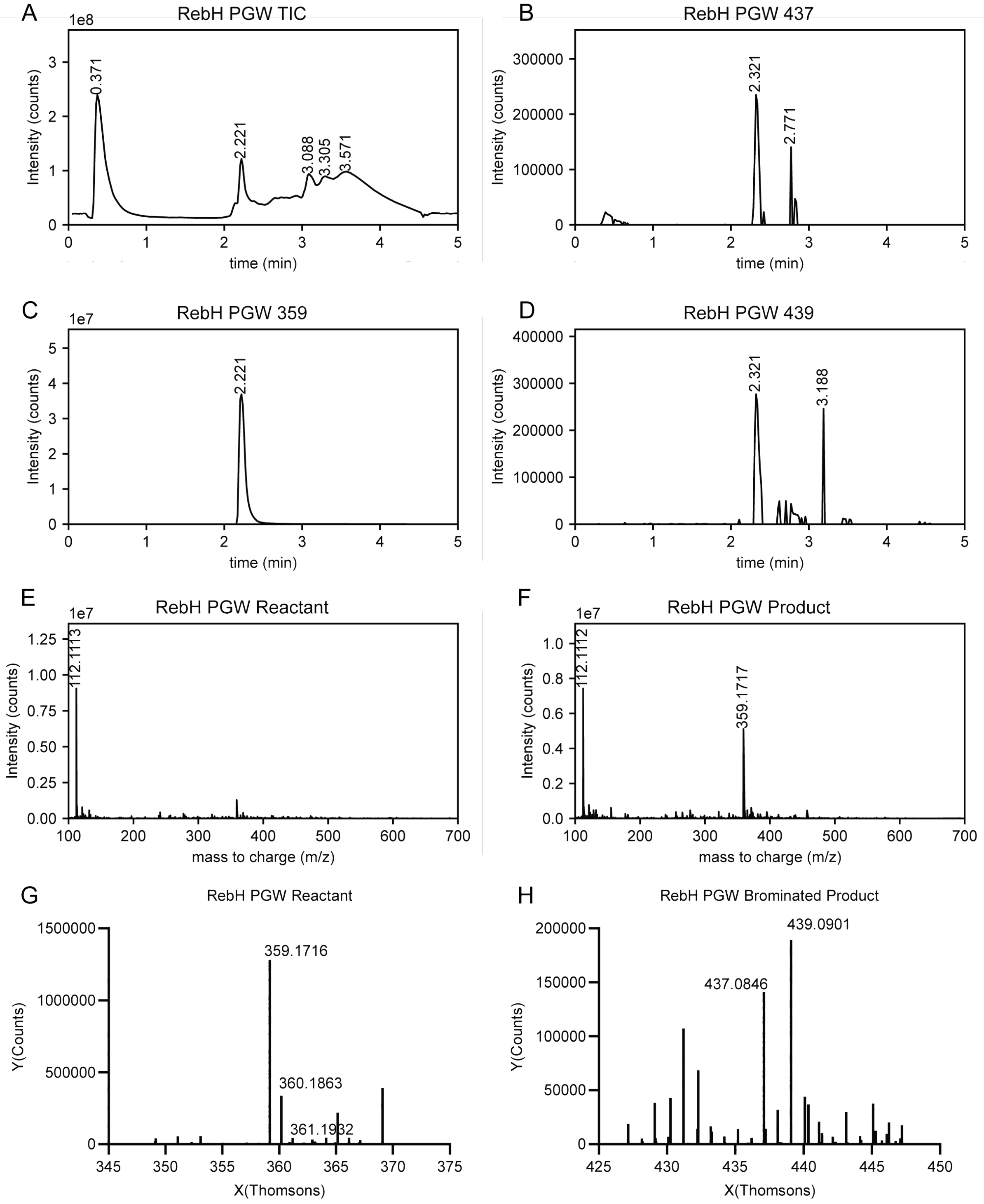
A
RebH PGW TIC
1e8
Intensity (counts)
time (min)
0.371
2.221
3.088
3.305
3.571
B
RebH PGW 437
2.321
2.771
C
RebH PGW 359
1e7
2.221
D
RebH PGW 439
2.321
3.188
E
RebH PGW Reactant
112.1113
mass to charge (m/z)
F
RebH PGW Product
112.1112
359.1717
G
RebH PGW Reactant
359.1716
360.1863
361.1932
Y(Counts)
X(Thomsons)
H
RebH PGW Brominated Product
439.0901
437.0846

#### RebH QGW

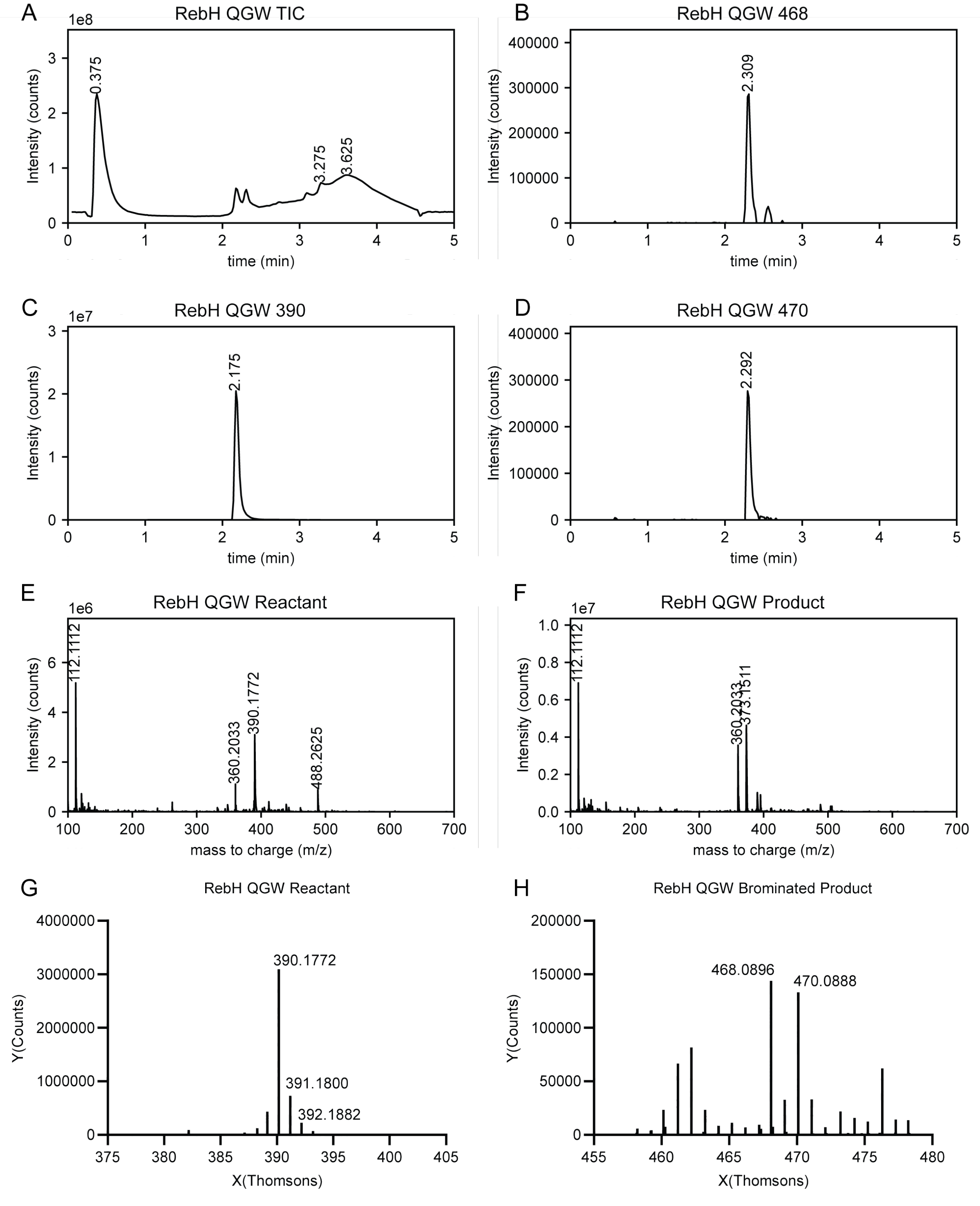
A
RebH QGW TIC
1e8
Intensity (counts)
time (min)
0.375
3.275
3.625
B
RebH QGW 468
Intensity (counts)
time (min)
2.309
C
RebH QGW 390
1e7
Intensity (counts)
time (min)
2.175
D
RebH QGW 470
Intensity (counts)
time (min)
2.292
E
RebH QGW Reactant
1e6
Intensity (counts)
mass to charge (m/z)
112.1112
360.2033
390.1772
488.2625
F
RebH QGW Product
1e7
Intensity (counts)
mass to charge (m/z)
112.1112
360.2033
373.1511
G
RebH QGW Reactant
Y(Counts)
X(Thomsons)
390.1772
391.1800
392.1882
H
RebH QGW Brominated Product
Y(Counts)
X(Thomsons)
468.0896
470.0888

#### RebH RGW

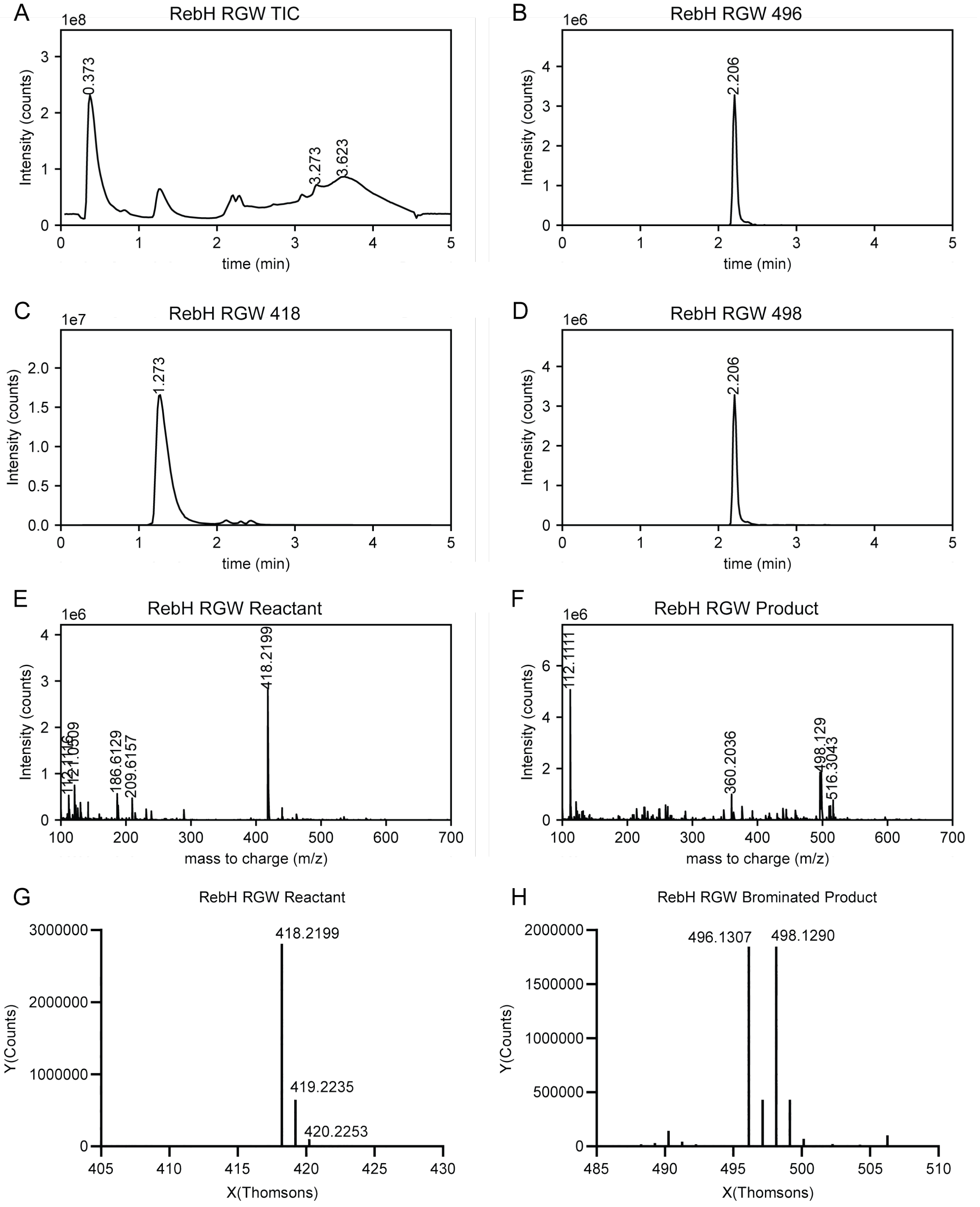

A
RebH RGW TIC
1e8
Intensity (counts)
time (min)
0.373
3.273
3.623
B
RebH RGW 496
1e6
Intensity (counts)
time (min)
2.206
C
RebH RGW 418
1e7
Intensity (counts)
time (min)
1.273
D
RebH RGW 498
1e6
Intensity (counts)
time (min)
2.206
E
RebH RGW Reactant
1e6
Intensity (counts)
mass to charge (m/z)
121.0509
186.6129
209.6157
418.2199
F
RebH RGW Product
1e6
Intensity (counts)
mass to charge (m/z)
112.1111
360.2036
498.129
516.3043
G
RebH RGW Reactant
Y(Counts)
X(Thomsons)
418.2199
419.2235
420.2253
H
RebH RGW Brominated Product
Y(Counts)
X(Thomsons)
496.1307
498.1290

#### RebH TGW

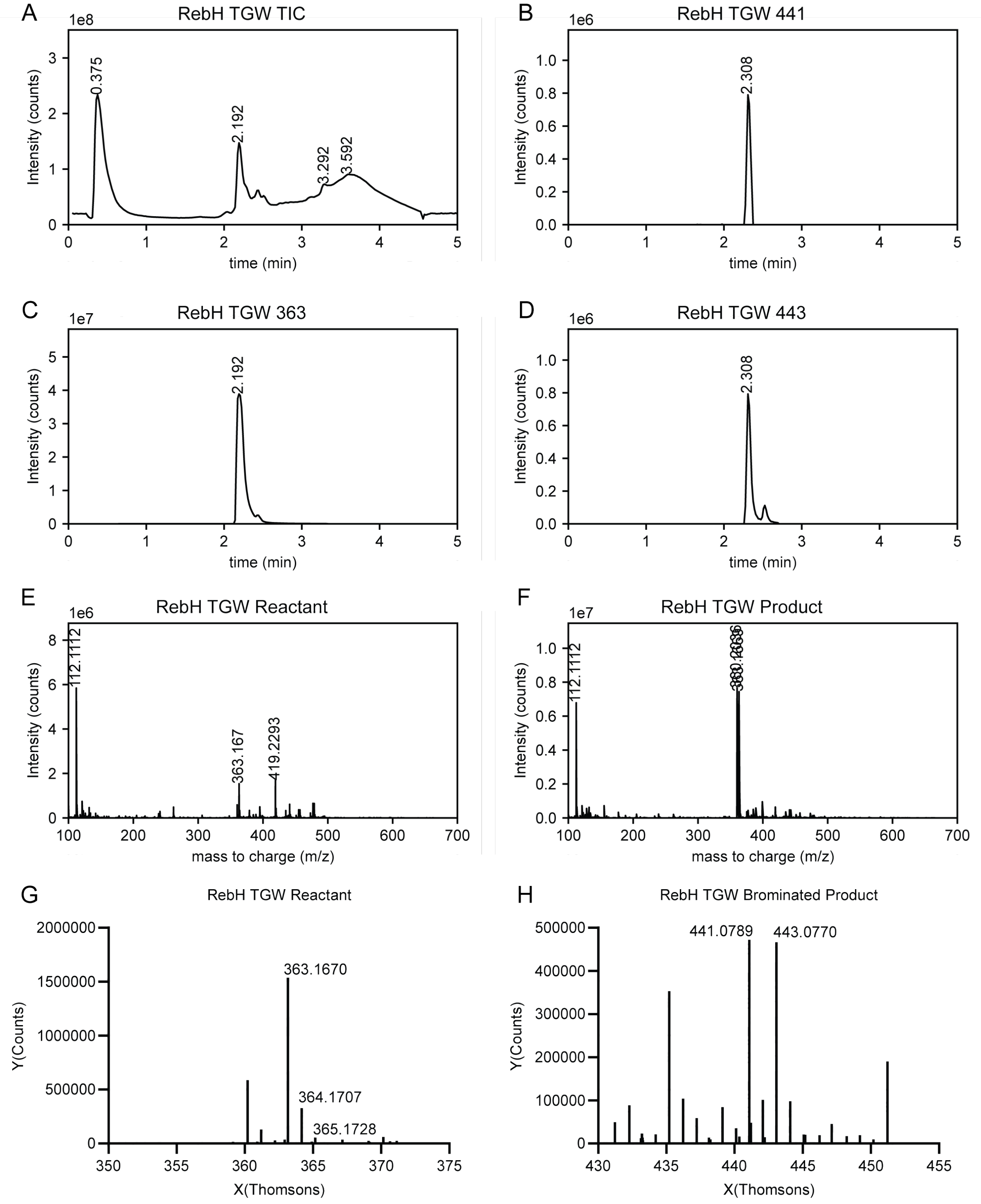
A
RebH TGW TIC
1e8
Intensity (counts)
time (min)
0.375
2.192
3.292
3.592
B
RebH TGW 441
1e6
Intensity (counts)
time (min)
2.308
C
RebH TGW 363
1e7
Intensity (counts)
time (min)
2.192
D
RebH TGW 443
1e6
Intensity (counts)
time (min)
2.308
E
RebH TGW Reactant
1e6
Intensity (counts)
mass to charge (m/z)
112.1112
363.167
419.2293
F
RebH TGW Product
1e7
Intensity (counts)
mass to charge (m/z)
112.1112
G
RebH TGW Reactant
Y(Counts)
X(Thomsons)
363.1670
364.1707
365.1728
H
RebH TGW Brominated Product
Y(Counts)
X(Thomsons)
441.0789
443.0770

#### RebH VGW

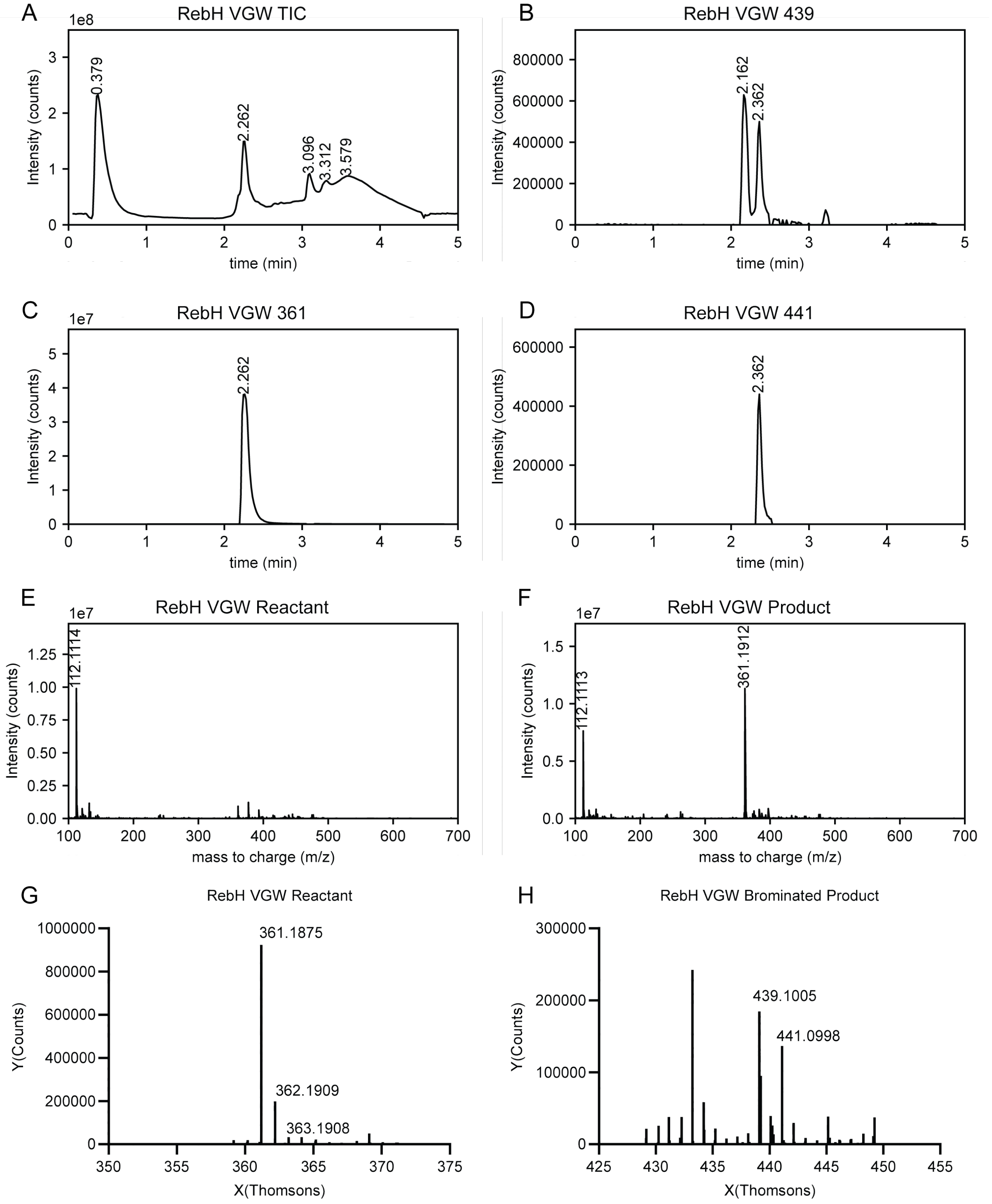

A
RebH VGW TIC
1e8
Intensity (counts)
time (min)
0.379
2.262
3.096
3.312
3.579
B
RebH VGW 439
Intensity (counts)
time (min)
2.162
2.362
C
RebH VGW 361
1e7
Intensity (counts)
time (min)
2.262
D
RebH VGW 441
Intensity (counts)
time (min)
2.362
E
RebH VGW Reactant
1e7
Intensity (counts)
mass to charge (m/z)
112.1114
F
RebH VGW Product
1e7
Intensity (counts)
mass to charge (m/z)
112.1113
361.1912
G
RebH VGW Reactant
Y(Counts)
X(Thomsons)
361.1875
362.1909
363.1908
H
RebH VGW Brominated Product
Y(Counts)
X(Thomsons)
439.1005
441.0998

#### RebH YGW

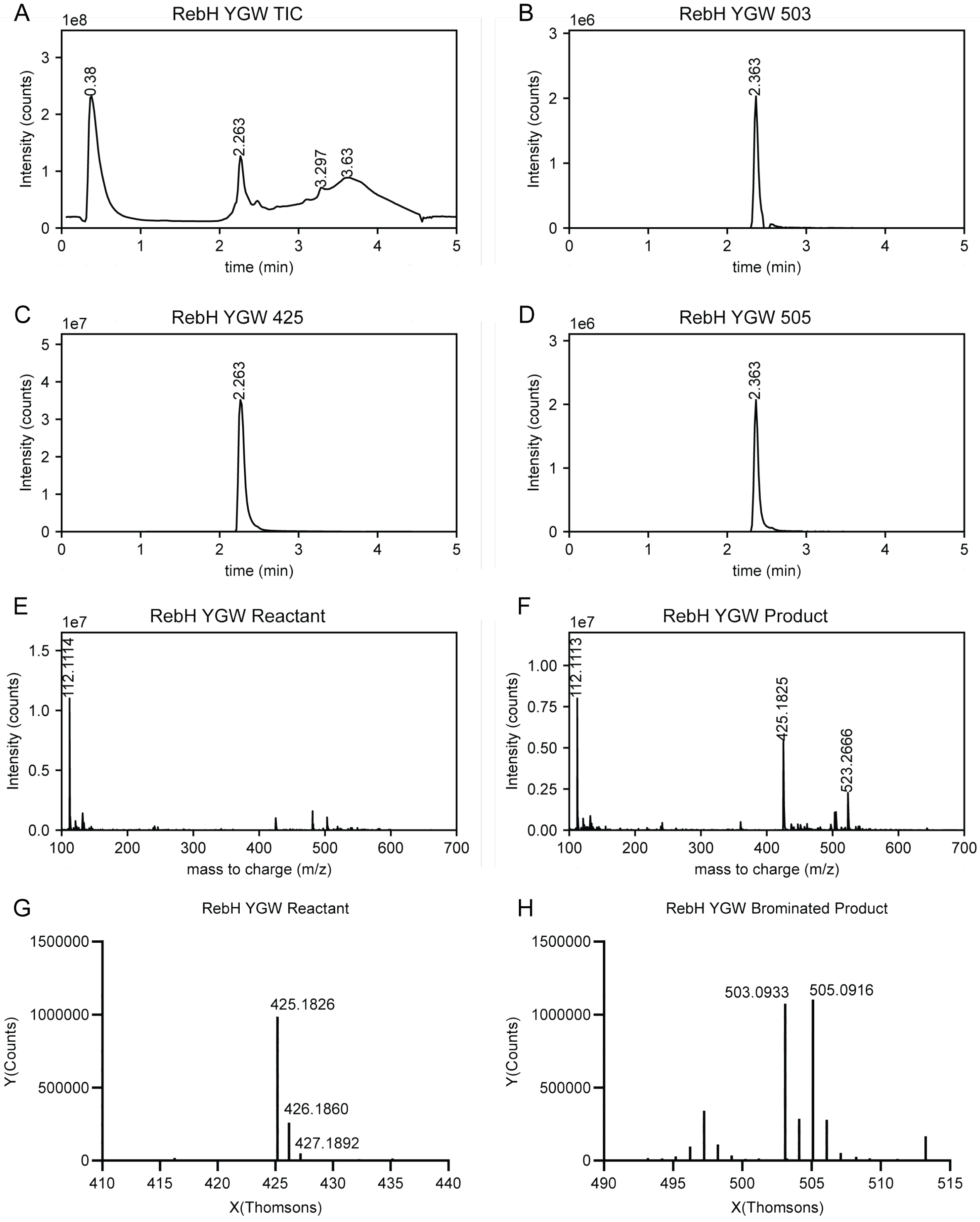
A
RebH YGW TIC
1e8
Intensity (counts)
time (min)
0.38
2.263
3.297
3.63
B
RebH YGW 503
1e6
2.363
C
RebH YGW 425
1e7
2.263
D
RebH YGW 505
1e6
2.363
E
RebH YGW Reactant
1e7
112.1114
mass to charge (m/z)
F
RebH YGW Product
1e7
112.1113
425.1825
523.2666
G
RebH YGW Reactant
Y(Counts)
X(Thomsons)
425.1826
426.1860
427.1892
H
RebH YGW Brominated Product
503.0933
505.0916

#### RebH GAW

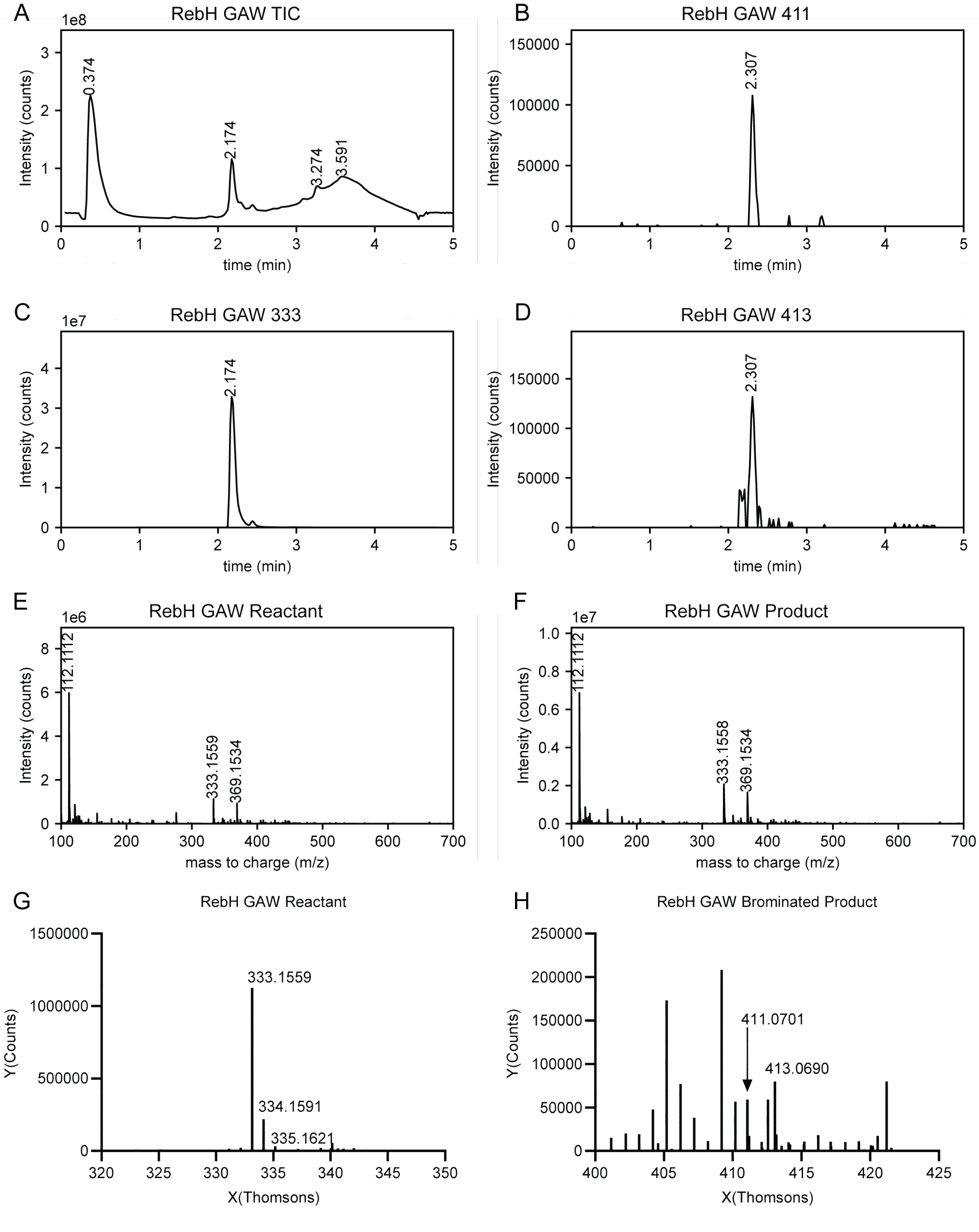
A
RebH GAW TIC
1e8
Intensity (counts)
time (min)
0.374
2.174
3.274
3.591
B
RebH GAW 411
Intensity (counts)
time (min)
2.307
C
RebH GAW 333
1e7
Intensity (counts)
time (min)
2.174
D
RebH GAW 413
Intensity (counts)
time (min)
2.307
E
RebH GAW Reactant
1e6
Intensity (counts)
mass to charge (m/z)
112.1112
333.1559
369.1534
F
RebH GAW Product
1e7
Intensity (counts)
mass to charge (m/z)
112.1112
333.1558
369.1534
G
RebH GAW Reactant
Y(Counts)
X(Thomsons)
333.1559
334.1591
335.1621
H
RebH GAW Brominated Product
Y(Counts)
X(Thomsons)
411.0701
413.0690

#### RebH GCW

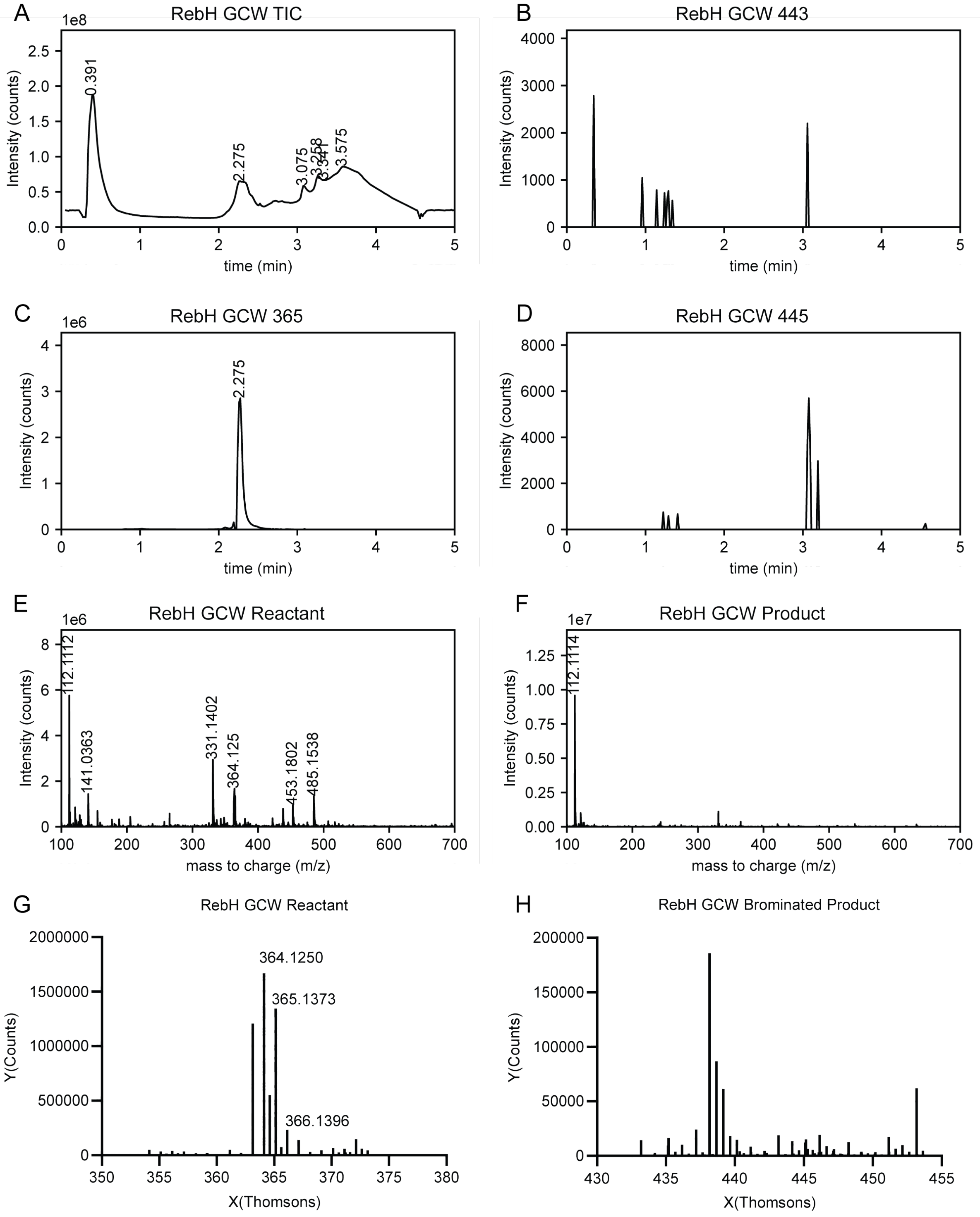

A
RebH GCW TIC
1e8
Intensity (counts)
time (min)
0.391
2.275
3.075
3.258
3.341
3.575
B
RebH GCW 443
Intensity (counts)
time (min)
C
RebH GCW 365
1e6
Intensity (counts)
time (min)
2.275
D
RebH GCW 445
Intensity (counts)
time (min)
E
RebH GCW Reactant
1e6
Intensity (counts)
mass to charge (m/z)
112.1112
141.0363
331.1402
364.125
453.1802
485.1538
F
RebH GCW Product
1e7
Intensity (counts)
mass to charge (m/z)
112.1114
G
RebH GCW Reactant
Y(Counts)
X(Thomsons)
364.1250
365.1373
366.1396
H
RebH GCW Brominated Product
Y(Counts)
X(Thomsons)

#### RebH GDW

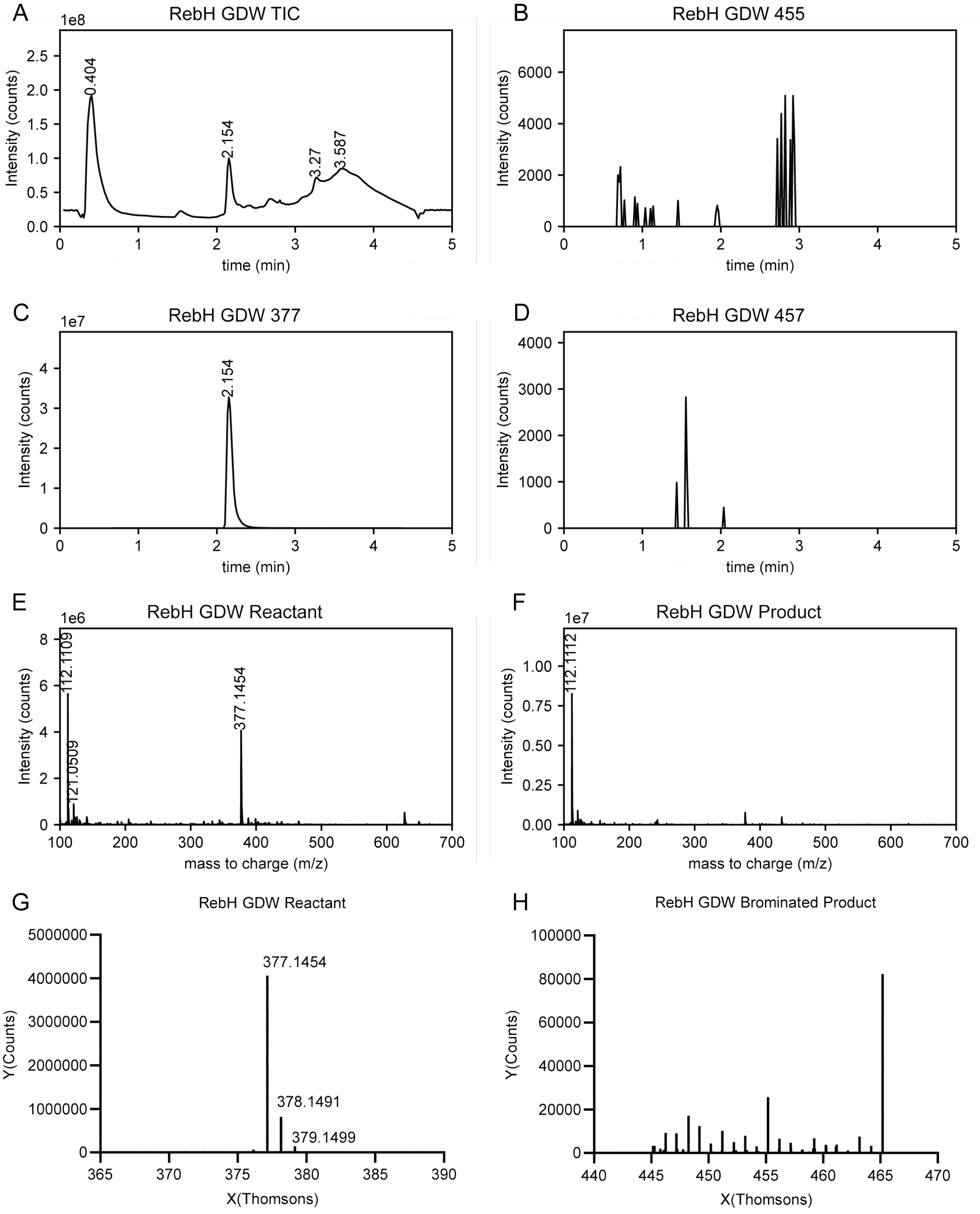

A
RebH GDW TIC
1e8
Intensity (counts)
time (min)
0.404
2.154
3.27
3.587
B
RebH GDW 455
Intensity (counts)
time (min)
C
RebH GDW 377
1e7
Intensity (counts)
time (min)
2.154
D
RebH GDW 457
Intensity (counts)
time (min)
E
RebH GDW Reactant
1e6
Intensity (counts)
mass to charge (m/z)
112.1109
121.0509
377.1454
F
RebH GDW Product
1e7
Intensity (counts)
mass to charge (m/z)
112.1112
G
RebH GDW Reactant
Y(Counts)
X(Thomsons)
377.1454
378.1491
379.1499
H
RebH GDW Brominated Product
Y(Counts)
X(Thomsons)

#### RebH GEW

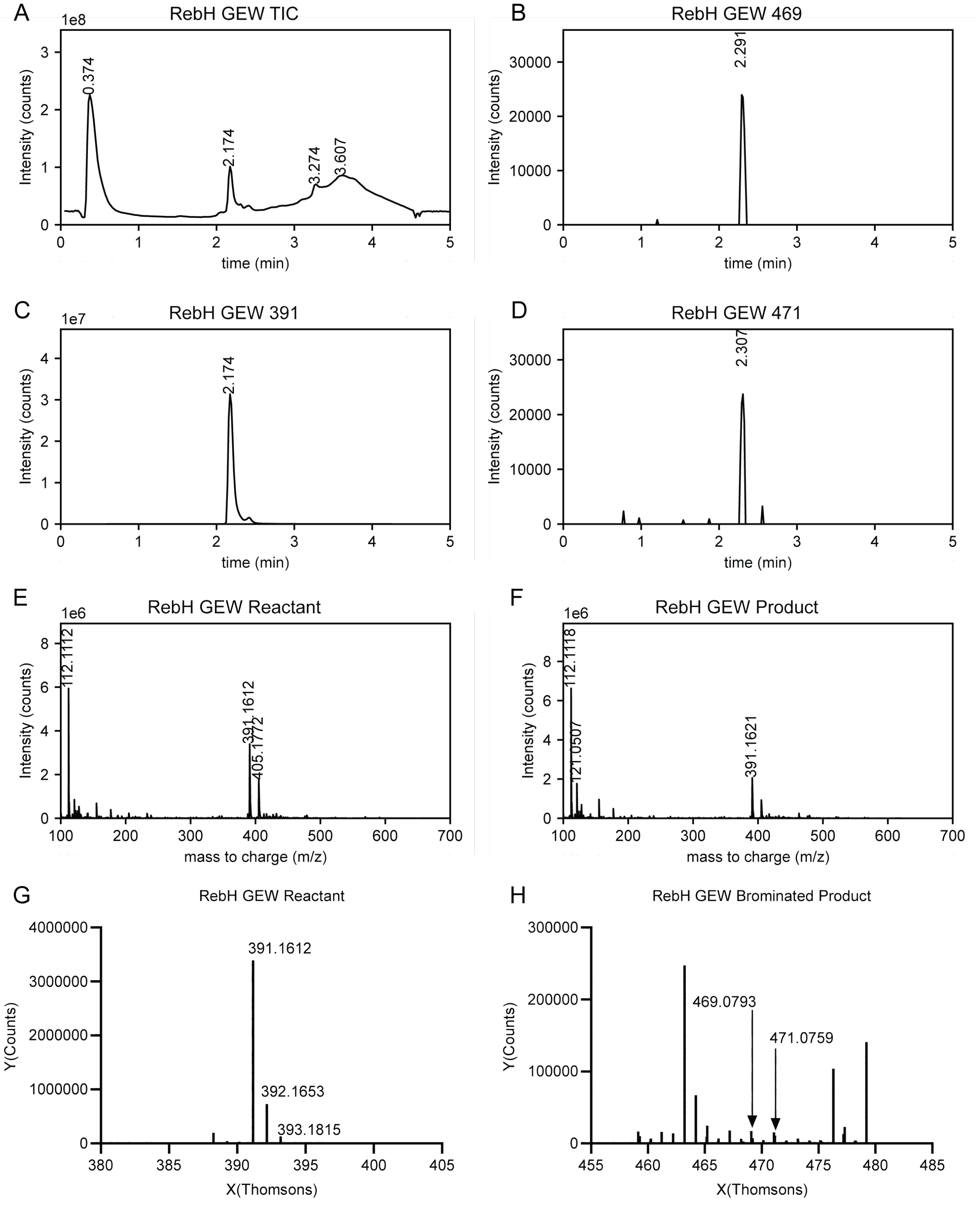

A
RebH GEW TIC
1e8
Intensity (counts)
time (min)
0.374
2.174
3.274
3.607
B
RebH GEW 469
Intensity (counts)
time (min)
2.291
C
RebH GEW 391
1e7
Intensity (counts)
time (min)
2.174
D
RebH GEW 471
Intensity (counts)
time (min)
2.307
E
RebH GEW Reactant
1e6
Intensity (counts)
mass to charge (m/z)
112.1112
391.1612
405.1772
F
RebH GEW Product
1e6
Intensity (counts)
mass to charge (m/z)
112.1118
121.0507
391.1621
G
RebH GEW Reactant
Y(Counts)
X(Thomsons)
391.1612
392.1653
393.1815
H
RebH GEW Brominated Product
Y(Counts)
X(Thomsons)
469.0793
471.0759

#### RebH GFW

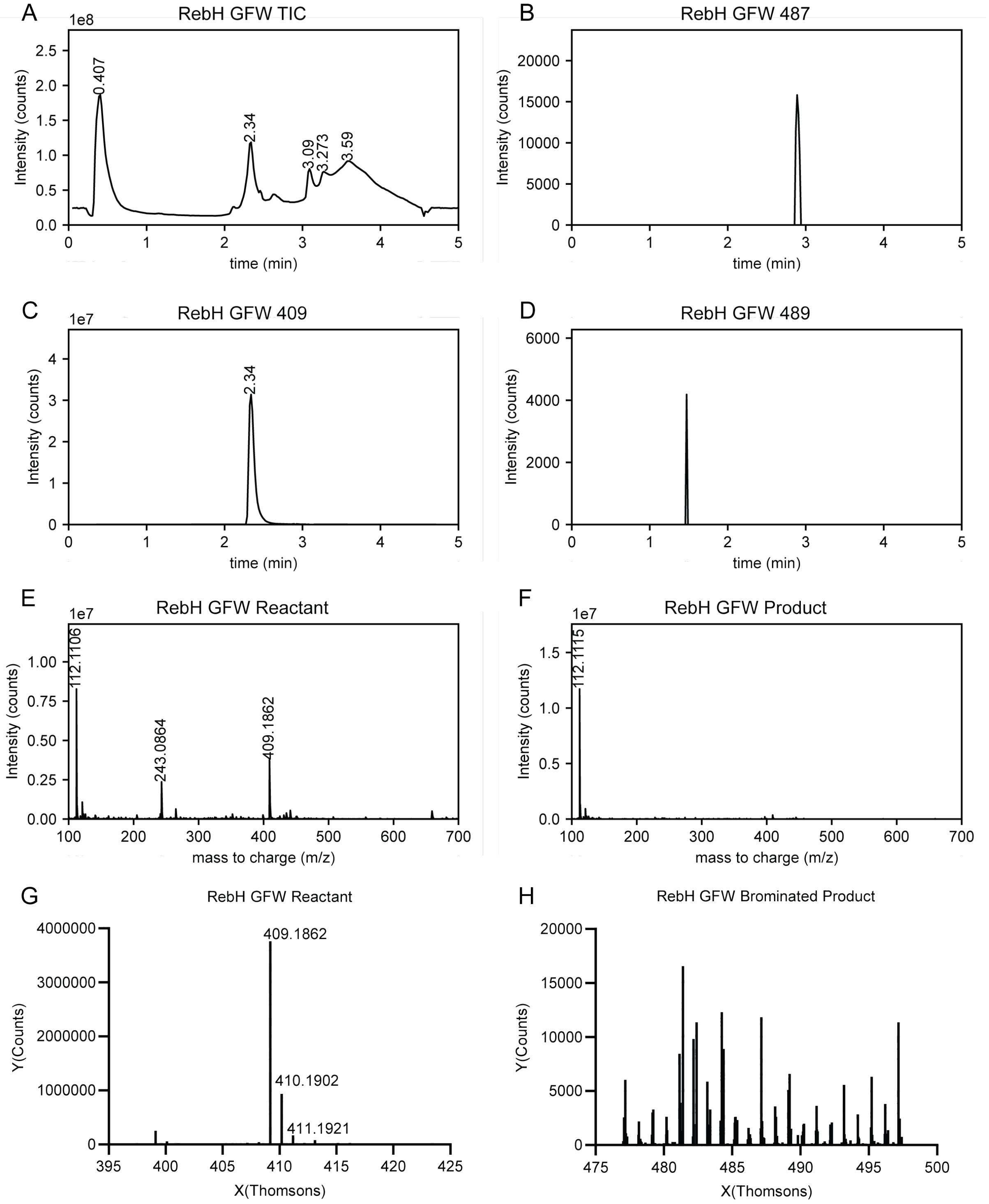

A
RebH GFW TIC
1e8
Intensity (counts)
time (min)
0.407
2.34
3.09
3.273
3.59
B
RebH GFW 487
Intensity (counts)
time (min)
C
RebH GFW 409
1e7
Intensity (counts)
time (min)
2.34
D
RebH GFW 489
Intensity (counts)
time (min)
E
RebH GFW Reactant
1e7
Intensity (counts)
mass to charge (m/z)
112.1106
243.0864
409.1862
F
RebH GFW Product
1e7
Intensity (counts)
mass to charge (m/z)
112.1115
G
RebH GFW Reactant
Y(Counts)
X(Thomsons)
409.1862
410.1902
411.1921
H
RebH GFW Brominated Product
Y(Counts)
X(Thomsons)

#### RebH GHW

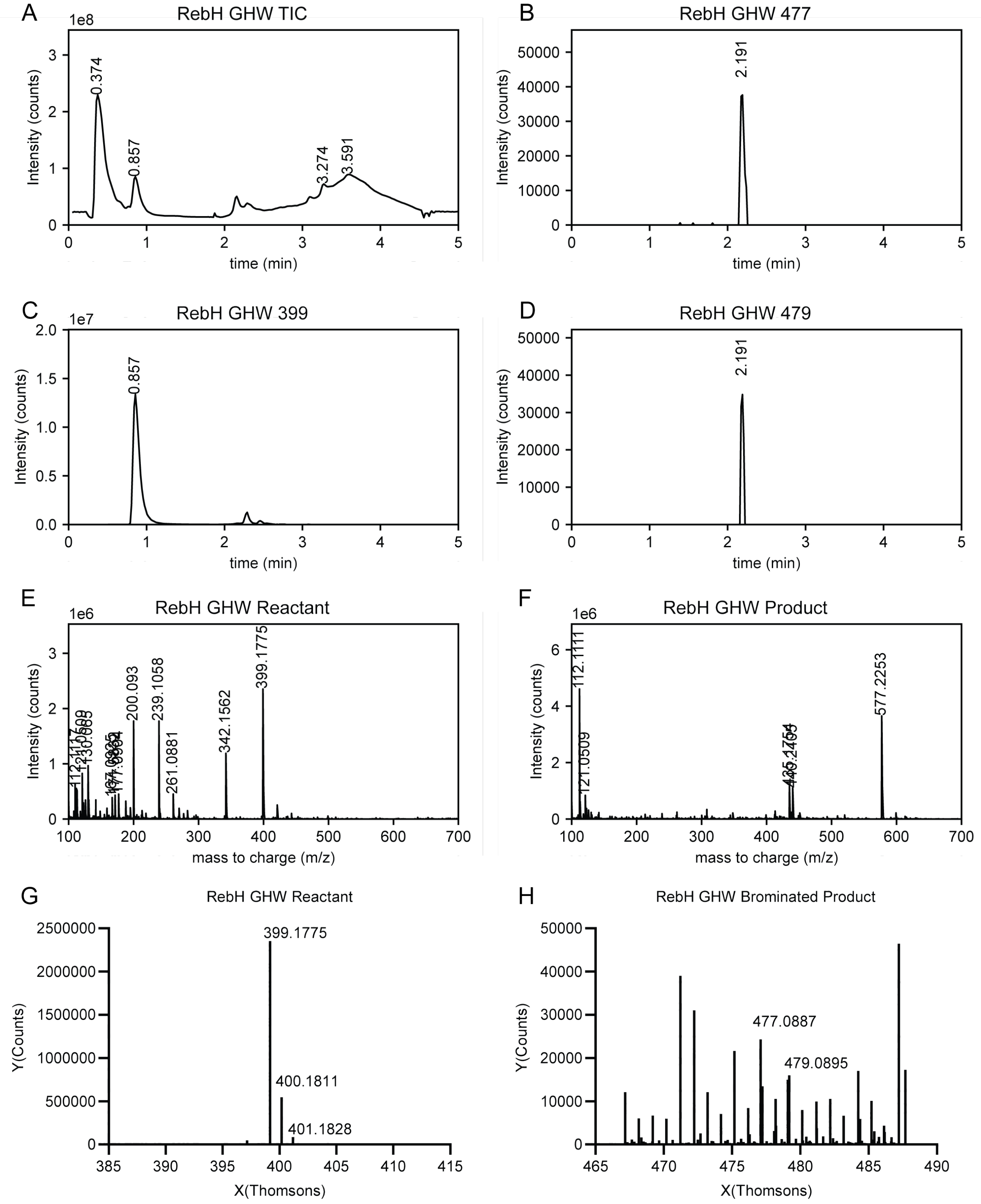

A
RebH GHW TIC
1e8
Intensity (counts)
time (min)
0.374
0.857
3.274
3.591
B
RebH GHW 477
Intensity (counts)
time (min)
2.191
C
RebH GHW 399
1e7
Intensity (counts)
time (min)
0.857
D
RebH GHW 479
Intensity (counts)
time (min)
2.191
E
RebH GHW Reactant
1e6
Intensity (counts)
mass to charge (m/z)
200.093
239.1058
261.0881
342.1562
399.1775
F
RebH GHW Product
1e6
Intensity (counts)
mass to charge (m/z)
112.1111
121.0509
577.2253
G
RebH GHW Reactant
Y(Counts)
X(Thomsons)
399.1775
400.1811
401.1828
H
RebH GHW Brominated Product
Y(Counts)
X(Thomsons)
477.0887
479.0895

#### RebH GKW

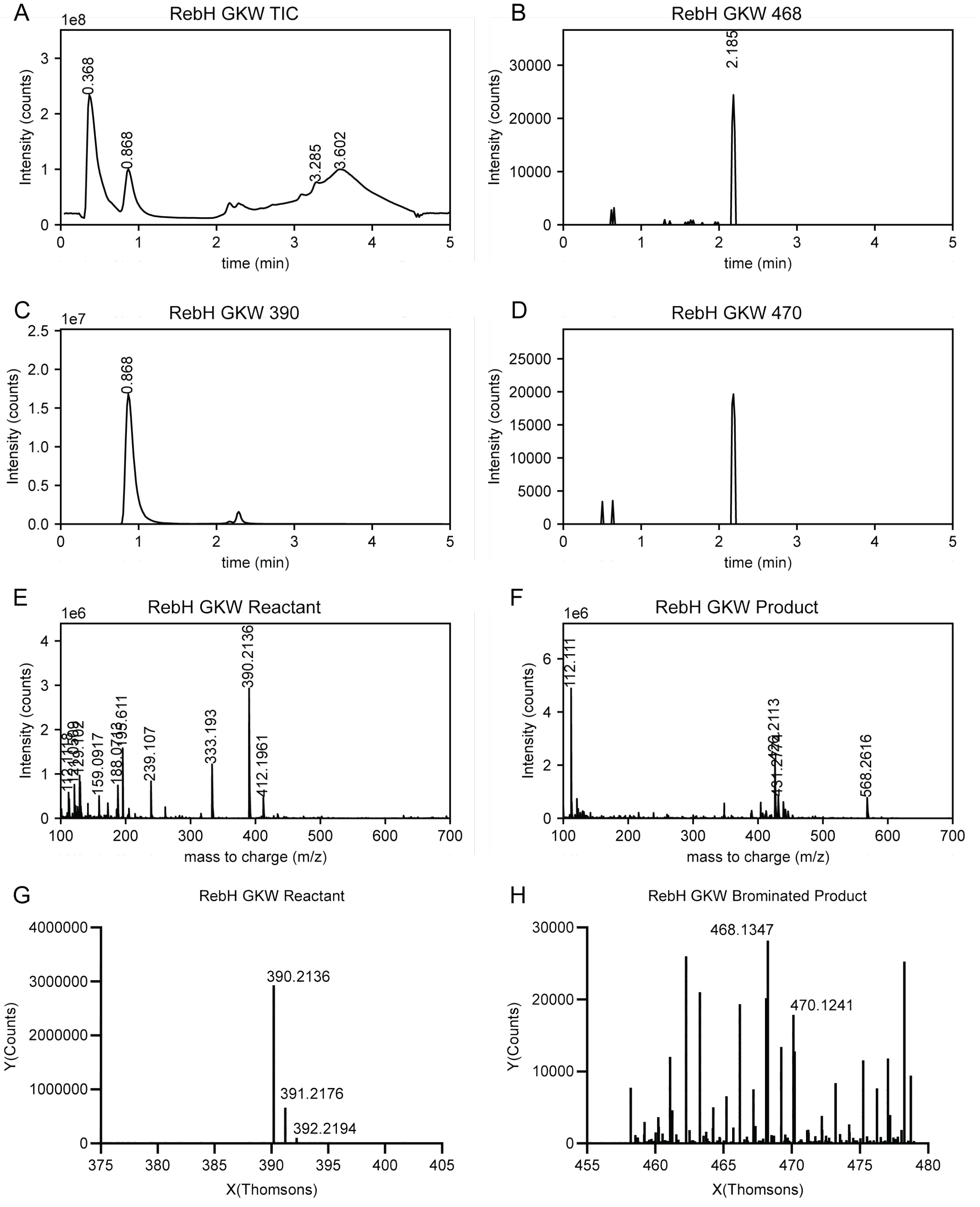

A
RebH GKW TIC
1e8
Intensity (counts)
time (min)
0.368
0.868
3.285
3.602
B
RebH GKW 468
Intensity (counts)
time (min)
2.185
C
RebH GKW 390
1e7
Intensity (counts)
time (min)
0.868
D
RebH GKW 470
Intensity (counts)
time (min)
E
RebH GKW Reactant
1e6
Intensity (counts)
mass to charge (m/z)
159.0917
239.107
333.193
390.2136
412.1961
F
RebH GKW Product
1e6
Intensity (counts)
mass to charge (m/z)
112.111
568.2616
G
RebH GKW Reactant
Y(Counts)
X(Thomsons)
390.2136
391.2176
392.2194
H
RebH GKW Brominated Product
Y(Counts)
X(Thomsons)
468.1347
470.1241

#### RebH GLW

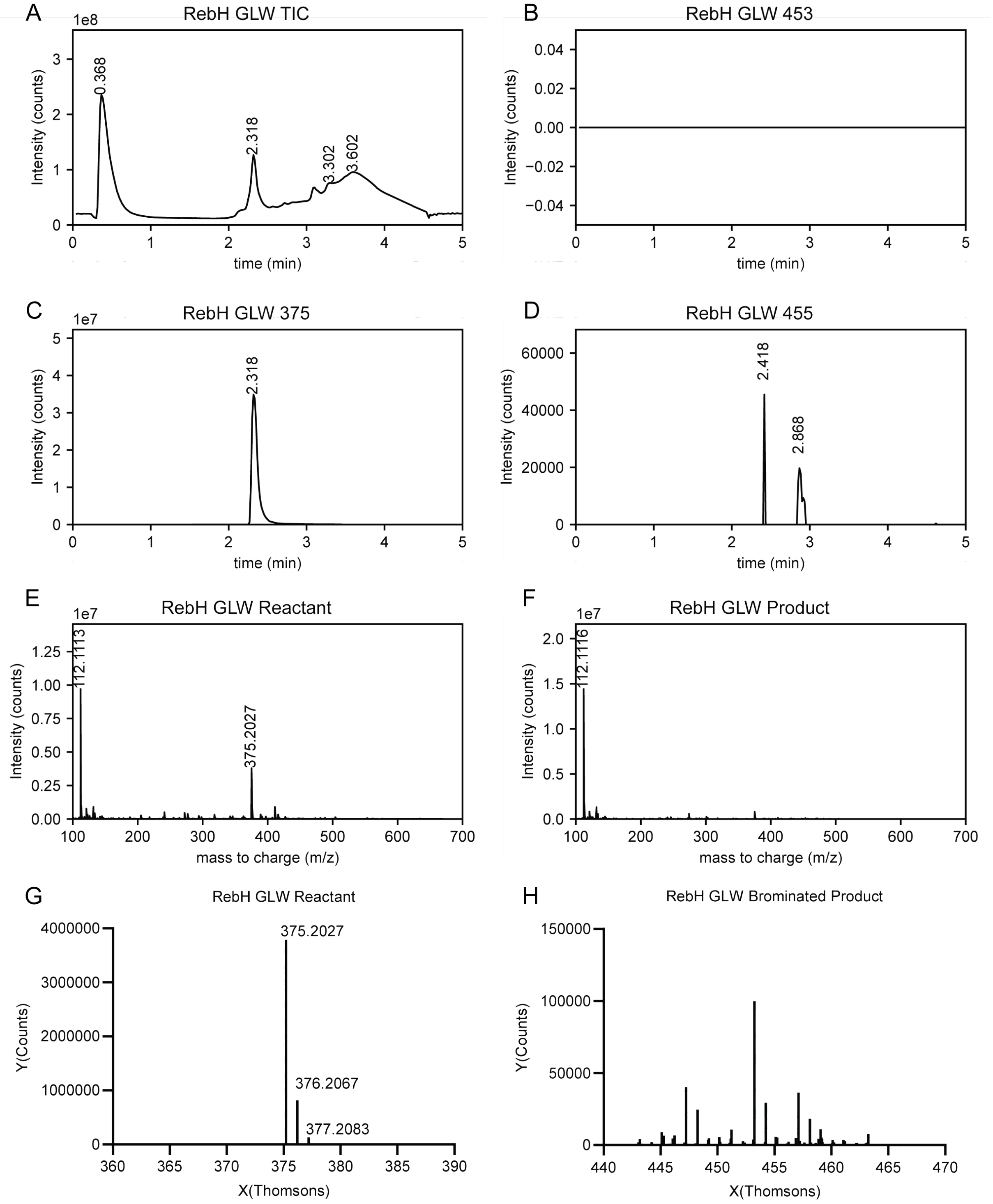

A
RebH GLW TIC
1e8
Intensity (counts)
time (min)
0.368
2.318
3.302
3.602
B
RebH GLW 453
Intensity (counts)
time (min)
C
RebH GLW 375
1e7
Intensity (counts)
time (min)
2.318
D
RebH GLW 455
Intensity (counts)
time (min)
2.418
2.868
E
RebH GLW Reactant
1e7
Intensity (counts)
mass to charge (m/z)
112.1113
375.2027
F
RebH GLW Product
1e7
Intensity (counts)
mass to charge (m/z)
112.1116
G
RebH GLW Reactant
Y(Counts)
X(Thomsons)
375.2027
376.2067
377.2083
H
RebH GLW Brominated Product
Y(Counts)
X(Thomsons)

#### RebH GMW

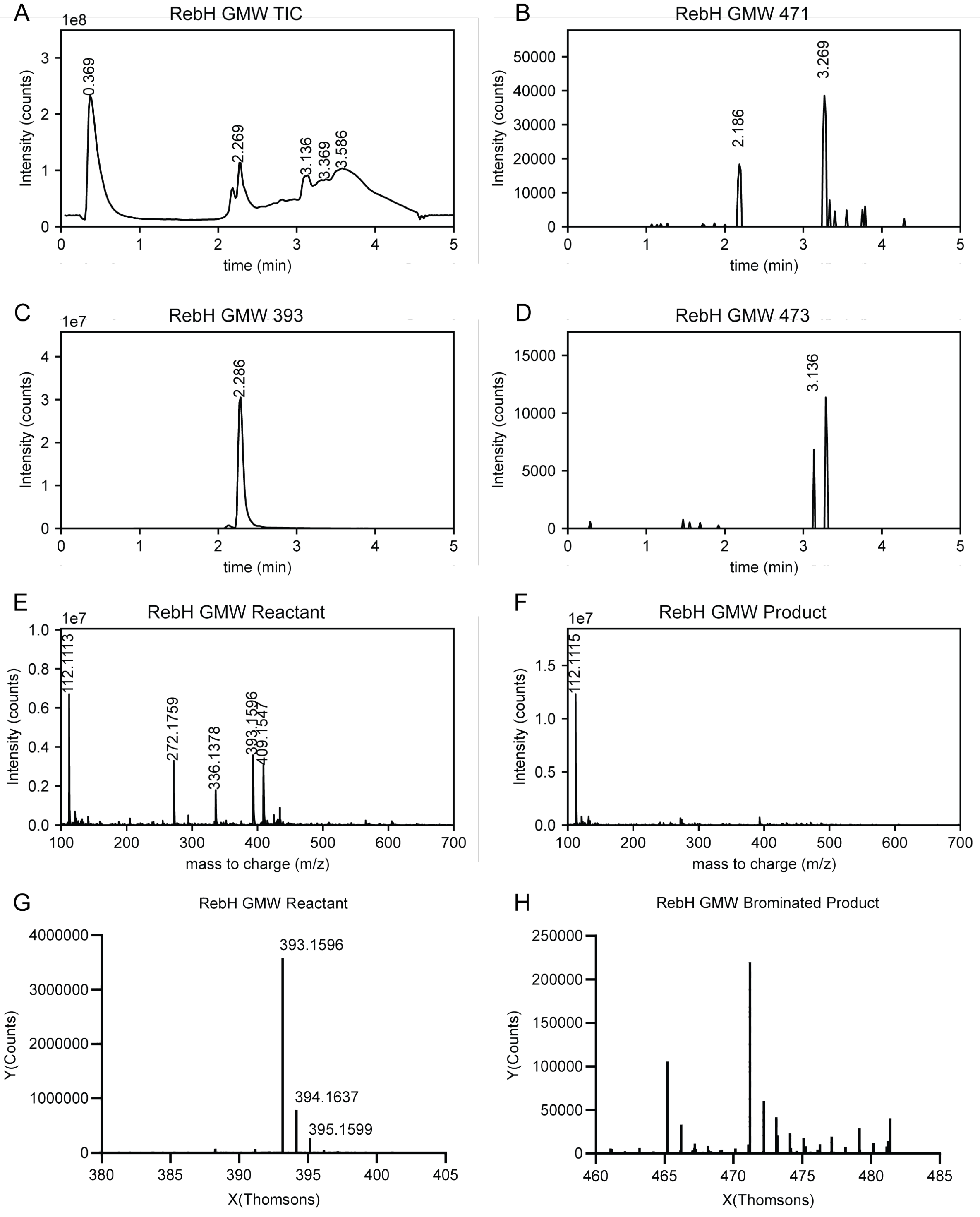

A
RebH GMW TIC
1e8
Intensity (counts)
time (min)
0.369
2.269
3.136
3.369
3.586
B
RebH GMW 471
Intensity (counts)
time (min)
2.186
3.269
C
RebH GMW 393
1e7
Intensity (counts)
time (min)
2.286
D
RebH GMW 473
Intensity (counts)
time (min)
3.136
E
RebH GMW Reactant
1e7
Intensity (counts)
mass to charge (m/z)
112.1113
272.1759
336.1378
393.1596
409.1547
F
RebH GMW Product
1e7
Intensity (counts)
mass to charge (m/z)
112.1115
G
RebH GMW Reactant
Y(Counts)
X(Thomsons)
393.1596
394.1637
395.1599
H
RebH GMW Brominated Product
Y(Counts)
X(Thomsons)

#### RebH GNW

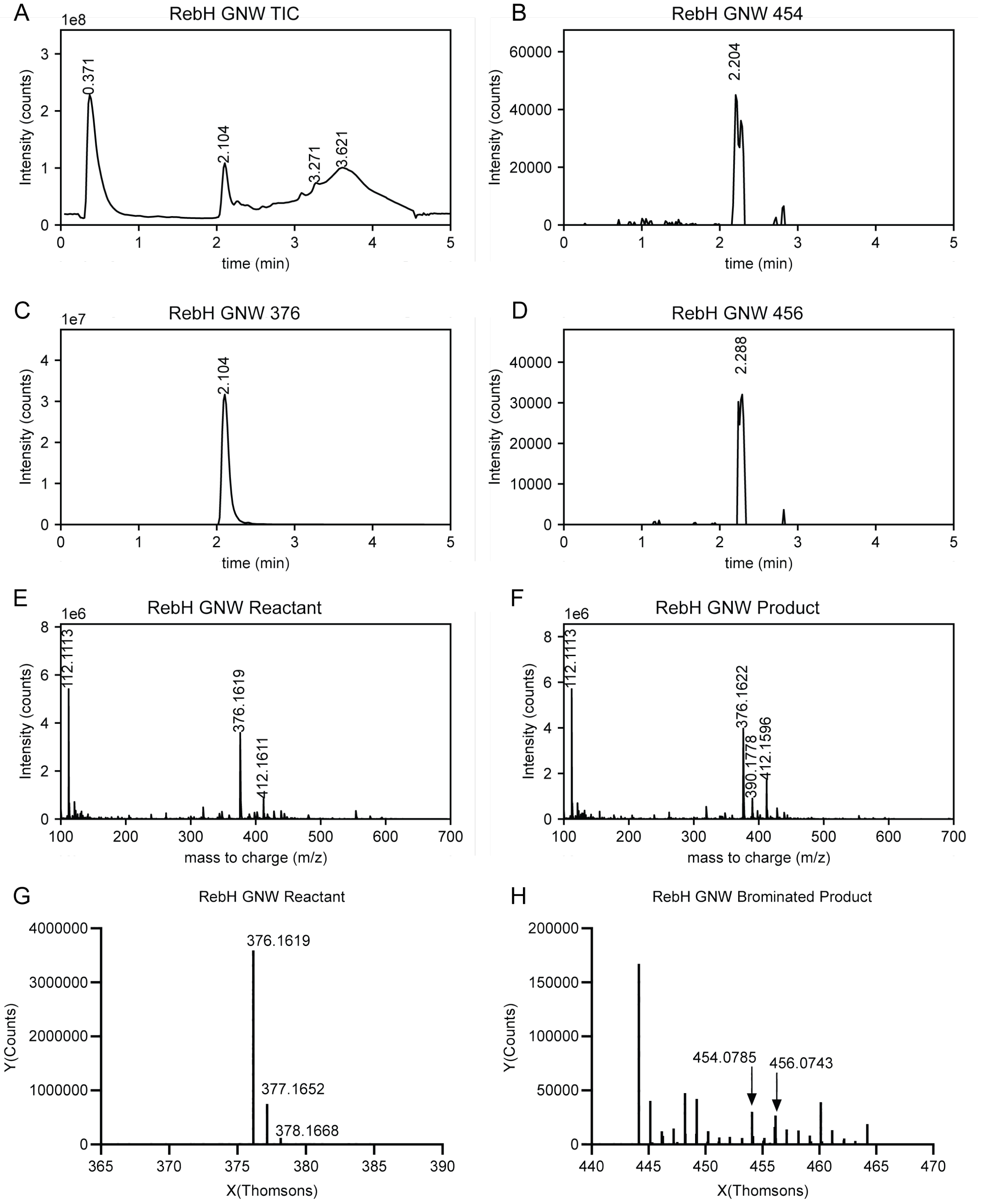

A
RebH GNW TIC
1e8
Intensity (counts)
time (min)
0.371
2.104
3.271
3.621
B
RebH GNW 454
Intensity (counts)
time (min)
2.204
C
RebH GNW 376
1e7
Intensity (counts)
time (min)
2.104
D
RebH GNW 456
Intensity (counts)
time (min)
2.288
E
RebH GNW Reactant
1e6
Intensity (counts)
mass to charge (m/z)
112.1113
376.1619
412.1611
F
RebH GNW Product
1e6
Intensity (counts)
mass to charge (m/z)
112.1113
376.1622
390.1778
412.1596
G
RebH GNW Reactant
Y(Counts)
X(Thomsons)
376.1619
377.1652
378.1668
H
RebH GNW Brominated Product
Y(Counts)
X(Thomsons)
454.0785
456.0743

#### RebH GPW

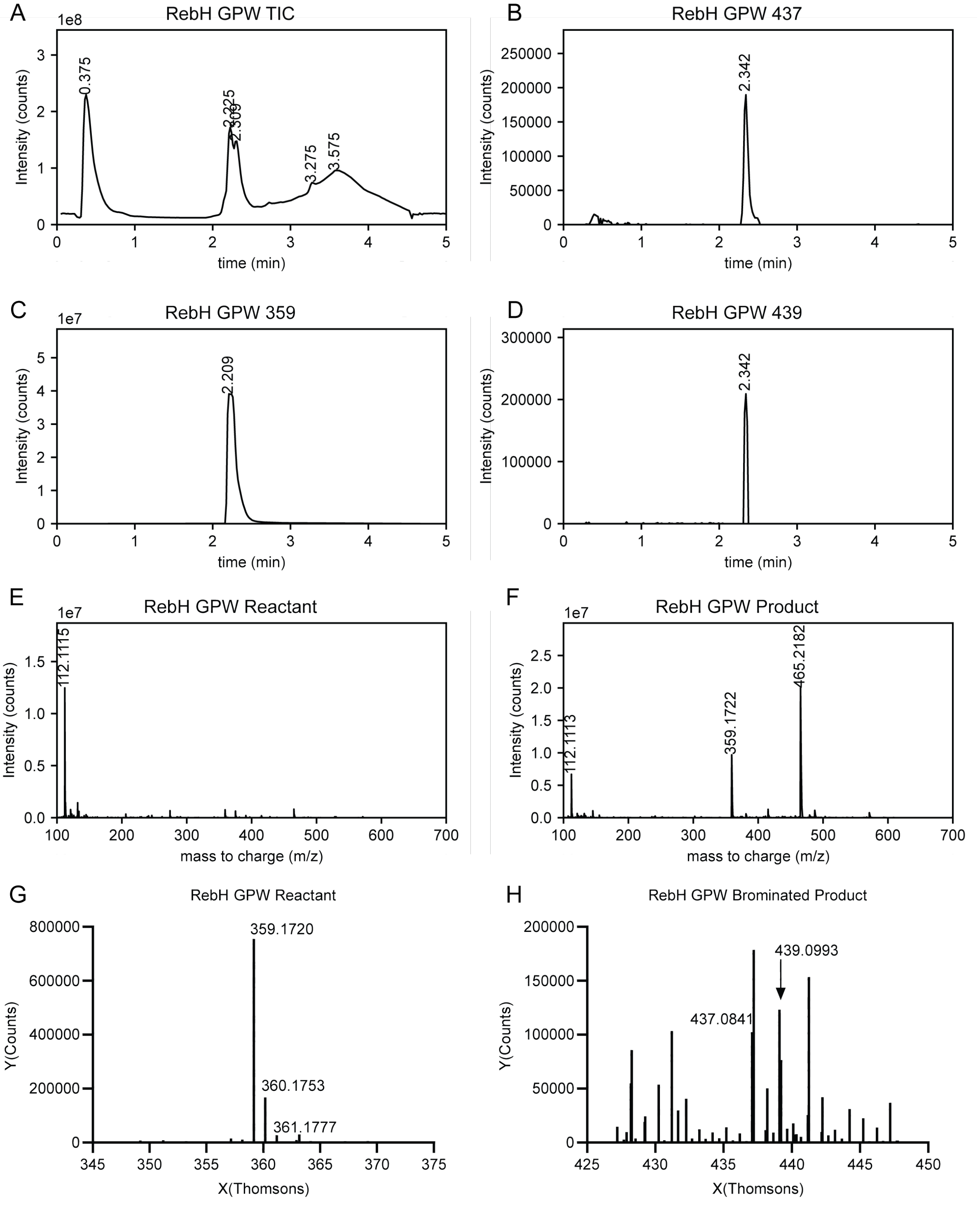
A
RebH GPW TIC
1e8
Intensity (counts)
time (min)
0.375
2.225
2.309
3.275
3.575
B
RebH GPW 437
Intensity (counts)
time (min)
2.342
C
RebH GPW 359
1e7
Intensity (counts)
time (min)
2.209
D
RebH GPW 439
Intensity (counts)
time (min)
2.342
E
RebH GPW Reactant
1e7
Intensity (counts)
mass to charge (m/z)
112.1115
F
RebH GPW Product
1e7
Intensity (counts)
mass to charge (m/z)
112.1113
359.1722
465.2182
G
RebH GPW Reactant
Y(Counts)
X(Thomsons)
359.1720
360.1753
361.1777
H
RebH GPW Brominated Product
Y(Counts)
X(Thomsons)
439.0993
437.0841

#### RebH GQW

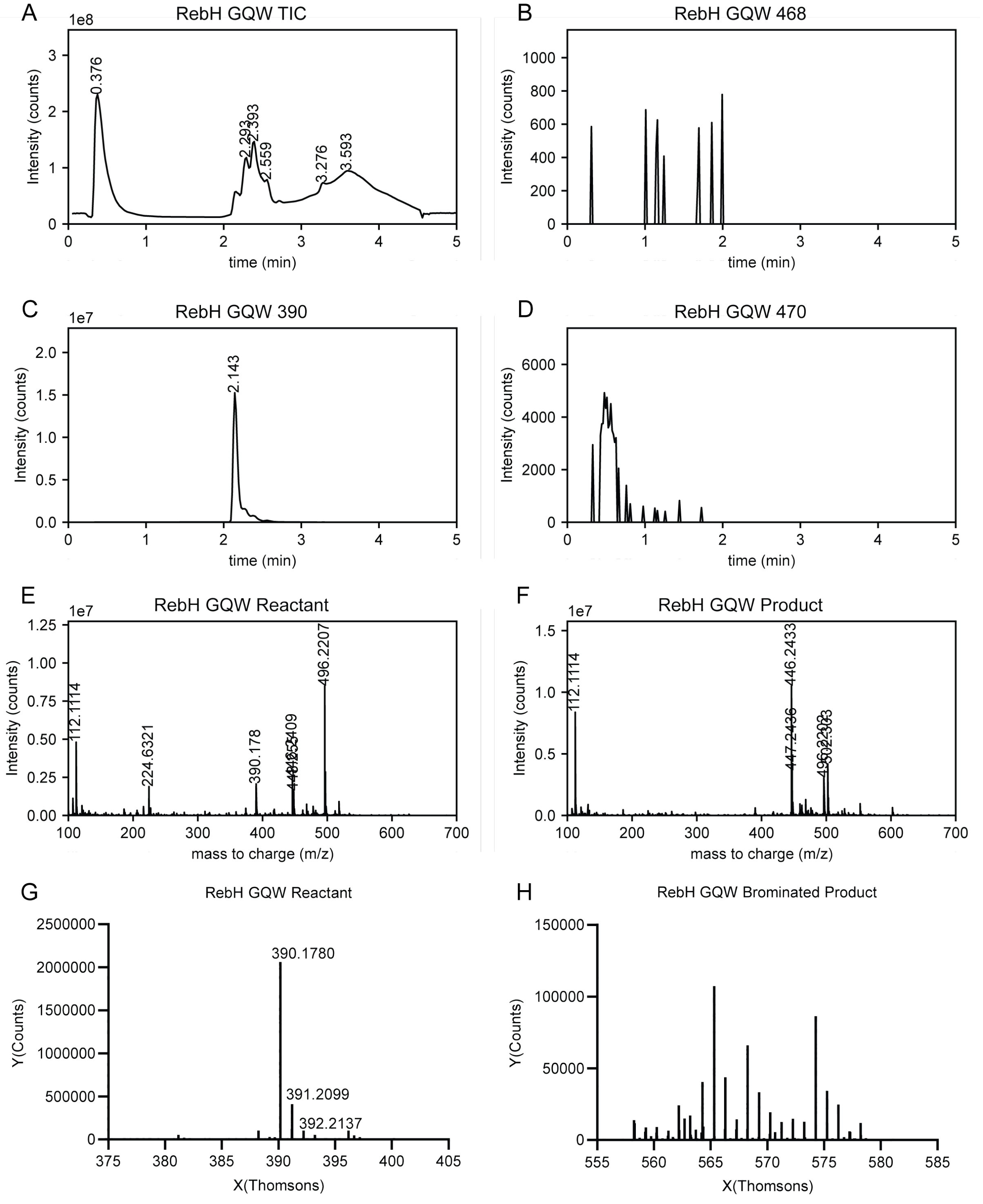
A
RebH GQW TIC
1e8
Intensity (counts)
time (min)
0.376
2.293
2.393
2.559
3.276
3.593
B
RebH GQW 468
Intensity (counts)
time (min)
C
RebH GQW 390
1e7
Intensity (counts)
time (min)
2.143
D
RebH GQW 470
Intensity (counts)
time (min)
E
RebH GQW Reactant
1e7
Intensity (counts)
mass to charge (m/z)
112.1114
224.6321
390.178
496.2207
F
RebH GQW Product
1e7
Intensity (counts)
mass to charge (m/z)
112.1114
446.2433
447.2436
G
RebH GQW Reactant
Y(Counts)
X(Thomsons)
390.1780
391.2099
392.2137
H
RebH GQW Brominated Product
Y(Counts)
X(Thomsons)

**RebH GRW**

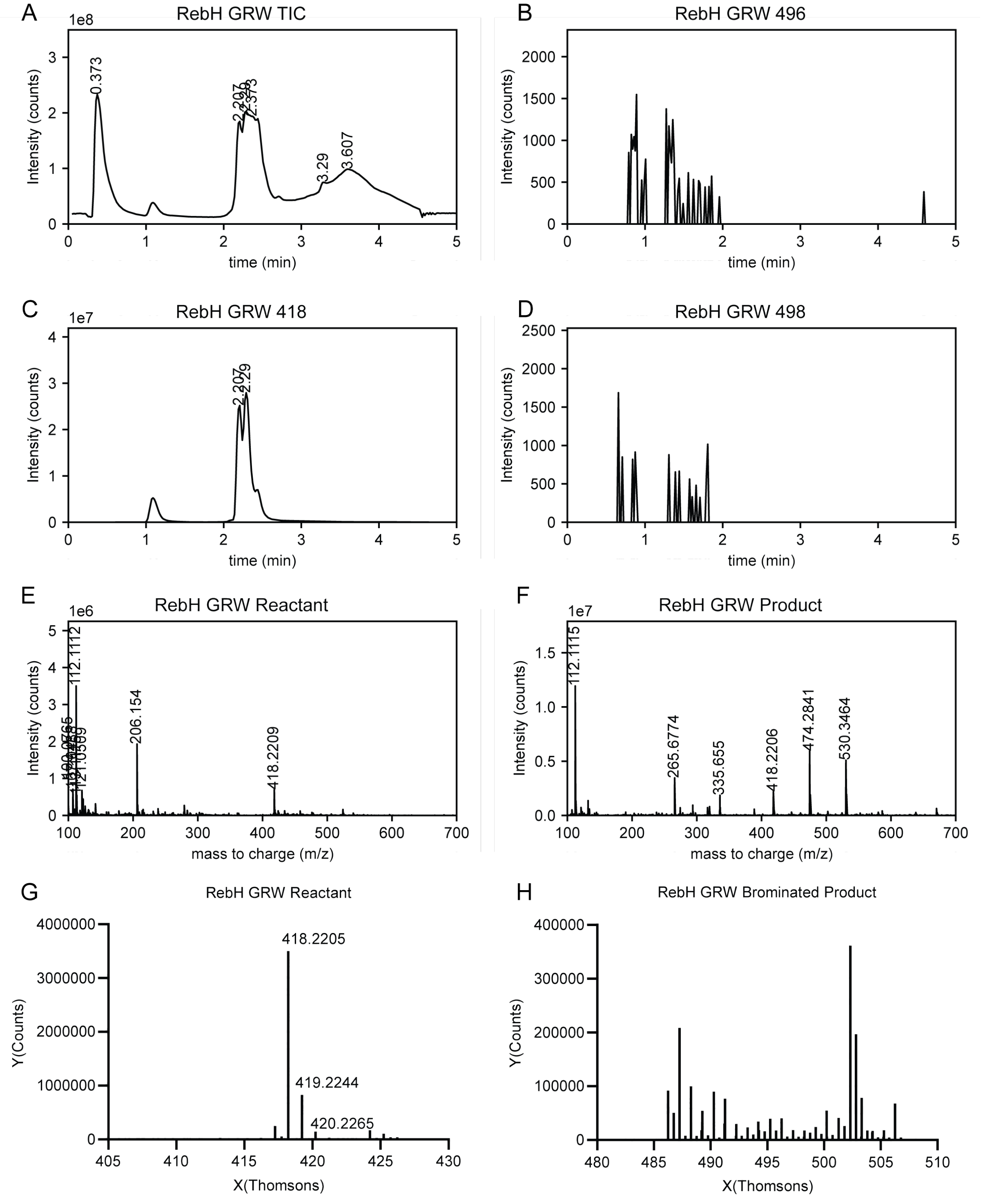
A
RebH GRW TIC
1e8
0.373
2.207
2.373
3.29
3.607
Intensity (counts)
time (min)
B
RebH GRW 496
Intensity (counts)
time (min)
C
RebH GRW 418
1e7
2.207
2.29
Intensity (counts)
time (min)
D
RebH GRW 498
Intensity (counts)
time (min)
E
RebH GRW Reactant
1e6
112.1112
206.154
418.2209
Intensity (counts)
mass to charge (m/z)
F
RebH GRW Product
1e7
112.1115
265.6774
335.655
418.2206
474.2841
530.3464
Intensity (counts)
mass to charge (m/z)
G
RebH GRW Reactant
418.2205
419.2244
420.2265
Y(Counts)
X(Thomsons)
H
RebH GRW Brominated Product
Y(Counts)
X(Thomsons)

#### RebH GTW

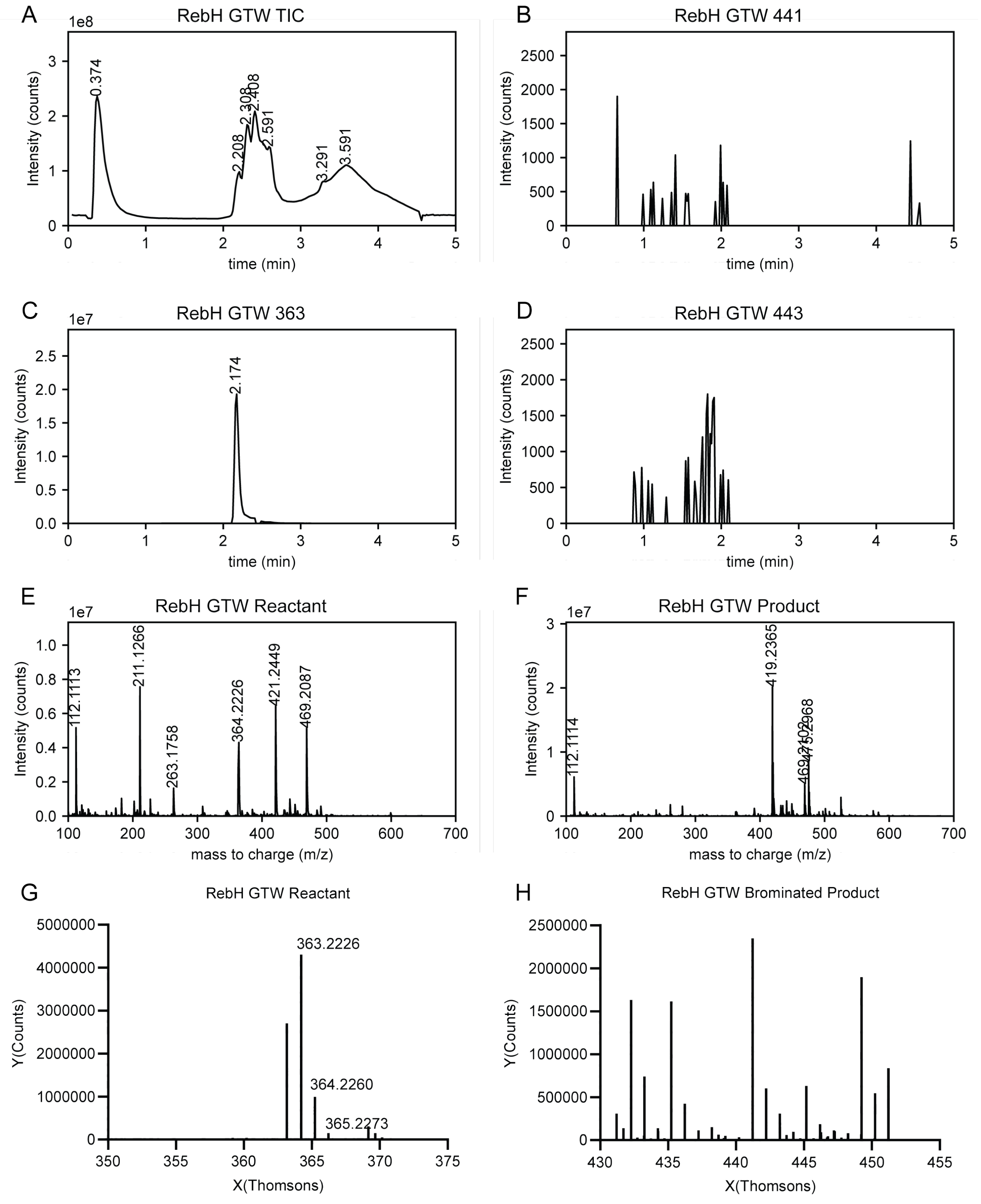
A
RebH GTW TIC
1e8
Intensity (counts)
time (min)
0.374
2.208
2.308
2.408
2.591
3.291
3.591
B
RebH GTW 441
Intensity (counts)
time (min)
C
RebH GTW 363
1e7
Intensity (counts)
time (min)
2.174
D
RebH GTW 443
Intensity (counts)
time (min)
E
RebH GTW Reactant
1e7
Intensity (counts)
mass to charge (m/z)
112.1113
211.1266
263.1758
364.2226
421.2449
469.2087
F
RebH GTW Product
1e7
Intensity (counts)
mass to charge (m/z)
112.1114
419.2365
G
RebH GTW Reactant
Y(Counts)
X(Thomsons)
363.2226
364.2260
365.2273
H
RebH GTW Brominated Product
Y(Counts)
X(Thomsons)

### RebH GVW

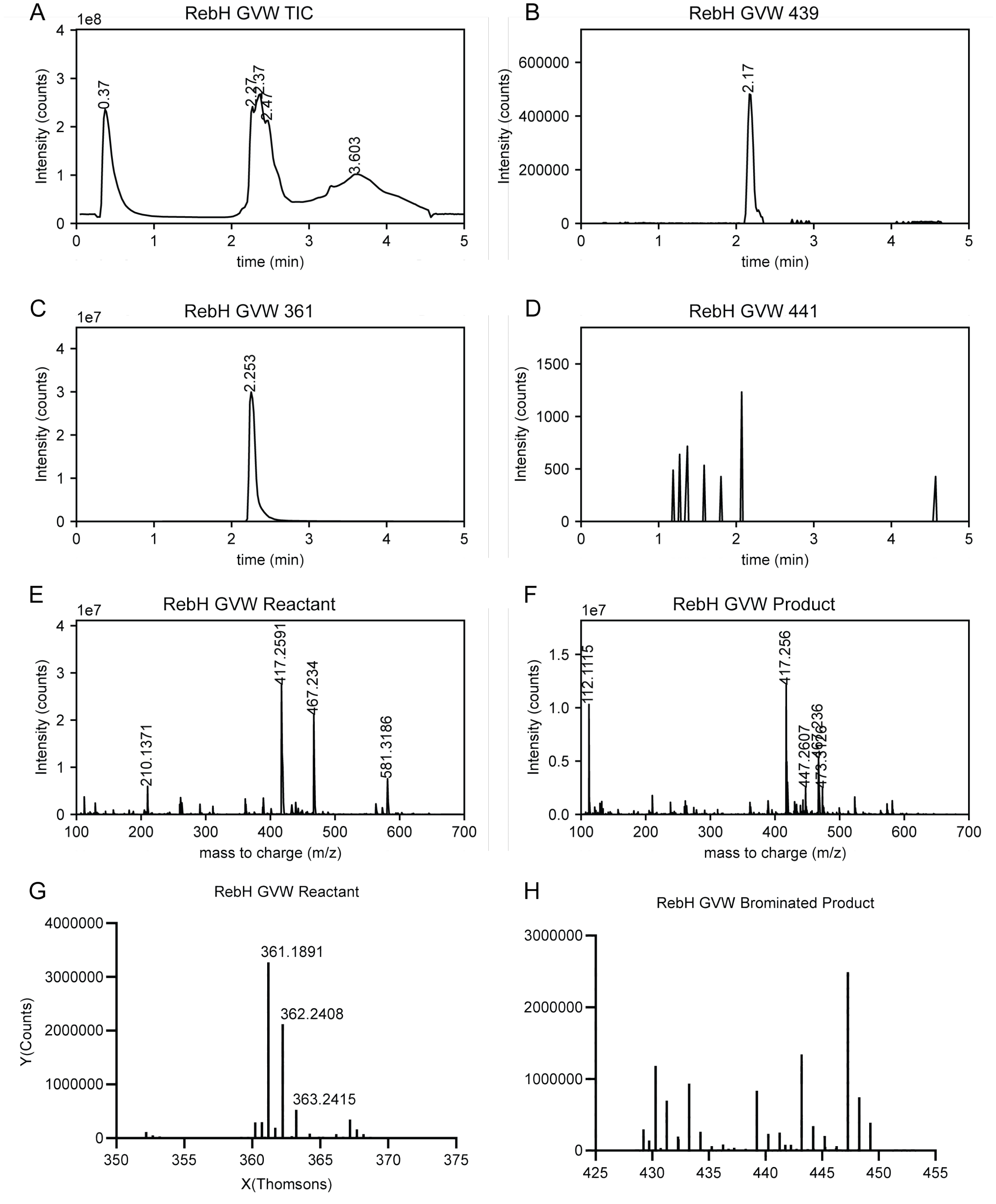
A
RebH GVW TIC
1e8
Intensity (counts)
time (min)
0.37
2.27
2.37
2.47
3.603
B
RebH GVW 439
Intensity (counts)
time (min)
2.17
C
RebH GVW 361
1e7
Intensity (counts)
time (min)
2.253
D
RebH GVW 441
Intensity (counts)
time (min)
E
RebH GVW Reactant
1e7
Intensity (counts)
mass to charge (m/z)
210.1371
417.2591
467.234
581.3186
F
RebH GVW Product
1e7
Intensity (counts)
mass to charge (m/z)
112.1115
417.256
447.2607
467.236
473.3126
G
RebH GVW Reactant
Y(Counts)
X(Thomsons)
361.1891
362.2408
363.2415
H
RebH GVW Brominated Product

#### RebH GYW

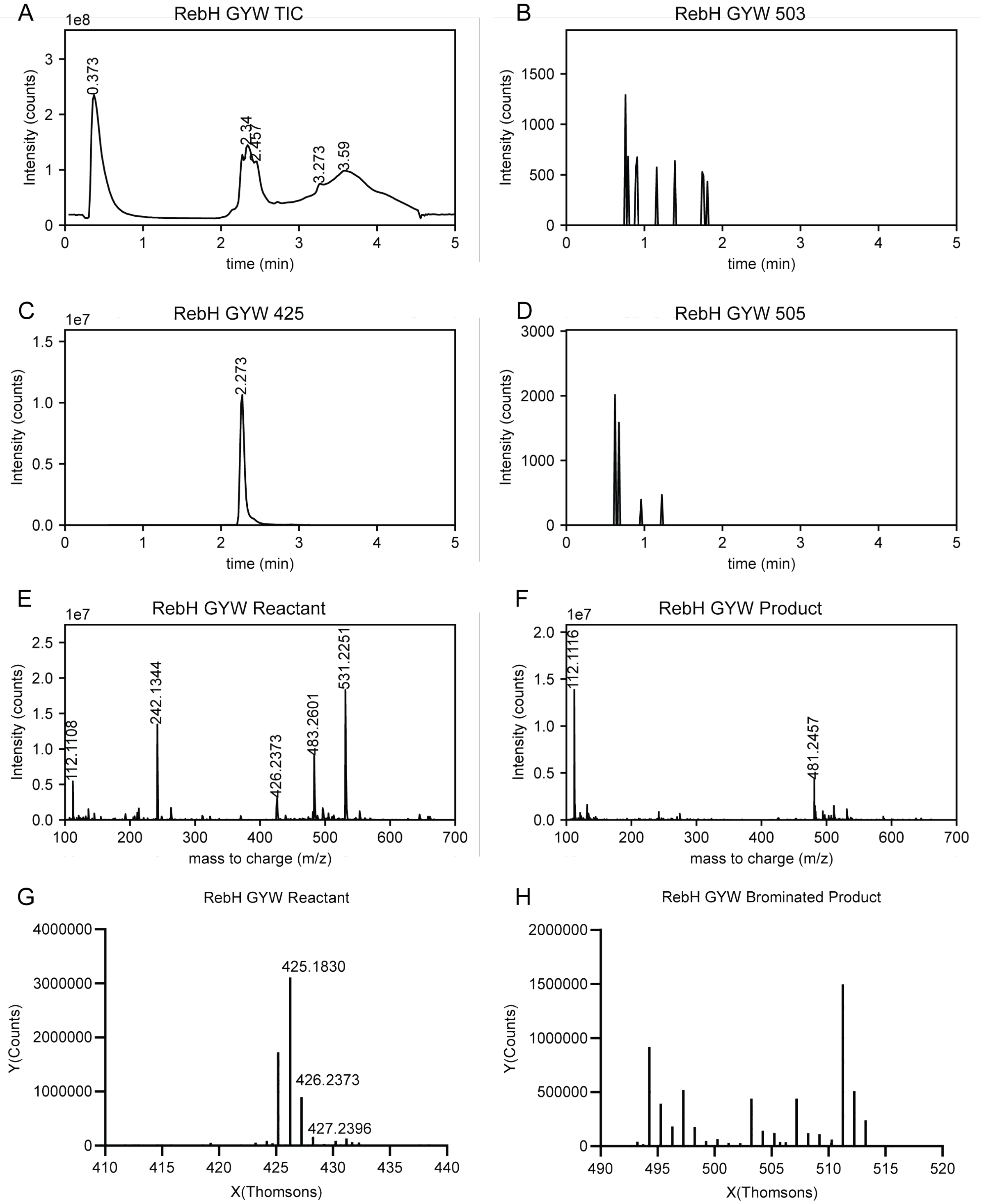
A
RebH GYW TIC
1e8
Intensity (counts)
time (min)
0.373
2.34
2.457
3.273
3.59
B
RebH GYW 503
Intensity (counts)
time (min)
C
RebH GYW 425
1e7
Intensity (counts)
time (min)
2.273
D
RebH GYW 505
Intensity (counts)
time (min)
E
RebH GYW Reactant
1e7
Intensity (counts)
mass to charge (m/z)
112.1108
242.1344
426.2373
483.2601
531.2251
F
RebH GYW Product
1e7
Intensity (counts)
mass to charge (m/z)
112.1116
481.2457
G
RebH GYW Reactant
Y(Counts)
X(Thomsons)
425.1830
426.2373
427.2396
H
RebH GYW Brominated Product
Y(Counts)
X(Thomsons)

#### 4V Brominated Tripeptide Mass Spectra

#### 4V WAG

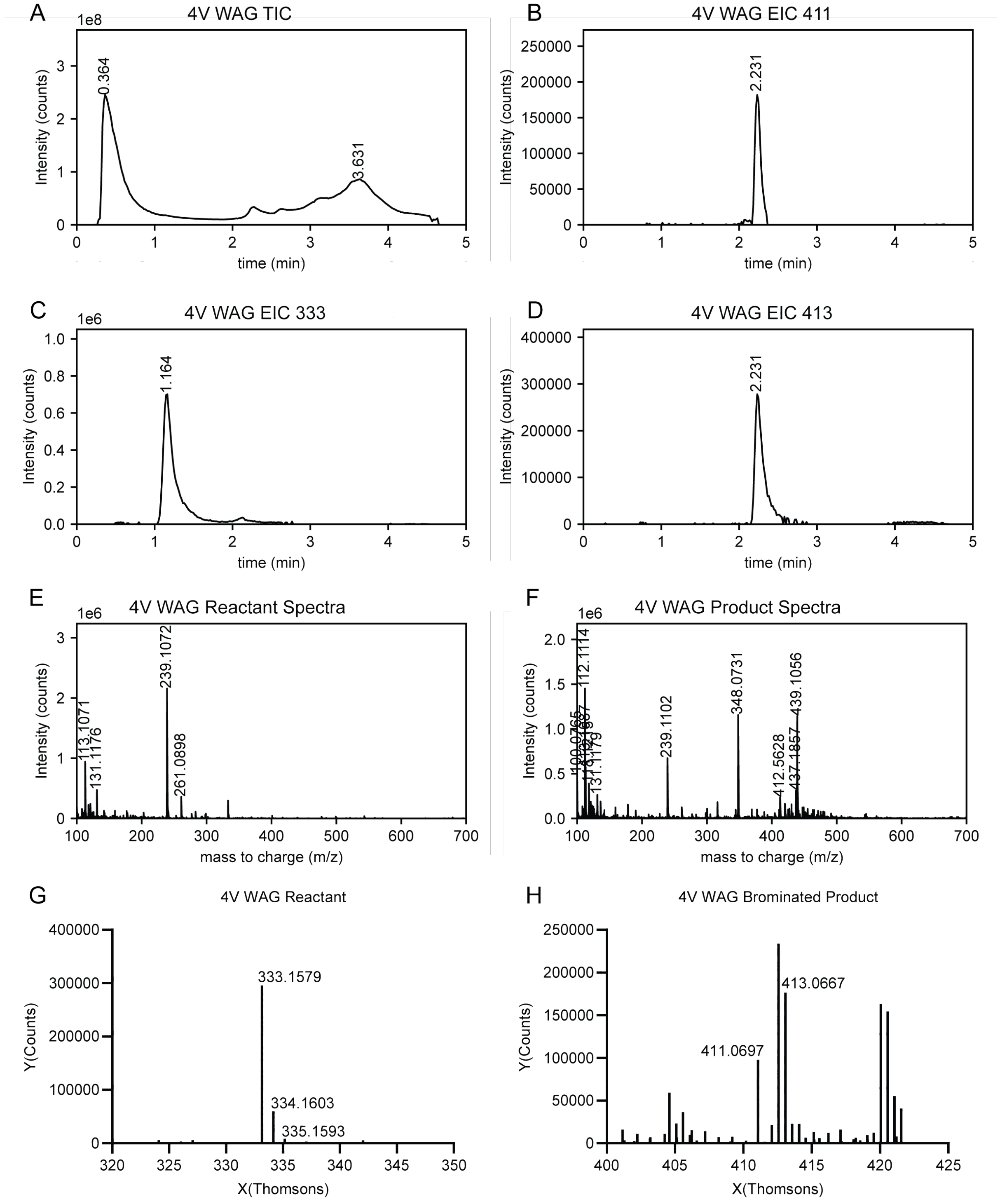
A
4V WAG TIC
1e8
Intensity (counts)
time (min)
0.364
3.631
B
4V WAG EIC 411
Intensity (counts)
time (min)
2.231
C
4V WAG EIC 333
1e6
Intensity (counts)
time (min)
1.164
D
4V WAG EIC 413
Intensity (counts)
time (min)
2.231
E
4V WAG Reactant Spectra
1e6
Intensity (counts)
mass to charge (m/z)
239.1072
113.1071
131.1176
261.0898
F
4V WAG Product Spectra
1e6
Intensity (counts)
mass to charge (m/z)
112.1114
100.0765
131.1179
239.1102
348.0731
439.1056
412.5628
437.1857
G
4V WAG Reactant
Y(Counts)
X(Thomsons)
333.1579
334.1603
335.1593
H
4V WAG Brominated Product
Y(Counts)
X(Thomsons)
413.0667
411.0697

#### 4V WCG

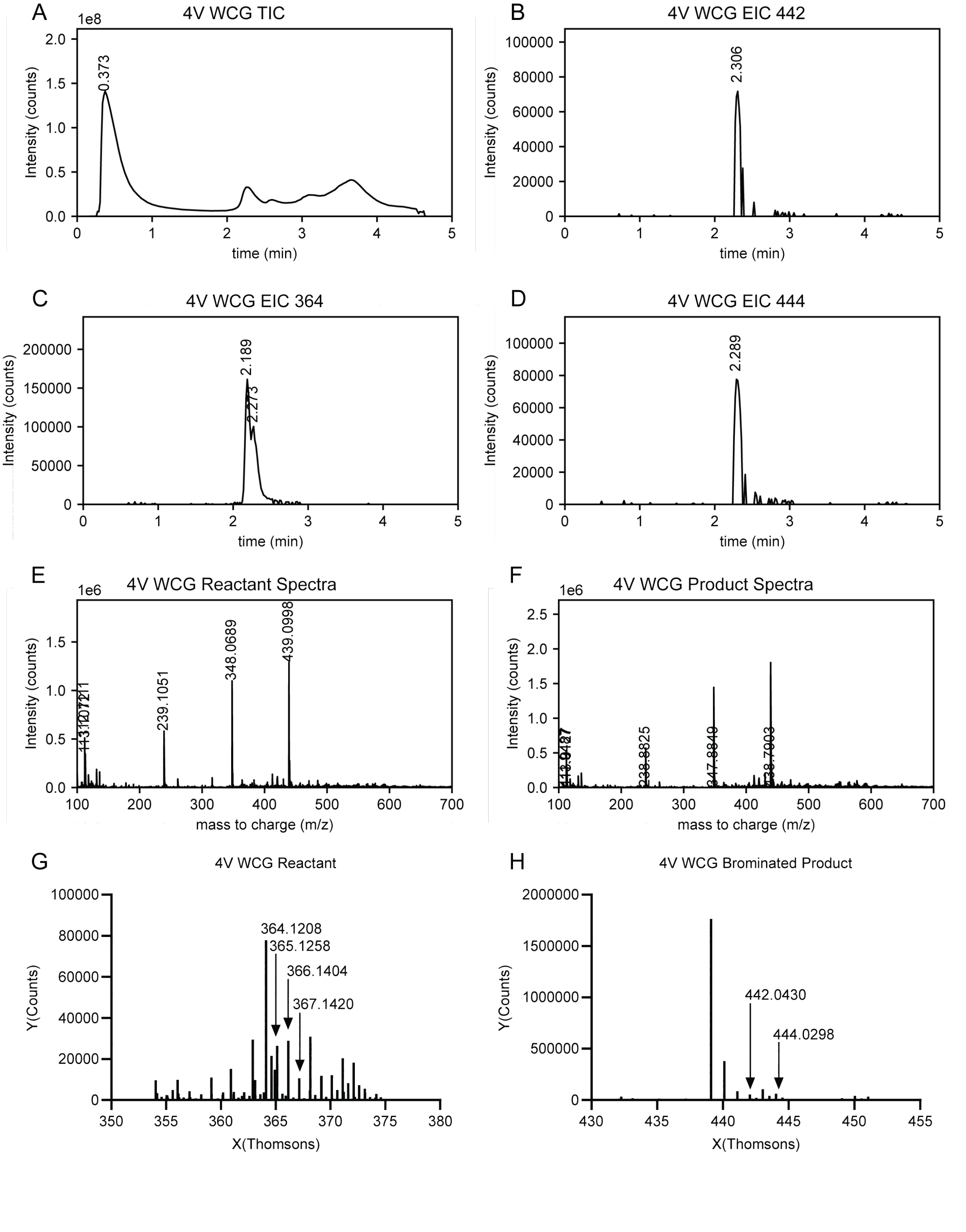

A
4V WCG TIC
1e8
Intensity (counts)
time (min)
0.373
B
4V WCG EIC 442
Intensity (counts)
time (min)
2.306
C
4V WCG EIC 364
Intensity (counts)
time (min)
2.189
2.273
D
4V WCG EIC 444
Intensity (counts)
time (min)
2.289
E
4V WCG Reactant Spectra
1e6
Intensity (counts)
mass to charge (m/z)
239.1051
348.0689
439.0998
F
4V WCG Product Spectra
1e6
Intensity (counts)
mass to charge (m/z)
238.8825
347.8849
438.7903
G
4V WCG Reactant
Y(Counts)
X(Thomsons)
364.1208
365.1258
366.1404
367.1420
H
4V WCG Brominated Product
Y(Counts)
X(Thomsons)
442.0430
444.0298

#### 4V WDG

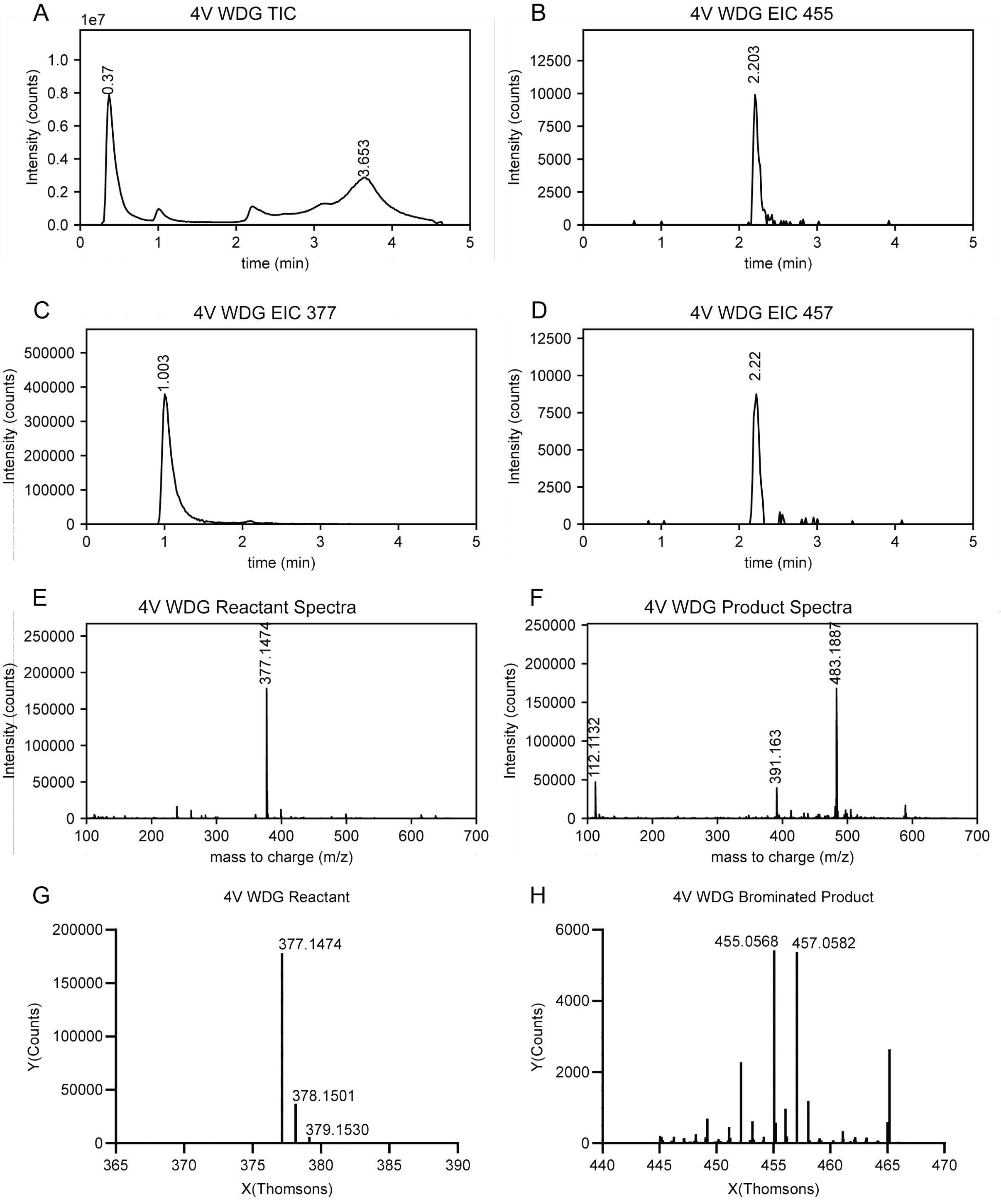
A
4V WDG TIC
1e7
Intensity (counts)
time (min)
0.37
3.653
B
4V WDG EIC 455
Intensity (counts)
time (min)
2.203
C
4V WDG EIC 377
Intensity (counts)
time (min)
1.003
D
4V WDG EIC 457
Intensity (counts)
time (min)
2.22
E
4V WDG Reactant Spectra
Intensity (counts)
mass to charge (m/z)
377.1474
F
4V WDG Product Spectra
Intensity (counts)
mass to charge (m/z)
112.1132
391.163
483.1887
G
4V WDG Reactant
Y(Counts)
X(Thomsons)
377.1474
378.1501
379.1530
H
4V WDG Brominated Product
Y(Counts)
X(Thomsons)
455.0568
457.0582

#### 4V WEG

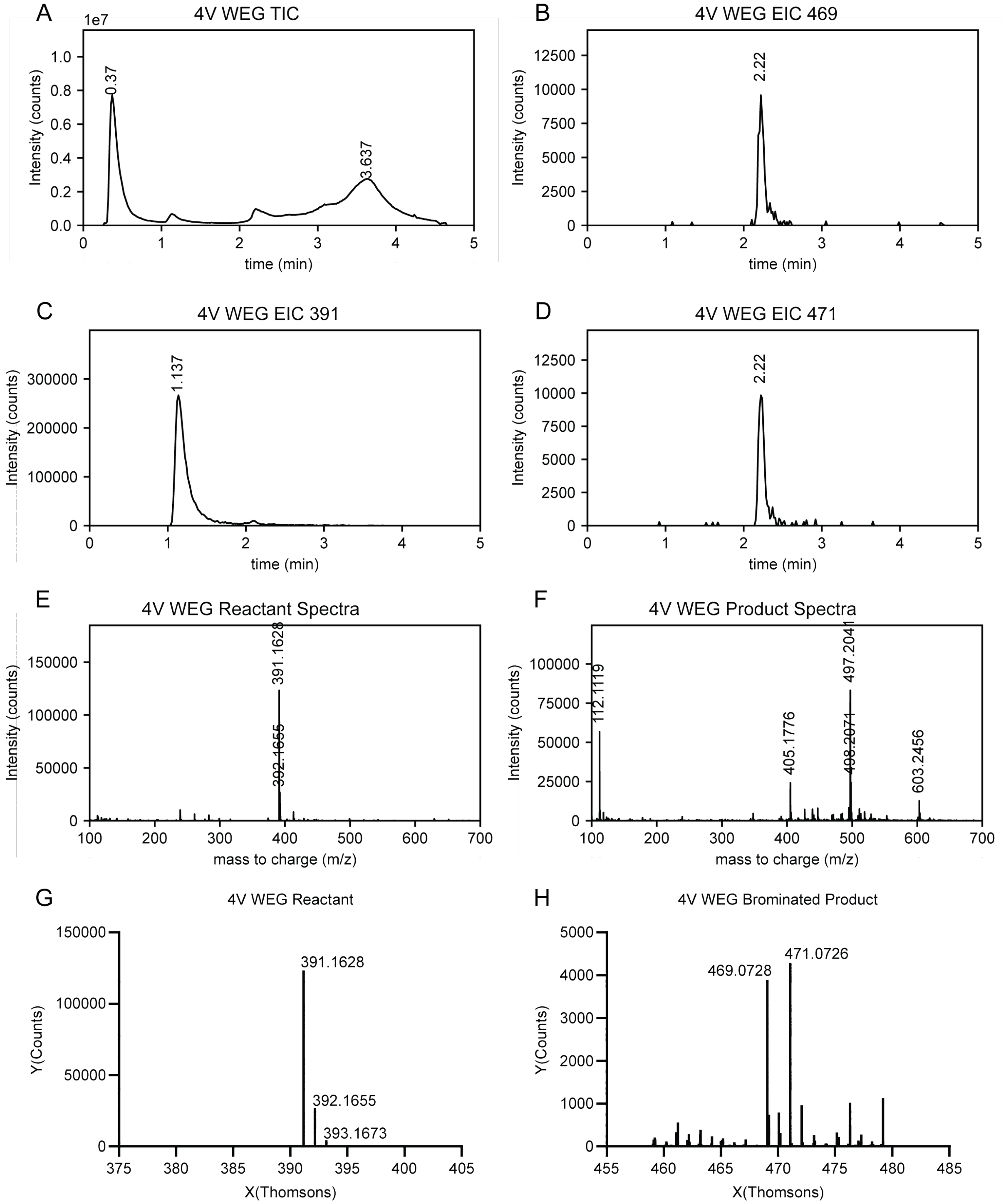

A
4V WEG TIC
1e7
Intensity (counts)
time (min)
0.37
3.637
B
4V WEG EIC 469
Intensity (counts)
time (min)
2.22
C
4V WEG EIC 391
Intensity (counts)
time (min)
1.137
D
4V WEG EIC 471
Intensity (counts)
time (min)
2.22
E
4V WEG Reactant Spectra
Intensity (counts)
mass to charge (m/z)
391.1628
392.1655
F
4V WEG Product Spectra
Intensity (counts)
mass to charge (m/z)
112.1119
405.1776
497.2041
498.2071
603.2456
G
4V WEG Reactant
Y(Counts)
X(Thomsons)
391.1628
392.1655
393.1673
H
4V WEG Brominated Product
Y(Counts)
X(Thomsons)
469.0728
471.0726

#### 4V WFG

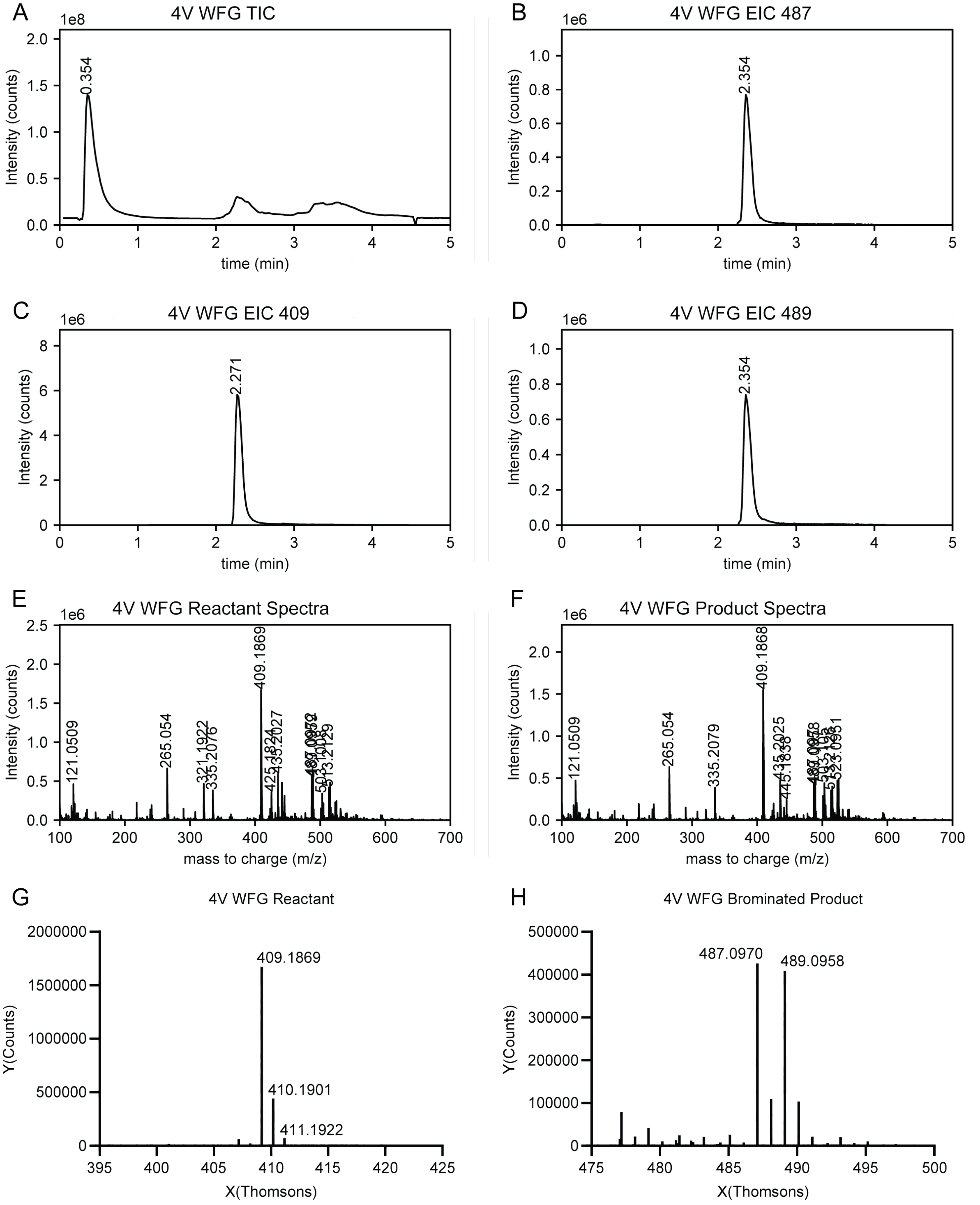

A
4V WFG TIC
1e8
Intensity (counts)
time (min)
0.354
B
4V WFG EIC 487
1e6
Intensity (counts)
time (min)
2.354
C
4V WFG EIC 409
1e6
Intensity (counts)
time (min)
2.271
D
4V WFG EIC 489
1e6
Intensity (counts)
time (min)
2.354
E
4V WFG Reactant Spectra
1e6
Intensity (counts)
mass to charge (m/z)
121.0509
265.054
321.1922
335.2076
409.1869
425.1824
435.2027
513.2129
F
4V WFG Product Spectra
1e6
Intensity (counts)
mass to charge (m/z)
121.0509
265.054
335.2079
409.1868
435.2025
445.1838
523.0951
G
4V WFG Reactant
Y(Counts)
X(Thomsons)
409.1869
410.1901
411.1922
H
4V WFG Brominated Product
Y(Counts)
X(Thomsons)
487.0970
489.0958

#### 4V WHG

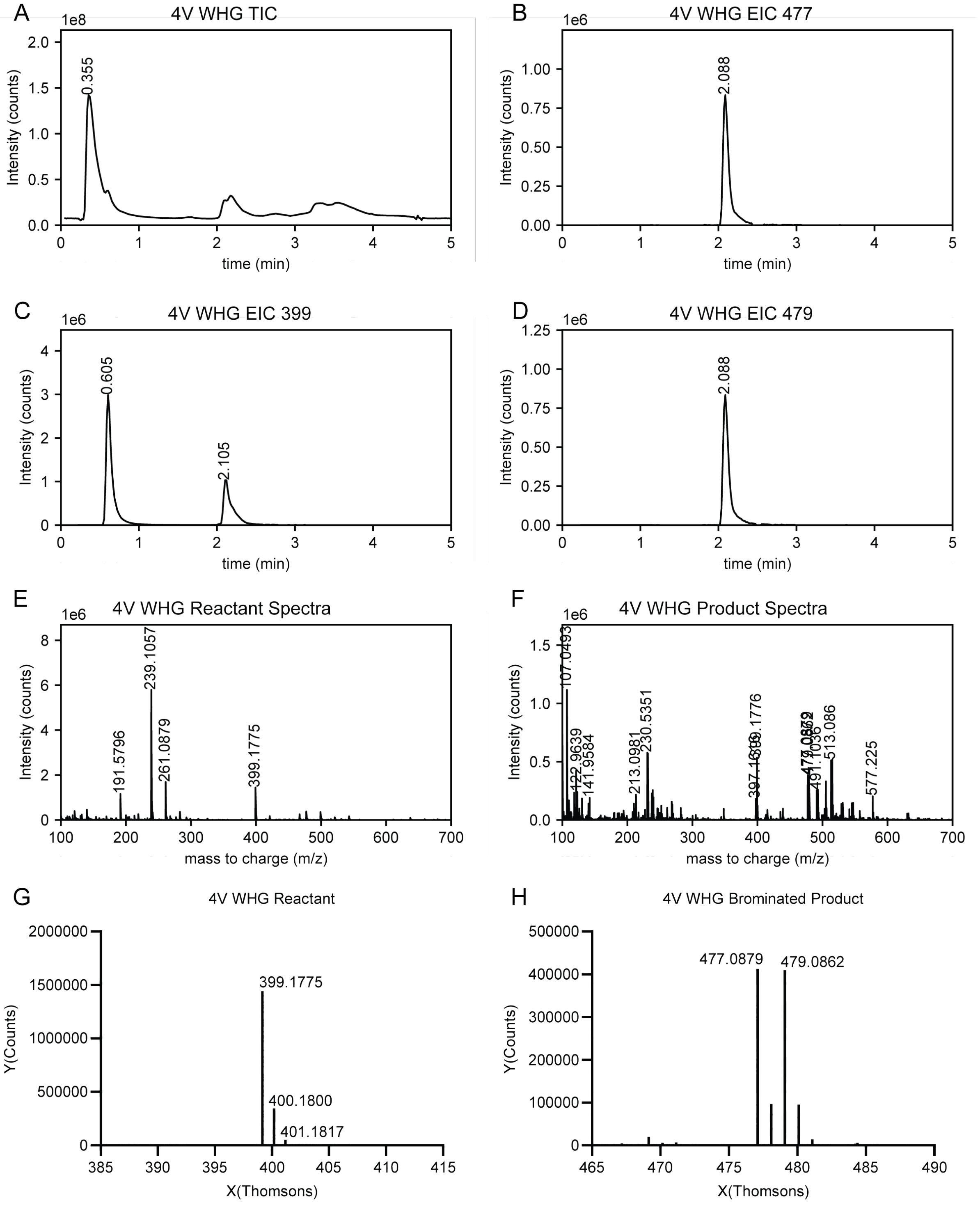
A
4V WHG TIC
1e8
Intensity (counts)
time (min)
0.355
B
4V WHG EIC 477
1e6
Intensity (counts)
time (min)
2.088
C
4V WHG EIC 399
1e6
Intensity (counts)
time (min)
0.605
2.105
D
4V WHG EIC 479
1e6
Intensity (counts)
time (min)
2.088
E
4V WHG Reactant Spectra
1e6
Intensity (counts)
mass to charge (m/z)
239.1057
191.5796
261.0879
399.1775
F
4V WHG Product Spectra
1e6
Intensity (counts)
mass to charge (m/z)
107.0493
122.9639
141.9584
213.0981
230.5351
491.1036
513.086
577.225
G
4V WHG Reactant
Y(Counts)
X(Thomsons)
399.1775
400.1800
401.1817
H
4V WHG Brominated Product
Y(Counts)
X(Thomsons)
477.0879
479.0862

#### 4V WKG

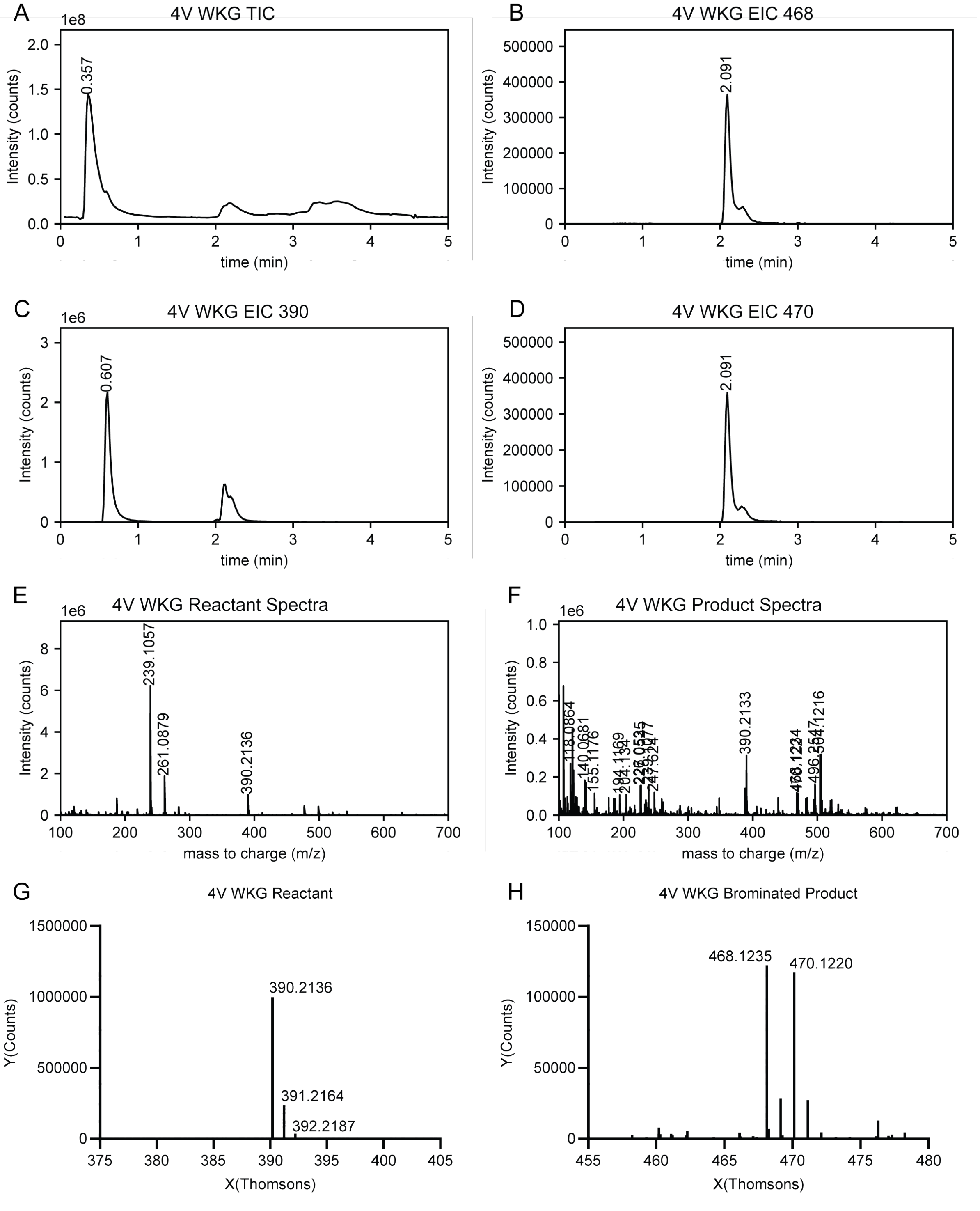

A
4V WKG TIC
1e8
Intensity (counts)
time (min)
0.357
B
4V WKG EIC 468
Intensity (counts)
time (min)
2.091
C
4V WKG EIC 390
1e6
Intensity (counts)
time (min)
0.607
D
4V WKG EIC 470
Intensity (counts)
time (min)
2.091
E
4V WKG Reactant Spectra
1e6
Intensity (counts)
mass to charge (m/z)
239.1057
261.0879
390.2136
F
4V WKG Product Spectra
1e6
Intensity (counts)
mass to charge (m/z)
118.0864
140.0681
155.1176
390.2133
G
4V WKG Reactant
Y(Counts)
X(Thomsons)
390.2136
391.2164
392.2187
H
4V WKG Brominated Product
Y(Counts)
X(Thomsons)
468.1235
470.1220

#### 4V WLG

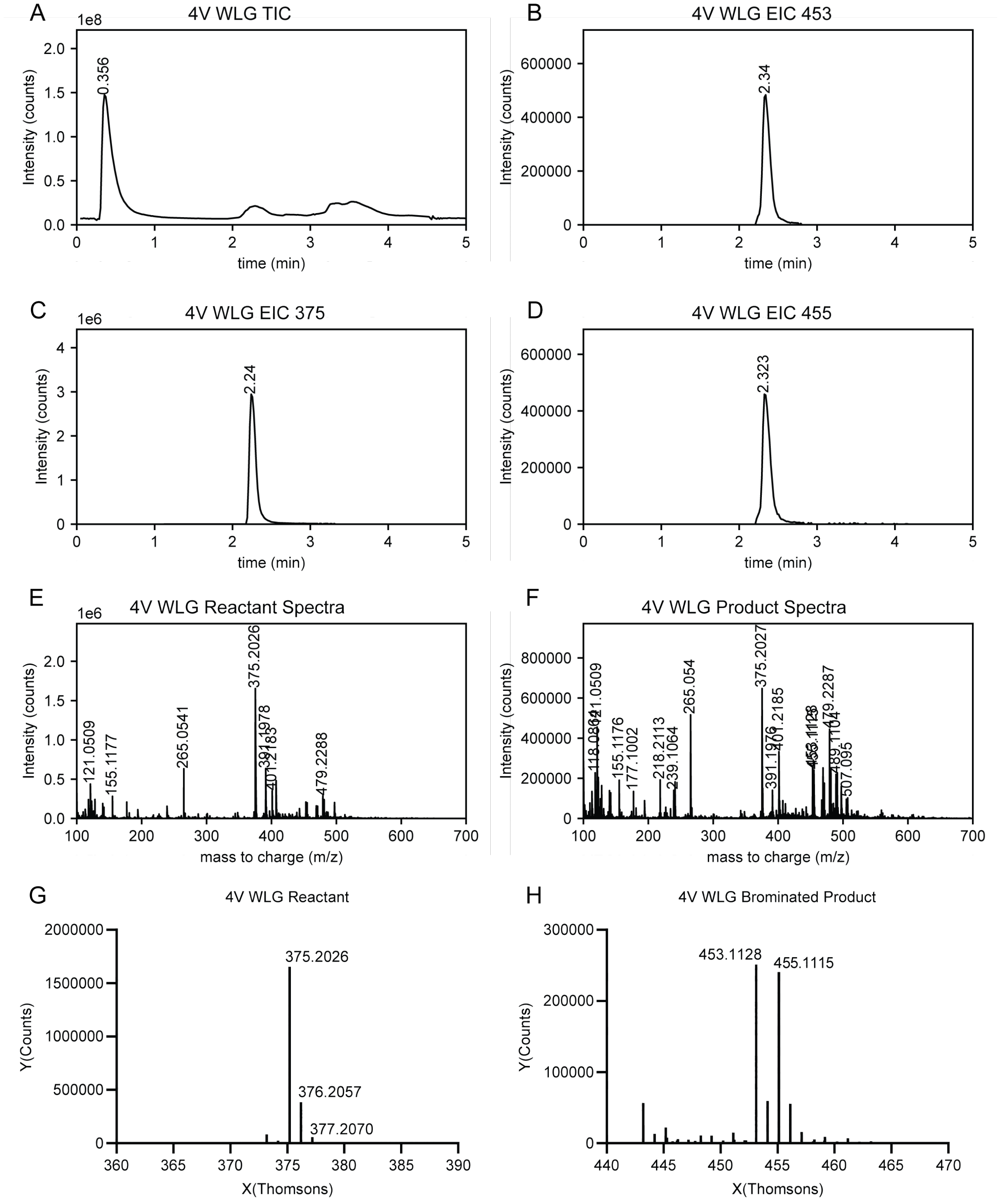
A
4V WLG TIC
1e8
Intensity (counts)
time (min)
0.356
B
4V WLG EIC 453
Intensity (counts)
time (min)
2.34
C
4V WLG EIC 375
1e6
Intensity (counts)
time (min)
2.24
D
4V WLG EIC 455
Intensity (counts)
time (min)
2.323
E
4V WLG Reactant Spectra
1e6
Intensity (counts)
mass to charge (m/z)
121.0509
155.1177
265.0541
375.2026
391.1978
401.2183
479.2288
F
4V WLG Product Spectra
Intensity (counts)
mass to charge (m/z)
155.1176
177.1002
218.2113
239.1064
265.054
375.2027
391.1976
401.2185
479.2287
489.1104
507.095
G
4V WLG Reactant
Y(Counts)
X(Thomsons)
375.2026
376.2057
377.2070
H
4V WLG Brominated Product
Y(Counts)
X(Thomsons)
453.1128
455.1115

#### 4V WMG

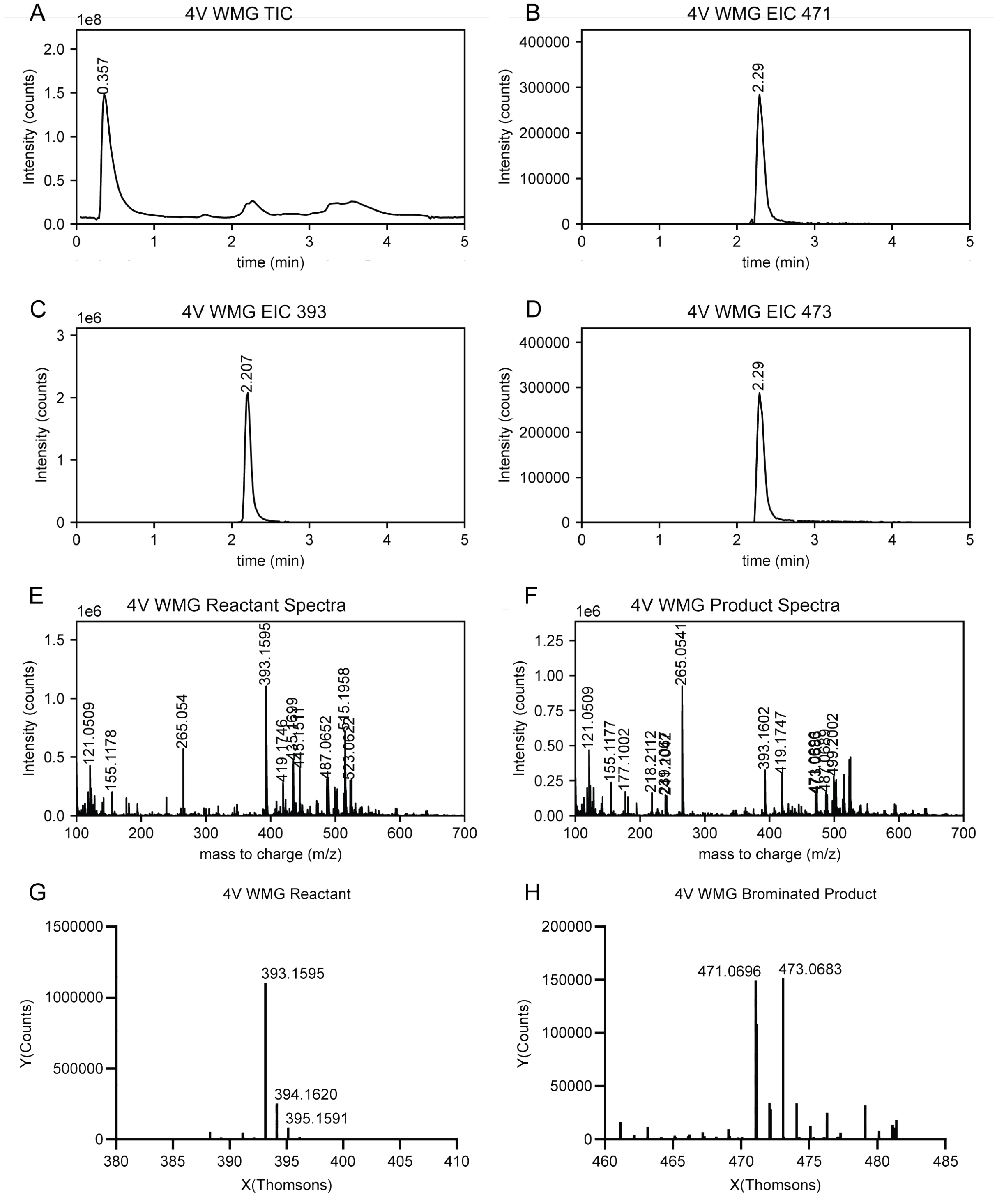

A
4V WMG TIC
1e8
0.357
Intensity (counts)
time (min)
B
4V WMG EIC 471
2.29
Intensity (counts)
time (min)
C
4V WMG EIC 393
1e6
2.207
Intensity (counts)
time (min)
D
4V WMG EIC 473
2.29
Intensity (counts)
time (min)
E
4V WMG Reactant Spectra
1e6
121.0509
155.1178
265.054
393.1595
419.1746
487.0652
515.1958
523.0622
Intensity (counts)
mass to charge (m/z)
F
4V WMG Product Spectra
1e6
121.0509
155.1177
177.1002
218.2112
265.0541
393.1602
419.1747
499.2002
Intensity (counts)
mass to charge (m/z)
G
4V WMG Reactant
393.1595
394.1620
395.1591
Y(Counts)
X(Thomsons)
H
4V WMG Brominated Product
471.0696
473.0683
Y(Counts)
X(Thomsons)

#### 4V WNG

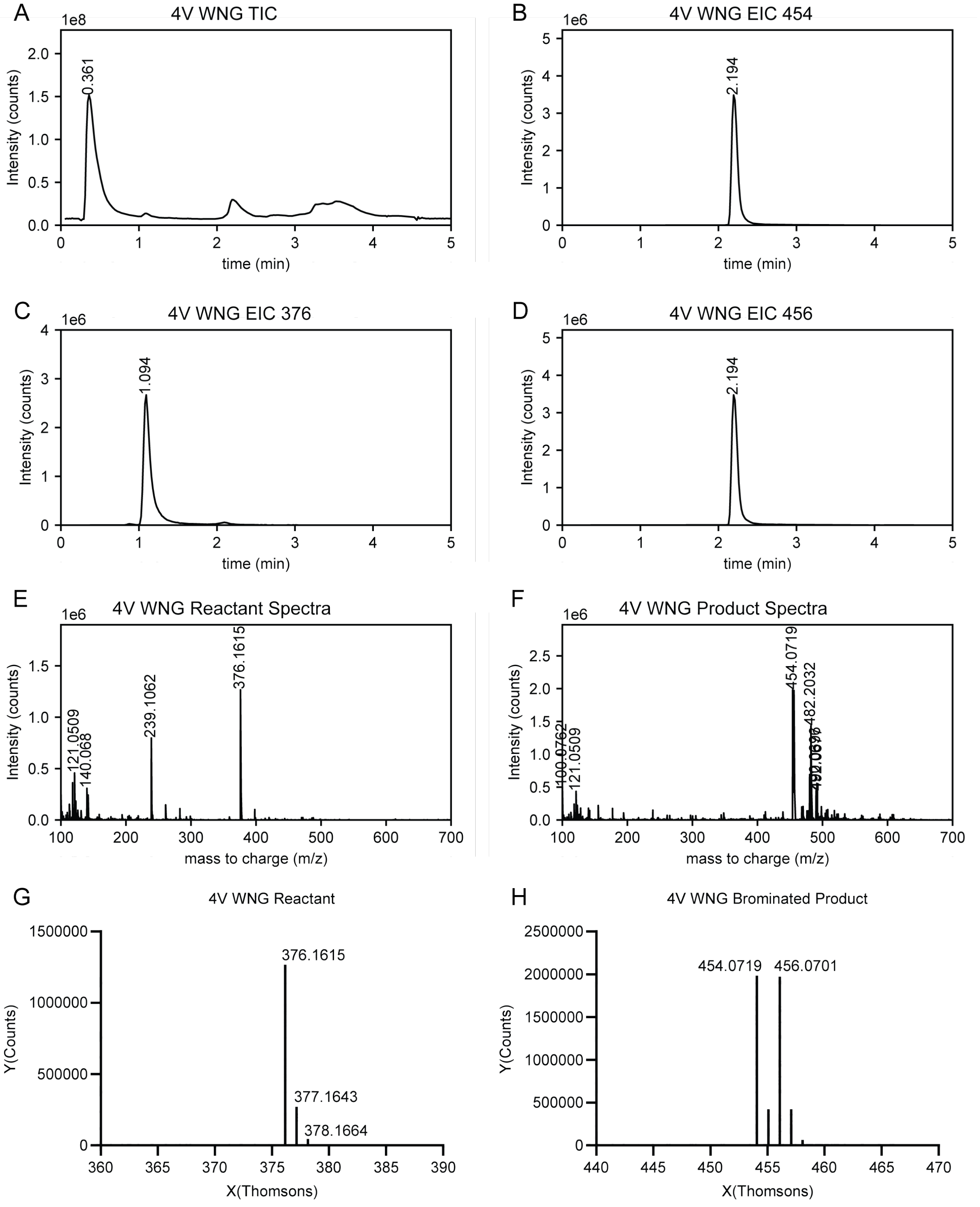
A
4V WNG TIC
1e8
0.361
Intensity (counts)
time (min)
B
4V WNG EIC 454
1e6
2.194
C
4V WNG EIC 376
1.094
D
4V WNG EIC 456
2.194
E
4V WNG Reactant Spectra
376.1615
239.1062
121.0509
140.068
mass to charge (m/z)
F
4V WNG Product Spectra
454.0719
482.2032
121.0509
G
4V WNG Reactant
376.1615
377.1643
378.1664
Y(Counts)
X(Thomsons)
H
4V WNG Brominated Product
454.0719
456.0701

**4V WPG**

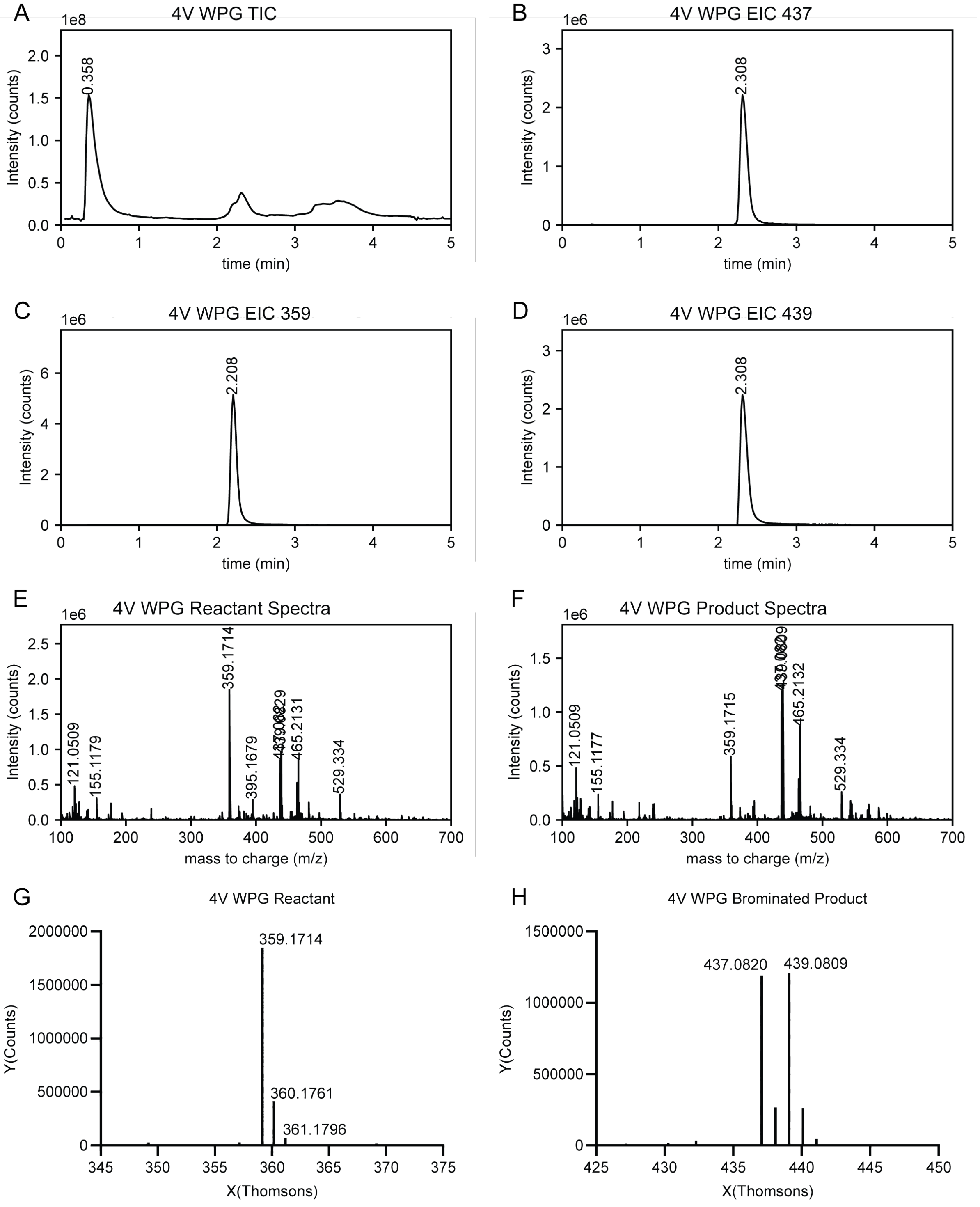
A
4V WPG TIC
1e8
Intensity (counts)
time (min)
0.358
B
4V WPG EIC 437
1e6
Intensity (counts)
time (min)
2.308
C
4V WPG EIC 359
1e6
Intensity (counts)
time (min)
2.208
D
4V WPG EIC 439
1e6
Intensity (counts)
time (min)
2.308
E
4V WPG Reactant Spectra
1e6
Intensity (counts)
mass to charge (m/z)
121.0509
155.1179
359.1714
395.1679
465.2131
529.334
F
4V WPG Product Spectra
1e6
Intensity (counts)
mass to charge (m/z)
121.0509
155.1177
359.1715
465.2132
529.334
G
4V WPG Reactant
Y(Counts)
X(Thomsons)
359.1714
360.1761
361.1796
H
4V WPG Brominated Product
Y(Counts)
X(Thomsons)
437.0820
439.0809

#### 4V WQG

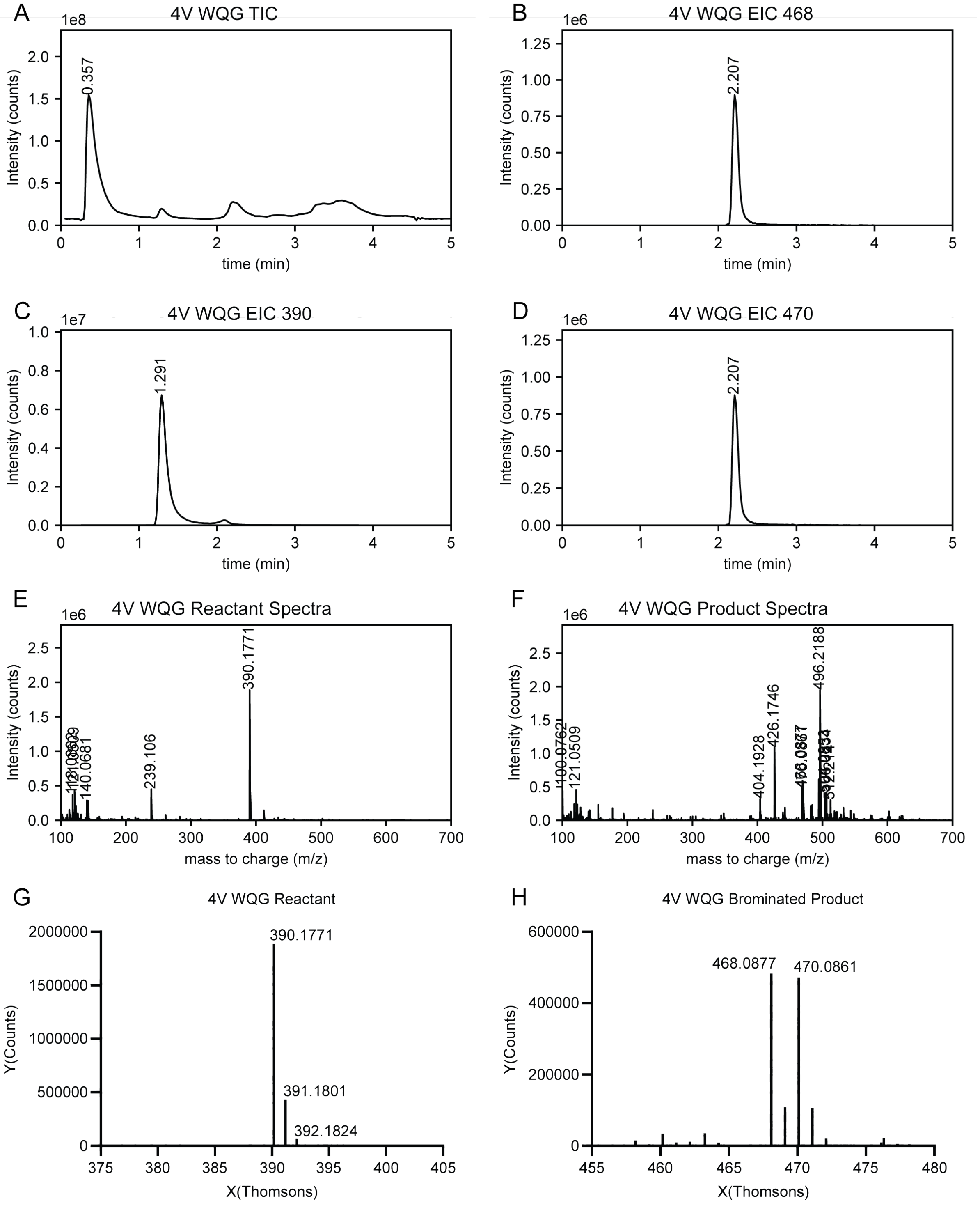

A
4V WQG TIC
1e8
0.357
Intensity (counts)
time (min)
B
4V WQG EIC 468
1e6
2.207
Intensity (counts)
time (min)
C
4V WQG EIC 390
1e7
1.291
Intensity (counts)
time (min)
D
4V WQG EIC 470
1e6
2.207
Intensity (counts)
time (min)
E
4V WQG Reactant Spectra
1e6
390.1771
239.106
140.0681
Intensity (counts)
mass to charge (m/z)
F
4V WQG Product Spectra
1e6
496.2188
426.1746
404.1928
121.0509
Intensity (counts)
mass to charge (m/z)
G
4V WQG Reactant
390.1771
391.1801
392.1824
Y(Counts)
X(Thomsons)
H
4V WQG Brominated Product
468.0877
470.0861
Y(Counts)
X(Thomsons)

#### 4V WRG

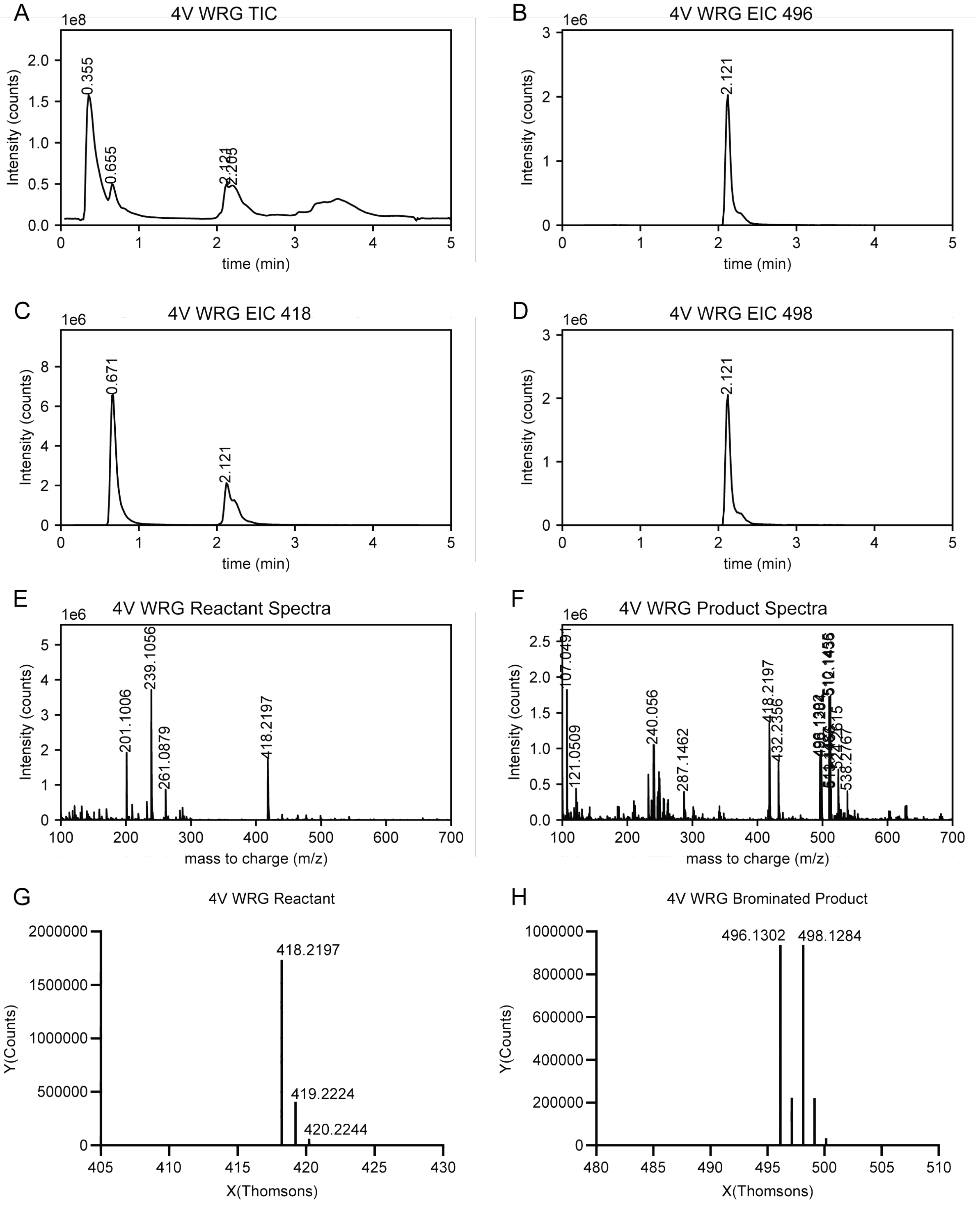

A
4V WRG TIC
1e8
Intensity (counts)
time (min)
0.355
0.655
2.121
2.265
B
4V WRG EIC 496
1e6
2.121
C
4V WRG EIC 418
0.671
2.121
D
4V WRG EIC 498
2.121
E
4V WRG Reactant Spectra
mass to charge (m/z)
201.1006
239.1056
261.0879
418.2197
F
4V WRG Product Spectra
107.0491
121.0509
240.056
287.1462
418.2197
432.2356
538.2767
G
4V WRG Reactant
Y(Counts)
X(Thomsons)
418.2197
419.2224
420.2244
H
4V WRG Brominated Product
496.1302
498.1284

#### 4V WTG

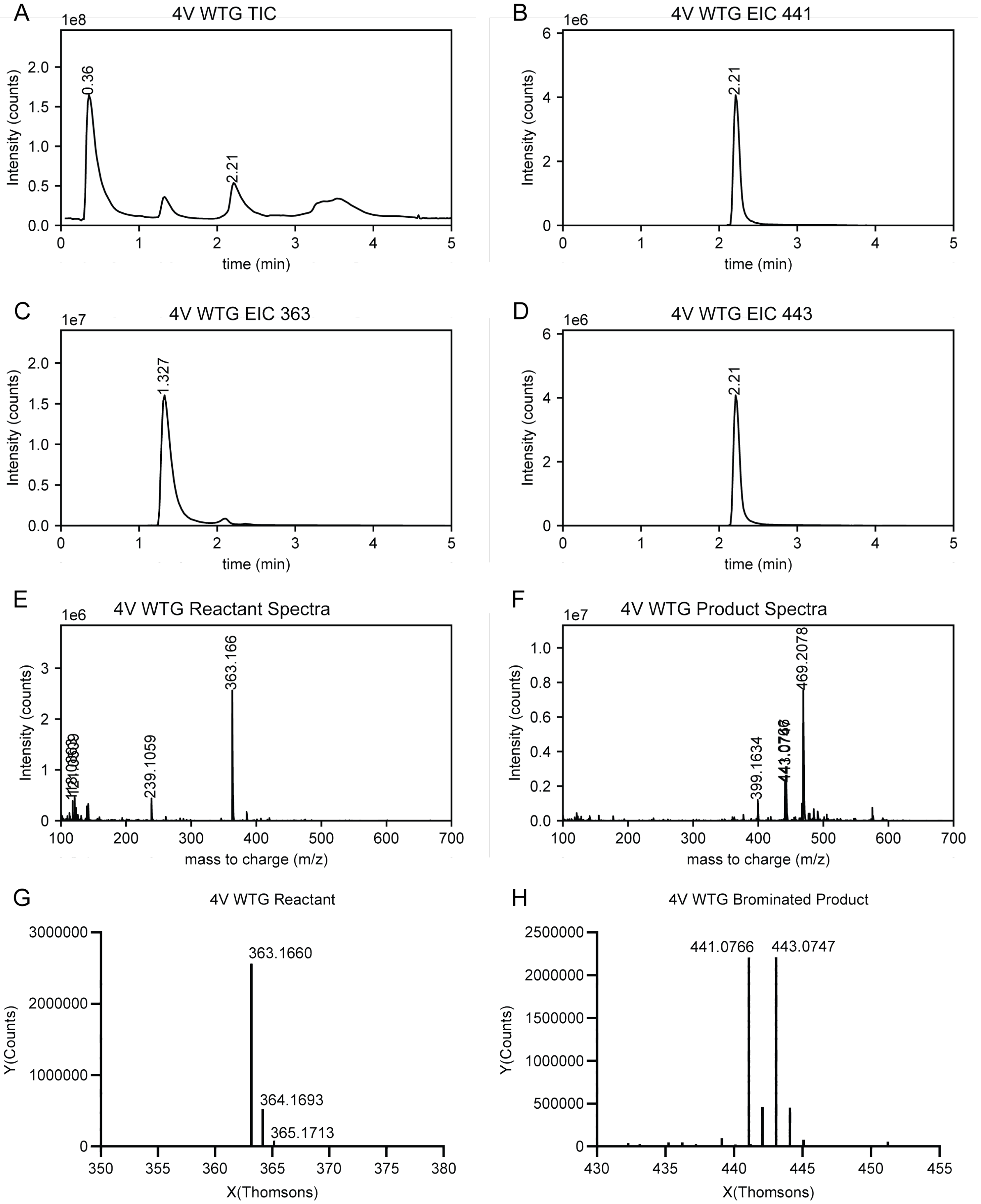

A
4V WTG TIC
1e8
Intensity (counts)
time (min)
0.36
2.21
B
4V WTG EIC 441
1e6
2.21
C
4V WTG EIC 363
1e7
1.327
D
4V WTG EIC 443
1e6
2.21
E
4V WTG Reactant Spectra
1e6
mass to charge (m/z)
363.166
239.1059
F
4V WTG Product Spectra
1e7
469.2078
399.1634
G
4V WTG Reactant
Y(Counts)
X(Thomsons)
363.1660
364.1693
365.1713
H
4V WTG Brominated Product
441.0766
443.0747

#### 4V WVG

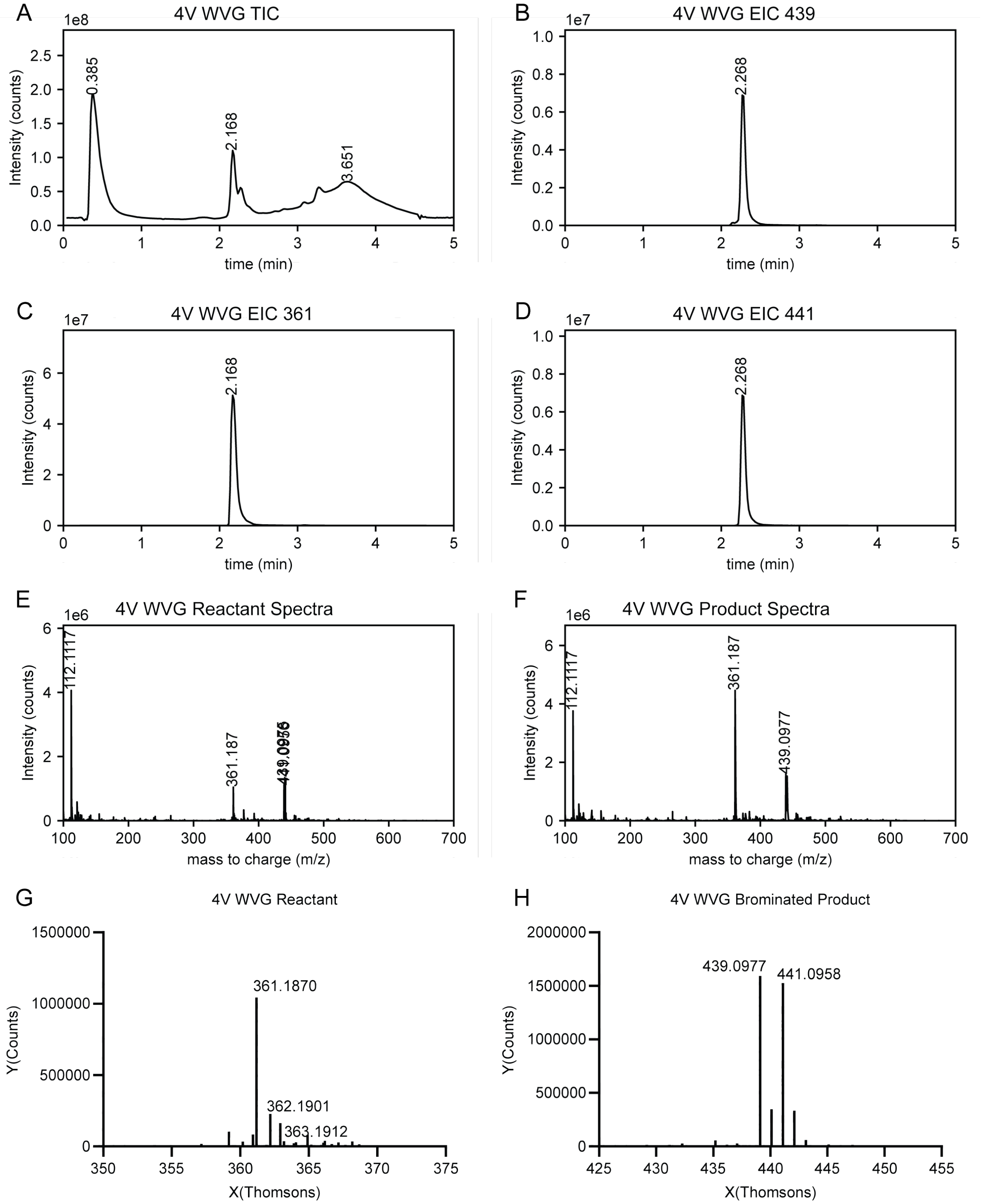
A
4V WVG TIC
1e8
Intensity (counts)
time (min)
0.385
2.168
3.651
B
4V WVG EIC 439
1e7
Intensity (counts)
time (min)
2.268
C
4V WVG EIC 361
1e7
Intensity (counts)
time (min)
2.168
D
4V WVG EIC 441
1e7
Intensity (counts)
time (min)
2.268
E
4V WVG Reactant Spectra
1e6
Intensity (counts)
mass to charge (m/z)
112.1117
361.187
F
4V WVG Product Spectra
1e6
Intensity (counts)
mass to charge (m/z)
112.1117
361.187
439.0977
G
4V WVG Reactant
Y(Counts)
X(Thomsons)
361.1870
362.1901
363.1912
H
4V WVG Brominated Product
Y(Counts)
X(Thomsons)
439.0977
441.0958

#### 4V WYG

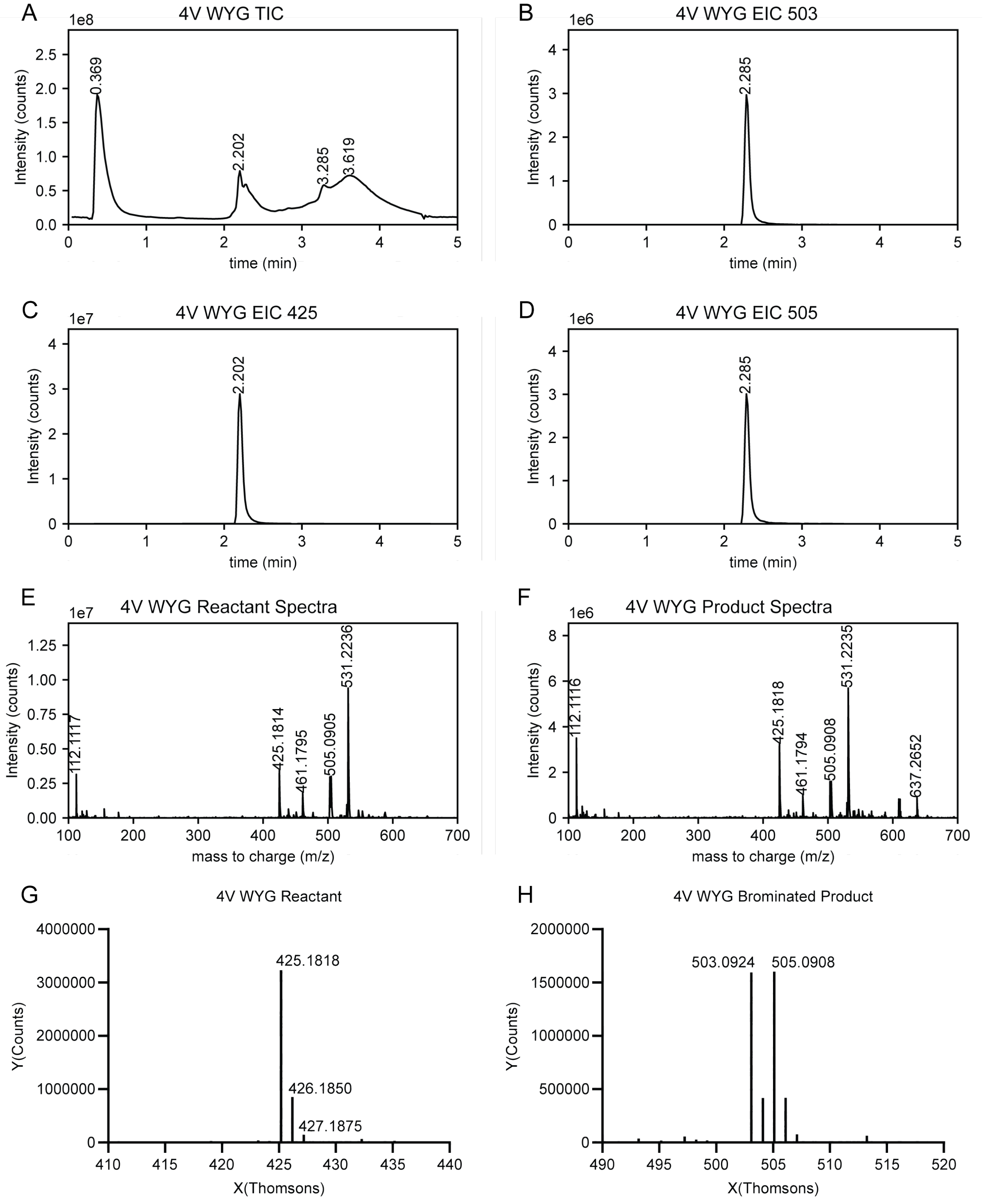

A
4V WYG TIC
1e8
Intensity (counts)
time (min)
0.369
2.202
3.285
3.619
B
4V WYG EIC 503
1e6
2.285
C
4V WYG EIC 425
1e7
2.202
D
4V WYG EIC 505
1e6
2.285
E
4V WYG Reactant Spectra
1e7
mass to charge (m/z)
112.1117
425.1814
461.1795
505.0905
531.2236
F
4V WYG Product Spectra
1e6
112.1116
425.1818
461.1794
505.0908
531.2235
637.2652
G
4V WYG Reactant
Y(Counts)
X(Thomsons)
425.1818
426.1850
427.1875
H
4V WYG Brominated Product
503.0924
505.0908

#### 4V WGA

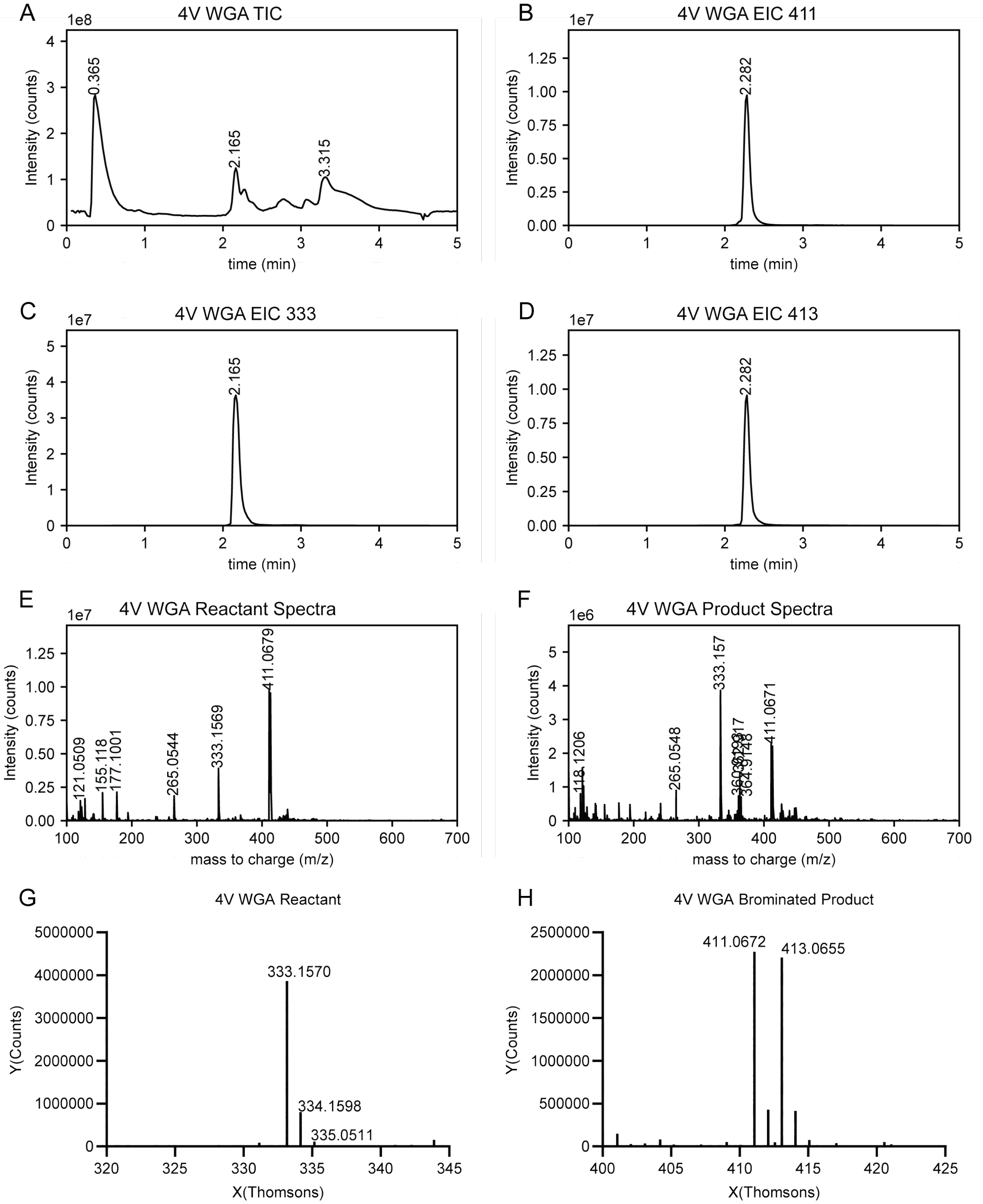
A
4V WGA TIC
1e8
Intensity (counts)
time (min)
0.365
2.165
3.315
B
4V WGA EIC 411
1e7
2.282
C
4V WGA EIC 333
2.165
D
4V WGA EIC 413
2.282
E
4V WGA Reactant Spectra
mass to charge (m/z)
121.0509
155.118
177.1001
265.0544
333.1569
411.0679
F
4V WGA Product Spectra
1e6
118.1206
265.0548
333.157
364.9148
411.0671
G
4V WGA Reactant
Y(Counts)
X(Thomsons)
333.1570
334.1598
335.0511
H
4V WGA Brominated Product
411.0672
413.0655

#### 4V WGC

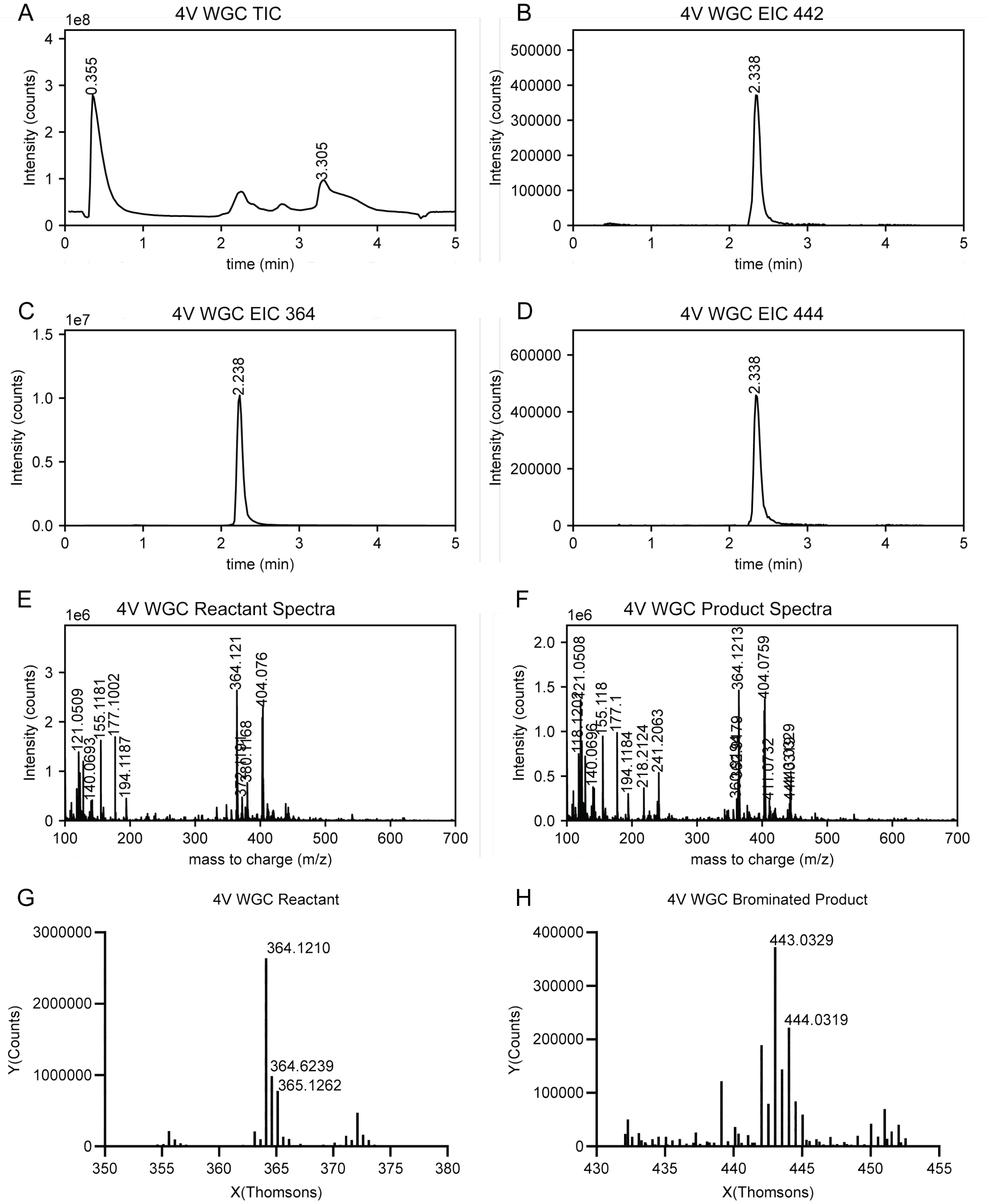

A
4V WGC TIC
1e8
Intensity (counts)
time (min)
0.355
3.305
B
4V WGC EIC 442
Intensity (counts)
time (min)
2.338
C
4V WGC EIC 364
1e7
Intensity (counts)
time (min)
2.238
D
4V WGC EIC 444
Intensity (counts)
time (min)
2.338
E
4V WGC Reactant Spectra
1e6
Intensity (counts)
mass to charge (m/z)
121.0509
140.0693
155.1181
177.1002
194.1187
364.121
380.1168
404.076
F
4V WGC Product Spectra
1e6
Intensity (counts)
mass to charge (m/z)
121.0508
140.0696
155.118
177.1
194.1184
218.2124
241.2063
364.1213
404.0759
411.0732
G
4V WGC Reactant
Y(Counts)
X(Thomsons)
364.1210
364.6239
365.1262
H
4V WGC Brominated Product
Y(Counts)
X(Thomsons)
443.0329
444.0319

#### 4V WGD

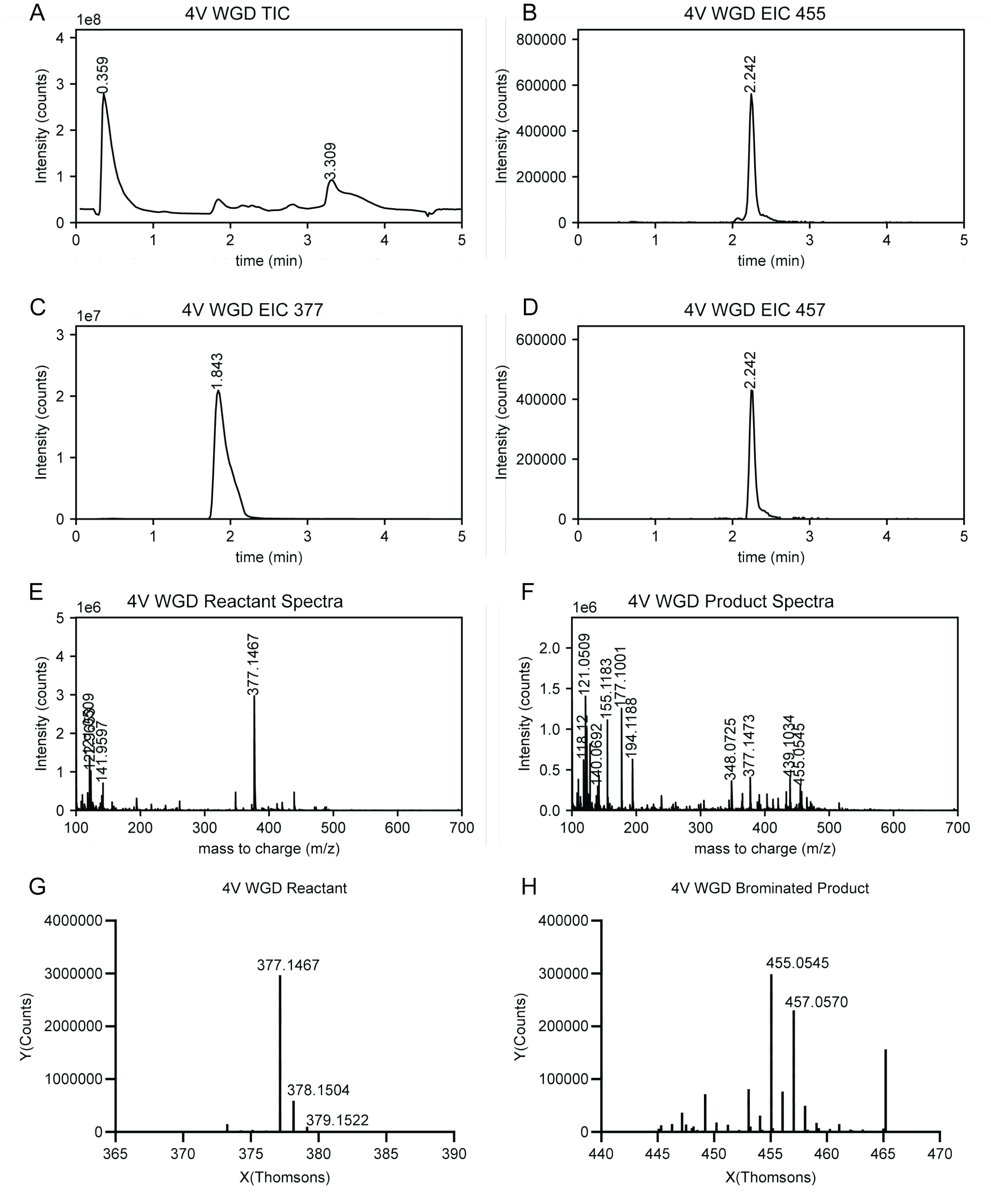
A
4V WGD TIC
0.359
3.309
Intensity (counts)
time (min)
B
4V WGD EIC 455
2.242
C
4V WGD EIC 377
1.843
D
4V WGD EIC 457
2.242
E
4V WGD Reactant Spectra
377.1467
141.9597
mass to charge (m/z)
F
4V WGD Product Spectra
121.0509
155.1183
177.1001
194.1188
118.12
140.0692
348.0725
377.1473
439.1034
455.0545
G
4V WGD Reactant
377.1467
378.1504
379.1522
Y(Counts)
X(Thomsons)
H
4V WGD Brominated Product
455.0545
457.0570

#### 4V WGE

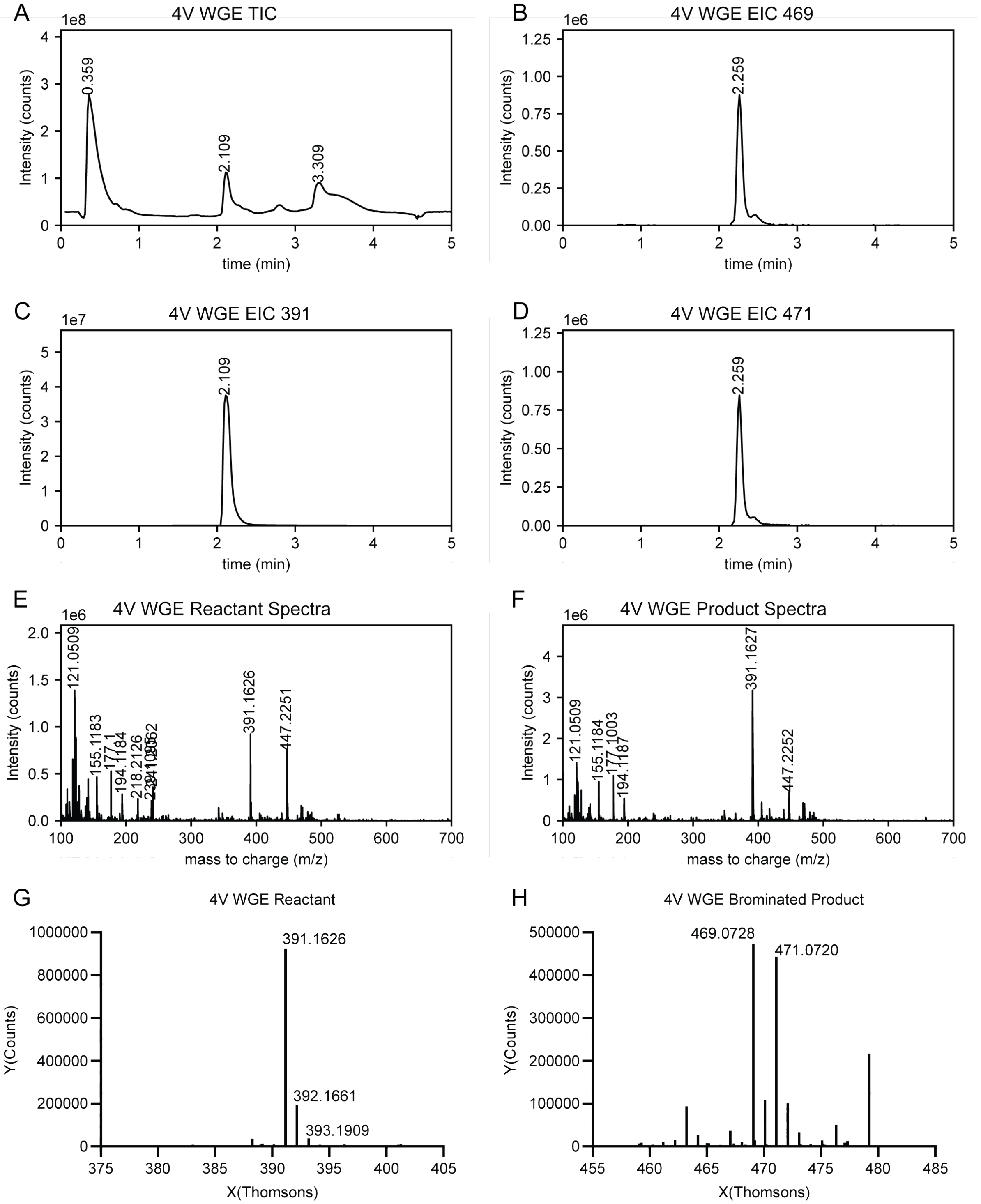

A
4V WGE TIC
B
4V WGE EIC 469
C
4V WGE EIC 391
D
4V WGE EIC 471
E
4V WGE Reactant Spectra
F
4V WGE Product Spectra
G
4V WGE Reactant
H
4V WGE Brominated Product
Intensity (counts)
time (min)
mass to charge (m/z)
Y(Counts)
X(Thomsons)
0.359
2.109
3.309
2.259
121.0509
155.1183
391.1626
447.2251
391.1627
447.2252
155.1184
177.1003
194.1187
392.1661
393.1909
469.0728
471.0720

#### 4V WGF

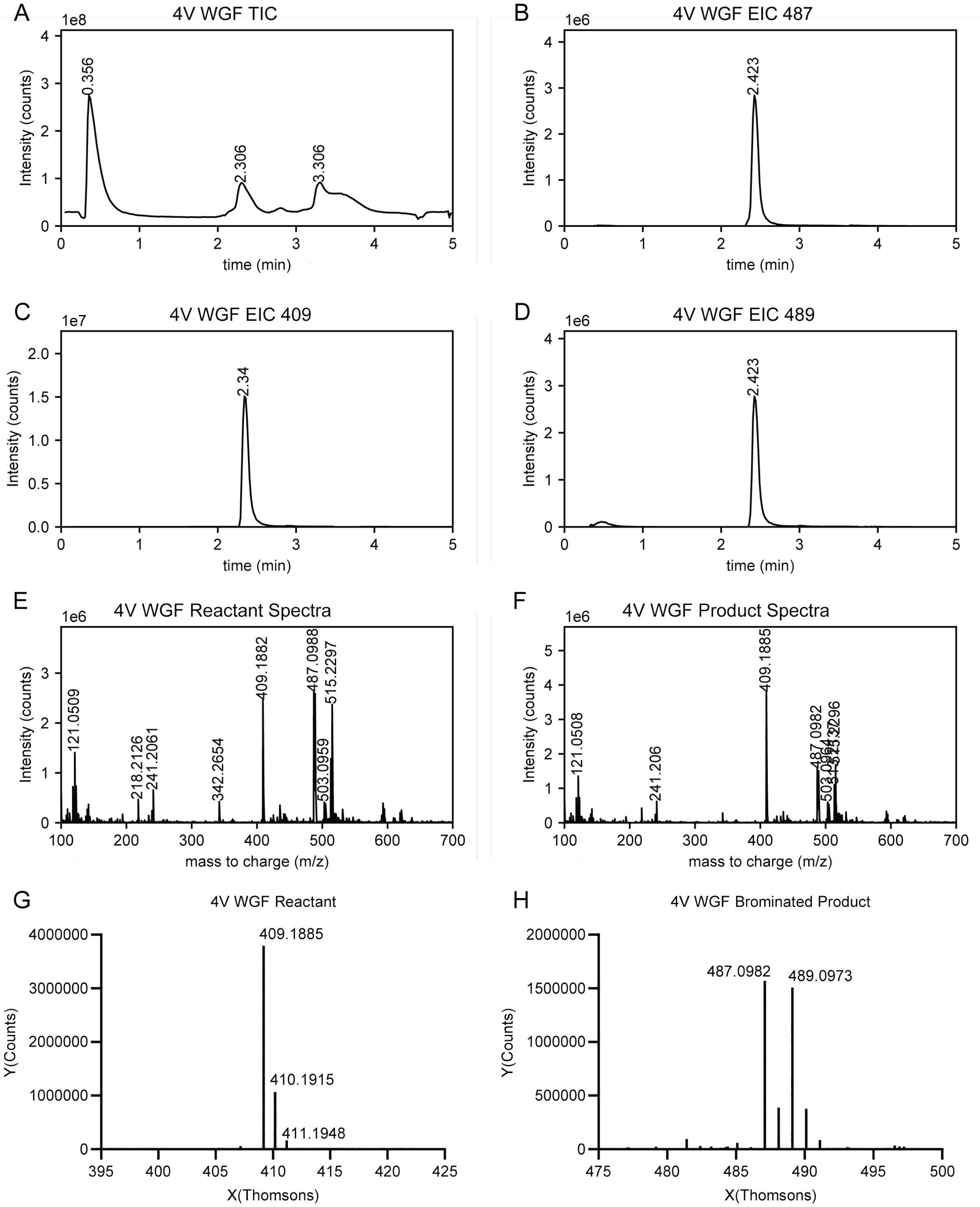
A
4V WGF TIC
1e8
Intensity (counts)
time (min)
0.356
2.306
3.306
B
4V WGF EIC 487
1e6
Intensity (counts)
time (min)
2.423
C
4V WGF EIC 409
1e7
Intensity (counts)
time (min)
2.34
D
4V WGF EIC 489
1e6
Intensity (counts)
time (min)
2.423
E
4V WGF Reactant Spectra
1e6
Intensity (counts)
mass to charge (m/z)
121.0509
218.2126
241.2061
342.2654
409.1882
487.0988
503.0959
515.2297
F
4V WGF Product Spectra
1e6
Intensity (counts)
mass to charge (m/z)
121.0508
241.206
409.1885
487.0982
G
4V WGF Reactant
Y(Counts)
X(Thomsons)
409.1885
410.1915
411.1948
H
4V WGF Brominated Product
Y(Counts)
X(Thomsons)
487.0982
489.0973

#### 4V WGH

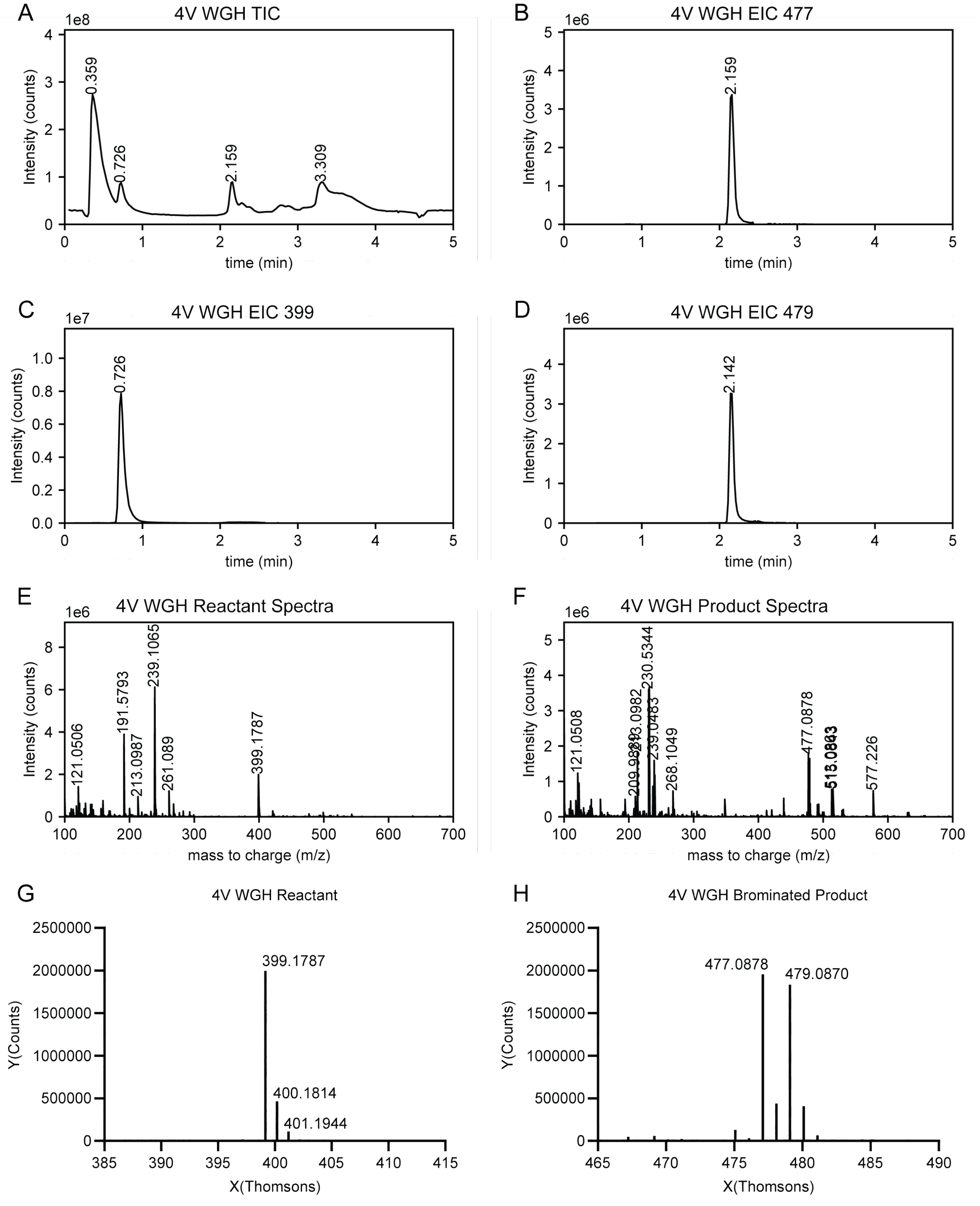
A
4V WGH TIC
1e8
Intensity (counts)
time (min)
0.359
0.726
2.159
3.309
B
4V WGH EIC 477
1e6
Intensity (counts)
time (min)
2.159
C
4V WGH EIC 399
1e7
Intensity (counts)
time (min)
0.726
D
4V WGH EIC 479
1e6
Intensity (counts)
time (min)
2.142
E
4V WGH Reactant Spectra
1e6
Intensity (counts)
mass to charge (m/z)
121.0506
191.5793
213.0987
239.1065
261.089
399.1787
F
4V WGH Product Spectra
1e6
Intensity (counts)
mass to charge (m/z)
121.0508
230.5344
239.0483
268.1049
477.0878
577.226
G
4V WGH Reactant
Y(Counts)
X(Thomsons)
399.1787
400.1814
401.1944
H
4V WGH Brominated Product
Y(Counts)
X(Thomsons)
477.0878
479.0870

#### 4V WGK

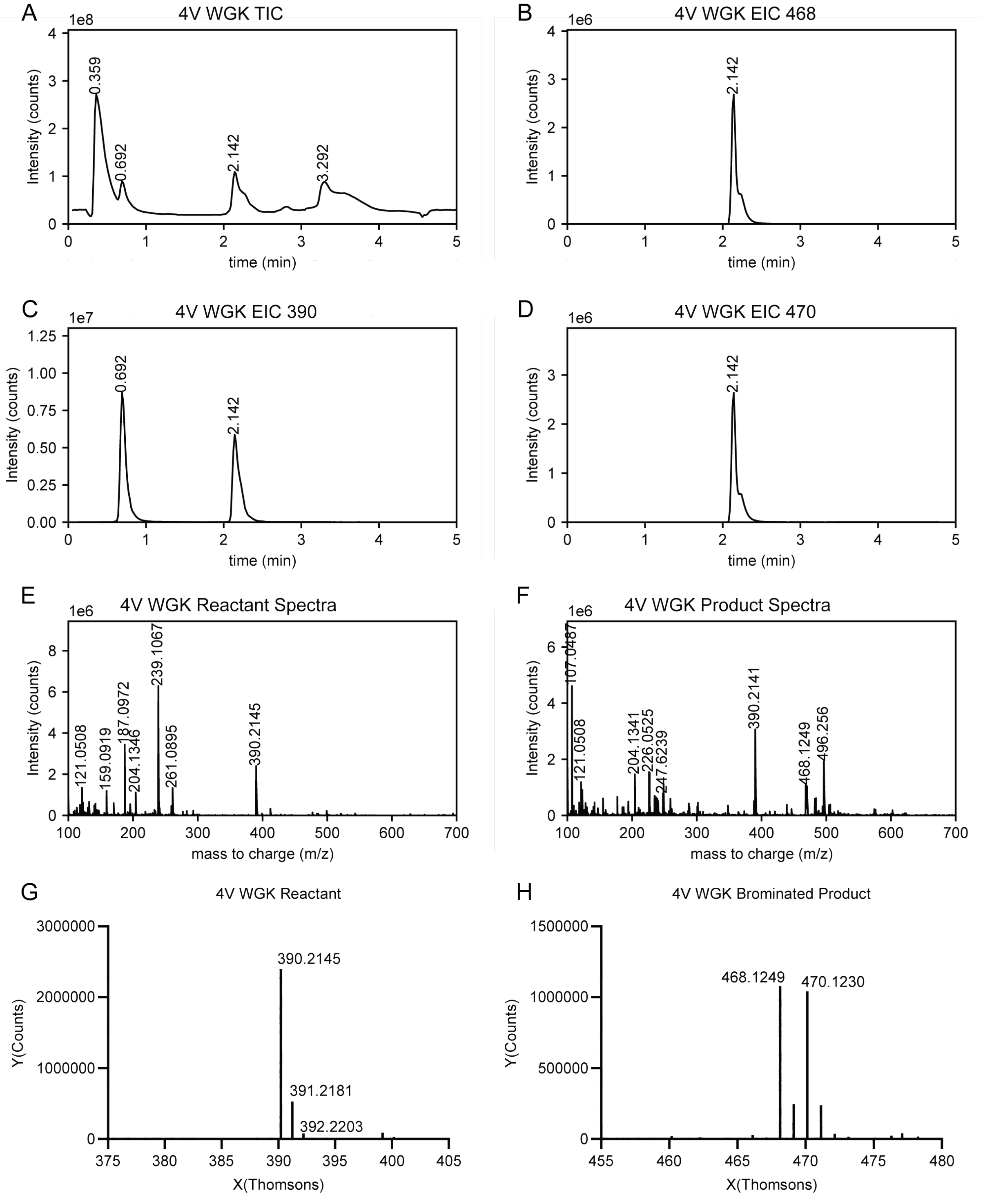

A
4V WGK TIC
1e8
Intensity (counts)
time (min)
0.359
0.692
2.142
3.292
B
4V WGK EIC 468
1e6
Intensity (counts)
time (min)
2.142
C
4V WGK EIC 390
1e7
Intensity (counts)
time (min)
0.692
2.142
D
4V WGK EIC 470
1e6
Intensity (counts)
time (min)
2.142
E
4V WGK Reactant Spectra
1e6
Intensity (counts)
mass to charge (m/z)
121.0508
159.0919
187.0972
204.1346
239.1067
261.0895
390.2145
F
4V WGK Product Spectra
1e6
Intensity (counts)
mass to charge (m/z)
107.0487
121.0508
204.1341
226.0525
247.6239
390.2141
468.1249
496.256
G
4V WGK Reactant
Y(Counts)
X(Thomsons)
390.2145
391.2181
392.2203
H
4V WGK Brominated Product
Y(Counts)
X(Thomsons)
468.1249
470.1230

#### 4V WGL

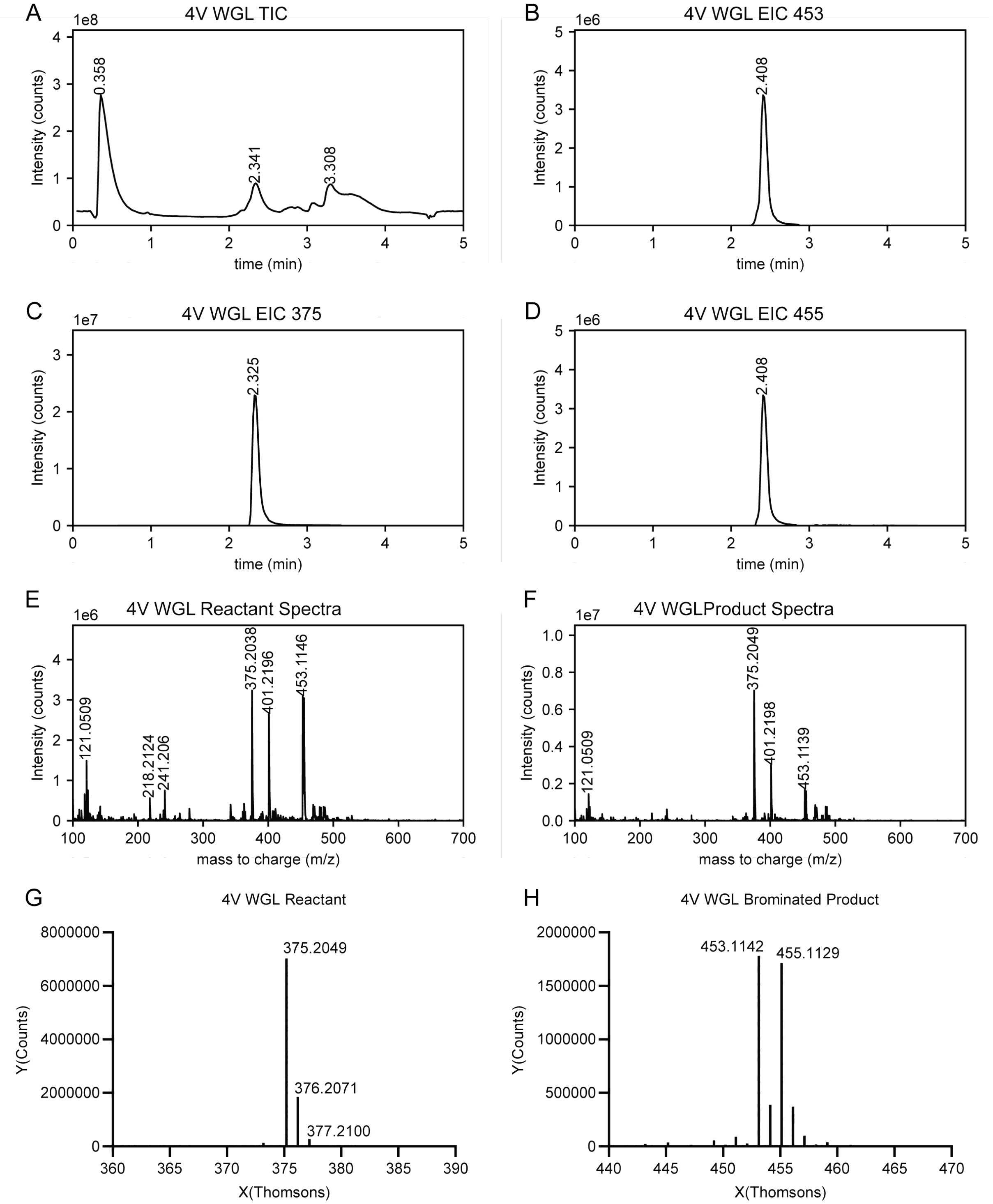
A
4V WGL TIC
1e8
0.358
2.341
3.308
Intensity (counts)
time (min)
B
4V WGL EIC 453
1e6
2.408
Intensity (counts)
time (min)
C
4V WGL EIC 375
1e7
2.325
Intensity (counts)
time (min)
D
4V WGL EIC 455
1e6
2.408
Intensity (counts)
time (min)
E
4V WGL Reactant Spectra
1e6
121.0509
218.2124
241.206
375.2038
401.2196
453.1146
Intensity (counts)
mass to charge (m/z)
F
4V WGLProduct Spectra
1e7
121.0509
375.2049
401.2198
453.1139
Intensity (counts)
mass to charge (m/z)
G
4V WGL Reactant
375.2049
376.2071
377.2100
Y(Counts)
X(Thomsons)
H
4V WGL Brominated Product
453.1142
455.1129
Y(Counts)
X(Thomsons)

#### 4V WGM

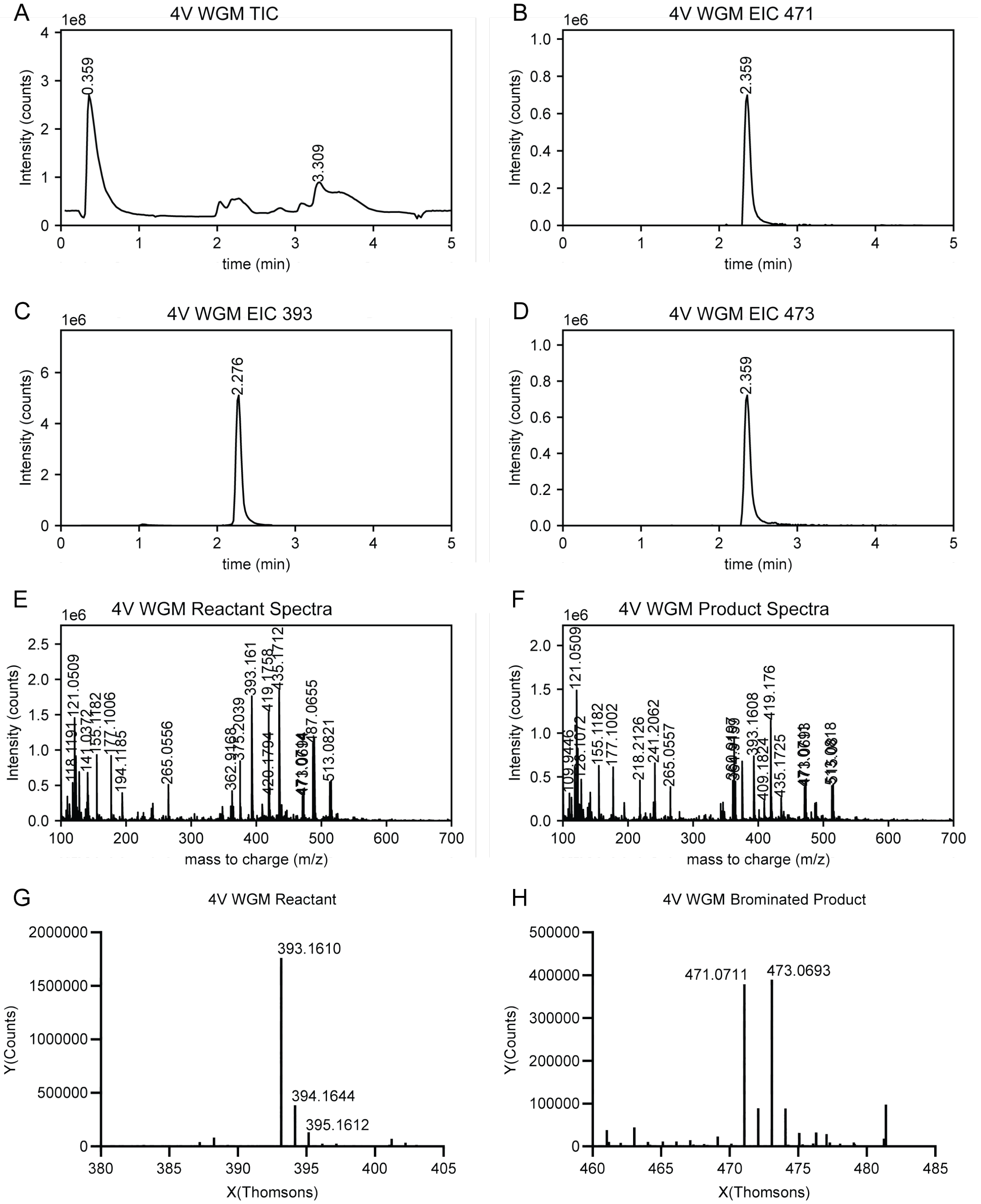
A
4V WGM TIC
1e8
Intensity (counts)
time (min)
0.359
3.309
B
4V WGM EIC 471
1e6
2.359
C
4V WGM EIC 393
2.276
D
4V WGM EIC 473
2.359
E
4V WGM Reactant Spectra
mass to charge (m/z)
121.0509
141.0372
155.1182
177.1006
194.1185
265.0556
362.9168
375.2039
393.161
419.1758
435.1712
420.1794
487.0655
513.0821
F
4V WGM Product Spectra
121.0509
155.1182
177.1002
218.2126
241.2062
265.0557
393.1608
419.176
409.1824
435.1725
G
4V WGM Reactant
Y(Counts)
X(Thomsons)
393.1610
394.1644
395.1612
H
4V WGM Brominated Product
471.0711
473.0693

#### 4V WGN

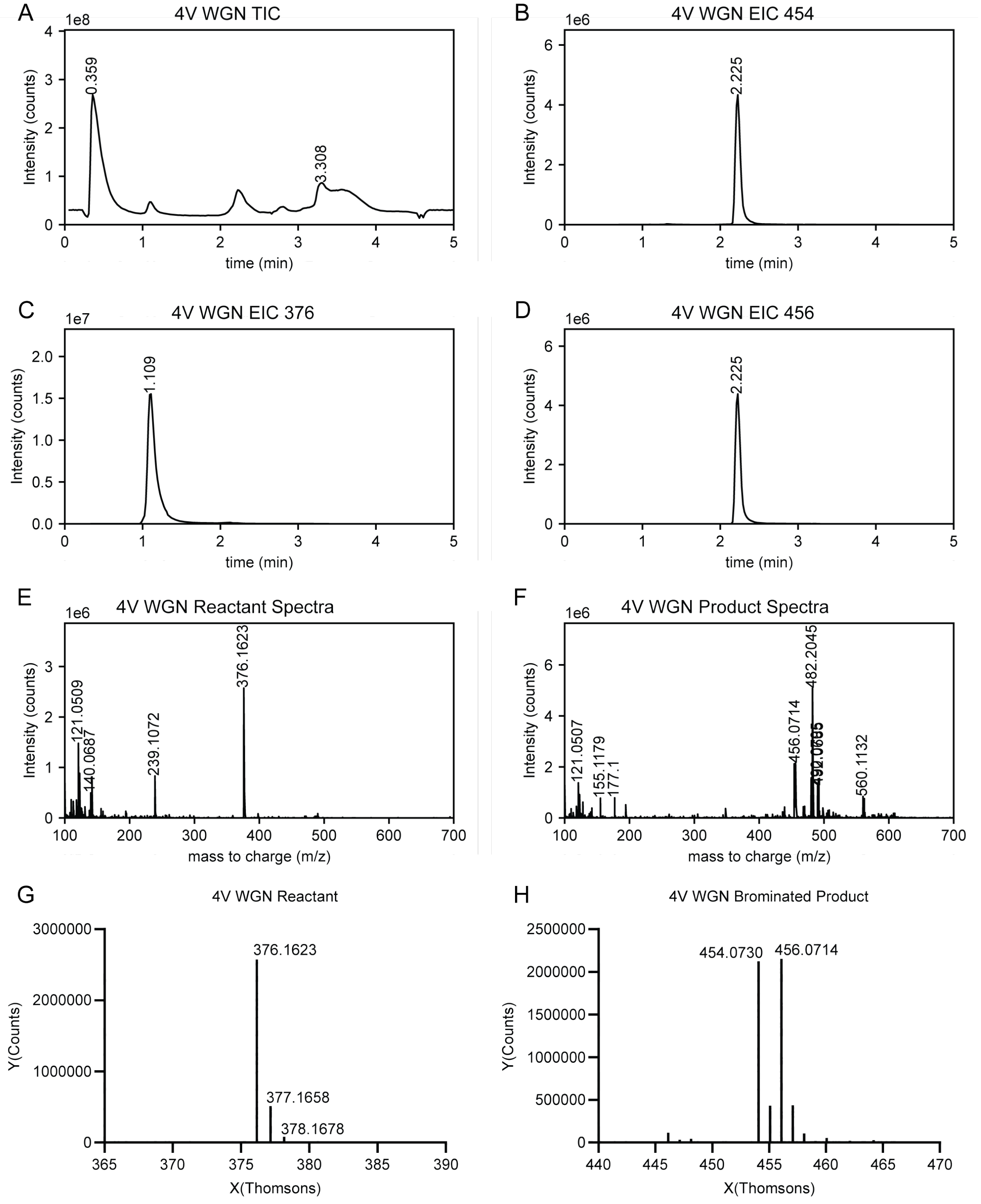
A
4V WGN TIC
1e8
Intensity (counts)
time (min)
0.359
3.308
B
4V WGN EIC 454
1e6
2.225
C
4V WGN EIC 376
1e7
1.109
D
4V WGN EIC 456
1e6
2.225
E
4V WGN Reactant Spectra
1e6
mass to charge (m/z)
121.0509
140.0687
239.1072
376.1623
F
4V WGN Product Spectra
1e6
121.0507
155.1179
177.1
456.0714
482.2045
560.1132
G
4V WGN Reactant
Y(Counts)
X(Thomsons)
376.1623
377.1658
378.1678
H
4V WGN Brominated Product
454.0730
456.0714

#### 4V WGP

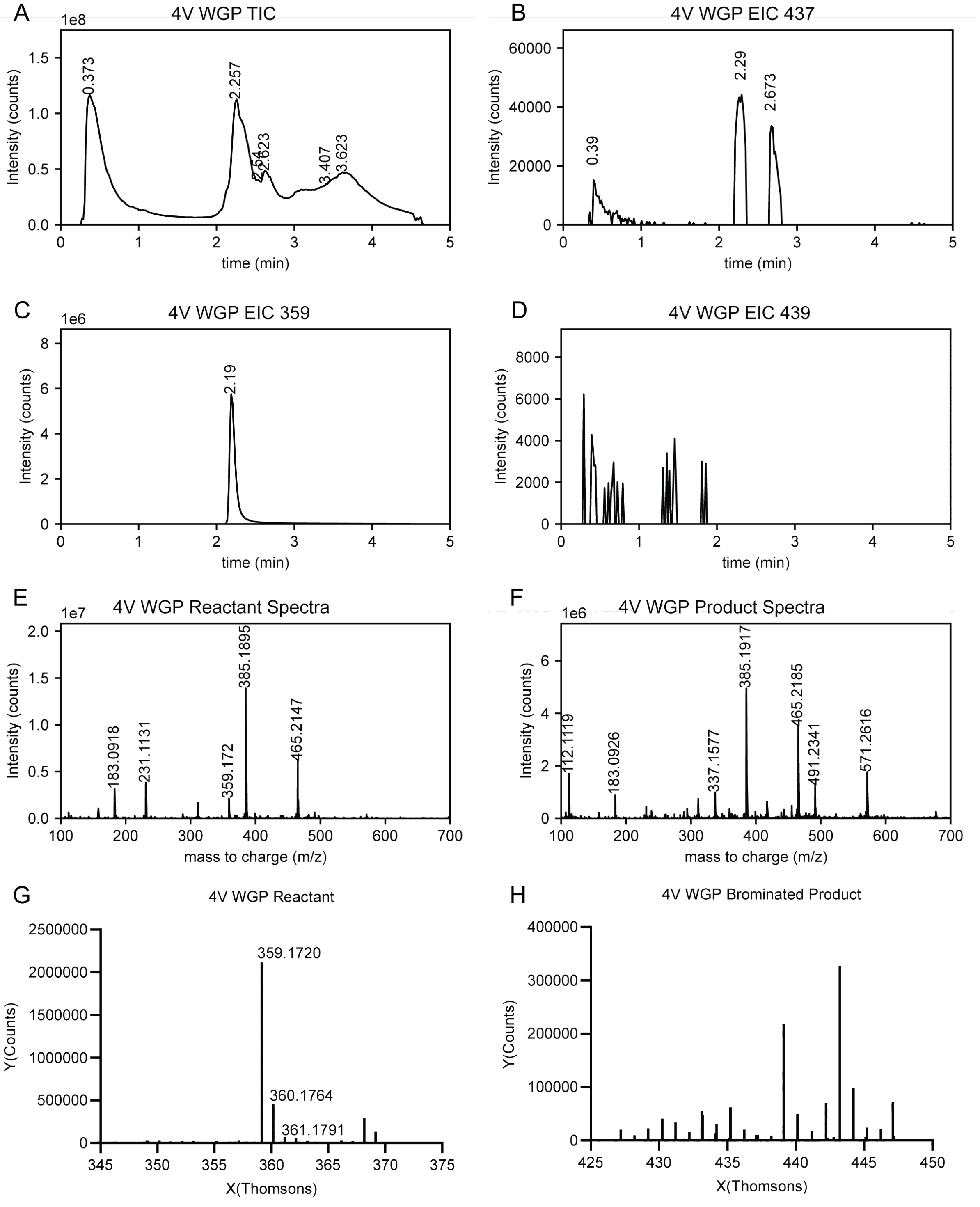

A
4V WGP TIC
1e8
Intensity (counts)
time (min)
0.373
2.257
2.54
2.623
3.407
3.623
B
4V WGP EIC 437
Intensity (counts)
time (min)
0.39
2.29
2.673
C
4V WGP EIC 359
1e6
Intensity (counts)
time (min)
2.19
D
4V WGP EIC 439
Intensity (counts)
time (min)
E
4V WGP Reactant Spectra
1e7
Intensity (counts)
mass to charge (m/z)
183.0918
231.1131
359.172
385.1895
465.2147
F
4V WGP Product Spectra
1e6
Intensity (counts)
mass to charge (m/z)
112.1119
183.0926
337.1577
385.1917
465.2185
491.2341
571.2616
G
4V WGP Reactant
Y(Counts)
X(Thomsons)
359.1720
360.1764
361.1791
H
4V WGP Brominated Product
Y(Counts)
X(Thomsons)

#### 4V WGQ

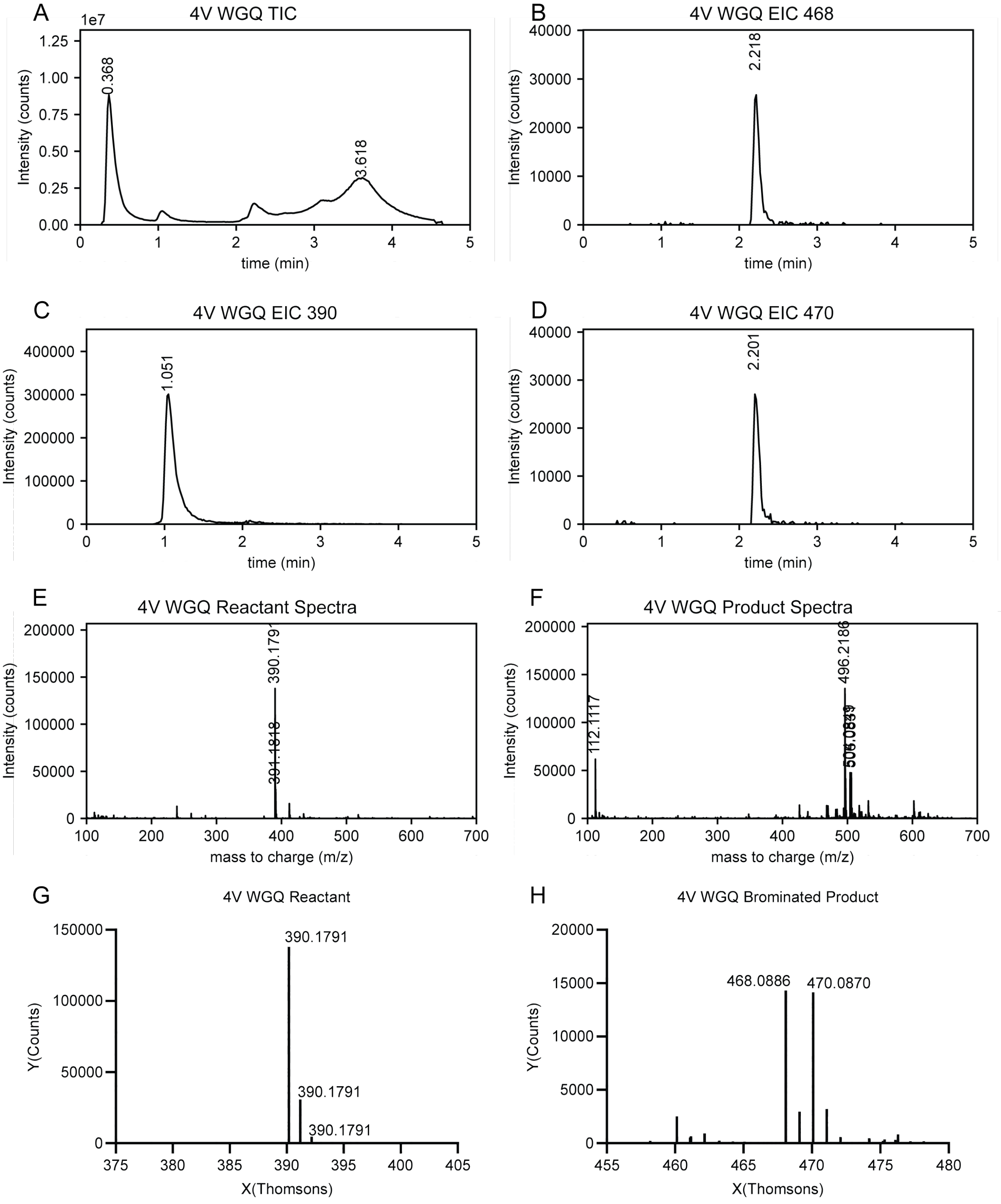

A
4V WGQ TIC
1e7
Intensity (counts)
time (min)
0.368
3.618
B
4V WGQ EIC 468
2.218
C
4V WGQ EIC 390
1.051
D
4V WGQ EIC 470
2.201
E
4V WGQ Reactant Spectra
390.1791
391.1818
mass to charge (m/z)
F
4V WGQ Product Spectra
496.2186
112.1117
G
4V WGQ Reactant
390.1791
390.1791
390.1791
Y(Counts)
X(Thomsons)
H
4V WGQ Brominated Product
468.0886
470.0870

**4V WGR**

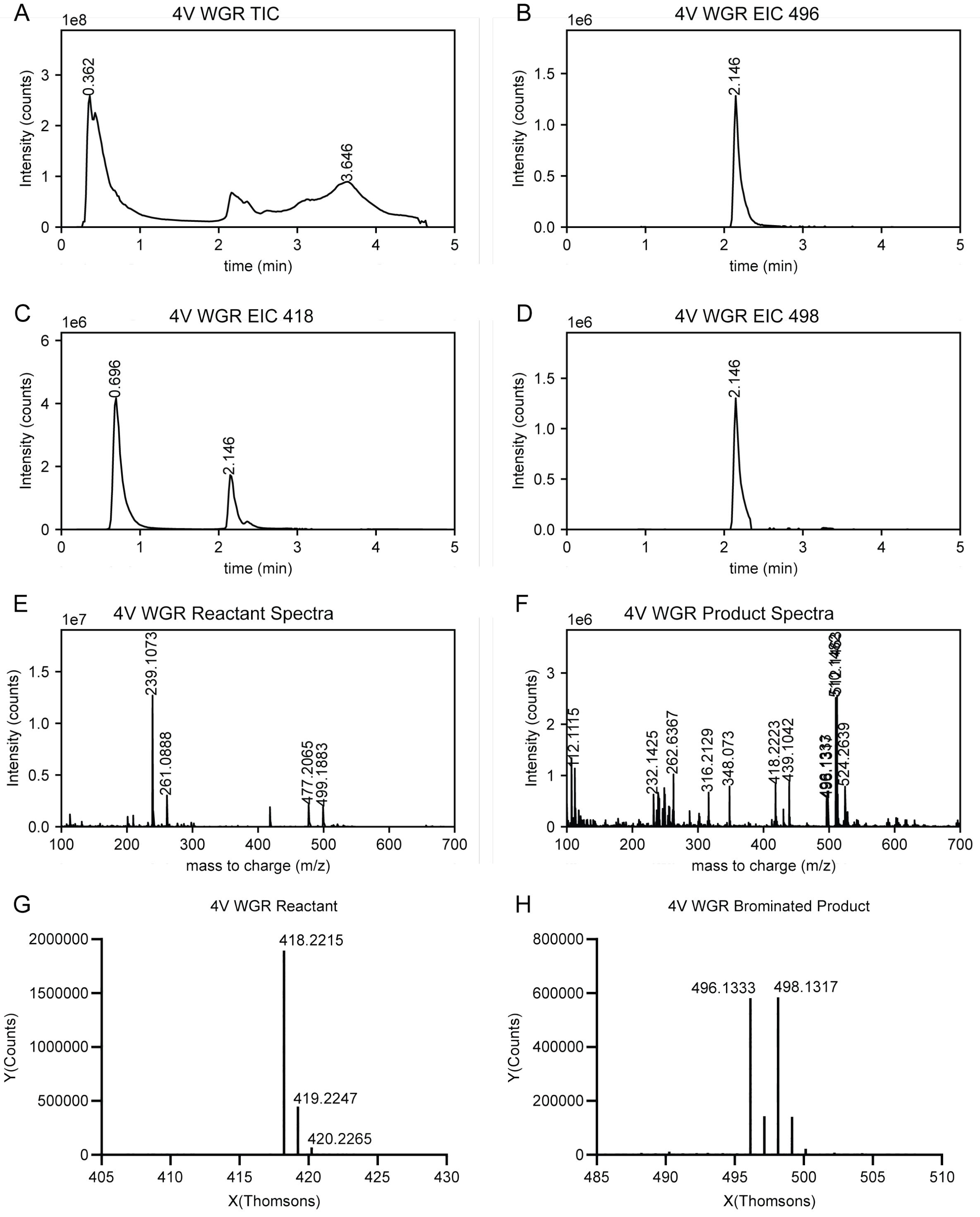

A
4V WGR TIC
1e8
Intensity (counts)
time (min)
0.362
3.646
B
4V WGR EIC 496
1e6
2.146
C
4V WGR EIC 418
0.696
2.146
D
4V WGR EIC 498
2.146
E
4V WGR Reactant Spectra
1e7
mass to charge (m/z)
239.1073
261.0888
477.2065
499.1883
F
4V WGR Product Spectra
112.1115
232.1425
262.6367
316.2129
348.073
418.2223
439.1042
510.1462
524.2639
G
4V WGR Reactant
Y(Counts)
X(Thomsons)
418.2215
419.2247
420.2265
H
4V WGR Brominated Product
496.1333
498.1317

**4V WGT**

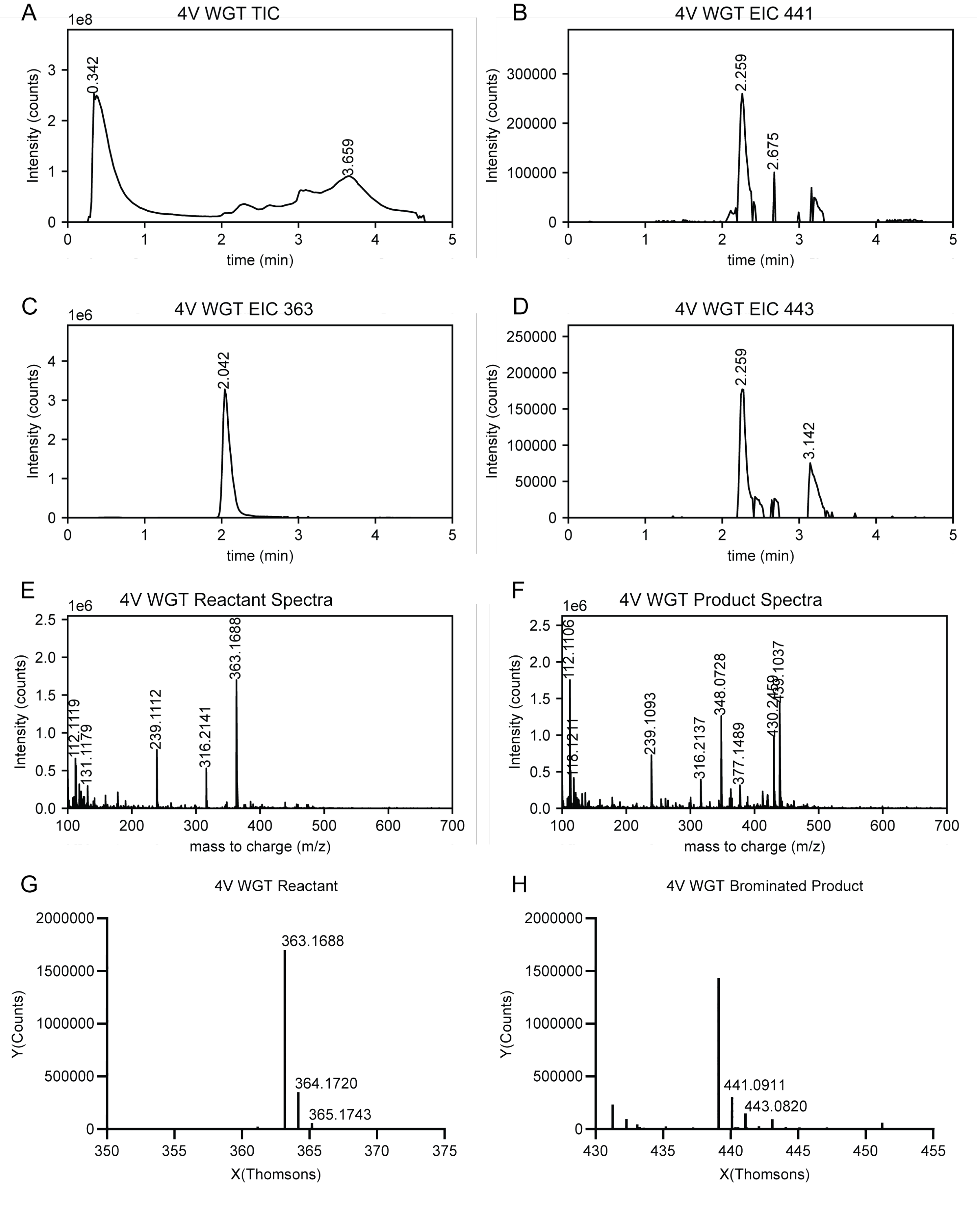

A
4V WGT TIC
1e8
Intensity (counts)
time (min)
0.342
3.659
B
4V WGT EIC 441
Intensity (counts)
time (min)
2.259
2.675
C
4V WGT EIC 363
1e6
Intensity (counts)
time (min)
2.042
D
4V WGT EIC 443
Intensity (counts)
time (min)
2.259
3.142
E
4V WGT Reactant Spectra
1e6
Intensity (counts)
mass to charge (m/z)
112.1119
131.1179
239.1112
316.2141
363.1688
F
4V WGT Product Spectra
1e6
Intensity (counts)
mass to charge (m/z)
112.1106
118.1211
239.1093
316.2137
348.0728
377.1489
430.2459
G
4V WGT Reactant
Y(Counts)
X(Thomsons)
363.1688
364.1720
365.1743
H
4V WGT Brominated Product
Y(Counts)
X(Thomsons)
441.0911
443.0820

#### 4V WGV

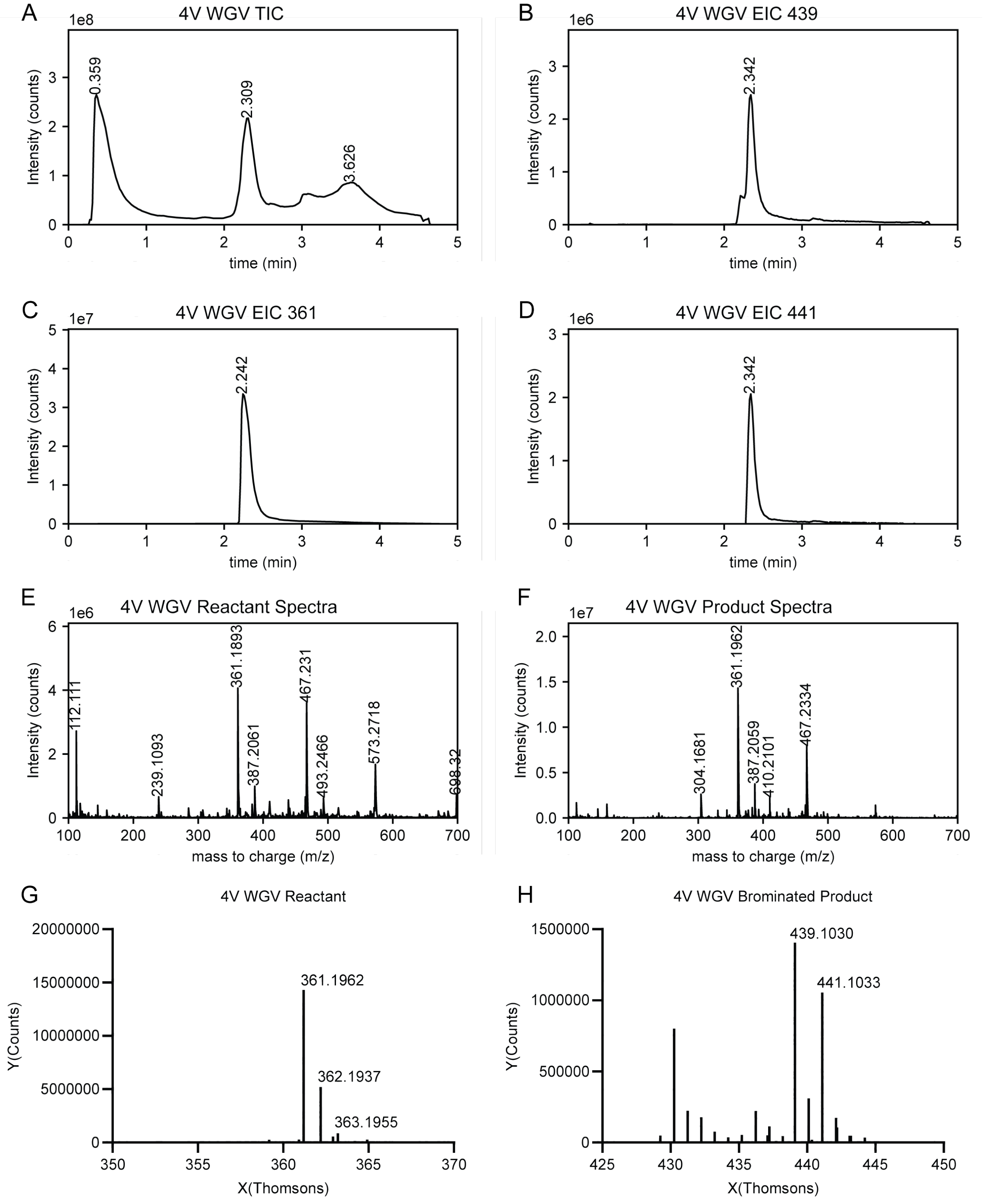
A
4V WGV TIC
1e8
0.359
2.309
3.626
Intensity (counts)
time (min)
B
4V WGV EIC 439
1e6
2.342
Intensity (counts)
time (min)
C
4V WGV EIC 361
1e7
2.242
Intensity (counts)
time (min)
D
4V WGV EIC 441
1e6
2.342
Intensity (counts)
time (min)
E
4V WGV Reactant Spectra
1e6
112.111
239.1093
361.1893
387.2061
467.231
493.2466
573.2718
698.32
Intensity (counts)
mass to charge (m/z)
F
4V WGV Product Spectra
1e7
304.1681
361.1962
387.2059
410.2101
467.2334
Intensity (counts)
mass to charge (m/z)
G
4V WGV Reactant
361.1962
362.1937
363.1955
Y(Counts)
X(Thomsons)
H
4V WGV Brominated Product
439.1030
441.1033
Y(Counts)
X(Thomsons)

#### 4V WGY

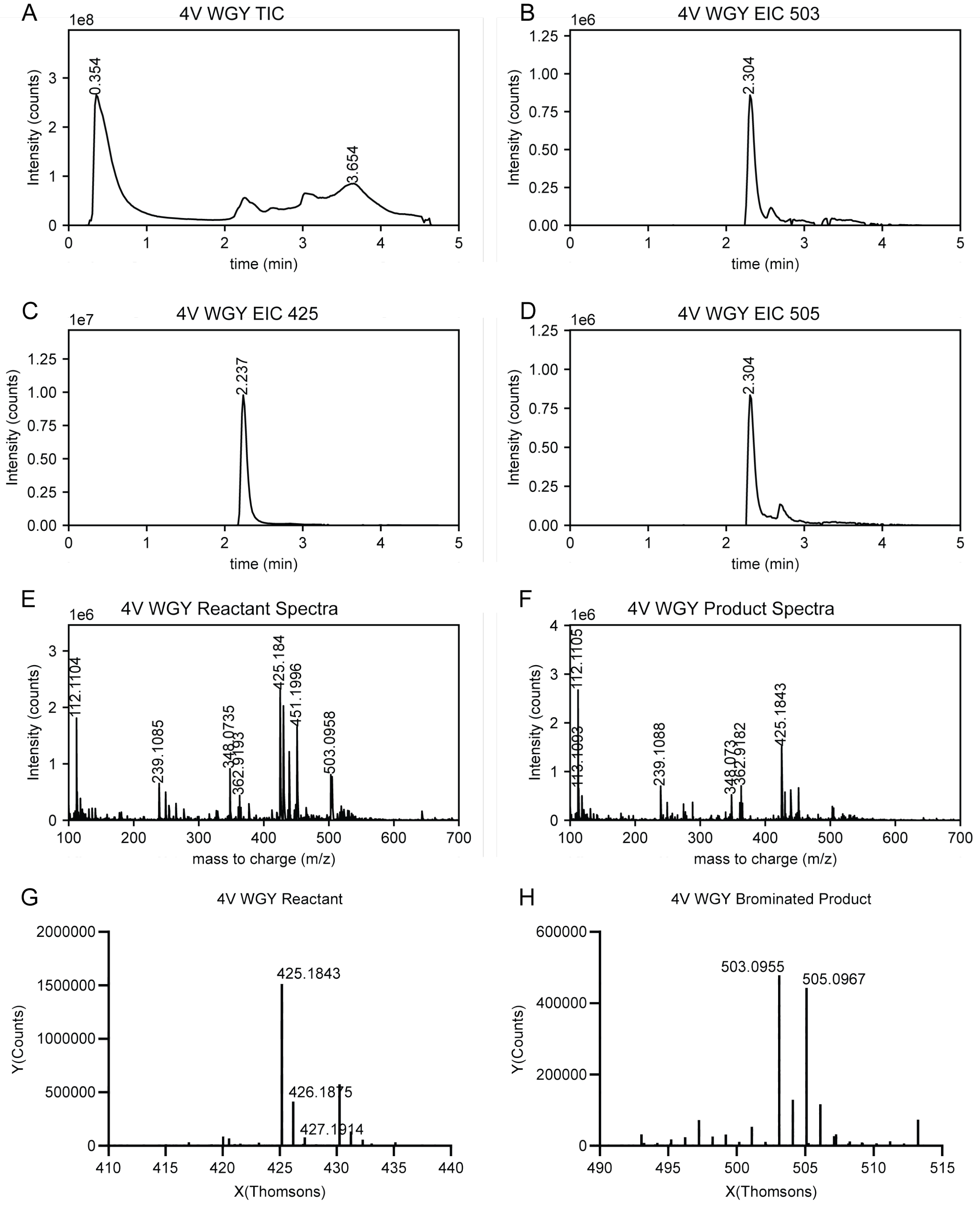
A
4V WGY TIC
1e8
Intensity (counts)
time (min)
0.354
3.654
B
4V WGY EIC 503
1e6
Intensity (counts)
time (min)
2.304
C
4V WGY EIC 425
1e7
Intensity (counts)
time (min)
2.237
D
4V WGY EIC 505
1e6
Intensity (counts)
time (min)
2.304
E
4V WGY Reactant Spectra
1e6
Intensity (counts)
mass to charge (m/z)
112.1104
239.1085
348.0735
362.9193
425.184
451.1996
503.0958
F
4V WGY Product Spectra
1e6
Intensity (counts)
mass to charge (m/z)
112.1105
113.1093
239.1088
348.073
362.9182
425.1843
G
4V WGY Reactant
Y(Counts)
X(Thomsons)
425.1843
426.1875
427.1914
H
4V WGY Brominated Product
Y(Counts)
X(Thomsons)
503.0955
505.0967

#### 4V AWG

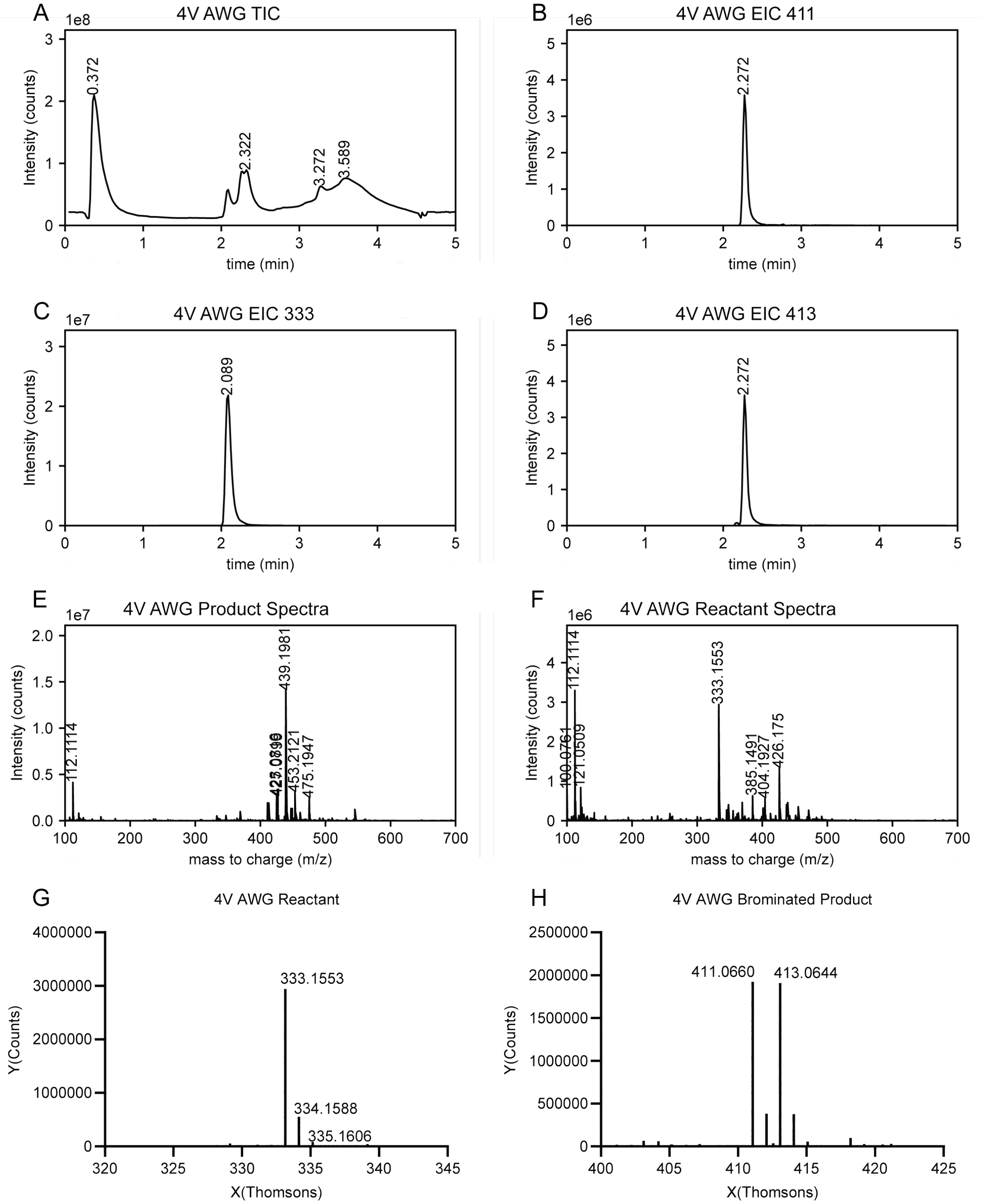

A
4V AWG TIC
1e8
Intensity (counts)
time (min)
0.372
2.322
3.272
3.589
B
4V AWG EIC 411
1e6
2.272
C
4V AWG EIC 333
1e7
2.089
D
4V AWG EIC 413
1e6
2.272
E
4V AWG Product Spectra
1e7
mass to charge (m/z)
112.1114
439.1981
453.2121
475.1947
F
4V AWG Reactant Spectra
1e6
112.1114
100.0761
121.0509
333.1553
385.1491
404.1927
426.175
G
4V AWG Reactant
Y(Counts)
X(Thomsons)
333.1553
334.1588
335.1606
H
4V AWG Brominated Product
411.0660
413.0644

#### 4V CWG

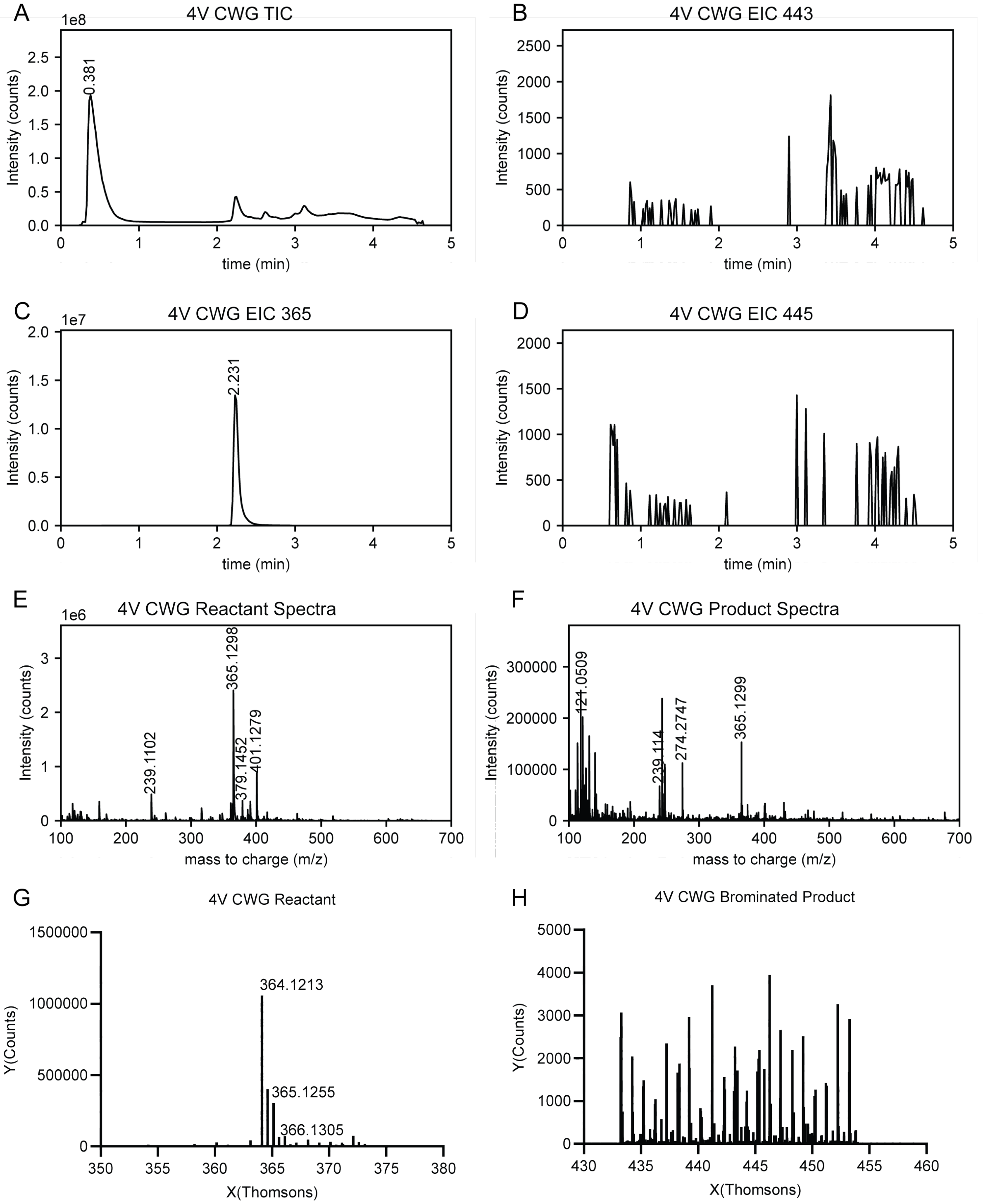

A
4V CWG TIC
1e8
0.381
Intensity (counts)
time (min)
B
4V CWG EIC 443
Intensity (counts)
time (min)
C
4V CWG EIC 365
1e7
2.231
Intensity (counts)
time (min)
D
4V CWG EIC 445
Intensity (counts)
time (min)
E
4V CWG Reactant Spectra
1e6
365.1298
239.1102
379.1452
401.1279
Intensity (counts)
mass to charge (m/z)
F
4V CWG Product Spectra
121.0509
239.114
274.2747
365.1299
Intensity (counts)
mass to charge (m/z)
G
4V CWG Reactant
364.1213
365.1255
366.1305
Y(Counts)
X(Thomsons)
H
4V CWG Brominated Product
Y(Counts)
X(Thomsons)

### 4V DWG

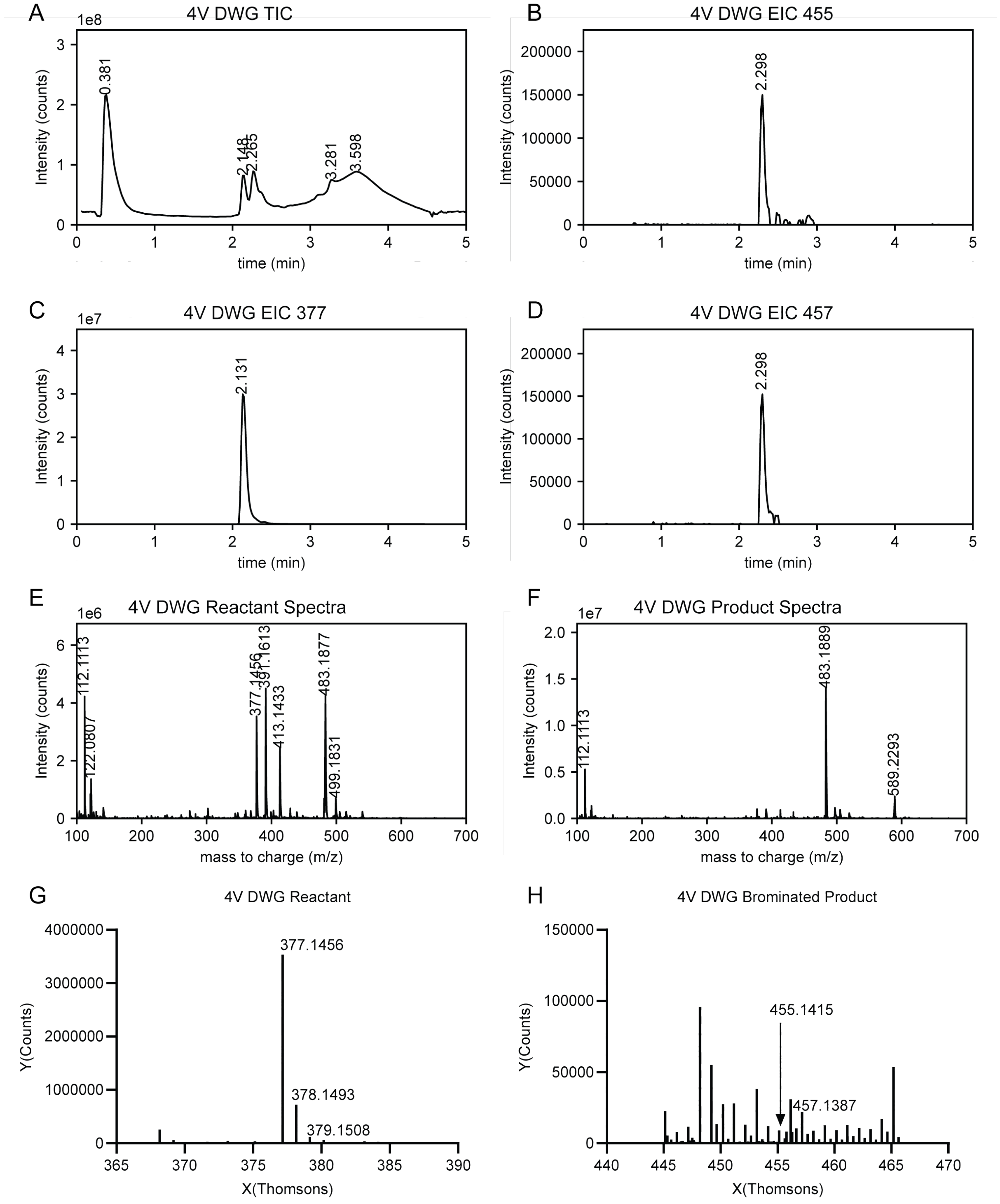

A
4V DWG TIC
1e8
0.381
2.148
2.265
3.281
3.598
Intensity (counts)
time (min)
B
4V DWG EIC 455
2.298
Intensity (counts)
time (min)
C
4V DWG EIC 377
1e7
2.131
Intensity (counts)
time (min)
D
4V DWG EIC 457
2.298
Intensity (counts)
time (min)
E
4V DWG Reactant Spectra
1e6
112.1113
122.0807
377.1456
391.1613
413.1433
483.1877
499.1831
Intensity (counts)
mass to charge (m/z)
F
4V DWG Product Spectra
1e7
112.1113
483.1889
589.2293
Intensity (counts)
mass to charge (m/z)
G
4V DWG Reactant
377.1456
378.1493
379.1508
Y(Counts)
X(Thomsons)
H
4V DWG Brominated Product
455.1415
457.1387
Y(Counts)
X(Thomsons)

#### 4V EWG

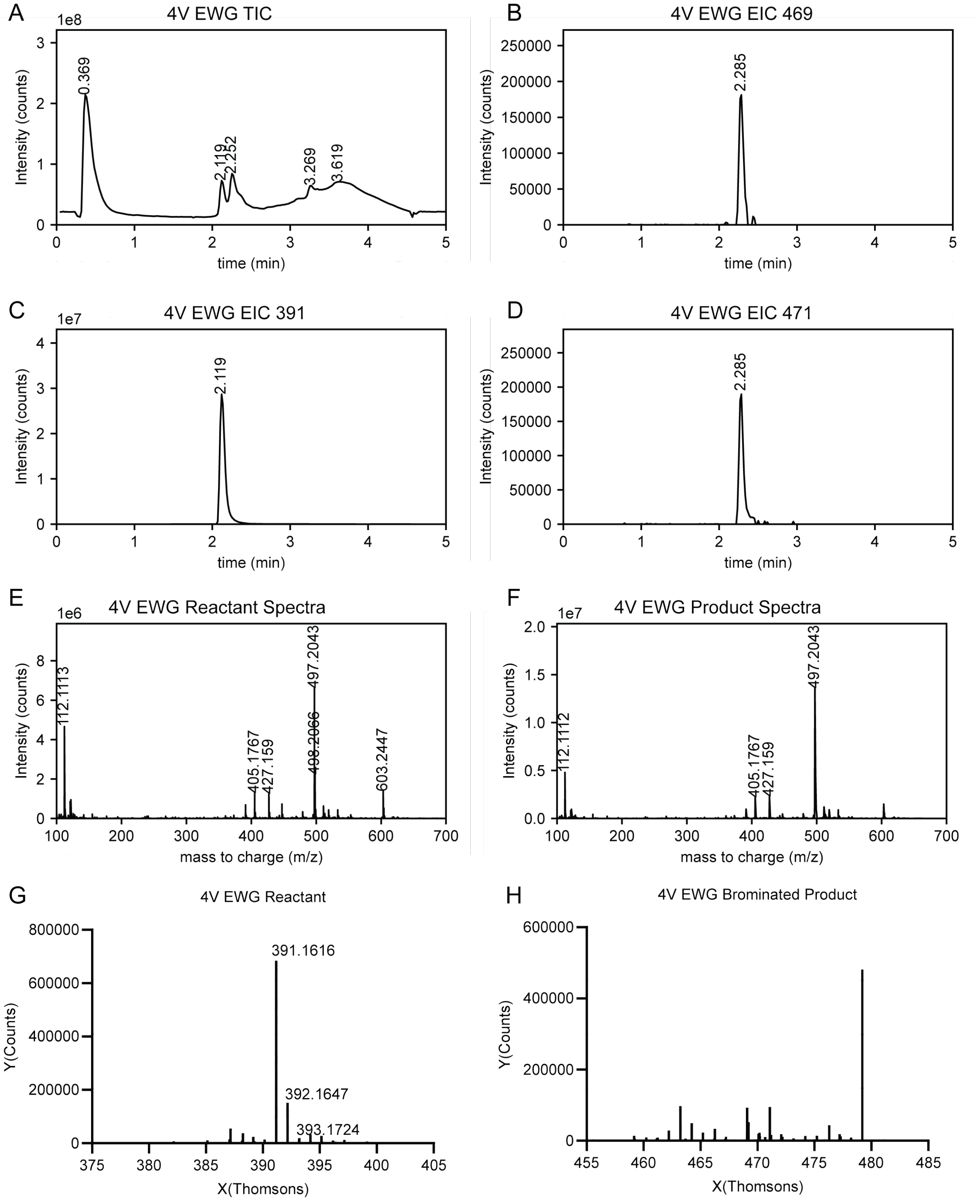
A
4V EWG TIC
1e8
Intensity (counts)
time (min)
0.369
2.119
2.252
3.269
3.619
B
4V EWG EIC 469
Intensity (counts)
time (min)
2.285
C
4V EWG EIC 391
1e7
Intensity (counts)
time (min)
2.119
D
4V EWG EIC 471
Intensity (counts)
time (min)
2.285
E
4V EWG Reactant Spectra
1e6
Intensity (counts)
mass to charge (m/z)
112.1113
405.1767
427.159
497.2043
498.2066
603.2447
F
4V EWG Product Spectra
1e7
Intensity (counts)
mass to charge (m/z)
112.1112
405.1767
427.159
497.2043
G
4V EWG Reactant
Y(Counts)
X(Thomsons)
391.1616
392.1647
393.1724
H
4V EWG Brominated Product
Y(Counts)
X(Thomsons)

#### 4V FWG

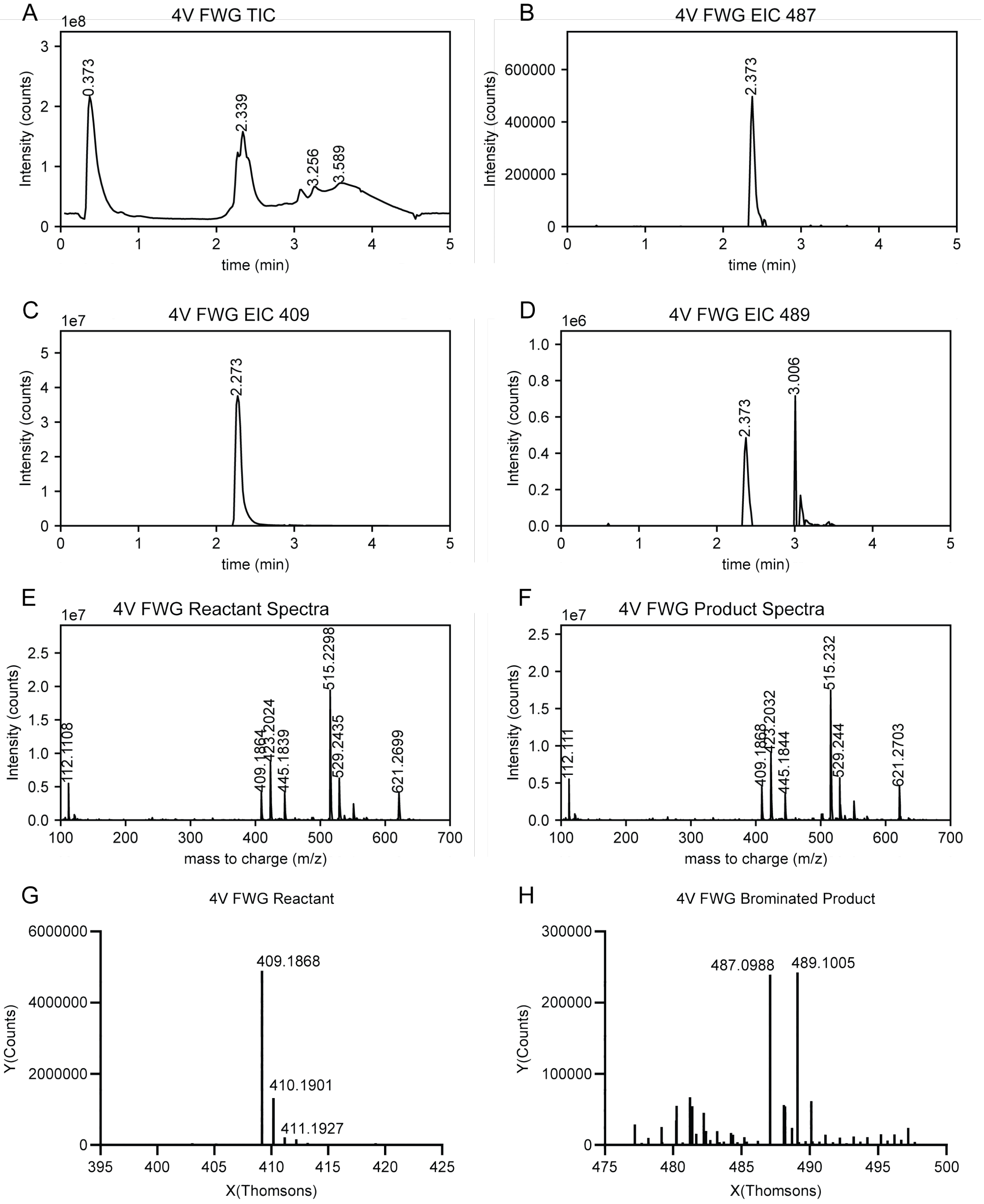

A
4V FWG TIC
1e8
Intensity (counts)
time (min)
0.373
2.339
3.256
3.589
B
4V FWG EIC 487
Intensity (counts)
time (min)
2.373
C
4V FWG EIC 409
1e7
Intensity (counts)
time (min)
2.273
D
4V FWG EIC 489
1e6
Intensity (counts)
time (min)
2.373
3.006
E
4V FWG Reactant Spectra
1e7
Intensity (counts)
mass to charge (m/z)
112.1108
409.1864
423.2024
445.1839
515.2298
529.2435
621.2699
F
4V FWG Product Spectra
1e7
Intensity (counts)
mass to charge (m/z)
112.111
409.1868
423.2032
445.1844
515.232
529.244
621.2703
G
4V FWG Reactant
Y(Counts)
X(Thomsons)
409.1868
410.1901
411.1927
H
4V FWG Brominated Product
Y(Counts)
X(Thomsons)
487.0988
489.1005

**4V HWG**

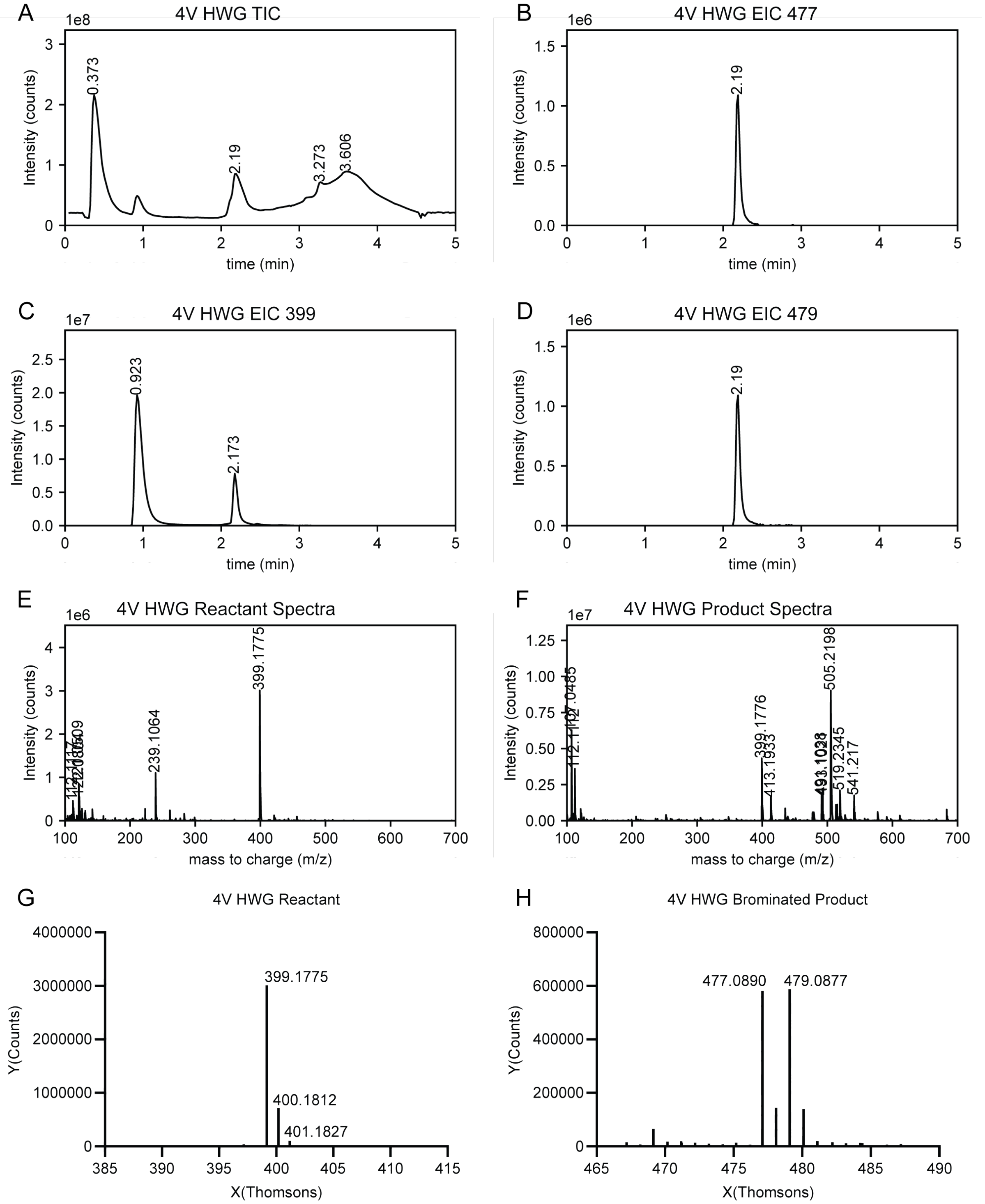
A
4V HWG TIC
1e8
Intensity (counts)
time (min)
0.373
2.19
3.273
3.606
B
4V HWG EIC 477
1e6
Intensity (counts)
time (min)
2.19
C
4V HWG EIC 399
1e7
Intensity (counts)
time (min)
0.923
2.173
D
4V HWG EIC 479
1e6
Intensity (counts)
time (min)
2.19
E
4V HWG Reactant Spectra
1e6
Intensity (counts)
mass to charge (m/z)
239.1064
399.1775
F
4V HWG Product Spectra
1e7
Intensity (counts)
mass to charge (m/z)
399.1776
413.1933
505.2198
519.2345
541.217
G
4V HWG Reactant
Y(Counts)
X(Thomsons)
399.1775
400.1812
401.1827
H
4V HWG Brominated Product
Y(Counts)
X(Thomsons)
477.0890
479.0877

#### 4V KWG

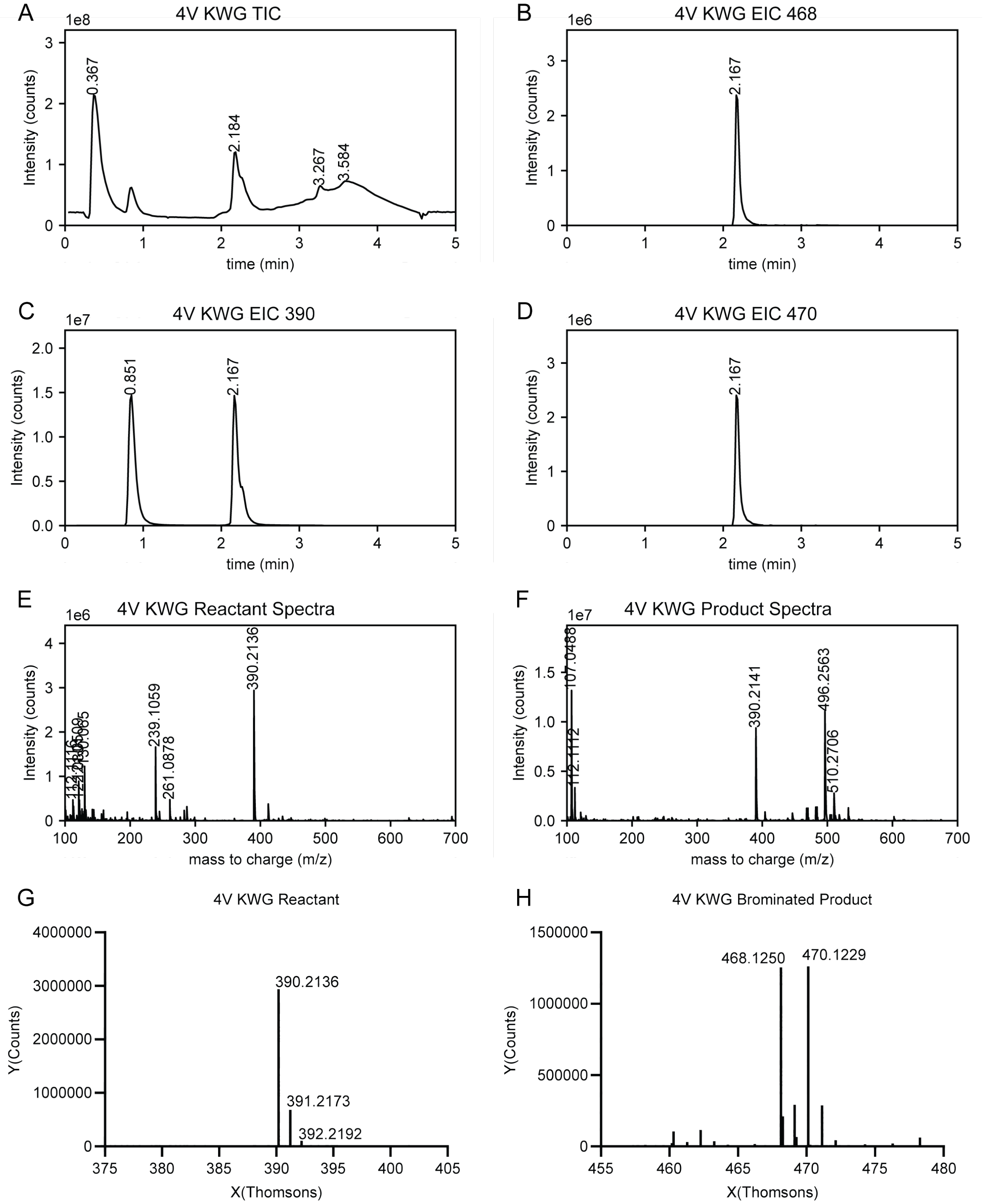
A
4V KWG TIC
1e8
Intensity (counts)
time (min)
0.367
2.184
3.267
3.584
B
4V KWG EIC 468
1e6
Intensity (counts)
time (min)
2.167
C
4V KWG EIC 390
1e7
Intensity (counts)
time (min)
0.851
2.167
D
4V KWG EIC 470
1e6
Intensity (counts)
time (min)
2.167
E
4V KWG Reactant Spectra
1e6
Intensity (counts)
mass to charge (m/z)
390.2136
239.1059
261.0878
F
4V KWG Product Spectra
1e7
Intensity (counts)
mass to charge (m/z)
107.0488
112.1112
390.2141
496.2563
510.2706
G
4V KWG Reactant
Y(Counts)
X(Thomsons)
390.2136
391.2173
392.2192
H
4V KWG Brominated Product
Y(Counts)
X(Thomsons)
468.1250
470.1229

#### 4V LWG

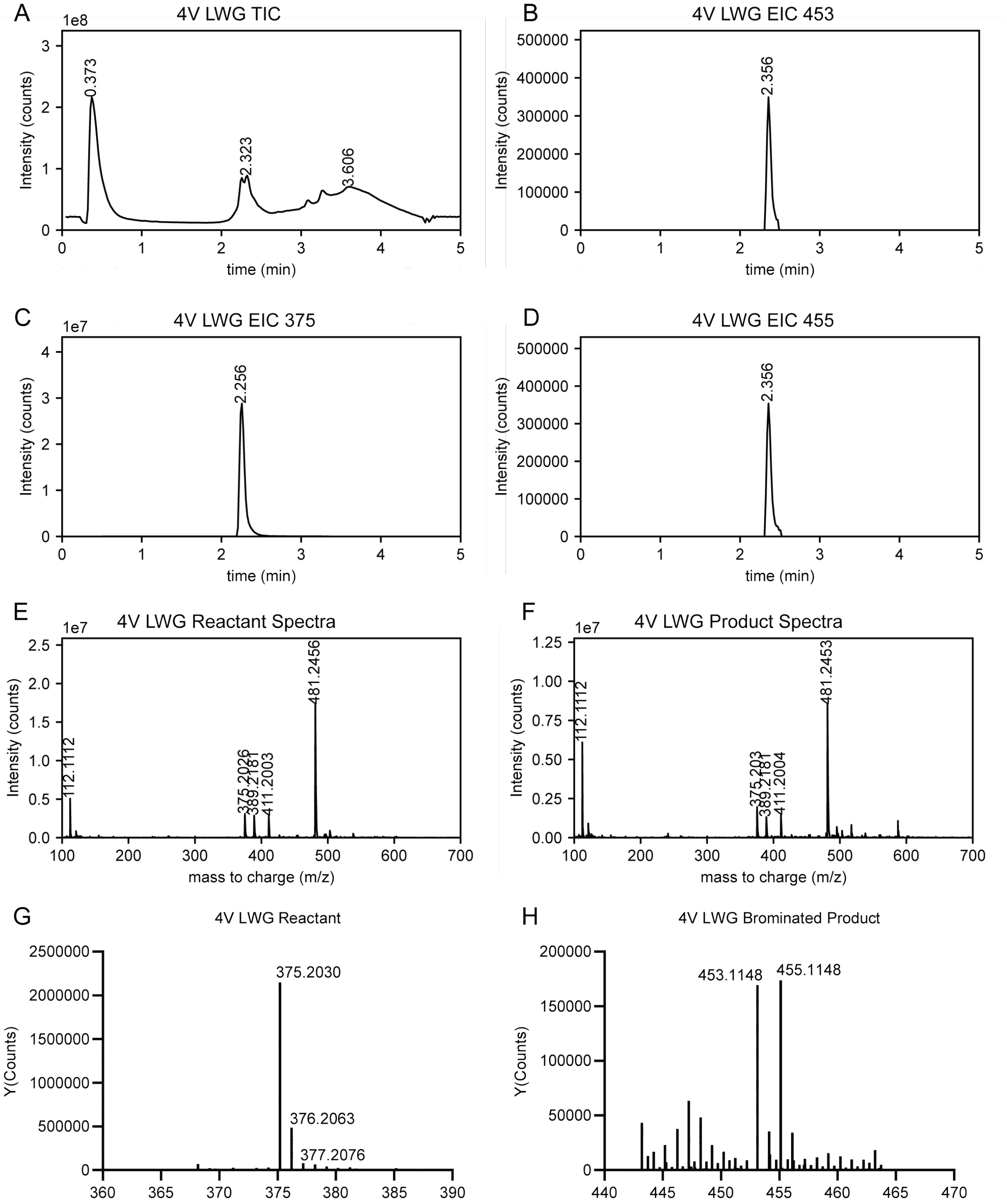
A
4V LWG TIC
1e8
Intensity (counts)
time (min)
0.373
2.323
3.606
B
4V LWG EIC 453
Intensity (counts)
time (min)
2.356
C
4V LWG EIC 375
1e7
Intensity (counts)
time (min)
2.256
D
4V LWG EIC 455
Intensity (counts)
time (min)
2.356
E
4V LWG Reactant Spectra
1e7
Intensity (counts)
mass to charge (m/z)
112.1112
375.2026
389.2181
411.2003
481.2456
F
4V LWG Product Spectra
1e7
Intensity (counts)
mass to charge (m/z)
112.1112
375.203
389.2181
411.2004
481.2453
G
4V LWG Reactant
Y(Counts)
X(Thomsons)
375.2030
376.2063
377.2076
H
4V LWG Brominated Product
Y(Counts)
X(Thomsons)
453.1148
455.1148

#### 4V MWG

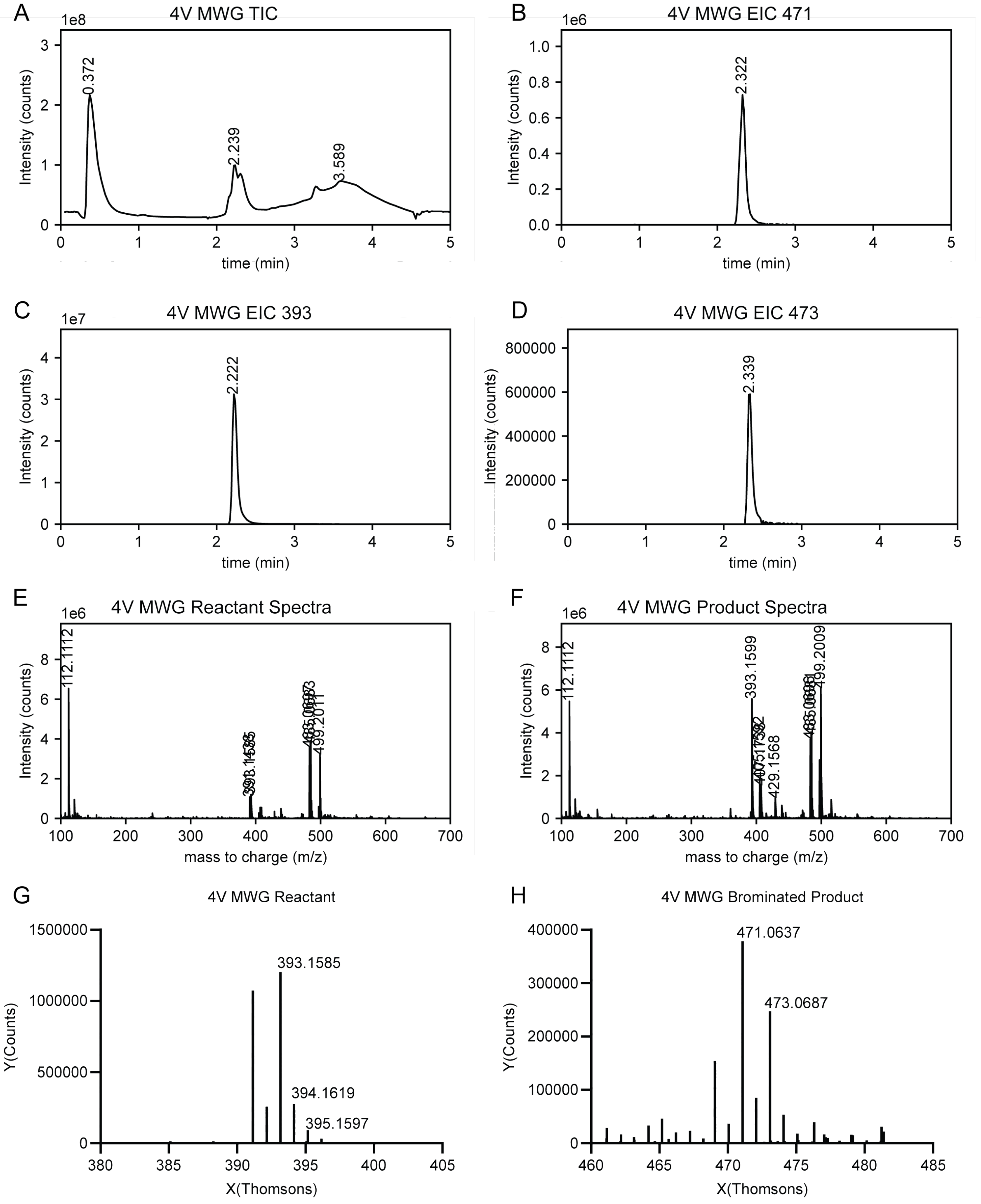
A
4V MWG TIC
1e8
Intensity (counts)
time (min)
0.372
2.239
3.589
B
4V MWG EIC 471
1e6
Intensity (counts)
time (min)
2.322
C
4V MWG EIC 393
1e7
Intensity (counts)
time (min)
2.222
D
4V MWG EIC 473
Intensity (counts)
time (min)
2.339
E
4V MWG Reactant Spectra
1e6
Intensity (counts)
mass to charge (m/z)
112.1112
499.2011
F
4V MWG Product Spectra
1e6
Intensity (counts)
mass to charge (m/z)
112.1112
393.1599
499.2009
429.1568
G
4V MWG Reactant
Y(Counts)
X(Thomsons)
393.1585
394.1619
395.1597
H
4V MWG Brominated Product
Y(Counts)
X(Thomsons)
471.0637
473.0687

#### 4V NWG

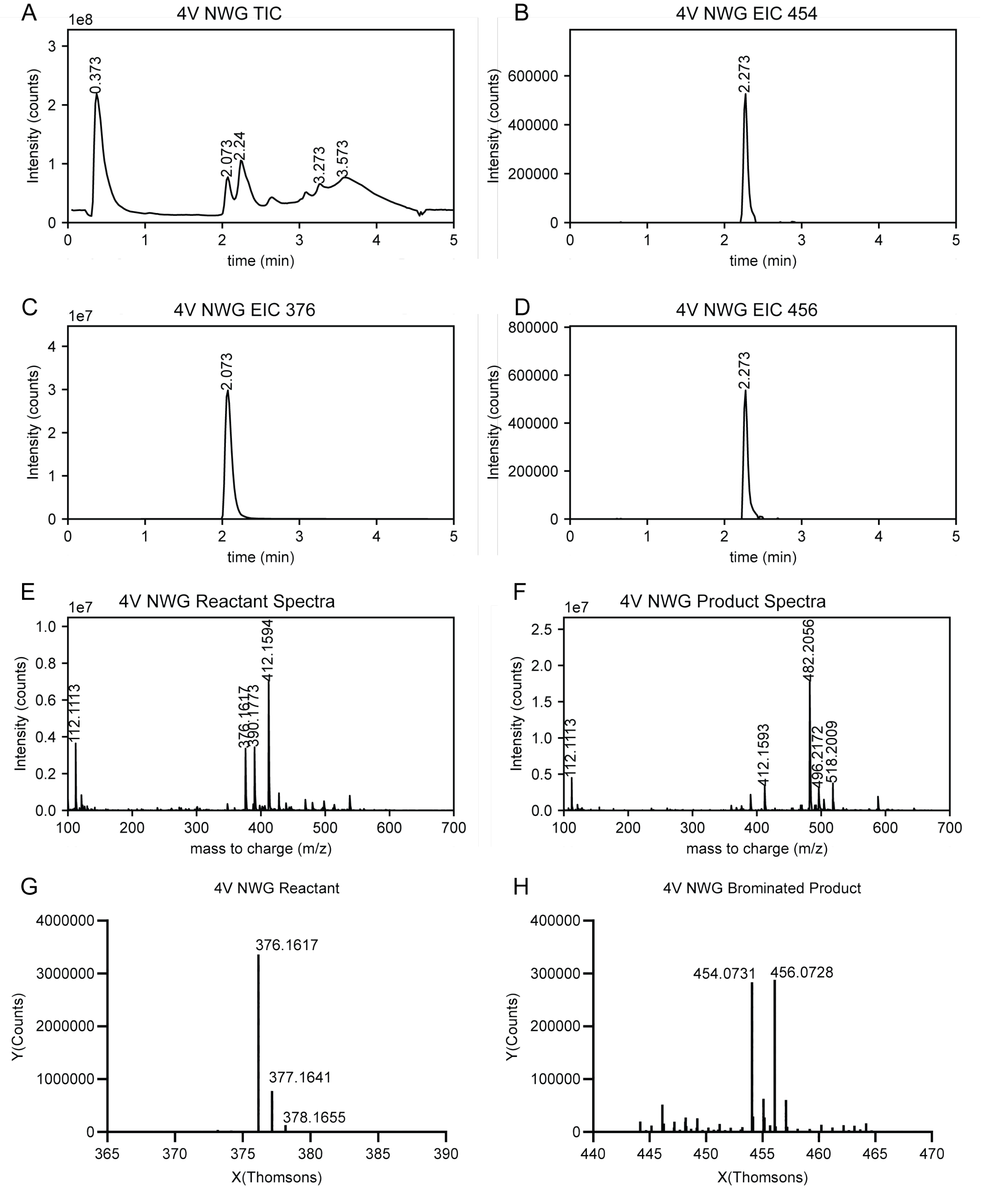

A
4V NWG TIC
1e8
Intensity (counts)
time (min)
0.373
2.073
2.24
3.273
3.573
B
4V NWG EIC 454
Intensity (counts)
time (min)
2.273
C
4V NWG EIC 376
1e7
Intensity (counts)
time (min)
2.073
D
4V NWG EIC 456
Intensity (counts)
time (min)
2.273
E
4V NWG Reactant Spectra
1e7
Intensity (counts)
mass to charge (m/z)
112.1113
376.1617
390.1773
412.1594
F
4V NWG Product Spectra
1e7
Intensity (counts)
mass to charge (m/z)
112.1113
412.1593
482.2056
496.2172
518.2009
G
4V NWG Reactant
Y(Counts)
X(Thomsons)
376.1617
377.1641
378.1655
H
4V NWG Brominated Product
Y(Counts)
X(Thomsons)
454.0731
456.0728

#### 4V PWG

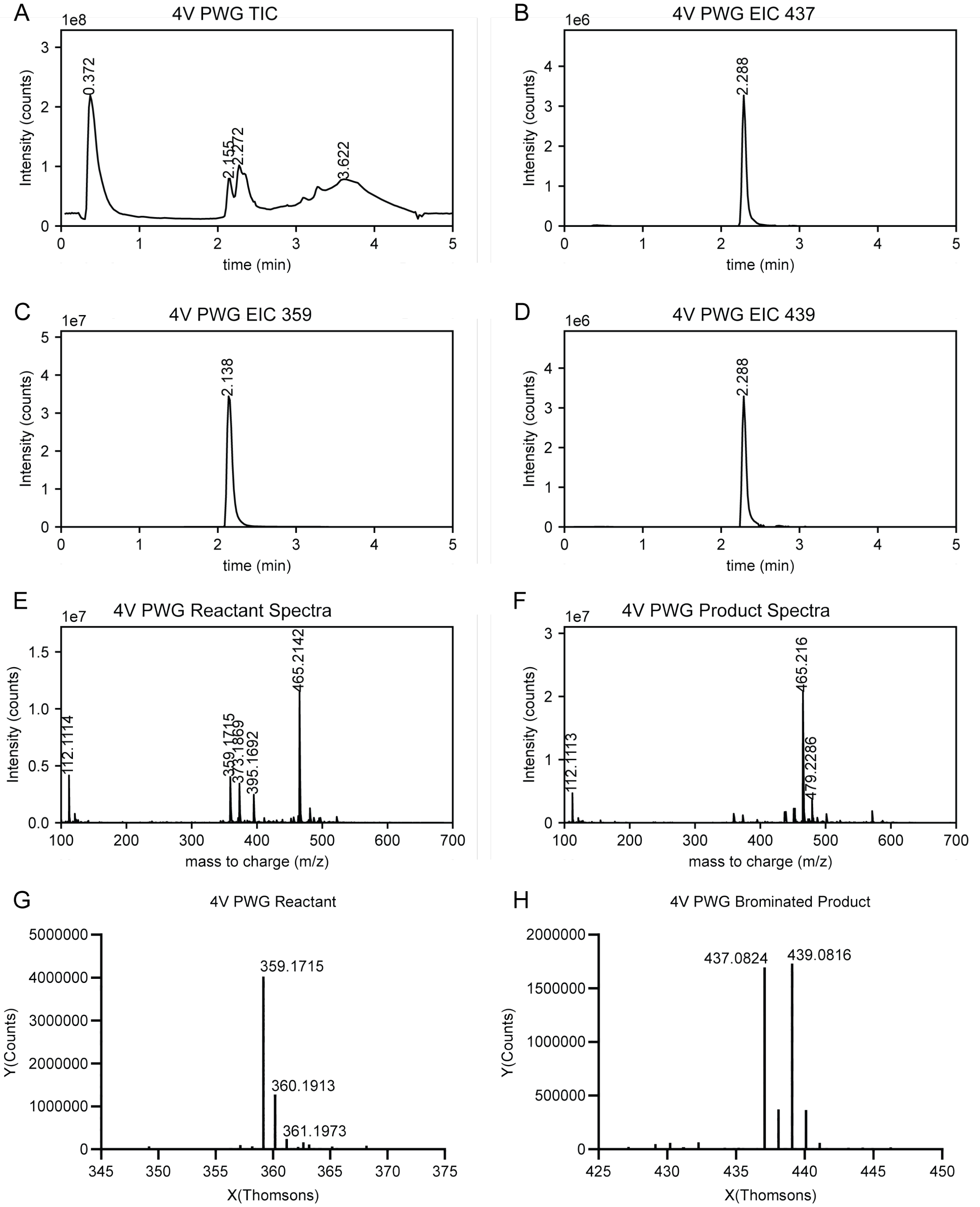

A
4V PWG TIC
1e8
Intensity (counts)
time (min)
0.372
2.155
2.272
3.622
B
4V PWG EIC 437
1e6
Intensity (counts)
time (min)
2.288
C
4V PWG EIC 359
1e7
Intensity (counts)
time (min)
2.138
D
4V PWG EIC 439
1e6
Intensity (counts)
time (min)
2.288
E
4V PWG Reactant Spectra
1e7
Intensity (counts)
mass to charge (m/z)
112.1114
359.1715
373.1869
395.1692
465.2142
F
4V PWG Product Spectra
1e7
Intensity (counts)
mass to charge (m/z)
112.1113
465.216
479.2286
G
4V PWG Reactant
Y(Counts)
X(Thomsons)
359.1715
360.1913
361.1973
H
4V PWG Brominated Product
Y(Counts)
X(Thomsons)
437.0824
439.0816

#### 4V QWG

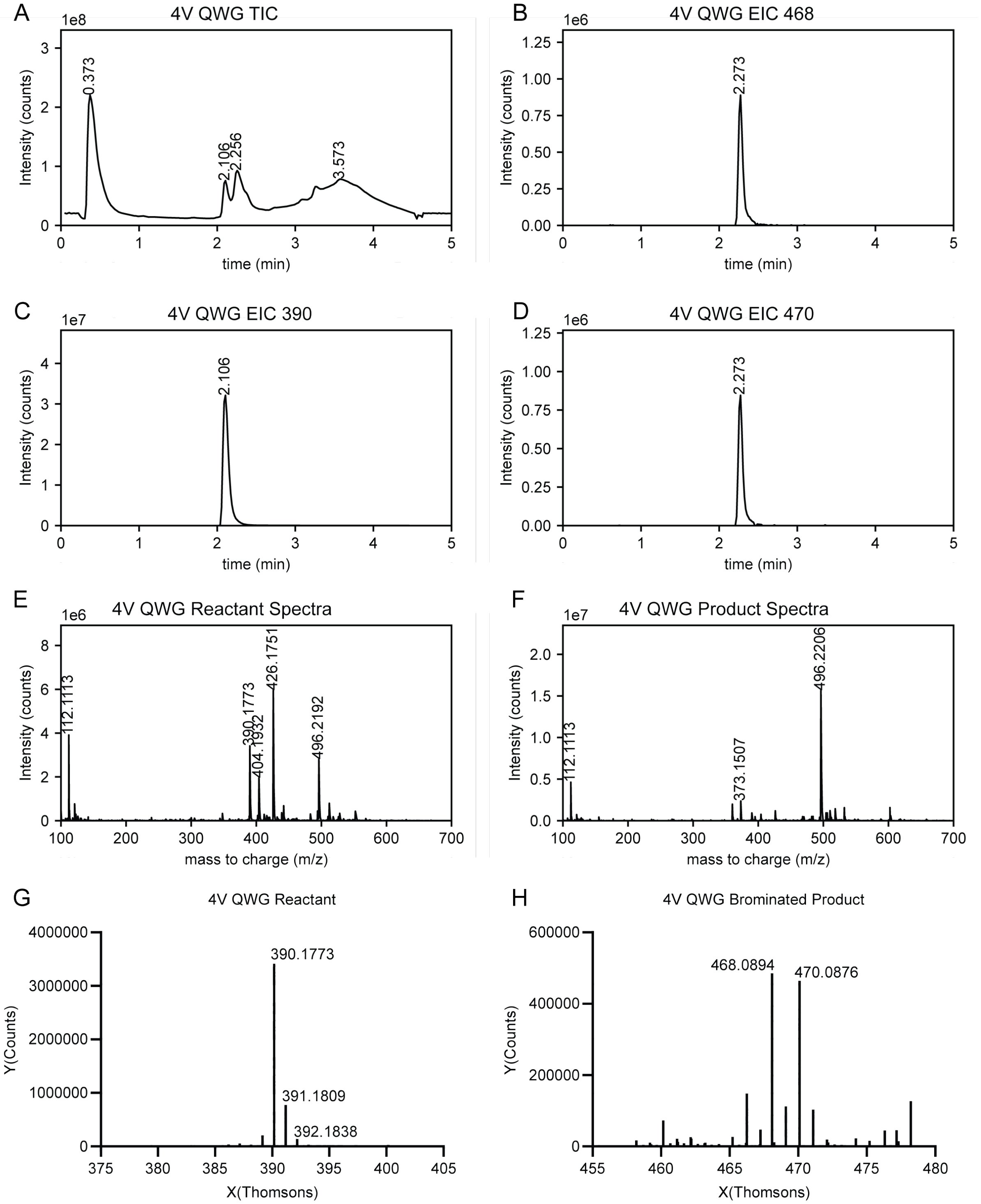

A
4V QWG TIC
1e8
Intensity (counts)
time (min)
0.373
2.106
2.256
3.573
B
4V QWG EIC 468
1e6
Intensity (counts)
time (min)
2.273
C
4V QWG EIC 390
1e7
Intensity (counts)
time (min)
2.106
D
4V QWG EIC 470
1e6
Intensity (counts)
time (min)
2.273
E
4V QWG Reactant Spectra
1e6
Intensity (counts)
mass to charge (m/z)
112.1113
390.1773
404.1932
426.1751
496.2192
F
4V QWG Product Spectra
1e7
Intensity (counts)
mass to charge (m/z)
112.1113
373.1507
496.2206
G
4V QWG Reactant
Y(Counts)
X(Thomsons)
390.1773
391.1809
392.1838
H
4V QWG Brominated Product
Y(Counts)
X(Thomsons)
468.0894
470.0876

#### 4V RWG

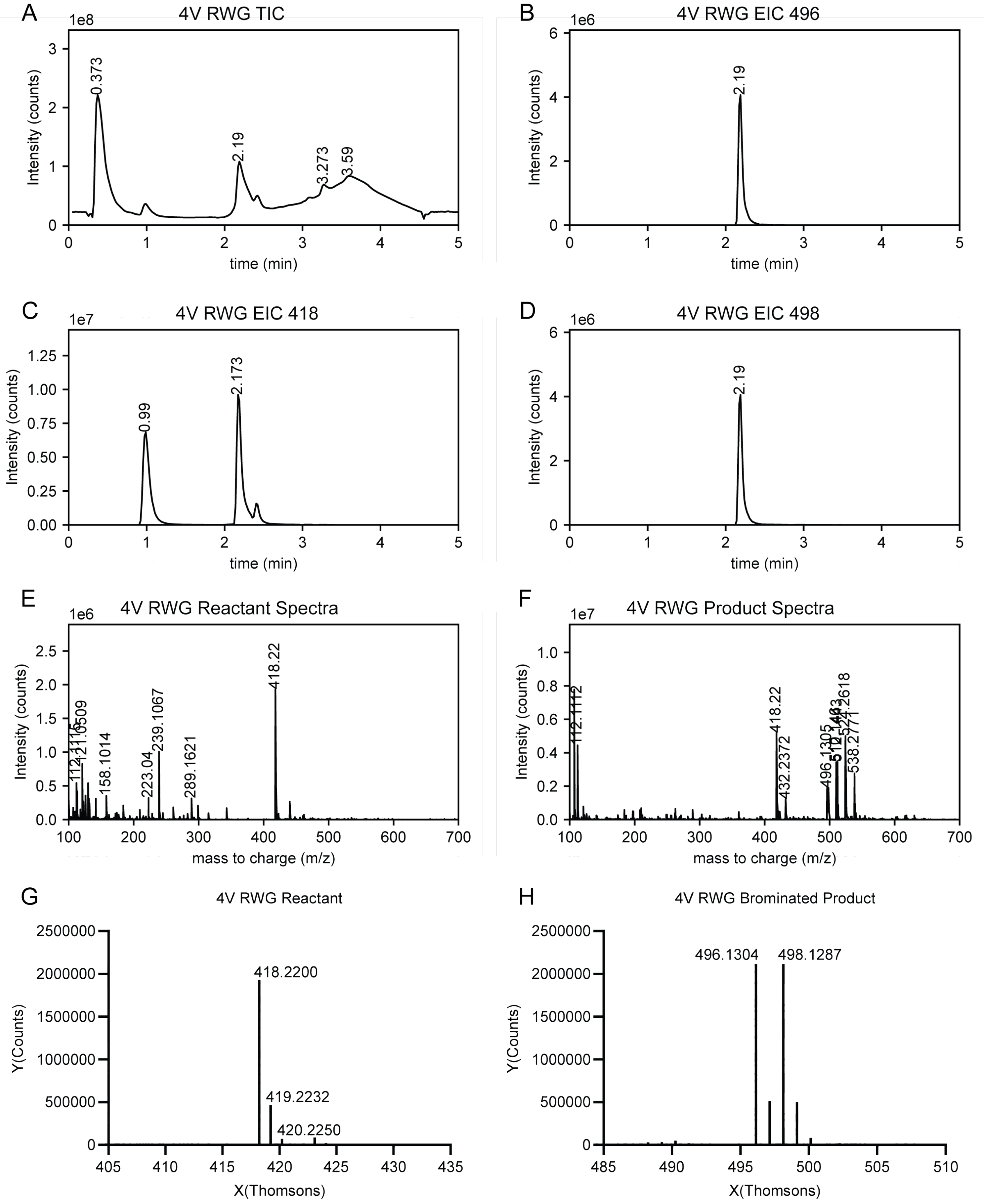

A
4V RWG TIC
1e8
0.373
2.19
3.273
3.59
Intensity (counts)
time (min)
B
4V RWG EIC 496
1e6
2.19
Intensity (counts)
time (min)
C
4V RWG EIC 418
1e7
0.99
2.173
Intensity (counts)
time (min)
D
4V RWG EIC 498
1e6
2.19
Intensity (counts)
time (min)
E
4V RWG Reactant Spectra
1e6
418.22
239.1067
289.1621
223.04
158.1014
121.0509
Intensity (counts)
mass to charge (m/z)
F
4V RWG Product Spectra
1e7
112.1112
418.22
432.2372
496.1305
538.2771
Intensity (counts)
mass to charge (m/z)
G
4V RWG Reactant
418.2200
419.2232
420.2250
Y(Counts)
X(Thomsons)
H
4V RWG Brominated Product
496.1304
498.1287
Y(Counts)
X(Thomsons)

#### 4V TWG

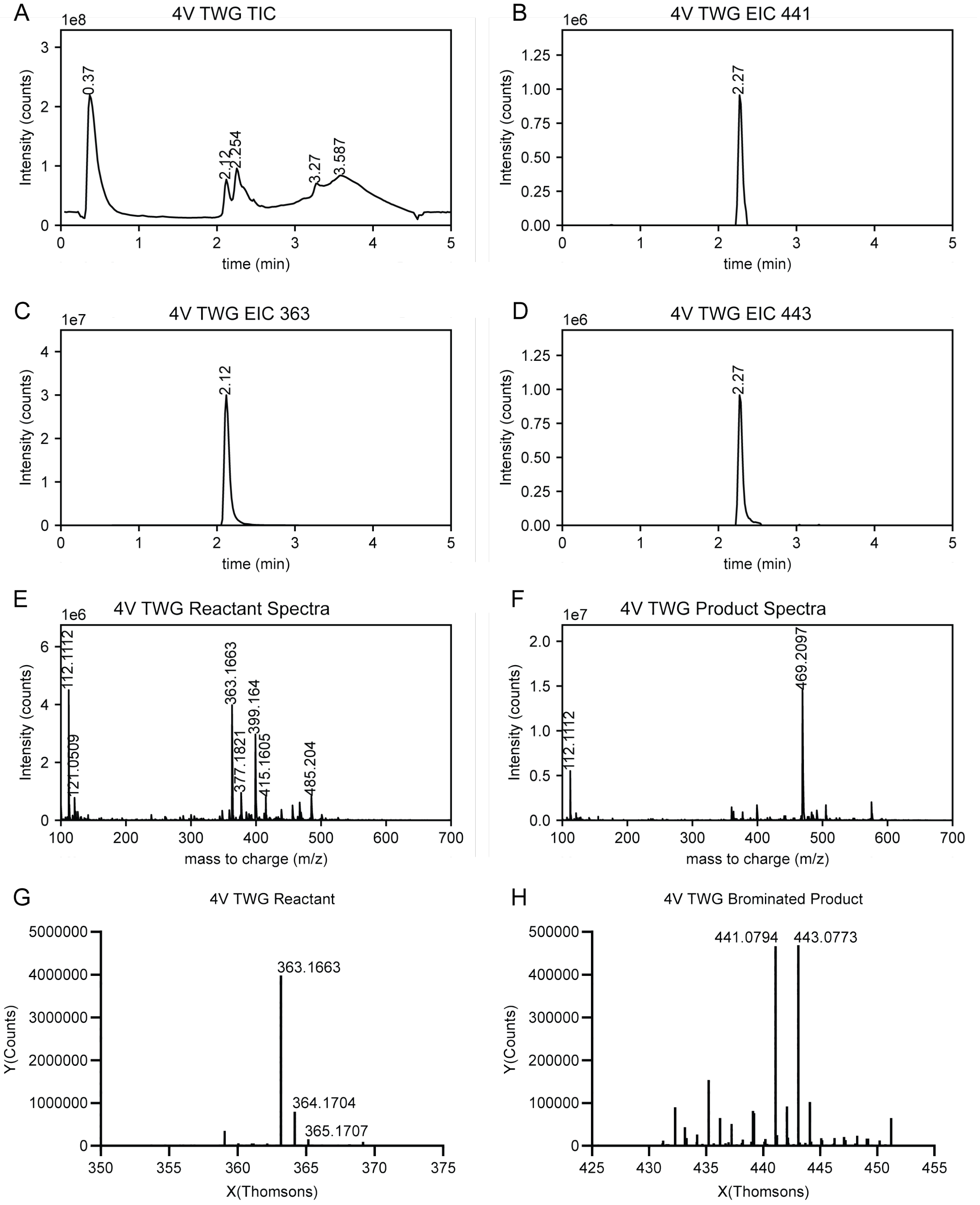
A
4V TWG TIC
1e8
Intensity (counts)
time (min)
0.37
2.12
2.254
3.27
3.587
B
4V TWG EIC 441
1e6
Intensity (counts)
time (min)
2.27
C
4V TWG EIC 363
1e7
Intensity (counts)
time (min)
2.12
D
4V TWG EIC 443
1e6
Intensity (counts)
time (min)
2.27
E
4V TWG Reactant Spectra
1e6
Intensity (counts)
mass to charge (m/z)
112.1112
121.0509
363.1663
377.1821
399.164
415.1605
485.204
F
4V TWG Product Spectra
1e7
Intensity (counts)
mass to charge (m/z)
469.2097
112.1112
G
4V TWG Reactant
Y(Counts)
X(Thomsons)
363.1663
364.1704
365.1707
H
4V TWG Brominated Product
Y(Counts)
X(Thomsons)
441.0794
443.0773

#### 4V VWG

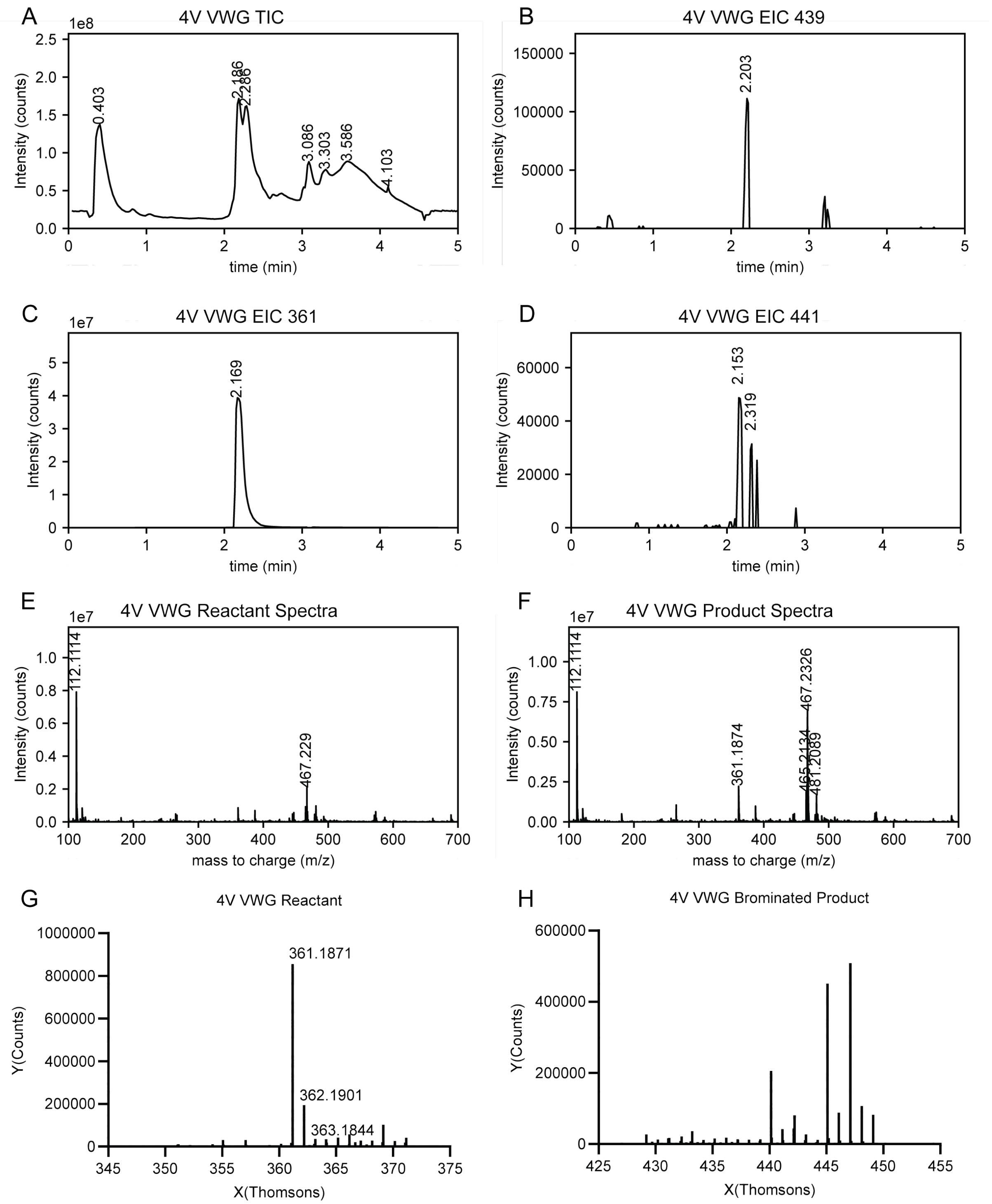

A
4V VWG TIC
1e8
Intensity (counts)
time (min)
0.403
2.186
2.286
3.086
3.303
3.586
4.103
B
4V VWG EIC 439
Intensity (counts)
time (min)
2.203
C
4V VWG EIC 361
1e7
Intensity (counts)
time (min)
2.169
D
4V VWG EIC 441
Intensity (counts)
time (min)
2.153
2.319
E
4V VWG Reactant Spectra
1e7
Intensity (counts)
mass to charge (m/z)
112.1114
467.229
F
4V VWG Product Spectra
1e7
Intensity (counts)
mass to charge (m/z)
112.1114
361.1874
467.2326
465.2134
481.2089
G
4V VWG Reactant
Y(Counts)
X(Thomsons)
361.1871
362.1901
363.1844
H
4V VWG Brominated Product
Y(Counts)
X(Thomsons)

#### 4V YWG

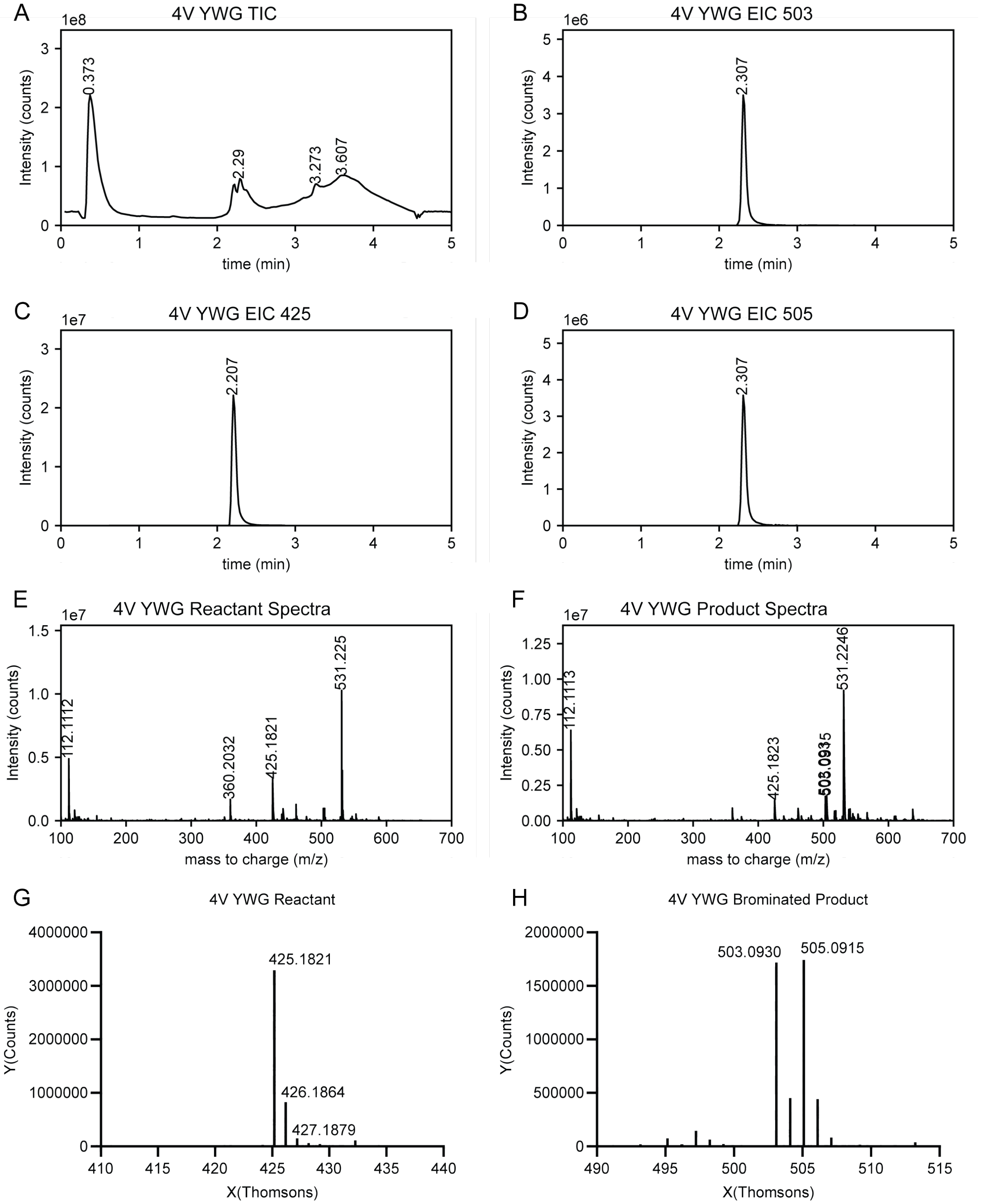

A
4V YWG TIC
1e8
Intensity (counts)
time (min)
0.373
2.29
3.273
3.607
B
4V YWG EIC 503
1e6
Intensity (counts)
time (min)
2.307
C
4V YWG EIC 425
1e7
Intensity (counts)
time (min)
2.207
D
4V YWG EIC 505
1e6
Intensity (counts)
time (min)
2.307
E
4V YWG Reactant Spectra
1e7
Intensity (counts)
mass to charge (m/z)
112.1112
360.2032
425.1821
531.225
F
4V YWG Product Spectra
1e7
Intensity (counts)
mass to charge (m/z)
112.1113
425.1823
531.2246
G
4V YWG Reactant
Y(Counts)
X(Thomsons)
425.1821
426.1864
427.1879
H
4V YWG Brominated Product
Y(Counts)
X(Thomsons)
503.0930
505.0915

#### 4V GWA

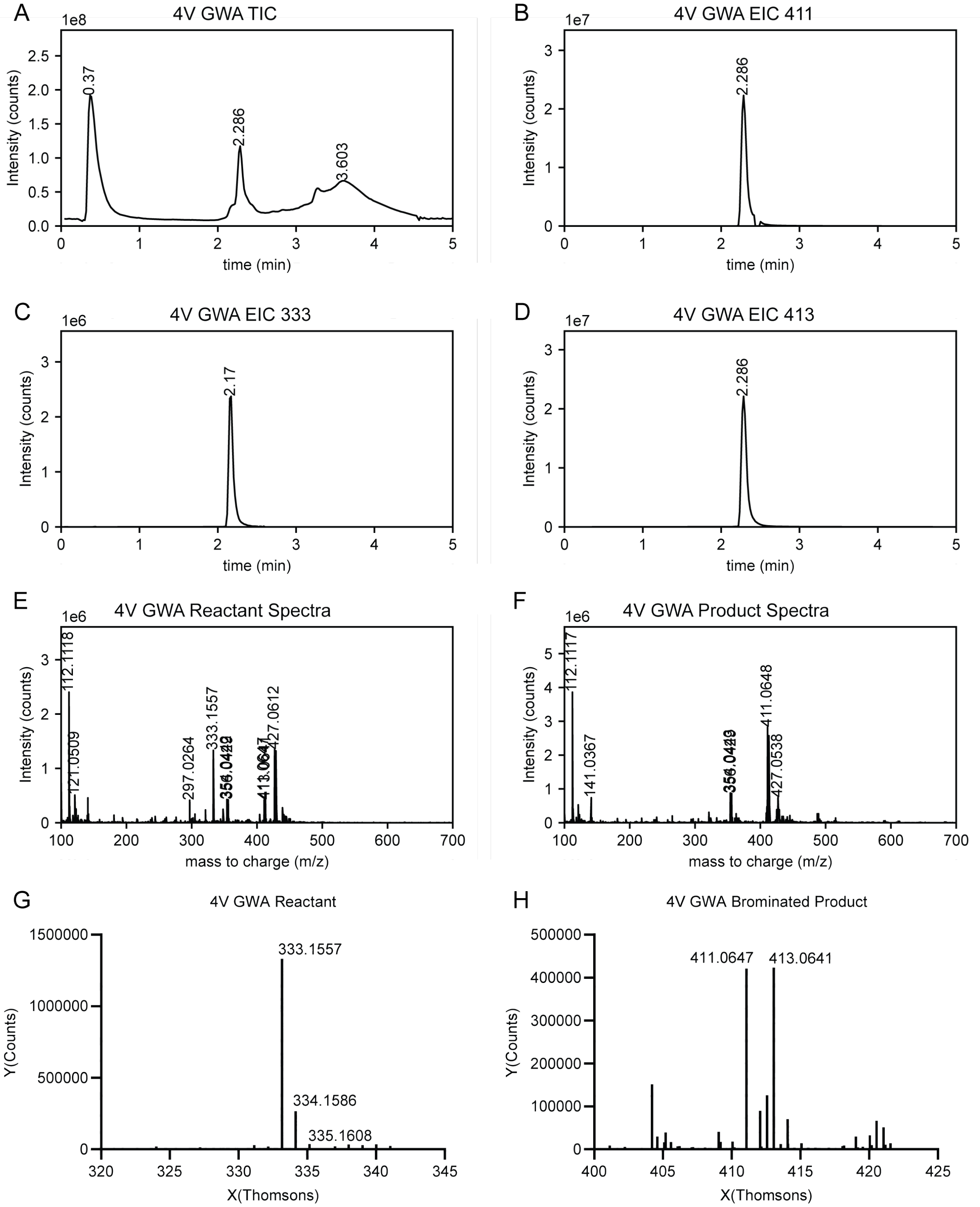

A
4V GWA TIC
1e8
Intensity (counts)
0.37
2.286
3.603
time (min)
B
4V GWA EIC 411
1e7
2.286
C
4V GWA EIC 333
1e6
2.17
D
4V GWA EIC 413
1e7
2.286
E
4V GWA Reactant Spectra
1e6
112.1118
121.0509
297.0264
333.1557
427.0612
mass to charge (m/z)
F
4V GWA Product Spectra
1e6
112.1117
141.0367
411.0648
427.0538
G
4V GWA Reactant
Y(Counts)
333.1557
334.1586
335.1608
X(Thomsons)
H
4V GWA Brominated Product
411.0647
413.0641

**4V GWC**

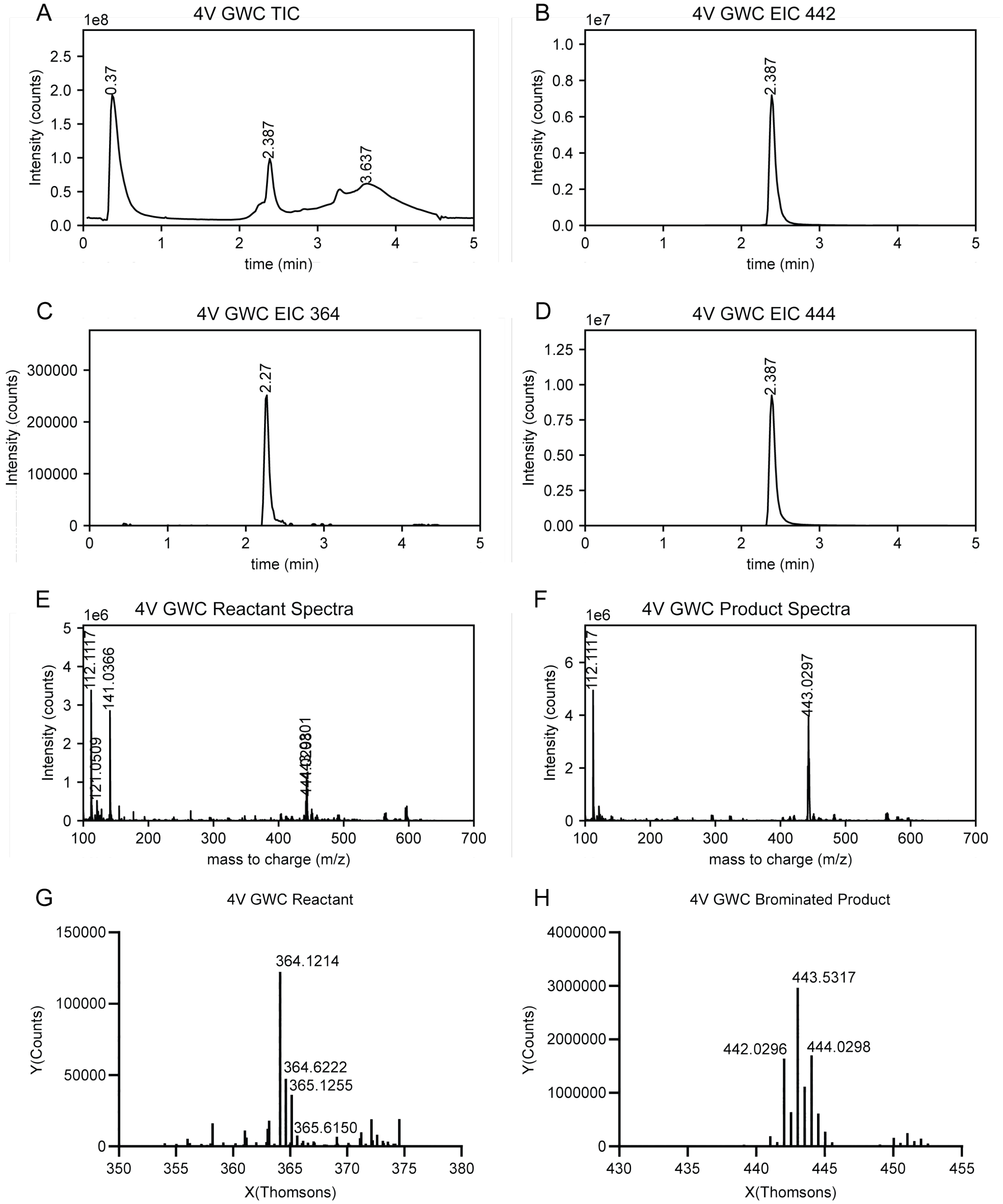
A
4V GWC TIC
1e8
Intensity (counts)
time (min)
0.37
2.387
3.637
B
4V GWC EIC 442
1e7
Intensity (counts)
time (min)
2.387
C
4V GWC EIC 364
Intensity (counts)
time (min)
2.27
D
4V GWC EIC 444
1e7
Intensity (counts)
time (min)
2.387
E
4V GWC Reactant Spectra
1e6
Intensity (counts)
mass to charge (m/z)
112.1117
141.0366
121.0509
F
4V GWC Product Spectra
1e6
Intensity (counts)
mass to charge (m/z)
112.1117
443.0297
G
4V GWC Reactant
Y(Counts)
X(Thomsons)
364.1214
364.6222
365.1255
365.6150
H
4V GWC Brominated Product
Y(Counts)
X(Thomsons)
443.5317
442.0296
444.0298

#### 4V GWD

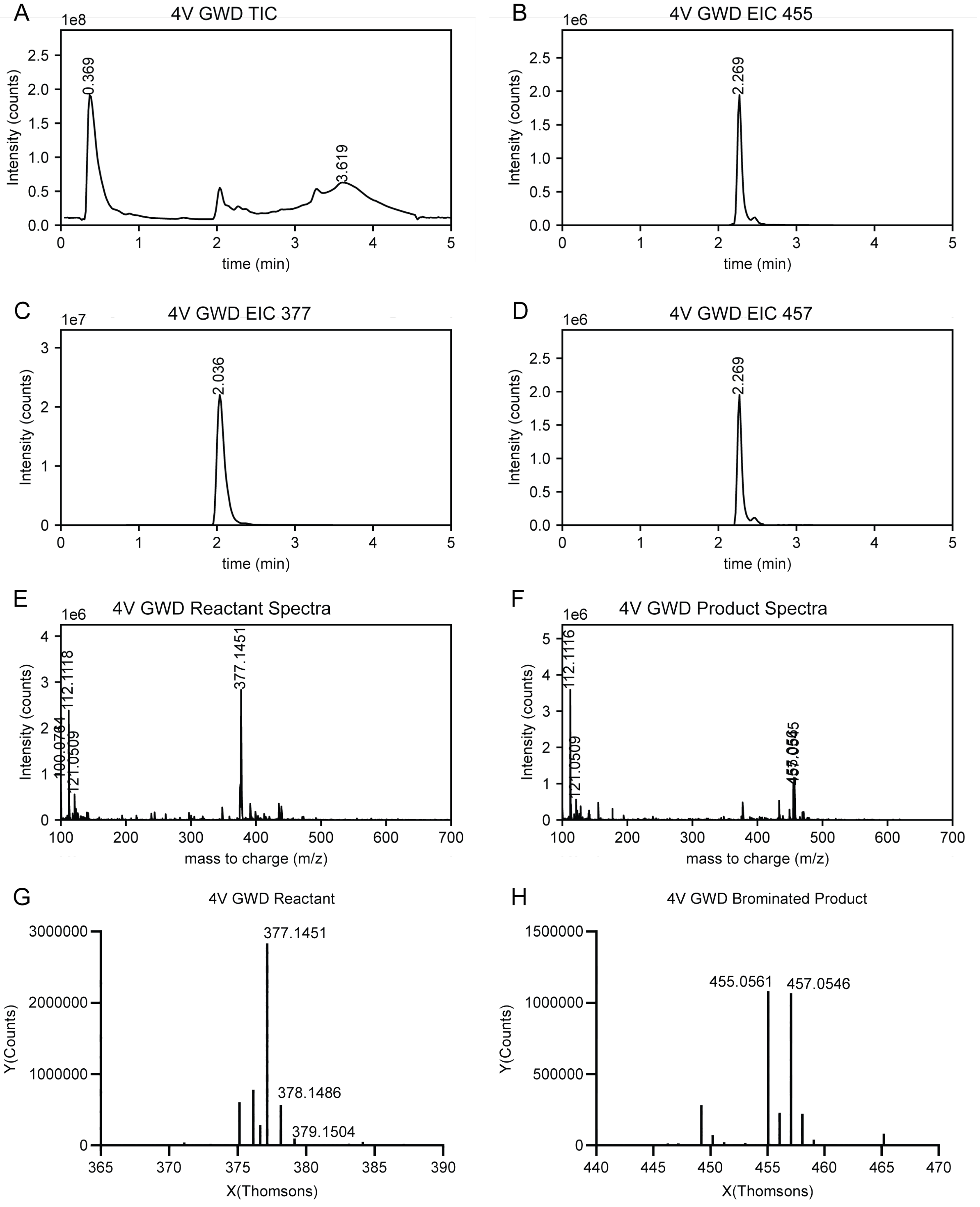
A
4V GWD TIC
1e8
Intensity (counts)
time (min)
0.369
3.619
B
4V GWD EIC 455
1e6
2.269
C
4V GWD EIC 377
1e7
2.036
D
4V GWD EIC 457
1e6
2.269
E
4V GWD Reactant Spectra
mass to charge (m/z)
377.1451
112.1118
121.0509
F
4V GWD Product Spectra
112.1116
121.0509
G
4V GWD Reactant
Y(Counts)
X(Thomsons)
377.1451
378.1486
379.1504
H
4V GWD Brominated Product
455.0561
457.0546

#### 4V GWE

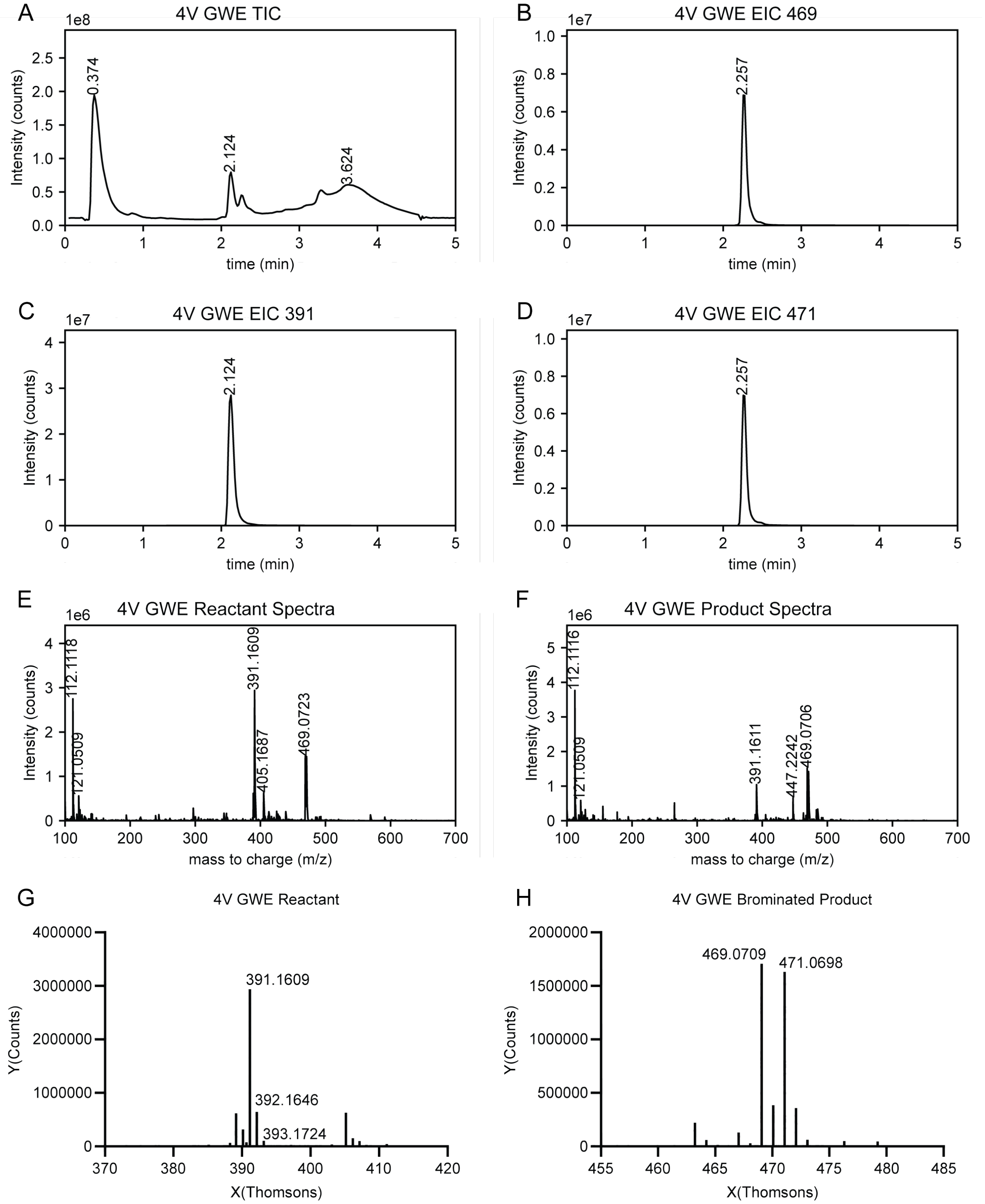

A
4V GWE TIC
B
4V GWE EIC 469
C
4V GWE EIC 391
D
4V GWE EIC 471
E
4V GWE Reactant Spectra
F
4V GWE Product Spectra
G
4V GWE Reactant
H
4V GWE Brominated Product
Intensity (counts)
time (min)
mass to charge (m/z)
Y(Counts)
X(Thomsons)
0.374
2.124
3.624
2.257
112.1118
121.0509
391.1609
405.1687
469.0723
112.1116
391.1611
447.2242
469.0706
392.1646
393.1724
469.0709
471.0698

**4V GWF**

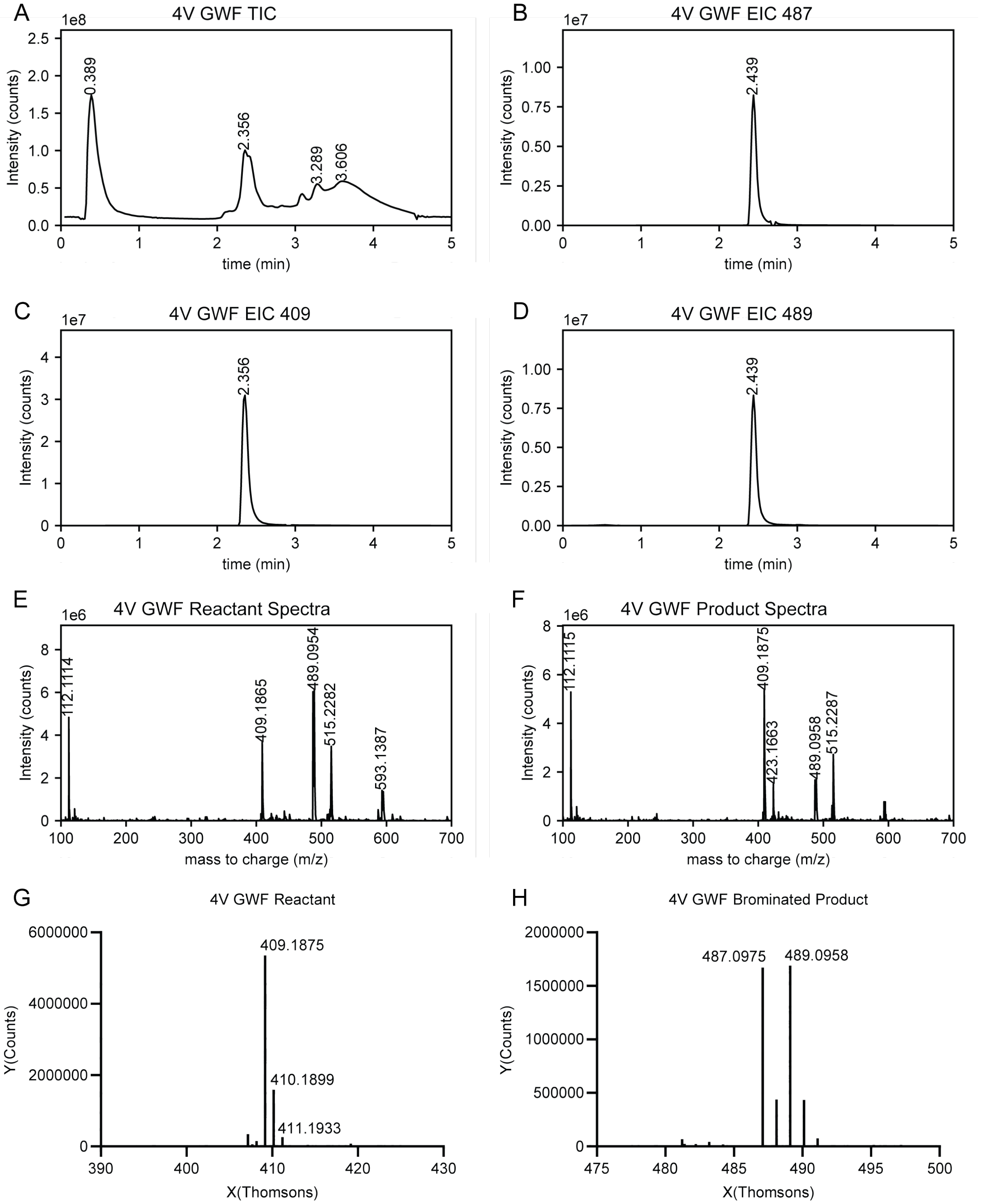
A
4V GWF TIC
1e8
0.389
2.356
3.289
3.606
Intensity (counts)
time (min)
B
4V GWF EIC 487
1e7
2.439
Intensity (counts)
time (min)
C
4V GWF EIC 409
1e7
2.356
Intensity (counts)
time (min)
D
4V GWF EIC 489
1e7
2.439
Intensity (counts)
time (min)
E
4V GWF Reactant Spectra
1e6
112.1114
409.1865
489.0954
515.2282
593.1387
Intensity (counts)
mass to charge (m/z)
F
4V GWF Product Spectra
1e6
112.1115
409.1875
423.1663
489.0958
515.2287
Intensity (counts)
mass to charge (m/z)
G
4V GWF Reactant
409.1875
410.1899
411.1933
Y(Counts)
X(Thomsons)
H
4V GWF Brominated Product
487.0975
489.0958
Y(Counts)
X(Thomsons)

#### 4V GWH

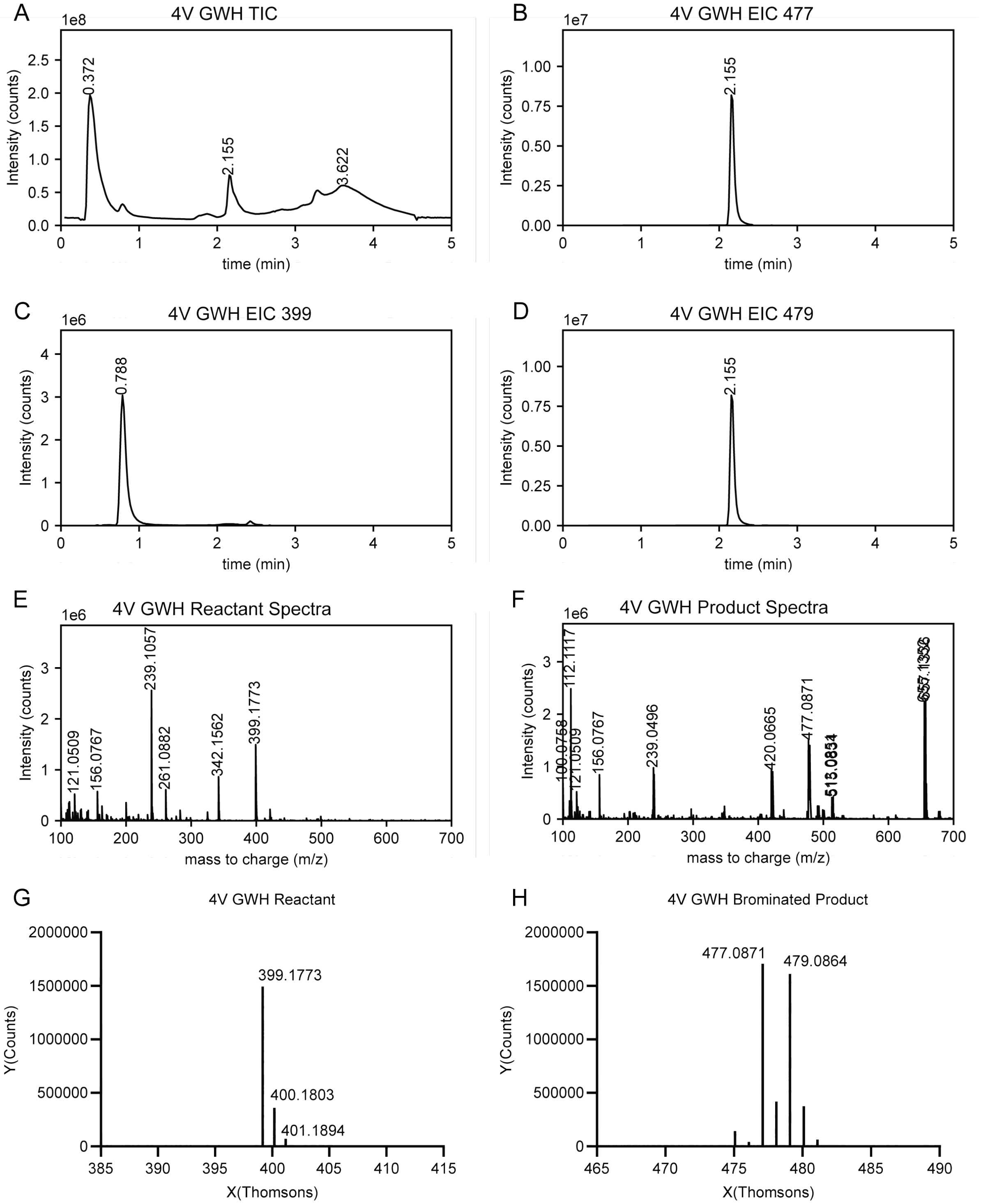

A
4V GWH TIC
1e8
Intensity (counts)
time (min)
0.372
2.155
3.622
B
4V GWH EIC 477
1e7
Intensity (counts)
time (min)
2.155
C
4V GWH EIC 399
1e6
Intensity (counts)
time (min)
0.788
D
4V GWH EIC 479
1e7
Intensity (counts)
time (min)
2.155
E
4V GWH Reactant Spectra
1e6
Intensity (counts)
mass to charge (m/z)
121.0509
156.0767
239.1057
261.0882
342.1562
399.1773
F
4V GWH Product Spectra
1e6
Intensity (counts)
mass to charge (m/z)
100.0758
112.1117
121.0509
156.0767
239.0496
420.0665
477.0871
G
4V GWH Reactant
Y(Counts)
X(Thomsons)
399.1773
400.1803
401.1894
H
4V GWH Brominated Product
Y(Counts)
X(Thomsons)
477.0871
479.0864

### 4V GWK

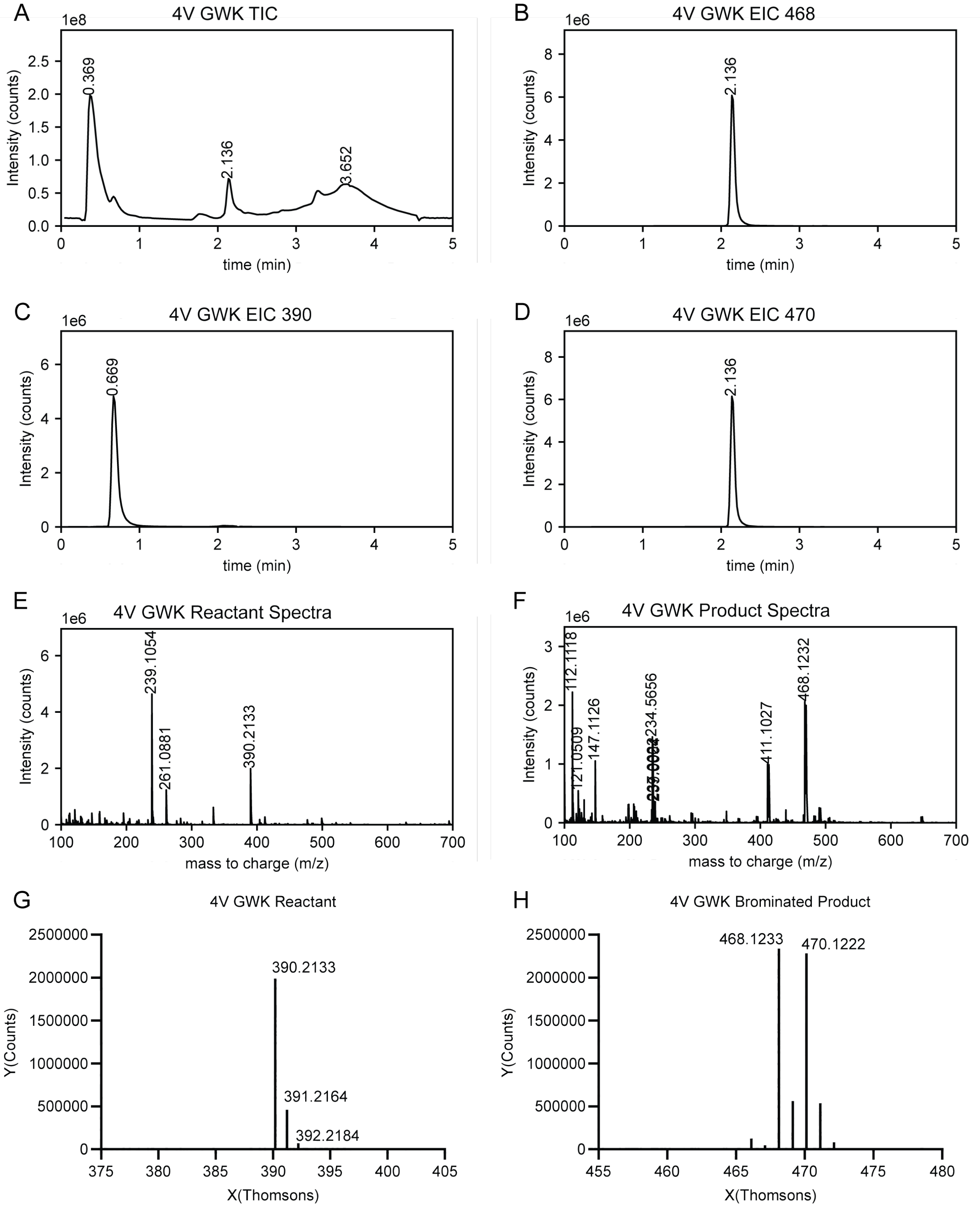
A
4V GWK TIC
1e8
Intensity (counts)
time (min)
0.369
2.136
3.652
B
4V GWK EIC 468
1e6
Intensity (counts)
time (min)
2.136
C
4V GWK EIC 390
1e6
Intensity (counts)
time (min)
0.669
D
4V GWK EIC 470
1e6
Intensity (counts)
time (min)
2.136
E
4V GWK Reactant Spectra
1e6
Intensity (counts)
mass to charge (m/z)
239.1054
261.0881
390.2133
F
4V GWK Product Spectra
1e6
Intensity (counts)
mass to charge (m/z)
112.1118
121.0509
147.1126
234.5656
411.1027
468.1232
G
4V GWK Reactant
Y(Counts)
X(Thomsons)
390.2133
391.2164
392.2184
H
4V GWK Brominated Product
Y(Counts)
X(Thomsons)
468.1233
470.1222

#### 4V GWL

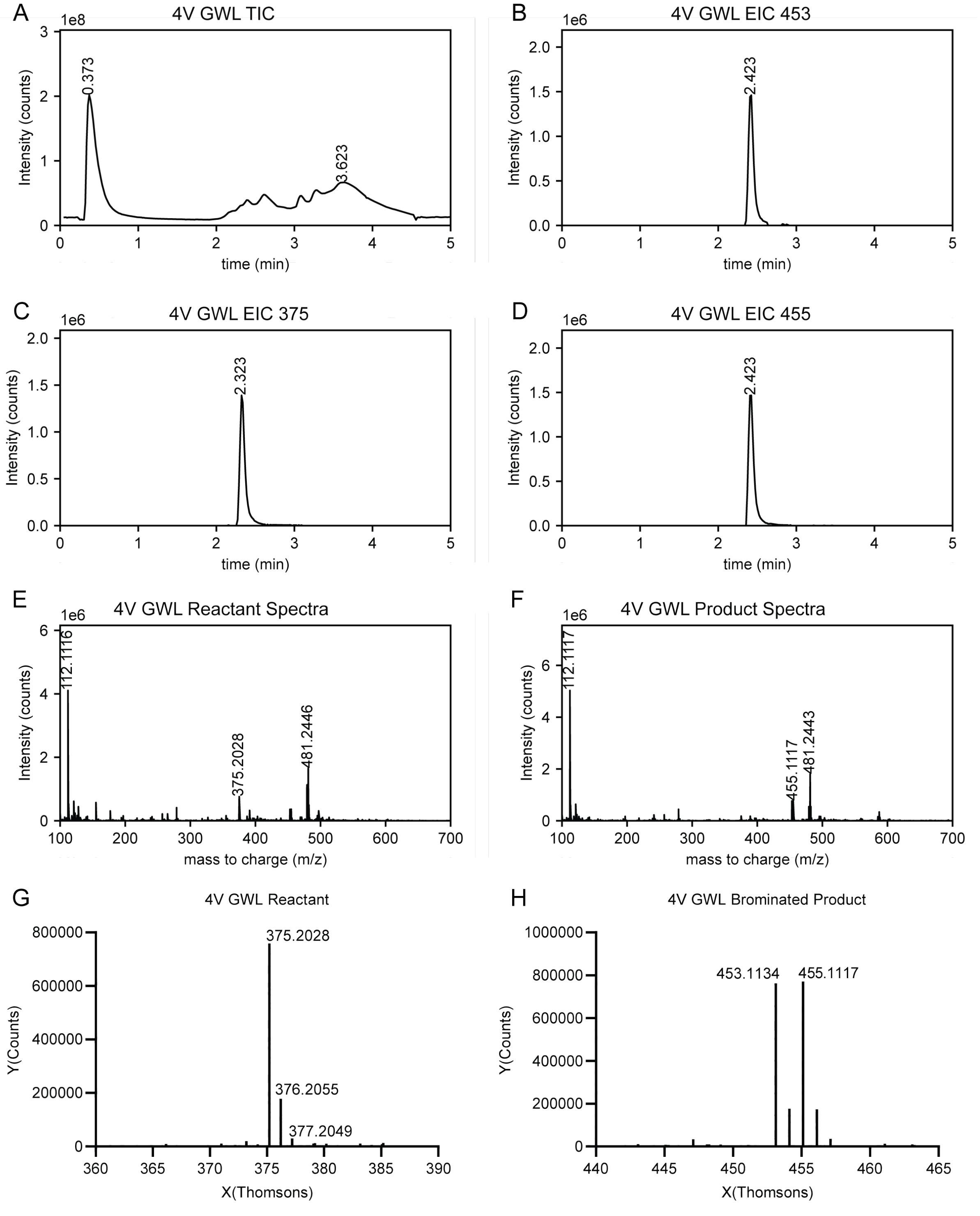

A
4V GWL TIC
1e8
Intensity (counts)
time (min)
0.373
3.623
B
4V GWL EIC 453
1e6
2.423
C
4V GWL EIC 375
1e6
2.323
D
4V GWL EIC 455
1e6
2.423
E
4V GWL Reactant Spectra
1e6
mass to charge (m/z)
112.1116
375.2028
481.2446
F
4V GWL Product Spectra
1e6
112.1117
455.1117
481.2443
G
4V GWL Reactant
Y(Counts)
X(Thomsons)
375.2028
376.2055
377.2049
H
4V GWL Brominated Product
453.1134
455.1117

#### 4V GWM

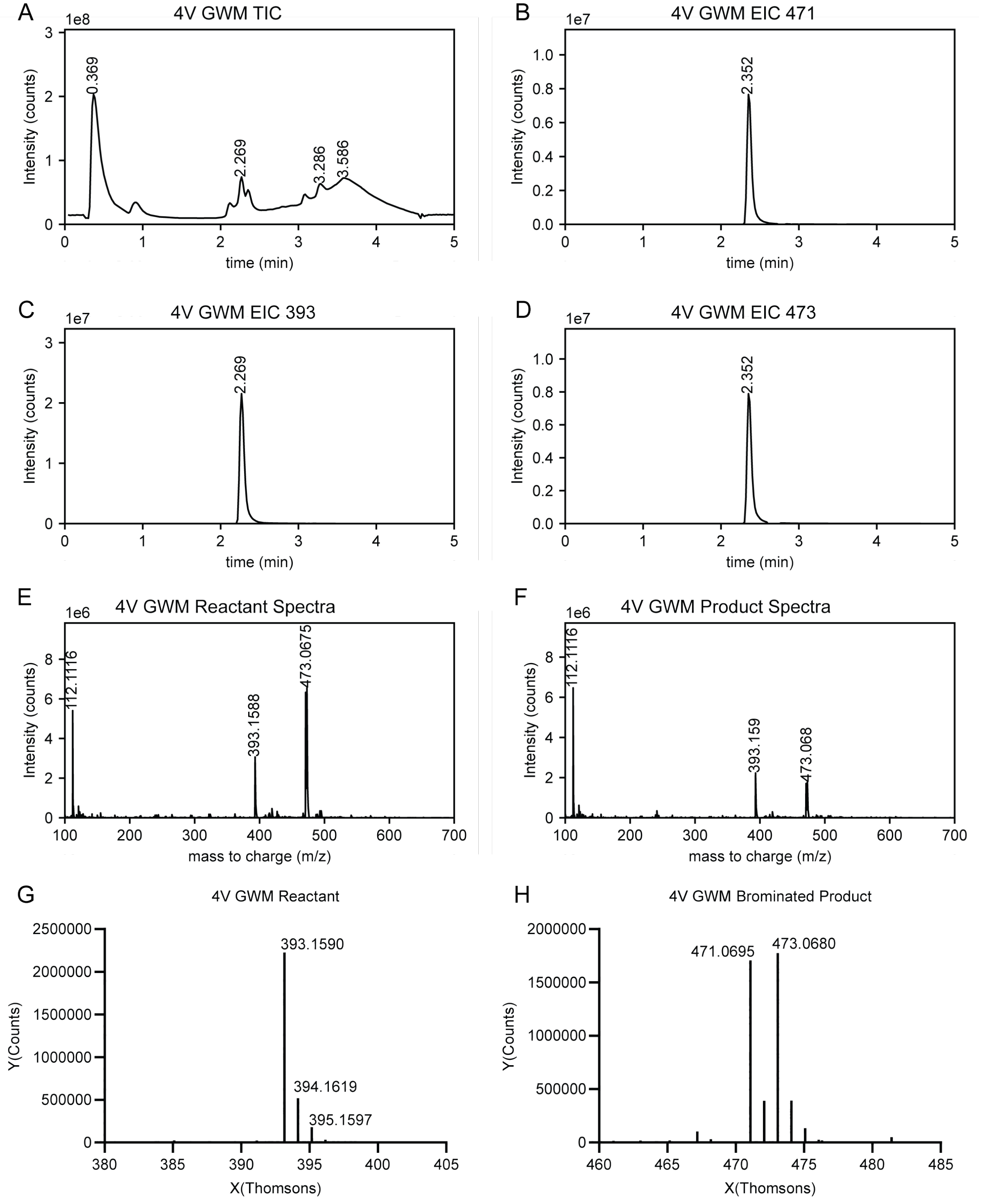
A
4V GWM TIC
1e8
Intensity (counts)
time (min)
0.369
2.269
3.286
3.586
B
4V GWM EIC 471
1e7
Intensity (counts)
time (min)
2.352
C
4V GWM EIC 393
1e7
Intensity (counts)
time (min)
2.269
D
4V GWM EIC 473
1e7
Intensity (counts)
time (min)
2.352
E
4V GWM Reactant Spectra
1e6
Intensity (counts)
mass to charge (m/z)
112.1116
393.1588
473.0675
F
4V GWM Product Spectra
1e6
Intensity (counts)
mass to charge (m/z)
112.1116
393.159
473.068
G
4V GWM Reactant
Y(Counts)
X(Thomsons)
393.1590
394.1619
395.1597
H
4V GWM Brominated Product
Y(Counts)
X(Thomsons)
471.0695
473.0680

#### 4V GWN

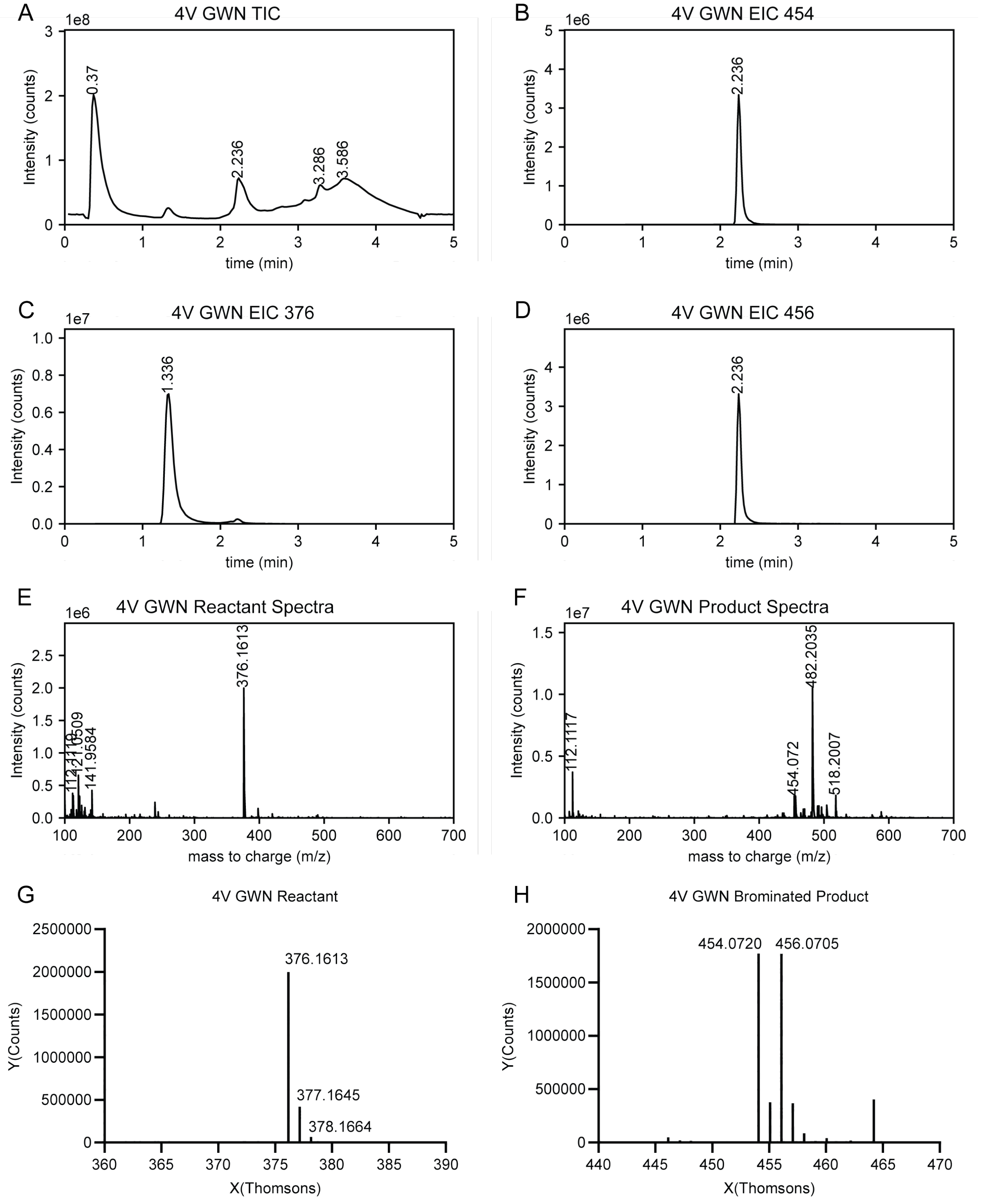

A
4V GWN TIC
1e8
Intensity (counts)
time (min)
0.37
2.236
3.286
3.586
B
4V GWN EIC 454
1e6
Intensity (counts)
time (min)
2.236
C
4V GWN EIC 376
1e7
Intensity (counts)
time (min)
1.336
D
4V GWN EIC 456
1e6
Intensity (counts)
time (min)
2.236
E
4V GWN Reactant Spectra
1e6
Intensity (counts)
mass to charge (m/z)
376.1613
121.0509
141.9584
F
4V GWN Product Spectra
1e7
Intensity (counts)
mass to charge (m/z)
482.2035
112.1117
454.072
518.2007
G
4V GWN Reactant
Y(Counts)
X(Thomsons)
376.1613
377.1645
378.1664
H
4V GWN Brominated Product
Y(Counts)
X(Thomsons)
454.0720
456.0705

#### 4V GWP

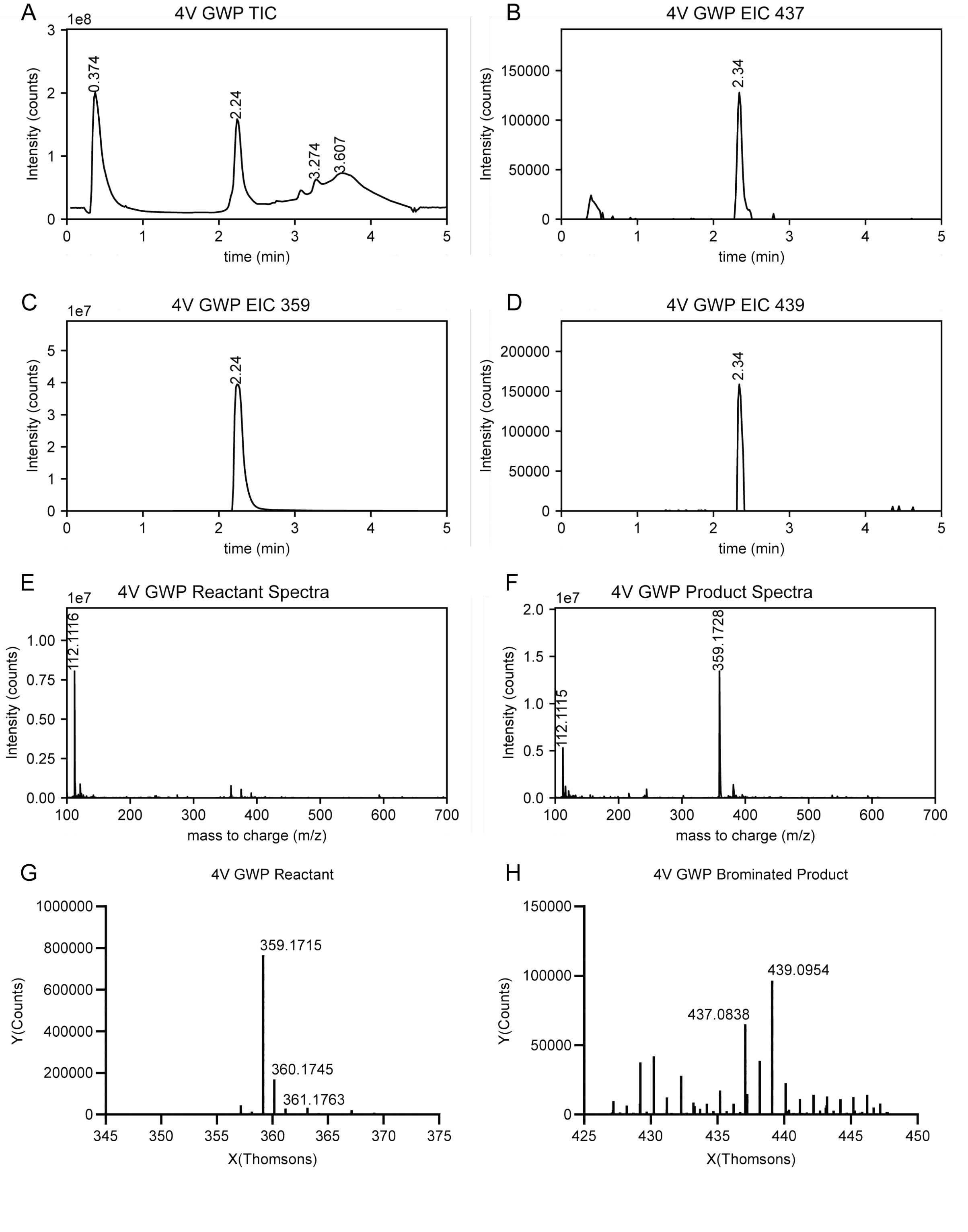

A
4V GWP TIC
1e8
Intensity (counts)
time (min)
0.374
2.24
3.274
3.607
B
4V GWP EIC 437
Intensity (counts)
time (min)
2.34
C
4V GWP EIC 359
1e7
Intensity (counts)
time (min)
2.24
D
4V GWP EIC 439
Intensity (counts)
time (min)
2.34
E
4V GWP Reactant Spectra
1e7
Intensity (counts)
mass to charge (m/z)
112.1116
F
4V GWP Product Spectra
1e7
Intensity (counts)
mass to charge (m/z)
112.1115
359.1728
G
4V GWP Reactant
Y(Counts)
X(Thomsons)
359.1715
360.1745
361.1763
H
4V GWP Brominated Product
Y(Counts)
X(Thomsons)
437.0838
439.0954

#### 4V GWQ

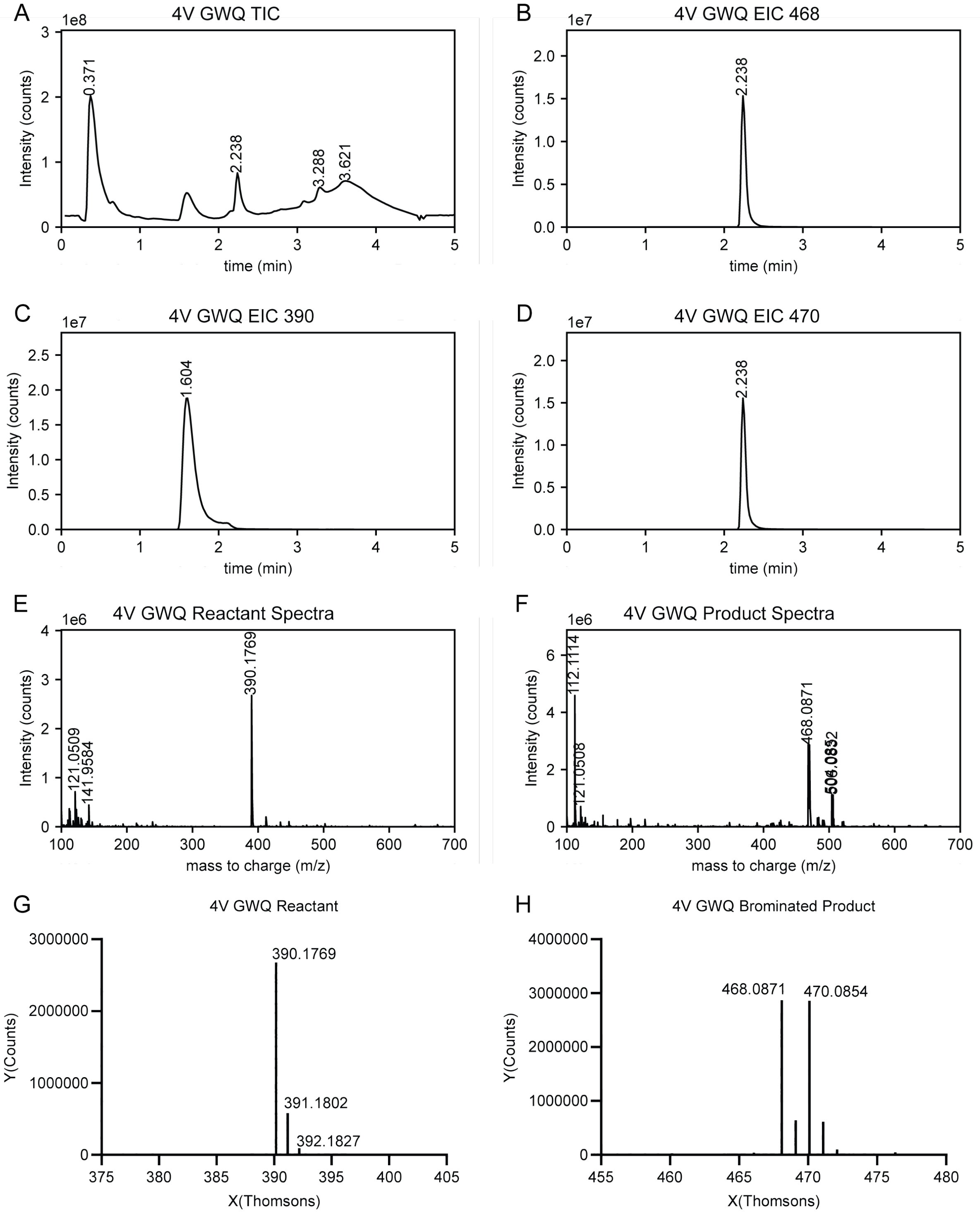
A
4V GWQ TIC
1e8
Intensity (counts)
time (min)
0.371
2.238
3.288
3.621
B
4V GWQ EIC 468
1e7
Intensity (counts)
time (min)
2.238
C
4V GWQ EIC 390
1e7
Intensity (counts)
time (min)
1.604
D
4V GWQ EIC 470
1e7
Intensity (counts)
time (min)
2.238
E
4V GWQ Reactant Spectra
1e6
Intensity (counts)
mass to charge (m/z)
390.1769
121.0509
141.9584
F
4V GWQ Product Spectra
1e6
Intensity (counts)
mass to charge (m/z)
112.1114
121.0508
468.0871
G
4V GWQ Reactant
Y(Counts)
X(Thomsons)
390.1769
391.1802
392.1827
H
4V GWQ Brominated Product
Y(Counts)
X(Thomsons)
468.0871
470.0854

#### 4V GWR

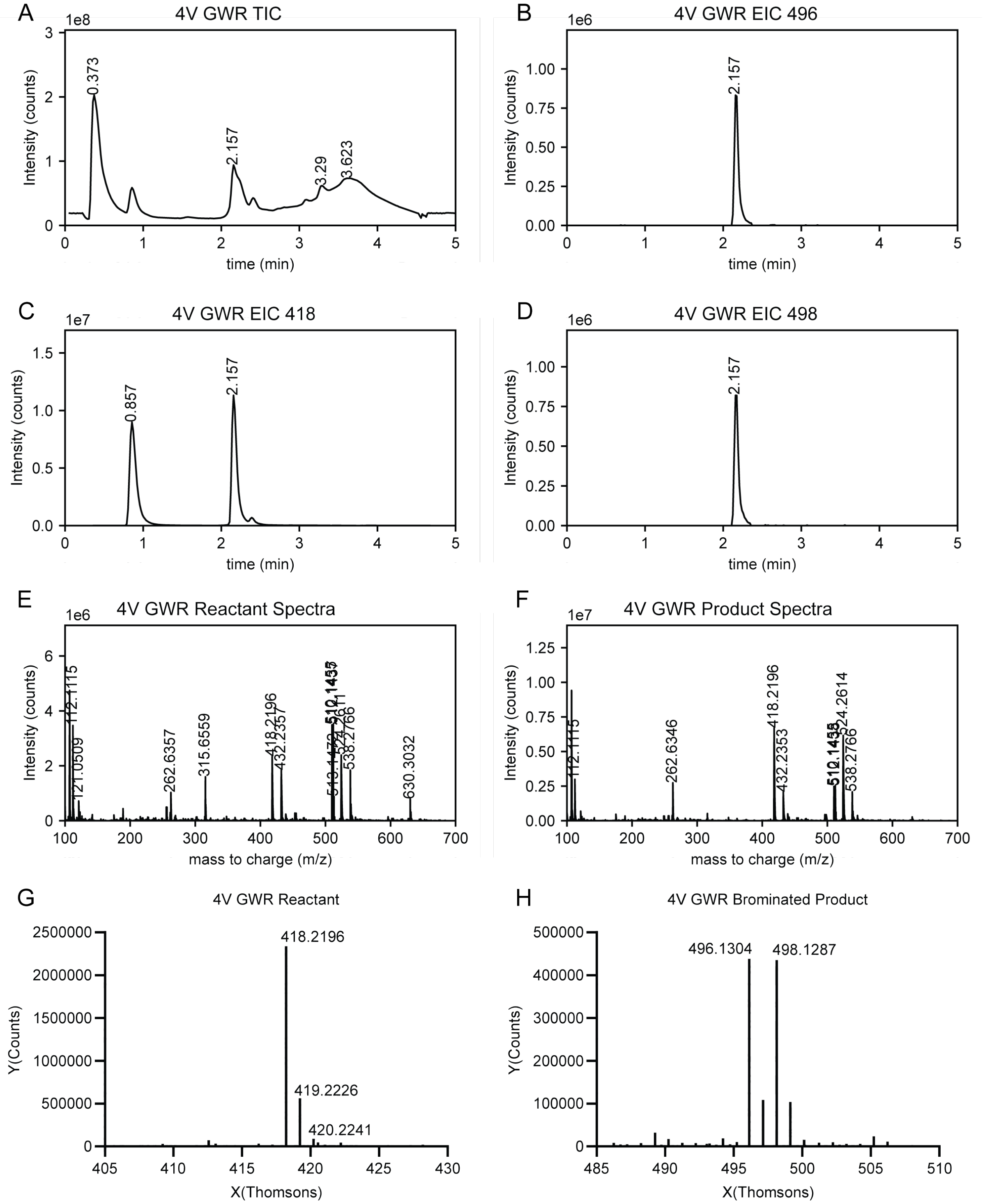

A
4V GWR TIC
1e8
Intensity (counts)
time (min)
0.373
2.157
3.29
3.623
B
4V GWR EIC 496
1e6
Intensity (counts)
time (min)
2.157
C
4V GWR EIC 418
1e7
Intensity (counts)
time (min)
0.857
2.157
D
4V GWR EIC 498
1e6
Intensity (counts)
time (min)
2.157
E
4V GWR Reactant Spectra
1e6
Intensity (counts)
mass to charge (m/z)
112.1115
121.0509
262.6357
315.6559
418.2196
432.2357
538.2766
630.3032
F
4V GWR Product Spectra
1e7
Intensity (counts)
mass to charge (m/z)
112.1115
262.6346
418.2196
432.2353
524.2614
538.2766
G
4V GWR Reactant
Y(Counts)
X(Thomsons)
418.2196
419.2226
420.2241
H
4V GWR Brominated Product
Y(Counts)
X(Thomsons)
496.1304
498.1287

**4V GWT**

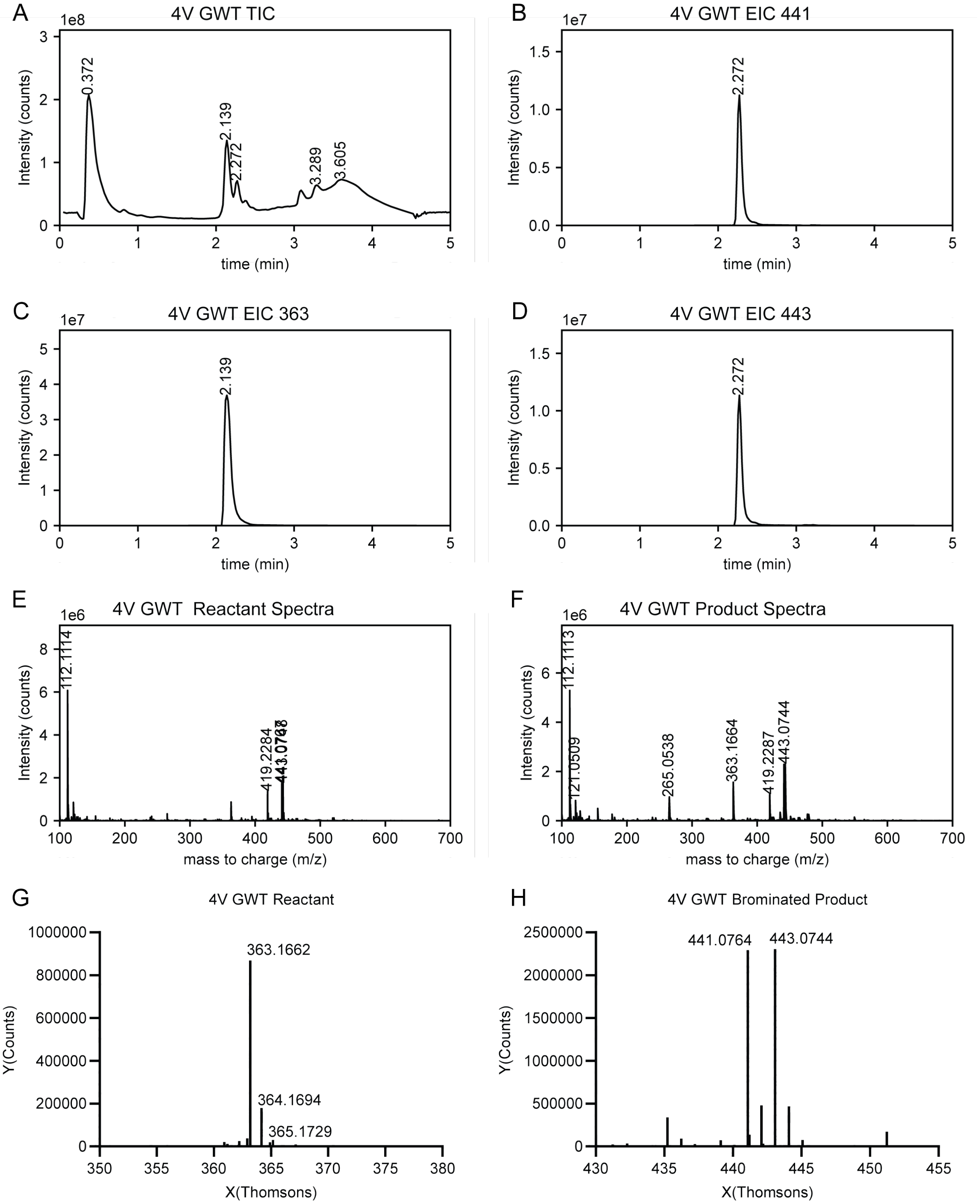

A
4V GWT TIC
1e8
Intensity (counts)
time (min)
0.372
2.139
2.272
3.289
3.605
B
4V GWT EIC 441
1e7
Intensity (counts)
time (min)
2.272
C
4V GWT EIC 363
1e7
Intensity (counts)
time (min)
2.139
D
4V GWT EIC 443
1e7
Intensity (counts)
time (min)
2.272
E
4V GWT Reactant Spectra
1e6
Intensity (counts)
mass to charge (m/z)
112.1114
419.2284
F
4V GWT Product Spectra
1e6
Intensity (counts)
mass to charge (m/z)
112.1113
121.0509
265.0538
363.1664
419.2287
443.0744
G
4V GWT Reactant
Y(Counts)
X(Thomsons)
363.1662
364.1694
365.1729
H
4V GWT Brominated Product
Y(Counts)
X(Thomsons)
441.0764
443.0744

#### 4V GWV

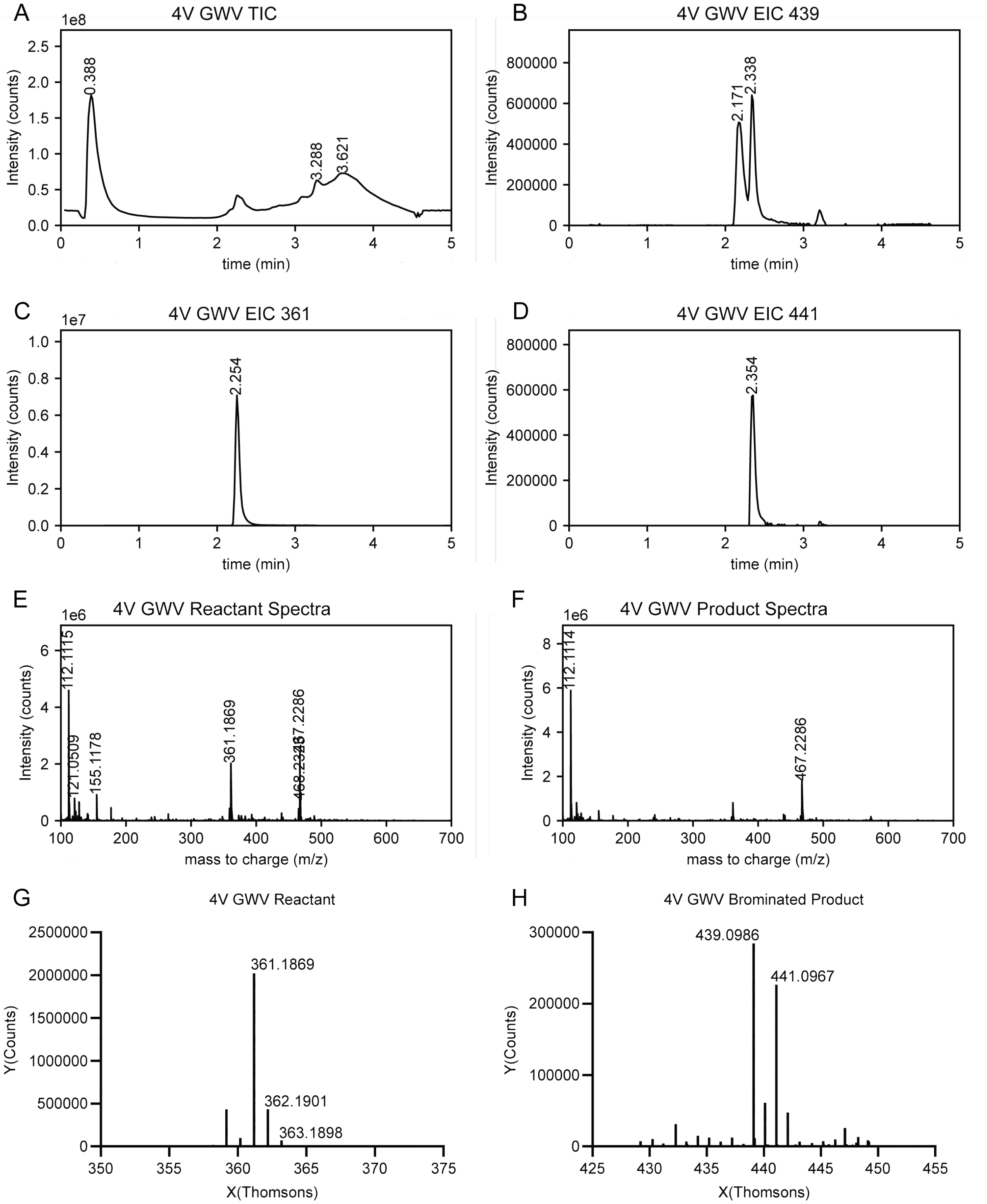
A
4V GWV TIC
1e8
Intensity (counts)
time (min)
0.388
3.288
3.621
B
4V GWV EIC 439
Intensity (counts)
time (min)
2.171
2.338
C
4V GWV EIC 361
1e7
Intensity (counts)
time (min)
2.254
D
4V GWV EIC 441
Intensity (counts)
time (min)
2.354
E
4V GWV Reactant Spectra
1e6
Intensity (counts)
mass to charge (m/z)
112.1115
121.0509
155.1178
361.1869
467.2286
F
4V GWV Product Spectra
1e6
Intensity (counts)
mass to charge (m/z)
112.1114
467.2286
G
4V GWV Reactant
Y(Counts)
X(Thomsons)
361.1869
362.1901
363.1898
H
4V GWV Brominated Product
Y(Counts)
X(Thomsons)
439.0986
441.0967

#### 4V GWY

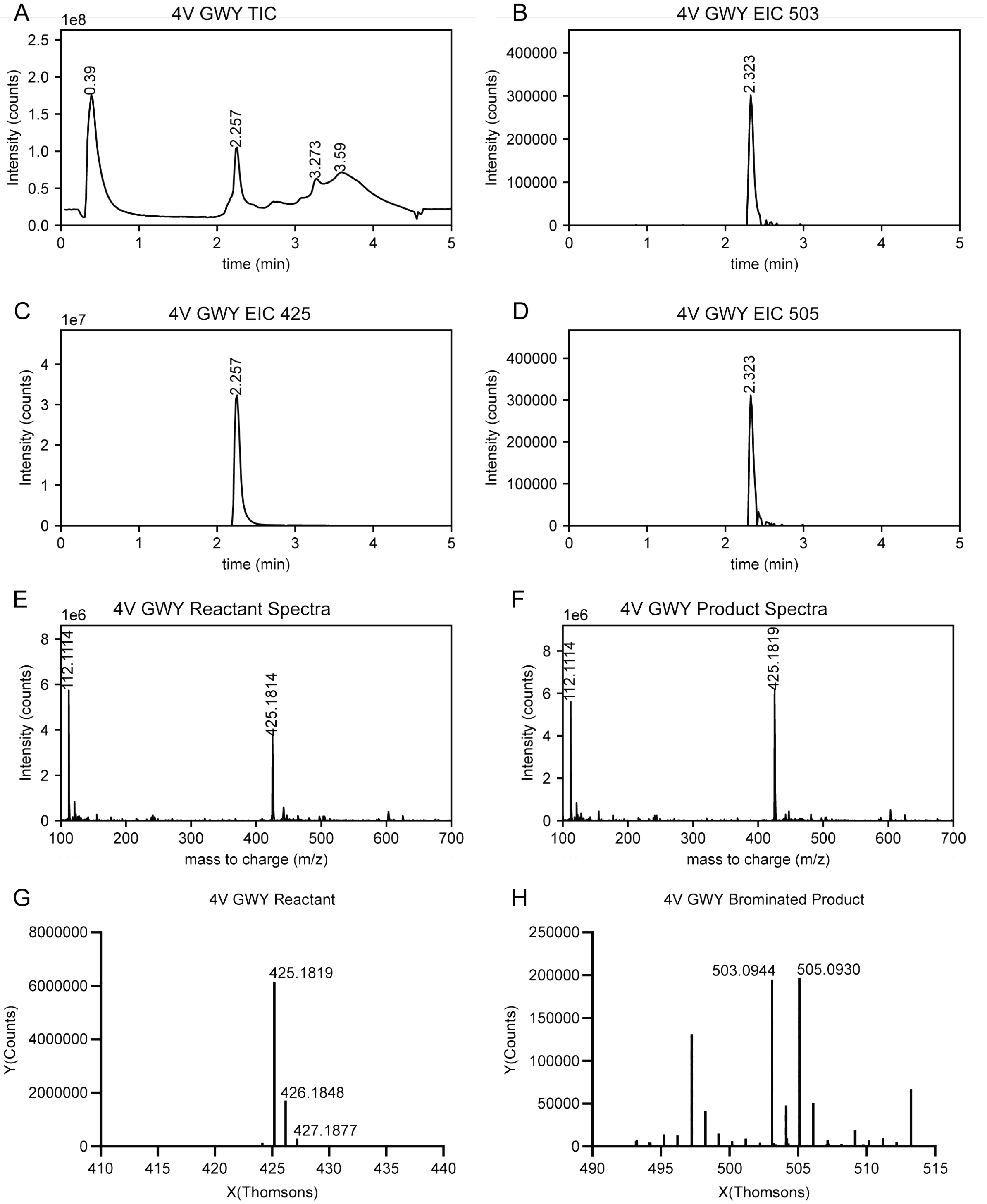

A
4V GWY TIC
1e8
Intensity (counts)
time (min)
0.39
2.257
3.273
3.59
B
4V GWY EIC 503
Intensity (counts)
time (min)
2.323
C
4V GWY EIC 425
1e7
Intensity (counts)
time (min)
2.257
D
4V GWY EIC 505
Intensity (counts)
time (min)
2.323
E
4V GWY Reactant Spectra
1e6
Intensity (counts)
mass to charge (m/z)
112.1114
425.1814
F
4V GWY Product Spectra
1e6
Intensity (counts)
mass to charge (m/z)
112.1114
425.1819
G
4V GWY Reactant
Y(Counts)
X(Thomsons)
425.1819
426.1848
427.1877
H
4V GWY Brominated Product
Y(Counts)
X(Thomsons)
503.0944
505.0930

#### 4V AGW

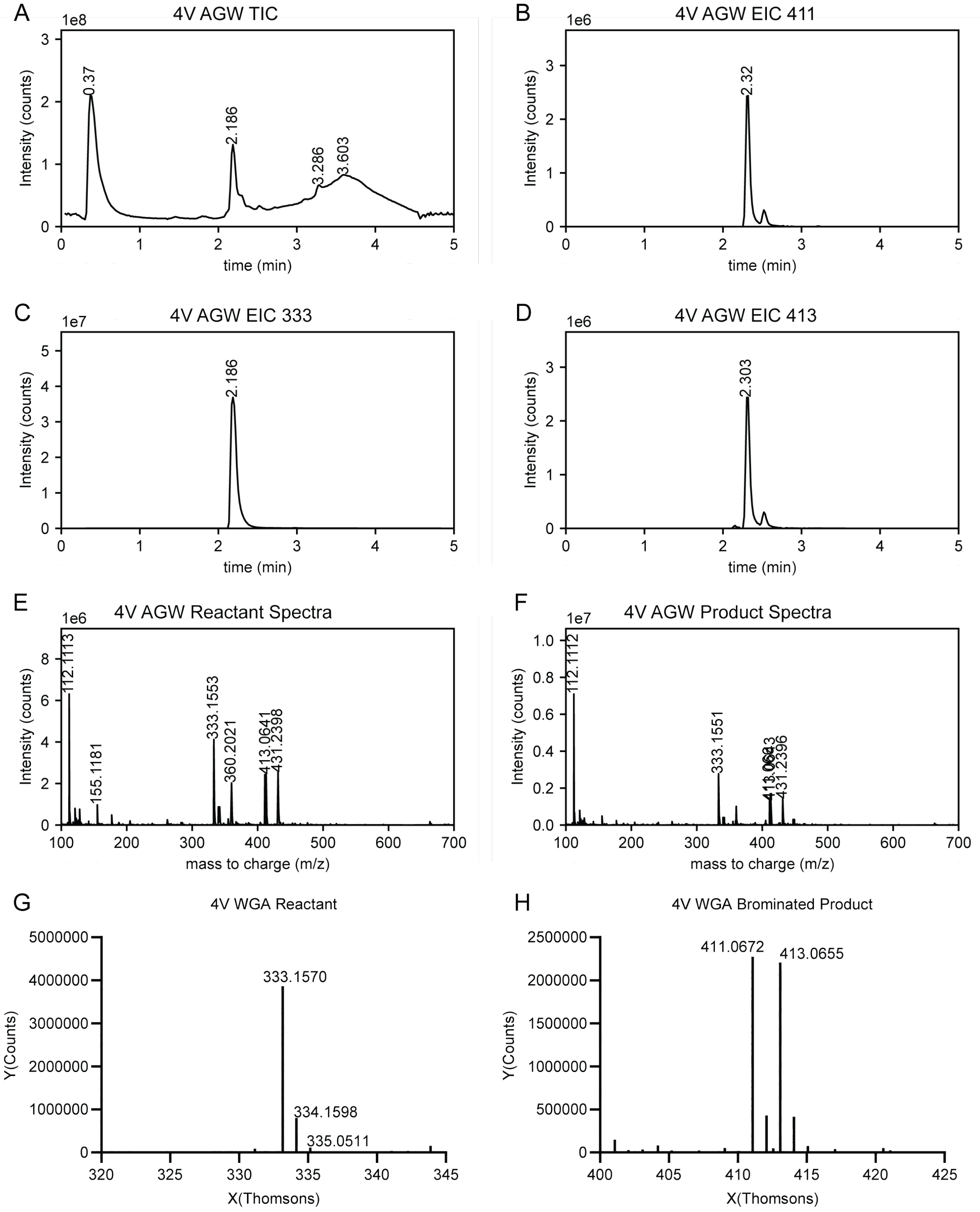

A
4V AGW TIC
1e8
Intensity (counts)
time (min)
0.37
2.186
3.286
3.603
B
4V AGW EIC 411
1e6
Intensity (counts)
time (min)
2.32
C
4V AGW EIC 333
1e7
Intensity (counts)
time (min)
2.186
D
4V AGW EIC 413
1e6
Intensity (counts)
time (min)
2.303
E
4V AGW Reactant Spectra
1e6
Intensity (counts)
mass to charge (m/z)
112.1113
155.1181
333.1553
360.2021
413.0641
431.2398
F
4V AGW Product Spectra
1e7
Intensity (counts)
mass to charge (m/z)
112.1112
333.1551
431.2396
G
4V WGA Reactant
Y(Counts)
X(Thomsons)
333.1570
334.1598
335.0511
H
4V WGA Brominated Product
Y(Counts)
X(Thomsons)
411.0672
413.0655

#### 4V CGW

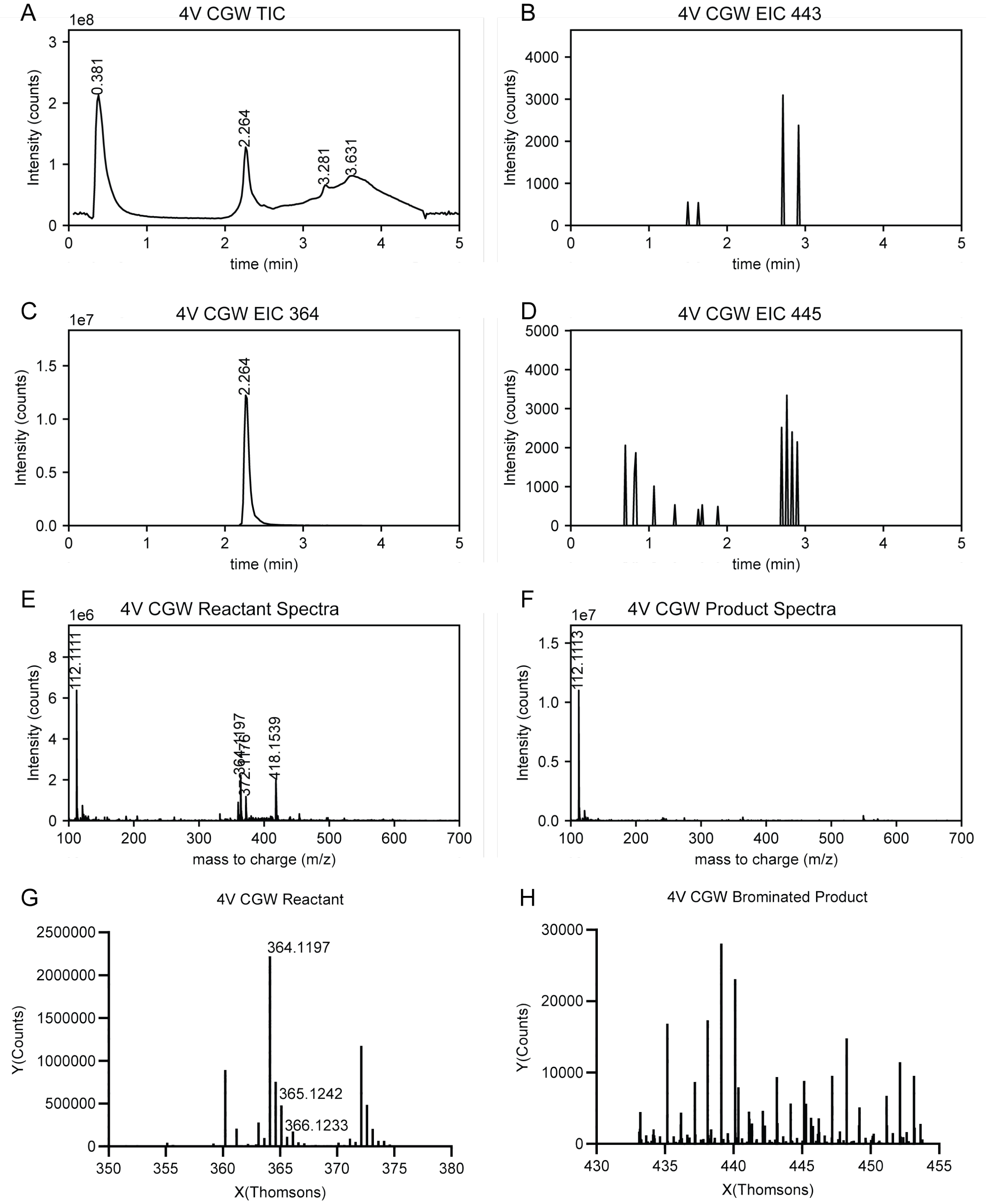

A
4V CGW TIC
1e8
Intensity (counts)
time (min)
0.381
2.264
3.281
3.631
B
4V CGW EIC 443
Intensity (counts)
time (min)
C
4V CGW EIC 364
1e7
Intensity (counts)
time (min)
2.264
D
4V CGW EIC 445
Intensity (counts)
time (min)
E
4V CGW Reactant Spectra
1e6
Intensity (counts)
mass to charge (m/z)
112.1111
364.1197
372.1176
418.1539
F
4V CGW Product Spectra
1e7
Intensity (counts)
mass to charge (m/z)
112.1113
G
4V CGW Reactant
Y(Counts)
X(Thomsons)
364.1197
365.1242
366.1233
H
4V CGW Brominated Product
Y(Counts)
X(Thomsons)

#### 4V DGW

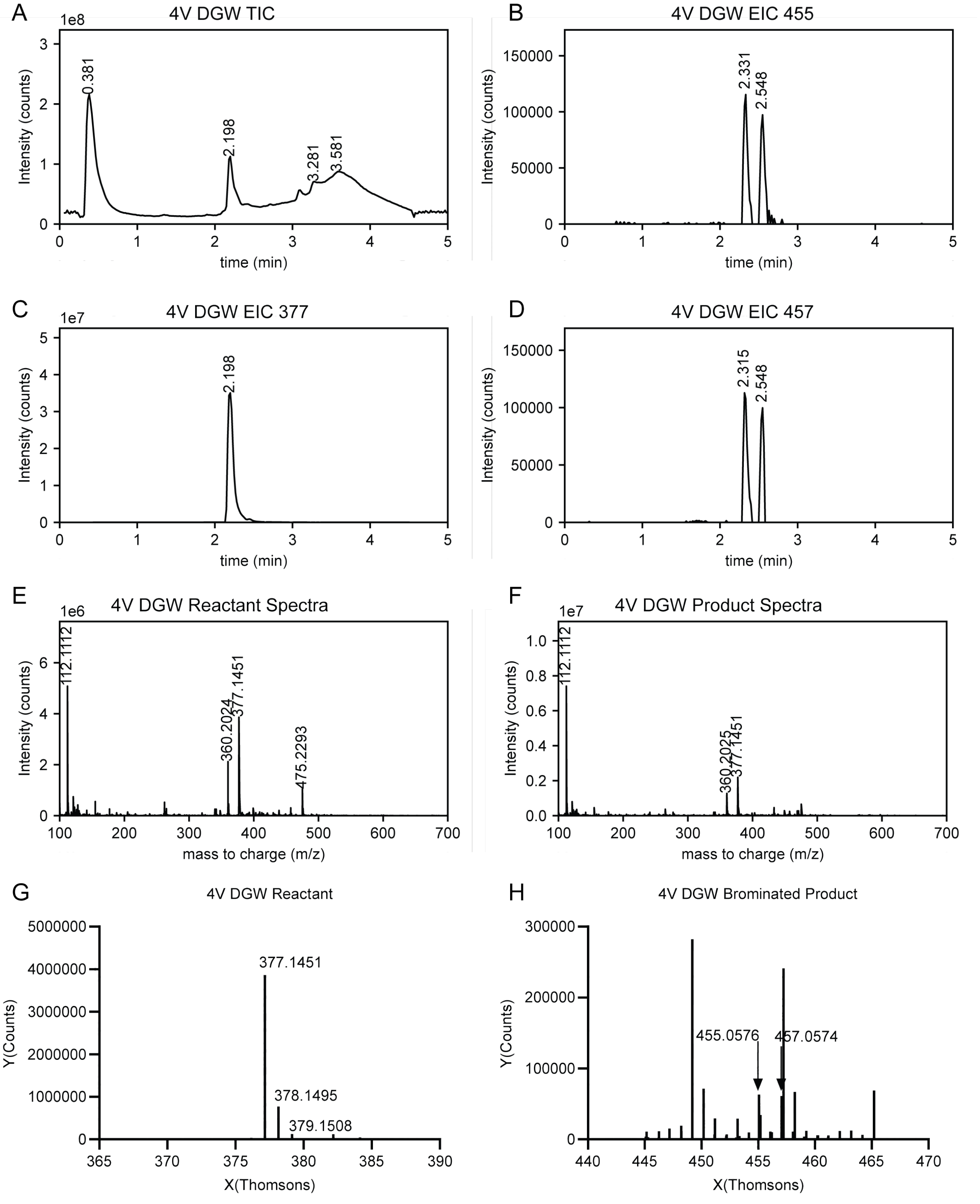
A
4V DGW TIC
1e8
Intensity (counts)
time (min)
0.381
2.198
3.281
3.581
B
4V DGW EIC 455
Intensity (counts)
time (min)
2.331
2.548
C
4V DGW EIC 377
1e7
Intensity (counts)
time (min)
2.198
D
4V DGW EIC 457
Intensity (counts)
time (min)
2.315
2.548
E
4V DGW Reactant Spectra
1e6
Intensity (counts)
mass to charge (m/z)
112.1112
360.2024
377.1451
475.2293
F
4V DGW Product Spectra
1e7
Intensity (counts)
mass to charge (m/z)
112.1112
360.2025
377.1451
G
4V DGW Reactant
Y(Counts)
X(Thomsons)
377.1451
378.1495
379.1508
H
4V DGW Brominated Product
Y(Counts)
X(Thomsons)
455.0576
457.0574

#### 4V EGW

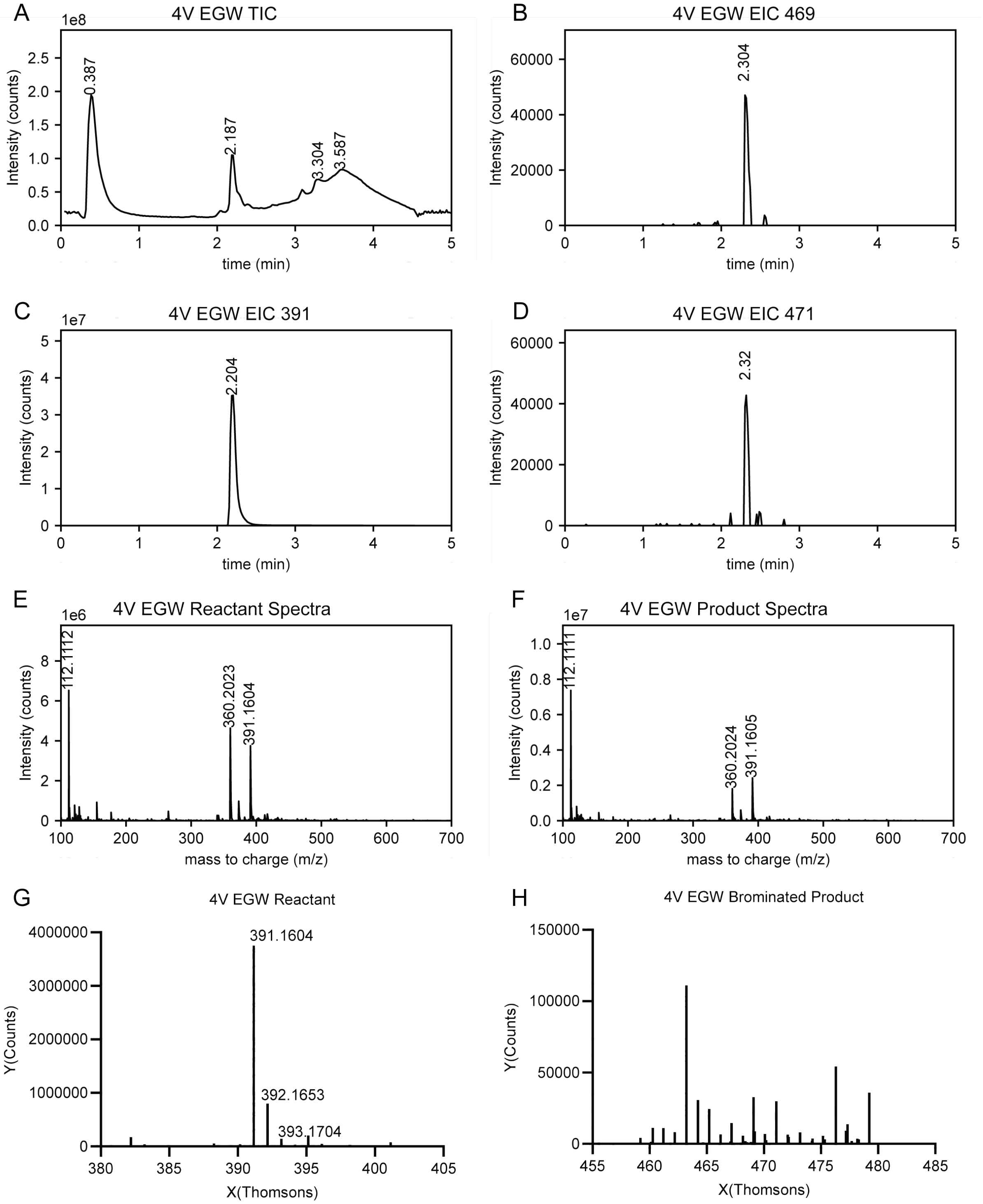

A
4V EGW TIC
B
4V EGW EIC 469
C
4V EGW EIC 391
D
4V EGW EIC 471
E
4V EGW Reactant Spectra
F
4V EGW Product Spectra
G
4V EGW Reactant
H
4V EGW Brominated Product
Intensity (counts)
time (min)
mass to charge (m/z)
Y(Counts)
X(Thomsons)
0.387
2.187
3.304
3.587
2.304
2.204
2.32
112.1112
360.2023
391.1604
112.1111
360.2024
391.1605
392.1653
393.1704

#### 4V FGW

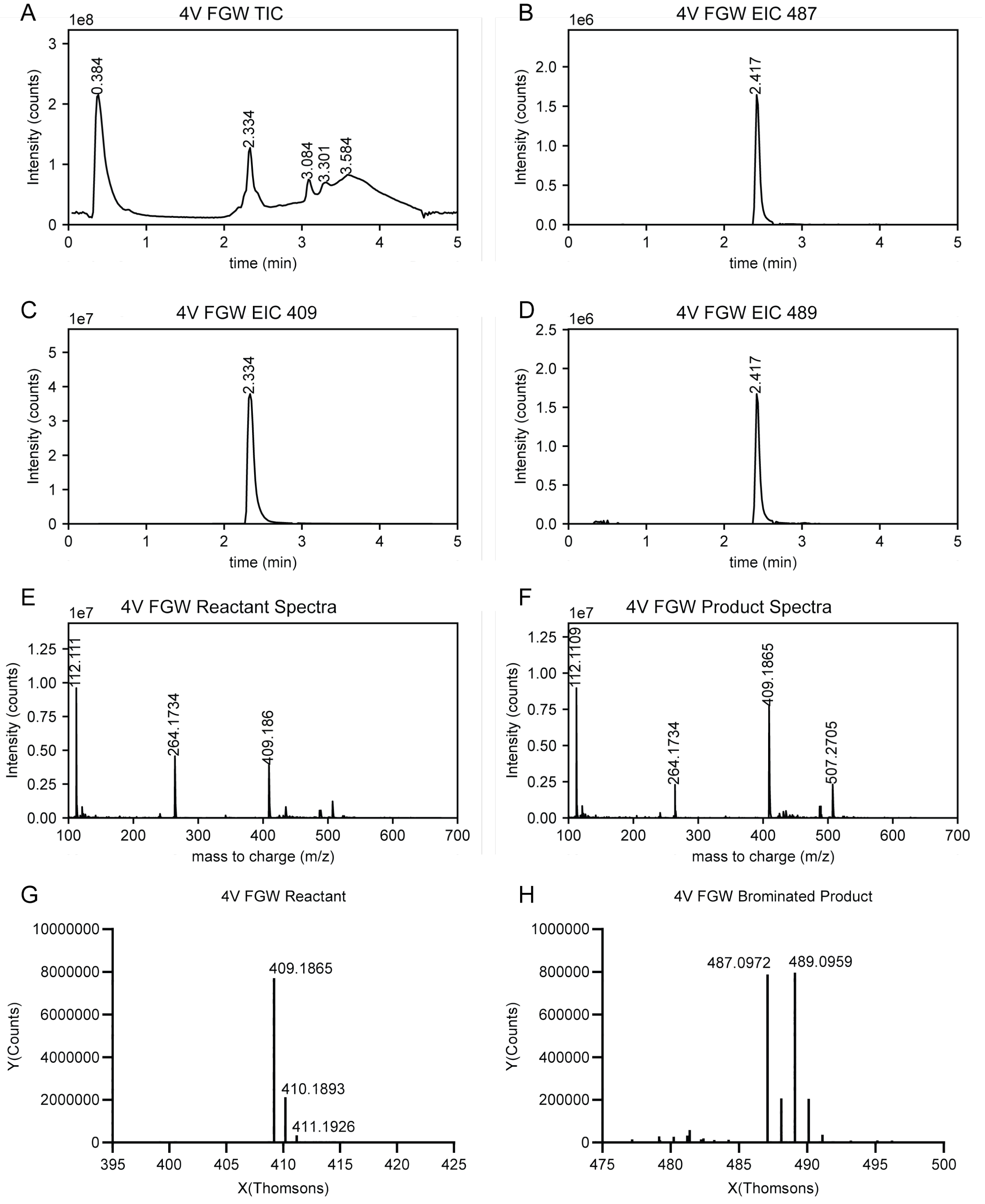
A
4V FGW TIC
1e8
Intensity (counts)
time (min)
0.384
2.334
3.084
3.301
3.584
B
4V FGW EIC 487
1e6
Intensity (counts)
time (min)
2.417
C
4V FGW EIC 409
1e7
Intensity (counts)
time (min)
2.334
D
4V FGW EIC 489
1e6
Intensity (counts)
time (min)
2.417
E
4V FGW Reactant Spectra
1e7
Intensity (counts)
mass to charge (m/z)
112.111
264.1734
409.186
F
4V FGW Product Spectra
1e7
Intensity (counts)
mass to charge (m/z)
112.1109
264.1734
409.1865
507.2705
G
4V FGW Reactant
Y(Counts)
X(Thomsons)
409.1865
410.1893
411.1926
H
4V FGW Brominated Product
Y(Counts)
X(Thomsons)
487.0972
489.0959

#### 4V HGW

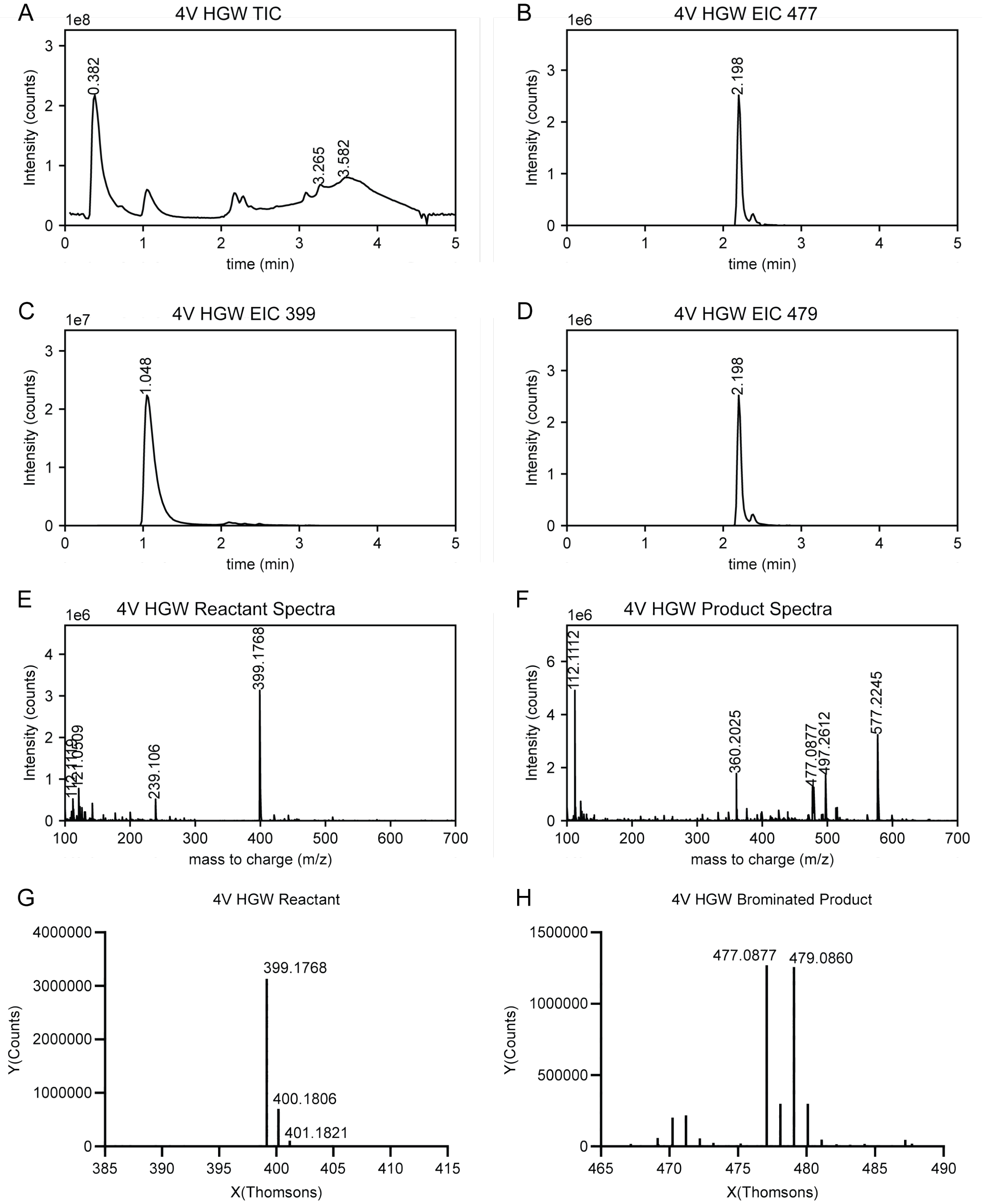

A
4V HGW TIC
1e8
Intensity (counts)
time (min)
0.382
3.265
3.582
B
4V HGW EIC 477
1e6
Intensity (counts)
time (min)
2.198
C
4V HGW EIC 399
1e7
Intensity (counts)
time (min)
1.048
D
4V HGW EIC 479
1e6
Intensity (counts)
time (min)
2.198
E
4V HGW Reactant Spectra
1e6
Intensity (counts)
mass to charge (m/z)
399.1768
239.106
F
4V HGW Product Spectra
1e6
Intensity (counts)
mass to charge (m/z)
112.1112
360.2025
477.0877
497.2612
577.2245
G
4V HGW Reactant
Y(Counts)
X(Thomsons)
399.1768
400.1806
401.1821
H
4V HGW Brominated Product
Y(Counts)
X(Thomsons)
477.0877
479.0860

#### 4V KGW

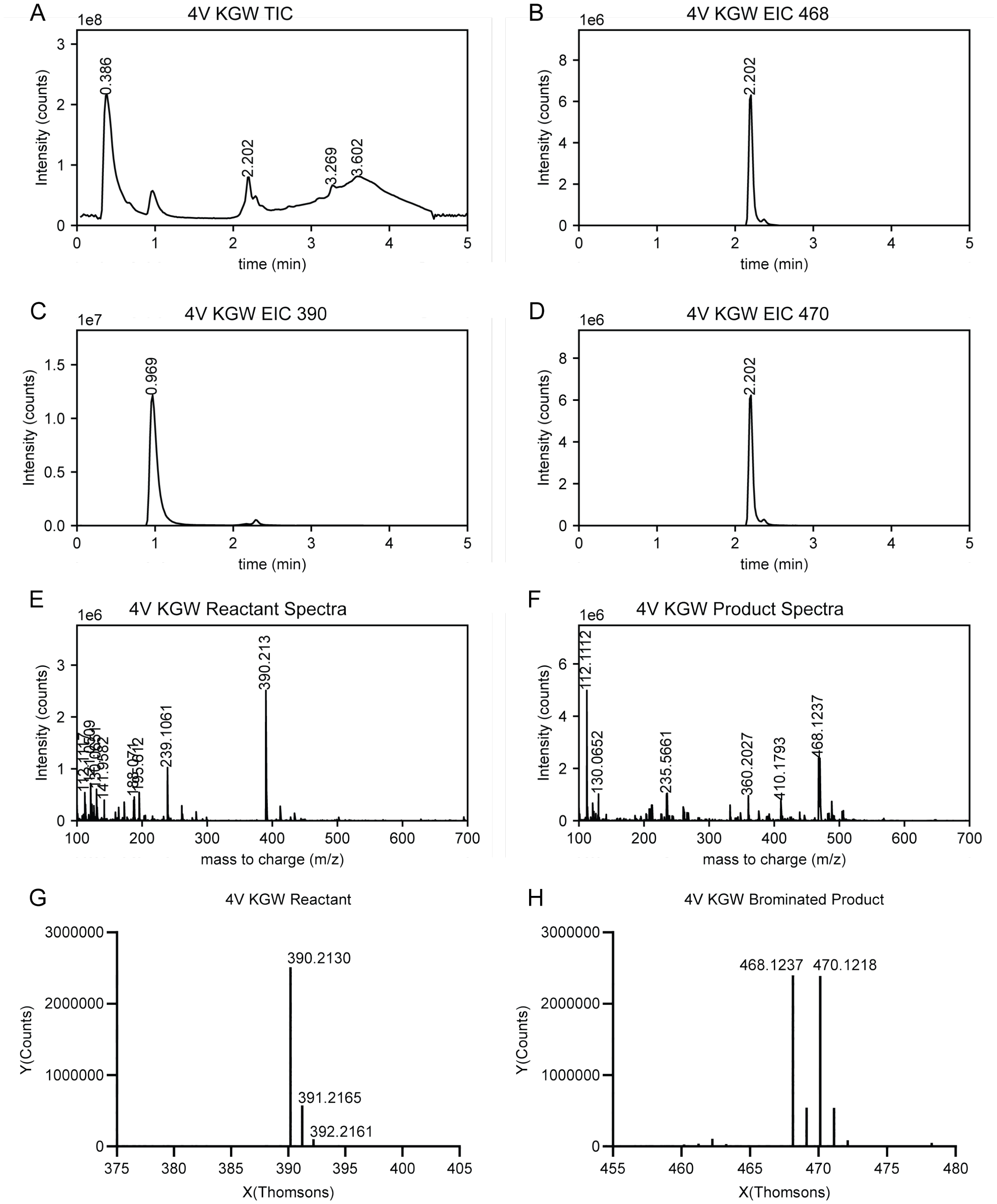

A
4V KGW TIC
1e8
Intensity (counts)
time (min)
0.386
2.202
3.269
3.602
B
4V KGW EIC 468
1e6
Intensity (counts)
time (min)
2.202
C
4V KGW EIC 390
1e7
Intensity (counts)
time (min)
0.969
D
4V KGW EIC 470
1e6
Intensity (counts)
time (min)
2.202
E
4V KGW Reactant Spectra
1e6
Intensity (counts)
mass to charge (m/z)
390.213
239.1061
F
4V KGW Product Spectra
1e6
Intensity (counts)
mass to charge (m/z)
112.1112
130.0652
235.5661
360.2027
410.1793
468.1237
G
4V KGW Reactant
Y(Counts)
X(Thomsons)
390.2130
391.2165
392.2161
H
4V KGW Brominated Product
Y(Counts)
X(Thomsons)
468.1237
470.1218

#### 4V LGW

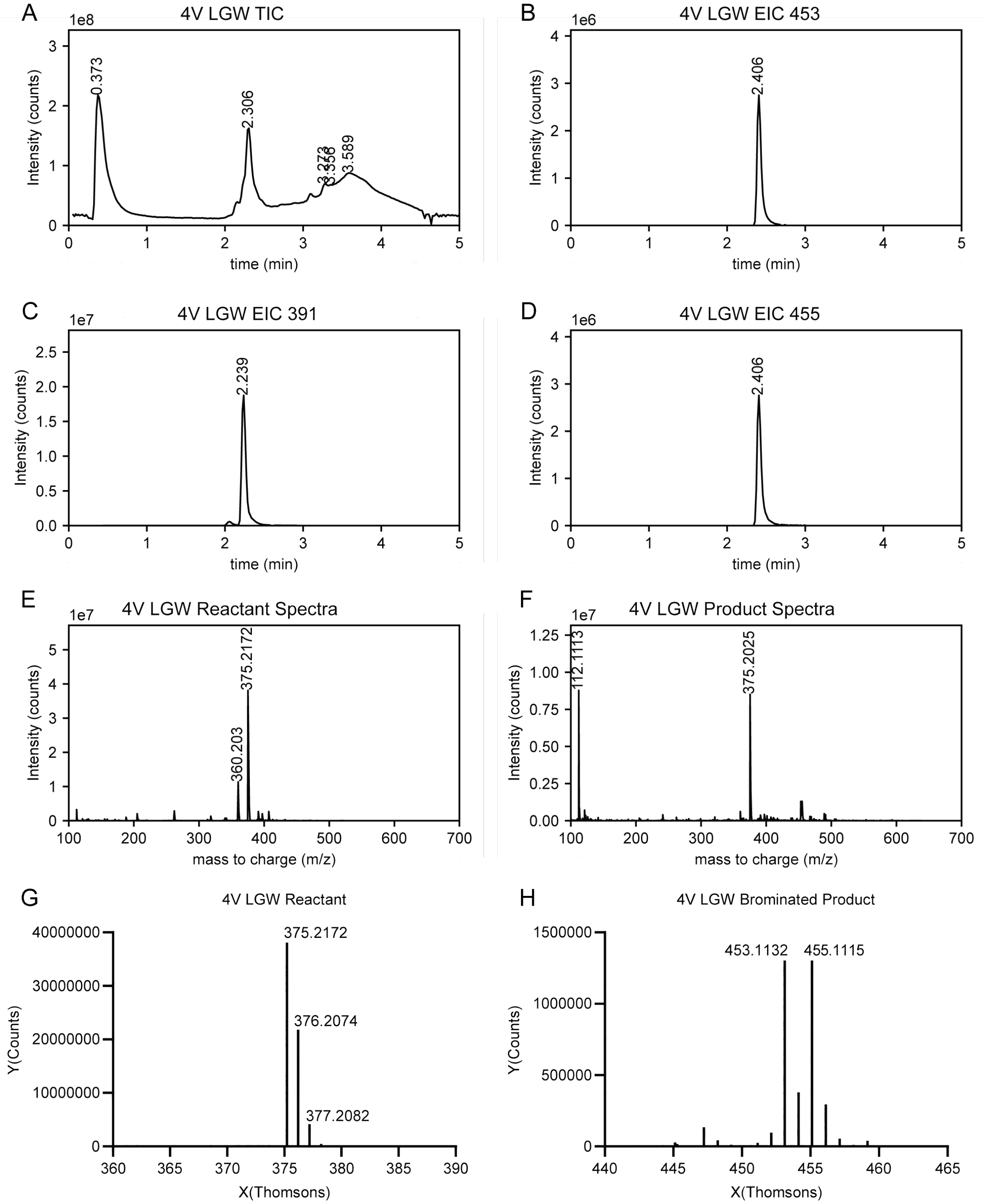
A
4V LGW TIC
1e8
Intensity (counts)
time (min)
0.373
2.306
3.589
B
4V LGW EIC 453
1e6
2.406
C
4V LGW EIC 391
1e7
2.239
D
4V LGW EIC 455
1e6
2.406
E
4V LGW Reactant Spectra
1e7
mass to charge (m/z)
375.2172
360.203
F
4V LGW Product Spectra
1e7
112.1113
375.2025
G
4V LGW Reactant
Y(Counts)
X(Thomsons)
375.2172
376.2074
377.2082
H
4V LGW Brominated Product
453.1132
455.1115

#### 4V MGW

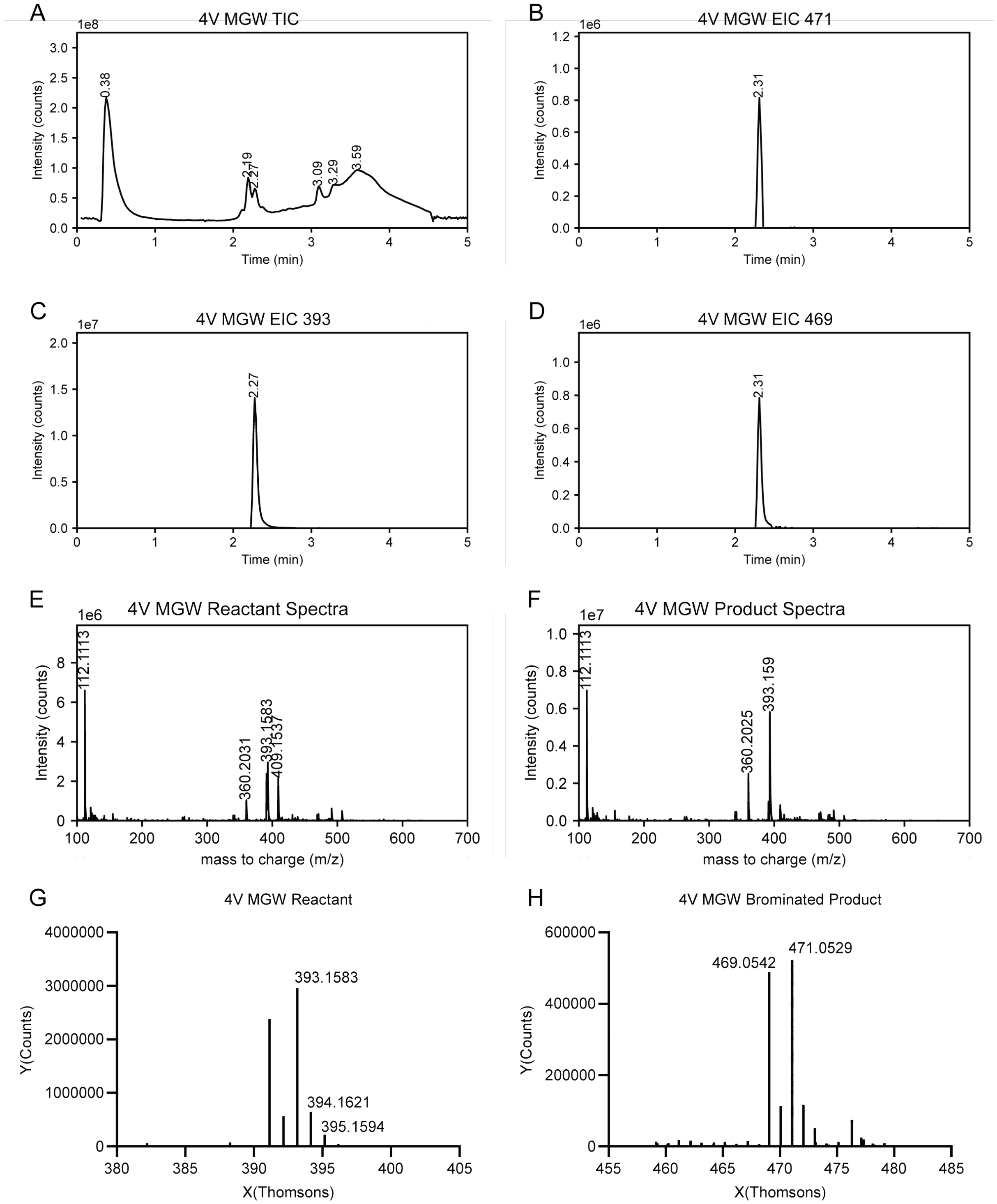
A
4V MGW TIC
1e8
Intensity (counts)
Time (min)
0.38
2.19
2.27
3.09
3.29
3.59
B
4V MGW EIC 471
1e6
2.31
C
4V MGW EIC 393
1e7
2.27
D
4V MGW EIC 469
1e6
2.31
E
4V MGW Reactant Spectra
1e6
112.1113
360.2031
393.1583
409.1537
mass to charge (m/z)
F
4V MGW Product Spectra
1e7
112.1113
360.2025
393.159
G
4V MGW Reactant
Y(Counts)
393.1583
394.1621
395.1594
X(Thomsons)
H
4V MGW Brominated Product
469.0542
471.0529

#### 4V NGW

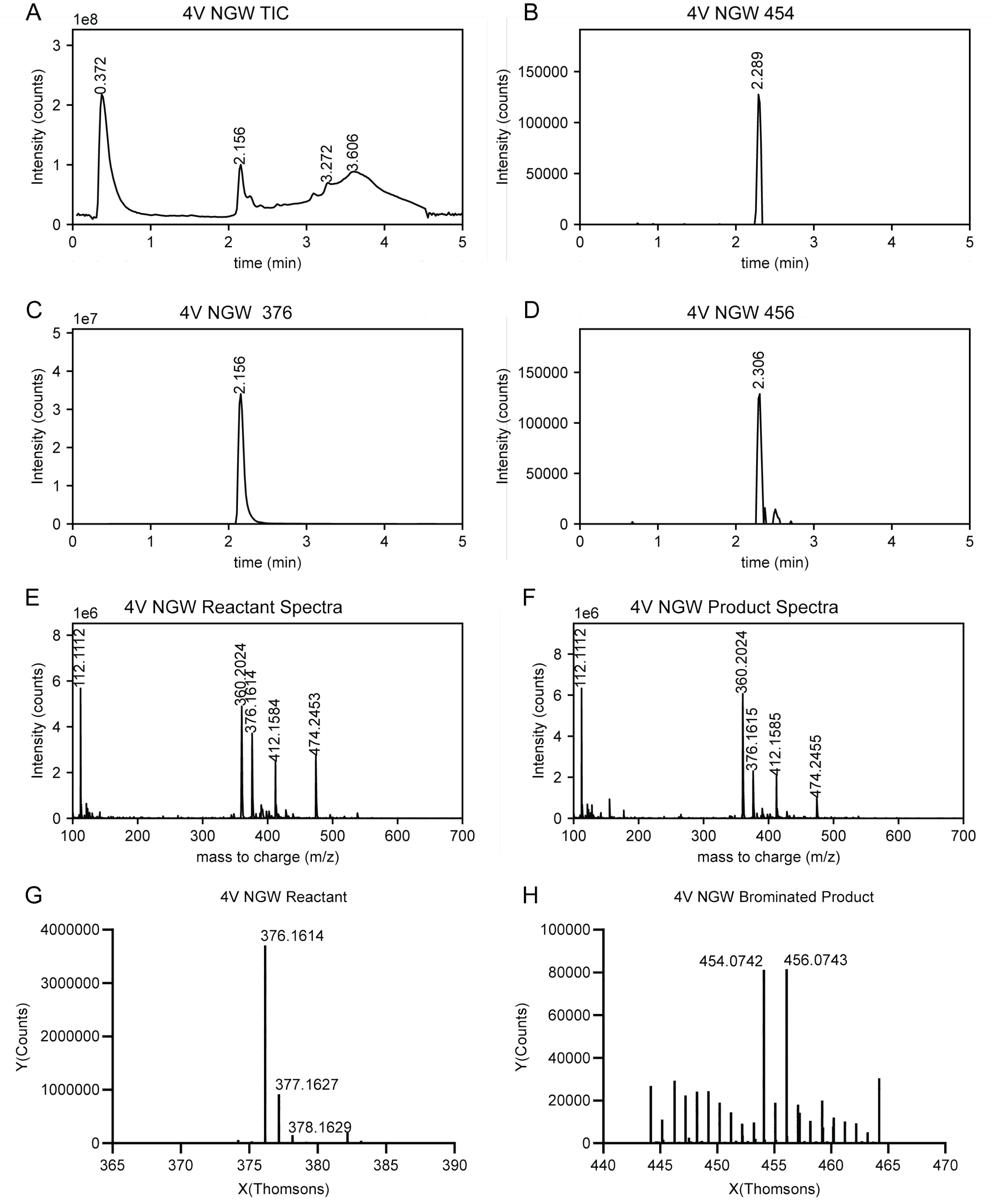
A
4V NGW TIC
1e8
Intensity (counts)
time (min)
0.372
2.156
3.272
3.606
B
4V NGW 454
Intensity (counts)
time (min)
2.289
C
4V NGW 376
1e7
Intensity (counts)
time (min)
2.156
D
4V NGW 456
Intensity (counts)
time (min)
2.306
E
4V NGW Reactant Spectra
1e6
Intensity (counts)
mass to charge (m/z)
112.1112
360.2024
376.1614
412.1584
474.2453
F
4V NGW Product Spectra
1e6
Intensity (counts)
mass to charge (m/z)
112.1112
360.2024
376.1615
412.1585
474.2455
G
4V NGW Reactant
Y(Counts)
X(Thomsons)
376.1614
377.1627
378.1629
H
4V NGW Brominated Product
Y(Counts)
X(Thomsons)
454.0742
456.0743

#### 4V PGW

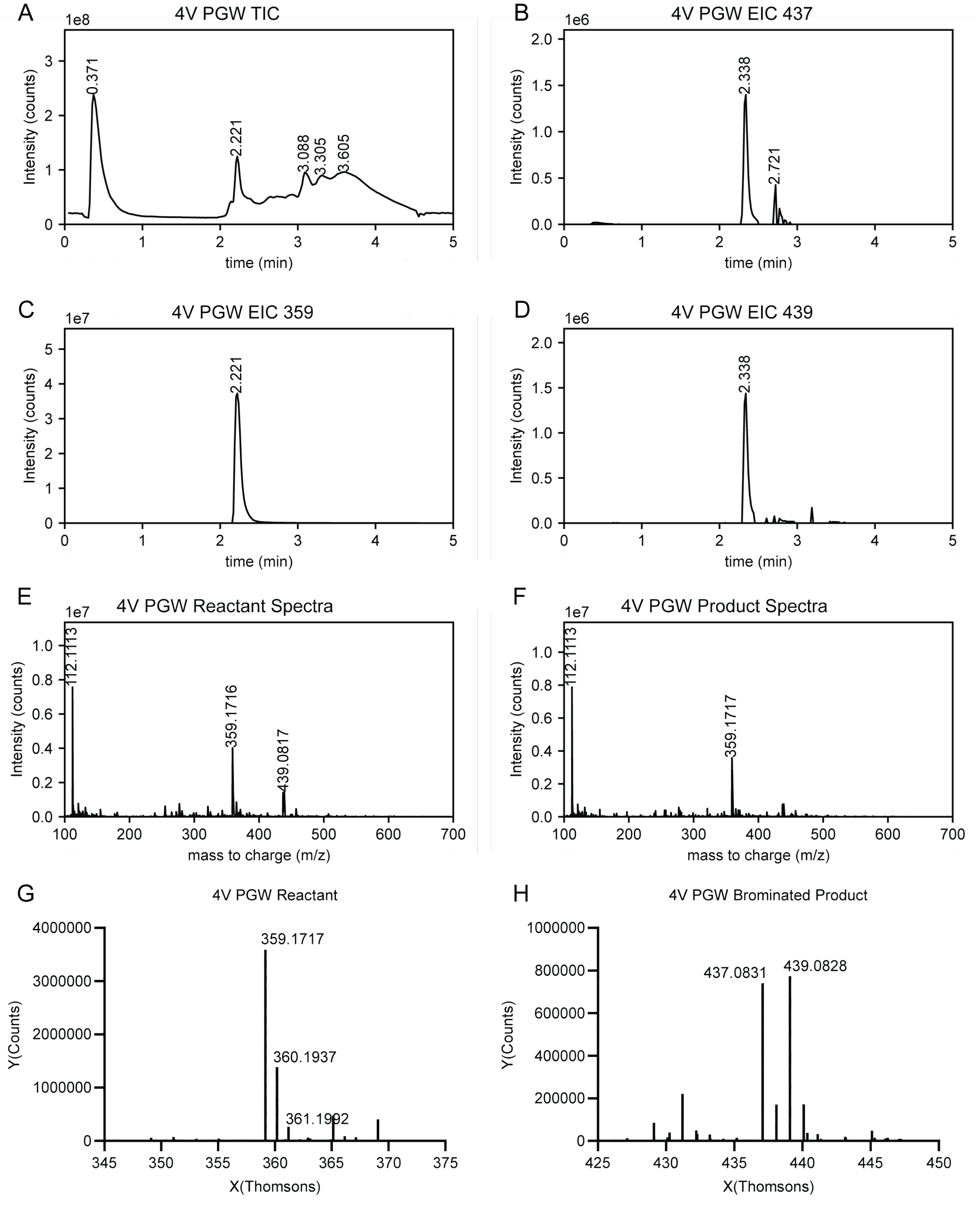

A
4V PGW TIC
1e8
Intensity (counts)
time (min)
0.371
2.221
3.088
3.305
3.605
B
4V PGW EIC 437
1e6
2.338
2.721
C
4V PGW EIC 359
1e7
2.221
D
4V PGW EIC 439
1e6
2.338
E
4V PGW Reactant Spectra
1e7
112.1113
359.1716
439.0817
mass to charge (m/z)
F
4V PGW Product Spectra
1e7
112.1113
359.1717
G
4V PGW Reactant
359.1717
360.1937
361.1992
Y(Counts)
X(Thomsons)
H
4V PGW Brominated Product
437.0831
439.0828

#### 4V QGW

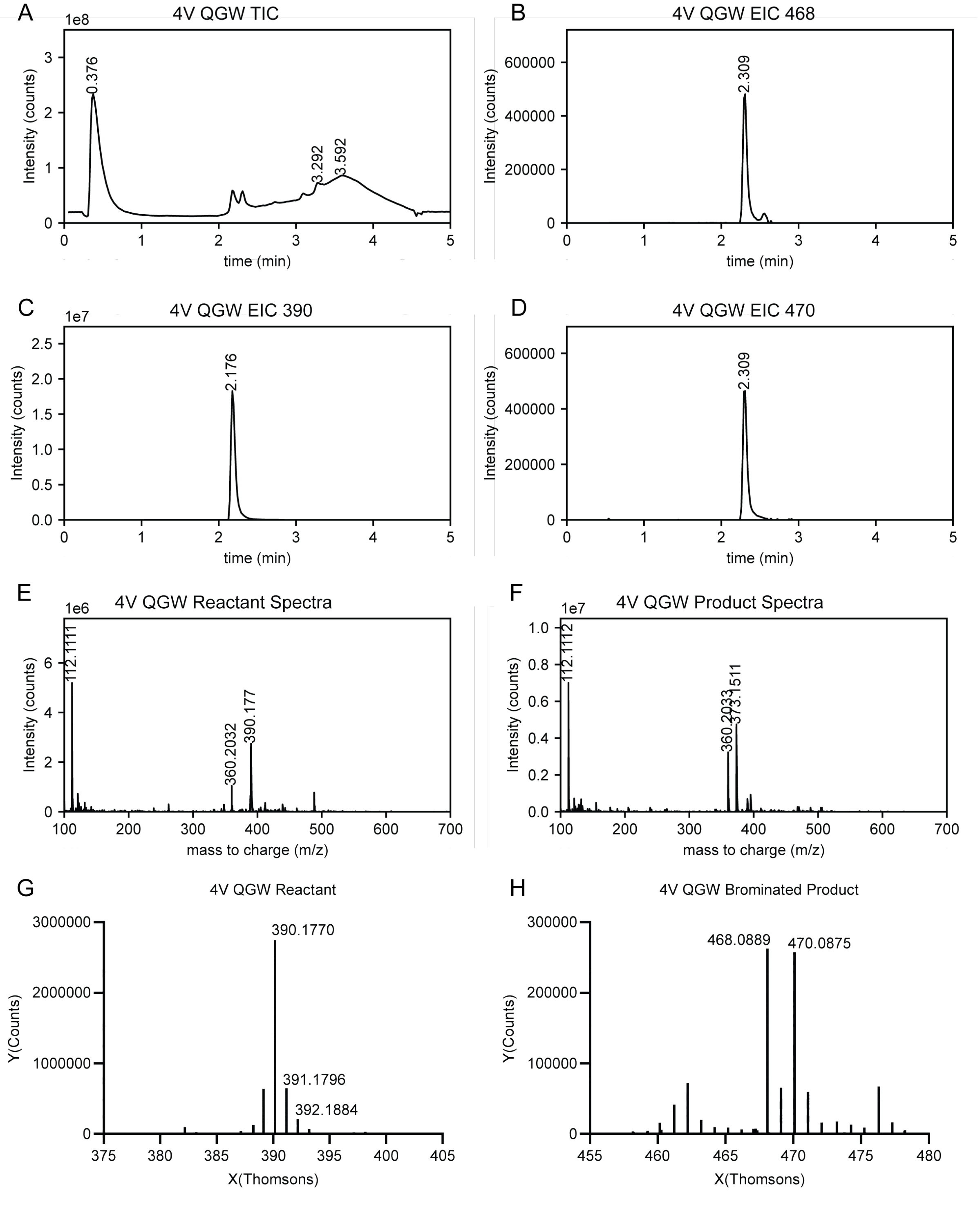
A
4V QGW TIC
1e8
Intensity (counts)
time (min)
0.376
3.292
3.592
B
4V QGW EIC 468
Intensity (counts)
time (min)
2.309
C
4V QGW EIC 390
1e7
Intensity (counts)
time (min)
2.176
D
4V QGW EIC 470
Intensity (counts)
time (min)
2.309
E
4V QGW Reactant Spectra
1e6
Intensity (counts)
mass to charge (m/z)
112.1111
360.2032
390.177
F
4V QGW Product Spectra
1e7
Intensity (counts)
mass to charge (m/z)
112.1112
360.2033
373.1511
G
4V QGW Reactant
Y(Counts)
X(Thomsons)
390.1770
391.1796
392.1884
H
4V QGW Brominated Product
Y(Counts)
X(Thomsons)
468.0889
470.0875

#### 4V RGW

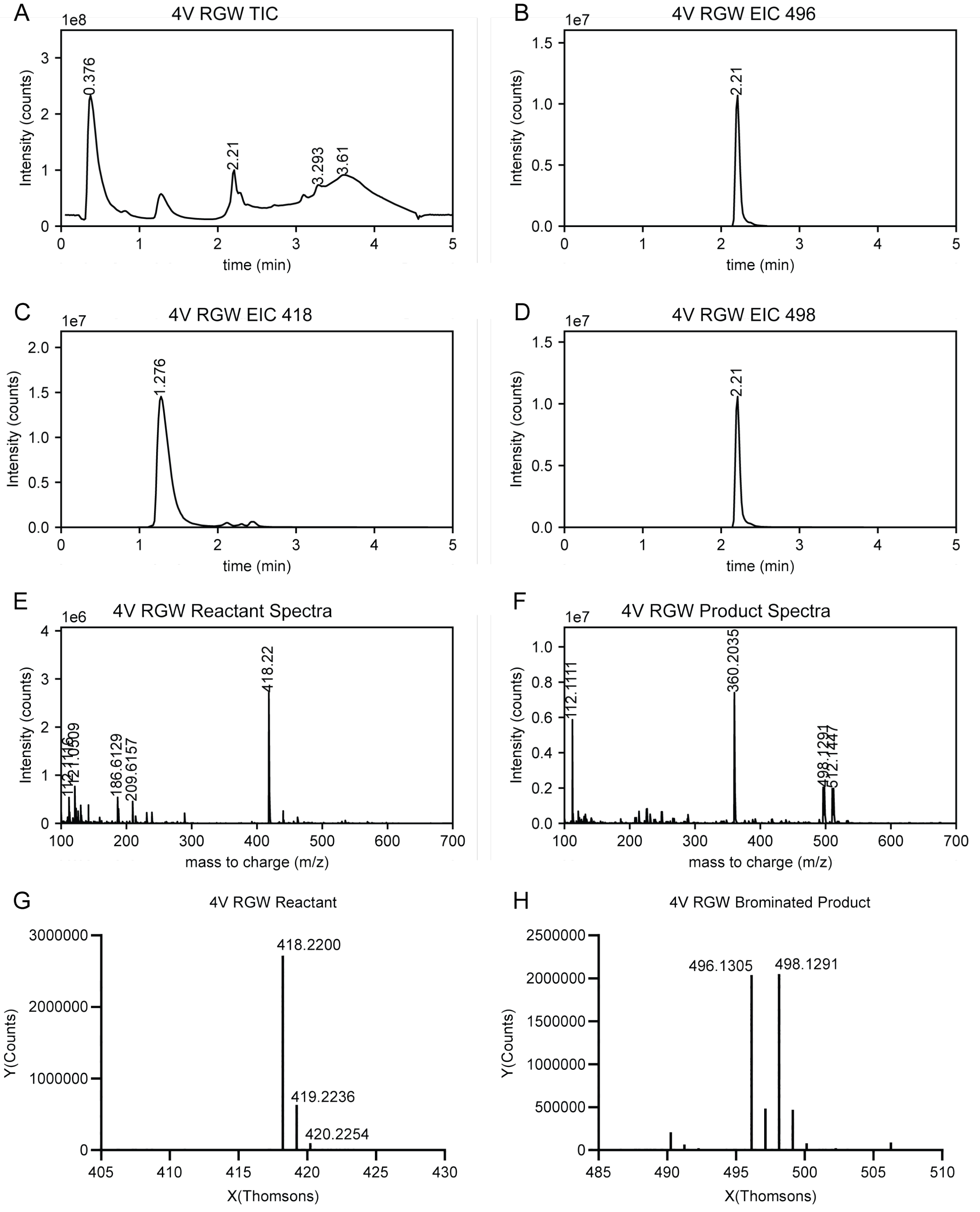
A
4V RGW TIC
1e8
Intensity (counts)
time (min)
0.376
2.21
3.293
3.61
B
4V RGW EIC 496
1e7
Intensity (counts)
time (min)
2.21
C
4V RGW EIC 418
1e7
Intensity (counts)
time (min)
1.276
D
4V RGW EIC 498
1e7
Intensity (counts)
time (min)
2.21
E
4V RGW Reactant Spectra
1e6
Intensity (counts)
mass to charge (m/z)
121.0509
186.6129
209.6157
418.22
F
4V RGW Product Spectra
1e7
Intensity (counts)
mass to charge (m/z)
112.1111
360.2035
498.1291
512.1447
G
4V RGW Reactant
Y(Counts)
X(Thomsons)
418.2200
419.2236
420.2254
H
4V RGW Brominated Product
Y(Counts)
X(Thomsons)
496.1305
498.1291

#### 4V TGW

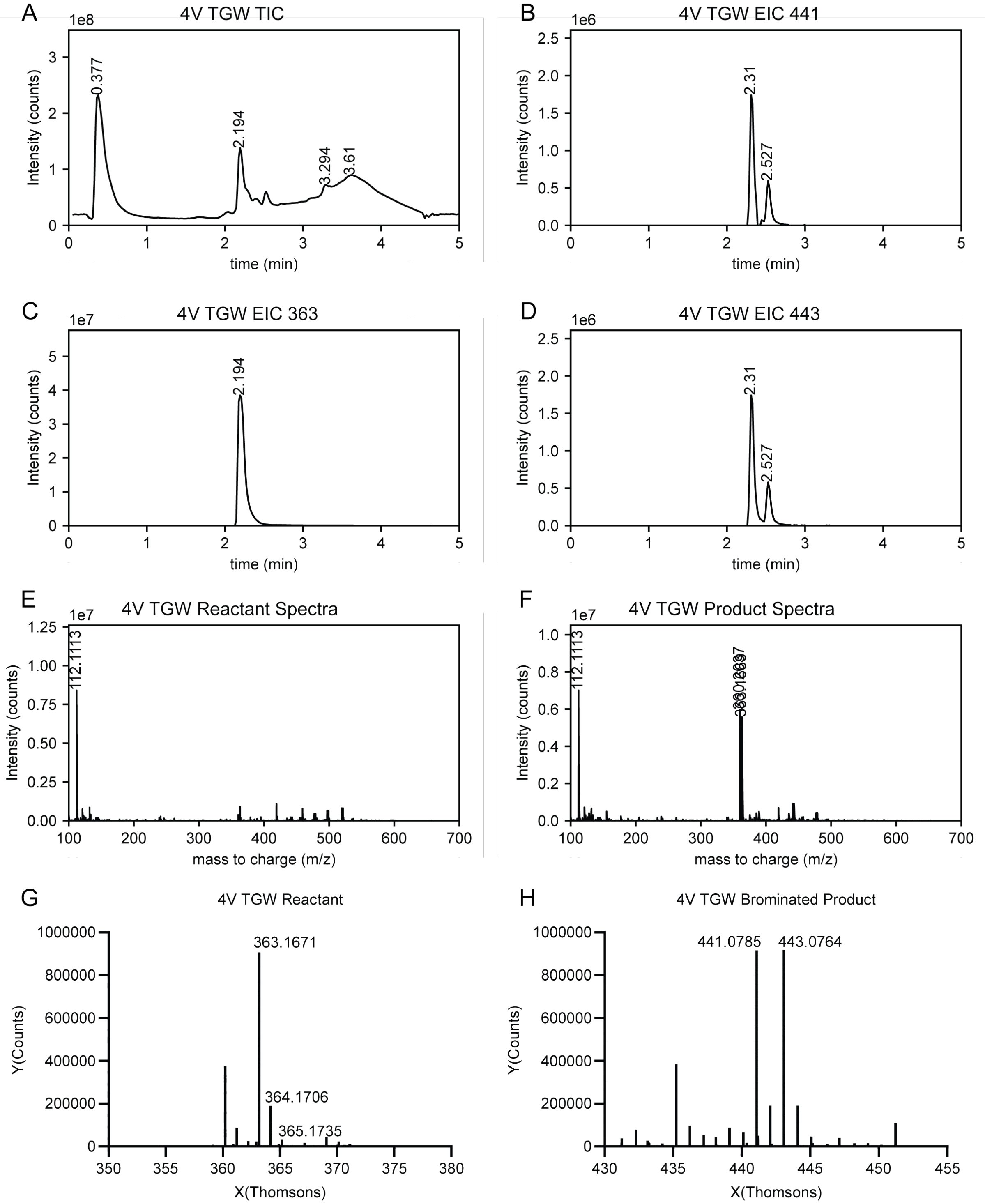
A
4V TGW TIC
1e8
Intensity (counts)
time (min)
0.377
2.194
3.294
3.61
B
4V TGW EIC 441
1e6
Intensity (counts)
time (min)
2.31
2.527
C
4V TGW EIC 363
1e7
Intensity (counts)
time (min)
2.194
D
4V TGW EIC 443
1e6
Intensity (counts)
time (min)
2.31
2.527
E
4V TGW Reactant Spectra
1e7
Intensity (counts)
mass to charge (m/z)
112.1113
F
4V TGW Product Spectra
1e7
Intensity (counts)
mass to charge (m/z)
112.1113
G
4V TGW Reactant
Y(Counts)
X(Thomsons)
363.1671
364.1706
365.1735
H
4V TGW Brominated Product
Y(Counts)
X(Thomsons)
441.0785
443.0764

#### 4V VGW

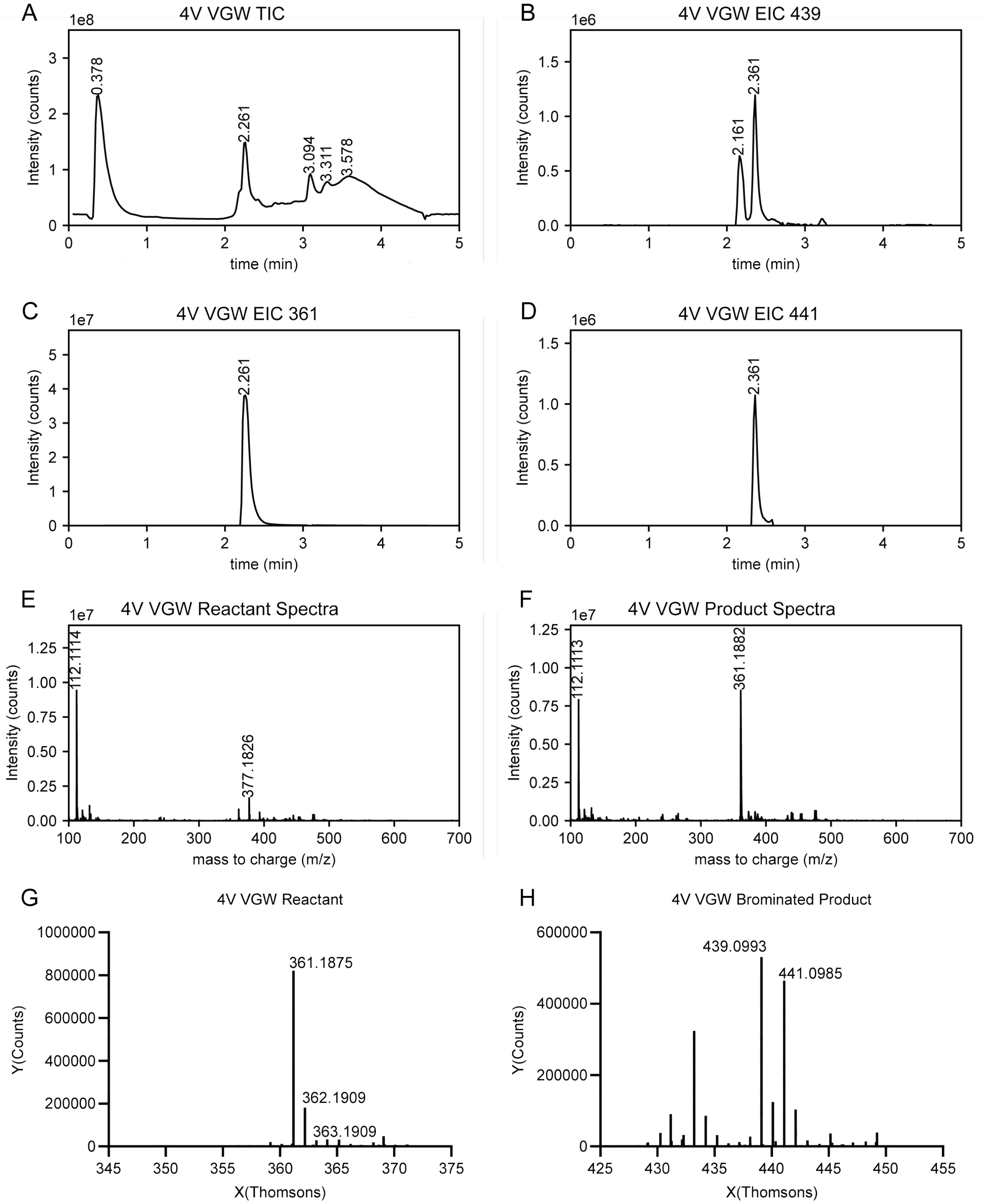
A
4V VGW TIC
1e8
Intensity (counts)
time (min)
0.378
2.261
3.094
3.311
3.578
B
4V VGW EIC 439
1e6
2.161
2.361
C
4V VGW EIC 361
1e7
2.261
D
4V VGW EIC 441
1e6
2.361
E
4V VGW Reactant Spectra
1e7
112.1114
377.1826
mass to charge (m/z)
F
4V VGW Product Spectra
1e7
112.1113
361.1882
G
4V VGW Reactant
361.1875
362.1909
363.1909
Y(Counts)
X(Thomsons)
H
4V VGW Brominated Product
439.0993
441.0985

#### 4V YGW

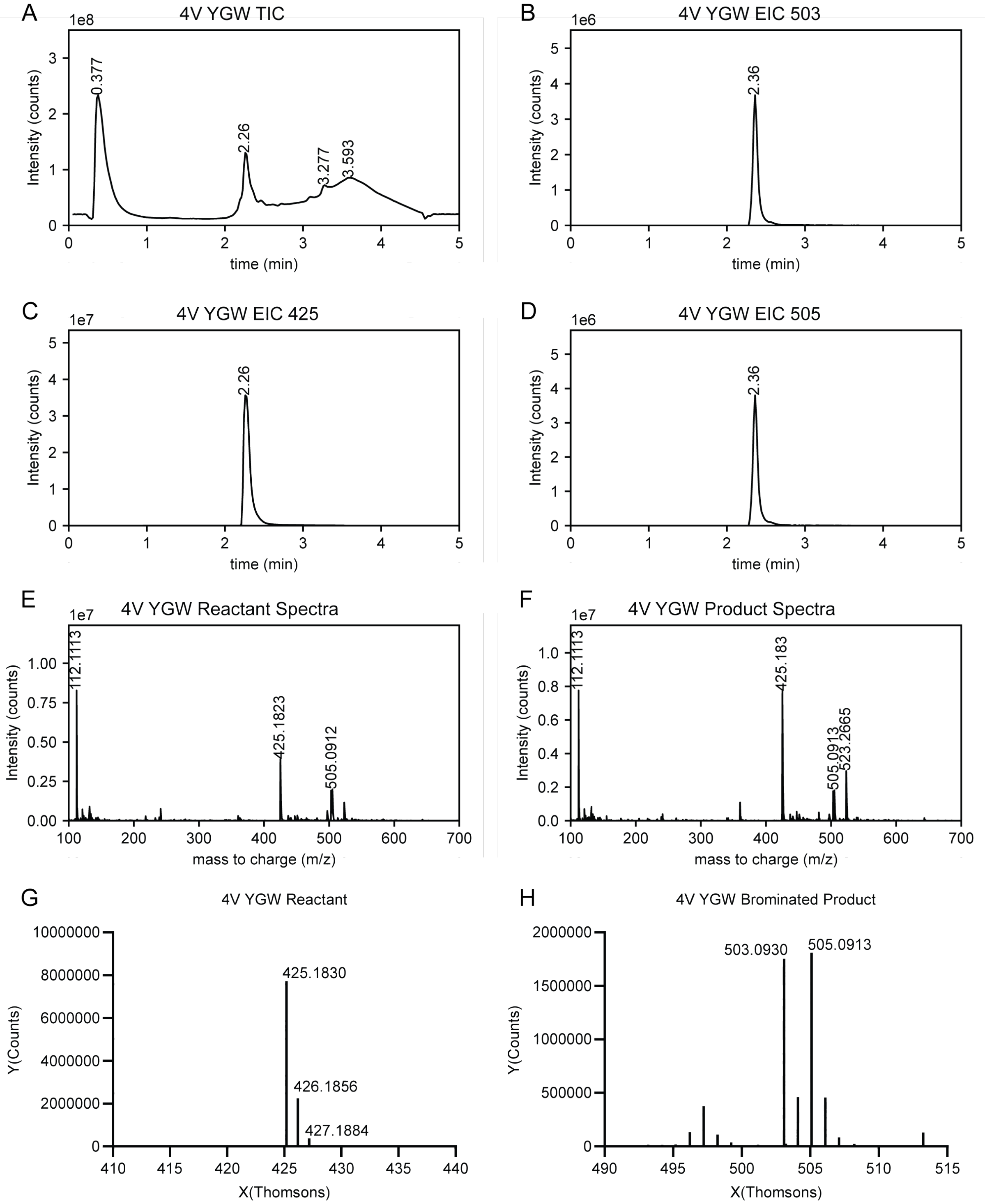

A
4V YGW TIC
1e8
Intensity (counts)
time (min)
0.377
2.26
3.277
3.593
B
4V YGW EIC 503
1e6
Intensity (counts)
time (min)
2.36
C
4V YGW EIC 425
1e7
Intensity (counts)
time (min)
2.26
D
4V YGW EIC 505
1e6
Intensity (counts)
time (min)
2.36
E
4V YGW Reactant Spectra
1e7
Intensity (counts)
mass to charge (m/z)
112.1113
425.1823
505.0912
F
4V YGW Product Spectra
1e7
Intensity (counts)
mass to charge (m/z)
112.1113
425.183
505.0913
523.2665
G
4V YGW Reactant
Y(Counts)
X(Thomsons)
425.1830
426.1856
427.1884
H
4V YGW Brominated Product
Y(Counts)
X(Thomsons)
503.0930
505.0913

#### 4V GAW

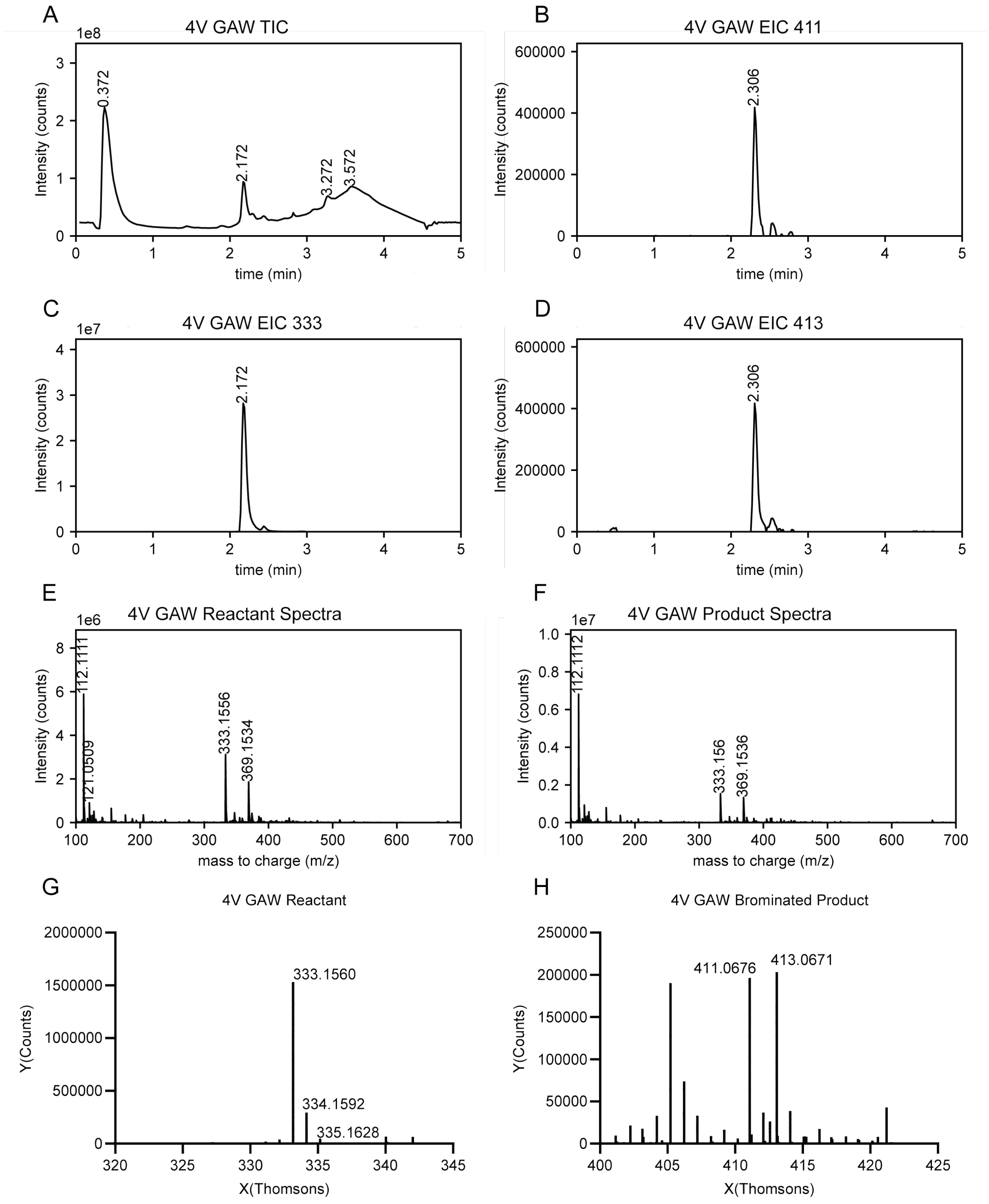

A
4V GAW TIC
1e8
Intensity (counts)
time (min)
0.372
2.172
3.272
3.572
B
4V GAW EIC 411
Intensity (counts)
time (min)
2.306
C
4V GAW EIC 333
1e7
Intensity (counts)
time (min)
2.172
D
4V GAW EIC 413
Intensity (counts)
time (min)
2.306
E
4V GAW Reactant Spectra
1e6
Intensity (counts)
mass to charge (m/z)
112.1111
121.0509
333.1556
369.1534
F
4V GAW Product Spectra
1e7
Intensity (counts)
mass to charge (m/z)
112.1112
333.156
369.1536
G
4V GAW Reactant
Y(Counts)
X(Thomsons)
333.1560
334.1592
335.1628
H
4V GAW Brominated Product
Y(Counts)
X(Thomsons)
411.0676
413.0671

#### 4V GCW

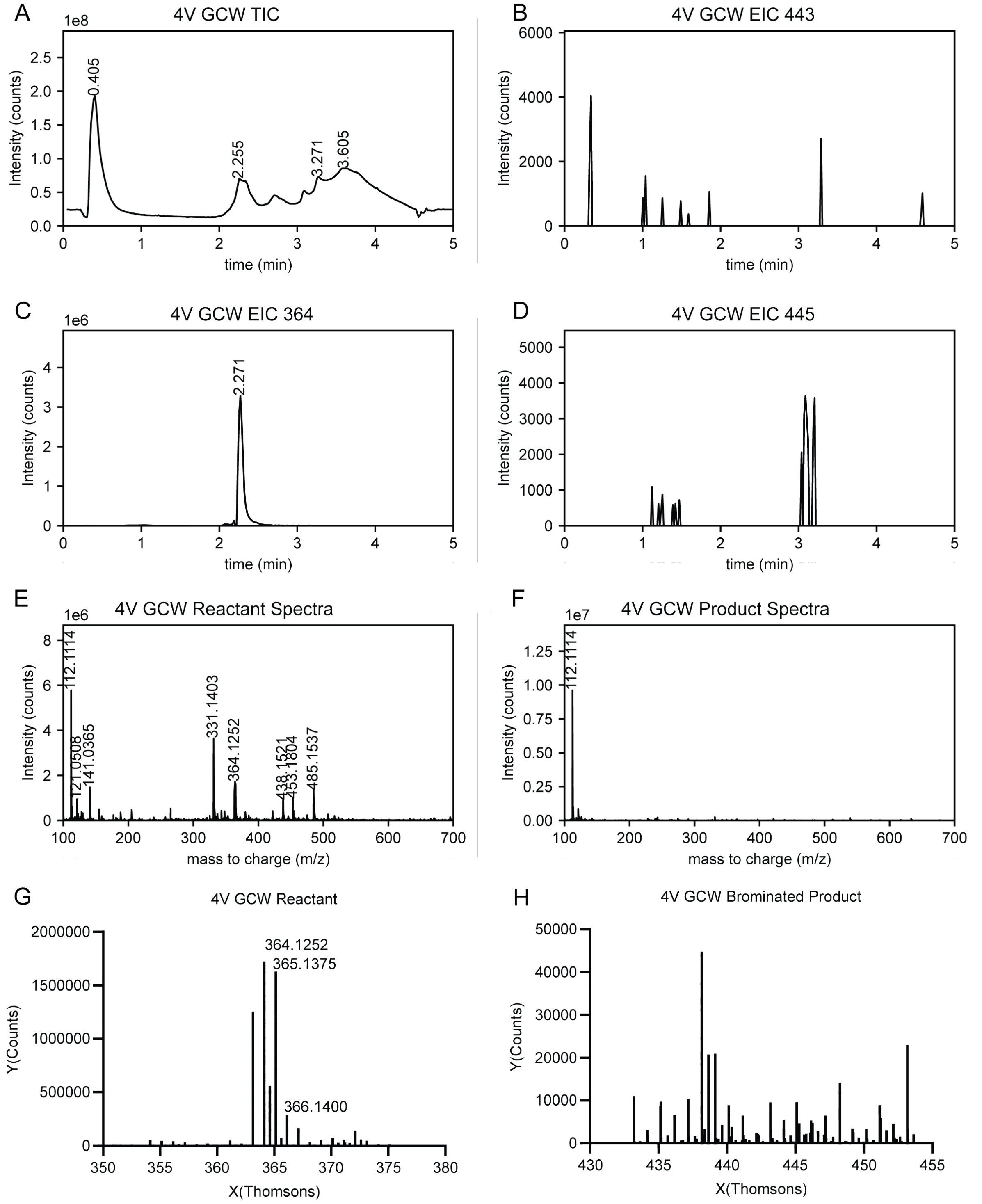
A
4V GCW TIC
1e8
Intensity (counts)
time (min)
0.405
2.255
3.271
3.605
B
4V GCW EIC 443
Intensity (counts)
time (min)
C
4V GCW EIC 364
1e6
Intensity (counts)
time (min)
2.271
D
4V GCW EIC 445
Intensity (counts)
time (min)
E
4V GCW Reactant Spectra
1e6
Intensity (counts)
mass to charge (m/z)
112.1114
121.0508
141.0365
331.1403
364.1252
438.1521
453.1804
485.1537
F
4V GCW Product Spectra
1e7
Intensity (counts)
mass to charge (m/z)
112.1114
G
4V GCW Reactant
Y(Counts)
X(Thomsons)
364.1252
365.1375
366.1400
H
4V GCW Brominated Product
Y(Counts)
X(Thomsons)

### 4V GDW

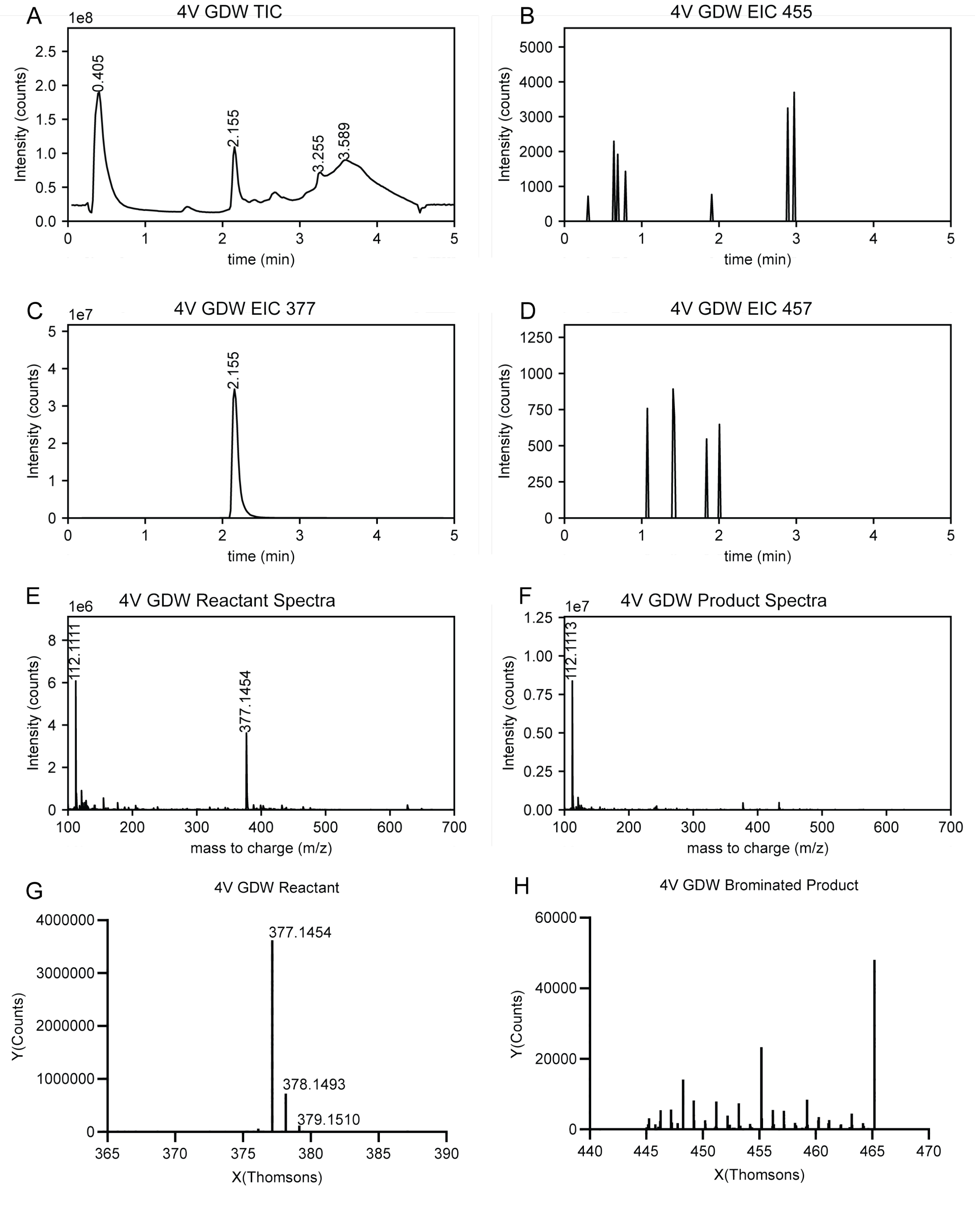

A
4V GDW TIC
1e8
Intensity (counts)
time (min)
0.405
2.155
3.255
3.589
B
4V GDW EIC 455
Intensity (counts)
time (min)
C
4V GDW EIC 377
1e7
Intensity (counts)
time (min)
2.155
D
4V GDW EIC 457
Intensity (counts)
time (min)
E
4V GDW Reactant Spectra
1e6
Intensity (counts)
mass to charge (m/z)
112.1111
377.1454
F
4V GDW Product Spectra
1e7
Intensity (counts)
mass to charge (m/z)
112.1113
G
4V GDW Reactant
Y(Counts)
X(Thomsons)
377.1454
378.1493
379.1510
H
4V GDW Brominated Product
Y(Counts)
X(Thomsons)

#### 4V GEW

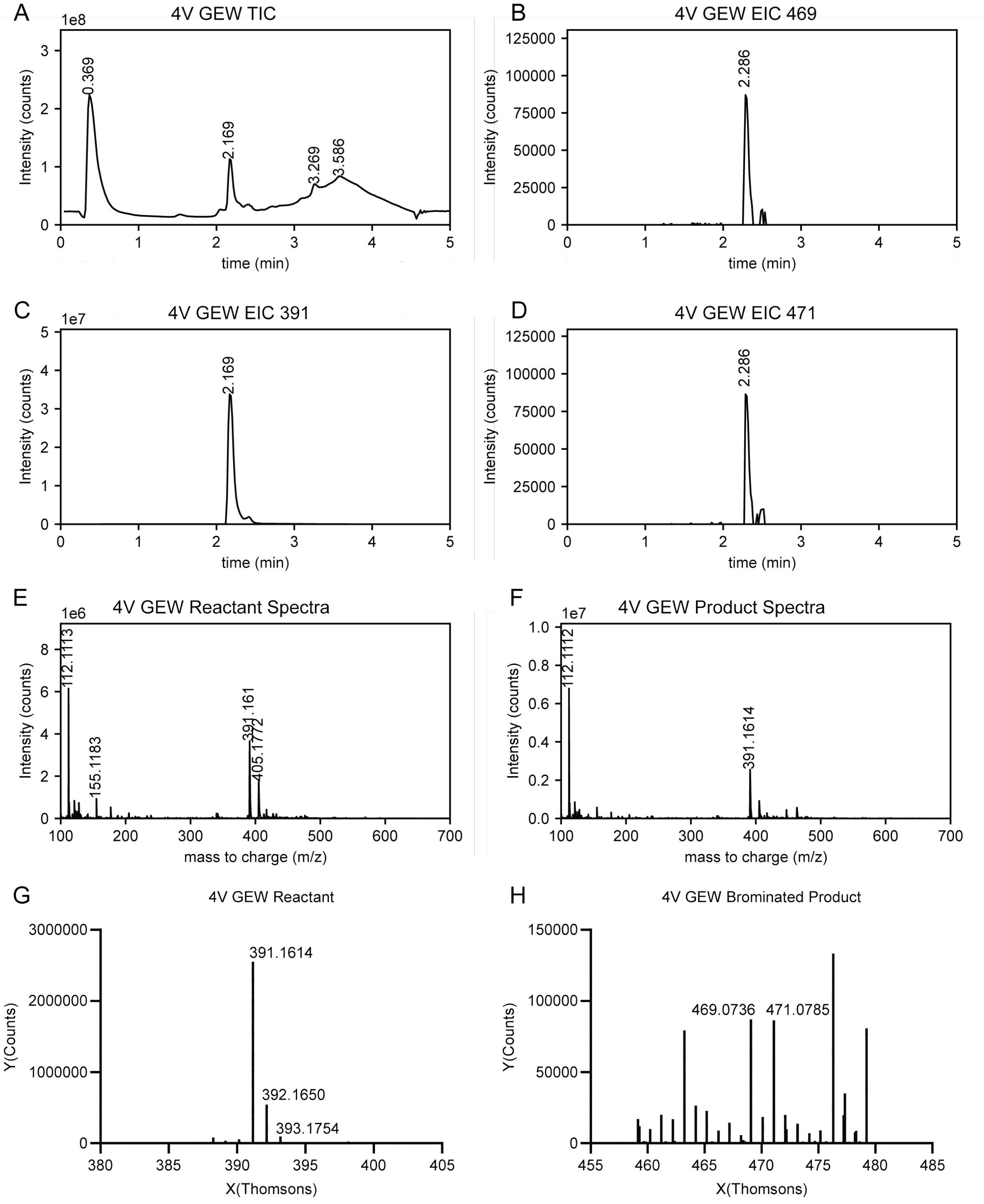

A
4V GEW TIC
1e8
Intensity (counts)
time (min)
0.369
2.169
3.269
3.586
B
4V GEW EIC 469
Intensity (counts)
time (min)
2.286
C
4V GEW EIC 391
1e7
Intensity (counts)
time (min)
2.169
D
4V GEW EIC 471
Intensity (counts)
time (min)
2.286
E
4V GEW Reactant Spectra
1e6
Intensity (counts)
mass to charge (m/z)
112.1113
155.1183
391.161
405.1772
F
4V GEW Product Spectra
1e7
Intensity (counts)
mass to charge (m/z)
112.1112
391.1614
G
4V GEW Reactant
Y(Counts)
X(Thomsons)
391.1614
392.1650
393.1754
H
4V GEW Brominated Product
Y(Counts)
X(Thomsons)
469.0736
471.0785

### 4V GFW

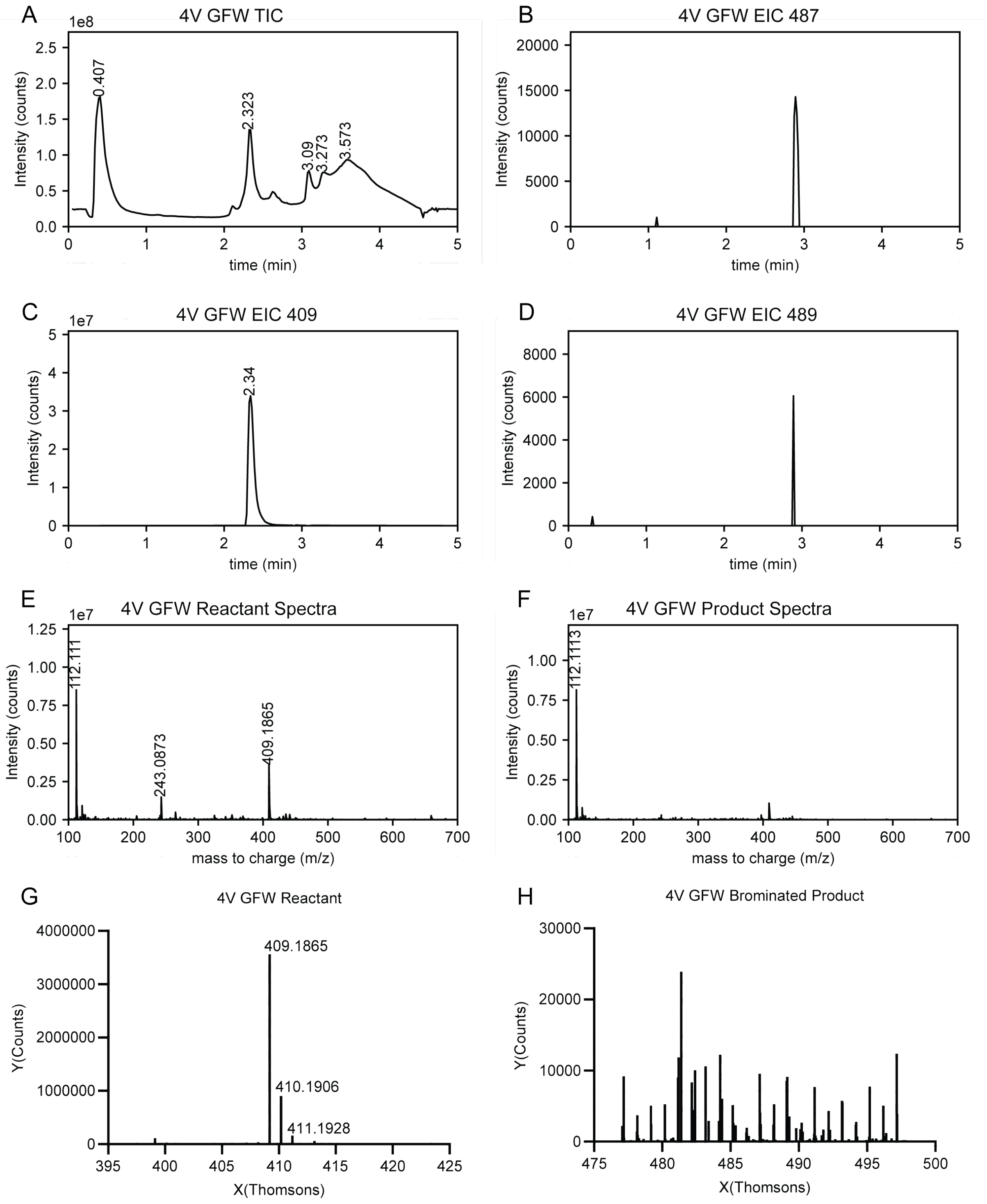
A
4V GFW TIC
1e8
Intensity (counts)
time (min)
0.407
2.323
3.09
3.273
3.573
B
4V GFW EIC 487
Intensity (counts)
time (min)
C
4V GFW EIC 409
1e7
Intensity (counts)
time (min)
2.34
D
4V GFW EIC 489
Intensity (counts)
time (min)
E
4V GFW Reactant Spectra
1e7
Intensity (counts)
mass to charge (m/z)
112.111
243.0873
409.1865
F
4V GFW Product Spectra
1e7
Intensity (counts)
mass to charge (m/z)
112.1113
G
4V GFW Reactant
Y(Counts)
X(Thomsons)
409.1865
410.1906
411.1928
H
4V GFW Brominated Product
Y(Counts)
X(Thomsons)

#### 4V GHW

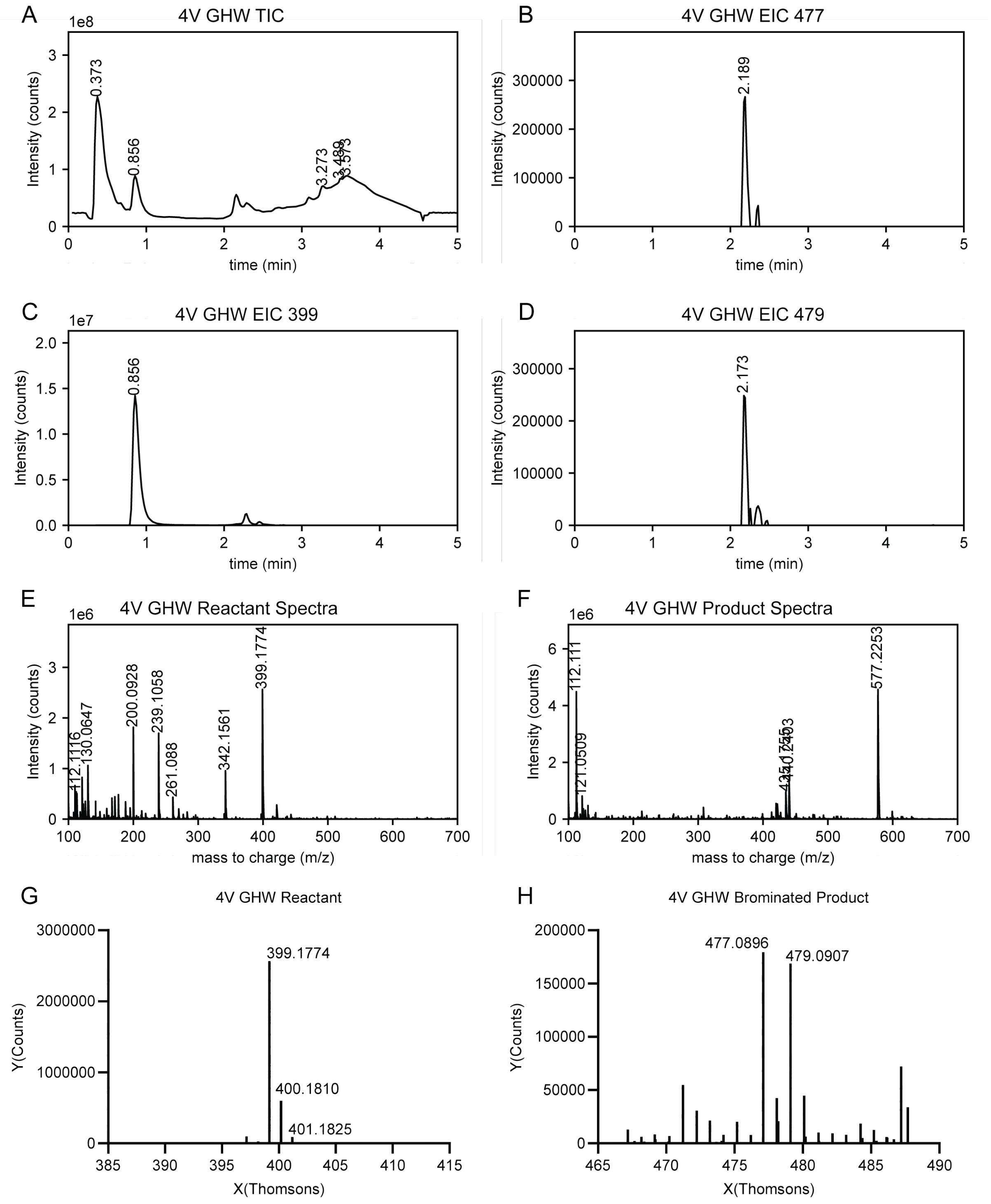
A
4V GHW TIC
1e8
Intensity (counts)
time (min)
0.373
0.856
3.273
B
4V GHW EIC 477
Intensity (counts)
time (min)
2.189
C
4V GHW EIC 399
1e7
Intensity (counts)
time (min)
0.856
D
4V GHW EIC 479
Intensity (counts)
time (min)
2.173
E
4V GHW Reactant Spectra
1e6
Intensity (counts)
mass to charge (m/z)
112.1116
130.0647
200.0928
239.1058
261.088
342.1561
399.1774
F
4V GHW Product Spectra
1e6
Intensity (counts)
mass to charge (m/z)
112.111
121.0509
577.2253
G
4V GHW Reactant
Y(Counts)
X(Thomsons)
399.1774
400.1810
401.1825
H
4V GHW Brominated Product
Y(Counts)
X(Thomsons)
477.0896
479.0907

**4V GKW**

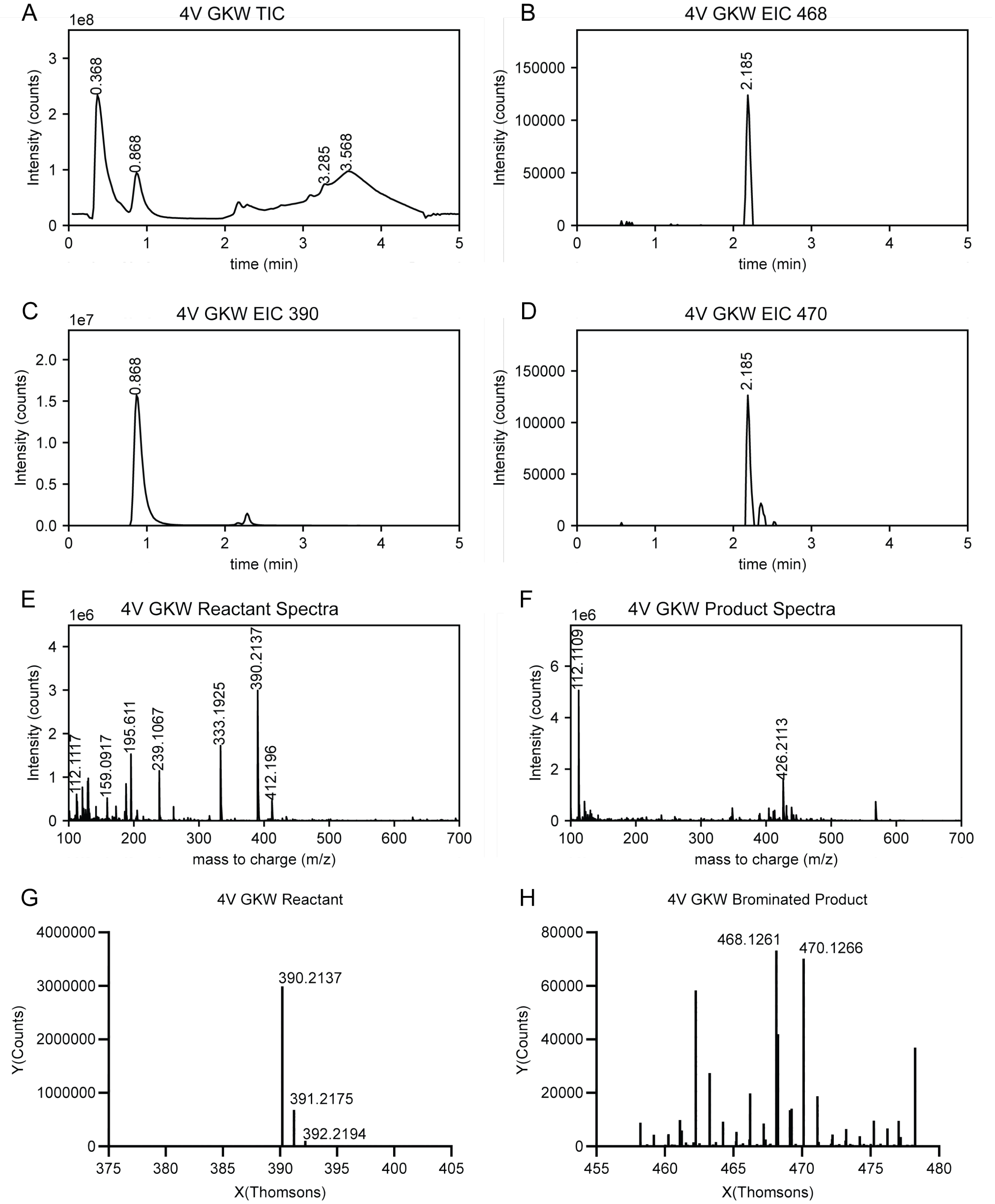

A
4V GKW TIC
1e8
Intensity (counts)
time (min)
0.368
0.868
3.285
3.568
B
4V GKW EIC 468
Intensity (counts)
time (min)
2.185
C
4V GKW EIC 390
1e7
Intensity (counts)
time (min)
0.868
D
4V GKW EIC 470
Intensity (counts)
time (min)
2.185
E
4V GKW Reactant Spectra
1e6
Intensity (counts)
mass to charge (m/z)
112.1117
159.0917
195.611
239.1067
333.1925
390.2137
412.196
F
4V GKW Product Spectra
1e6
Intensity (counts)
mass to charge (m/z)
112.1109
426.2113
G
4V GKW Reactant
Y(Counts)
X(Thomsons)
390.2137
391.2175
392.2194
H
4V GKW Brominated Product
Y(Counts)
X(Thomsons)
468.1261
470.1266

#### 4V GLW

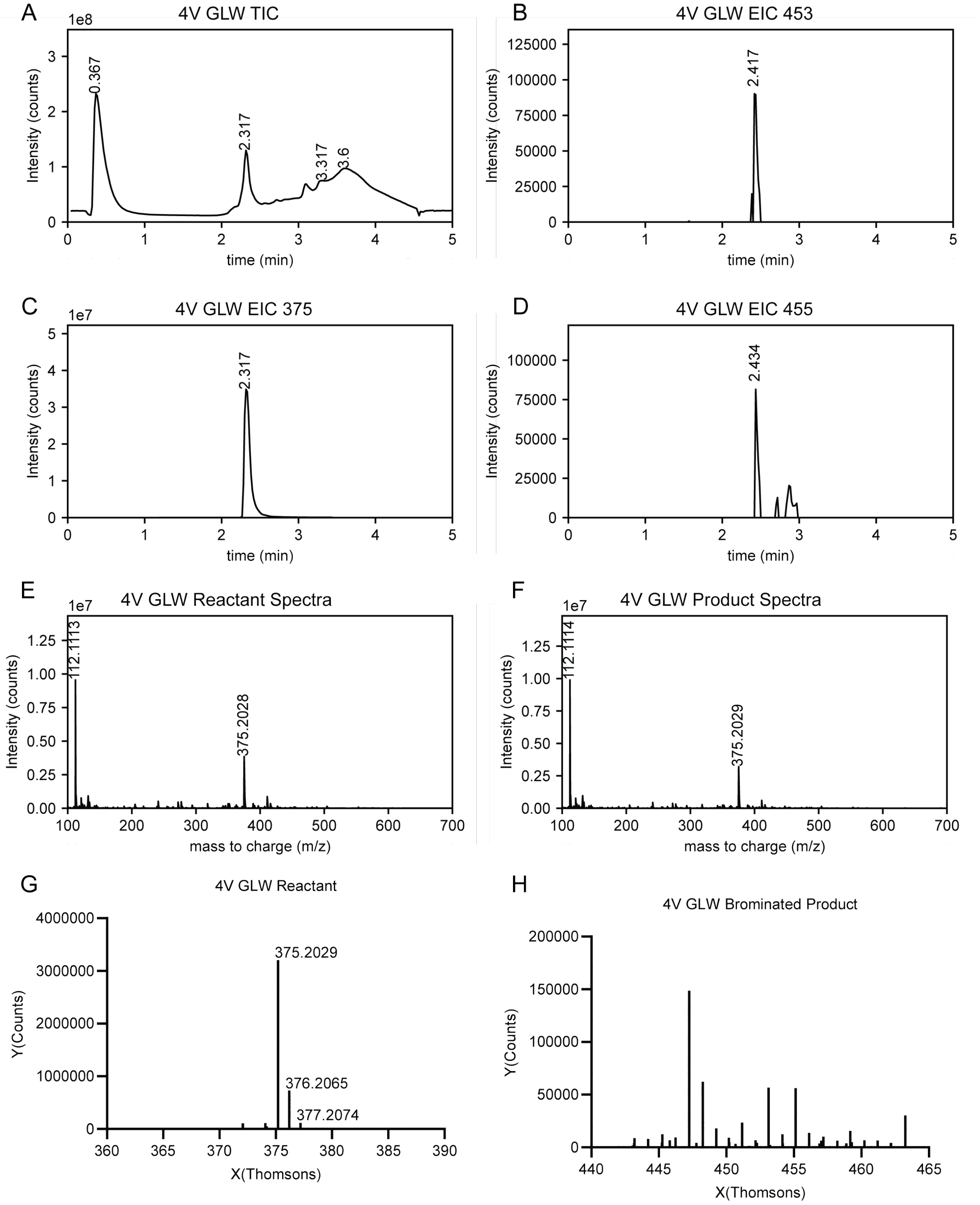

A
4V GLW TIC
1e8
Intensity (counts)
time (min)
0.367
2.317
3.317
3.6
B
4V GLW EIC 453
Intensity (counts)
time (min)
2.417
C
4V GLW EIC 375
1e7
Intensity (counts)
time (min)
2.317
D
4V GLW EIC 455
Intensity (counts)
time (min)
2.434
E
4V GLW Reactant Spectra
1e7
Intensity (counts)
mass to charge (m/z)
112.1113
375.2028
F
4V GLW Product Spectra
1e7
Intensity (counts)
mass to charge (m/z)
112.1114
375.2029
G
4V GLW Reactant
Y(Counts)
X(Thomsons)
375.2029
376.2065
377.2074
H
4V GLW Brominated Product
Y(Counts)
X(Thomsons)

#### 4V GMW

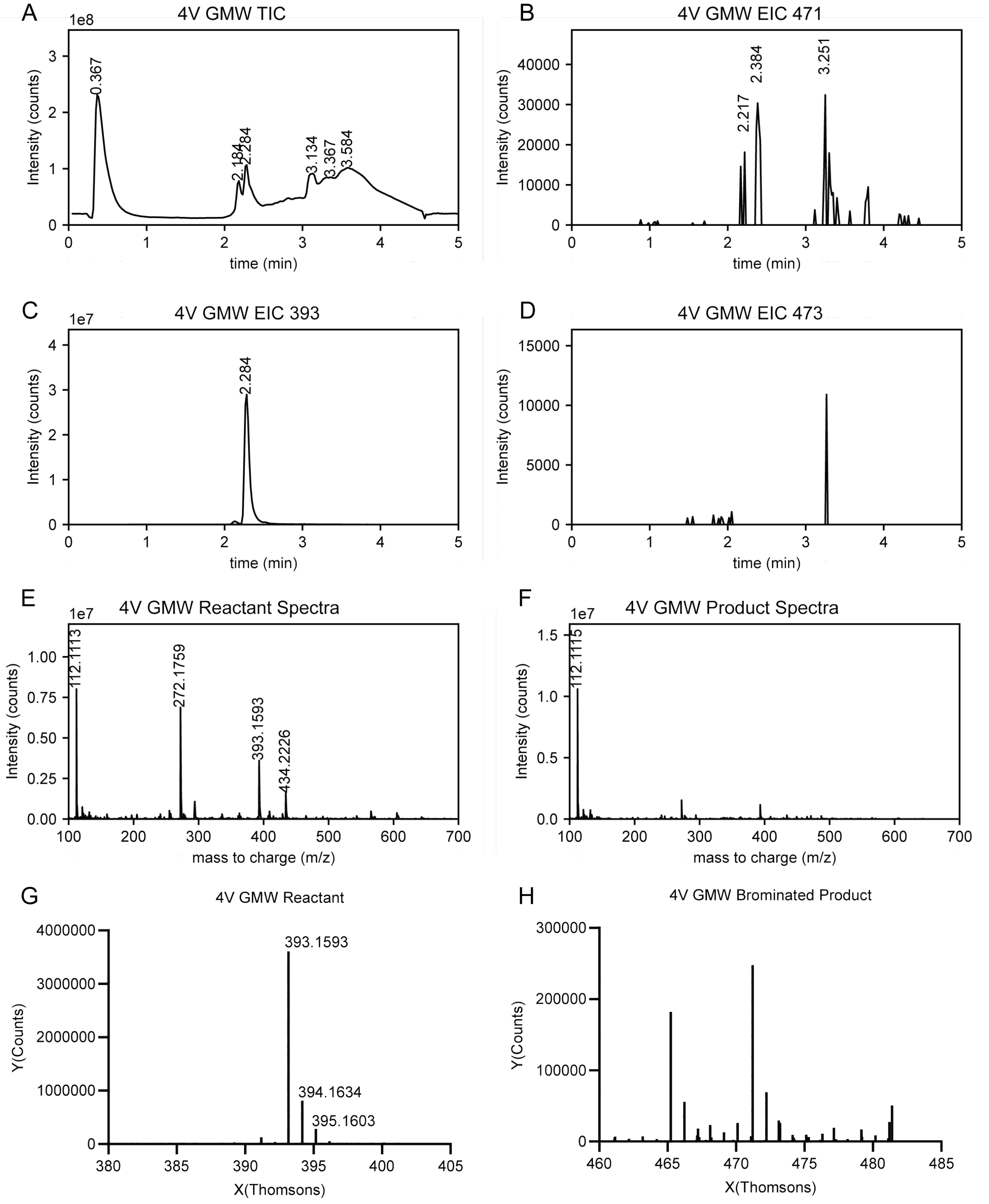
A
4V GMW TIC
1e8
0.367
2.184
2.284
3.134
3.367
3.584
Intensity (counts)
time (min)
B
4V GMW EIC 471
2.217
2.384
3.251
Intensity (counts)
time (min)
C
4V GMW EIC 393
1e7
2.284
Intensity (counts)
time (min)
D
4V GMW EIC 473
Intensity (counts)
time (min)
E
4V GMW Reactant Spectra
1e7
112.1113
272.1759
393.1593
434.2226
Intensity (counts)
mass to charge (m/z)
F
4V GMW Product Spectra
1e7
112.1115
Intensity (counts)
mass to charge (m/z)
G
4V GMW Reactant
393.1593
394.1634
395.1603
Y(Counts)
X(Thomsons)
H
4V GMW Brominated Product
Y(Counts)
X(Thomsons)

#### 4V GNW

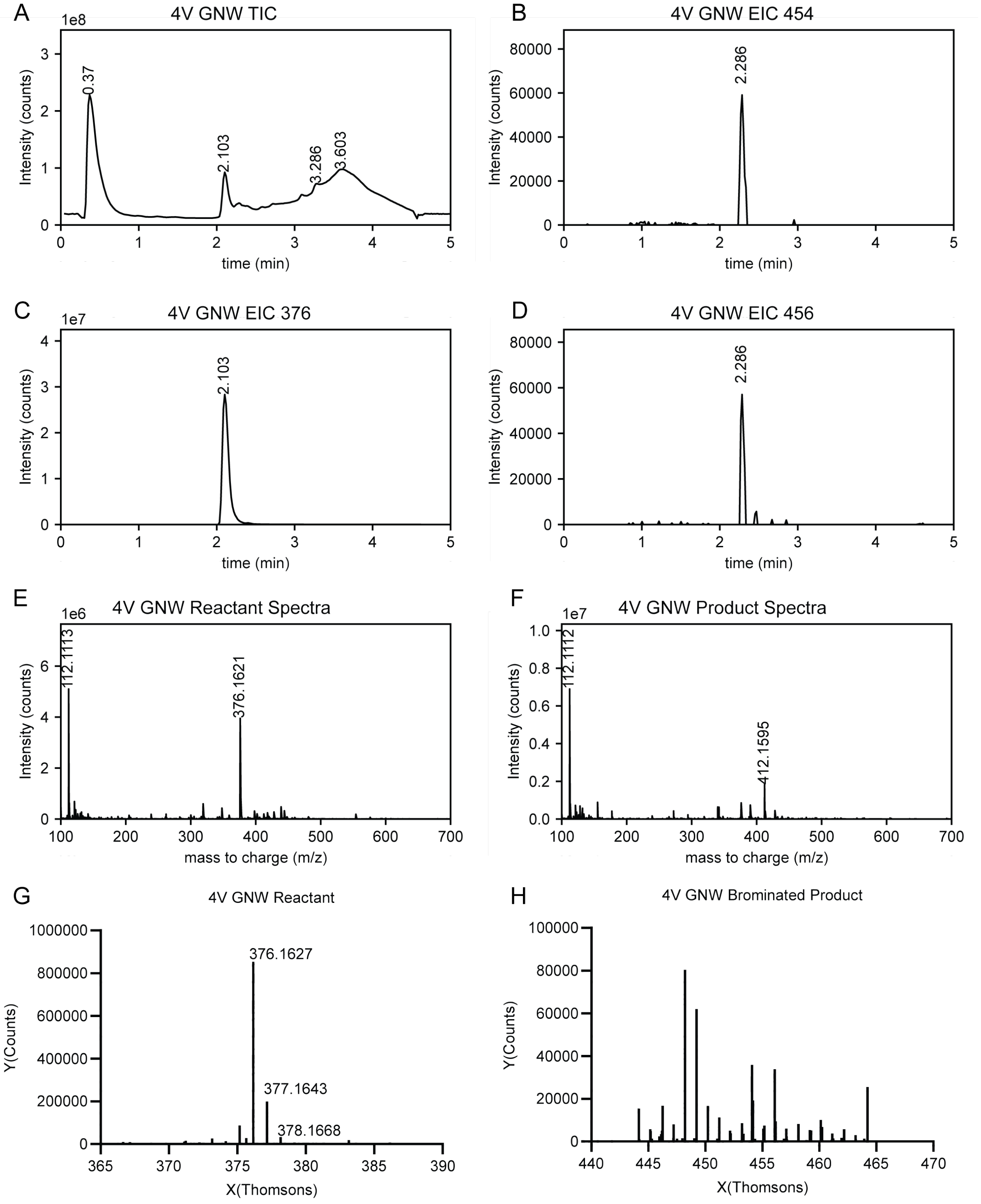

A
4V GNW TIC
1e8
Intensity (counts)
time (min)
0.37
2.103
3.286
3.603
B
4V GNW EIC 454
2.286
C
4V GNW EIC 376
1e7
2.103
D
4V GNW EIC 456
2.286
E
4V GNW Reactant Spectra
1e6
112.1113
376.1621
mass to charge (m/z)
F
4V GNW Product Spectra
1e7
112.1112
412.1595
G
4V GNW Reactant
376.1627
377.1643
378.1668
Y(Counts)
X(Thomsons)
H
4V GNW Brominated Product

#### 4V GPW

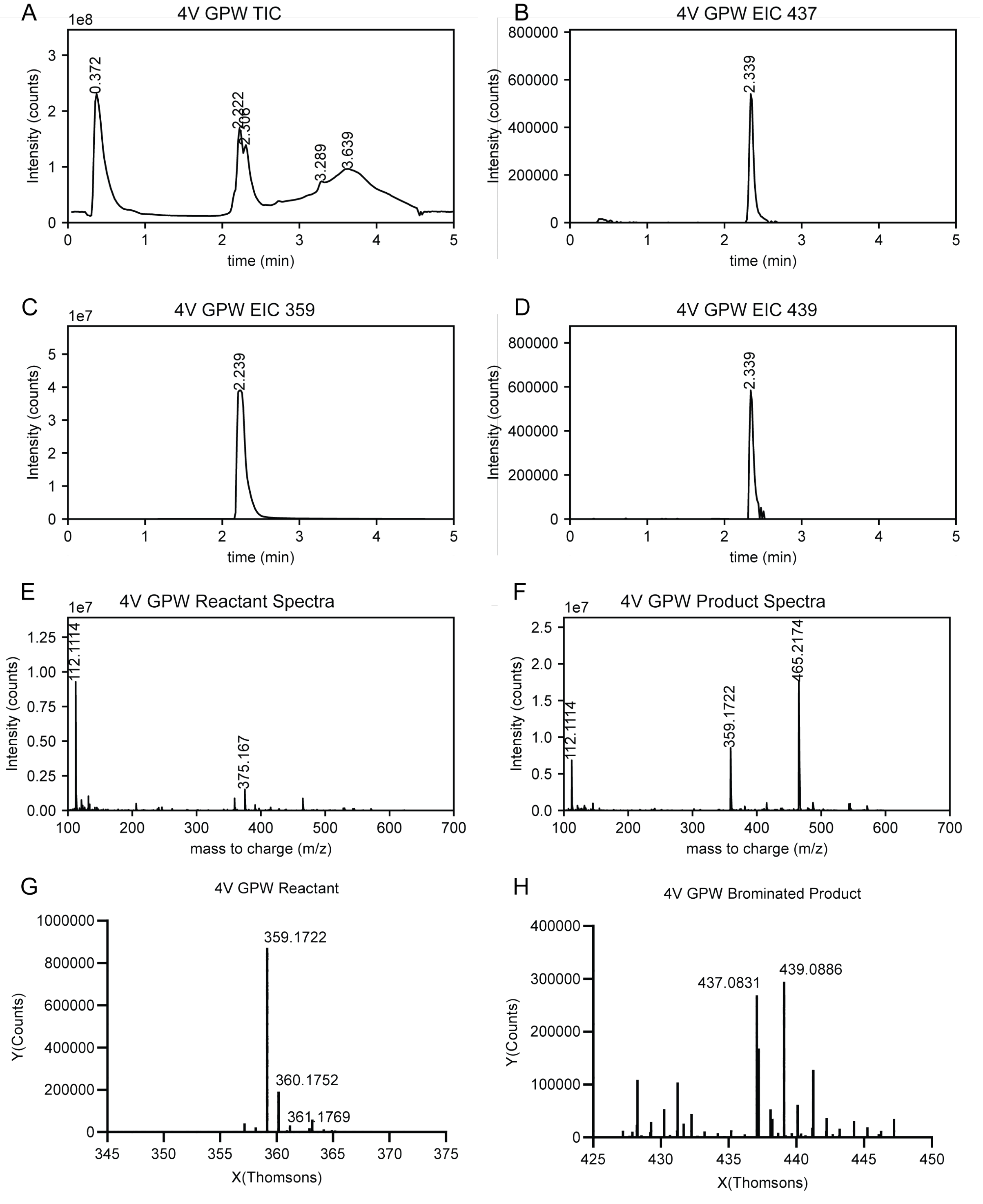
A
4V GPW TIC
1e8
Intensity (counts)
0.372
2.222
2.306
3.289
3.639
time (min)
B
4V GPW EIC 437
2.339
Intensity (counts)
time (min)
C
4V GPW EIC 359
1e7
2.239
Intensity (counts)
time (min)
D
4V GPW EIC 439
2.339
Intensity (counts)
time (min)
E
4V GPW Reactant Spectra
1e7
112.1114
375.167
Intensity (counts)
mass to charge (m/z)
F
4V GPW Product Spectra
1e7
112.1114
359.1722
465.2174
Intensity (counts)
mass to charge (m/z)
G
4V GPW Reactant
359.1722
360.1752
361.1769
Y(Counts)
X(Thomsons)
H
4V GPW Brominated Product
437.0831
439.0886
Y(Counts)
X(Thomsons)

#### 4V GQW

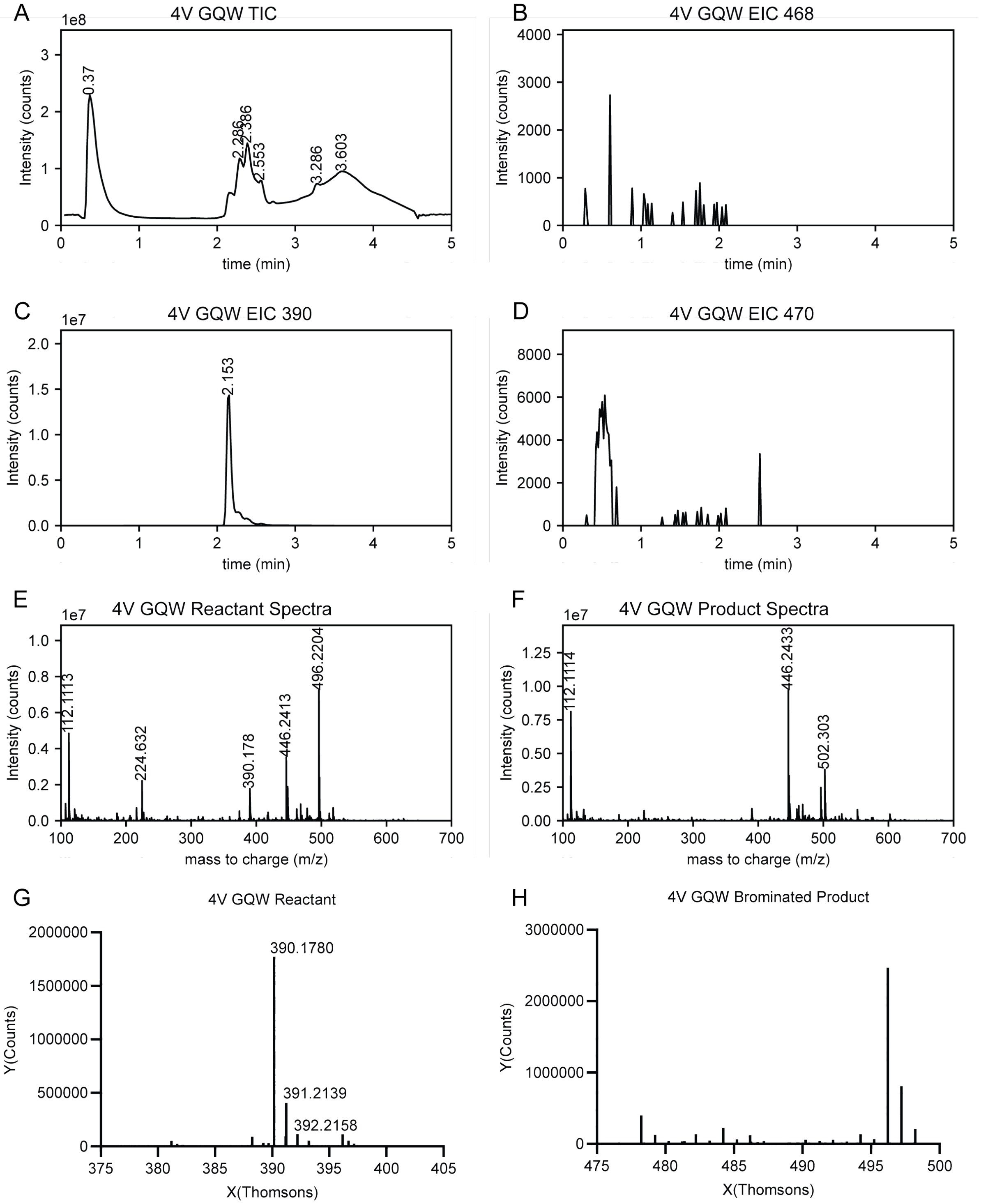

A
4V GQW TIC
1e8
Intensity (counts)
time (min)
0.37
2.286
2.386
2.553
3.286
3.603
B
4V GQW EIC 468
Intensity (counts)
time (min)
C
4V GQW EIC 390
1e7
Intensity (counts)
time (min)
2.153
D
4V GQW EIC 470
Intensity (counts)
time (min)
E
4V GQW Reactant Spectra
1e7
Intensity (counts)
mass to charge (m/z)
112.1113
224.632
390.178
446.2413
496.2204
F
4V GQW Product Spectra
1e7
Intensity (counts)
mass to charge (m/z)
112.1114
446.2433
502.303
G
4V GQW Reactant
Y(Counts)
X(Thomsons)
390.1780
391.2139
392.2158
H
4V GQW Brominated Product
Y(Counts)
X(Thomsons)

#### 4V GRW

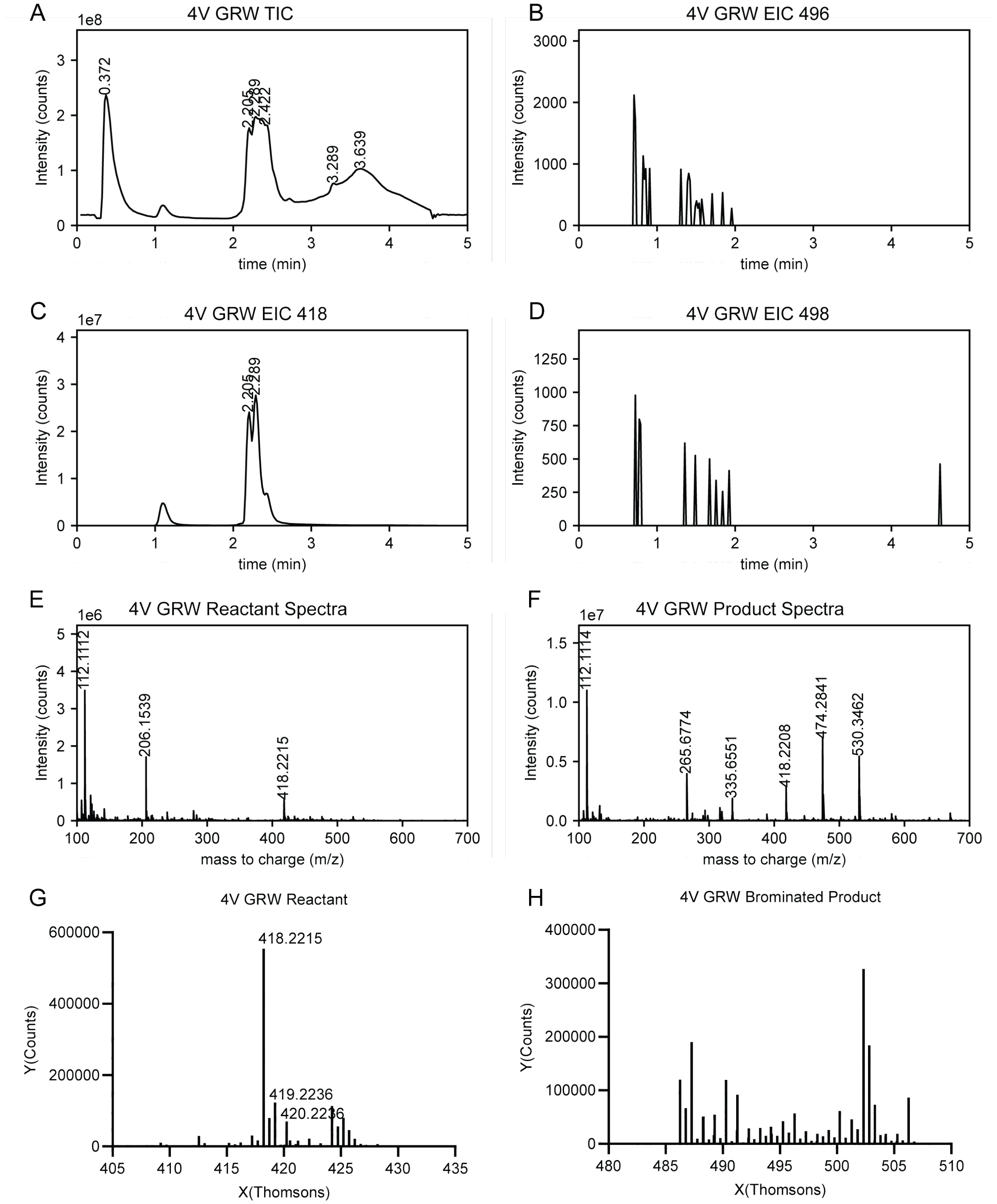

A
4V GRW TIC
1e8
Intensity (counts)
time (min)
0.372
2.205
2.289
2.422
3.289
3.639
B
4V GRW EIC 496
Intensity (counts)
time (min)
C
4V GRW EIC 418
1e7
Intensity (counts)
time (min)
2.205
2.289
D
4V GRW EIC 498
Intensity (counts)
time (min)
E
4V GRW Reactant Spectra
1e6
Intensity (counts)
mass to charge (m/z)
112.1112
206.1539
418.2215
F
4V GRW Product Spectra
1e7
Intensity (counts)
mass to charge (m/z)
112.1114
265.6774
335.6551
418.2208
474.2841
530.3462
G
4V GRW Reactant
Y(Counts)
X(Thomsons)
418.2215
419.2236
420.2236
H
4V GRW Brominated Product
Y(Counts)
X(Thomsons)

#### 4V GTW

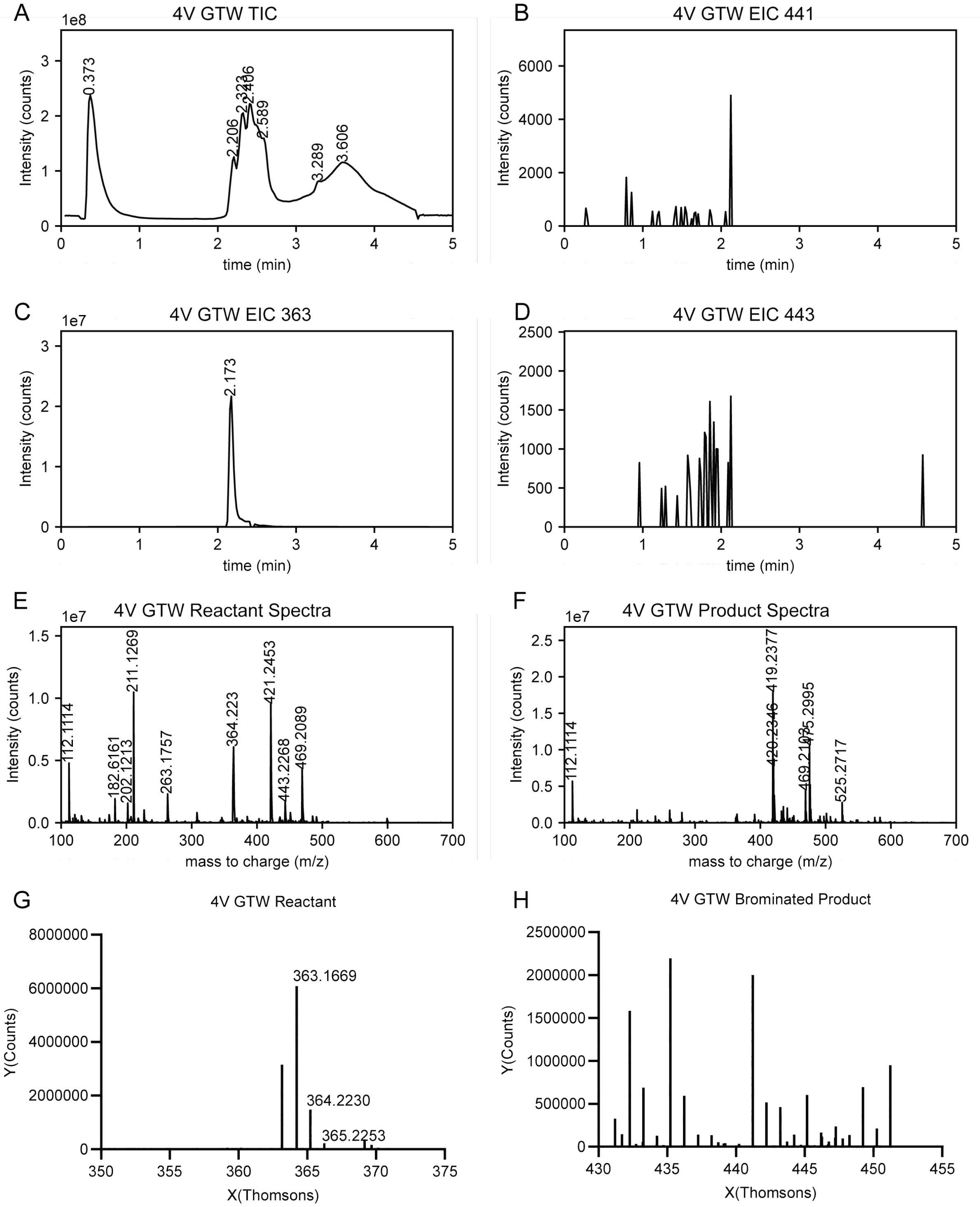

A
4V GTW TIC
1e8
Intensity (counts)
time (min)
0.373
2.206
2.323
2.406
2.589
3.289
3.606
B
4V GTW EIC 441
Intensity (counts)
time (min)
C
4V GTW EIC 363
1e7
Intensity (counts)
time (min)
2.173
D
4V GTW EIC 443
Intensity (counts)
time (min)
E
4V GTW Reactant Spectra
1e7
Intensity (counts)
mass to charge (m/z)
112.1114
182.6161
202.1213
211.1269
263.1757
364.223
421.2453
443.2268
469.2089
F
4V GTW Product Spectra
1e7
Intensity (counts)
mass to charge (m/z)
112.1114
419.2377
469.2107
475.2995
525.2717
G
4V GTW Reactant
Y(Counts)
X(Thomsons)
363.1669
364.2230
365.2253
H
4V GTW Brominated Product
Y(Counts)
X(Thomsons)

**4V GVW**

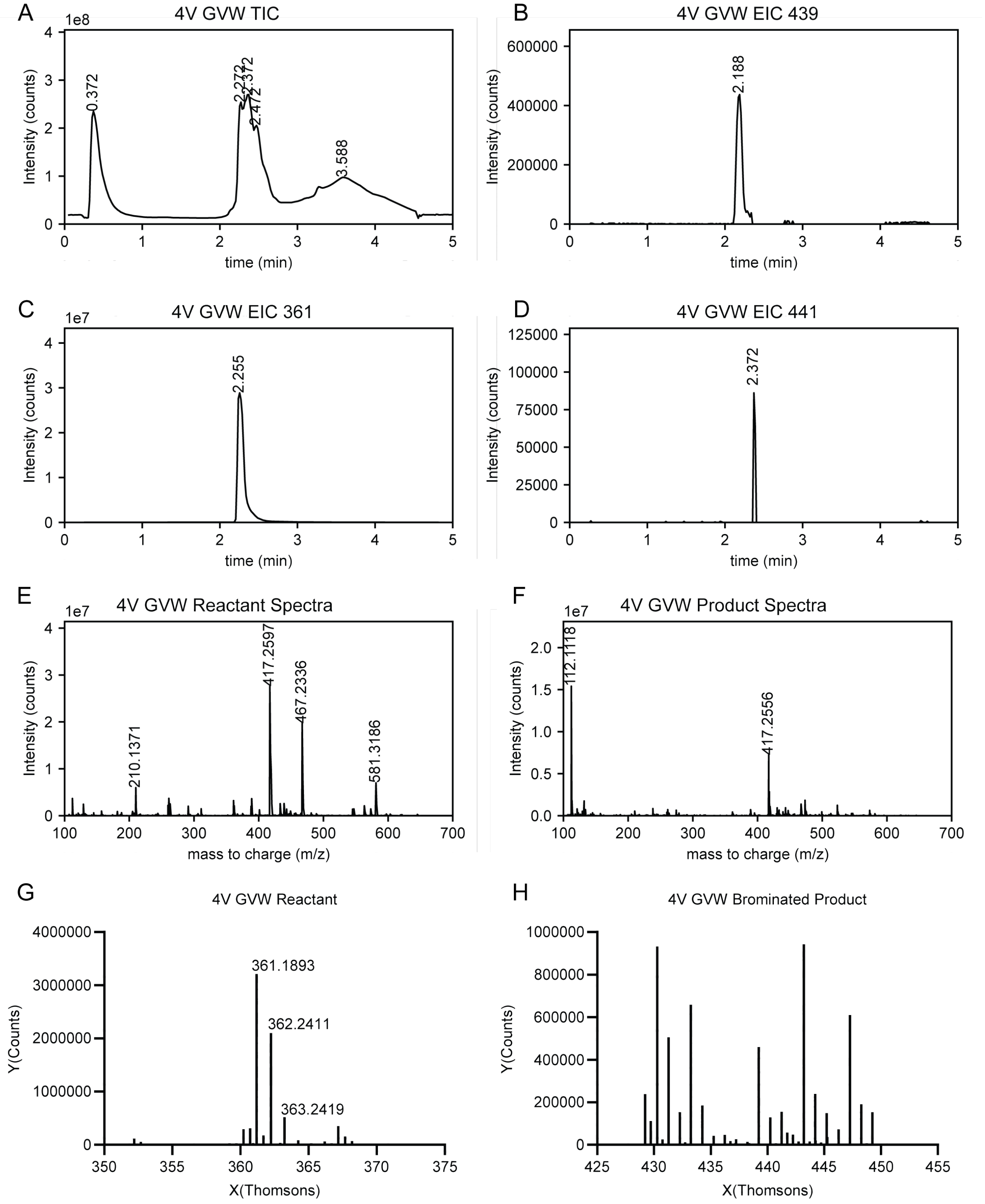
A
4V GVW TIC
1e8
0.372
2.272
2.372
2.472
3.588
Intensity (counts)
time (min)
B
4V GVW EIC 439
2.188
Intensity (counts)
time (min)
C
4V GVW EIC 361
1e7
2.255
Intensity (counts)
time (min)
D
4V GVW EIC 441
2.372
Intensity (counts)
time (min)
E
4V GVW Reactant Spectra
1e7
210.1371
417.2597
467.2336
581.3186
Intensity (counts)
mass to charge (m/z)
F
4V GVW Product Spectra
1e7
112.1118
417.2556
Intensity (counts)
mass to charge (m/z)
G
4V GVW Reactant
361.1893
362.2411
363.2419
Y(Counts)
X(Thomsons)
H
4V GVW Brominated Product
Y(Counts)
X(Thomsons)

**4V GYW**

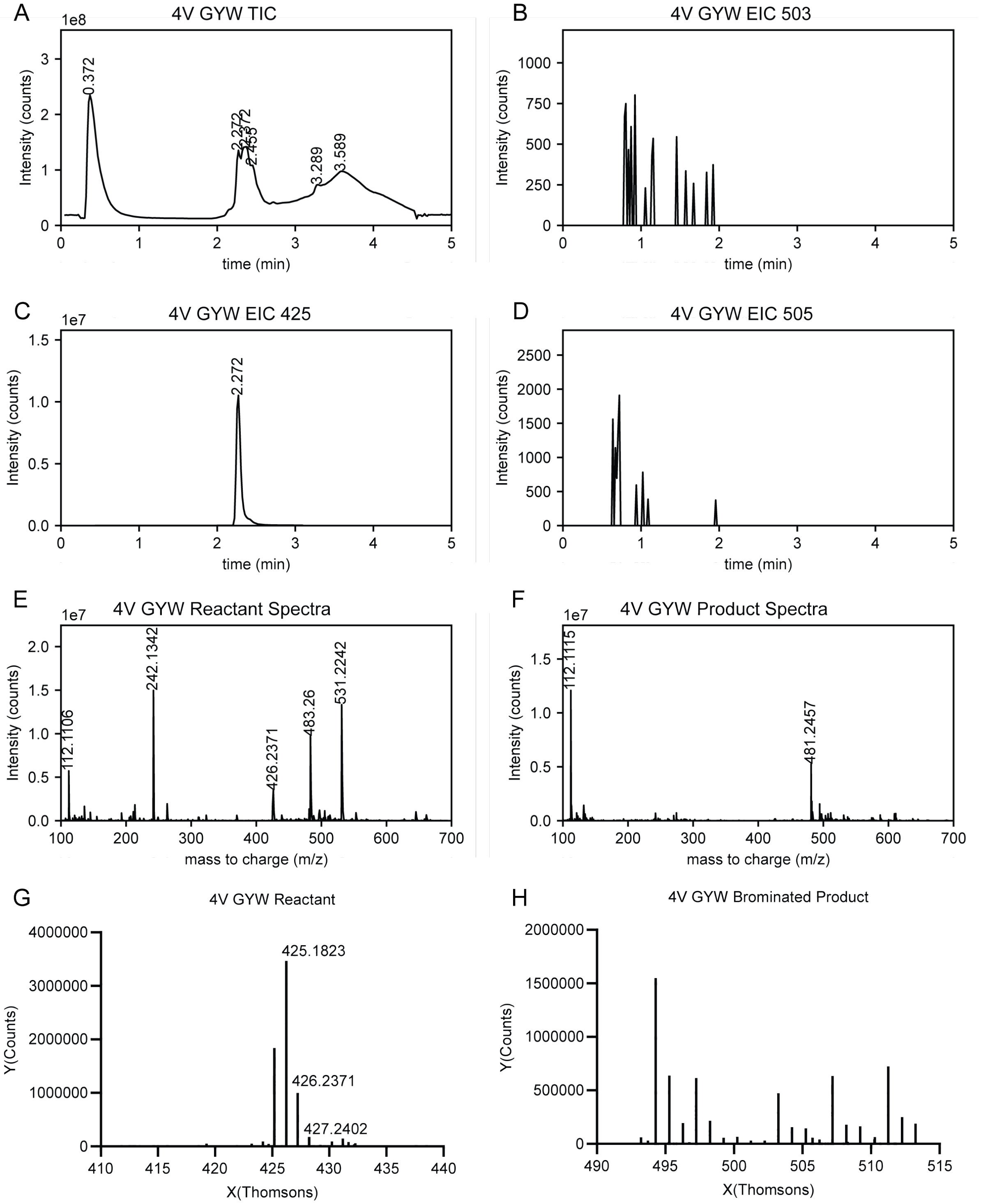
A
4V GYW TIC
1e8
Intensity (counts)
time (min)
0.372
2.272
2.372
2.453
3.289
3.589
B
4V GYW EIC 503
Intensity (counts)
time (min)
C
4V GYW EIC 425
1e7
Intensity (counts)
time (min)
2.272
D
4V GYW EIC 505
Intensity (counts)
time (min)
E
4V GYW Reactant Spectra
1e7
Intensity (counts)
mass to charge (m/z)
112.1106
242.1342
426.2371
483.26
531.2242
F
4V GYW Product Spectra
1e7
Intensity (counts)
mass to charge (m/z)
112.1115
481.2457
G
4V GYW Reactant
Y(Counts)
X(Thomsons)
425.1823
426.2371
427.2402
H
4V GYW Brominated Product
Y(Counts)
X(Thomsons)

#### SMC Modified 4V Brominated Endomorphin-1 Mass Spectra

#### 4V D-Endomorphin-1

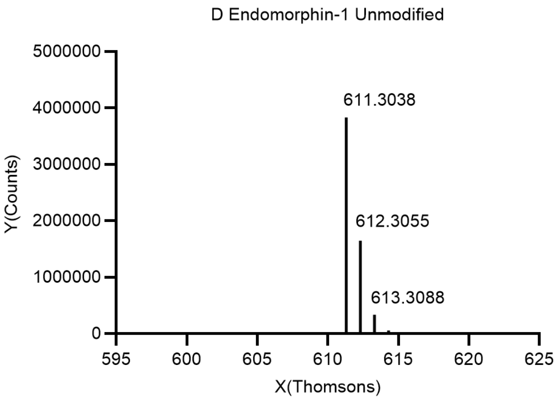


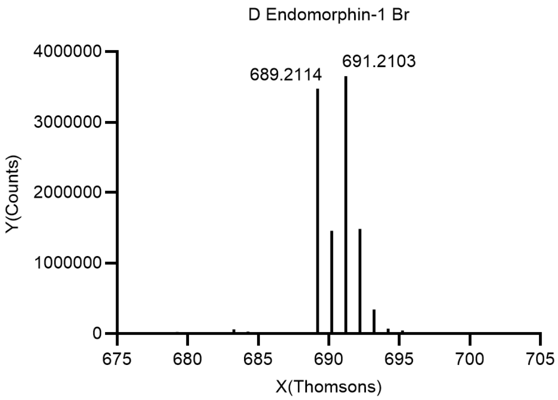


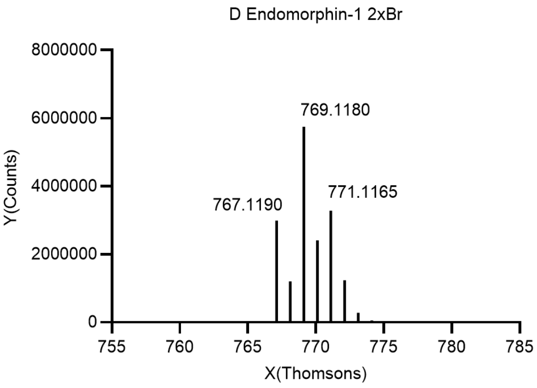


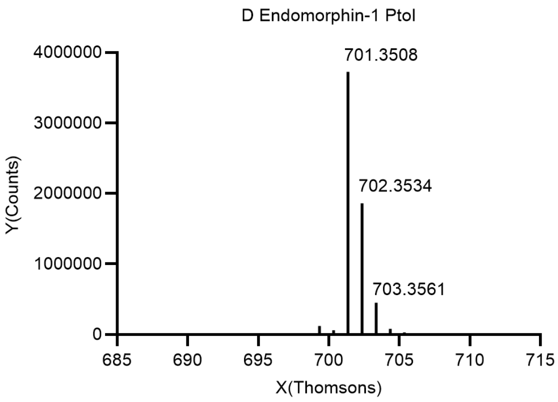


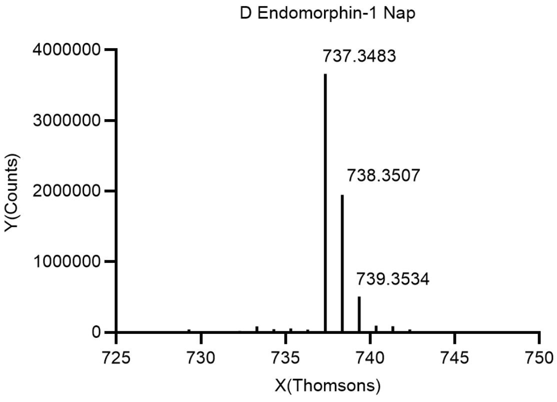


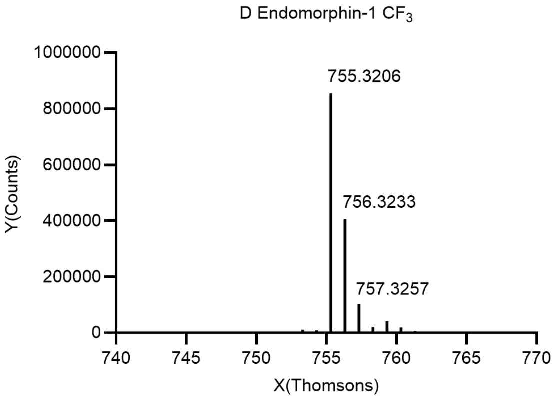

#### 4V L-Endomorphin-1

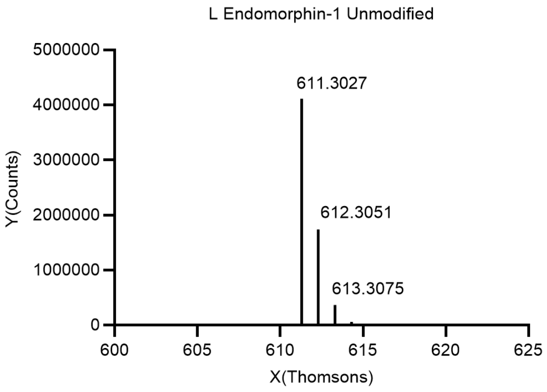


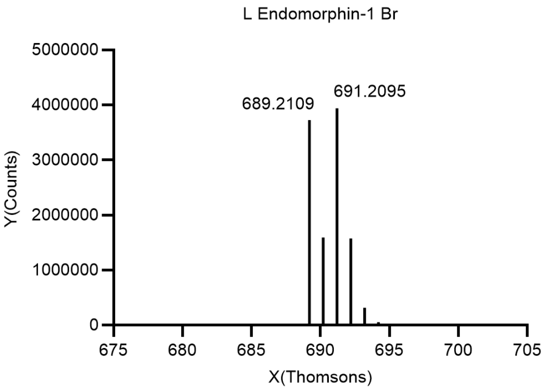


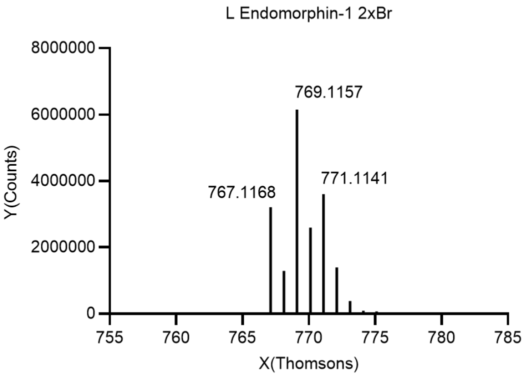


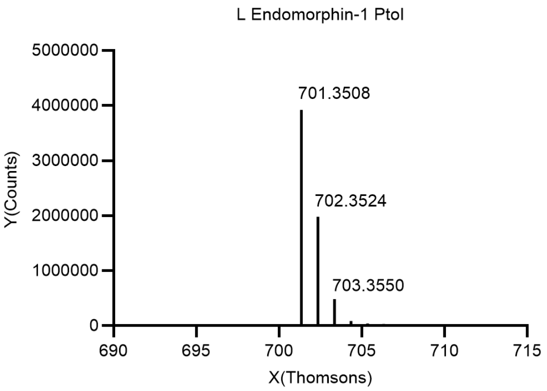


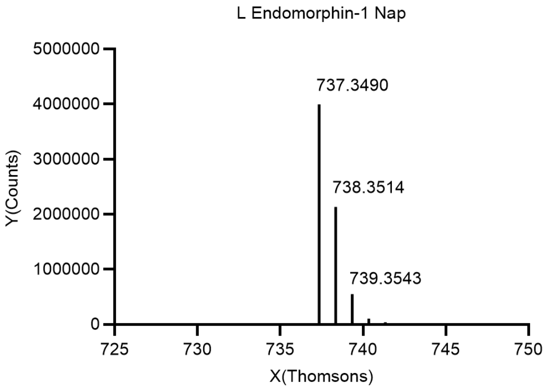


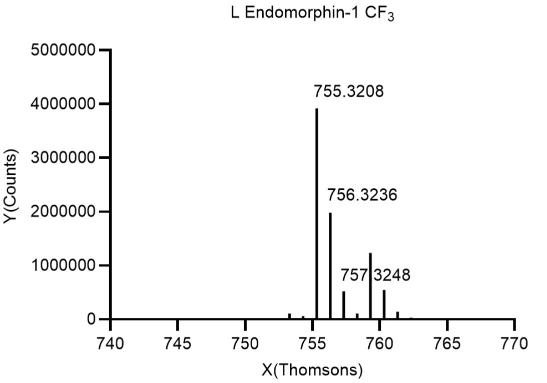

#### 4V Bioactive Peptide Bromination Mass Spectra

#### 4V L-Endomorphin-1

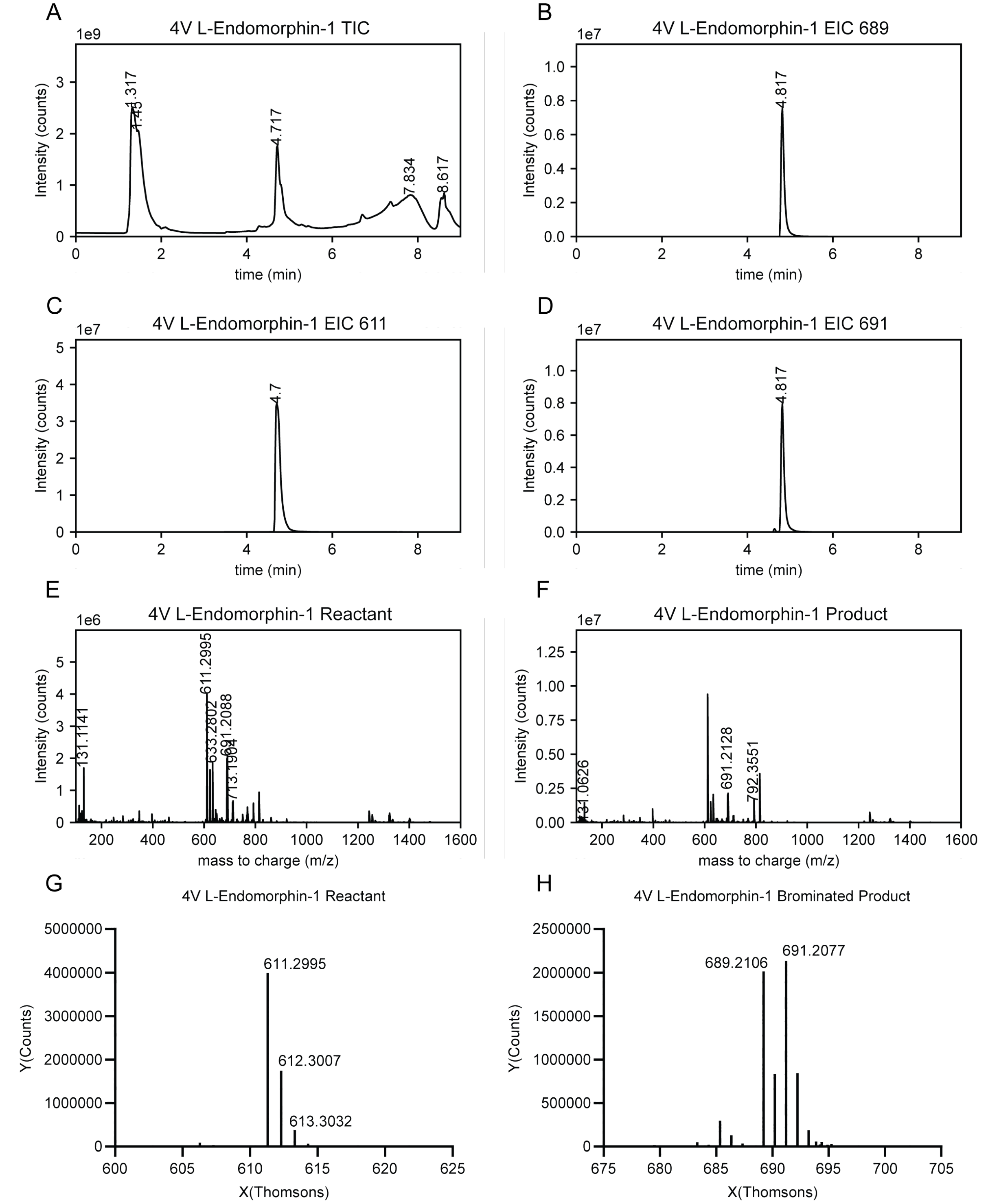

#### 4V D-Endomorphin-1

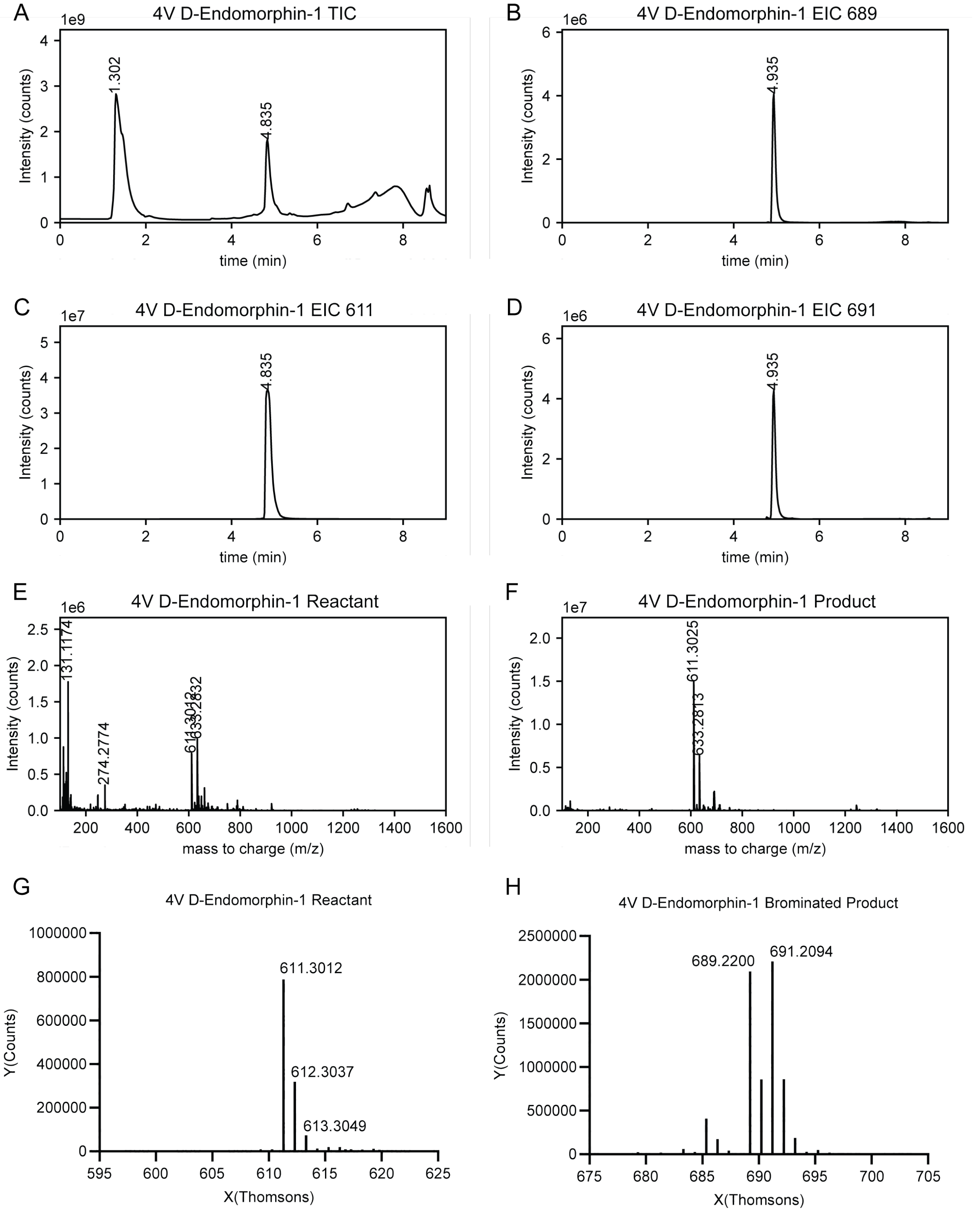

#### 4V Human Galanin

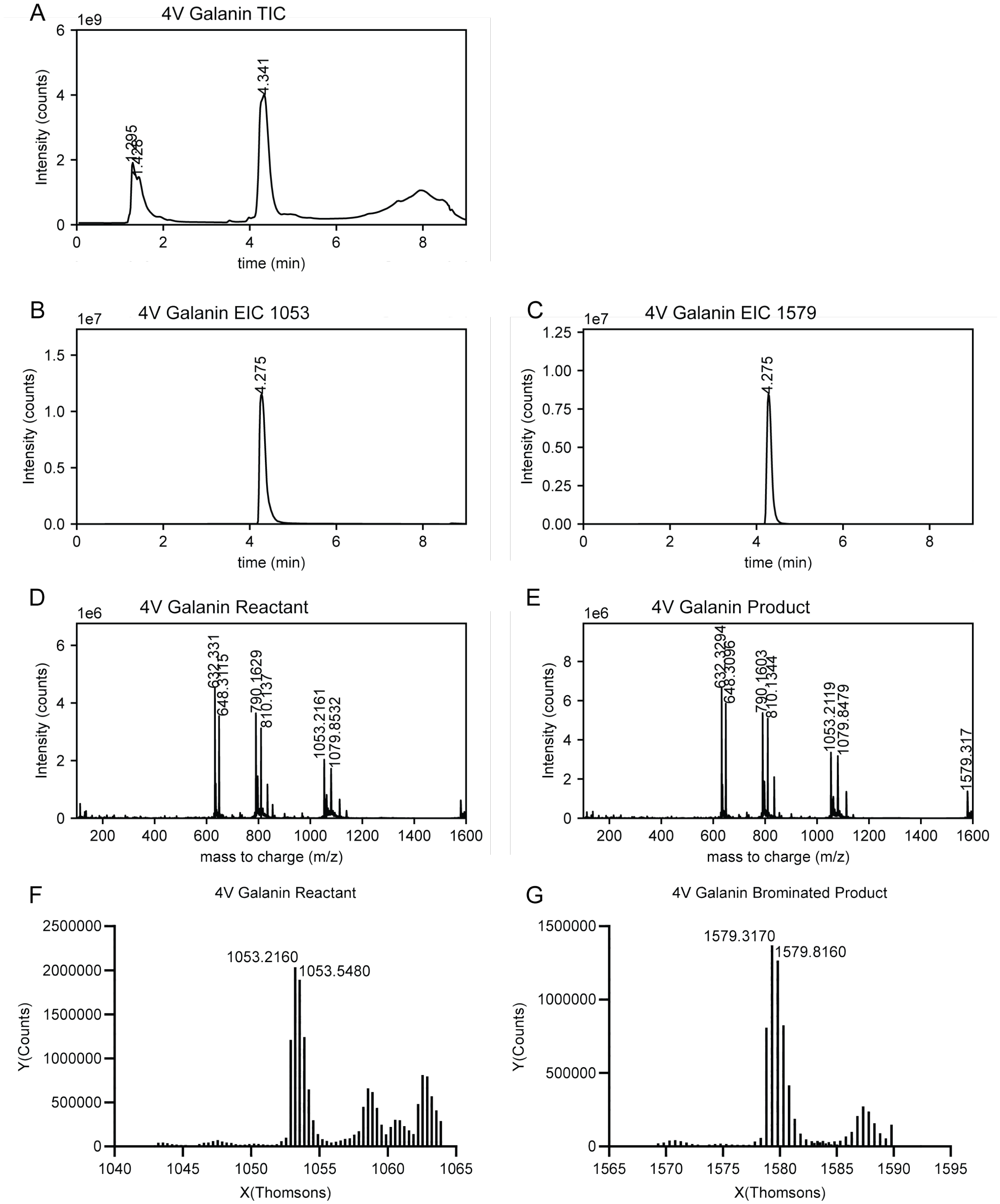

#### 4V Human Neuropeptide W23

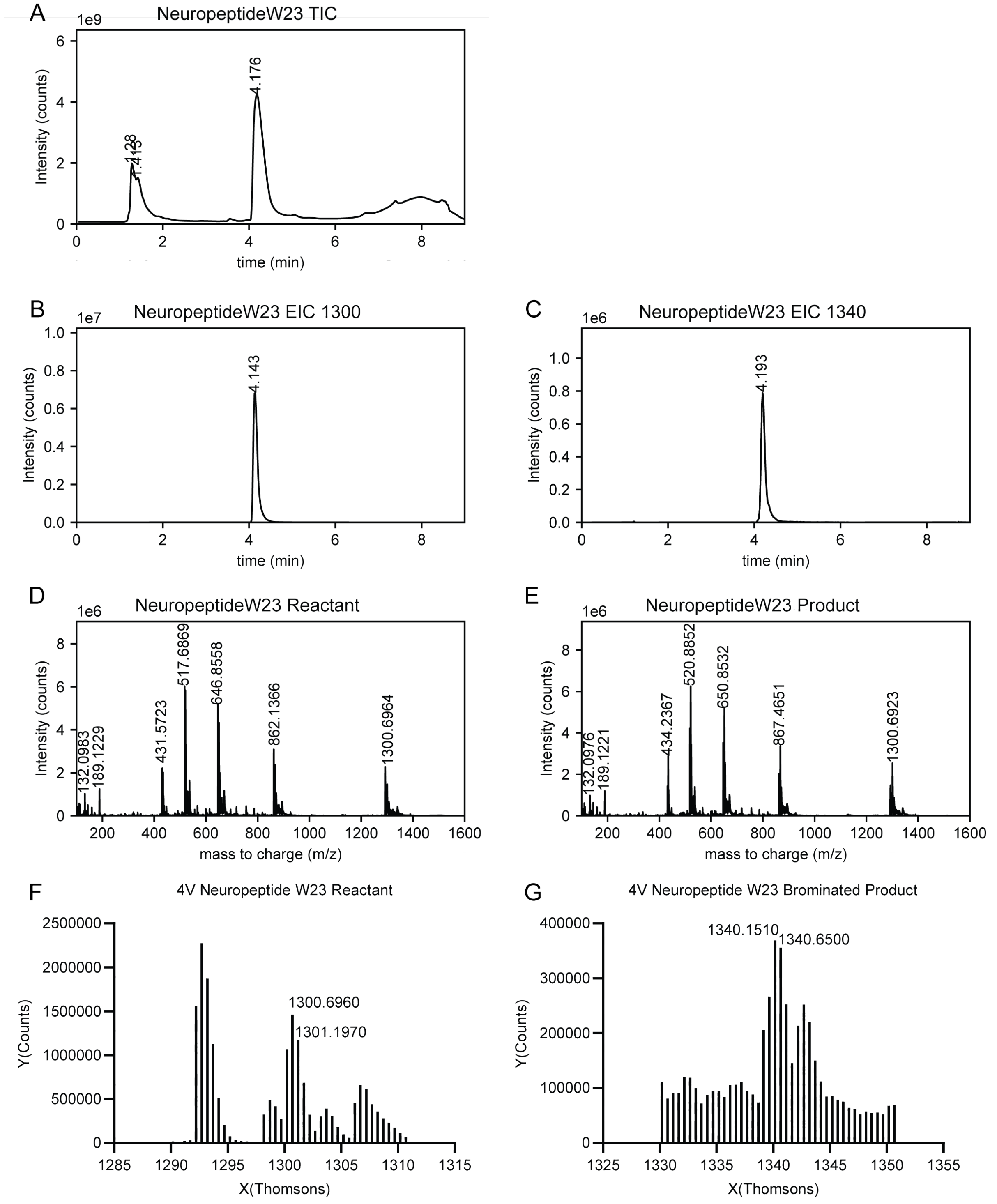

## 4V $(RW)_3$

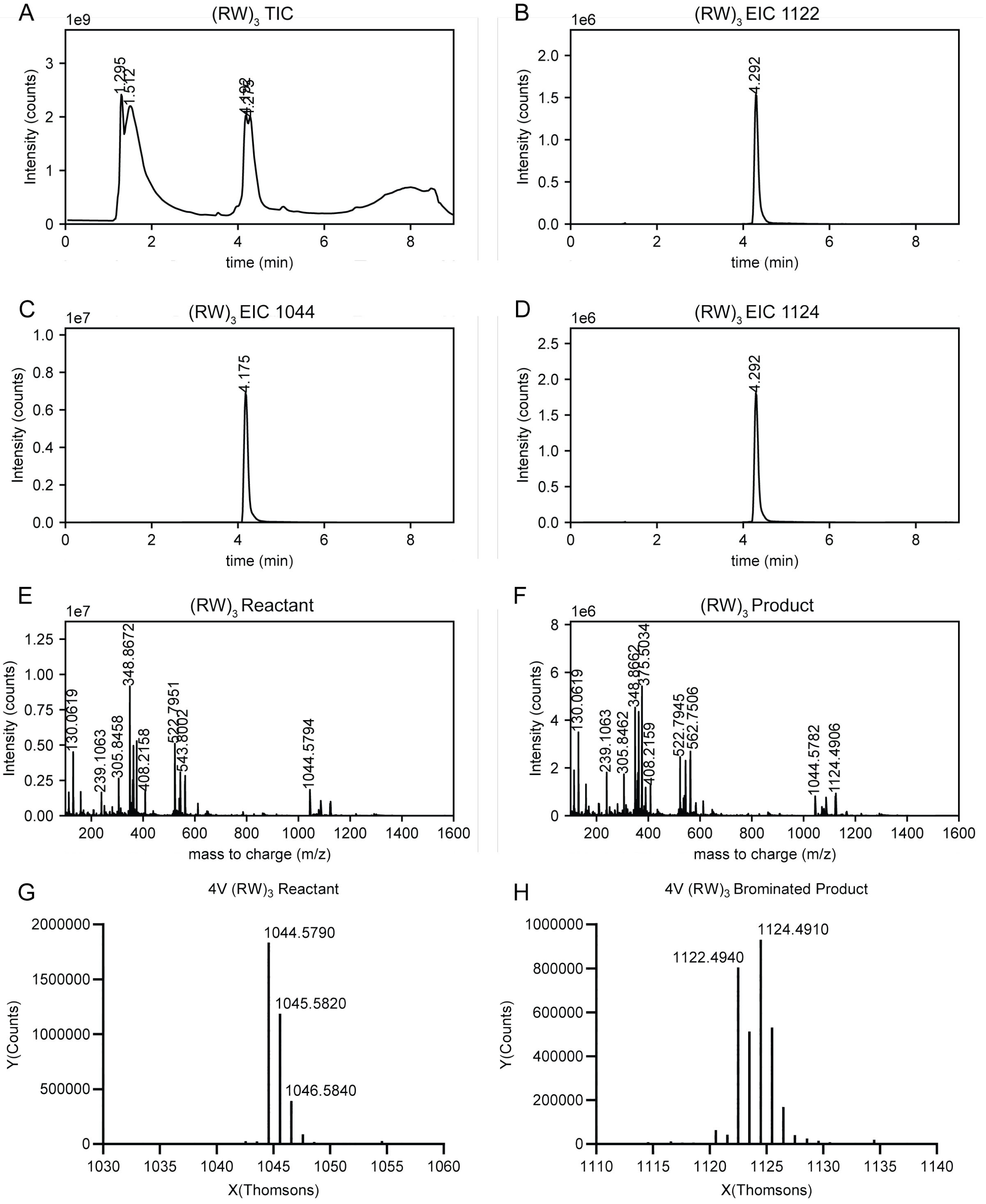

A
(RW)3 TIC
1e9
Intensity (counts)
time (min)
1.295
1.512
B
(RW)3 EIC 1122
1e6
4.292
C
(RW)3 EIC 1044
1e7
4.175
D
(RW)3 EIC 1124
1e6
4.292
E
(RW)3 Reactant
1e7
mass to charge (m/z)
130.0619
239.1063
305.8458
348.8672
408.2158
522.7951
543.8002
1044.5794
F
(RW)3 Product
1e6
130.0619
239.1063
305.8462
348.8662
375.5034
408.2159
522.7945
562.7506
1044.5782
1124.4906
G
4V (RW)3 Reactant
Y(Counts)
X(Thomsons)
1044.5790
1045.5820
1046.5840
H
4V (RW)3 Brominated Product
1122.4940
1124.4910

## 4V $(RW)_4$

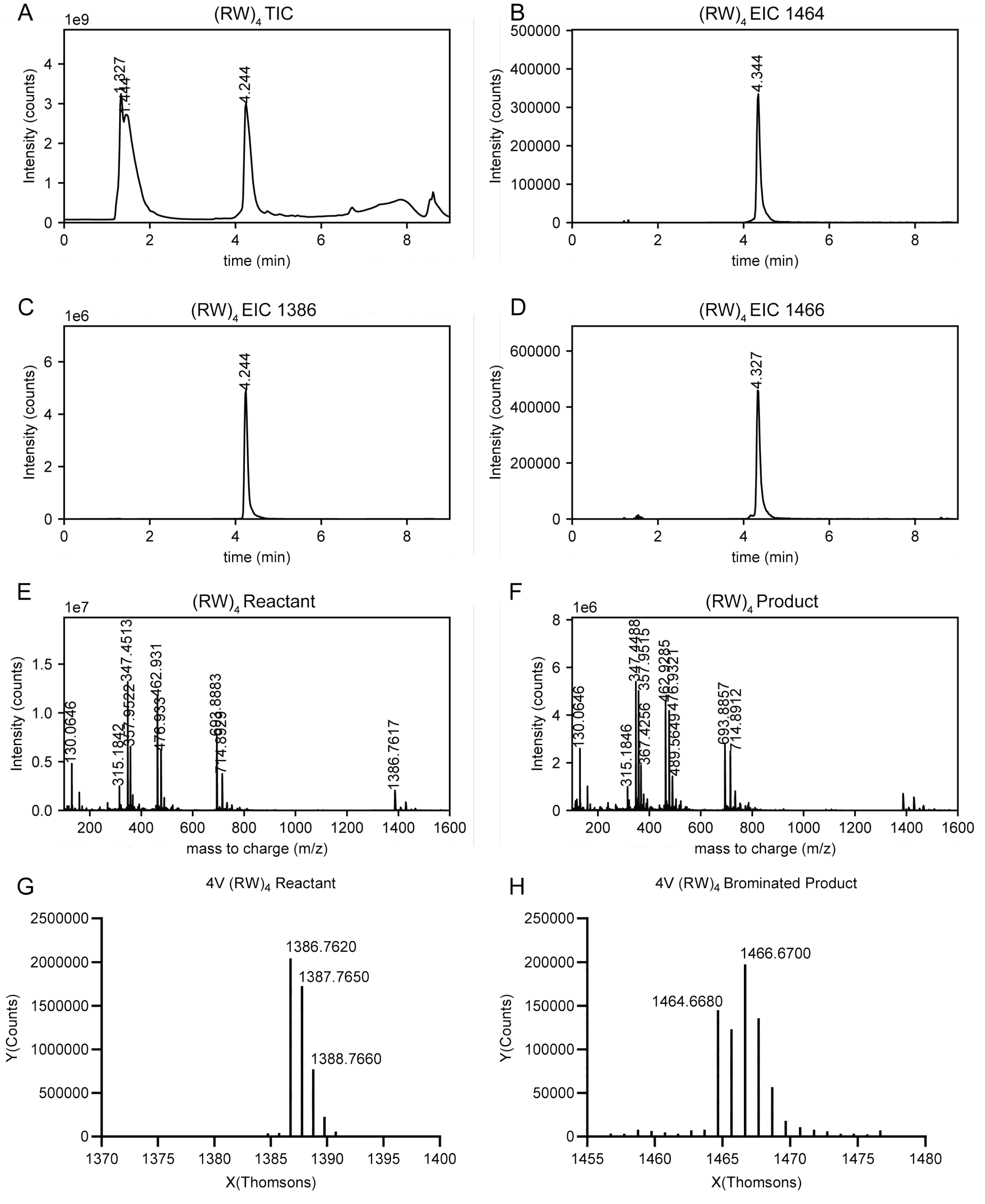

A
(RW)4 TIC
B
(RW)4 EIC 1464
C
(RW)4 EIC 1386
D
(RW)4 EIC 1466
E
(RW)4 Reactant
F
(RW)4 Product
G
4V (RW)4 Reactant
1386.7620
1387.7650
1388.7660
H
4V (RW)4 Brominated Product
1464.6680
1466.6700
Intensity (counts)
time (min)
mass to charge (m/z)
Y(Counts)
X(Thomsons)

#### 4V Human Spexin-1

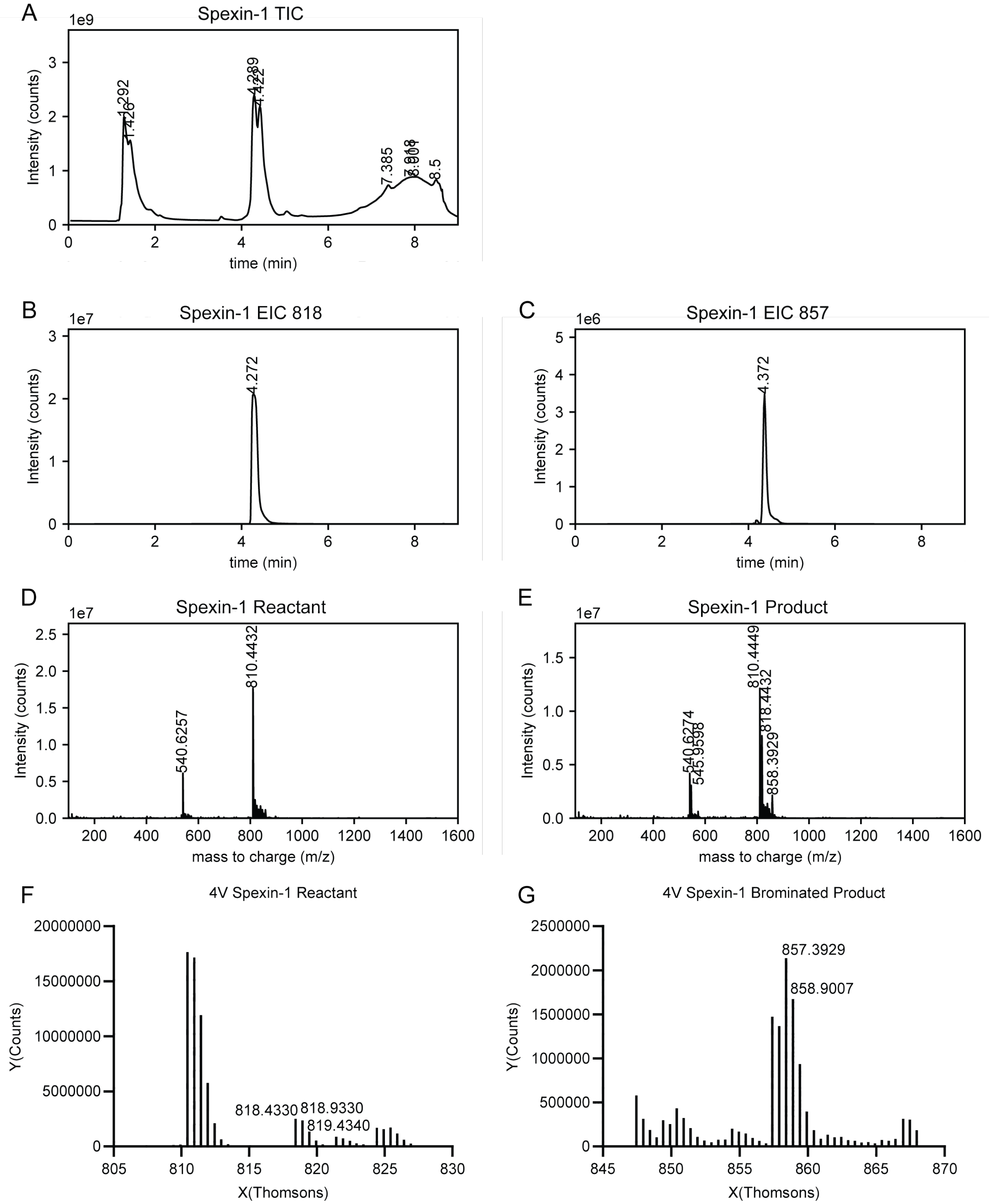

#### 4V Transportan

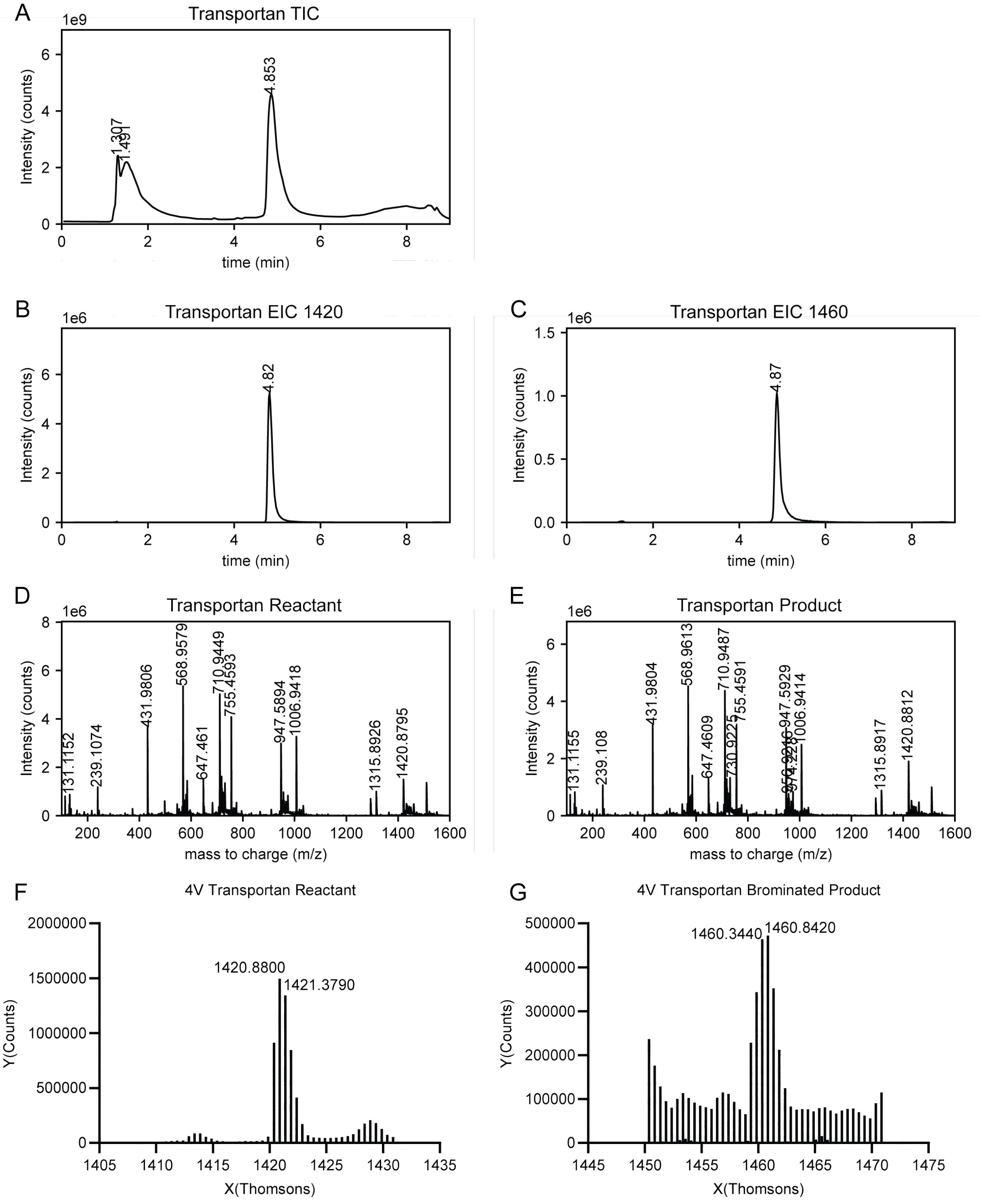

#### Amino Acid Calculated and Found Mass List

| Enzyme | Substrate | Charge State | Calculated Product | Calculated Reactant | Found Product | Found Reactant |
|---|---|---|---|---|---|---|
| No RebH-Br | D-W | $[M+H^+]$ | 239.0582, 241.0552 | 205.0972 | Not found | 205.1030 |
| No RebH-Br | L-W | $[M+H^+]$ | 239.0582, 241.0552 | 205.0972 | Not found | 205.1028 |
| No RebH-Cl | D-W | $[M+H^+]$ | 283.0077, 285.0056 | 205.0972 | Not found | 205.1001 |
| No RebH-Cl | L-W | $[M+H^+]$ | 283.0077, 285.0056 | 205.0972 | Not found | 205.0995 |
| RebH-Br | D-W | $[M+H^+]$ | 283.0077, 285.0056 | 205.0972 | 283.0125, 285.0105 | Not found |
| RebH-Br | L-W | $[M+H^+]$ | 283.0077, 285.0056 | 205.0972 | 283.0102, 285.0081 | 205.0976 |
| RebH-Cl | D-W | $[M+H^+]$ | 239.0582, 241.0552 | 205.0972 | 239.0787, 241.0642 | 205.0995 |
| RebH-Cl | L-W | $[M+H^+]$ | 239.0582, 241.0552 | 205.0972 | 239.0669, 241.0593 | Not found |
| 4V-Br | D-W | $[M+H^+]$ | 283.0077, 285.0056 | 205.0972 | 283.0090, 285.0070 | 205.0985 |
| 4V-Br | L-W | $[M+H^+]$ | 283.0077, 285.0056 | 205.0972 | 283.0092, 285.0072 | 205.0970 |
| 4V-Cl | D-W | $[M+H^+]$ | 239.0582, 241.0552 | 205.0972 | 239.0768, 241.0630 | 205.0995 |
| 4V-Cl | L-W | $[M+H^+]$ | 239.0582, 241.0552 | 205.0972 | 239.0682, 241.0597 | 205.0997 |

#### Glycine Peptides of Increasing Length Calculated and Found Mass List

| Enzyme | Substrate | Charge State | Calculated Product | Calculated Reactant | Found Product | Found Reactant |
|---|---|---|---|---|---|---|
| RebH | GGW | $[M+H^+]$ | 396.0666, 398.0645 | 318.1561 | 396.0699, 398.0679 | 318.1582 |
| RebH | GGGW | $[M+H^+]$ | 453.088, 455.0860 | 375.1775 | 453.0968, 455.0860 | 371.464 |
| RebH | GGGGW | $[M+H^+]$ | 510.1095, 512.1075 | 432.1990 | 510.1168, 512.1150 | 432.2077 |
| RebH | GGGGGW | $[M+H^+]$ | 567.1310, 569.1289 | 489.2205 | 567.1385, 569.1369 | 489.2313 |
| RebH | WGG | $[M+H^+]$ | 396.0666, 398.0645 | 318.1561 | 396.0674, 398.0656 | 318.1577 |
| RebH | WGGG | $[M+H^+]$ | 453.088, 455.0860 | 375.1775 | 453.0890, 455.0875 | 375.1809 |
| RebH | WGGGG | $[M+H^+]$ | 510.1095, 512.1075 | 432.1990 | 510.1103, 512.1085 | 432.2003 |
| RebH | WGGGGG | $[M+H^+]$ | 567.1310, 569.1289 | 489.2205 | 567.1318, 569.1301 | 489.2221 |
| RebH | GWG | $[M+H^+]$ | 396.0666, 398.0645 | 318.1561 | 396.0689, 398.0669 | 318.1408 |
| RebH | GGWGG | $[M+H^+]$ | 510.1095, 512.1075 | 432.1990 | 510.1142, 512.1125 | 432.2012 |
| 4V | GGW | $[M+H^+]$ | 396.0666, 398.0645 | 318.1561 | 396.0698, 398.0681 | Not found |
| 4V | GGGW | $[M+H^+]$ | 453.088, 455.0860 | 375.1775 | 453.0972, 455.0954 | 375.1908 |
| 4V | GGGGW | $[M+H^+]$ | 510.1095, 512.1075 | 432.1990 | 510.1168, 512.1151 | 432.2071 |
| 4V | GGGGGW | $[M+H^+]$ | 567.1310, 569.1289 | 489.2205 | 567.1379, 569.1363 | 489.2312 |
| 4V | WGG | $[M+H^+]$ | 396.0666, 398.0645 | 318.1561 | 396.0678, 398.0659 | 318.1577 |
| 4V | WGGG | $[M+H^+]$ | 453.088, 455.0860 | 375.1775 | 453.0893, 455.0874 | 375.1813 |
| 4V | WGGGG | $[M+H^+]$ | 510.1095, 512.1075 | 432.1990 | 510.1105, 512.1089 | 432.2053 |
| 4V | WGGGGG | $[M+H^+]$ | 567.1310, 569.1289 | 489.2205 | 567.1322, 569.1305 | 489.2223 |
| 4V | GWG | $[M+H^+]$ | 396.0666, 398.0645 | 318.1561 | 396.0705, 398.0681 | Not found |
| 4V | GGWGG | $[M+H^+]$ | 510.1095, 512.1075 | 432.1990 | 510.1126, 512.1107 | 432.2022 |
| no RebH | GGW | $[M+H^+]$ | 396.0666, 398.0645 | 318.1561 | Not found | 318.1573 |
| no RebH | GGGW | $[M+H^+]$ | 453.088, 455.0860 | 375.1775 | Not found | 375.1833 |
| no RebH | GGGGW | $[M+H^+]$ | 510.1095, 512.1075 | 432.1990 | Not found | 432.2011 |

| Enzyme | Substrate | Charge State | Calculated Product | Calculated Reactant | Found Product | Found Reactant |
|---|---|---|---|---|---|---|
| no RebH | GGGGGW | [M+H$^+$] | 567.1310, 569.1289 | 489.2205 | Not found | 489.2242 |
| no RebH | WGG | [M+H$^+$] | 396.0666, 398.0645 | 318.1561 | Not found | 318.1621 |
| no RebH | WGGG | [M+H$^+$] | 453.088, 455.0860 | 375.1775 | Not found | 375.1942 |
| no RebH | WGGGG | [M+H$^+$] | 510.1095, 512.1075 | 432.1990 | Not found | 432.2200 |
| no RebH | WGGGGG | [M+H$^+$] | 567.1310, 569.1289 | 489.2205 | Not found | 489.2560 |
| no RebH | GWG | [M+H$^+$] | 396.0666, 398.0645 | 318.1561 | Not found | 318.1598 |
| no RebH | GGWGG | [M+H$^+$] | 510.1095, 512.1075 | 432.1990 | Not found | 432.2226 |
| 4V | GW-COOH | [M+H$^+$] | 340.0291, 342.0271 | 262.1186 | 340.0339, 342.0320 | 262.1228 |
| 4V | WG-COOH | [M+H$^+$] | 340.0291, 342.0271 | 262.1186 | 340.0320, 342.0301 | 262.1215 |
| 4V | GGW-COOH | [M+H$^+$] | 397.0506, 399.0485 | 319.1401 | 397.0526, 399.0507 | 319.1447 |
| 4V | WGG-COOH | [M+H$^+$] | 397.0506, 399.0485 | 319.1401 | 397.0523, 399.0509 | 319.1417 |
| RebH | GW-COOH | [M+H$^+$] | 340.0291, 342.0271 | 262.1186 | 340.0334, 342.0314 | 262.1222 |
| RebH | WG-COOH | [M+H$^+$] | 340.0291, 342.0271 | 262.1186 | 340.0327, 342.0308 | 262.1207 |
| RebH | GGW-COOH | [M+H$^+$] | 397.0506, 399.0485 | 319.1401 | 397.0536, 399.0517 | 319.1444 |
| RebH | WGG-COOH | [M+H$^+$] | 397.0506, 399.0485 | 319.1401 | 397.0521, 399.0505 | 319.1415 |
| No RebH | GW-COOH | [M+H$^+$] | 340.0291, 342.0271 | 262.1186 | 262.1216 | Not found |
| No RebH | WG-COOH | [M+H$^+$] | 340.0291, 342.0271 | 262.1186 | 262.1212 | Not found |
| No RebH | GGW-COOH | [M+H$^+$] | 397.0506, 399.0485 | 319.1401 | 319.1433 | Not found |
| No RebH | WGG-COOH | [M+H$^+$] | 397.0506, 399.0485 | 319.1401 | 319.1435 | Not found |

#### Dipeptides Calculated and Found Mass List

| Enzyme | Dipeptide Substrate | Charge State | Calculated Product | Calculated Reactant | Found Product | Found Reactant |
|---|---|---|---|---|---|---|
| RebH | AW | [M+H⁺] | 353.0535, 355.0535 | 275.1430 | 353.0607, 355.0588 | 275.1514 |
| RebH | CW | [M+H⁺] | 385.0328, 387.0308 | 307.1223 | 385.0317, 387.0318 | 306.1146 |
| RebH | DW | [M+H⁺] | 397.0506, 399.0485 | 319.1401 | 397.0516, 399.0551 | 319.1414 |
| RebH | EW | [M+H⁺] | 411.0662, 413.0642 | 333.1557 | 411.0673, 413.0654 | 333.1573 |
| RebH | FW | [M+H⁺] | 429.0921, 431.0900 | 351.1816 | 429.0934, 431.0917 | 351.1857 |
| RebH | GW | [M+H⁺] | 339.0451, 341.0431 | 261.1346 | 339.0458, 341.0486 | 261.1252 |
| RebH | HW | [M+H⁺] | 419.0826, 421.0805 | 341.1721 | 419.0835, 421.0848 | 341.1738 |
| RebH | IW | [M+H⁺] | 395.1077, 397.1057 | 317.4125 | 395.1107, 397.1088 | 317.2114 |
| RebH | KW | [M+H⁺] | 410.1186, 412.1166 | 332.2081 | 410.1206, 412.1180 | 332.2095 |
| RebH | LW | [M+H⁺] | 395.1077, 397.1057 | 317.1972 | 395.1095, 397.1077 | 317.2016 |
| RebH | MW | [M+H⁺] | 413.0641, 415.0621 | 335.1536 | 411.0523, 413.0505 | 333.1425 |
| RebH | NW | [M+H⁺] | 396.0666, 398.0645 | 318.1561 | 396.0661, 398.0661 | 318.1576 |
| RebH | PW | [M+H⁺] | 379.0764, 381.0744 | 301.1659 | 379.0791, 381.0772 | 301.1975 |
| RebH | QW | [M+H⁺] | 410.0822, 412.0802 | 332.1717 | 410.9339, 412.9128 | 332.7982 |
| RebH | RW | [M+H⁺] | 438.1248, 440.1227 | 360.2142 | 438.1269, 440.1244 | 360.2159 |
| RebH | SW | [M+H⁺] | 369.0557, 371.0536 | 291.1452 | 369.0577, 371.0554 | 291.1471 |
| RebH | TW | [M+H⁺] | 383.0713, 385.0693 | 305.1608 | 383.0728, 385.0711 | 305.1635 |
| RebH | VW | [M+H⁺] | 381.0921, 383.0900 | 303.1816 | 381.0957, 383.0933 | 303.1847 |
| RebH | YW | [M+H⁺] | 445.087, 447.0849 | 367.1765 | 445.0896, 447.0871 | 367.1805 |
| RebH | WA | [M+H⁺] | 353.0535, 355.0535 | 275.1430 | 353.0616, 355.0598 | 275.1542 |
| RebH | WC | [M+H⁺] | 385.0328, 387.0308 | 307.1223 | 384.0291, 386.0259 | 306.1175 |
| RebH | WD | [M+H⁺] | 397.0506, 399.0485 | 319.1401 | 397.0516, 399.0500 | 319.1415 |
| RebH | WE | [M+H⁺] | 411.0662, 413.0642 | 333.1557 | 411.0673, 413.0654 | 333.1577 |
| RebH | WF | [M+H⁺] | 429.0921, 431.0900 | 351.1816 | 429.0933, 431.0917 | 351.1921 |

| Enzyme | Dipeptide Substrate | Charge State | Calculated Product | Calculated Reactant | Found Product | Found Reactant |
|---|---|---|---|---|---|---|
| RebH | WG | [M+H+] | 339.0451, 341.0431 | 261.1346 | 339.0460, 341.0441 | 261.1359 |
| RebH | WH | [M+H+] | 419.0826, 421.0805 | 341.1721 | 419.0840, 421.0821 | 341.1731 |
| RebH | WI | [M+H+] | 395.1077, 397.1057 | 317.4125 | 395.1113, 397.1095 | 317.2063 |
| RebH | WK | [M+H+] | 410.1186, 412.1166 | 332.2081 | 410.1196, 412.1175 | 332.2088 |
| RebH | WL | [M+H+] | 395.1077, 397.1057 | 317.1972 | 395.1087, 397.1070 | 317.1995 |
| RebH | WM | [M+H+] | 413.0641, 415.0621 | 335.1536 | 413.0649, 415.0649 | 335.1554 |
| RebH | WN | [M+H+] | 396.0666, 398.0645 | 318.1561 | 396.0675, 398.0656 | 318.1568 |
| RebH | WP | [M+H+] | 379.0764, 381.0744 | 301.1659 | 379.0778, 381.0759 | 301.1725 |
| RebH | WQ | [M+H+] | 410.0822, 412.0802 | 332.1717 | 410.0835, 412.0825 | 332.1745 |
| RebH | WR | [M+H+] | 438.1248, 440.1227 | 360.2142 | 438.1256, 440.1238 | 360.2158 |
| RebH | WS | [M+H+] | 369.0557, 371.0536 | 291.1452 | 369.0570, 371.0552 | 291.1464 |
| RebH | WT | [M+H+] | 383.0713, 385.0693 | 305.1608 | 383.0727, 385.0707 | 305.1620 |
| RebH | WV | [M+H+] | 381.0921, 383.0900 | 303.1816 | 381.0939, 383.0921 | 303.1938 |
| RebH | WY | [M+H+] | 445.087, 447.0849 | 367.1765 | 445.0882, 447.0873 | 367.1765 |
| 4V | AW | [M+H+] | 353.0535, 355.0535 | 275.1430 | 353.0613, 355.0601 | Not found |
| 4V | CW | [M+H+] | 385.0328, 387.0308 | 307.1223 | 385.0317, 387.0318 | 306.1163 |
| 4V | DW | [M+H+] | 397.0506, 399.0485 | 319.1401 | 397.0520, 399.0501 | 319.1414 |
| 4V | EW | [M+H+] | 411.0662, 413.0642 | 333.1557 | 411.0676, 413.0658 | 333.1570 |
| 4V | FW | [M+H+] | 429.0921, 431.0900 | 351.1816 | 429.0942, 431.0924 | 351.1835 |
| 4V | GW | [M+H+] | 339.0451, 341.0431 | 261.1346 | 339.0466, 341.0449 | 261.1275 |
| 4V | HW | [M+H+] | 419.0826, 421.0805 | 341.1721 | 419.0831, 431.0817 | 341.1731 |
| 4V | IW | [M+H+] | 395.1077, 397.1057 | 317.4125 | 395.1209, 397.1190 | 317.2108 |
| 4V | KW | [M+H+] | 410.1186, 412.1166 | 332.2081 | 410.1206, 412.1188 | 332.2095 |
| 4V | LW | [M+H+] | 395.1077, 397.1057 | 317.1972 | 395.1122, 397.1104 | 317.2026 |
| 4V | MW | [M+H+] | 413.0641, 415.0621 | 335.1536 | 411.0527, 413.0510 | 333.1392 |
| 4V | NW | [M+H+] | 396.0666, 398.0645 | 318.1561 | 396.0681, 398.0663 | 318.1584 |

| Enzyme | Dipeptide Substrate | Charge State | Calculated Product | Calculated Reactant | Found Product | Found Reactant |
|---|---|---|---|---|---|---|
| 4V | PW | [M+H+] | 379.0764, 381.0744 | 301.1659 | 379.0781, 381.0763 | 301.1771 |
| 4V | QW | [M+H+] | 410.0822, 412.0802 | 332.1717 | 410.0518, 412.0518 | 332.1282 |
| 4V | RW | [M+H+] | 438.1248, 440.1227 | 360.2142 | 438.1266, 440.1246 | 360.2142 |
| 4V | SW | [M+H+] | 369.0557, 371.0536 | 291.1452 | 369.0588, 371.0572 | 291.1469 |
| 4V | TW | [M+H+] | 383.0713, 385.0693 | 305.1608 | 383.0735, 385.0715 | 305.163 |
| 4V | VW | [M+H+] | 381.0921, 383.0900 | 303.1816 | 381.0955, 383.0900 | 303.1840 |
| 4V | YW | [M+H+] | 445.087, 447.0849 | 367.1765 | 445.0893, 447.0872 | 367.1789 |
| 4V | WA | [M+H+] | 353.0535, 355.0535 | 275.1430 | 353.0616, 355.0598 | 275.1542 |
| 4V | WC | [M+H+] | 385.0328, 387.0308 | 307.1223 | 384.0291, 386.0259 | 306.0047 |
| 4V | WD | [M+H+] | 397.0506, 399.0485 | 319.1401 | 397.0516, 399.0500 | 319.1415 |
| 4V | WE | [M+H+] | 411.0662, 413.0642 | 333.1557 | 411.0674, 413.0658 | 333.1583 |
| 4V | WF | [M+H+] | 429.0921, 431.0900 | 351.1816 | 429.0938, 431.0922 | 351.1868 |
| 4V | WG | [M+H+] | 339.0451, 341.0431 | 261.1346 | 339.0461, 341.0442 | 261.1349 |
| 4V | WH | [M+H+] | 419.0826, 421.0805 | 341.1721 | 419.0838, 421.0818 | 341.1733 |
| 4V | WI | [M+H+] | 395.1077, 397.1057 | 317.4125 | 395.1119, 397.1101 | 317.2085 |
| 4V | WK | [M+H+] | 410.1186, 412.1166 | 332.2081 | 410.1197, 412.1177 | 332.2088 |
| 4V | WL | [M+H+] | 395.1077, 397.1057 | 317.1972 | 395.1096, 397.1078 | 317.2005 |
| 4V | WM | [M+H+] | 413.0641, 415.0621 | 335.1536 | 413.0649, 415.0635 | 335.1564 |
| 4V | WN | [M+H+] | 396.0666, 398.0645 | 318.1561 | 396.0673, 398.0655 | 318.1568 |
| 4V | WP | [M+H+] | 379.0764, 381.0744 | 301.1659 | 379.0775, 381.0756 | 301.1696 |
| 4V | WQ | [M+H+] | 410.0822, 412.0802 | 332.1717 | 410.0835, 412.0817 | 332.1752 |
| 4V | WR | [M+H+] | 438.1248, 440.1227 | 360.2142 | 438.1262, 440.1244 | 360.2156 |
| 4V | WS | [M+H+] | 369.0557, 371.0536 | 291.1452 | 369.0576, 371.0557 | 291.1461 |
| 4V | WT | [M+H+] | 383.0713, 385.0693 | 305.1608 | 383.0723, 385.0704 | 305.1619 |
| 4V | WV | [M+H+] | 381.0921, 383.0900 | 303.1816 | 381.0939, 383.0921 | 303.1905 |
| 4V | WY | [M+H+] | 445.087, 447.0849 | 367.1765 | 445.0879, 447.0863 | 367.1788 |

| Enzyme | Dipeptide Substrate | Charge State | Calculated Product | Calculated Reactant | Found Product | Found Reactant |
|---|---|---|---|---|---|---|
| no RebH | AW | [M+H+] | 353.0535, 355.0535 | 275.1430 | Not found | 275.1524 |
| no RebH | CW | [M+H+] | 385.0328, 387.0308 | 307.1223 | Not found | 306.1249 |
| no RebH | DW | [M+H+] | 397.0506, 399.0485 | 319.1401 | Not found | 319.1658 |
| no RebH | EW | [M+H+] | 411.0662, 413.0642 | 333.1557 | Not found | 333.1626 |
| no RebH | FW | [M+H+] | 429.0921, 431.0900 | 351.1816 | Not found | 351.2258 |
| no RebH | GW | [M+H+] | 339.0451, 341.0431 | 261.1346 | Not found | 261.1361 |
| no RebH | HW | [M+H+] | 419.0826, 421.0805 | 341.1721 | Not found | 341.1755 |
| no RebH | IW | [M+H+] | 395.1077, 397.1057 | 317.4125 | Not found | 317.2348 |
| no RebH | KW | [M+H+] | 410.1186, 412.1166 | 332.2081 | Not found | 332.2174 |
| no RebH | LW | [M+H+] | 395.1077, 397.1057 | 317.1972 | Not found | 317.2315 |
| no RebH | MW | [M+H+] | 413.0641, 415.0621 | 335.1536 | Not found | 335.1783 |
| no RebH | NW | [M+H+] | 396.0666, 398.0645 | 318.1561 | Not found | 318.1636 |
| no RebH | PW | [M+H+] | 379.0764, 381.0744 | 301.1659 | Not found | 301.1886 |
| no RebH | QW | [M+H+] | 410.0822, 412.0802 | 332.1717 | Not found | 332.1740 |
| no RebH | RW | [M+H+] | 438.1248, 440.1227 | 360.2142 | Not found | 360.2177 |
| no RebH | SW | [M+H+] | 369.0557, 371.0536 | 291.1452 | Not found | 291.1489 |
| no RebH | TW | [M+H+] | 383.0713, 385.0693 | 305.1608 | Not found | 305.1634 |
| no RebH | VW | [M+H+] | 381.0921, 383.0900 | 303.1816 | Not found | 303.1883 |
| no RebH | YW | [M+H+] | 445.087, 447.0849 | 367.1765 | Not found | 367.1817 |
| no RebH | WA | [M+H+] | 353.0535, 355.0535 | 275.1430 | Not found | 275.1644 |
| no RebH | WC | [M+H+] | 385.0328, 387.0308 | 307.1223 | Not found | 306.1192 |
| no RebH | WD | [M+H+] | 397.0506, 399.0485 | 319.1401 | Not found | 319.1574 |
| no RebH | WE | [M+H+] | 411.0662, 413.0642 | 333.1557 | Not found | 333.1640 |
| no RebH | WF | [M+H+] | 429.0921, 431.0900 | 351.1816 | Not found | 351.2145 |
| no RebH | WG | [M+H+] | 339.0451, 341.0431 | 261.1346 | Not found | 261.1382 |
| no RebH | WH | [M+H+] | 419.0826, 421.0805 | 341.1721 | Not found | 341.1949 |

| Enzyme | Dipeptide Substrate | Charge State | Calculated Product | Calculated Reactant | Found Product | Found Reactant |
|---|---|---|---|---|---|---|
| no RebH | WI | [M+H+] | 395.1077, 397.1057 | 317.4125 | Not found | 317.2390 |
| no RebH | WK | [M+H+] | 410.1186, 412.1166 | 332.2081 | Not found | 332.2246 |
| no RebH | WL | [M+H+] | 395.1077, 397.1057 | 317.1972 | Not found | 317.2425 |
| no RebH | WM | [M+H+] | 413.0641, 415.0621 | 335.1536 | Not found | 335.1795 |
| no RebH | WN | [M+H+] | 396.0666, 398.0645 | 318.1561 | Not found | 318.1615 |
| no RebH | WP | [M+H+] | 379.0764, 381.0744 | 301.1659 | Not found | 301.2009 |
| no RebH | WQ | [M+H+] | 410.0822, 412.0802 | 332.1717 | Not found | 332.1890 |
| no RebH | WR | [M+H+] | 438.1248, 440.1227 | 360.2142 | Not found | 360.2190 |
| no RebH | WS | [M+H+] | 369.0557, 371.0536 | 291.1452 | Not found | 291.1550 |
| no RebH | WT | [M+H+] | 383.0713, 385.0693 | 305.1608 | Not found | 305.1687 |
| no RebH | WV | [M+H+] | 381.0921, 383.0900 | 303.1816 | Not found | 303.2172 |
| no RebH | WY | [M+H+] | 445.087, 447.0849 | 367.1765 | Not found | 367.1937 |

#### Tripeptide Calculated and Found Mass List

| Enzyme | Tripeptide Substrate | Charge State | Calculated Product | Calculated Reactant | Found Product | Found Reactant |
|---|---|---|---|---|---|---|
| No RebH | WGA | $[M+H^+]$ | 411.0662, 413.0642 | 333.1557 | Not found | 333.1684 |
| No RebH | WGC | $[M+H^+]$ | 443.0383, 445.0363 | 365.1278 | Not found | 364.1255 |
| No RebH | WGD | $[M+H^+]$ | 455.0561, 457.0540 | 377.1456 | Not found | 377.1495 |
| No RebH | WGE | $[M+H^+]$ | 469.0717, 471.0697 | 391.1612 | Not found | 391.1686 |
| No RebH | WGF | $[M+H^+]$ | 487.0975, 489.0955 | 409.1870 | Not found | 409.1883 |
| No RebH | WGH | $[M+H^+]$ | 477.0880, 479.0860 | 399.1775 | Not found | 399.1788 |
| No RebH | WGK | $[M+H^+]$ | 468.1241, 470.1220 | 390.2136 | Not found | 390.2144 |
| No RebH | WGL | $[M+H^+]$ | 453.1132, 455.1111 | 375.2027 | Not found | 375.2092 |
| No RebH | WGM | $[M+H^+]$ | 471.0696, 473.0676 | 393.1591 | Not found | 393.1611 |
| No RebH | WGN | $[M+H^+]$ | 454.0721, 456.0700 | 376.1615 | Not found | 376.1646 |
| No RebH | WGP | $[M+H^+]$ | 437.0819, 439.0798 | 359.1714 | Not found | 359.1738 |
| No RebH | WGQ | $[M+H^+]$ | 468.0877, 470.0857 | 390.1772 | Not found | 390.1802 |
| No RebH | WGR | $[M+H^+]$ | 496.1302, 498.1282 | 418.2197 | Not found | 418.2226 |
| No RebH | WGT | $[M+H^+]$ | 441.0768, 443.0748 | 363.1663 | Not found | 363.1695 |
| No RebH | WGV | $[M+H^+]$ | 439.0975, 441.0955 | 361.1870 | Not found | 361.2068 |
| No RebH | WGY | $[M+H^+]$ | 503.0925, 505.0904 | 425.1819 | Not found | 425.1858 |
| No RebH | WAG | $[M+H^+]$ | 411.0662, 413.0642 | 333.1557 | Not found | 333.1638 |
| No RebH | WCG | $[M+H^+]$ | 443.0383, 445.0363 | 365.1278 | Not found | 365.1305 |
| No RebH | WDG | $[M+H^+]$ | 455.0561, 457.0540 | 377.1456 | Not found | 377.1494 |
| No RebH | WEG | $[M+H^+]$ | 469.0717, 471.0697 | 391.1612 | Not found | 391.1643 |
| No RebH | WFG | $[M+H^+]$ | 487.0975, 489.0955 | 409.1870 | Not found | 409.1870 |
| No RebH | WHG | $[M+H^+]$ | 477.0880, 479.0860 | 399.1775 | Not found | 399.1771 |
| No RebH | WKG | $[M+H^+]$ | 468.1241, 470.1220 | 390.2136 | Not found | 390.2131 |
| No RebH | WLG | $[M+H^+]$ | 453.1132, 455.1111 | 375.2027 | Not found | 375.2026 |

| Enzyme | Tripeptide Substrate | Charge State | Calculated Product | Calculated Reactant | Found Product | Found Reactant |
|---|---|---|---|---|---|---|
| No RebH | WMG | [M+H+] | 471.0696, 473.0676 | 393.1591 | Not found | 393.1599 |
| No RebH | WNG | [M+H+] | 454.0721, 456.0700 | 376.1615 | Not found | 376.1612 |
| No RebH | WPG | [M+H+] | 437.0819, 439.0798 | 359.1714 | Not found | 359.1714 |
| No RebH | WQG | [M+H+] | 468.0877, 470.0857 | 390.1772 | Not found | 390.1770 |
| No RebH | WRG | [M+H+] | 496.1302, 498.1282 | 418.2197 | Not found | 418.2196 |
| No RebH | WTG | [M+H+] | 441.0768, 443.0748 | 363.1663 | Not found | 363.1663 |
| No RebH | WVG | [M+H+] | 439.0975, 441.0955 | 361.1870 | Not found | 361.1925 |
| No RebH | WYG | [M+H+] | 503.0925, 505.0904 | 425.1819 | Not found | 425.1843 |
| No RebH | GWA | [M+H+] | 411.0662, 413.0642 | 333.1557 | Not found | 333.1577 |
| No RebH | GWC | [M+H+] | 443.0383, 445.0363 | 365.1278 | Not found | 364.1216 |
| No RebH | GWD | [M+H+] | 455.0561, 457.0540 | 377.1456 | Not found | 377.1464 |
| No RebH | GWE | [M+H+] | 469.0717, 471.0697 | 391.1612 | Not found | 391.1641 |
| No RebH | GWF | [M+H+] | 487.0975, 489.0955 | 409.1870 | Not found | 409.1913 |
| No RebH | GWH | [M+H+] | 477.0880, 479.0860 | 399.1775 | Not found | 399.1773 |
| No RebH | GWK | [M+H+] | 468.1241, 470.1220 | 390.2136 | Not found | 390.2133 |
| No RebH | GWL | [M+H+] | 453.1132, 455.1111 | 375.2027 | Not found | 375.2022 |
| No RebH | GWM | [M+H+] | 471.0696, 473.0676 | 393.1591 | Not found | 393.1590 |
| No RebH | GWN | [M+H+] | 454.0721, 456.0700 | 376.1615 | Not found | 376.1611 |
| No RebH | GWP | [M+H+] | 437.0819, 439.0798 | 359.1714 | Not found | 359.1833 |
| No RebH | GWQ | [M+H+] | 468.0877, 470.0857 | 390.1772 | Not found | 390.1784 |
| No RebH | GWR | [M+H+] | 496.1302, 498.1282 | 418.2197 | Not found | 418.2196 |
| No RebH | GWT | [M+H+] | 441.0768, 443.0748 | 363.1663 | Not found | 363.1727 |
| No RebH | GWV | [M+H+] | 439.0975, 441.0955 | 361.1870 | Not found | 361.1865 |
| No RebH | GWY | [M+H+] | 503.0925, 505.0904 | 425.1819 | Not found | 425.1848 |
| No RebH | AWG | [M+H+] | 411.0662, 413.0642 | 333.1557 | Not found | 333.1567 |
| No RebH | CWG | [M+H+] | 443.0383, 445.0363 | 365.1278 | Not found | 364.1235 |

| Enzyme | Tripeptide Substrate | Charge State | Calculated Product | Calculated Reactant | Found Product | Found Reactant |
|---|---|---|---|---|---|---|
| No RebH | DWG | [M+H$^+$] | 455.0561, 457.0540 | 377.1456 | Not found | 377.1479 |
| No RebH | EWG | [M+H$^+$] | 469.0717, 471.0697 | 391.1612 | Not found | 391.1636 |
| No RebH | FWG | [M+H$^+$] | 487.0975, 489.0955 | 409.1870 | Not found | 409.1922 |
| No RebH | HWG | [M+H$^+$] | 477.0880, 479.0860 | 399.1775 | Not found | 399.1786 |
| No RebH | KWG | [M+H$^+$] | 468.1241, 470.1220 | 390.2136 | Not found | 390.2138 |
| No RebH | LWG | [M+H$^+$] | 453.1132, 455.1111 | 375.2027 | Not found | 375.2049 |
| No RebH | MWG | [M+H$^+$] | 471.0696, 473.0676 | 393.1591 | Not found | 393.1603 |
| No RebH | NWG | [M+H$^+$] | 454.0721, 456.0700 | 376.1615 | Not found | 376.1643 |
| No RebH | PWG | [M+H$^+$] | 437.0819, 439.0798 | 359.1714 | Not found | 359.1723 |
| No RebH | QWG | [M+H$^+$] | 468.0877, 470.0857 | 390.1772 | Not found | 390.1794 |
| No RebH | RWG | [M+H$^+$] | 496.1302, 498.1282 | 418.2197 | Not found | 418.2204 |
| No RebH | TWG | [M+H$^+$] | 441.0768, 443.0748 | 363.1663 | Not found | 363.1695 |
| No RebH | VWG | [M+H$^+$] | 439.0975, 441.0955 | 361.1870 | Not found | 361.1963 |
| No RebH | YWG | [M+H$^+$] | 503.0925, 505.0904 | 425.1819 | Not found | 425.1846 |
| No RebH | GAW | [M+H$^+$] | 411.0662, 413.0642 | 333.1557 | Not found | 333.1603 |
| No RebH | GCW | [M+H$^+$] | 443.0383, 445.0363 | 365.1278 | Not found | 364.1278 |
| No RebH | GDW | [M+H$^+$] | 455.0561, 457.0540 | 377.1456 | Not found | 377.1471 |
| No RebH | GEW | [M+H$^+$] | 469.0717, 471.0697 | 391.1612 | Not found | 391.1706 |
| No RebH | GFW | [M+H$^+$] | 487.0975, 489.0955 | 409.1870 | Not found | 409.1860 |
| No RebH | GHW | [M+H$^+$] | 477.0880, 479.0860 | 399.1775 | Not found | 399.1786 |
| No RebH | GKW | [M+H$^+$] | 468.1241, 470.1220 | 390.2136 | Not found | 390.2146 |
| No RebH | GLW | [M+H$^+$] | 453.1132, 455.1111 | 375.2027 | Not found | 375.2108 |
| No RebH | GMW | [M+H$^+$] | 471.0696, 473.0676 | 393.1591 | Not found | 393.1617 |
| No RebH | GNW | [M+H$^+$] | 454.0721, 456.0700 | 376.1615 | Not found | 376.1700 |
| No RebH | GPW | [M+H$^+$] | 437.0819, 439.0798 | 359.1714 | Not found | 359.1879 |
| No RebH | GQW | [M+H$^+$] | 468.0877, 470.0857 | 390.1772 | Not found | 390.1789 |

| Enzyme | Tripeptide Substrate | Charge State | Calculated Product | Calculated Reactant | Found Product | Found Reactant |
|---|---|---|---|---|---|---|
| No RebH | GRW | [M+H+] | 496.1302, 498.1282 | 418.2197 | Not found | 418.2209 |
| No RebH | GTW | [M+H+] | 441.0768, 443.0748 | 363.1663 | Not found | 363.1677 |
| No RebH | GVW | [M+H+] | 439.0975, 441.0955 | 361.1870 | Not found | 361.1925 |
| No RebH | GYW | [M+H+] | 503.0925, 505.0904 | 425.1819 | Not found | 425.1826 |
| No RebH | AGW | [M+H+] | 411.0662, 413.0642 | 333.1557 | Not found | 333.1620 |
| No RebH | CGW | [M+H+] | 443.0383, 445.0363 | 365.1278 | Not found | 364.1204 |
| No RebH | DGW | [M+H+] | 455.0561, 457.0540 | 377.1456 | Not found | 377.1516 |
| No RebH | EGW | [M+H+] | 469.0717, 471.0697 | 391.1612 | Not found | 391.1659 |
| No RebH | FGW | [M+H+] | 487.0975, 489.0955 | 409.1870 | Not found | 409.1922 |
| No RebH | HGW | [M+H+] | 477.0880, 479.0860 | 399.1775 | Not found | 399.1786 |
| No RebH | KGW | [M+H+] | 468.1241, 470.1220 | 390.2136 | Not found | 390.2136 |
| No RebH | LGW | [M+H+] | 453.1132, 455.1111 | 375.2027 | Not found | 375.2186 |
| No RebH | MGW | [M+H+] | 471.0696, 473.0676 | 393.1591 | Not found | 393.1589 |
| No RebH | NGW | [M+H+] | 454.0721, 456.0700 | 376.1615 | Not found | 376.1636 |
| No RebH | PGW | [M+H+] | 437.0819, 439.0798 | 359.1714 | Not found | 359.1791 |
| No RebH | QGW | [M+H+] | 468.0877, 470.0857 | 390.1772 | Not found | 390.1789 |
| No RebH | RGW | [M+H+] | 496.1302, 498.1282 | 418.2197 | Not found | 418.2211 |
| No RebH | TGW | [M+H+] | 441.0768, 443.0748 | 363.1663 | Not found | 363.1763 |
| No RebH | VGW | [M+H+] | 439.0975, 441.0955 | 361.1870 | Not found | 361.1988 |
| No RebH | YGW | [M+H+] | 503.0925, 505.0904 | 425.1819 | Not found | 425.1909 |
| RebH | WGA | [M+H+] | 411.0662, 413.0642 | 333.1557 | 411.0677, 413.0654 | 333.1628 |
| RebH | WGC | [M+H+] | 443.0383, 445.0363 | 365.1278 | 442.0329, 444.0284 | 364.1219 |
| RebH | WGD | [M+H+] | 455.0561, 457.0540 | 377.1456 | 455.0728, 457.0716 | 377.1494 |
| RebH | WGE | [M+H+] | 469.0717, 471.0697 | 391.1612 | 469.0732, 471.0719 | 391.1689 |
| RebH | WGF | [M+H+] | 487.0975, 489.0955 | 409.1870 | 487.0976, 489.0967 | 409.1883 |
| RebH | WGH | [M+H+] | 477.0880, 479.0860 | 399.1775 | 477.0877, 479.0863 | 399.1796 |

| Enzyme | Tripeptide Substrate | Charge State | Calculated Product | Calculated Reactant | Found Product | Found Reactant |
|---|---|---|---|---|---|---|
| RebH | WGK | [M+H+] | 468.1241, 470.1220 | 390.2136 | 468.1244, 470.1231 | 390.2141 |
| RebH | WGL | [M+H+] | 453.1132, 455.1111 | 375.2027 | 453.1147, 455.1137 | 375.2063 |
| RebH | WGM | [M+H+] | 471.0696, 473.0676 | 393.1591 | 471.0726, 473.0708 | 393.1610 |
| RebH | WGN | [M+H+] | 454.0721, 456.0700 | 376.1615 | 454.0736, 456.0720 | 376.1636 |
| RebH | WGP | [M+H+] | 437.0819, 439.0798 | 359.1714 | Not found | 359.1749 |
| RebH | WGQ | [M+H+] | 468.0877, 470.0857 | 390.1772 | 468.0919, 470.0893 | 390.1804 |
| RebH | WGR | [M+H+] | 496.1302, 498.1282 | 418.2197 | 496.1337, 498.1310 | 418.2210 |
| RebH | WGT | [M+H+] | 441.0768, 443.0748 | 363.1663 | 441.0990, 443.1414 | 363.1692 |
| RebH | WGV | [M+H+] | 439.0975, 441.0955 | 361.1870 | 439.1079, 441.1145 | 361.2066 |
| RebH | WGY | [M+H+] | 503.0925, 505.0904 | 425.1819 | 503.0995, 505.0970 | 425.1851 |
| RebH | WAG | [M+H+] | 411.0662, 413.0642 | 333.1557 | 411.0704, 413.0685 | 333.1599 |
| RebH | WCG | [M+H+] | 443.0383, 445.0363 | 365.1278 | Not found | 364.1251 |
| RebH | WDG | [M+H+] | 455.0561, 457.0540 | 377.1456 | 455.0595, 457.0873 | 377.1488 |
| RebH | WEG | [M+H+] | 469.0717, 471.0697 | 391.1612 | 469.0751, 471.0731 | 391.1678 |
| RebH | WFG | [M+H+] | 487.0975, 489.0955 | 409.1870 | Not found | 409.1868 |
| RebH | WHG | [M+H+] | 477.0880, 479.0860 | 399.1775 | Not found | 399.1773 |
| RebH | WKG | [M+H+] | 468.1241, 470.1220 | 390.2136 | 468.1232, 470.1219 | 390.2134 |
| RebH | WLG | [M+H+] | 453.1132, 455.1111 | 375.2027 | 453.1132, 455.1135 | 375.2024 |
| RebH | WMG | [M+H+] | 471.0696, 473.0676 | 393.1591 | Not found | 393.1594 |
| RebH | WNG | [M+H+] | 454.0721, 456.0700 | 376.1615 | 454.0718, 456.0700 | 376.1614 |
| RebH | WPG | [M+H+] | 437.0819, 439.0798 | 359.1714 | Not found | 359.1714 |
| RebH | WQG | [M+H+] | 468.0877, 470.0857 | 390.1772 | 468.0878, 470.0861 | 390.1770 |
| RebH | WRG | [M+H+] | 496.1302, 498.1282 | 418.2197 | 496.1295, 498.1278 | 418.2194 |
| RebH | WTG | [M+H+] | 441.0768, 443.0748 | 363.1663 | 441.0772, 443.0753 | 363.1665 |
| RebH | WVG | [M+H+] | 439.0975, 441.0955 | 361.1870 | 439.0984, 441.0959 | 361.1938 |
| RebH | WYG | [M+H+] | 503.0925, 505.0904 | 425.1819 | 503.0923, 505.0912 | 425.1832 |

| Enzyme | Tripeptide Substrate | Charge State | Calculated Product | Calculated Reactant | Found Product | Found Reactant |
|---|---|---|---|---|---|---|
| RebH | GWA | [M+H$^+$] | 411.0662, 413.0642 | 333.1557 | 411.0671, 413.0652 | 333.1563 |
| RebH | GWC | [M+H$^+$] | 443.0383, 445.0363 | 365.1278 | 442.0307, 444.0291 | 364.1227 |
| RebH | GWD | [M+H$^+$] | 455.0561, 457.0540 | 377.1456 | 455.0558, 457.0545 | 377.1466 |
| RebH | GWE | [M+H$^+$] | 469.0717, 471.0697 | 391.1612 | 469.0726, 471.0702 | 391.1616 |
| RebH | GWF | [M+H$^+$] | 487.0975, 489.0955 | 409.1870 | 487.0980, 489.0967 | 409.1900 |
| RebH | GWH | [M+H$^+$] | 477.0880, 479.0860 | 399.1775 | 477.0878, 479.0860 | 399.1772 |
| RebH | GWK | [M+H$^+$] | 468.1241, 470.1220 | 390.2136 | 468.1243, 470.1220 | 390.2140 |
| RebH | GWL | [M+H$^+$] | 453.1132, 455.1111 | 375.2027 | 453.1129, 455.1194 | 375.2024 |
| RebH | GWM | [M+H$^+$] | 471.0696, 473.0676 | 393.1591 | 471.0697, 473.0709 | 393.1591 |
| RebH | GWN | [M+H$^+$] | 454.0721, 456.0700 | 376.1615 | 454.0718, 456.0703 | 376.1610 |
| RebH | GWP | [M+H$^+$] | 437.0819, 439.0798 | 359.1714 | Not found | 359.1788 |
| RebH | GWQ | [M+H$^+$] | 468.0877, 470.0857 | 390.1772 | 468.0879, 470.0861 | 390.1775 |
| RebH | GWR | [M+H$^+$] | 496.1302, 498.1282 | 418.2197 | 496.1305, 498.1287 | 418.2193 |
| RebH | GWT | [M+H$^+$] | 441.0768, 443.0748 | 363.1663 | 441.0770, 443.0751 | 363.1695 |
| RebH | GWV | [M+H$^+$] | 439.0975, 441.0955 | 361.1870 | Not found | 361.1869 |
| RebH | GWY | [M+H$^+$] | 503.0925, 505.0904 | 425.1819 | Not found | 425.1852 |
| RebH | AWG | [M+H$^+$] | 411.0662, 413.0642 | 333.1557 | 411.0659, 413.0641 | 333.1560 |
| RebH | CWG | [M+H$^+$] | 443.0383, 445.0363 | 365.1278 | Not found | 364.1233 |
| RebH | DWG | [M+H$^+$] | 455.0561, 457.0540 | 377.1456 | Not found | 377.1476 |
| RebH | EWG | [M+H$^+$] | 469.0717, 471.0697 | 391.1612 | Not found | 391.1640 |
| RebH | FWG | [M+H$^+$] | 487.0975, 489.0955 | 409.1870 | 487.0981, 489.0991 | 409.1923 |
| RebH | HWG | [M+H$^+$] | 477.0880, 479.0860 | 399.1775 | 477.0888, 479.0876 | 399.1787 |
| RebH | KWG | [M+H$^+$] | 468.1241, 470.1220 | 390.2136 | 468.1251, 470.1229 | 390.2140 |
| RebH | LWG | [M+H$^+$] | 453.1132, 455.1111 | 375.2027 | 453.1153, 455.1174 | 375.2053 |
| RebH | MWG | [M+H$^+$] | 471.0696, 473.0676 | 393.1591 | 471.0669, 473.0712 | 393.1606 |
| RebH | NWG | [M+H$^+$] | 454.0721, 456.0700 | 376.1615 | 454.0727, 456.0722 | 376.1624 |

| Enzyme | Tripeptide Substrate | Charge State | Calculated Product | Calculated Reactant | Found Product | Found Reactant |
|---|---|---|---|---|---|---|
| RebH | PWG | [M+H+] | 437.0819, 439.0798 | 359.1714 | 437.0835, 439.0842 | 359.1772 |
| RebH | QWG | [M+H+] | 468.0877, 470.0857 | 390.1772 | 468.0891, 470.0874 | 390.1800 |
| RebH | RWG | [M+H+] | 496.1302, 498.1282 | 418.2197 | 496.1313, 498.1290 | 418.2200 |
| RebH | TWG | [M+H+] | 441.0768, 443.0748 | 363.1663 | 441.0783, 443.775 | 363.1695 |
| RebH | VWG | [M+H+] | 439.0975, 441.0955 | 361.1870 | Not found | 361.1954 |
| RebH | YWG | [M+H+] | 503.0925, 505.0904 | 425.1819 | 503.0934, 505.0935 | 425.1835 |
| RebH | GAW | [M+H+] | 411.0662, 413.0642 | 333.1557 | 411.0701, 413.0708 | 333.1582 |
| RebH | GCW | [M+H+] | 443.0383, 445.0363 | 365.1278 | Not found | 364.1233 |
| RebH | GDW | [M+H+] | 455.0561, 457.0540 | 377.1456 | Not found | 377.1489 |
| RebH | GEW | [M+H+] | 469.0717, 471.0697 | 391.1612 | Not found | 391.1651 |
| RebH | GFW | [M+H+] | 487.0975, 489.0955 | 409.1870 | Not found | 409.1890 |
| RebH | GHW | [M+H+] | 477.0880, 479.0860 | 399.1775 | Not found | 399.1780 |
| RebH | GKW | [M+H+] | 468.1241, 470.1220 | 390.2136 | Not found | 390.2146 |
| RebH | GLW | [M+H+] | 453.1132, 455.1111 | 375.2027 | Not found | 375.2101 |
| RebH | GMW | [M+H+] | 471.0696, 473.0676 | 393.1591 | Not found | 393.1637 |
| RebH | GNW | [M+H+] | 454.0721, 456.0700 | 376.1615 | Not found | 376.1661 |
| RebH | GPW | [M+H+] | 437.0819, 439.0798 | 359.1714 | Not found | 359.1936 |
| RebH | GQW | [M+H+] | 468.0877, 470.0857 | 390.1772 | Not found | 390.1781 |
| RebH | GRW | [M+H+] | 496.1302, 498.1282 | 418.2197 | Not found | 418.2241 |
| RebH | GTW | [M+H+] | 441.0768, 443.0748 | 363.1663 | Not found | 363.1676 |
| RebH | GVW | [M+H+] | 439.0975, 441.0955 | 361.1870 | Not found | 361.1972 |
| RebH | GYW | [M+H+] | 503.0925, 505.0904 | 425.1819 | Not found | 425.1822 |
| RebH | AGW | [M+H+] | 411.0662, 413.0642 | 333.1557 | 411.0662, 413.0645 | 333.1629 |
| RebH | CGW | [M+H+] | 443.0383, 445.0363 | 365.1278 | Not found | 364.1204 |
| RebH | DGW | [M+H+] | 455.0561, 457.0540 | 377.1456 | Not found | 377.1513 |
| RebH | EGW | [M+H+] | 469.0717, 471.0697 | 391.1612 | Not found | 391.1662 |

| Enzyme | Tripeptide Substrate | Charge State | Calculated Product | Calculated Reactant | Found Product | Found Reactant |
|---|---|---|---|---|---|---|
| RebH | FGW | $[M+H^+]$ | 487.0975, 489.0955 | 409.1870 | 487.0974, 489.0971 | 409.1952 |
| RebH | HGW | $[M+H^+]$ | 477.0880, 479.0860 | 399.1775 | 477.0885, 479.0869 | 399.1790 |
| RebH | KGW | $[M+H^+]$ | 468.1241, 470.1220 | 390.2136 | 468.2494, 470.1296 | 390.2137 |
| RebH | LGW | $[M+H^+]$ | 453.1132, 455.1111 | 375.2027 | 453.1144, 455.1125 | 375.2190 |
| RebH | MGW | $[M+H^+]$ | 471.0696, 473.0676 | 393.1591 | 469.0537, 471.0530 | 393.1599 |
| RebH | NGW | $[M+H^+]$ | 454.0721, 456.0700 | 376.1615 | 454.0722, 456.0719 | 376.1646 |
| RebH | PGW | $[M+H^+]$ | 437.0819, 439.0798 | 359.1714 | 437.0845, 439.0923 | 359.1784 |
| RebH | QGW | $[M+H^+]$ | 468.0877, 470.0857 | 390.1772 | 468.0886, 470.0876 | 390.1786 |
| RebH | RGW | $[M+H^+]$ | 496.1302, 498.1282 | 418.2197 | 496.1307, 498.1286 | 418.2211 |
| RebH | TGW | $[M+H^+]$ | 441.0768, 443.0748 | 363.1663 | 441.0787, 443.0769 | 363.1803 |
| RebH | VGW | $[M+H^+]$ | 439.0975, 441.0955 | 361.1870 | 439.0998, 441.0980 | 361.2013 |
| RebH | YGW | $[M+H^+]$ | 503.0925, 505.0904 | 425.1819 | 503.0936, 505.0922 | 425.1887 |
| 4V | WGA | $[M+H^+]$ | 411.0662, 413.0642 | 333.1557 | 411.0680, 413.0661 | 333.1615 |
| 4V | WGC | $[M+H^+]$ | 443.0383, 445.0363 | 365.1278 | 442.0323, 444.0324 | 364.1216 |
| 4V | WGD | $[M+H^+]$ | 455.0561, 457.0540 | 377.1456 | 455.0558, 457.0562 | 377.1471 |
| 4V | WGE | $[M+H^+]$ | 469.0717, 471.0697 | 391.1612 | 469.0737, 471.0723 | 391.1708 |
| 4V | WGF | $[M+H^+]$ | 487.0975, 489.0955 | 409.1870 | 487.0974, 489.0960 | 409.1882 |
| 4V | WGH | $[M+H^+]$ | 477.0880, 479.0860 | 399.1775 | 477.0887, 479.0875 | 399.1790 |
| 4V | WGK | $[M+H^+]$ | 468.1241, 470.1220 | 390.2136 | 468.1246, 470.1235 | 390.2150 |
| 4V | WGL | $[M+H^+]$ | 453.1132, 455.1111 | 375.2027 | 453.1141, 455.1123 | 375.2055 |
| 4V | WGM | $[M+H^+]$ | 471.0696, 473.0676 | 393.1591 | 471.0741, 473.0704 | 393.1611 |
| 4V | WGN | $[M+H^+]$ | 454.0721, 456.0700 | 376.1615 | 454.0723, 456.0707 | 376.1634 |
| 4V | WGP | $[M+H^+]$ | 437.0819, 439.0798 | 359.1714 | Not found | 359.1724 |
| 4V | WGQ | $[M+H^+]$ | 468.0877, 470.0857 | 390.1772 | 468.0793, 470.0782 | 390.1794 |
| 4V | WGR | $[M+H^+]$ | 496.1302, 498.1282 | 418.2197 | 496.1405, 498.1198 | 418.2196 |
| 4V | WGT | $[M+H^+]$ | 441.0768, 443.0748 | 363.1663 | 436.7723, 438.7520 | 359.6270 |

| Enzyme | Tripeptide Substrate | Charge State | Calculated Product | Calculated Reactant | Found Product | Found Reactant |
|---|---|---|---|---|---|---|
| 4V | WGV | [M+H+] | 439.0975, 441.0955 | 361.1870 | 434.8193, 436.7983 | 357.6727 |
| 4V | WGY | [M+H+] | 503.0925, 505.0904 | 425.1819 | 498.1668, 500.1463 | 421.0393 |
| 4V | WAG | [M+H+] | 411.0662, 413.0642 | 333.1557 | 407.0605, 409.0393 | 333.1580 |
| 4V | WCG | [M+H+] | 443.0383, 445.0363 | 365.1278 | 442.0369, 444.0278 | 364.1187 |
| 4V | WDG | [M+H+] | 455.0561, 457.0540 | 377.1456 | 455.0595, 457.0585 | 377.1378 |
| 4V | WEG | [M+H+] | 469.0717, 471.0697 | 391.1612 | 469.0694, 471.0729 | 391.1679 |
| 4V | WFG | [M+H+] | 487.0975, 489.0955 | 409.1870 | 487.0973, 489.0959 | 409.1868 |
| 4V | WHG | [M+H+] | 477.0880, 479.0860 | 399.1775 | 477.0872, 479.0863 | 399.1772 |
| 4V | WKG | [M+H+] | 468.1241, 470.1220 | 390.2136 | 468.1238, 470.1215 | 390.2137 |
| 4V | WLG | [M+H+] | 453.1132, 455.1111 | 375.2027 | 453.1128, 455.1115 | 375.2024 |
| 4V | WMG | [M+H+] | 471.0696, 473.0676 | 393.1591 | 471.1651, 473.0687 | 393.1596 |
| 4V | WNG | [M+H+] | 454.0721, 456.0700 | 376.1615 | 454.0717, 456.0699 | 376.1614 |
| 4V | WPG | [M+H+] | 437.0819, 439.0798 | 359.1714 | 437.0822, 439.0810 | 359.1712 |
| 4V | WQG | [M+H+] | 468.0877, 470.0857 | 390.1772 | 468.0879, 470.0864 | 390.1767 |
| 4V | WRG | [M+H+] | 496.1302, 498.1282 | 418.2197 | 496.1295, 498.1283 | 418.2191 |
| 4V | WTG | [M+H+] | 441.0768, 443.0748 | 363.1663 | 441.0764, 443.0745 | 363.1662 |
| 4V | WVG | [M+H+] | 439.0975, 441.0955 | 361.1870 | 439.0970, 441.0953 | 361.1916 |
| 4V | WYG | [M+H+] | 503.0925, 505.0904 | 425.1819 | 503.0920, 505.0905 | 425.1840 |
| 4V | GWA | [M+H+] | 411.0662, 413.0642 | 333.1557 | 411.0674, 413.0656 | 333.1556 |
| 4V | GWC | [M+H+] | 443.0383, 445.0363 | 365.1278 | 442.0320, 444.0317 | 364.1211 |
| 4V | GWD | [M+H+] | 455.0561, 457.0540 | 377.1456 | 455.0558, 457.0560 | 377.1464 |
| 4V | GWE | [M+H+] | 469.0717, 471.0697 | 391.1612 | 469.0715, 471.0697 | 391.1624 |
| 4V | GWF | [M+H+] | 487.0975, 489.0955 | 409.1870 | 487.0972, 489.0955 | 490.1895 |
| 4V | GWH | [M+H+] | 477.0880, 479.0860 | 399.1775 | 477.0879, 479.0860 | 399.1773 |
| 4V | GWK | [M+H+] | 468.1241, 470.1220 | 390.2136 | 468.1237, 470.1221 | 390.2130 |
| 4V | GWL | [M+H+] | 453.1132, 455.1111 | 375.2027 | 453.1139, 455.1127 | 375.2032 |

| Enzyme | Tripeptide Substrate | Charge State | Calculated Product | Calculated Reactant | Found Product | Found Reactant |
|---|---|---|---|---|---|---|
| 4V | GWM | [M+H$^+$] | 471.0696, 473.0676 | 393.1591 | 471.0694, 743.0675 | 393.1598 |
| 4V | GWN | [M+H$^+$] | 454.0721, 456.0700 | 376.1615 | 454.0717, 456.0702 | 376.1610 |
| 4V | GWP | [M+H$^+$] | 437.0819, 439.0798 | 359.1714 | 437.0821, 439.0901 | 359.1843 |
| 4V | GWQ | [M+H$^+$] | 468.0877, 470.0857 | 390.1772 | 468.0875, 470.0857 | 390.1777 |
| 4V | GWR | [M+H$^+$] | 496.1302, 498.1282 | 418.2197 | 496.1306, 498.1288 | 418.2192 |
| 4V | GWT | [M+H$^+$] | 441.0768, 443.0748 | 363.1663 | 441.0765, 443.0747 | 363.1713 |
| 4V | GWV | [M+H$^+$] | 439.0975, 441.0955 | 361.1870 | 439.0982, 441.0961 | 361.1866 |
| 4V | GWY | [M+H$^+$] | 503.0925, 505.0904 | 425.1819 | 503.0935, 505.0917 | 425.1853 |
| 4V | AWG | [M+H$^+$] | 411.0662, 413.0642 | 333.1557 | 411.0660, 413.0645 | 333.1562 |
| 4V | CWG | [M+H$^+$] | 443.0383, 445.0363 | 365.1278 | Not found | 364.1208 |
| 4V | DWG | [M+H$^+$] | 455.0561, 457.0540 | 377.1456 | Not found | 377.1471 |
| 4V | EWG | [M+H$^+$] | 469.0717, 471.0697 | 391.1612 | Not found | 391.1640 |
| 4V | FWG | [M+H$^+$] | 487.0975, 489.0955 | 409.1870 | 487.0979, 489.0993 | 409.1946 |
| 4V | HWG | [M+H$^+$] | 477.0880, 479.0860 | 399.1775 | 477.0890, 479.0873 | 399.1786 |
| 4V | KWG | [M+H$^+$] | 468.1241, 470.1220 | 390.2136 | 468.1254, 470.1234 | 390.2140 |
| 4V | LWG | [M+H$^+$] | 453.1132, 455.1111 | 375.2027 | 453.1141, 455.1143 | 375.2056 |
| 4V | MWG | [M+H$^+$] | 471.0696, 473.0676 | 393.1591 | 471.0688, 473.0687 | 393.1622 |
| 4V | NWG | [M+H$^+$] | 454.0721, 456.0700 | 376.1615 | 454.0727, 456.0733 | 376.1645 |
| 4V | PWG | [M+H$^+$] | 437.0819, 439.0798 | 359.1714 | 437.0822, 439.0812 | 359.1757 |
| 4V | QWG | [M+H$^+$] | 468.0877, 470.0857 | 390.1772 | 468.0883, 470.0878 | 390.1810 |
| 4V | RWG | [M+H$^+$] | 496.1302, 498.1282 | 418.2197 | 496.1301, 498.1284 | 418.2200 |
| 4V | TWG | [M+H$^+$] | 441.0768, 443.0748 | 363.1663 | 441.0782, 443.0762 | 363.1693 |
| 4V | VWG | [M+H$^+$] | 439.0975, 441.0955 | 361.1870 | Not found | 361.2049 |
| 4V | YWG | [M+H$^+$] | 503.0925, 505.0904 | 425.1819 | 503.0928, 505.0911 | 425.1837 |
| 4V | GAW | [M+H$^+$] | 411.0662, 413.0642 | 333.1557 | 411.0719, 413.0633 | 333.1584 |
| 4V | GCW | [M+H$^+$] | 443.0383, 445.0363 | 365.1278 | Not found | 364.1227 |

| Enzyme | Tripeptide Substrate | Charge State | Calculated Product | Calculated Reactant | Found Product | Found Reactant |
|---|---|---|---|---|---|---|
| 4V | GDW | [M+H+] | 455.0561, 457.0540 | 377.1456 | Not found | 377.1495 |
| 4V | GEW | [M+H+] | 469.0717, 471.0697 | 391.1612 | 469.0735, 471.0712 | 391.1664 |
| 4V | GFW | [M+H+] | 487.0975, 489.0955 | 409.1870 | Not found | 409.1895 |
| 4V | GHW | [M+H+] | 477.0880, 479.0860 | 399.1775 | 477.0893, 479.0885 | 399.1782 |
| 4V | GKW | [M+H+] | 468.1241, 470.1220 | 390.2136 | 468.1269, 470.1241 | 390.2145 |
| 4V | GLW | [M+H+] | 453.1132, 455.1111 | 375.2027 | Not found | 375.2094 |
| 4V | GMW | [M+H+] | 471.0696, 473.0676 | 393.1591 | Not found | 393.1624 |
| 4V | GNW | [M+H+] | 454.0721, 456.0700 | 376.1615 | Not found | 376.1642 |
| 4V | GPW | [M+H+] | 437.0819, 439.0798 | 359.1714 | 437.0825, 439.0888 | 359.1920 |
| 4V | GQW | [M+H+] | 468.0877, 470.0857 | 390.1772 | Not found | 390.1788 |
| 4V | GRW | [M+H+] | 496.1302, 498.1282 | 418.2197 | Not found | 418.2228 |
| 4V | GTW | [M+H+] | 441.0768, 443.0748 | 363.1663 | Not found | 363.1680 |
| 4V | GVW | [M+H+] | 439.0975, 441.0955 | 361.1870 | 439.2318, 441.2380 | 361.1960 |
| 4V | GYW | [M+H+] | 503.0925, 505.0904 | 425.1819 | Not found | 425.1817 |
| 4V | AGW | [M+H+] | 411.0662, 413.0642 | 333.1557 | 411.0657, 413.0641 | 333.1607 |
| 4V | CGW | [M+H+] | 443.0383, 445.0363 | 365.1278 | Not found | 364.1200 |
| 4V | DGW | [M+H+] | 455.0561, 457.0540 | 377.1456 | 455.0561, 457.0548 | 377.1488 |
| 4V | EGW | [M+H+] | 469.0717, 471.0697 | 391.1612 | Not found | 391.1638 |
| 4V | FGW | [M+H+] | 487.0975, 489.0955 | 409.1870 | 487.0969, 489.0953 | 409.1938 |
| 4V | HGW | [M+H+] | 477.0880, 479.0860 | 399.1775 | 477.0895, 479.0882 | 399.1784 |
| 4V | KGW | [M+H+] | 468.1241, 470.1220 | 390.2136 | 468.1239, 470.1219 | 390.2133 |
| 4V | LGW | [M+H+] | 453.1132, 455.1111 | 375.2027 | 453.1138, 455.1123 | 375.2144 |
| 4V | MGW | [M+H+] | 471.0696, 473.0676 | 393.1591 | 469.0542, 471.0527 | 391.1434 |
| 4V | NGW | [M+H+] | 454.0721, 456.0700 | 376.1615 | 454.0726, 456.0743 | 376.1664 |
| 4V | PGW | [M+H+] | 437.0819, 439.0798 | 359.1714 | 437.0833, 439.0848 | 359.1807 |
| 4V | QGW | [M+H+] | 468.0877, 470.0857 | 390.1772 | 468.0889, 470.0874 | 390.1782 |

| Enzyme | Tripeptide Substrate | Charge State | Calculated Product | Calculated Reactant | Found Product | Found Reactant |
|---|---|---|---|---|---|---|
| 4V | RGW | [M+H$^+$] | 496.1302, 498.1282 | 418.2197 | 496.1308, 498.1292 | 418.2208 |
| 4V | TGW | [M+H$^+$] | 441.0768, 443.0748 | 363.1663 | 441.0777, 443.0760 | 363.1782 |
| 4V | VGW | [M+H$^+$] | 439.0975, 441.0955 | 361.1870 | 439.0996, 441.0975 | 361.2014 |
| 4V | YGW | [M+H$^+$] | 503.0925, 505.0904 | 425.1819 | 503.0929, 505.0912 | 425.1864 |

**Bioactive Peptide Found and Calculated Mass List**

| Enzyme | Bioactive Peptide | Charge State | Calculated Product | Calculated Reactant | Found Product | Found Reactant |
|---|---|---|---|---|---|---|
| 4V | D-endomorphin-1 | [M+H⁺] | 689.2082, 691.2061 | 611.2976 | 689.2143, 691.2086 | 611.2986 |
| 4V | L-endomorphin-1 | [M+H⁺] | 689.2082, 691.2061 | 611.2976 | 689.2081, 691.2061 | 611.3119 |
| 4V | Galanin | [M+2H⁺] | 1619.2400 | 1579.2858 | 1619.2902 | 1579.2850 |
| 4V | Galanin | [M+3H⁺] | 1079.1631 | 1053.1930 | 1079.8611 | 1053.1956 |
| 4V | Neuropeptide W23 | [M+2H⁺ +MetOx] | 1340.6453 | 1300.6910 | 1340.1724 | 1300.7113 |
| 4V | Neuropeptide W23 | [M+3H⁺ +MetOx] | 893.4333 | 867.4631 | 893.4534 | 867.4780 |
| 4V | Neuropeptide W23 | [M+4H⁺ +MetOx] | 670.8263 | 650.8492 | 670.8386 | 520.8948 |
| 4V | $(RW)_3$ | [M+H⁺] | 1122.4856, 1124.4836 | 1044.5751 | 1122.4918, 1124.4906 | 1044.5806 |
| 4V | $(RW)_3$ | [M+2H⁺] | 561.7464, 562.7454 | 522.7912 | 561.7513, 562.7505 | 522.7990 |
| 4V | $(RW)_3$ | [M+3H⁺] | 374.8334, 375.4994 | 348.8632 | 374.8376, 375.5036 | 348.8742 |
| 4V | $(RW)_4$ | [M+1H⁺] | 1464.6660, 1466.6640 | 1386.7555 | 1464.6654, 1466.6671 | 1386.7616 |
| 4V | $(RW)_4$ | [M+2H⁺] | 732.8367, 733.8356 | 693.8814 | 732.8428, 733.8409 | 693.8604 |
| 4V | $(RW)_4$ | [M+3H⁺] | 488.8935, 489.5595 | 462.6234 | 488.8979, 489.5642 | 462.9352 |
| 4V | $(RW)_4$ | [M+4H⁺] | 366.9220, 367.4215 | 347.4443 | 366.9261, 367.4252 | 347.4560 |
| 4V | Spexin-1 | [M+H⁺+ MetOx] | 857.3745 | 818.4192 | 857.3779 | 818.5085 |
| 4V | Spexin-1 | [M+2H⁺ +MetOx] | 571.9187 | 545.9486 | 571.9235 | 546.0001 |
| 4V | Transportan | [M+2H⁺] | 1460.8238 | 1420.8695 | 1460.8439 | 1420.822 |
| 4V | Transportan | [M+3H⁺] | 973.5523 | 947.5821 | 973.8994 | 947.5948 |
| 4V | Transportan | [M+4H⁺] | 730.9155 | 710.9384 | 730.9276 | 710.9512 |
| No Br | D-endomorphin-1 | [M+H⁺] | 689.2082, 691.2061 | 611.2976 | Not found | 611.3203 |
| No Br | L-endomorphin-1 | [M+H⁺] | 689.2082, 691.2061 | 611.2976 | Not found | 611.3113 |
| No Br | Galanin | [M+2H⁺] | 1619.2400 | 1579.2858 | Not found | 1579.2881 |
| No Br | Galanin | [M+3H⁺] | 1079.1631 | 1053.1930 | Not found | 1053.1940 |
| No Br | Neuropeptide W23 | [M+2H⁺ +MetOx] | 1340.6453 | 1300.6910 | Not found | 1300.7135 |
| No Br | Neuropeptide W23 | [M+3H⁺ +MetOx] | 893.4333 | 867.4631 | Not found | 867.4796 |

| Enzyme | Bioactive Peptide | Charge State | Calculated Product | Calculated Reactant | Found Product | Found Reactant |
|---|---|---|---|---|---|---|
| No Br | Neuropeptide W23 | $[M+4H^{+}$ $+MetOx]$ | 670.8263 | 650.8492 | Not found | 650.8646 |
| No Br | $(RW)_3$ | $[M+H^{+}]$ | 1122.4856, 1124.4836 | 1044.5751 | Not found | 1044.5801 |
| No Br | $(RW)_3$ | $[M+2H^{+}]$ | 561.7464, 562.7454 | 522.7912 | Not found | 522.7997 |
| No Br | $(RW)_3$ | $[M+3H^{+}]$ | 374.8334, 375.4994 | 348.8632 | Not found | 348.8756 |
| No Br | $(RW)_4$ | $[M+1H^{+}]$ | 1464.6660, 1466.6640 | 1386.7555 | Not found | 1386.7739 |
| No Br | $(RW)_4$ | $[M+2H^{+}]$ | 732.8367, 733.8356 | 693.8814 | Not found | 693.8986 |
| No Br | $(RW)_4$ | $[M+3H^{+}]$ | 488.8935, 489.5595 | 462.6234 | Not found | 462.9417 |
| No Br | $(RW)_4$ | $[M+4H^{+}]$ | 366.9220, 367.4215 | 347.4443 | Not found | 347.4609 |
| No Br | Spexin-1 | $[M+H^{+}+$ $MetOx]$ | 857.3745 | 818.4192 | Not found | 818.5014 |
| No Br | Spexin-1 | $[M+2H^{+}$ $+MetOx]$ | 571.9187 | 545.9486 | Not found | 545.9942 |
| 4V | Transportan | $[M+2H^{+}]$ | 1460.8238 | 1420.8695 | Not found | 1420.8914 |
| 4V | Transportan | $[M+3H^{+}]$ | 973.5523 | 947.5821 | Not found | 947.6007 |
| 4V | Transportan | $[M+4H^{+}]$ | 730.9155 | 710.9384 | Not found | 710.9548 |
